# Global profiling of human blood ILC subtypes reveals that NK cells produce homeostatic cytokine amphiregulin and sheds light on HIV-1 pathogenesis

**DOI:** 10.1101/2021.04.20.440368

**Authors:** Yetao Wang, Lawrence Lifshitz, Noah J. Silverstein, Esther Mintzer, Kevin Luk, Pam St. Louis, Michael A Brehm, Scot A. Wolfe, Steven G. Deeks, Jeremy Luban

**Affiliations:** Hospital for Skin Diseases (Institute of Dermatology), Chinese Academy of Medical Sciences and Peking Union Medical College, Nanjing, China.; Key Laboratory of Basic and Translational Research on Immune-Mediated Skin Diseases, Chinese Academy of Medical Sciences, Nanjing, China.; Jiangsu Key Laboratory of Molecular Biology for Skin Diseases and STIs, Institute of Dermatology, Chinese Academy of Medical Sciences and Peking Union Medical College, Nanjing, China; Program in Molecular Medicine, University of Massachusetts Medical School, Worcester, MA, USA; Department of Molecular, Cell and Cancer Biology, University of Massachusetts Medical School, Worcester, MA, USA; Diabetes Center of Excellence, University of Massachusetts Medical School, Worcester, MA, USA; Department of Medicine, University of California, San Francisco, CA, USA; Department of Biochemistry and Molecular Pharmacology, University of Massachusetts Medical School, Worcester, MA, USA; Broad Institute of MIT and Harvard, Cambridge, MA, USA; Ragon Institute of MGH, MIT, and Harvard, Cambridge, MA, USA; Massachusetts Consortium on Pathogen Readiness, Boston, MA, USA

## Abstract

The interrelatedness of human blood innate lymphoid cell (ILC) subsets, and how they are perturbed by HIV-1, remains unclear. Transcriptional and chromatin profiling separated blood ILCs into ILC2s, ILCPs, one cluster that included CD56^dim^ and CD56^−^NK cells, and CD56^hi^NK cells that have features of both CD56^dim/–^NK cells and ILCs. In contrast to mice, human NK cells expressed tissue repair protein amphiregulin (AREG), with greater production by CD56^hi^NK cells than by ILCs. AREG was induced by TCF7/WNT signaling, IL-2, or IL-15, but not by inflammatory cytokines, and was inhibited by TGFB1, a cytokine elevated in people living with HIV-1. NK cell knockout of the TGFB1-stimulated WNT antagonist RUNX3 increased AREG production. In people living with HIV-1, AREG^+^NK cell percentage correlated with numbers of ILCs and CD4^+^T cells, and correlated inversely with inflammatory cytokine IL-6. RNA-Seq showed increased antiviral gene expression in all ILC subsets from people who were HIV-1 viremic, and increased expression of anti-inflammatory gene MYDGF in CD56^hi^NK cells from elite controllers. Functionally-defective CD56^−^NK cells were increased in people living with HIV-1 in inverse correlation with CD56^dim^NK cells, ILCs, and CD4^+^T cells. Experiments with human PBMCs *ex vivo* and in humanized mice revealed that CD4^+^T cells and their production of IL-2 prevented CD56^dim^ transition to CD56^−^NK cells by activating mTOR, and, in people living with HIV-1, plasma IL-2 correlated with CD4^+^T cell number but not with CD8^+^T cells. These studies clarify how ILC subsets are interrelated and provide insight into how HIV-1 infection disrupts NK cells, including homeostatic functions of NK cells discovered here.

**Graphical Abstract:** 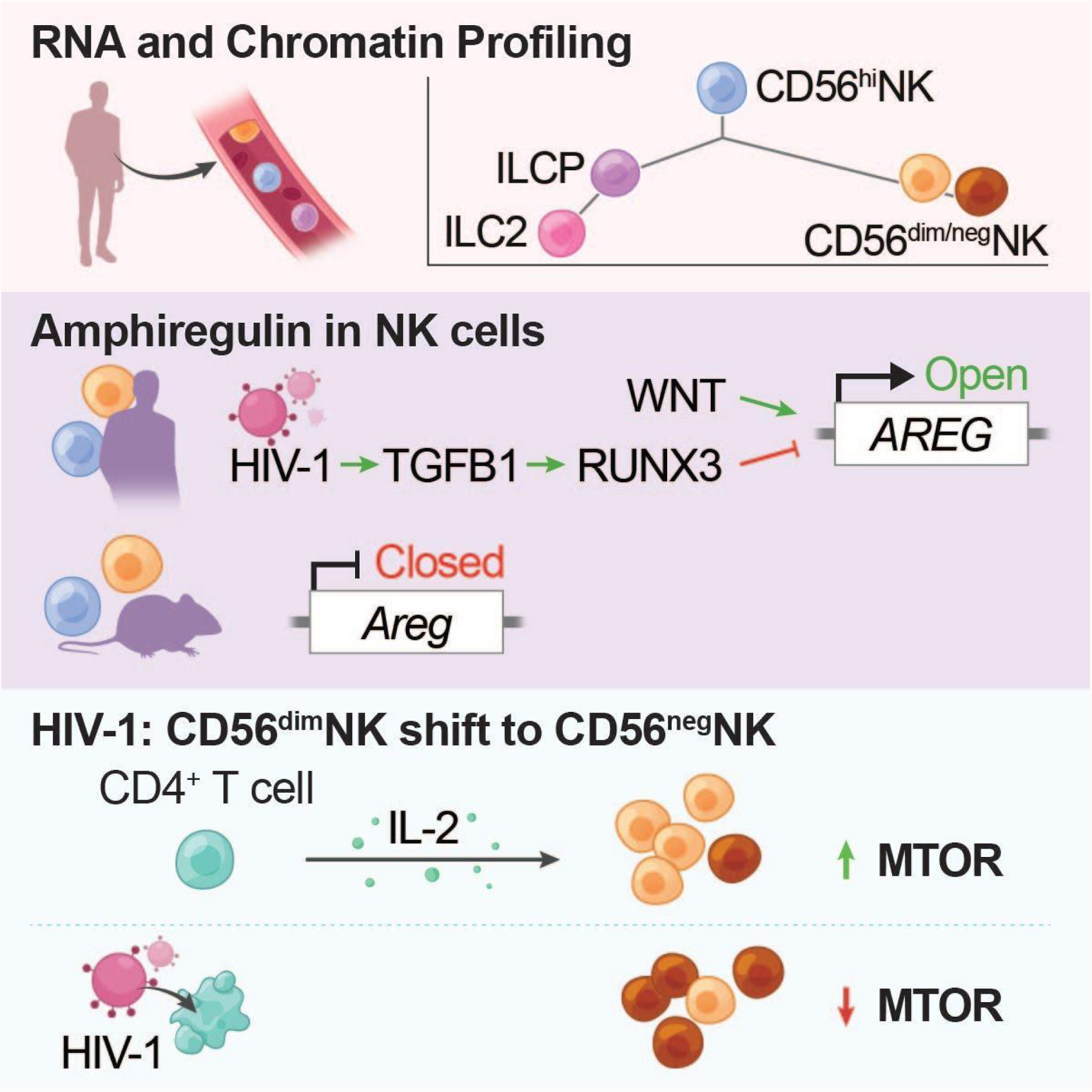

## Introduction

Innate lymphoid cells (ILCs) are a diverse population of cells which contribute to a broad range of biological functions, including tissue homeostasis and repair, inflammation, and protection from infection (1–3). All ILCs bear the leukocyte common antigen CD45, but lack markers of well-characterized cellular lineages, including T cells, B cells, hematopoietic stem cells, mast cells, and myeloid cells (4–6).

CD127, the cell surface protein IL-7Rα encoded by IL7R, defines a heterogeneous subset of cytokine-producing ILCs, with developmental and functional parallels to CD4^+^ helper T cells (7, 8). Specifically, ILC1s are analogs of T_H_1 cells, which are defined by the transcription factor TBX21, and produce IFN-γ. Similar to T_H_2 cells, ILC2s are defined by GATA3^+^, bear cell surface marker CRTH2^+^, and produce IL-4, IL-5, and IL-13. ILC3s are counterparts of T_H_17 cells, which express RORγT and produce IL-17 and IL-22.

Natural killer (NK) cells are a subset of ILCs, more akin to CD8^+^ T cells in that they have cytolytic activity against tumor cells and virus-infected cells mediated by perforin and granzymes (7, 8). NK cells bear proteins that distinguish them from the other ILCs, including CD56, CD16, and the transcription factor EOMES (6–9).

Despite diversity in gene expression, protein production, and phenotype, all human ILC subsets can be generated experimentally from a common innate lymphoid cell precursor (ILCP) found in the blood (10). Consistent with shared derivation from a common precursor, the canonical markers described above do not always discriminate between the ILC subsets. For example, unlike the vast majority of CD56^dim^NK cells, CD56^hi^NK cells bear CD127, CD117, and TCF7 (6), proteins typical of ILCs.

Perturbation of ILC and NK cell subsets has been reported in association with Crohn’s disease, psoriasis, chronic obstructive pulmonary disease, non-small cell lung cancer, or infection with any of several viruses, including human immunodeficiency virus type 1 (HIV-1), human cytomegalovirus (HCMV), hepatitis C virus (HCV), and influenza A virus (6, 11, 12). People living with HIV-1 have permanent depletion of ILC2s in the blood, and of ILC3s in the intestinal lamina propria, even after viremia has been suppressed by antiviral therapy (6, 13, 14).

CD56^hi^NK cells are increased in the blood of people living with HCV (20), and in people with melanoma (15), breast cancer (16), and Sjögren’s syndrome (17). A subset of NK cells that are TCF7+ are increased in people living with HIV-1 (6). The numbers of a functionally-defective Lin^−^CD56^−^CD16^+^NK cell subset are also expanded by viral infection (HIV-1, HCV, HCMV, and hantavirus), or by autoimmune disease (12, 18, 19). These CD56^−^NK cells appear to be derived from CD56^dim^NK cells in that the expansion of CD56^−^NK cells is accompanied by a decrease in CD56^dim^NK cells (12, 18, 19), and when CD56^−^NK cells from people with untreated HIV-1 infection are incubated *ex vivo* with exogenous IL-2, CD56 becomes detectable, though cytolytic function is not restored to the level of CD56^dim^ cells (19).

Taken together, these observations concerning human blood ILCs and NK cells under conditions of normal homeostasis and in the context of pathogenic inflammation, indicate that much remains to be clarified regarding the relationship between these cell types. Here, global transcriptional and chromatin profiling was used to investigate the relationships between blood ILC and NK cell subsets, and the changes in these cells in people living with HIV-1 infection, including people who were untreated/viremic, treated with antiretroviral therapy (ART), spontaneous controllers, or elite controllers. These studies provide insight into the relationship between these innate lymphoid cell types, reveal previously unappreciated functions of NK cells, and demonstrate how HIV-1 infection perturbs them.

## Results

### CD127 and CD56 identify four discrete cell populations among Lin^−^ human PBMCs

To identify ILC subsets in human peripheral blood mononuclear cells (PBMCs), 14 lineage antibodies (against CD3, CD4, TCRαβ, TCRγδ, CD19, CD20, CD22, CD34, FcεRIα, CD11c, CD303, CD123, CD1a, and CD14) were used to exclude T cells, B cells, monocytes, and other cells from defined lineages (Supplemental Table 1). Steady-state human blood innate lymphoid cell populations are composed of Lin^−^CD127^+^CD56^−^ ILCs and Lin^−^CD56^+^ NK cells (Figure 1A) (5, 6, 11). However, Lin^−^CD127^−^CD56^−^ cells that appear to be distinct from ILCs are often apparent (Figure 1A). To better characterize this population, and to clarify its relatedness to ILCs, Lin^−^CD45^+^CD56^−^PBMCs from five HIV-1-negative donors were sorted into CD127^−^ and CD127^+^ subpopulations, and each was subjected to bulk RNA-Seq (Figure 1A and Supplemental Figure 1A). 401 and 241 differentially expressed genes (DEgenes) were enriched among the CD127^−^ and CD127^+^ cells, respectively (Figure 1B and Supplemental Table 2).

**Figure 1.**
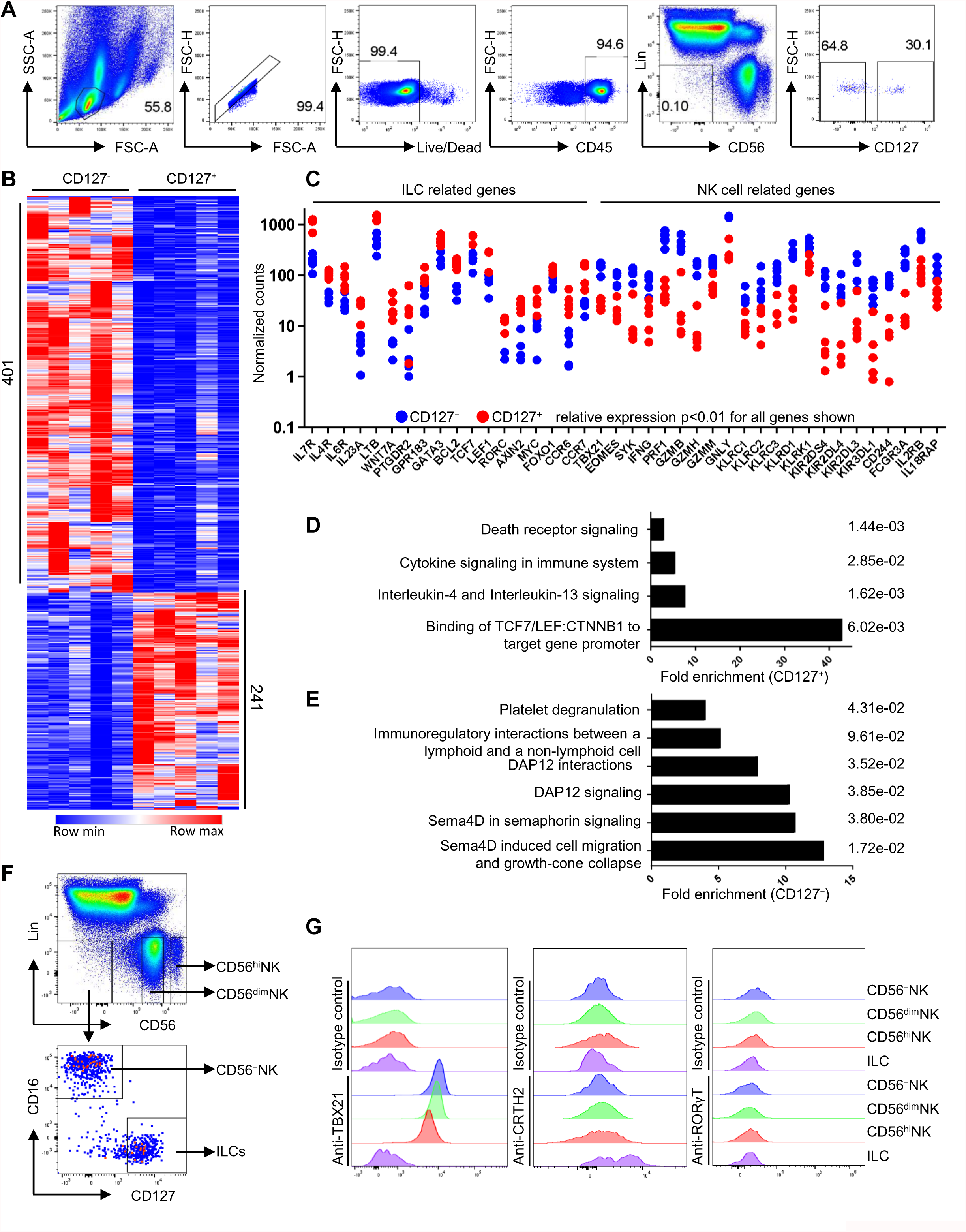
Characterization of Lin^−^ lymphoid cells among human PBMCs. (A) CD127^+^ and CD127^−^ cells from Lin^−^CD56^−^ population after gating on lymphoid, singlet, live, CD45^+^ cells of PBMCs. Lineage (Lin) markers include antibodies against: CD3, CD4, TCRαβ, TCRγδ, CD19, CD20, CD22, CD34, FcεRIα, CD11c, CD303, CD123, CD1a, and CD14. (B) Heatmap of differentially expressed genes by RNA-Seq, sorted Lin^−^CD56^−^CD16^−^CD127^+^ versus Lin^−^CD56^−^CD16^−^CD127^−^ cells, from PBMCs of five donors (log2 fold change >1, padj<0.01 determined by DESeq2, the post-sort profiles are shown in Supplemental Figure 1). (C) Normalized counts of Lin^−^CD56^−^CD127^+^ (red) and Lin^−^CD56^−^CD127^−^ (blue) cells related genes from (B) by DESeq2 (n=5). (D, E) Reactome analysis based on enriched transcripts of Lin^−^CD56^−^CD127^+^ cells (D) or Lin^−^CD56^−^CD127^−^ cells (E). (F) Gating of Lin^−^CD56^hi^NK, Lin^−^CD56^dim^NK, Lin^−^CD56^−^NK cells and Lin^−^CD56^−^CD16^−^CD127^+^ILCs. (G) The indicated populations in (F) were detected with isotype controls or antibodies against TBX21, CRTH2 and RORγT. All data were generated using blood from healthy HIV-1-negative donors.

Consistent with previous reports (6, 10), Lin^−^CD45^+^CD56^−^CD127^+^ blood cells were enriched for genes specific to ILC2s, or shared with ILCPs, including IL7R (CD127), IL4R, IL6R, PTGDR2 (CRTH2), CCR6, CCR7, GATA3 and TCF7. Gene ontology (GO) analysis highlighted the association of expressed genes with cytokine production (Figure 1C, 1D and Supplemental Table 2).

The Lin^−^CD45^+^CD56^−^CD127^−^ population was enriched for mRNAs typical of NK cells, including TBX21, EOMES, IFNG, KLRD1 (CD94), 2B4 (CD244), FCGR3A, PRF1, GZMB, GZMH, GZMM, GNLY, and killer Ig-like receptors (KIRs) (KIR2DL3, KIR2DL4, KIR2DS4, KIR3DL1, KIR3DL2, and KIR3DL3) (Figure 1C and Supplemental Table 2), and express NK cell-specific receptors (such as NKG2C and NKp44) that signal via DAP12 (20, 21) (Figure 1E and Supplemental Table 2). ARHGEF12, ERBB2, RHOC, and MYL9 were well-expressed on these cells (Figure 1E and Supplemental Table 2); the latter genes are required for signaling by SEMA4D, a protein that regulates NK cell killing activity and IFN-γ production (22–25). TBX21 and CD16 proteins were detected by flow cytometry on the majority of Lin^−^CD45^+^CD56^−^CD127^−^ cells (Figure 1F and 1G), consistent with the fact that, despite undetectable CD56 protein, this population produces proteins typical of *bona fide* NK cells (6). Conversely, flow cytometry of Lin^−^CD56^−^CD16^−^CD127^+^ILCs failed to detect NK cell-associated proteins, including EOMES, CD94, KIR3DL1, KIR2DL1, KIR2DL4, NKp44, NKp46, NKp80, NKG2A, and NKG2C (Supplemental Figure 1B). However, ∼10% of ILCs were NKG2D positive (Supplemental Figure 1B), consistent with a previous study (26).

Taking together the transcriptional profiling and flow cytometry analysis, Lin^−^PBMCs can be divided into four main ILC subsets, Lin^−^CD56^hi^NK cells, Lin^−^ CD56^dim^NK cells, Lin^−^CD56^−^NK cells, and Lin^−^CD56^−^CD16^−^CD127^+^ILCs, the latter including CD127^+^CRTH2^+^ILC2s and CD127^+^CRTH2^−^CD117^+^ILCPs (Figure 1F, 1G and Supplemental Figure 1C).

### Global transcription and epigenetic features of the four ILC subsets in human blood

To better characterize the four ILC subsets in human blood, PBMCs from four HIV-1-negative blood donors were sorted into ILCs (Lin^−^CD56^−^CD16^−^CD127^+^), CD56^−^NK cells (Lin^−^CD56^−^CD127^−^CD16^+^), CD56^dim^NK cells (Lin^−^CD56^dim^), and CD56^hi^NK cells (Lin^−^CD56^hi^). Each population was then separately subjected to bulk RNA-Seq (Supplemental Figure 1D).

When compared with ILCs, CD56^−^, CD56^dim^, and CD56^hi^ NK cells exhibited 1,128, 1,236, and 910 DEgenes, respectively (Figure 2A and Supplemental Table 3). In contrast with CD56^−^ and CD56^dim^ NK cells, CD56^hi^NK cells expressed lower levels of GZMB and GZMH, and shared many transcripts with ILCs, including IL7R (CD127), KIT, IL4R, IL1RL1 (IL33R), IL18R1, CCR7, GPR183, TCF7, and MYC (Supplemental Figure 2A and Supplemental Table 3) (27). Surprisingly, AREG, the gene encoding the epidermal growth factor (EGF)-like amphiregulin that is produced by ILC2s and promotes tissue repair and homeostasis (28), was expressed in all NK cell subsets, with highest levels in CD56^hi^ NK cells (Figure 2B and Supplemental Table 3). In contrast, multiple mouse datasets showed that AREG was expressed by ILC2s, but not by other ILC subsets, nor by NK cells (Supplemental Figure 2B-2E and Supplemental Supplemental Table 4) (29–31), indicating a major difference in AREG regulation between mouse and human.

**Figure 2.**
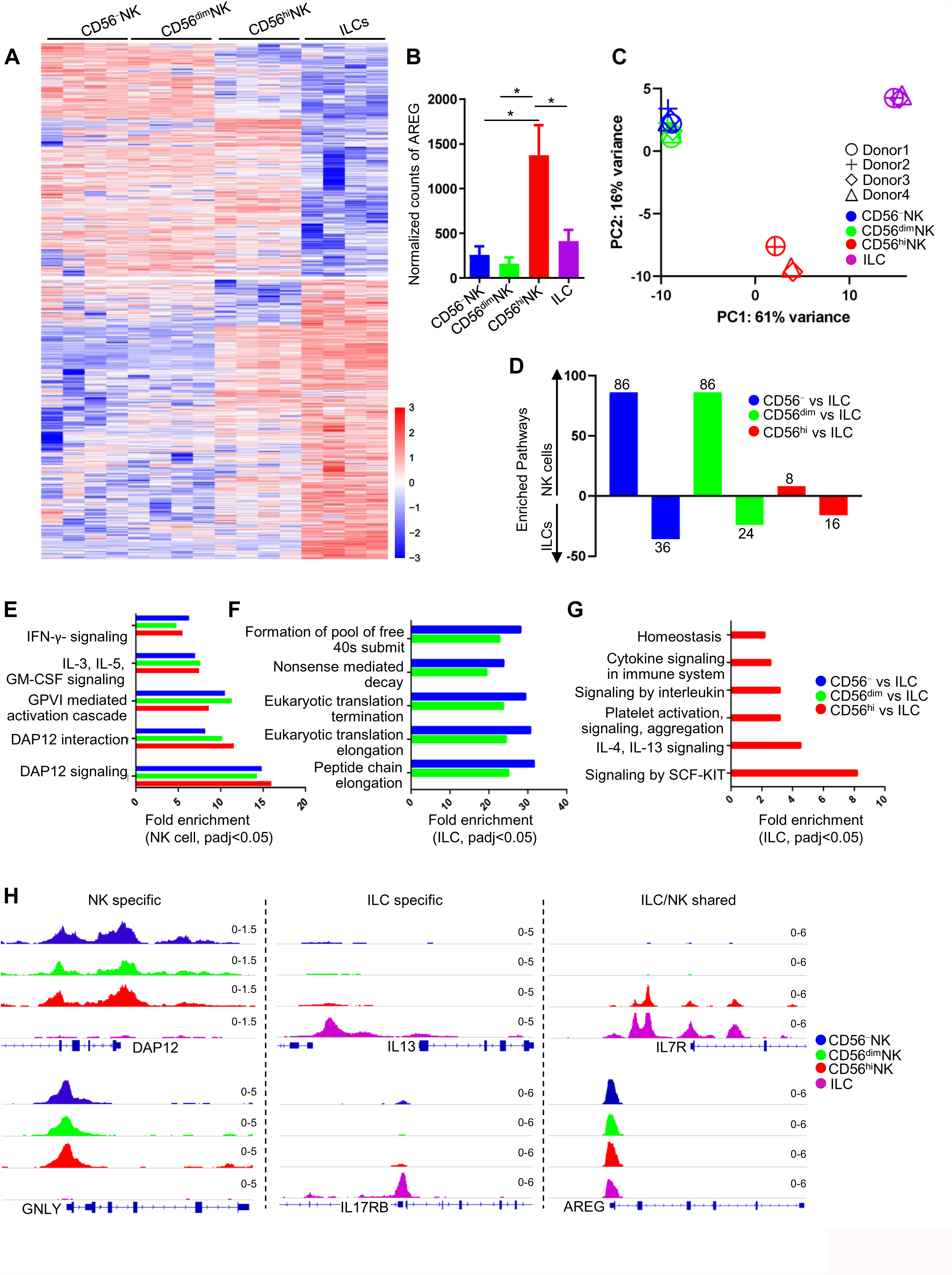
Transcriptional and chromatin accessibility analysis of human blood NK cells and ILCs. (A) Heatmap of differentially expressed genes by RNA-Seq, sorted CD56^hi^NK, CD56^dim^NK, CD56^−^NK cells and ILCs, from PBMCs of 4 donors (log2 fold change >1, p<0.01 determined by DESeq2, the post-sort profiles are shown in Supplemental Figure 1). (B) Normalized counts of AREG from (A) by DESeq2. (C) PCA based on RNA-Seq data of indicated populations. (D) Number of enriched pathways based on differentially expressed genes of indicated NK subsets versus ILCs by Go Enrichment Analysis. (E) Enriched pathways shared by all NK subsets as compared with ILCs. (F) Shared enriched pathways of ILCs compared with CD56^dim^ or CD56^−^NK cells. (G) Enriched pathways of ILCs compared with CD56^hi^NK cells. (H) ATAC-Seq analysis of sorted CD56^hi^NK, CD56^dim^NK, CD56^−^NK cells and ILCs at indicated gene loci (representative of two donors). All data were generated using blood from healthy HIV-1-negative donors.

Principal component analysis confirmed that the human blood ILC transcriptome was distinct from that of all three NK cell subsets, that CD56^dim^ and CD56^−^ NK cell transcriptomes were nearly indistinguishable from each other, and that the CD56^hi^NK cell transcriptome was distinct from that of the other NK subsets (Figure 2C).

Reactome analysis showed that, as compared with ILCs, 86 classified pathways were enriched in CD56^−^ and CD56^dim^ NK cells, whereas only 8 pathways were enriched in CD56^hi^NK cells (Figure 2D and Supplemental Table 3). All NK cell subsets, though, were distinguished from ILCs by five pathways, which highlight the fundamental role of IFN-γ and DAP12 signaling in the three NK cell subsets (Figure 2E, Supplemental Figure 2F and Supplemental Table 3). Typical of cells with a primary role in secretion of cytokines, the ILCs had high-level expression of genes encoding ribosomal proteins, and proteins involved in translation initiation and elongation (Figure 2F, Supplemental Figure 2G and Supplemental Table 3). The ILCs were further distinguished from CD56^hi^NK cells by higher expression of IL-4, IL-13, and SCF-KIT pathways, consistent with established functions of ILC2s and ILCPs (Figure 2G, Supplemental Figure 2H and Supplemental Table 3).

The global transcriptional profiles of the ILCs and NK cell subsets were mirrored by the chromatin accessibility of genes, as determined by ATAC-Seq. DAP12, GNLY, EOMES, TBX21, NKG7, ILR2B, and CST7 loci were more accessible to Tn5 transposase in NK cell subsets, whereas IL13, IL17RB, PTGDR2, TNFRSF25, SOCS3, IL23A, and IL32 loci were more accessible in ILCs (Figure 2H and Supplemental Figure 3A). The IL7R promoter was open in both ILCs and CD56^hi^NK cells, but not in CD56^dim/–^ NK cells, and the AREG promoter was accessible in all NK subsets, as well as in ILCs (Figure 2H). In contrast to these results with human cells, the mouse Areg promoter was only accessible in mouse ILC2s, but not in other ILC subsets or in NK cells (Supplemental Figure 3B). Interestingly, despite undetectable RORγT protein in blood ILCs (Figure 1G), which are primarily ILC2s in phenotype, the chromatin at the RORC promoter was open (Supplemental Figure 3A), consistent with the phenotypic plasticity of these cells and their potential to acquire an ILC3 phenotype (10).

### Single cell transcriptional analysis of human blood ILCs and NK cells

To clarify the relationship among different ILC subsets, PBMCs from three HIV-1-negative blood donors were sorted into ILCs (Lin^−^CD45^+^CD56^−^CD127^+^), CD56^−^NK cells (Lin^−^CD45^+^CD56^−^CD127^−^CD16^+^), CD56^dim^NK cells (Lin^−^CD45^+^CD56^dim^), and CD56^hi^NK cells (Lin^−^CD45^+^CD56^hi^). Equal numbers of each subset were pooled and subjected to single cell RNA-Seq (Supplemental Figure 1D). Transcriptomes from 5,210 individual cells were analyzed, and 96% of the cells fit within one of four main clusters (Figure 3A), though surprisingly, not the same four subsets that had been sorted.

**Figure 3.**
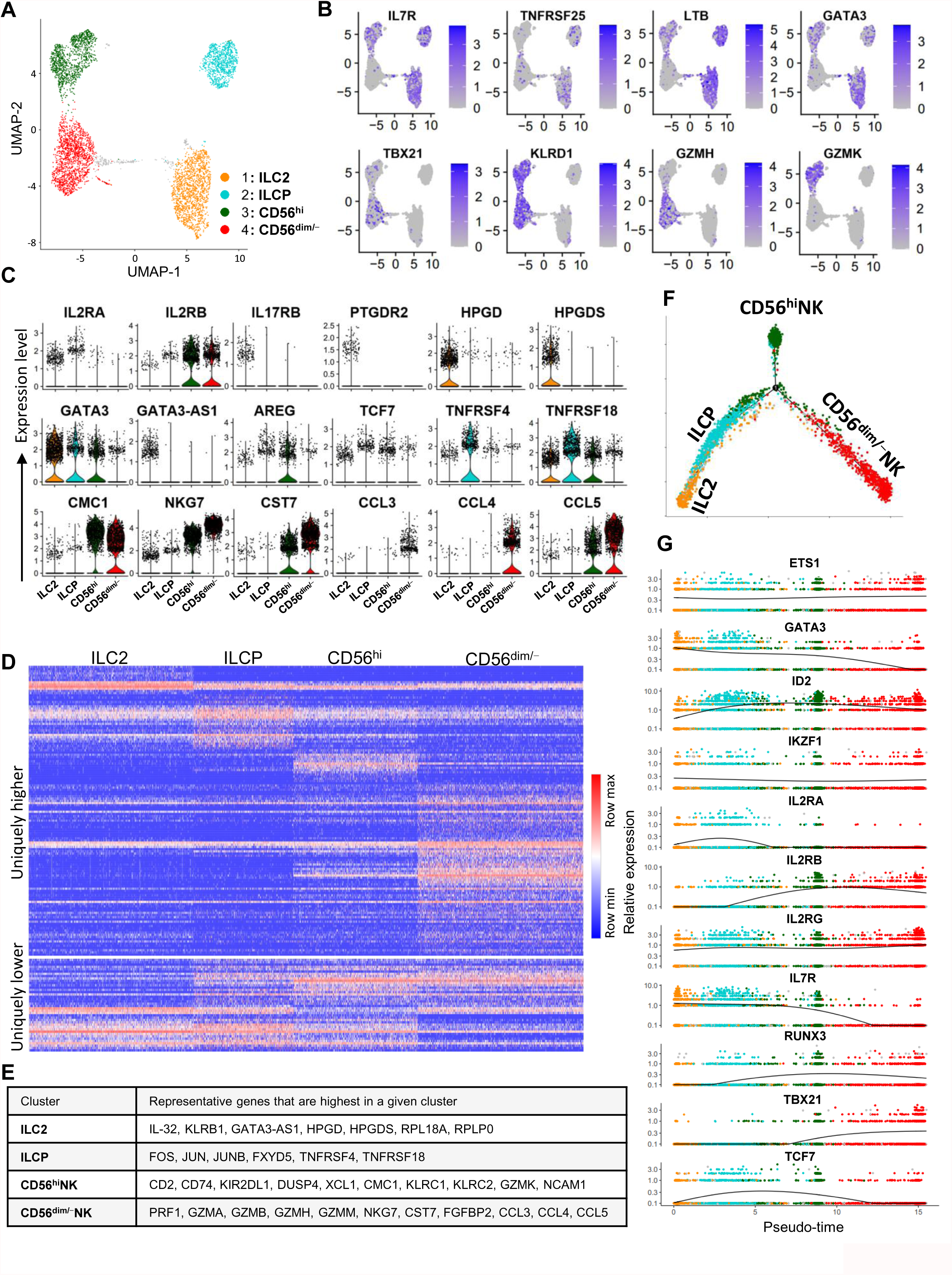
Single cell transcriptome analysis of blood NK cells and ILCs. (A) Uniform manifold approximation and projection (UMAP) of single cell RNA-Seq of sorted CD56^hi^NK, CD56^dim^NK, CD56^−^NK cells and ILCs pooled from 3 donors. (B, C) Expression and density of the indicated genes within UMAP. (D) Heatmap of uniquely higher or lower expressed genes of NK cell or ILC clusters. (E) Representative genes of indicated clusters. (F) Minimum spanning tree based on the transcriptome of individual cells from (A) showing pseudotime trajectory (black line, cells are color coded by clusters). (G) the expression of genes that are enriched by NK cells and/or ILCs along the pseudotime trajectory. All data were generated using blood from healthy HIV-1-negative donors.

The sorted Lin^−^CD45^+^CD56^−^CD127^+^ ILCs formed two distinct clusters. Cluster one was enriched for expression of genes that define ILC2s, including IL7R (CD127), TCF7, PTGDR2 (CRTH2), GATA3, IL2RA, TNFRSF25, LTB, IL17RB (Figure 3B-3E and Supplemental Table 5). GATA3-AS1, a gene which increases transcription of IL-5, IL-13, and GATA3 (32), was also enriched in this cluster, as were HPGD and HPGDS, genes encoding prostaglandin D2 biosynthetic enzymes required for cytokine production by ILC2s (Figure 3C-3E) (33).

Cluster two lacked critical ILC2-associated transcripts, and had higher expression of TNFRSF4 and TNFRSF18, genes that define ILCPs (10). FOS, JUN, and JUNB, genes required for cell survival, proliferation, and development (34), were also enriched in this ILCP cluster (Figure 3C-3E and Supplemental Table 5).

Cluster three was defined as CD56^hi^NK cells, based on expression of GZMK (6), and on enrichment for transcripts from both NK cells and ILCs, including TBX21, KLRD1, IL7R, LTB, GATA3, and TCF7 (Figure 3B-3E and Supplemental Table 5). Consistent with the PCA analysis of bulk RNA-Seq data (Figure 2C), cluster four consisted of both CD56^dim^ and CD56^−^ NK cells, despite the fact that the two subsets had been separated by flow cytometry based on CD56 positivity. The CD56^dim/–^ cluster was distinguished from CD56^hi^NK cells by exclusively expressing GZMH, and higher levels of CCL3, CCL4, and CCL5. The NK cell signature genes KLRD1, CMC1, NKG7, and CST7 were shared by the CD56^hi^ and CD56^dim/–^ NK cell clusters (Figure 3B-3E and Supplemental Table 5) (35). Consistent with bulk RNA-Seq data (Supplemental Figure 2 and Supplemental Table 3), AREG mRNA was detected in all four clusters (Figure 3C).

Pseudotime analysis (36, 37) was used to determine how the four clusters of blood ILCs are related to each other. ILCs and ILCPs formed one branch, and CD56^−^ and CD56^dim^ NK cells formed a second distinct branch (Figure 3F). ETS1, ID2, IKZF1, and IL2RG, genes essential for ILC and NK cell development (38–40), were expressed along the full trajectory from ILCs/ILCPs to CD56^dim/–^NK cells (Figure 3G). Consistent with CD56^hi^NK cells exhibiting epigenetic and transcriptional features of both ILCs (IL7R, KIT, IL4R, IL1RL1, CCR7, GPR183, MYC and TCF7), and NK cells (TBX21, NCAM1, KLRD1, GZMB, GNLY, and KIRs) (Figure 2 and Supplemental Table 3), pseudotime analysis placed TBX21^+^CD56^hi^NK cells along the trajectory at the junction between TBX21-negative ILCs/ILCPs and TBX21^+^CD56^dim/–^ NK cells (Figure 3G), demonstrating that CD56^hi^NK cells occupy an intermediate state between these clusters.

### Gene expression profiling of NK cell subsets and ILCs from people living with HIV-1

To assess the effect of HIV-1 infection on the individual innate lymphoid cell subsets, CD56^hi^, CD56^dim^, and CD56^−^NK cells, and ILCs, were sorted from PBMCs donated by 10 HIV-1-negative people, 10 people living with HIV-1 who were viremic, 10 people living with HIV-1 who were treated with ART, and 10 people living with HIV-1 who were elite controllers (undetectable HIV-1 viral load without ART). lLCs were permanently depleted in people living with HIV-1, including those individuals who were treated with ART or who were controllers (Figure 7A) (6). Therefore, total ILCs were sorted here due to limiting numbers of ILCPs. In people living with HIV-1 the biggest shifts in NK cell subsets were in people who were viremic, in whom the numbers of CD56^−^NK cells were increased and the numbers of CD56^dim^NK cells were decreased most significantly (Figure 7C) (6). RNA-Seq was performed on each sorted cell population and PCA showed that the cell populations grouped most strongly according to cell type rather than according to HIV-1 infection status (Figure 4A).

**Figure 4.**
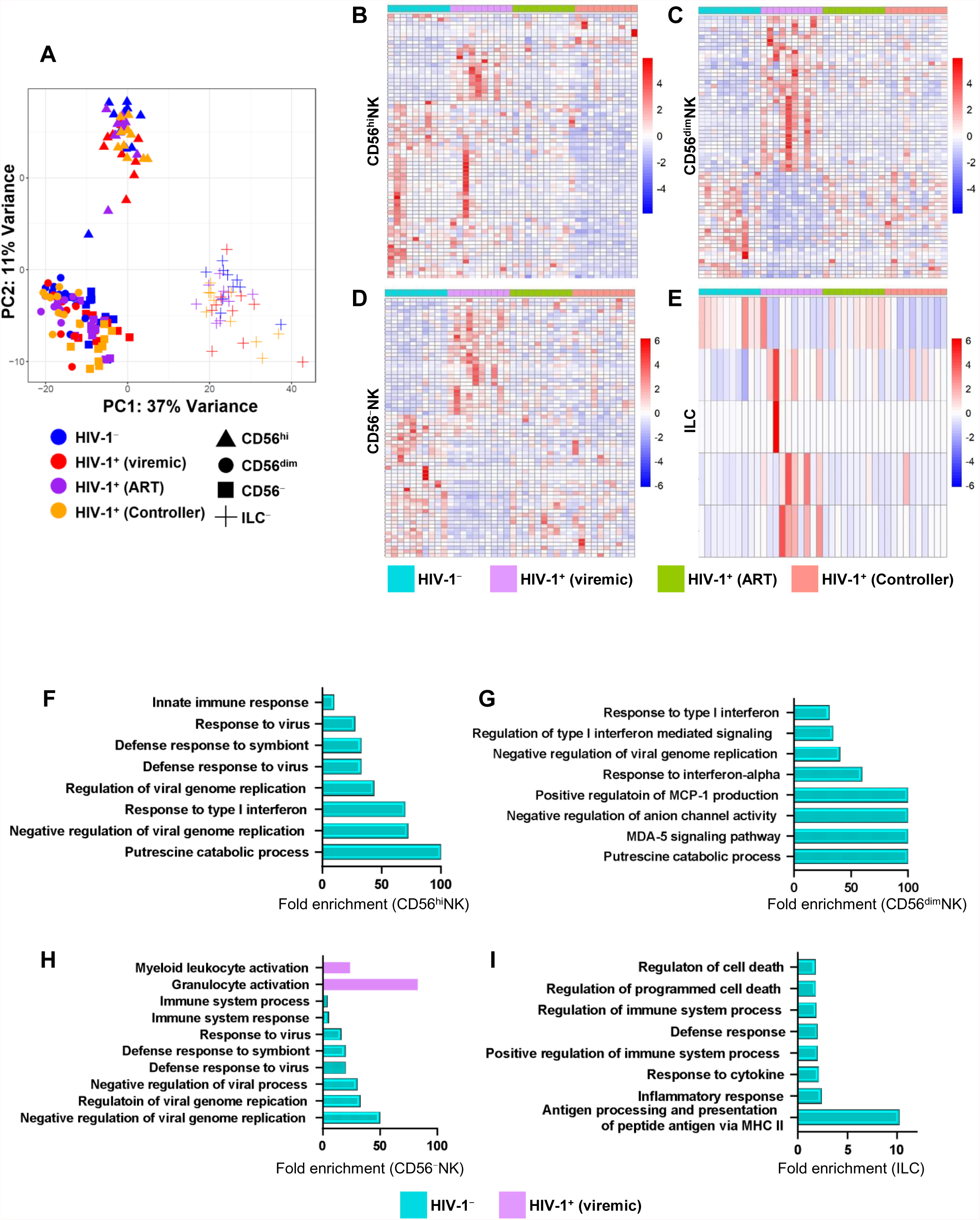
The effect of HIV-1 infection on NK cells and ILCs. (A) PCA based on RNA-Seq data of sorted NK cell subsets and ILCs from PBMCs of HIV-1 negative and different groups of HIV-1^+^ donors (each group 10 donors), post-sort profiles are shown in Supplemental Figure 1. (B-E) Heatmap of differentially expressed genes of sorted CD56^hi^NK (B), CD56^dim^NK (C), CD56^−^NK cells (D) and ILCs (E) from PBMCs of HIV-1^−^ (n=10), HIV-1^+^ viremic (n=10), HIV-1^+^ ART suppressed (n=10), and HIV-1^+^ spontaneous controllers (n=10) (log2 fold change>1, padj<0.05 determined by DESeq2). (F-I) Enriched pathways of CD56^hi^NK (F), CD56^dim^NK (G), CD56^−^NK cells (H) and ILCs (I) of HIV-1^−^ and HIV-1^+^ viremic people. Cohort is described in Supplemental Table 7.

The biggest effects of HIV-1 infection on NK cell gene expression were detected in viremic individuals and were evident in all three NK cell subsets (Figure 4B-4D and Supplemental Table 6). Among the differentially expressed genes were ISGs with antiviral activity, including ISG15, ISG20, IFI6, MX1, IFI44, IFI44L, IFIT1, IFIT3, and IFITM3 (Figure 4F-4H and Supplemental Table 6). The expression of certain genes important for NK cell function, including IL2RB, IL18RAP, SYK and FCER1G, was decreased in CD56^−^NK cells (Supplemental Table 6). Additionally, IRF7 and SAT1, two genes important for antiviral responses that were upregulated in CD56^hi^ and CD56^dim^ NK cells from HIV-1-viremic individuals, were not upregulated in CD56^−^NK cells (Supplemental Table 6) (41, 42). Among NK cell cytotoxicity-associated genes, GZMA and GZMB were modestly upregulated in CD56^hi^NK cells from HIV-1^+^ people under ART, whereas, GZMB was downregulated in CD56^−^NK cells from HIV-1^+^ viremic and ART groups (Supplemental Figure 4A). Compared with HIV-1-negative people, CD56^hi^NK cells from elite controllers upregulated MYDGF, which encodes a paracrine protein that suppresses inflammation and blunts endothelial injury by inhibiting tissue recruitment of neutrophils and other leukocytes (43, 44) (Supplemental Figure 4A and Supplemental Table 6).

Few DEgenes were detected in sorted, live ILCs from people living with HIV-1 when an adjusted p value of < 0.05 was used as a significance cut-off (Figure 4E and Supplemental Table 6). Unlike the NK cells, ILCs were depleted in all groups of people living with HIV-1, with more severe depletion in people who were untreated and viremic (6) (Figure 7A). Expression of cell death-associated genes, including CDKN1A, HIF1a, TPD52L1, and MADD, was increased in people with HIV-1 viremia when a less stringent cutoff for DEgenes was used (unadjusted p<0.05), along with genes such as HLA-DRA, HLA-DRB1, HLA-DPB1, and CD74 that are involved in antigen presentation (Figure 4I and Supplemental Table 6), a known role of ILCs during inflammation (13, 45, 46). These results suggest that ILCs are depleted by inflammation-induced apoptosis that accompanies HIV-1 infection.

### Human blood NK cells are major producers of amphiregulin

Analysis of the RNA-Seq data revealed high-level expression of the homeostatic cytokine gene AREG in human blood NK cells (Figure 2B and 3C). This was surprising given the lack of Areg expression in mouse NK cells (Supplemental Figure 2B). Cell-associated amphiregulin protein was readily detected by flow cytometry in blood ILCs and in all NK subsets from HIV-1 negative donors after stimulation with PMA and ionomycin (Figure 5A). In CD56^hi^NK cells the amphiregulin protein signal was even higher than in ILCs, whether it was assessed for mean fluorescence intensity or for percent positive cells (Figure 4A and Supplemental Figure 4B).

**Figure 5.**
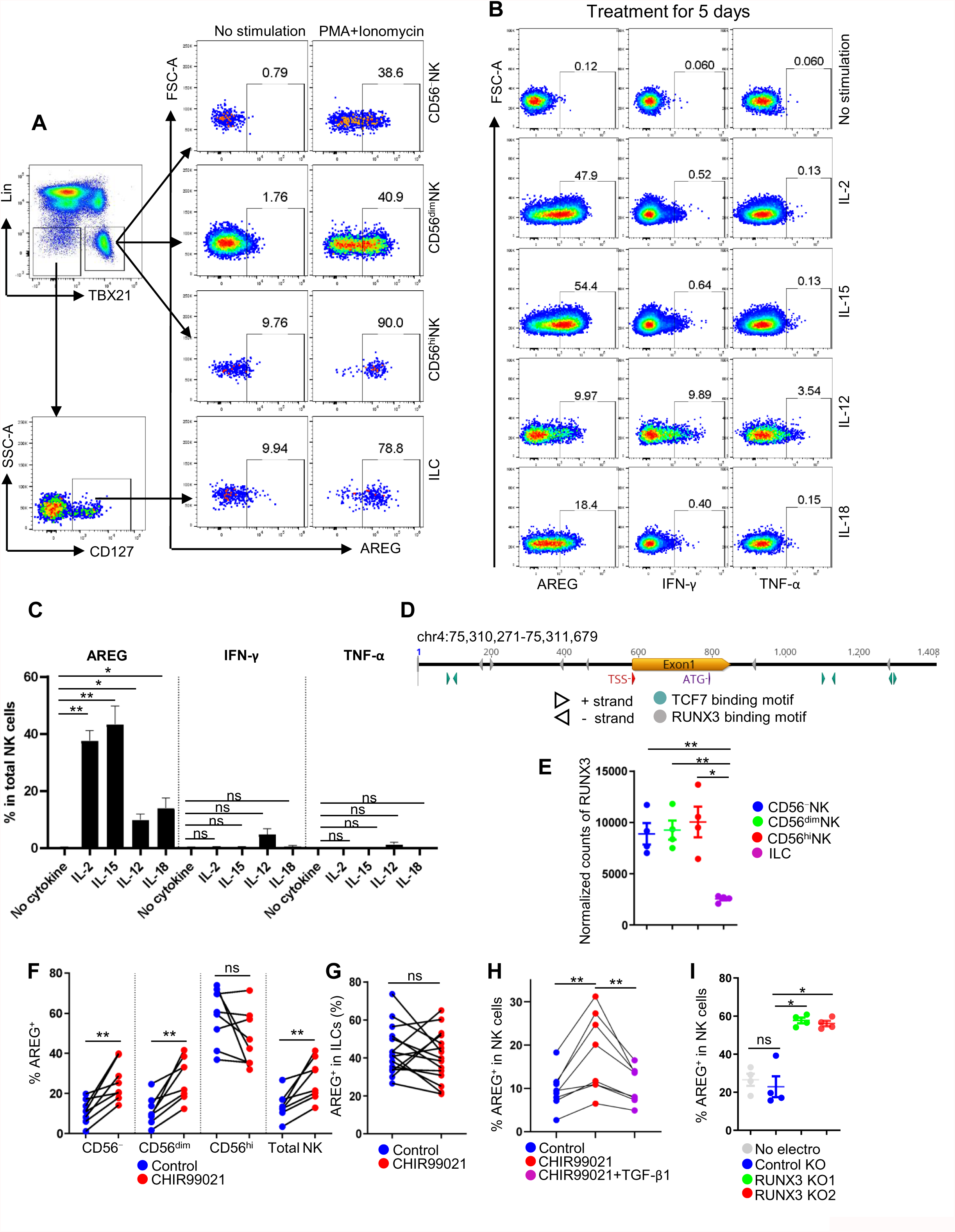
Human NK cells are major AREG producers among lineage negative PBMCs. (A) PBMCs were stimulated with PMA and ionomycin for 3 hrs and then intracellular AREG was detected within the different subsets of NK cells and ILCs by flow cytometry. **(B)** PBMCs were cultured in MACS NK medium at 37 °C in 5% CO_2_, 50 ng/ml IL-2, 50 ng/ml IL-15, 10 ng/ml IL-12 or 50 ng/ml IL-18 for 5 days, then AREG, IFN-γ or TNF-α from different subsets of NK cells and ILCs were detected. (C) The AREG^+^, IFN-γ^+^ or TNF-α^+^ NK cells detected in (B) (n=4). (D) Schematic map of AREG open chromatin region in NK cells as detected by ATAC-Seq. TCF7 and RUNX3 binding motifs were identified by JASPAR motif analysis (http://jaspar.genereg.net/). (E) Normalized counts of RUNX3 by DESeq2 (n=4). (F, G) PBMCs were treated with CHIR99021 (10 uM) in RPMI 1640 at 37 °C in 5% CO_2_ for 48 hrs, then stimulated with PMA+ionomycin. The percentage of AREG^+^NK cells (F) (n=8) and ILCs (G) (n=15) are shown. (H) As in (F), PBMCs were additionally treated with CHIR99021 and TGF-β1 (50 ng/ml) before AREG^+^NK cells were detected by flow cytometry (n=8). (I) AREG^+^NK cells from control or RUNX3 knockout groups were detected after IL-12+IL-15+IL-18 stimulation in MACS NK medium at 37°C in 5% CO_2_ for 16 hrs (n=4). Data are mean ± s.e.m., (C, E, F-I), two-tailed paired *t*-test. *p<0.05, **p<0.01, ***p<0.001. All data were generated using blood from healthy HIV-1-negative donors.

None of twenty-one stimuli that included cytokines, toll-like receptor agonists, and HIV-1-infected CD4^+^T cells, induced AREG protein in NK cells after 16 hrs (Supplemental Figure 4C and 4D). However, treatment with either IL-2 or IL-15 alone for 5 days induced AREG protein (Figure 5B and 5C). IL-12 or IL-18 also induced AREG, but the effect was not as strong as with IL-2 or IL-15 (Figure 5B and 5C). Therefore, in contrast to the induction in NK cells of inflammatory cytokines IFN-γ and TNF-α, which requires stimulation with at least two inflammatory cytokines (47), treatment with IL-2 alone was sufficient to stimulate production of the homeostatic cytokine AREG in NK cells.

Consistent with the expression of AREG in human NK cells, ATAC-Seq showed that the chromatin region around the human AREG promoter (chr 4: 75,310,271-75,311,679) was open in ILCs and in the three subsets of NK cells examined here (Figure 2H). In contrast, the mouse Areg chromatin region (chr 5: 91,287,998-91,289,503) is closed in NK cells, though open in ILCs (Supplemental Figure 3B). Interestingly, the open chromatin region surrounding the human AREG promoter contains six TCF7-binding sites and CD56^hi^NK cells and ILCs had high-level expression of TCF7 and other genes in the WNT signaling pathway (Figure 5D, Supplemental Figure 2A and Supplemental Table 3). TCF7 is expressed in mouse ILCs but not in mouse NK cells, and the mouse Areg chromatin region has only one TCF7 binding site (http://jaspar.genereg.net/).

Transcription factor RUNX3, an antagonist of TCF7 and WNT signaling (48, 49), was expressed at a higher level in all NK cell subsets than in ILCs (Figure 5E), and six RUNX3 binding sites were present in the open chromatin region of AREG (Figure 5D). Consistent with the biological significance of these differences in TCF7 and RUNX3 expression, WNT agonist CHIR99021 upregulated AREG production in CD56^−^ and CD56^dim^NK cells, but not in CD56^hi^NK cells and ILCs (Figure 5F and 5G). Furthermore, TGF-β1, which activates RUNX3 signaling and is upregulated during HIV-1 infection (50–53), attenuated the CHIR99021-induced AREG upregulation in NK cells (Figure 5H). Finally, the effect of Cas12a/RNP-mediated knockout (54) of RUNX3 was assessed in primary blood NK cells from HIV-1 negative donors. The editing rate for either of two crRNAs was greater than 90% (Supplemental Figure 4E, 2 upper panels) and, after knockout, RUNX3 protein was undetectable in most of these cells by flow cytometry (Supplemental Figure 4E, lower panel). As compared to cells treated with Cas12a/RNP and a control crRNA targeting the AAVS1, gene editing by either of the two RUNX-specific crRNAs increased AREG production in NK cells (Figure 5I and Supplemental Figure 4F). These results demonstrate that AREG production in human NK cells and ILCs is differentially regulated by TCF7/WNT and RUNX3.

### AREG^+^ NK cells in people living with HIV-1

AREG-expressing ILCs maintain tissue homeostasis and limit the tissue damage that results from viral infection and the associated inflammation (14, 28, 55, 56). People living with HIV-1 infection have reduced numbers of ILCs, and this reduction correlates inversely with systemic inflammation (6). Compared with HIV-1-negative individuals, people living with HIV-1 who were not taking ART had a decreased percentage of AREG^+^ cells within each of the NK cell subsets (Figure 6A and Supplemental Table 7). This was also the case among spontaneous controllers who maintain viral load below 2,000 copies of HIV-1 gRNA/ml without ART (Figure 6A). In HIV-1^+^ individuals on ART, the percentage of AREG^+^ cells was decreased, but only among CD56^hi^NK cells and ILCs (Figure 6A). Consistent with the effect on percentage, the total number of AREG^+^NK cells was also decreased by HIV-1 infection (Figure 6B). The number of AREG^+^ILCs was decreased in HIV-1^+^ people who are viremic or under ART, although the percentage AREG+ in HIV-1^+^ viremic people was not changed (Figure 6A).

**Figure 6.**
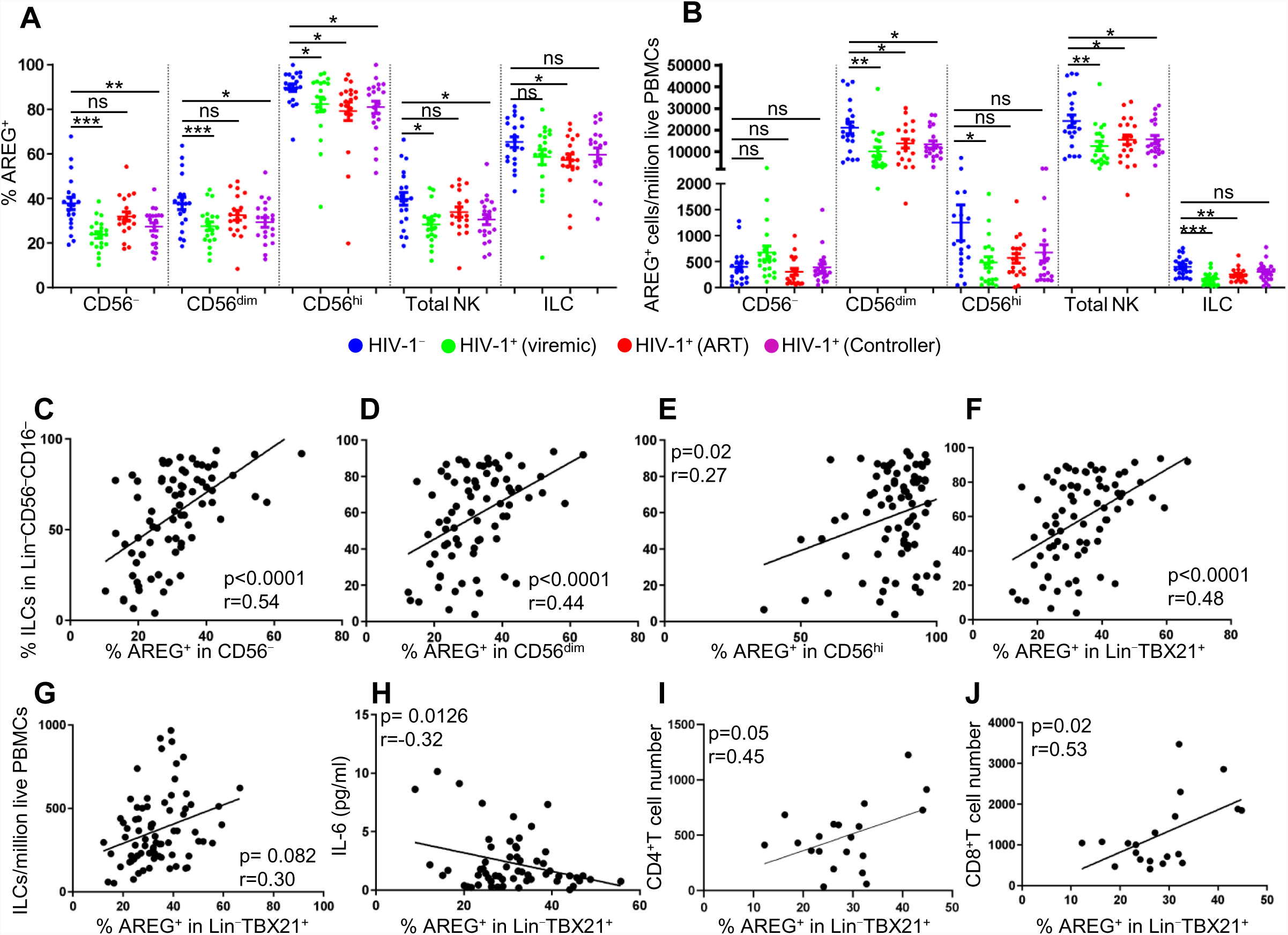
AREG^+^NK cells in people living with HIV-1. (A, B) PBMCs from HIV-1^−^ (n=20), HIV-1^+^ viremic (n=20), HIV-1^+^ ART suppressed (n=19), and HIV-1^+^ spontaneous controllers (n=20) were stimulated with PMA and ionomycin in RPMI 1640 at 37 °C in 5% CO_2_ for 3 hrs. The percentage (A) and the number (B) of AREG^+^ cells from are shown for the indicated populations. (C-F) Correlation of the percentage of AREG^+^ cells in CD56^−^ (C), CD56^dim^ (D), CD56^hi^ (E) or total NK cells (F), from (A), with the percentage of ILCs. (G) Correlation of the percentage of AREG^+^ cells from total NK cells with the number of ILCs. (H) Correlation of total AREG^+^NK cells with the plasma IL-6 in people living with HIV-1 (n=59). (I, J) Correlation of total AREG^+^NK cells with the plasma CD4^+^ (I) or CD8^+^ (J) T cell numbers in HIV-1^+^ viremic people (n=20). (C-J), correlation coefficient (r) by Pearson, zero slope p value determined by the F-test. (A, B), Two-tailed unpaired *t*-test. ns, not significant, *p<0.05, **p<0.01, ***p<0.001. (A-J), cohort described in Supplemental Table 7.

Interestingly, in people living with HIV-1, the percentage of AREG^+^NK cells correlated with the percentage and the number of ILCs (Figure 6C-6G), and there was inverse correlation with the plasma concentration of inflammatory cytokine IL-6 (Figure 6H). In people living with HIV-1 who are viremic, the number of CD4^+^T cells and CD8^+^T cells correlated with the AREG-positive fraction among total NK cells (Figure 6I and 6J). Though a trend was evident, statistical significance was not reached when the number of AREG^+^CD56^dim^NK or AREG^+^CD56^hi^NK cells was compared with CD4^+^T cells, perhaps due to the limited number of samples analyzed (Supplemental Figure 5A).

### CD56^dim^NK cells become CD56^−^ in the absence of IL-2-producing CD4^+^ T cells

The experiments above demonstrated that the majority of Lin^−^CD45^+^CD56^−^PBMCs are either ILCs or CD56^−^NK cells (Figure 1F). The stability of these innate lymphoid populations can be perturbed by autoimmune diseases or by the inflammation that accompanies viral infection (6, 11, 12). In people living with HIV-1, ILCs are permanently depleted (Figure 7A) (6, 13), and this reduction in ILCs correlated inversely with expansion of CD56^−^NK cells (Figure 7B, Supplemental Figure 5B and Supplemental Table 7). Additionally, among people living with HIV-1, CD56^−^NK cell expansion was greatest in HIV-1^+^ people who were not on ART and had persistent viremia (Figure 7C, Supplemental Figure 5C and Supplemental Table 7) (19).

**Figure 7.**
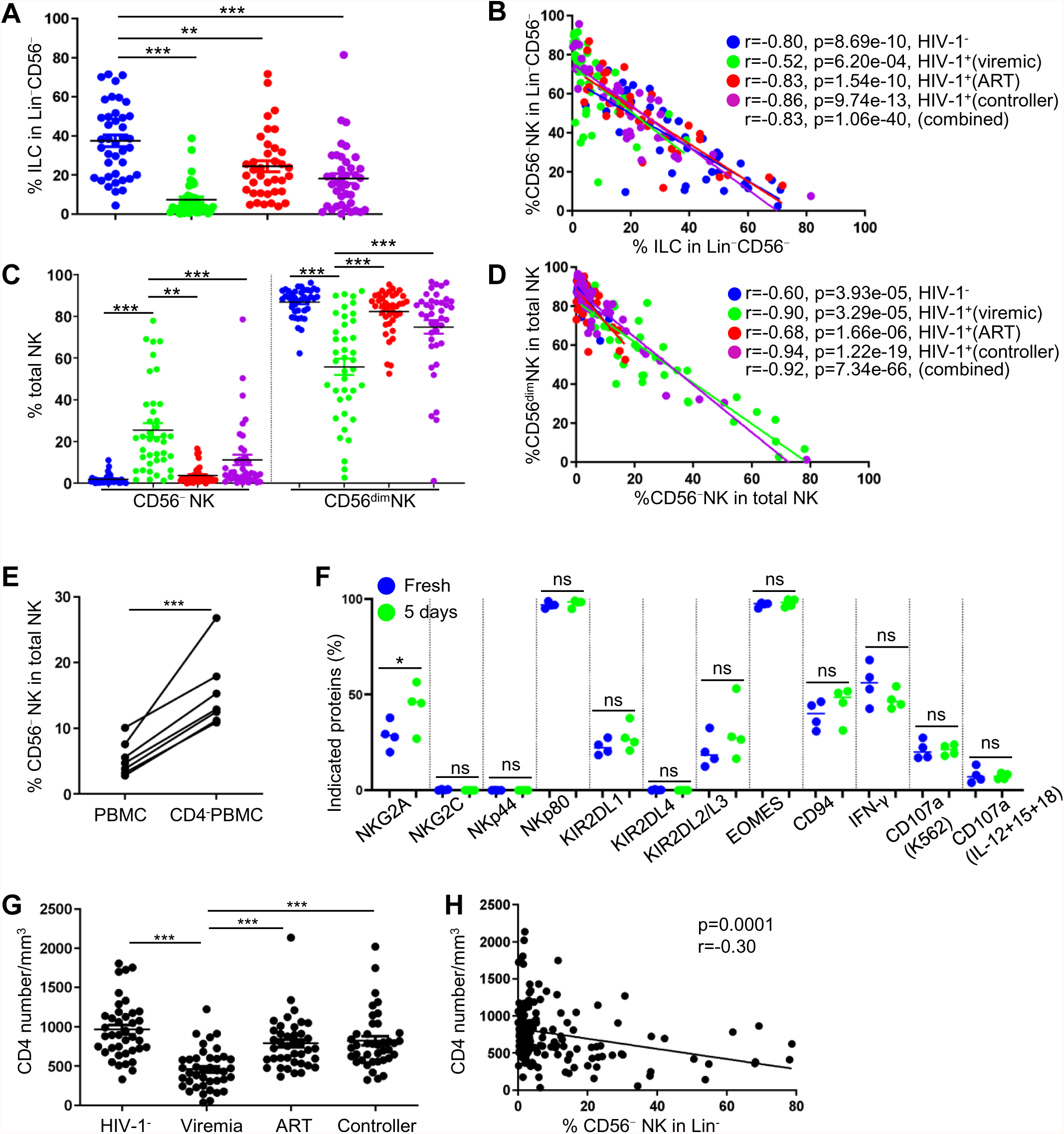
CD4^+^T cells maintain CD56^dim^ NK cells. (A) Percentage of ILCs among Lin^−^CD56^−^ cells from HIV-1-negative (n=40), HIV-1^+^ viremic (n=40), HIV-1^+^ ART suppressed (n=37), and HIV-1^+^ spontaneous controllers (n=40). (B) Correlation of ILCs and CD56^−^NK cells in the Lin^−^CD56^−^ population from (A). (C) Percentage of CD56^−^ and CD56^dim^ NK cells among total NK cells from the indicated groups (n=40 for each group). (D) Correlation between CD56^−^ and CD56^dim^NK cells from (C). (E) Percentage of CD56^−^NK cells among total NK cells from PBMCs and CD4^−^PBMCs after culture in RPMI 1640 at 37°C in 5% CO_2_ for 5 days (n=7). (F) Percentage of CD56^−^NK cells bearing the indicated proteins by flow cytometry, after detection in fresh PBMCs or after 5 days of culturing CD4^+^T cell-depleted PBMCs (n=4). (G) CD4^+^T cell number (counts/mm^3^) from the indicated groups (n=40 for each group). (H) Correlation between numbers of CD4^+^ T cells and CD56^−^NK cells (n=160). Data are mean ± s.e.m., (A, C, G), two-tailed unpaired *t*-test. (E), two-tailed paired *t*-test. **p<0.01, ***p<0.001. For (E), data were derived from healthy donors. For (A-D, G, H), the cohort of people living with HIV-1 was described in Supplemental Table 7.

The close similarity between CD56^−^NK cells and CD56^dim^NK cells revealed by our cell clustering algorithm (Figure 3), along with the inverse correlation in the numbers of these two cell populations (Figure 7D), suggested that CD56^dim^NK cells give rise to CD56^−^NK cells in the context of HIV-1 infection (Supplemental Figure 5D). To identify an experimental condition in tissue culture under which CD56^dim^NK cells give rise to CD56^−^NK cells, and to determine whether a specific cell type among PBMCs stabilizes CD56^dim^NK cells, PBMCs from HIV-1^−^ blood donors were maintained in culture for 5 days after selective depletion of either T cells (anti-CD3), CD4^+^T cells (anti-CD4), CD8^+^ T cells (anti-CD8), B cells (anti-CD19 and -CD20), monocytes and macrophages (anti-CD14 and -CD11b), stem cells (anti-CD34), myeloid cells (anti-CD33), or dendritic cells (anti-DC-SIGN and anti-BDCA3). Depletion of CD3^+^T cells, or of CD4^+^T cells, but not of any of the other cell types, increased the proportion of CD56^−^NK cells among Lin^−^TBX21^+^ cells in the cultures (Figure 7E and Supplemental Figure 6A and 6B). When CD56^−^NK cells from freshly isolated PBMCs were compared with CD56^−^NK cells isolated after 5 days of culturing PBMCs depleted of CD4^+^T cells, the levels of NKG2C, NKp44, NKp80, KIR2DL1, KIR2DL4, KIR2DL2/L3, EOMES, and CD94 were comparable (Figure 7F), as were IFN-γ production and degranulation in response to various stimuli and experimental conditions (Figure 7F), indicating that CD56^−^NK cells from these two sources are phenotypically and functionally similar. Consistent with this *ex vivo* experiment, CD56^−^NK cells were maximally increased in HIV-1^+^ individuals who were not treated with ART, in whom CD4^+^T cells were most severely depleted (Figure 7G and Supplemental Table 7). This observation was further supported by the inverse correlation between the numbers of CD56^−^NK cells and CD4^+^ T cells (Figure 7H).

Since CD4^+^T cells maintain NK cell physiology by secreting IL-2 (57, 58), the effect of IL-2 was tested next. Addition of exogenous IL-2 to the PBMC cultures that had been depleted of CD4^+^T cells maintained the stability of CD56^dim^NK cells, and prevented the increase of CD56^−^NK cells (Figure 8A, 8B and Supplemental Figure 6C). Similar effect was also observed with IL-15 (Figure 8C). The effect of IL-2 was counteracted by anti-IL-2 antibody (Figure 8B, 8D and Supplemental Figure 6D) or by inhibition of IL-2 signaling by STAT3 or JAK3 inhibitors, without compromising cell survival (Supplemental Figure 6E and 6F). This inhibition was not seen with inhibitors of STAT6, AKT, or ERK1/2 (Supplemental Figure 6E and 6F). Additionally, TGF-β1 which suppresses IL-2 production by CD4^+^T cells during HIV-1 infection (50–53, 59), increased the percentage of CD56^−^NK cells and decreased IFN-γ production by NK cells (Figure 8E and 8F); both effects of TGF-β1 were counteracted by exogenous IL-2 (Figure 8E and 8F).

**Figure 8.**
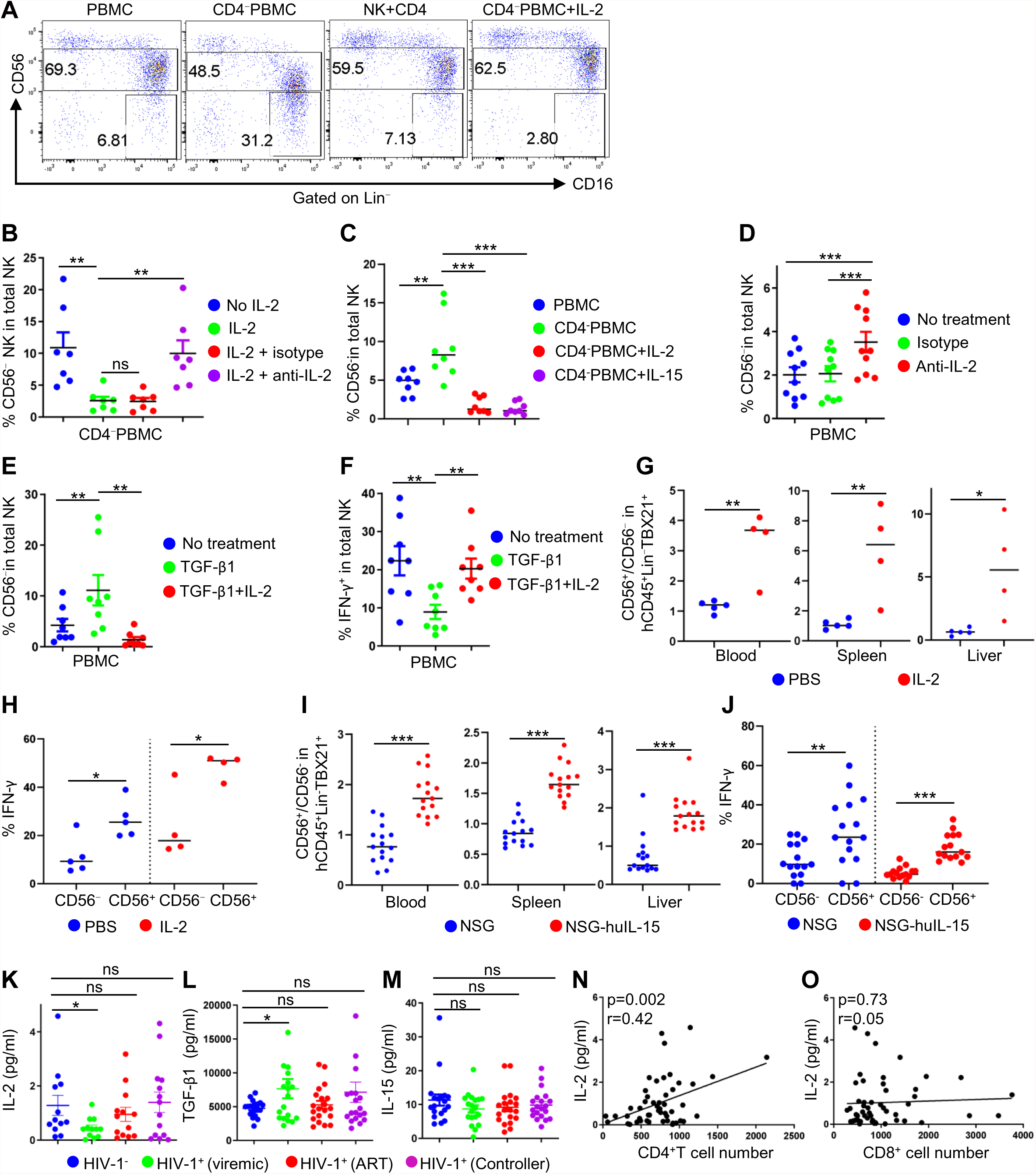
Effect of CD4^+^T cells and IL-2 on NK cell populations. (A) CD56^dim^ and CD56^−^ NK cells were detected after 5 days culture in RPMI 1640 at 37°C in 5% CO_2_ under the indicated conditions. (B) Percentage of CD56^−^NK cells among total NK cells from CD4^−^PBMCs cultured as in (A), in the presence or absence of IL-2 (10 ng/ml) or IL-2 neutralizing antibody (4 ug/ml) (n=7). (C) Percentage of CD56^−^NK cells from PBMCs or CD4^−^PBMCs cultured as in (A) in the presence or absence of IL-2 (10 ng/ml), or IL-15 (10 ng/ml) (n=8). (D) Percentage of CD56^−^NK cells among total NK cells in PBMCs after culturing as in (A) with isotype control or IL-2 neutralizing antibody (4 ug/ml) (n=10). (E, F) Percentage of CD56^−^NK (E) or IFN-γ^+^ NK (F) cells among total NK cells after culturing PBMCs as in (A) with TGF-β1 (50 ng/ml) or TGF-β1 (50 ng/ml) + IL-2 (20 ng/ml) (n=8). (G, H) NSG mice reconstituted with human CD34^+^ hematopoietic stem cells were injected intravenously with PBS (n=5) or with hIL-2-rAAV (n=4). Human NK cells from blood, spleen and liver were assessed 6 weeks later for CD56 (G) or IFN-γ production after 16 hrs stimulation with IL-12, IL-15, and IL-18 (H). (I, J) NSG (n=15) or human IL-15 transgenic NSG (NSG-huIL-15) mice (n=15) were reconstituted with human CD34^+^ hematopoietic stem cells for 6 weeks. Blood, spleen, and liver were harvested for detection of CD56 (I) or IFN-γ production after 16 hrs stimulation with IL-12, IL-15, and IL-18 (J). (K-L) Plasma concentration of IL-2 (HIV-1^−^ (n=12), viremic (n=12), ART suppressed (n=13), controllers (n=14)) (K), TGF-β1 (HIV-1^−^ (n=20), viremic (n=19), ART suppressed (n=20), controllers (n=20)) (L) and IL-15 (HIV-1^−^ (n=19), viremic (n=19), ART suppressed (n=20), controllers (n=20)) (M), detected by Luminex. (N, O), The correlation of plasma IL-2 with numbers of CD4^+^ T cells (N) or CD8^+^T cells (O) (n=51). Data are mean ± s.e.m., (B-F), two-tailed paired *t*-test. (G-M), two-tailed unpaired *t*-test. ns, not significant, *p<0.05, **p<0.01, ***p<0.001. For (A-F), data were derived from healthy donors. For (K-O), cohort was described in Supplemental Table 7.

To test the importance of IL-2 in an *in vivo* model, NOD-scid Il2rg^null^ (NSG) mice reconstituted with human CD34^+^ hematopoietic stem cells were injected intravenously with PBS, or with recombinant adeno-associated virus expressing IL-2 (hIL-2-rAAV). 6 weeks later, the ratio of CD56^+^ to CD56^−^ NK cells was greater in animals injected with hIL-2-rAAV than in controls (Figure 8G and Supplemental Figure 6G), and the NK cells from the hIL-2-rAAV-treated mice showed higher IFN-γ production than did cells from the control mice (Figure 8H and Supplemental Figure 6H). The importance of IL-15 was also tested *in vivo* by reconstituting NSG and human IL-15 transgenic NSG (NSG-huIL-15) mice with human CD34^+^ hematopoietic stem cells for 6 weeks. As compared with human NK cells generated in NSG mice, human NK cells from NSG-huIL-15 mice exhibited a higher ratio of CD56^+^ to CD56^−^ NK cells, and the CD56^+^NK cells produced more IFN-γ than did the CD56^−^NK cells (Figure 8I and 8J). Thus, IL-2 and IL-15 are each capable of maintaining functional CD56^dim^NK cells *in vivo*. To complement the above experiments, the plasma concentrations of IL-2, IL-15, and TGF-β1 were measured in people living with HIV-1. In people who were untreated and HIV-1 viremic, IL-2 levels were decreased and TGF-β1 levels were increased (Figure 8K and 8L). Additionally, the plasma IL-2 concentration correlated with the number of CD4^+^T cells, but not with CD8^+^T cells (Figure 8N and 8O). IL-15 levels were not detectably changed by HIV-1 infection (Figure 8M). These results are consistent with the clinical studies showing that treatment with exogenous IL-2 restored CD56^dim^NK cells in HIV-1^+^ viremic people (60), and suggest that a defective CD4^+^T cell–IL-2 axis, but not IL-15, is responsible for decreased stability of CD56^dim^NK cells in people living with HIV-1 infection.

### Metabolic difference between CD56^dim^ and CD56^−^NK cells

Comparison of bulk RNA-Seq data from CD56^dim^ and CD56^−^NK cells sorted from HIV-1 negative donors, revealed that genes associated with immune function (CD6, TRAF3, and IRAK2), and glycolysis and oxidative phosphorylation (NCAM1, MRPL24, ACAT2, B3GAT1, DGKK), were enriched in CD56^dim^NK cells (Figure 9A and Supplemental Table 8) (61). Genes regulating transcription, protein modification, and membrane trafficking (TXNDC5, PIGL, GORASP1, ARAP3 and AGAP1) were also enriched in CD56^dim^NK cells, whereas genes encoding proteins that inhibit AKT, G-protein signaling, transcription, and NK cell activation and survival (PRMT6, RGS1, ZBTB46, MED20, GNAQ) were enriched in CD56^−^NK cells (Figure 9A and Supplemental Table 8).

**Figure 9.**
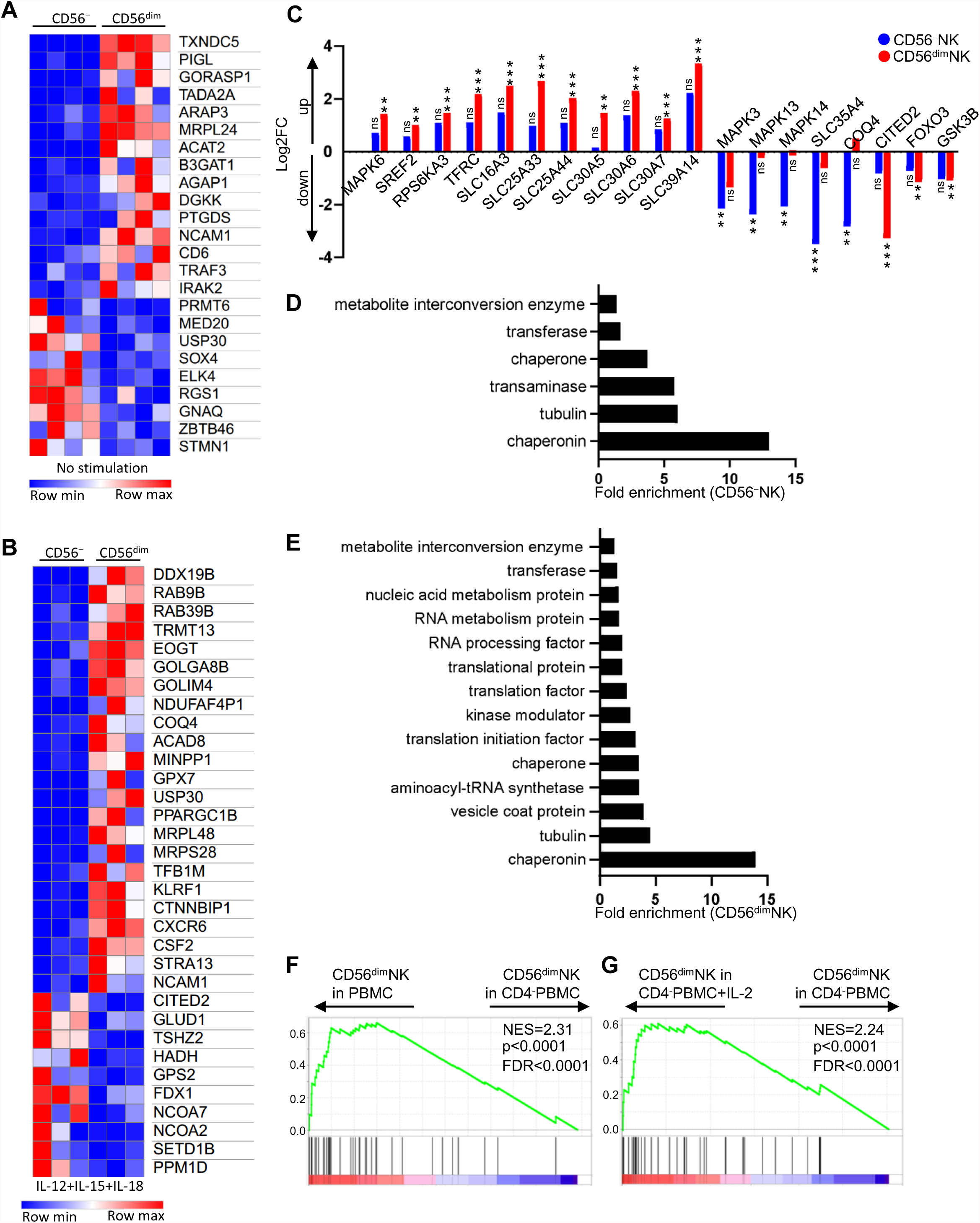
CD56^dim^NK cells are distinguished from CD56^−^NK cells by metabolic gene expression. (A, B) Heatmap of metabolism and immune related genes that were differentially expressed between CD56^−^ and CD56^dim^ NK cells directly sorted from PBMCs, (A) before (n=4), or (B) after IL-12, IL-15, and IL-18 stimulation in RPMI 1640 at 37°C in 5% CO_2_ for 16 hrs (n=3) (log2 fold change >1, p<0.01 determined by DESeq2). (C) Log2 fold change of mTOR signaling-related genes, showing cells stimulated with IL-12, IL-15, and IL-18 versus unstimulated CD56^−^ or CD56^dim^ NK cells. ns, not significant, **p<0.01, ***p<0.001. (D, E) Gene enrichment in CD56^−^ or CD56^dim^NK cells after stimulation with IL-12, IL-15, and IL-18. (F, G) GSEAs comparing the CD56^dim^NK cell expression signature in PBMCs (determined by RNA-seq, CD56^dim^ versus CD56^−^NK cells, Table S8) from sorted CD56^dim^NK cells in PBMC versus CD56^dim^NK cells in CD4^−^PBMC (F) or CD56^dim^NK in CD4^−^PBMC with IL-2 versus CD56^dim^NK in CD4^−^PBMC (G) after culture in RPMI 1640 at 37 °C in 5% CO_2_ for 5 days. All data were generated using blood from healthy HIV-1-negative donors.

Upon stimulation with IL-12, IL-15, and IL-18, as compared with CD56^−^NK cells, CD56^dim^NK cells had greater activation of genes involved in oxidative phosphorylation, translation, membrane trafficking, and immune activation, including DDX19B, RAB9B, RAB39B, TRMT13, GOLGA8B, NDUFAF4P1, COQ4, ACAD8, USP30, PPARGC1B, KLRF1, CXCR6, and STRA13 (Figure 9B and Supplemental Table 8). Of note, the mitochondrial deubiquitinase USP30, which is important for protecting against depolarization-induced cell death (62), was also upregulated by stimulation to a greater extent in CD56^dim^NK cells. In contrast, genes encoding proteins that inhibit AKT-mTOR activity and transcription (CITED2 and TSHZ2), and genes associated with fatty acid and cholesterol catabolism (HADH and FDX1), all features of functionally impaired NK cells, were enriched in CD56^−^NK cells (Figure 9B and Supplemental Table 8).

The findings above indicate that CD56^dim^NK cells exhibit higher metabolic fitness upon stimulation than do CD56^−^NK cells. Consistent with this, multiple activators and targets of mTOR were upregulated, including MAPK6, SREF2, RPS6KA3, TFRC and SLC transporters. mTOR inhibitors, such as CITED2, FOXO3, and GSK3B, were downregulated by stimulation in CD56^dim^NK cells (Figure 9C and Supplemental Table 8). Stimulation of CD56^−^NK cells downregulated mTOR activators or targets, such as MAPK3, MAPK13, MAPK14, SLC35A4 and COQ4 (Figure 9C and Supplemental Table 8). Biological processes that are critical for NK cell effector function were only enriched in CD56^dim^NK cells (Figure 9D, 9E and Supplemental Table 8).

To better assess the effects of CD4^+^T cells and IL-2 on cultured CD56^dim^NK cells, RNA-Seq was performed on CD56^dim^NK cells which had been sorted from 5 day cultures of PBMCs, PBMCs depleted of CD4^+^T cells, or PBMCs depleted of CD4^+^T cells, but supplemented with exogenous IL-2. Gene set enrichment analysis (GSEA) revealed that CD56^dim^NK cells from the full PBMC culture, or from the CD4^−^PBMC culture supplemented with IL-2, but not from the CD4^−^PBMC culture, were enriched for transcripts that are highly expressed in sorted CD56^dim^NK cells, as compared with sorted CD56^neg^NK cells from PBMC culture (Figure 9F, 9G and Supplemental Table 9). The convergence of RNA-Seq data from the freshly sorted cells, and the cells cultured under different conditions, indicates that maintenance of metabolically healthy and functional CD56^dim^NK cells is dependent upon IL-2-producing CD4^+^T cells.

### IL-2 and IL-15 prevents NK cell functional defects caused by mTOR inhibition

The above results indicate that CD56^dim^ and CD56^−^NK cells have distinct metabolic profiles, with the former having greater mTOR activity (Figure 9). Glycolysis, oxidative phosphorylation, and RNA and protein synthesis, all processes regulated by mTOR signaling, are required for NK cell killing activity and cytokine production (61, 63), and IL-2 induced AKT-mTOR activation is critical for NK cell effector function (64, 65). In fact, IL-2 treatment increased the activity of multiple mTOR components in NK cells from HIV-1 negative donors, as evidenced by enhanced phosphorylation of mTOR itself on Ser2448, AKT on Ser473, 4EBP1 on Thr36 and Thr45, and S6 on Ser235 and Ser236, and increased surface levels of the transferrin receptor CD71 (Figure 10A), the synthesis of which is known to be positively regulated by mTOR (66). Depletion of CD4^+^T cells from cultured PBMCs of HIV-1 negative people resulted in downregulation of these markers (Figure 10B), consistent with the role of CD4^+^T cells and IL-2 in maintaining metabolic fitness of NK cells (Figure 9F). IL-15 showed similar effect as IL-2 in enhancing mTOR signaling activity (Figure 10C).

**Figure 10.**
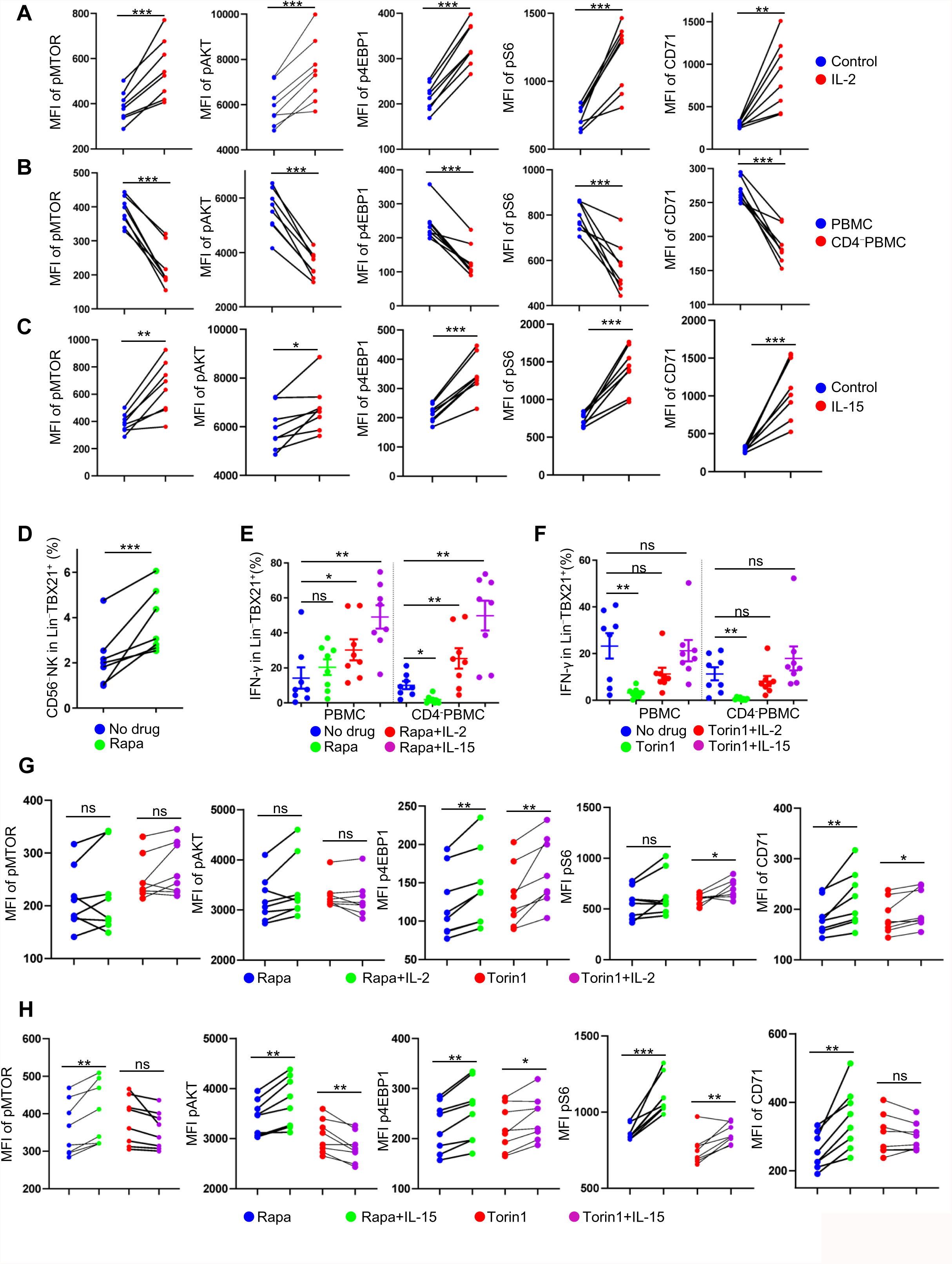
IL-2 overcomes the effects of CD4^+^T cell depletion or MTOR inhibition. (A) PBMCs were cultured at 37 °C in 5% CO_2_ with or without IL-2 (50 ng/ml) for 5 days, the phosphorylation of mTOR on Ser2448, AKT on Ser473, 4EBP1 on Thr36 and Thr45, and S6 on Ser235 and Ser236, and the surface CD71 from NK cells (Lin^−^TBX21^+^) were detected by flow cytometry (n=8). (B) As in (A), PBMCs or CD4^−^PBMCs were cultured for 5 days, the indicated targets from NK cells were detected (n=8). (C) As in (A), PBMCs were cultured with or without IL-15 (50 ng/ml) for 5 days, the indicated targets were detected (n=8). (D) PBMCs were treated with or without rapamycin (10 nM) in RPMI and cultured as in (A) for 5 days, the percentage of CD56^−^NK cells among the Lin^−^TBX21^+^ population is shown (n=7). (E, F) PBMCs or CD4^−^PBMCs were cultured in the presence or absence of rapamycin (10 nM) (F, n=8) or Torin 1 (250 nM) (G, n=8), with or without IL-2 (10 ng/ml) or IL-15 (10 ng/ml), for 5 days. Cells were then stimulated with IL-12, IL-15, and IL-18 for 16 hrs and IFN-γ was detected in Lin^−^TBX21^+^ cells. (G, H) PBMCs were cultured in the presence or absence of rapamycin (10 nM) (F) or Torin 1 (250 nM) combined with or without IL-2 (10 ng/ml) (G, n=8) or IL-15 (10 ng/ml) (H, n=8) for 5 days. The indicated targets from the Lin^−^TBX21^+^ population were detected as in (A). Data are mean ± s.e.m., two-tailed paired *t*-test. ns, not significant, *p<0.05, **p<0.01, ***p<0.001. All data were generated using blood from healthy HIV-1-negative donors.

When mTOR was inhibited by incubating PBMCs from HIV-1 negative people in rapamycin, the percentage of CD56^−^NK cells was increased (Figure 10D and Supplemental Figure 6I) and IFN-γ production by NK cells cultured without CD4^+^T cells was decreased (Figure 10E). Moreover, IL-2 or IL-15 treatment rescued IFN-γ production in the presence of rapamycin (Figure 10E). More potent inhibition by torin 1 (mTOR kinase inhibitor) led to impaired IFN-γ production by NK cells, even in the presence of CD4^+^T cells; however, this inhibition was overcome by addition to the culture of exogenous IL-2 or IL-15 (Figure 10F). Consistent with the above results, IL-2 or IL-15 treatment elevated the activity of multiple mTOR components and increased the level of CD71 in the presence of rapamycin or torin 1 (Figure 10G and 10H). Thus, NK cell functionality is dependent on mTOR, the activity of which can be maintained by IL-2 or IL-15.

## Discussion

While the vast majority of knowledge concerning ILCs and NK cells has come from studies in the mouse, significant species-specific differences have been detected in the development of innate lymphoid cells, the interrelatedness of their subsets, their gene regulation, and their function (27, 67). A detailed examination of the interrelationship between human NK cells and ILCs in the blood from uninfected people and in people living with HIV-1 is reported here.

Single cell RNA-Seq identified four discrete subpopulations of lineage-negative cells among human PBMCs: ILC2s, ILCPs, CD56^hi^NK cells, and a cluster which included both CD56^dim^ and CD56^−^NK cells (Figure 2C). Pseudotime analysis placed TBX21^+^CD56^hi^NK cells at a position between TBX21-negative ILCs/ILCPs and TBX21^+^CD56^dim/–^NK cells along the pseudotime trajectory (Figure 3F and 3G), revealing that CD56^hi^NK cells serve as a bridge connecting the phenotypically and functionally distinct ILCs and CD56^dim/–^NK cells. This is consistent with the fact that ILCPs not only have the potential to develop into different subsets of ILCs but also to develop into NK cells (10). An exact ILCP homolog with the same potential to develop into ILCs or NK cells has not been reported in the mouse (27).

In addition to bearing features associated with ILCs, including higher expression of IL7R, TCF7, and cytokines (Supplemental Figure 2A) (6), CD56^hi^NK cells are also distinguished from CD56^dim/–^ NK cells by their expression of GZMK (Figure 3B), but otherwise having lower levels of expression of cytotoxicity-related genes such as GZMB and GZMH (Supplemental Figure 2A). Interestingly, GZMK^+^CD8^+^T cells are distinguished from other CD8^+^ T cells by higher expression of IL7R, TCF7, and cytokines, and these cells have lower levels of GZMB, GZMH and decreased cytotoxic activity (68). Thus, GZMK^+^CD8^+^T cells appear to be the adaptive counterpart of CD56^hi^NK cells, representing an intermediate state between cytokine producing CD4^+^T cells and cytotoxic CD8^+^T cells.

Among people living with HIV-1 infection the biggest effect on gene expression in the blood innate lymphocyte subsets was detected in viremic individuals and was evident in all three NK cell subsets (Supplemental Table 6 and Figure 4B-4D), with more modest changes in people on ART or in elite controllers (Supplemental Table 6 and Figure 4B-4D). Each NK cell subset responded differently to HIV-1 infection. For example, genes important for NK cell function, including IL2RB, IL18RAP, SYK, FCER1G, IRF7, and SAT1, were dysregulated in CD56^−^NK cells, as compared with CD56^hi^ and CD56^dim^NK cells (Supplemental Table 6). HIV-1 infection is associated with permanent depletion of ILCs but not NK cells (6, 13). This greatly limits the yield of ILCs from blood and makes it difficult to dissect differential effects of HIV-1 on ILC2s from those on ILCPs. Nonetheless, bulk RNA-Seq on the mixed ILC population from people who were HIV-1 viremic indicated that genes associated with cell death pathways (CDKN1A, HIF1a, TPD52L1, and MADD) were enriched in ILCs (Figure 4I), but not in NK cells, providing a clue as to the mechanism by which HIV-1 depletes ILCs.

An important finding here was that human NK cells express AREG (Figure 5A, 5B and Supplemental Figure 4B), an epidermal growth factor family member that plays roles in tissue repair and immune tolerance (28). In humans, the AREG locus possesses multiple TCF7 binding sites and is open in NK cells. The mouse Areg locus does not have multiple TCF7 binding sites, and though mouse ILC2s express Areg, in mouse NK cells Areg is not expressed and the Areg locus is closed (Supplemental Figure 2B and 3B). The fact that the regulation of this gene is so different between the two species (Figure 5D) may explain why production of this homeostatic protein by NK cells has not been reported before. Interestingly, CD56^hi^NK cells from people living with HIV-1 who have undetectable HIV-1 in their blood in the absence of ART had elevated expression of MYDGF (Supplemental Figure 4A and Supplemental Table 6), a cytokine that prevents tissue damage by inhibiting recruitment of neutrophils and other leukocytes (43, 44). Like AREG, the MYDGF locus has binding sites for TCF7. Further investigation into the role of AREG and MYDGF production by NK cells may provide insight into the mechanisms that underlie the chronic inflammation and increased risk of cardiovascular disease in people living with HIV-1.

Unlike the production of IFN-γ in human NK cells, which occurs in response to 16 hrs stimulation with a mixture of inflammatory cytokines (Figure 10E and 10F), AREG protein production by NK cells occurred in response to stimulation with IL-2 alone, but for longer duration (Figure 5B and 5C). Perhaps this reflects gradual induction of TCF7/WNT or mTOR (Figure 10A and 10C) (6). Indeed, IL-2 treatment of NK cells for 5 days upregulated TCF7 protein and activated mTOR signaling (Figure 10A and 10C) (6). Pretreatment with a WNT agonist boosts AREG production by CD56^−^ and CD56^dim^NK cells but not by CD56^hi^NK cells and ILCs (Figure 5F and 5G); WNT signaling is already activated in CD56^hi^NK cells and ILCs (6) so treatment with the WNT agonist would not be expected to further increase AREG production in these cells. The locus-specific chromatin assays performed here demonstrate that TGF-β1 and RUNX3 negatively regulate AREG expression in human NK cells (Figure 5H and 5I). TGF-β1 is elevated in people living with HIV-1 (Figure 8L), inhibits IL-2 production by CD4^+^T cells (50–53), and AREG produced by NK cells was downregulated in people living with HIV-1 (Figure 6A and 6B). Perhaps effects on AREG expression explain in part the persistence of inflammation in the immunosuppressive environment associated with dysregulated TGF-β1.

It is well-established that the numbers of CD56^−^NK cells are increased in people living with HIV-1 who are viremic (18, 19) but the mechanism behind this effect has not been established. The importance of each PBMC cell type for NK cell phenotype was assessed *ex vivo*. Depletion of CD4^+^T cells alone resulted in decrease in CD56^dim^NK cells and concomitant increase in CD56^−^NK cells (Figures 7E, 8A, Supplemental Figure 6A and 6B), indicating that the former gave rise to the latter. Although macrophages play important roles in NK cell activation (69) depletion of CD14^+^ cells from the PBMCs that were put into culture did not increase the percentage of CD56^−^NK cells (Supplemental Figure 6A). The physiological relevance of this *ex vivo* assay was demonstrated by showing that CD56^−^NK cells generated by culturing in the absence of CD4^+^T cells have phenotype and function similar to their counterpart isolated directly from PBMCs (Figure 7F). Additionally, a series of experiments demonstrated that the basal levels of IL-2 secreted by the CD4^+^T cells in this *ex vivo* system was sufficient to maintain stable numbers of CD56^dim^NK cells (Figure 8B and 8C). Together with data on the effect of IL-2 in humanized mice (Figure 8G and 8H), and the fact that the increased numbers of these cells in HIV-1 infection correlate with decreases in plasma IL-2 (Figure 8K), the evidence support a model in which transition between CD56^−^ and CD56^dim^ NK cells is a dynamic process, explaining why even in HIV-1-negative blood donors some CD56^−^NK cells are detected (Figure 7C).

IL-15 could substitute for IL-2 in maintaining the stability of CD56^dim^NK cells in the ex vivo assay (Figure 8C, 8I and 8J). Unexpectedly, though, when combined with IL-12 and IL-18, IL-15 downregulated CD56 RNA (NCAM1) in CD56^dim^ and CD56^hi^NK cells, and downregulated CD56 protein in CD56^dim^NK cells (Supplemental Figure 6J and 6K). Nonetheless, the physiological role of IL-15 during HIV-1 infection is not clear, since no change in the plasma concentration of IL-15 was detected in people living with HIV-1 (Figure 8M). Finally, data generated with the *ex vivo* system revealed that, in response to IL-2 produced by CD4^+^T cells, mTOR within the NK cells is critical for maintaining the normal physiology of the CD56^dim^NK cell population (Figure 10).

## Acknowledgements

We thank the study participants who provided blood samples and their caretakers. M. Krone (UCSF) provided Institutional Review Board regulatory assistance, sample preparation, and record keeping. S. Maitland provided advice on the purification of programmable nuclease proteins. Ruijia Wang and Xue Wang assisted with data analysis. This research was supported by NIH grants R37AI147868 to J.L., R01HL150669 to S.A.W., R24OD026440, R01AI132963, and UC4DK104218 to M.A.B., F30HD100110 to N.J.S., and F31HL147482 to K.L. The UCSF-based SCOPE cohort was supported by the UCSF/Gladstone Institute of Virology and Immunology CFAR (P30 AI027763), and the CFAR Network of Integrated Systems (R24 AI067039). The content of this publication does not necessarily reflect the views or policies of DHHS, nor does the mention of trade names, commercial products, or organizations imply endorsement by the US Government.

## Author contributions

Y.W. and J.L. designed the experiments. Y.W. performed the experiments with assistance from L.L., N.J.S., E.M., K.L., P.S.L., M.A.B. and S.A.W.. Y.W. and J.L. analyzed the experimental data. Y.W., L.L., N.J.S. and J.L. analyzed the expression data. S.G.D. obtained and provided the clinical samples. Y.W. and J.L. wrote the manuscript, which was revised and approved by all authors.

## Methods

### Clinical samples

PBMCs of HIV-1^−^, HIV-1^+^ viremic, HIV-1^+^ on ART, HIV-1^+^, spontaneous controllers (viral load < 2,000 copies of HIV-1 gRNA/ml without ART) and HIV-1^+^ elite controllers (undetectable viral load without ART) used for flow cytometry, Luminex cytokine assay, and RNA-Seq, were from the University of California San Francisco SCOPE Cohort. The clinical characteristics of each individual in the SCOPE cohort was provided in Supplemental Table 7. All participants provided written informed consent for protocols that were included in study of cellular immunity in HIV-1 infection, in accordance with procedures approved by the University of Massachusetts Medical School (UMMS), and the University of California, San Francisco (UCSF) Institutional Review Boards. Consent was obtained from all participants the day of the procedure. All PBMCs from the SCOPE cohort were tested blindly with the code broken after analysis, and no samples were excluded from the analysis.

### Human mononuclear cell isolation and NK cell enrichment

80 ml of RPMI-1640 (Gibco) was added to each human peripheral blood leukopak. This was overlaid on Lymphoprep (STEMSELL, 07851) and centrifuged at 500 x g at room temperature for 30 minutes. Mononuclear cells were washed 3 times with MACS buffer (0.5% BSA and 2 mM EDTA in PBS), and either used immediately, or frozen in FBS containing 10% DMSO. NK cells were negatively enriched using the EasySep™ Human NK Cell Isolation Kit (STEMCELL, 17955), according to the manufacturer’s instructions.

### Flow cytometry

Cells were first stained with the Live and Dead violet viability kit (Invitrogen, L-34963). To detect surface molecules, cells were stained in MACS buffer with antibodies for 30 min at 4°C in the dark. For cytokine or transcription factor detection, Cells were fixed and permeabilized, and intracellular molecules were stained with specific antibodies in permeabilization buffer, using reagents from the Foxp3 staining kit (eBioscience, 00-5523-00). Fluorescence Minus One (FMO) and isotype controls for the antibodies used in this study were shown in Supplemental Figure 7.

### Stimulation Conditions

Specific conditions for stimulation are listed here. For cytokine detection, all cells were cultured at 37 °C containing 5% CO2. Conditions for each experiment are listed here.

Figure 5A: PMA (81 nM) and ionomycin (1.34 uM) (1:500, eBioscience, 00-4970-03) in RPMI 1640 for 3 hrs.

Figure 5B and 5C: 50 ng/ml IL-2, 50 ng/ml IL-15, 10 ng/ml IL-12 or 50 ng/ml IL-18 for 5 days in MACS NK medium (130-114-429, MACS).

Figure 5F–5H: CHIR99021 (10 uM) with or without TGF-β1 (50 ng/ml) in RPMI 1640 for 48 hrs, then stimulated with PMA and ionomycin for 3 hrs.

Figure 5I: 10 ng/ml IL-12 + 50 ng/ml IL-15 + 50 ng/ml IL-18 in MACS NK medium for 16 hrs.

Figure 6A and 6B: Stimulated as Figure 5A.

Figure 8A-8D: Untreated, or 10 ng/ml IL-2, or 10 ng/ml IL-2 + 4 ug/ml isotype control, or 10 ng/ml IL-2 + 4 ug/ml IL-2 neutralizing antibody, or 10 ng/ml IL-15, or 4 ug/ml IL-2 neutralizing antibody, or isotype in RPMI 1640 for 5 days.

Figure 8E and 8F: Untreated, or 50 ng/ml TGF-β1, or 50 ng/ml TGF-β1 + 20 ng/ml IL-2 in RPMI 1640 for 5 days (E), then 10 ng/ml IL-12 + 50 ng/ml IL-15 + 50 ng/ml IL-18 in RPMI 1640 for 16 hrs (F).

Figure 8H and 8J: 10 ng/ml IL-12 + 50 ng/ml IL-15 + 50 ng/ml IL-18 in RPMI 1640 for 16 hrs.

Figure 10A-10C: Untreated, or 50ng/ml IL-2, or 50 ng/ml IL-15 for 5 days.

Figure 10D-10H: Untreated, or rapamycin (10 nM), or Torin (250 nM), or combined with 10 ng/ml IL-2 or 10 ng/ml IL-15 in RPMI 1640 for 5 days (G, H), then 10 ng/ml IL-12 + 50 ng/ml IL-15 + 50 ng/ml IL-18 in RPMI 1640 for 16 hrs (E, F).

### Sorting of NK cells and ILCs

PBMCs were stained with a panel of lineage markers (antibodies against CD3, CD4, TCRαβ, TCRγδ, CD19, CD20, CD22, CD34, FcεRIα, CD11c, CD303, CD123, CD1a, and CD14), CD56, CD16 and CD127 (Supplemental Table 1). The CD56^hi^, CD56^dim^, CD56^−^ and ILCs were sorted as indicated in Supplemental Figure 1A and 1D using BD FACSAria IIu. Enrichment of the sorted cell populations was confirmed by flow cytometry before initiation of downstream experiments.

### Bulk RNA-Seq Library preparation

The sequencing library was prepared using CEL-Seq2 (74). RNA from sorted cells was extracted using TRIzol reagent (ThermoFisher, 15596018). 10 ng RNA was used for first strand cDNA synthesis using barcoded primers (the specific primers for each sample were listed in Supplemental Table 10). The second strand was synthesized by NEBNext Second Strand Synthesis Module (NEB, E6111L). The pooled dsDNA was purified with AMPure XP beads (Beckman Coulter, A63880), and subjected to in vitro transcription (IVT) using HiScribe T7 High Yield RNA Synthesis Kit (NEB, E2040S), then treated with ExoSAP-IT (Affymetrix, 78200). IVT RNA was fragmented using RNA fragmentation reagents (Ambion), and underwent another reverse transcription step using random hexamer RT primer-5’-GCC TTG GCA CCC GAG AAT TCC ANN NNN N-3’ to incorporate the second adapter. The final library was amplified with indexed primers: RP1 and RPI1 or RPI2 (as indicated in Supplemental Table 10), and the beads purified library was quantified with 4200 TapeStation (Agilent Technologies), and paired-end sequenced on Nextseq 500 V2 (Illumina), Read 1: 15 cycles; index 1: 6 cycles; Read 2: 60 cycles. Libraries of CD56^hi^, CD56^dim^, CD56^−^NK cells and ILCs from people who were HIV-1^−^, or HIV-1^+^ and viremic, under ART, or elite controllers, were sent to Novogene for sequencing by NovaSeq.

### Library preparation for single cell RNA-Seq

The sequencing library was prepared by Single Cell 3’ Reagent Kits v2 (10xgenomics, 120234). Isolated cells were 2x washed and resuspended in 1xPBS containing 0.05% BSA. Cell number and viability were measured by Bio-Rad TC 20 cell counter, cell concentration was adjusted around 1000-1500 cells/ul (viability>90%). Single cell suspension was loaded onto Chromium Controller (10xGenomics) to participate 3000-6000 single cells into gel beads in emulsions (GEMs). Libraries were constructed according to the instruction of single cell 3’ reagent kits v2. The yield and the quality of amplified cDNA were checked using High Sensitivity D5000 ScreenTape on TapeStation (Agilent Technologies). The final library was amplified using PCR cycles determined by cDNA quantification, and the quality of the library was checked again by TapeStation. The sequencing depth was controlled at more than 50,000 reads per cell. Sorted CD56^−^, CD56^dim^, CD56^hi^NK cells and ILCs from three donors were pooled and were sequenced as one sample, indexed using primers from the B2 position of the index plate.

### ATAC-Seq

Nuclei were precipitated after sorted CD56^hi^, CD56^dim^, CD56^−^ NK cells and ILCs were suspended in lysis buffer (10 mM Tris·Cl, pH 7.4; 10 mM NaCl; 3 mM MgCl_2_, 0.1% NP-40). 2.5 ul of Tn5 transposase (Nextera DNA Library Prep Kit, Illumina, FC-121-1030) was added per 50 ul reaction, for 30 min at 37°C. Released DNA fragments were purified by PCR product purification kit (Promega, A9282) and used to generate libraries. The barcode primers and common primers used for each sample from 2 donors were listed in Supplemental Table 10. Libraries were paired-end sequenced on Nextseq 500 V2 (Illumina) using Read 1: 42 cycles; Index 1: 8 cycles, and Read 2: 42 cycles.

### Bulk RNA-Seq Processing and Analysis

The pooled reads were separated by CEL-Seq2 barcodes and mapped to the hg19 genome using Tophat (75) (version 2.0.14, default parameters). ESAT (76) was used to quantify aligned reads using a transcript annotation file containing all RefSeq genes filtered to select only ‘NM’ transcripts, and extending the transcripts up to 1,000 bases past the end of the annotated 3’ end (*-wExt 1000, -task score3p*), multi-mapped reads (*-multimap ignore*) were discarded. DEBrowser was used to analyze the most varied genes, DESeq2 was used to perform differential expression analysis(77). Data were transformed using rlog within DEseq2, and prcomp was used to calculate the PCs for principal component analysis (PCA). Mouse RNA-seq data were downloaded from GSE77695, GSE109125, and GSE116092, and aligned to the mouse reference genome (mm10) using STAR (version 2.1.6) (78). Counts of reads aligned to RefSeq genes were quantified using RSEM (version 1.3.1) (79) and normalized using DEseq2.

### Single cell RNA-Seq processing and analysis

The Cell Ranger software package (10X Genomics) was used to perform sample demultiplexing, barcode processing, and single cell 3’ gene counting. Default settings were used. The filtered gene matrices generated by Cell Ranger were used as input into the open-source R package Seurat 3.0 (http://satijalab.org/seurat/) (80). FindNeighbors and FindClusters (using the default resolution of 0.6) were then run on the PCA reduction. FindMarkers was used to identify differentially expressed genes among clusters. Pseudotime analysis was done using monocle (version 2) (81). Cluster information (for color coding the cells for display) was imported from the Seurat analysis. The raw count data from the Seurat object was read into monocle and the standard workflow was followed: estimateSizeFactors, estimateDispersions, and detectGenes(min_expr=1) were called. Then, the dispersion table was calculated and subsetted to generate ordering genes with mean_expression >= 0.1 and dispersion_empirical >= dispersion_fit. The data was reduced to two dimensions and the cells were ordered (using the orderCells function). plot_cell_trajectory was used to create pseudotime trajectories and plot_genes_in_pseudotime (with its default settings, which rescales expression to relative values) used to show individual gene expression as a function of pseudotime.

### ATAC-Seq analysis

Paired-end reads were filtered with trimmomatic (version 0.32) (82), aligned with Bowtie2 (version 2.2.3) (83) to a reference genome hg19. The duplicates were removed using Picard’s MarkDuplicates (version 0.32). To be able to accurately call the peaks, each aligned read was first trimmed to the 9-bases at the 5’ end, the region where the Tn5 transposase cuts the DNA. To smooth the peaks, the start site of the trimmed reads were extended up and down stream for 10 bases. Peaks were called using these adjusted aligned reads with MACS2 (84). The adjusted aligned reads were converted to tdf files using IGVTools (IGVtools count –w5) (version 2.3.31) (85) for visualization. The Tn5 transposase accessible region was used as input for JASPAR motif analysis (http://jaspar.genereg.net/). Mouse ATAC-Seq data was downloaded from GSE77695, GSE100738, and GSE116091, and was analyzed as above using mm10 as the reference genome.

### Cas12a RNP-mediated knockout in NK cells

NK cells from PBMCs were isolated with EasySep™ Human NK Cell Isolation Kit (STEMCELL, 17955), and after culture in NK MACS Medium (MACS,130-114-429) for 7 days, the NK cells were ready for electroporation. Two Cas12a target sites (PAM underlined) within RUNX3 sequences were chosen for editing: 5’-<u>TTTC</u>ACCCTGACCATCACTGTGTTCAC-3’ and 5’-<u>CTTA</u>CCTCGCCCACTGCGGCCCACGAA-3’. The following Cas12a target site in adeno-associated virus integration site 1 (AAVS1) was used as a control chromosomal DNA target: 5’-<u>TTTA</u>TCTGTCCCCTCCACCCCACAGTG-3’. Chemically end-protected (Alt-R) AsCas12a crRNAs were synthesized for each target site by IDT. Enhanced AsCas12a (enAsCas12a) (86) coding sequence was engineered to include three nuclear localization signal sequences and cloned into a pet21a expression vector for expression in, and purification from *E. coli*, as previously described (54). For each electroporation, 200 pmol crRNA and 100 pmol recombinant enAsCas12a protein were mixed at room temperature for 20 min. 1 million NK cells were resuspended in 20 ul 4D nucleofector master mix (82% P3 + 18% supplement 1; Lonza, V4XP-3032) and then mixed with Cas12a RNPs for electroporation using program CM137 (87). The electroporated NK cells were ready for downstream experiments after culture in MACS medium for 5 days.

### *In vivo* treatment with IL-2

NOD-*Prkdc^scid^IL2rg^tm1Wjl^* (NSG) and NOD-*Prkdc^scid^IL2rg^tm1Wjl^ Tg(IL15)1Sz/SzJ* (NSG-Tg(Hu-IL15)) mice were purchased from The Jackson Laboratory (Bar Harbor, ME) and engrafted with human CD34^+^ hematopoietic stem cells derived from umbilical cord blood (UCB). UCB was obtained from donors that were consented under an approved IRB protocol at the UMass Memorial Medical Center, Department of General Obstetrics and Gynecology (Worcester, MA), and all samples used for engraftment were de-identified. UCB was processed as previously described and underwent CD3 T cell depletion (88). For engraftment, NSG or NSG-Tg(Hu-IL15) mice between 3 to 4 weeks of age received 100 cGy irradiation and were then injected IV with 100,000 CD34+ cells (89). Human immune system development was evaluated by flow cytometry at 6 weeks post-HSC injection.

The double-stranded (ds) adeno-associated virus (AAV) vectors were engineered and packaged as previously described (90). Briefly, full-length cDNA encoding human IL-2 were subcloned into a dsAAV plasmid (91) containing the murine preproinsulin II promoter. dsAAV vector packaging with serotype 8 capsid protein was produced by the Viral Vector Core at the University of Massachusetts Medical School Horae Gene Therapy Center (Worcester, MA, USA). HSC-engrafted NSG mice were intraperitoneally injected with 2.5 × 10^11^ particles of the purified AAV8-huIL-2 (AAV-IL-2). 6 weeks later, the blood, spleen, and liver were harvested for detection of CD56 or IFN-γ production from NK cells. NSG-Tg(Hu-IL15) were treated similarly except without injection of AAV vector.

## Statistical methods

Statistical test was performed using GraphPad Prism 9. The usage of paired or unpaired two-tailed student’s *t*-test was indicated in the figure legends. p<0.05 was considered as significant. Variance was estimated by calculating the mean ± s.e.m. in each group. Variances among groups of samples were compared using the F-test function in GraphPad Prism 9.

## Data and code availability

The data that support the findings of this study are available within the manuscript and in its supplementary information files. Bulk and single-cell RNA-Seq, and ATAC-Seq datasets generated by this study, can be found at: NCBI Gene Expression Omnibus (GEO): GSE168212, GSE203002. Areg expression and ATAC-Seq data generated by previous studies are from NCBI GEO:GSE77695, GSE109125, and GSE11609 (29–31).

