## Supplementary material for "Global profiling of human blood ILC subtypes reveals that NK cells produce homeostatic cytokine amphiregulin and sheds light on HIV-1 pathogenesis": Supp Figs 1-7

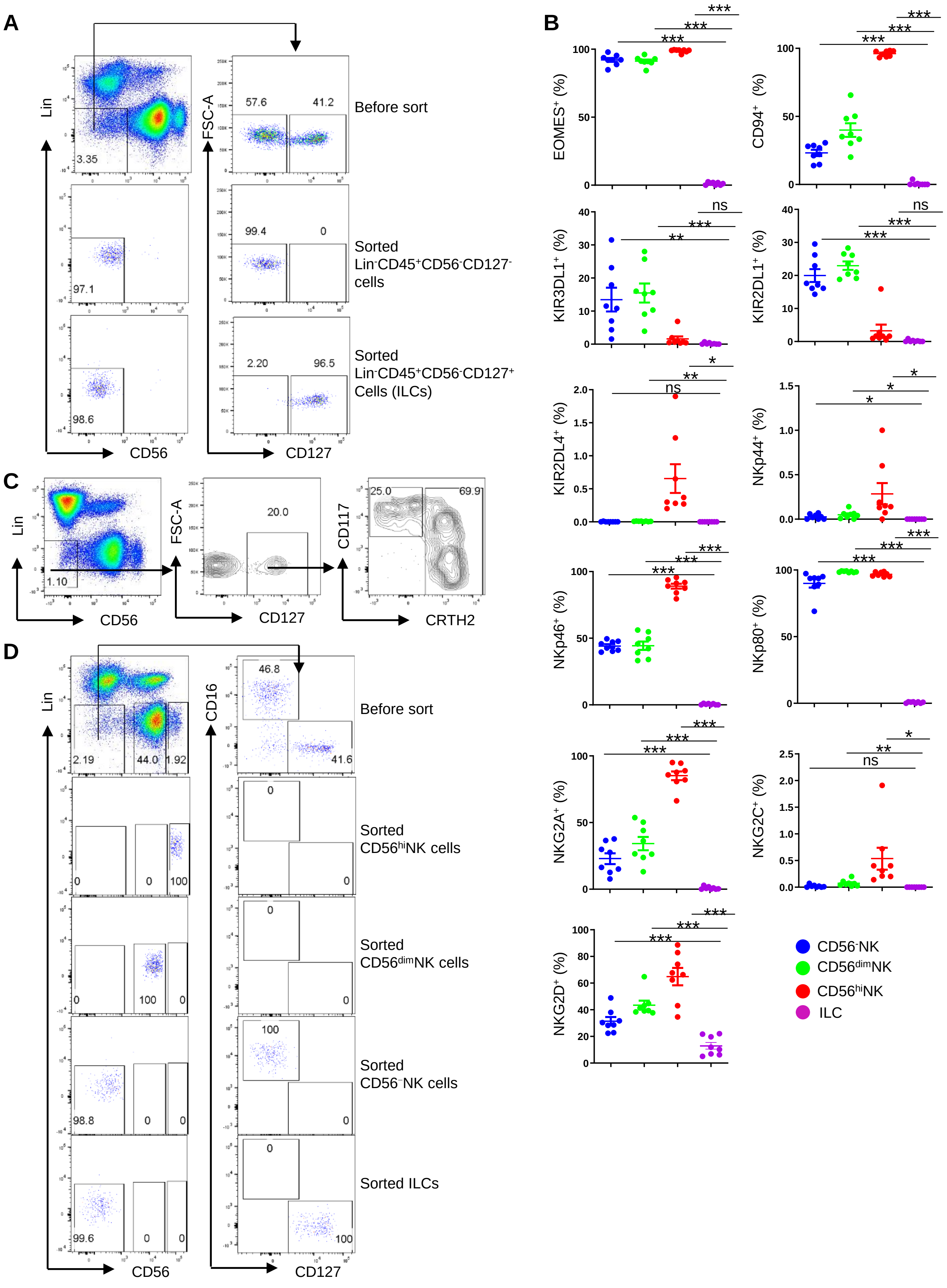
A
Lin
CD56
FSC-A
CD127
3.35
57.6
41.2
Before sort
97.1
99.4
0
Sorted
Lin-CD45+CD56-CD127-
cells
98.6
2.20
96.5
Sorted
Lin-CD45+CD56-CD127+
Cells (ILCs)
B
EOMES+ (%)
CD94+ (%)
KIR3DL1+ (%)
KIR2DL1+ (%)
KIR2DL4+ (%)
NKp44+ (%)
NKp46+ (%)
NKp80+ (%)
NKG2A+ (%)
NKG2C+ (%)
NKG2D+ (%)
***
**
*
ns
CD56-NK
CD56dimNK
CD56hiNK
ILC
C
Lin
CD56
1.10
FSC-A
CD127
20.0
CD117
CRTH2
25.0
69.9
D
Lin
CD56
CD16
CD127
2.19
44.0
1.92
46.8
41.6
Before sort
0
0
100
0
0
Sorted
CD56hiNK cells
0
100
0
0
0
Sorted
CD56dimNK cells
98.8
0
0
100
0
Sorted
CD56-NK cells
99.6
0
0
0
100
Sorted ILCs

**Supplemental Figure 1. Analysis of lineage negative PBMCs.**
(A) Sorting strategy for $Lin^{–}CD45^{+}CD56^{–}CD127^{–}$ and $Lin^{–}CD45^{+}CD56^{–}CD127^{+}$cells. (B) NK cell associated markers were detected in indicated populations by flow cytometry (n=8). (C) $Lin^{–}CD45^{+}CD56^{–}$PBMCs were detected with CD127, CD117 and CRTH2. (D) Sorting strategy for $CD56^{–}$, $CD56^{dim}$ and $CD56^{hi}$NK cells and ILCs. All data were generated using blood from healthy donors.

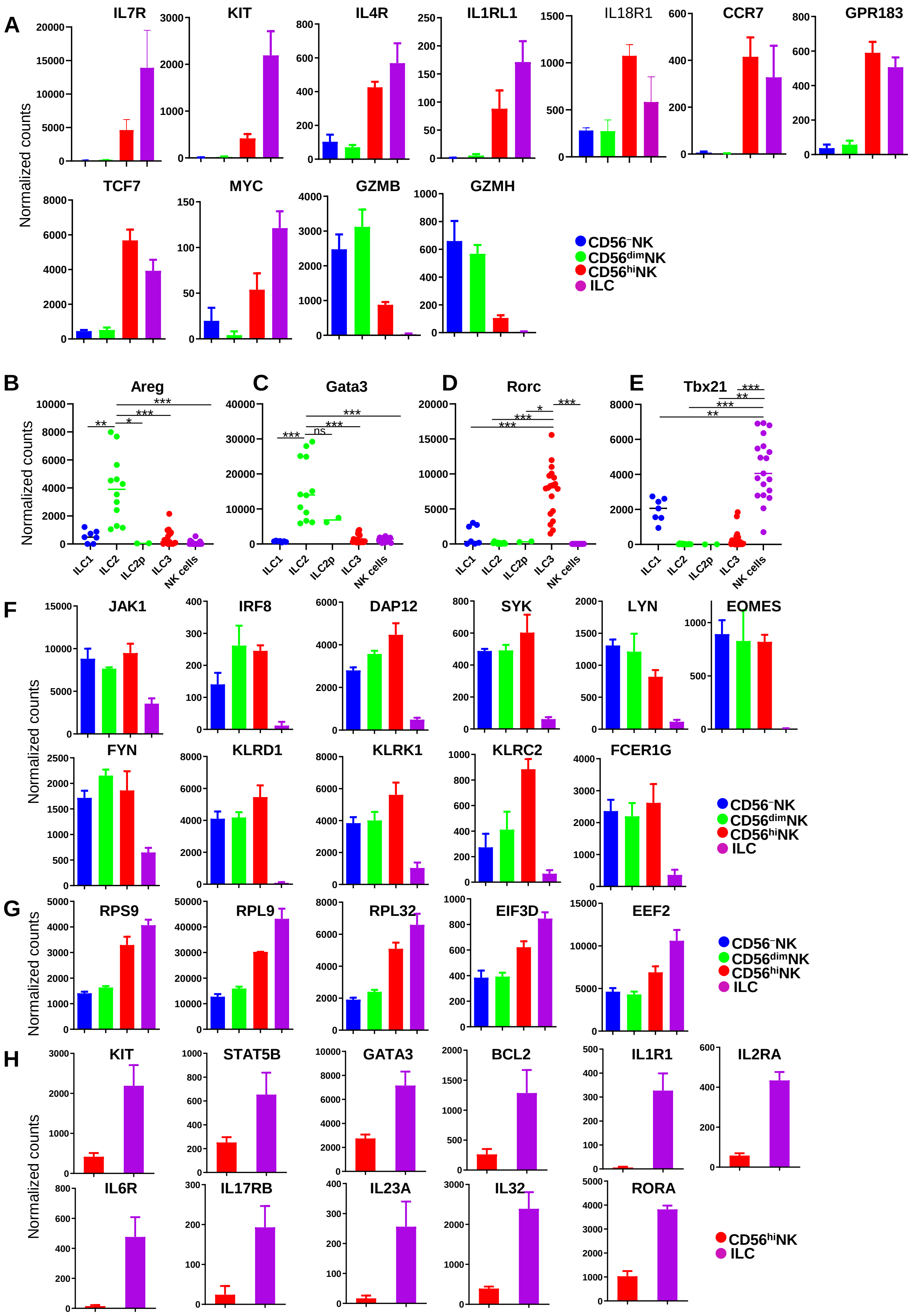
A
B
C
D
E
F
G
H
Normalized counts
IL7R
KIT
IL4R
IL1RL1
IL18R1
CCR7
GPR183
TCF7
MYC
GZMB
GZMH
Areg
Gata3
Rorc
Tbx21
ILC1
ILC2
ILC2p
ILC3
NK cells
JAK1
IRF8
DAP12
SYK
LYN
EOMES
FYN
KLRD1
KLRK1
KLRC2
FCER1G
RPS9
RPL9
RPL32
EIF3D
EEF2
STAT5B
GATA3
BCL2
IL1R1
IL2RA
IL6R
IL17RB
IL23A
IL32
RORA
CD56⁻NK
CD56dimNK
CD56hiNK
ILC

**Supplemental Figure 2. Representative transcripts of NK cells and ILCs**.

(A) Representative genes shared between CD56$^{hi}$NK cells and ILCs. (B-E) Normalized counts of Areg (B), Gata3 (C), Rorc (E), and Tbx21 (E), in mouse ILC1s (n=7), ILC2s (n=12), ILC2ps (n=2), ILC3s (n=19) and NK cells (n=19). Data are mean ± s.e.m., two-tailed unpaired *t*-test. *p<0.05, **p<0.01, ***p<0.001. (F) Enriched genes in all human NK cell subsets compared with ILCs. (G) Enriched genes in ILCs and CD56$^{hi}$NK cells compared with CD56$^{-}$ and CD56$^{dim}$NK cells. (H) Enriched genes in ILCs compared with CD56$^{hi}$NK cells. For (A, F-H)**,** Data were generated using blood from healthy donors. For (B-E), Mouse data was derived from GSE77695, GSE109125, and GSE116092. All counts were normalized by DESeq2.

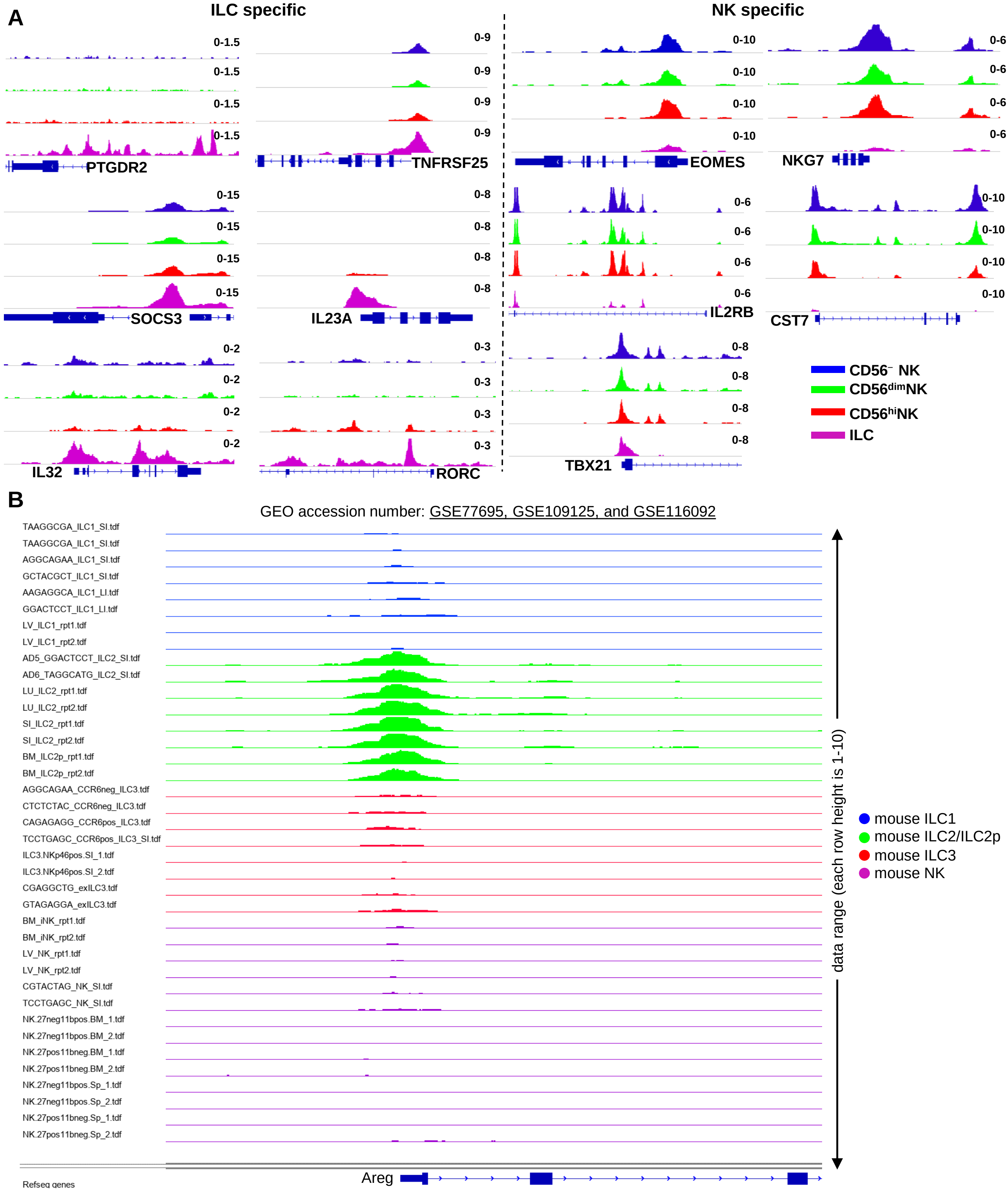


**Supplemental Figure 3. Chromatin accessibility in NK cells and ILCs.**
(A) ATAC-Seq signal of gene loci associated with human NK cells and ILCs from healthy donors. (B) ATAC-Seq signal from the Areg promoter region in mouse ILC1s, ILC2s, ILC2ps, ILC3s and NK cells, data was derived from GSE77695, GSE109125, and GSE116092.

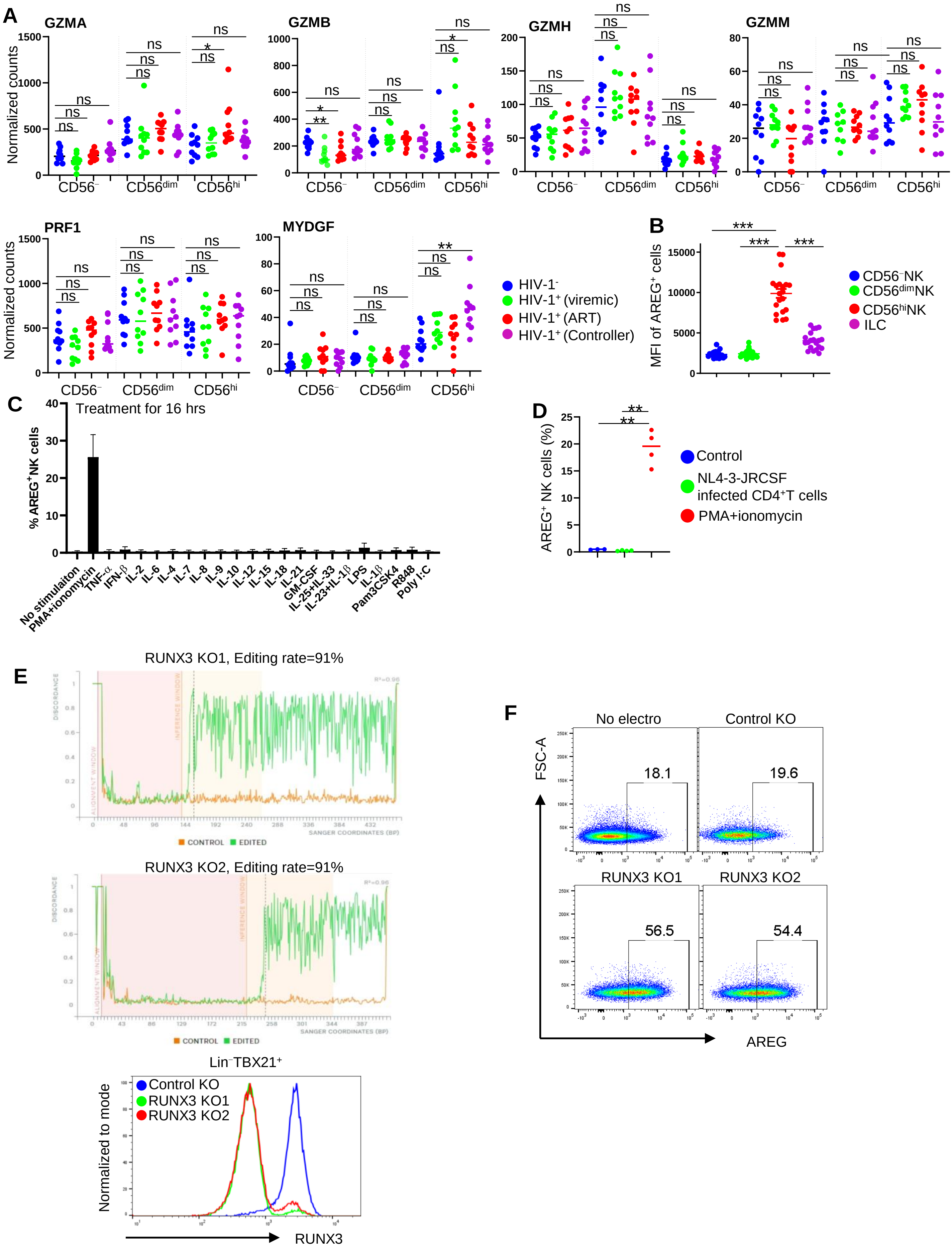
A
GZMA
GZMB
GZMH
GZMM
PRF1
MYDGF
Normalized counts
CD56⁻
CD56dim
CD56hi
ns
*
**
HIV-1⁻
HIV-1⁺ (viremic)
HIV-1⁺ (ART)
HIV-1⁺ (Controller)
B
MFI of AREG⁺ cells
***
CD56⁻NK
CD56dimNK
CD56hiNK
ILC
C
Treatment for 16 hrs
% AREG⁺NK cells
No stimulaiton
PMA+ionomycin
TNF-α
IFN-β
IL-2
IL-6
IL-4
IL-7
IL-8
IL-9
IL-10
IL-12
IL-15
IL-18
IL-21
GM-CSF
IL-25+IL-33
IL-23+IL-1β
LPS
IL-1β
Pam3CSK4
R848
Poly I:C
D
AREG⁺ NK cells (%)
Control
NL4-3-JRCSF infected CD4⁺T cells
PMA+ionomycin
E
RUNX3 KO1, Editing rate=91%
RUNX3 KO2, Editing rate=91%
DISCORDANCE
ALIGNMENT WINDOW
INFERENCE WINDOW
SANGER COORDINATES (BP)
CONTROL
EDITED
Lin⁻TBX21⁺
Normalized to mode
Control KO
RUNX3 KO1
RUNX3 KO2
RUNX3
F
No electro
Control KO
RUNX3 KO1
RUNX3 KO2
FSC-A
AREG
18.1
19.6
56.5
54.4

**Supplemental Figure 4. Human NK cells are the main producer of AREG.**

(A) Normalized counts of indicated genes from HIV-1$^{-}$ and different groups of HIV-1$^{+}$ individuals (n=10 for each group). (B) The MFI of AREG produced by indicated populations (n=20). (C) PBMCs from healthy donors were stimulated with indicated stimulus (PMA+ionomycin: 1:500, TNF-$\alpha$: 25 ng/ml, IFN-$\beta$: 25 ng/ml, IL-2: 50 ng/ml, IL-6: 50 ng/ml, IL-4: 1:50, IL-7: 50 ng/ml, IL-8: 50 ng/ml, IL-9: 50 ng/ml, IL-10: 50 ng/ml, IL-12: 50 ng/ml, IL-15: 50 ng/ml, IL-18: 50 ng/ml, IL-21: 50 ng/ml, GM-CSF: 1:50, IL-25+IL-33: 50 ng/ml each, IL-23+IL-1$\beta$: 50 ng/ml each, LPS: 100 ng/ml, IL-1$\beta$: 50 ng/ml, Pam3CSK4: 1 ug/ml, R848: 50 ng/ml, Poly I:C: 50 ng/ml) in RPMI 1640 at 37 °C in 5% $CO_2$ for 16 hrs, then AREG from Lin$^{-}$TBX21$^{+}$ cells was detected. (D) Beads purified NK cells were incubated with NL4-3-JRCSF infected CD4$^{+}$T cells or PMA+ionomycin in RPMI 1640 at 37 °C in 5% $CO_2$ for 3 hrs, AREG from NK cells were detected. (E) Detection of RUNX3 editing rate by ICE analysis (https://ice.synthego.com/) (upper 2 panels), and protein in NK cells by flow cytometry after control (AAVS1) or RUNX3 knockout (lower panel). (F) Detection of AREG$^{+}$NK cells from control or RUNX3 knockout groups after IL-12+IL-15+IL-18 stimulation in MACS NK medium at 37 °C in 5% $CO_2$ for 16 hrs. Data are mean ± s.e.m., *p<0.05, **p<0.01, ***p<0.001. (A, B, D), Two-tailed unpaired *t*-test. For (A), cohort was described in Supplemental Table 7.

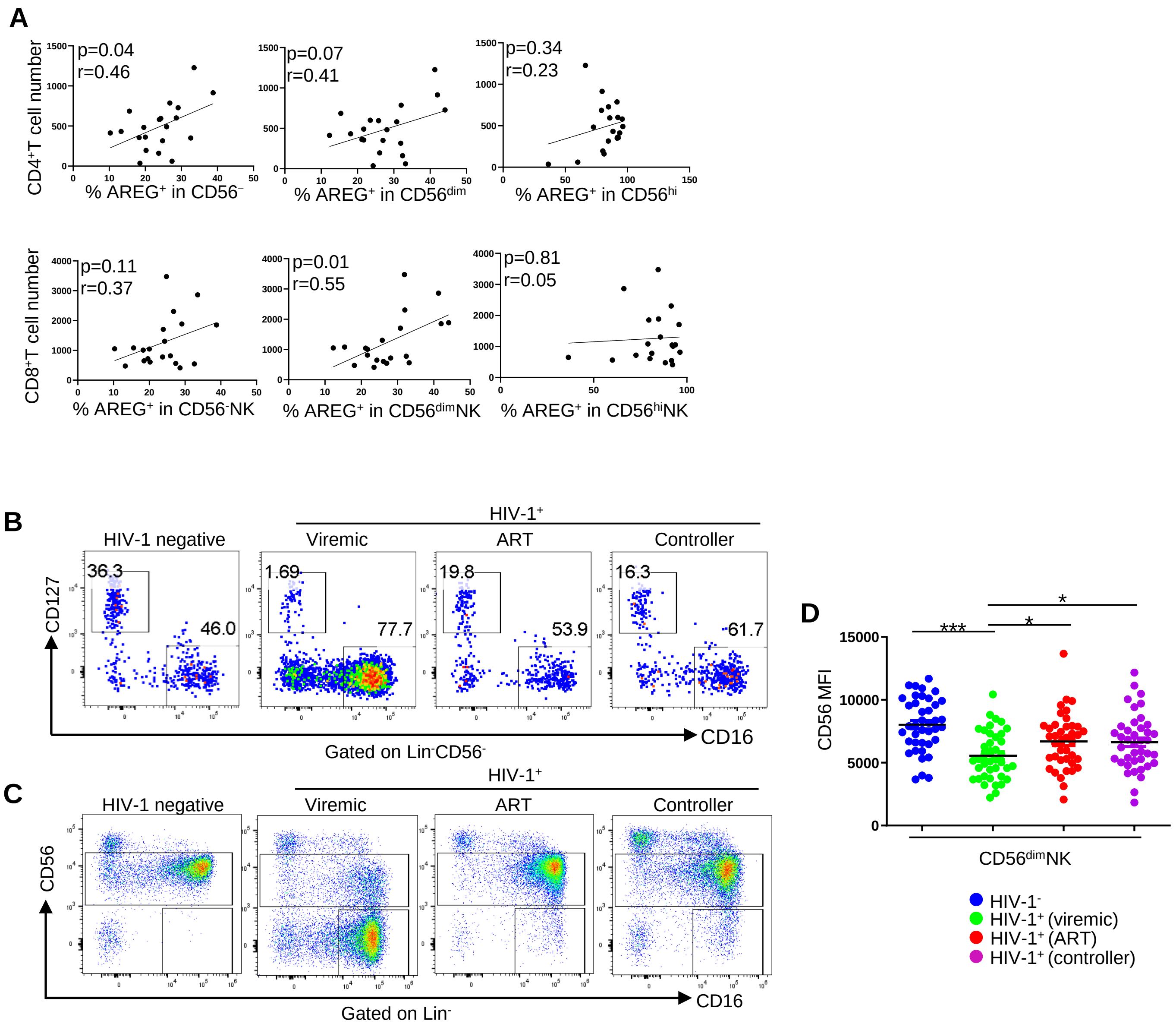


**Supplemental Figure 5. CD56⁻NK cells are expanded in HIV-1 infection.**
(A) The correlation of $AREG^+$ cells in different NK cell subsets with the number of $CD4^+T$ cells or $CD8^+T$ cells in HIV-$1^+$ viremic people (n=20). (B) Detection of ILCs and $CD56^-NK$ cells from indicated groups. (C) Detection of $CD56^{dim}$ and $CD56^-$ NK cells in indicated groups. (D) Detection of CD56 MFI in $CD56^{dim}NK$ cells in indicated groups (n=20 for each group). Data are mean ± s.e.m., *p<0.05, ***p<0.001. Cohort was described in Supplemental Table 7.

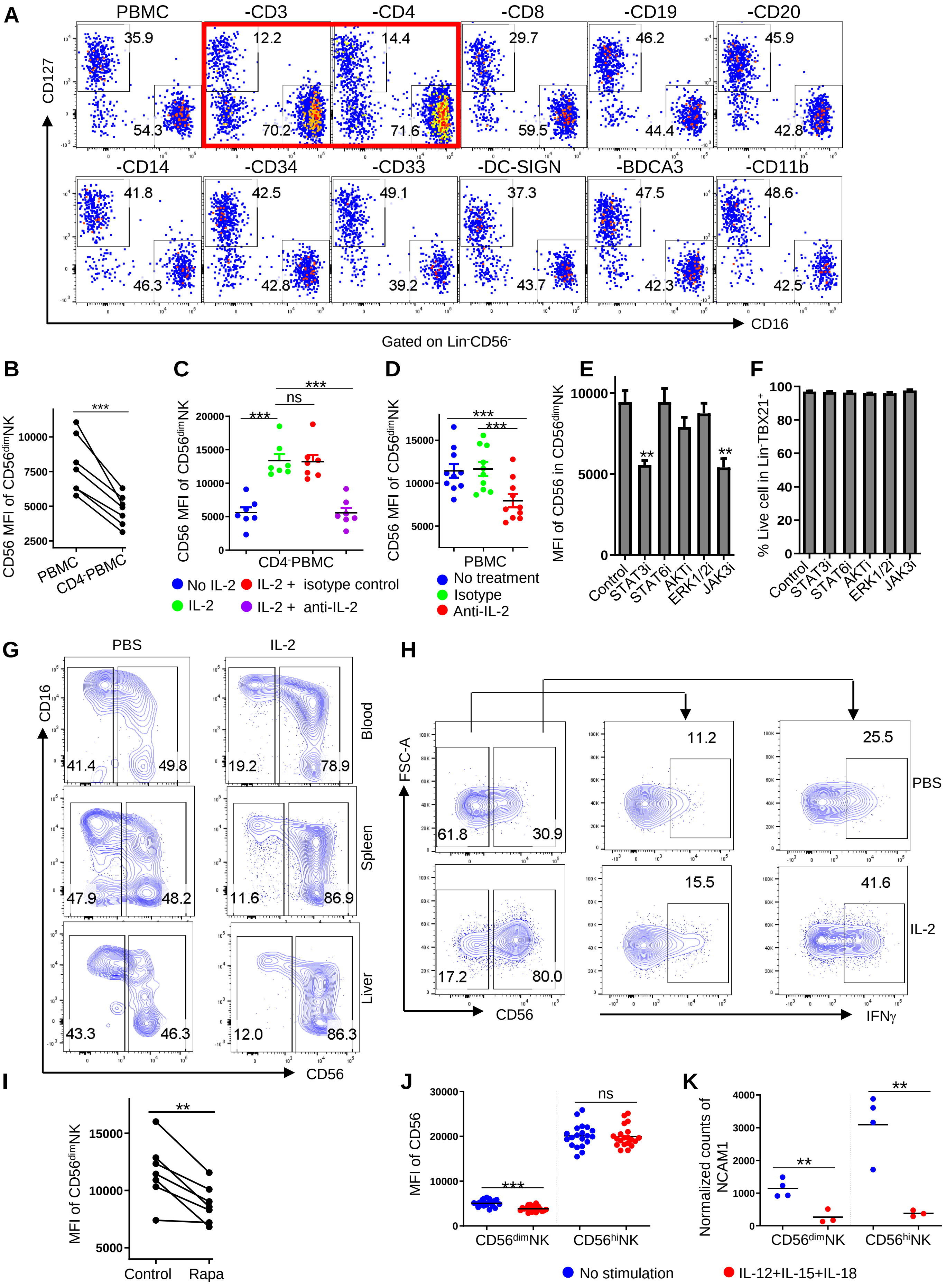
A
PBMC
-CD3
-CD4
-CD8
-CD19
-CD20
-CD14
-CD34
-CD33
-DC-SIGN
-BDCA3
-CD11b
CD127
CD16
Gated on Lin-CD56-
B
C
D
E
F
CD56 MFI of CD56dimNK
PBMC
CD4-PBMC
No IL-2
IL-2 + isotype control
IL-2
IL-2 + anti-IL-2
No treatment
Isotype
Anti-IL-2
MFI of CD56 in CD56dimNK
Control
STAT3i
STAT6i
AKTi
ERK1/2i
JAK3i
% Live cell in Lin-TBX21+
G
PBS
IL-2
Blood
Spleen
Liver
CD16
CD56
H
FSC-A
IFNγ
I
MFI of CD56dimNK
Control
Rapa
J
MFI of CD56
CD56dimNK
CD56hiNK
K
Normalized counts of NCAM1
No stimulation
IL-12+IL-15+IL-18

**Supplemental Figure 6. CD4$^+$T cells and IL-2 signaling are required for NK cell stability.**

(A) Specific cell types were depleted from PBMCs by magnetic beads based on surface markers as indicated, cultured for 5 days in RPMI 1640 at 37 °C in 5% $CO_2$, then Lin$^–$CD56$^–$PBMCs were detected with CD127 and CD16 by flow cytometry. (B) PBMC or CD4$^–$PBMCs were cultured as in (A), CD56 MFI in CD56$^{dim}$NK cells were detected (n=7). (C) CD56 MFI in CD56$^{dim}$NK cells was detected from CD4$^–$PBMCs in the presence or absence of IL-2 (10 ng/ml) combined with or without IL-2 neutralizing antibody (4 ug/ml) after 5 days culture as in (A) (n=7). (D) As cultured in (A), CD56 MFI in CD56$^{dim}$NK cells was detected from PBMCs after treatment with isotype or IL-2 neutralizing antibody (4 ug/ml) for 5 days (n=10). (E, F) As cultured in (A), CD56 MFI in CD56$^{dim}$NK cells (E) and live cell percentage in Lin$^–$TBX21$^+$ cells (F) were detected from PBMCs after treatment with c188-9 (STAT3i), AS-1517499 (STAT6i), MK-2206 (AKTi), Ravoxertinib (ERK1/2i) and CP690550 (JAK3i) for 5 days (n=4). (G, H) CD56 (G) or IFN-γ production (H) from human NK cells in humanized NSG mice were detected as Figure 8G and 8H. (I) As cultured in (A), PBMCs were treated with or without rapamycin (10 nM) for 5 days, the MFI of CD56 in CD56$^{dim}$NK cells was detected (n=7). (J) The CD56 MFI of CD56$^{dim}$ (n=20) and CD56$^{hi}$NK cells (n=20) from PBMCs stimulated with or without IL-12+IL-15+IL-18 for 16 hrs in RPMI 1640 at 37 °C. (K) The DESeq2 normalized counts of NCAM1 of sorted CD56$^{dim}$ (n=4) and CD56$^{hi}$NK cells (n=3) stimulated with or without IL-12+IL-15+IL-18 in RPMI 1640 at 37 °C for 16 hrs. Data are mean ± s.e.m., (B-F, I and J) two-tailed paired *t*-test, (K) two-tailed unpaired t-test. ns, not significant, $^{**}p<0.01$, $^{***}p<0.001$. All data were generated using blood from healthy donors.

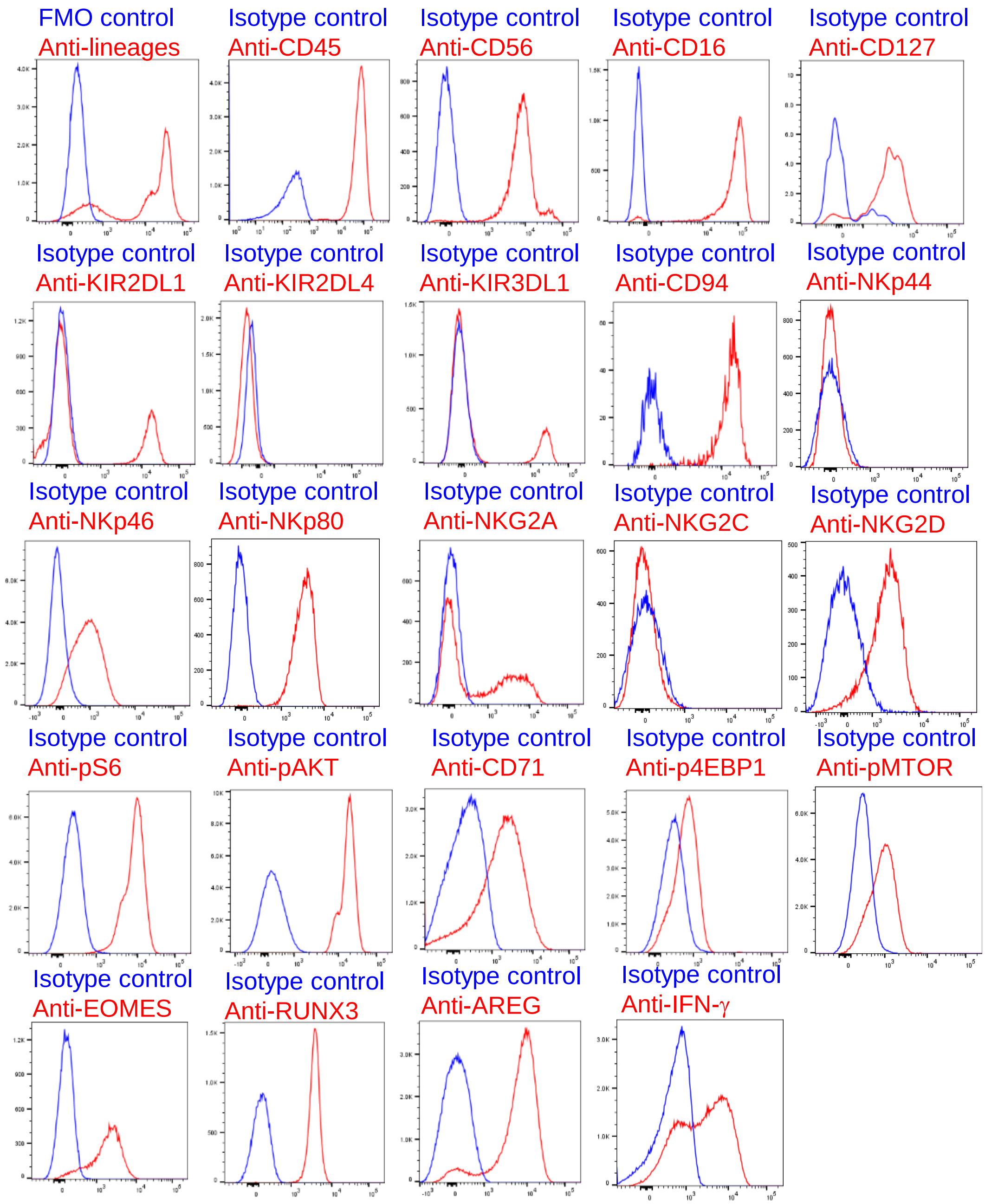


**Supplemental Figure 7. Confirming quality of antibodies used in this study.**
For lineage markers, PBMCs were stained with anti-lineage antibodies, FMO was used as negative control, signals from live singlets were detected. For CD45, PBMCs were stained with anti-CD45 antibody or isotype control, signals from live singlets were detected. For CD56, CD16, KIR2DL1, KIR2DL4, KIR3DL1, CD94, NKp44, NKp46, NKp80, NKG2A, NKG2C, NKG2D, pS6, pAKT, CD71, p4EBP1, pMTOR, EOMES and RUNX3, PBMCs were stained with specific antibodies or isotype controls, then signals from NK cells ($Lin^{-}TBX21^{+}$ or $Lin^{-}CD56^{+}$) were detected. For CD127, PBMCs were stained with anti-CD127 antibody or isotype control, signal from $Lin^{-}CD56^{-}CD16^{-}$ population was detected. For AREG and IFN-γ, PBMCs were stimulated with PMA+ionomycin in RPMI 1640 at 37 °C in 5% $CO_2$ for 3 hrs, then were stained with anti-AREG or anti-IFN-γ or isotype controls, signals from NK cells were detected. All PBMCs used were derived from healthy donors.
