## Supplementary material for "Global profiling of human blood ILC subtypes reveals that NK cells produce homeostatic cytokine amphiregulin and sheds light on HIV-1 pathogenesis": Suppl Tables 1-10

| Supplemental Table 1 | | |
|---|---|---|
| Reagents or resource | Source | Identifer |
| **Antibodies** | | |
| Anti-Human CD3 | BioLegend | Cat# 317306 Clone: OKT3, FITC |
| Anti-Human CD4 | BioLegend | Cat# 317408 Clone: OKT4, FITC |
| Anti-Human TCRα/β | BioLegend | Cat# 306706 Clone: IP26, FITC |
| Anti-Human TCRγ/δ | BioLegend | Cat# 331208 Clone: B1, FITC |
| Anti-Human CD19 | BioLegend | Cat# 302206 Clone: HIB19, FITC |
| Anti-Human CD20 | BioLegend | Cat# 302304 Clone: 2H7a, FITC |
| Anti-Human CD22 | BioLegend | Cat# 363508 Clone: S-HCL-1, FITC |
| Anti-Human CD14 | BioLegend | Cat# 325604 Clone: HCD14, FITC |
| Anti-Human CD34 | BioLegend | Cat# 343504 Clone: 581, FITC |
| Anti-Human FcεR1α | BioLegend | Cat# 334608 Clone: AER-37, FITC |
| Anti-Human CD1a | BioLegend | Cat# 300104 Clone: HI149, FITC |
| Anti-Human CD11c | BioLegend | Cat# 301604 Clone: 3.9, FITC |
| Anti-Human CD123 | BioLegend | Cat# 306014 Clone: 6H6, FITC |
| Anti-Human BDCA1 | BioLegend | Cat# 354208 Clone: 201A, FITC |
| Anti-Human RORγT | ebioscience | Cat# 12-6988-82 Clone: AFKJS-9, PE |
| Anti-Human TBX21 | ebioscience | Cat# 25-5825-82 Clone: ebio4B10, PE-Cy7 |
| Anti-Human CRTH2 | BioLegend | Cat# 350116 Clone: BM16, PerCP-Cy5.5 |
| Mouse IgG1 k Isotype control | ebioscience | Cat# 25-4714-80 Clone: P3.6.2.8.1, PE-Cy7 |
| Rat IgG2a k isotype control | ebioscience | Cat# 12-4321-80 Clone: eBR2a, PE |
| Rat IgG2a, k isotype control | BioLegend | Cat# 400532 Clone: RTK2758, PerCP-Cy5.5 |
| Anti-Human IFN-γ | BioLegend | Cat# 502512 Clone: 4S.B3, APC |
| Anti-Human IFN-γ | BioLegend | Cat# 502509 Clone: 4S.B3, PE |
| Anti-Human IFN-γ | BioLegend | Cat# 502528 Clone: 4S.B3, PE-Cy7 |
| Anti-Human AREG | ebioscience | Cat# 17-5370-42 Clone: AREG559, APC |
| Anti-Human CD16 | BioLegend | Cat# 980104 Clone: 3G8, APC |
| Anti-Human CD45 | BD | Cat# 560178 Clone: 2D1, APC-H7 |
| Anti-Human CD56 | BioLegend | Cat# 318306 Clone: HCD56, PE |
| Anti-Human CD56 | BioLegend | Cat# 318310 Clone: HCD56, APC |
| Anti-Human CD56 | BioLegend | Cat# 318322 Clone: HCD56, PerCP-Cy5.5 |
| Anti-Human CD117 | BioLegend | Cat# 313213 Clone: 104D2, PerCP-Cy5.5 |
| Anti-Human CD127 | BioLegend | Cat# 351316 Clone: A019D5, APC |
| Anti-Human CD127 | BioLegend | Cat# 351304 Clone: A019D5, PE |
| Anti-Human CD127 | BioLegend | Cat# 351320 Clone: A019D5, PE-Cy7 |
| Anti-Human CD71 | BioLegend | Cat# 334114 Clone: CY1G4, PerCP-Cy5.5 |
| Anti-Human IL-2 | R&D | Cat# MAB202 |
| Mouse IgG1 Isotype Control | R&D | Cat# MAB002 |
| Anti-Human pMTOR Ser 2448 | ebioscience | Cat# 12-9718-42 Clone: MRRBY, PE |
| Anti-Human pAKT Ser 473 | ebioscience | Cat# 17-9715-42 Clone: SDRNR, APC |
| Anti-Human p4EBP1 Thr 36, 45 | ebioscience | Cat# 12-9107-42 Clone: V3NTY24, PE |
| Anti-Human pS6 Ser 235, 236 | ebioscience | Cat# 12-9007-42 Clone: cupk43k, PE |
| Anti-Human EOMES | ebioscience | Cat# 12-4877-42 Clone: WD1982, PE |
| Anti-Human CD94 | BioLegend | Cat# 305516 Clone: DX22, PE-Cy7 |
| Anti-Human KIR3DL1 | BioLegend | Cat# 312719 Clone: DX9, PE-Cy7 |
| Anti-Human KIR2DL1 | BioLegend | Cat# 339505 Clone: HP-MX4, PE |
| Anti-Human KIR2DL4 | BioLegend | Cat# 347008 Clone: mAb 33 (33), APC |
| Anti-Human NKp44 | BioLegend | Cat# 325110 Clone: P44-8, APC |
| Anti-Human NKp46 | BioLegend | Cat# 331918 Clone: 9E2, APC |
| Anti-Human NKp80 | BioLegend | Cat# 346706 Clone:5D12, PE |
| Anti-Human NKG2A | MACS | Cat# 130-098-813 Clone: REA110, PE |
| Anti-Human NKG2C | BioLegend | Cat# 375108 Clone: S19004C, APC |
| Anti-Human NKG2D | BioLegend | Cat# 320824 Clone:1D11, APC-Cy7 |

| | | |
|---|---|---|
| Anti-Human CD107a | BioLegend | Cat# 328607 Clone: H4A3, PE |
| mouse IgG1 k isotype control | BioLegend | Cat# 400128 Clone: MOPC-21, APC-Cy7 |
| mouse IgG1 k isotype control | BioLegend | Cat# 400150 Clone: MOPC-21, PerCP-Cy5.5 |
| mouse IgG1 k isotype control | BioLegend | Cat# 400112 Clone: MOPC-21, PE |
| mouse IgG1 k isotype control | BioLegend | Cat# 400126 Clone: MOPC-21, PE-Cy7 |
| mouse IgG1 k isotype control | BioLegend | Cat# 400120 Clone: MOPC-21, APC |
| human IgG1 k isotype contorl | BioLegend | Cat# 403503 Clone: OA16A12, PE |
| mouse Ig2a k isotype control | BioLegend | Cat# 400212 Clone: MOPC-173, PE |
| mouse Ig2a k isotype control | BioLegend | Cat# 400220 Clone: MOPC-173, APC |
| mouse Ig2a k isotype control | BioLegend | Cat# 400251 Clone: MOPC-173, PerCP-Cy5.5 |
| **Biological Samples** | | |
| PBMCs | UCSF Medical Center | https://hividgm.ucsf.edu/scope-study |
| PBMCs | New York Biologics | https://www.newyorkbiologics.com/ |
| **Chemicals, Peptides, and Recombinant Proteins** | | |
| IL-2 | PeproTech | Cat# 200-02 |
| IL-12p70 | PeproTech | Cat# 200-12 |
| IL-15 | PeproTech | Cat# 200-15 |
| IL-18 | InvivoGen | Cat# rcyec-hil18 |
| TGF-b | PeproTech | Cat# 100-21 |
| Rapamycin | Millipore | Cat# 553211-500UG |
| Torin1 | TOCRIS | Cat# 4247 |
| CP-690550 | InvivoGen | Cat# tlrl-cp69 |
| CHIR99021 | TOCRIS | Cat# 4423 |
| C188-9 | Selleckchem | Cat# S8605 |
| AS-1517499 | Selleckchem | Cat# S8685 |
| MK-2206 | Selleckchem | Cat# S1078 |
| Ravoxertinib | Selleckchem | Cat# S7554 |
| cell stimulation cocktail | ebioscience | Cat# 00-4970-03 |
| protein transport inhibitor | ebioscience | Cat# 00-4980-03 |
| TRIzol reagent | Invitrogen | Cat# 15596018 |
| NEBNext Multiplex Oligos for Illumina | NEB | Cat# E6609S |
| AMPure XP beads | Beckman Culter | Cat# A63880 |
| enAsCAS12a | Wolfe Lab (University of Massachusetts Medical school) | N/A |
| ExoSAP-IT | Affymetrix | Cat# 78200 |
| Nextera Tn5 Transposase | Illumina | Cat# FC-121-1030 |
| SuperScript VILO Master Mix | Invitrogen | Cat# 11755050 |
| Critical Commercial Assays | | |
| Live and Dead violet viability kit | Invitrogen | Cat# L-34963 |
| Mycoplasma detection kit | Lonza | Cat# LT27-318 |
| NEBNext Ultra II DNA library Prep kit for Illumina | NEB | Cat# 7645L |
| PCR product purification kit | Promega | Cat# A9282 |
| P3 Primary Cell 4D-NucleofectorTM X Kit S | Lonza | Cat# V4XP-3032 |
| EasySep Human NK Cell Isolation Kit | STEMCELL | Cat#17955 |

| | | |
|---|---|---|
| NK MACS Medium human | MACS | Cat# 130-114-429 |
| Single Cell 3' Reagent Kits v2 | 10xGenomics | Cat# 120234 |
| NEBNext Ultra II Non directional Second Strand Synthesis Module | NEB | Cat# E6111L |
| HiScribe T7 High Yield RNA Synthesis Kit | NEB | Cat# E2040S |
| Deposited Data | | |
| Bulk and single cell RNA-Seq, ATAC-Seq data generated in this study | NCBI Gene Expression Omnibus | GSE168212 |
| Mouse RNA-Seq and ATAC-Seq generated by previous research | NCBI Gene Expression Omnibus | GSE77695, GSE109125, and GSE11609 |
| **Software and Algorithms** | | |
| FlowJo | FlowJo, LLC | https://www.flowjo.com/ |
| GraphPad Prism | GraphPad Software, Inc | https://www.graphpad.com/scientific-software/prism/ |
| Cell Ranger | 10xGENOMICS | https://support.10xgenomics.com/single-cell-gene-expression/software/downloads/latest? |
| Tophat | Kim et al., 2013 | https://ccb.jhu.edu/software/tophat/index.shtml |
| samtools | Li et al., 2009 | http://www.htslib.org |
| R Statistical Computing Software | The R Foundation | https://www.r-project.org/ |
| ESAT | Derr et al. 2016 | https://github.com/garber-lab/ESAT |
| DESeq2 | Love et al., 2014 | https://bioconductor.org/packages/release/bioc/html/DESeq2.html |
| EdgeR | Robinson and Oshlack, 2010 | https://bioconductor.org/packages/release/bioc/html/edgeR.html |
| Rtsne | Van Der Maaten, 2014 | https://cran.r-project.org/web/packages/Rtsne/index.html |
| Monocle | Trapnell et al., 2014 | https://bioconductor.org/packages/release/bioc/html/monocle.html |
| Bowtie2 | Langmead and Salzberg et al., 2012 | http://bowtie-bio.sourceforge.net/bowtie2/index.shtml |
| RSEM | Li and Dewey, 2011 | https://cran.r-project.org/web/packages/rsem/index.html |
| MACS2 | Zhang et al., 2008 | https://github.com/macs3-project/MACS/wiki/Install-macs2 |
| IGVTools | Robinson etal., 2011 | https://software.broadinstitute.org/software/igv/download |
| Trimmomatic | Bolger et al., 2014 | https://github.com/usadellab/Trimmomatic |
| STAR | Dobin et al., 2013 | https://cran.r-project.org/web/packages/stars/index.html |
| JASPAR | JASPAR | http://jaspar.genereg.net/ |
| GSEA | Aravind S, 2005 | https://genepattern.broadinstitute.org/gp/pages/index.jsf |
| Seurat 3.0 | Sturat and Butler, 2019 | http://satijalab.org/seurat/ |

| Supplemental Table 2 \| DEgenes of CD127-vs CD127+ and | | | |
|---|---|---|---|
| DEgenes of CD127$^{-}$vs CD127$^{+}$ | | | |
| ID | padj | log2FC | pvalue |
| NCR1 | 2.39E-07 | -5.7127 | 4.18E-09 |
| PDGFRB | 9.76E-09 | -5.3543 | 1.31E-10 |
| KRT86 | 0.00144 | -5.2819 | 8.93E-05 |
| CCNJL | 2.45E-09 | -5.2332 | 2.94E-11 |
| SLC1A7 | 4.32E-07 | -5.1574 | 8.09E-09 |
| IDO1 | 0.00767 | -5.144 | 0.000679 |
| RBPMS2 | 0.00078 | -5.0068 | 4.35E-05 |
| CMKLR1 | 7.31E-11 | -4.938 | 6.81E-13 |
| BAALC | 0.00017 | -4.9297 | 7.28E-06 |
| LILRA4 | 3.62E-05 | -4.8251 | 1.27E-06 |
| B3GNT7 | 1.31E-18 | -4.665 | 3.98E-21 |
| JCHAIN | 0.0002 | -4.6478 | 8.82E-06 |
| PRSS23 | 6.58E-22 | -4.6071 | 1.31E-24 |
| PODN | 0.00096 | -4.4908 | 5.55E-05 |
| LAIR2 | 9.37E-15 | -4.4208 | 5.52E-17 |
| ADGRG1 | 5.45E-32 | -4.411 | 1.56E-35 |
| PALLD | 1.62E-07 | -4.4106 | 2.72E-09 |
| SLCO4C1 | 5.24E-17 | -4.4051 | 2.29E-19 |
| KIFC3 | 3.39E-13 | -4.3864 | 2.39E-15 |
| GNAL | 3.38E-08 | -4.3824 | 5.04E-10 |
| KLRF1 | 5.02E-11 | -4.3472 | 4.53E-13 |
| CX3CR1 | 6.50E-09 | -4.3249 | 8.53E-11 |
| RAMP1 | 1.05E-05 | -4.322 | 3.04E-07 |
| GUCY1B3 | 6.91E-06 | -4.3106 | 1.86E-07 |
| CD38 | 5.01E-19 | -4.2555 | 1.48E-21 |
| SH2D1B | 1.34E-31 | -4.2367 | 6.38E-35 |
| NME8 | 2.83E-12 | -4.1978 | 2.26E-14 |
| CD160 | 3.39E-38 | -4.1658 | 3.22E-42 |
| GZMH | 5.96E-20 | -4.1344 | 1.59E-22 |
| CXCR2 | 5.01E-16 | -4.1161 | 2.43E-18 |
| MEIS1 | 0.00063 | -4.1134 | 3.39E-05 |
| SPON2 | 1.20E-30 | -4.0808 | 1.14E-33 |
| AKR1C3 | 3.70E-31 | -4.0444 | 2.11E-34 |
| BNC2 | 5.79E-10 | -4.043 | 6.27E-12 |
| LOC100129083 | 4.50E-05 | -4.0214 | 1.63E-06 |
| PROS1 | 0.00033 | -3.9893 | 1.61E-05 |
| CNTLN | 1.39E-05 | -3.9579 | 4.18E-07 |
| FGFBP2 | 1.18E-23 | -3.9323 | 2.02E-26 |
| ALOX12 | 0.00514 | -3.922 | 0.000423 |
| FCRL2 | 2.20E-06 | -3.9013 | 4.95E-08 |
| FAT4 | 2.31E-05 | -3.8832 | 7.47E-07 |
| MLC1 | 0.00073 | -3.8805 | 3.97E-05 |
| GFOD1 | 7.27E-18 | -3.8532 | 2.63E-20 |
| LGALS9B | 6.23E-05 | -3.8474 | 2.35E-06 |
| KIR3DL3 | 0.00273 | -3.8191 | 0.000193 |
| S1PR5 | 1.77E-33 | -3.8131 | 3.37E-37 |
| NMUR1 | 3.33E-18 | -3.8028 | 1.14E-20 |
| ATP9A | 3.07E-05 | -3.791 | 1.06E-06 |
| HAVCR2 | 6.48E-16 | -3.7821 | 3.33E-18 |
| SETBP1 | 1.71E-07 | -3.7815 | 2.90E-09 |
| TTC38 | 2.47E-19 | -3.7607 | 6.80E-22 |

| | | | |
|---|---|---|---|
| ADGRA2 | 2.89E-05 | -3.7543 | 9.81E-07 |
| MKI67 | 2.87E-13 | -3.7492 | 1.99E-15 |
| CCL3 | 0.00178 | -3.7334 | 0.000115 |
| GZMB | 0.00039 | -3.7323 | 1.94E-05 |
| AKAP5 | 3.87E-10 | -3.7305 | 4.09E-12 |
| ENC1 | 1.33E-13 | -3.7218 | 8.71E-16 |
| SYNGR1 | 2.35E-13 | -3.6979 | 1.58E-15 |
| RHOBTB3 | 2.84E-19 | -3.6192 | 8.10E-22 |
| GPR153 | 5.32E-06 | -3.6068 | 1.37E-07 |
| LGR6 | 2.13E-05 | -3.5839 | 6.74E-07 |
| FCGR3A | 1.94E-18 | -3.5801 | 6.27E-21 |
| ERBB2 | 0.00018 | -3.5682 | 7.99E-06 |
| IGLL5 | 0.00678 | -3.5633 | 0.000588 |
| KIR2DS4 | 0.00015 | -3.5306 | 6.36E-06 |
| PRF1 | 8.09E-29 | -3.5256 | 9.23E-32 |
| CCL4 | 1.20E-30 | -3.4928 | 1.03E-33 |
| IGFBP7 | 6.15E-13 | -3.4765 | 4.45E-15 |
| ASPM | 0.00097 | -3.4571 | 5.65E-05 |
| KIR3DL1 | 0.00073 | -3.4558 | 4.01E-05 |
| PPP1R14A | 0.00093 | -3.4521 | 5.36E-05 |
| DAB2 | 5.45E-14 | -3.4465 | 3.37E-16 |
| CHST2 | 2.79E-05 | -3.431 | 9.40E-07 |
| ASCL2 | 1.14E-06 | -3.3821 | 2.37E-08 |
| FCRL6 | 7.90E-32 | -3.372 | 3.01E-35 |
| LRRC16B | 0.00145 | -3.3662 | 9.03E-05 |
| ADAMTS1 | 0.00399 | -3.3659 | 0.000307 |
| CMTM5 | 0.00078 | -3.3594 | 4.37E-05 |
| CERCAM | 0.00252 | -3.2728 | 0.000175 |
| KIAA0226L | 6.75E-05 | -3.2665 | 2.55E-06 |
| C1orf21 | 9.49E-17 | -3.2593 | 4.24E-19 |
| ITGAX | 0.0001 | -3.2534 | 4.01E-06 |
| FASLG | 1.06E-06 | -3.2002 | 2.17E-08 |
| MYLK | 0.00461 | -3.1912 | 0.000369 |
| ITGB3 | 3.55E-08 | -3.1898 | 5.37E-10 |
| LGALS9C | 0.00028 | -3.1836 | 1.35E-05 |
| PLOD1 | 2.66E-17 | -3.1748 | 1.06E-19 |
| CD244 | 8.59E-15 | -3.1695 | 4.98E-17 |
| PDGFD | 2.09E-16 | -3.1541 | 9.55E-19 |
| FAXDC2 | 0.00042 | -3.1478 | 2.15E-05 |
| ATP8B4 | 2.73E-05 | -3.1464 | 9.11E-07 |
| XCL2 | 2.08E-17 | -3.1362 | 7.90E-20 |
| SYK | 2.96E-09 | -3.117 | 3.63E-11 |
| KLRD1 | 3.85E-18 | -3.0986 | 1.35E-20 |
| SLAMF7 | 4.09E-31 | -3.0488 | 3.11E-34 |
| CLIC3 | 1.17E-08 | -3.048 | 1.61E-10 |
| PCSK6 | 0.00291 | -3.0321 | 0.00021 |
| FGR | 1.38E-05 | -3.0211 | 4.11E-07 |
| HES6 | 2.96E-07 | -3.0197 | 5.29E-09 |
| MYOM2 | 1.84E-05 | -3.0166 | 5.76E-07 |
| ARVCF | 1.01E-05 | -3.0038 | 2.92E-07 |
| GNLY | 2.23E-18 | -2.9953 | 7.42E-21 |
| COLGALT2 | 3.03E-06 | -2.9893 | 7.27E-08 |
| CPNE5 | 0.00017 | -2.9675 | 7.55E-06 |
| KIR2DL4 | 0.00545 | -2.9621 | 0.000455 |

| | | | |
|---|---|---|---|
| KIR2DL3 | 0.00235 | -2.952 | 0.000162 |
| NUAK1 | 6.73E-05 | -2.9471 | 2.54E-06 |
| CTBP2 | 0.00058 | -2.9426 | 3.09E-05 |
| FEZ1 | 0.0019 | -2.9308 | 0.000125 |
| ZEB2 | 2.80E-13 | -2.9031 | 1.92E-15 |
| PIF1 | 0.00048 | -2.8862 | 2.45E-05 |
| MZB1 | 3.93E-05 | -2.8825 | 1.41E-06 |
| AOAH | 4.09E-31 | -2.882 | 2.84E-34 |
| IRF8 | 0.0019 | -2.865 | 0.000124 |
| OSBPL5 | 1.04E-21 | -2.8591 | 2.17E-24 |
| NKG7 | 8.57E-21 | -2.8556 | 2.12E-23 |
| CD22 | 3.03E-05 | -2.8497 | 1.03E-06 |
| B4GALT6 | 0.00039 | -2.8466 | 1.97E-05 |
| TAL1 | 0.00069 | -2.8457 | 3.73E-05 |
| GOLGA8J | 5.70E-13 | -2.8266 | 4.06E-15 |
| PLEK | 5.54E-05 | -2.8097 | 2.07E-06 |
| FAM179A | 1.45E-15 | -2.8009 | 7.89E-18 |
| ADAM28 | 5.35E-08 | -2.7899 | 8.50E-10 |
| PLCG2 | 1.97E-08 | -2.7739 | 2.83E-10 |
| LGALS2 | 0.0009 | -2.7709 | 5.14E-05 |
| MGST1 | 0.00152 | -2.7615 | 9.52E-05 |
| FOXP3 | 0.0042 | -2.7584 | 0.00033 |
| PTGDR | 2.38E-12 | -2.7557 | 1.88E-14 |
| GNAZ | 4.53E-05 | -2.7401 | 1.65E-06 |
| FBLN2 | 0.0028 | -2.7305 | 0.0002 |
| ABI3 | 7.04E-21 | -2.7229 | 1.64E-23 |
| SIGLEC7 | 0.00122 | -2.716 | 7.41E-05 |
| CXXC5 | 5.93E-10 | -2.7077 | 6.54E-12 |
| FAM129C | 0.00032 | -2.6751 | 1.57E-05 |
| ELOVL7 | 0.00102 | -2.6736 | 6.00E-05 |
| CEP78 | 1.49E-24 | -2.6644 | 2.27E-27 |
| GSN | 1.29E-05 | -2.6384 | 3.81E-07 |
| SRC | 0.00948 | -2.6367 | 0.000883 |
| DTHD1 | 7.63E-06 | -2.6272 | 2.08E-07 |
| DLG5 | 0.00011 | -2.6027 | 4.48E-06 |
| CD72 | 0.00436 | -2.5916 | 0.000345 |
| FSD1 | 0.00151 | -2.5809 | 9.45E-05 |
| GOLM1 | 0.00035 | -2.5771 | 1.70E-05 |
| KLRC3 | 2.08E-12 | -2.5703 | 1.62E-14 |
| TM6SF1 | 0.00069 | -2.5613 | 3.76E-05 |
| P2RX1 | 0.00555 | -2.5598 | 0.000465 |
| CBFA2T3 | 0.00288 | -2.5404 | 0.000207 |
| TBX21 | 6.03E-23 | -2.5332 | 1.09E-25 |
| SHTN1 | 0.00874 | -2.533 | 0.000804 |
| DERL3 | 0.00202 | -2.5173 | 0.000134 |
| KRT73 | 0.00321 | -2.516 | 0.000237 |
| YPEL1 | 9.96E-25 | -2.5061 | 1.42E-27 |
| SERPINE2 | 0.00496 | -2.5058 | 0.000404 |
| PTGS1 | 0.00058 | -2.5005 | 3.10E-05 |
| ZCCHC24 | 0.00047 | -2.481 | 2.41E-05 |
| GNGT2 | 0.00014 | -2.4775 | 5.60E-06 |
| FBXO6 | 5.36E-10 | -2.4745 | 5.76E-12 |
| TLE1 | 1.06E-06 | -2.4598 | 2.15E-08 |
| BANK1 | 0.00111 | -2.4562 | 6.63E-05 |

| | | | |
|---|---|---|---|
| TCL1A | 0.00209 | -2.4405 | 0.000142 |
| POU2AF1 | 0.00865 | -2.4308 | 0.00079 |
| TIE1 | 0.00489 | -2.4307 | 0.000396 |
| SPIB | 0.00021 | -2.4199 | 9.51E-06 |
| MYL9 | 0.00158 | -2.4197 | 9.97E-05 |
| GAS2L1 | 0.00516 | -2.4191 | 0.000426 |
| SLC15A4 | 5.11E-25 | -2.4135 | 6.31E-28 |
| FCRL3 | 0.00052 | -2.4111 | 2.68E-05 |
| FCRL1 | 0.00371 | -2.377 | 0.000282 |
| KRT72 | 1.84E-05 | -2.3726 | 5.78E-07 |
| IRF4 | 4.04E-07 | -2.3672 | 7.53E-09 |
| TSPAN13 | 0.00473 | -2.3635 | 0.00038 |
| KLRC4 | 0.00329 | -2.3623 | 0.000244 |
| ADGRG5 | 3.46E-06 | -2.3456 | 8.54E-08 |
| RGS3 | 7.84E-07 | -2.3437 | 1.56E-08 |
| CST7 | 7.64E-16 | -2.3405 | 4.00E-18 |
| IL2RB | 2.91E-16 | -2.3044 | 1.38E-18 |
| FAM49A | 0.00071 | -2.2745 | 3.86E-05 |
| YES1 | 0.00165 | -2.266 | 0.000105 |
| PTPN12 | 2.42E-17 | -2.2577 | 9.43E-20 |
| BCL11A | 7.65E-06 | -2.2488 | 2.09E-07 |
| TST | 0.00277 | -2.2392 | 0.000198 |
| MCTP2 | 8.29E-15 | -2.2224 | 4.73E-17 |
| GK5 | 5.53E-24 | -2.2192 | 8.94E-27 |
| BZRAP1 | 7.80E-08 | -2.2188 | 1.28E-09 |
| MEF2C | 0.00455 | -2.2075 | 0.000363 |
| KLRC2 | 0.00015 | -2.2014 | 6.36E-06 |
| VANGL1 | 1.13E-05 | -2.2008 | 3.31E-07 |
| HIC1 | 0.00158 | -2.2007 | 9.97E-05 |
| CD93 | 0.00407 | -2.1873 | 0.000315 |
| FHL3 | 5.04E-07 | -2.1861 | 9.69E-09 |
| ST3GAL4 | 8.05E-06 | -2.1838 | 2.26E-07 |
| RGS18 | 0.00255 | -2.1652 | 0.000178 |
| GZMA | 1.35E-30 | -2.1644 | 1.41E-33 |
| TCF4 | 0.00109 | -2.1564 | 6.50E-05 |
| PAX5 | 0.0002 | -2.1528 | 8.96E-06 |
| ANKRD9 | 0.00455 | -2.1512 | 0.000362 |
| USP28 | 2.53E-21 | -2.1466 | 5.53E-24 |
| PAFAH2 | 8.33E-14 | -2.1349 | 5.23E-16 |
| IFNG | 5.20E-05 | -2.1264 | 1.93E-06 |
| METTL7A | 0.00047 | -2.1264 | 2.41E-05 |
| KYNU | 0.00845 | -2.1223 | 0.000764 |
| CCDC28B | 0.00028 | -2.114 | 1.31E-05 |
| BPGM | 3.13E-08 | -2.1112 | 4.62E-10 |
| CENPE | 1.91E-06 | -2.1029 | 4.26E-08 |
| TUBA8 | 0.00683 | -2.0996 | 0.000595 |
| UCK2 | 0.00152 | -2.0994 | 9.52E-05 |
| CMC1 | 1.42E-10 | -2.0918 | 1.40E-12 |
| LPCAT1 | 9.78E-12 | -2.0896 | 8.28E-14 |
| KIAA0101 | 0.00026 | -2.0831 | 1.22E-05 |
| GP6 | 0.00655 | -2.071 | 0.000563 |
| CD300A | 1.04E-15 | -2.0584 | 5.56E-18 |
| TOX | 8.55E-12 | -2.0564 | 7.07E-14 |
| TRIO | 2.73E-05 | -2.0534 | 9.15E-07 |

| | | | |
|---|---|---|---|
| EOMES | 2.90E-09 | -2.0476 | 3.53E-11 |
| CENPF | 1.14E-06 | -2.0377 | 2.36E-08 |
| TOP2A | 0.00047 | -2.0293 | 2.40E-05 |
| LAT2 | 0.00026 | -2.0161 | 1.24E-05 |
| RHOC | 3.07E-05 | -2.0071 | 1.06E-06 |
| CXorf21 | 0.00591 | -1.9831 | 0.000498 |
| SLC2A8 | 0.00525 | -1.9515 | 0.000434 |
| EFHD2 | 1.74E-08 | -1.9427 | 2.47E-10 |
| F2R | 1.89E-10 | -1.9361 | 1.91E-12 |
| CD9 | 0.001 | -1.9287 | 5.85E-05 |
| PEAK1 | 5.69E-06 | -1.9109 | 1.48E-07 |
| METRNL | 7.09E-09 | -1.9089 | 9.44E-11 |
| EVI5 | 0.00998 | -1.8966 | 0.000943 |
| KIR3DL2 | 1.09E-10 | -1.8774 | 1.05E-12 |
| IL12RB2 | 0.00352 | -1.8757 | 0.000265 |
| CLU | 1.28E-05 | -1.8734 | 3.77E-07 |
| GSAP | 4.12E-08 | -1.8702 | 6.43E-10 |
| WIPI1 | 0.00011 | -1.8565 | 4.23E-06 |
| SUSD1 | 7.68E-10 | -1.8544 | 8.69E-12 |
| LILRB1 | 0.00283 | -1.8463 | 0.000202 |
| PDLIM1 | 6.46E-06 | -1.8459 | 1.71E-07 |
| KCTD10 | 9.69E-11 | -1.8457 | 9.31E-13 |
| GOLGA8N | 6.78E-05 | -1.8403 | 2.58E-06 |
| TMIGD2 | 1.67E-06 | -1.8293 | 3.66E-08 |
| HDAC9 | 0.00021 | -1.8253 | 9.66E-06 |
| APMAP | 2.82E-17 | -1.8234 | 1.15E-19 |
| TESC | 0.00665 | -1.82 | 0.000575 |
| MPP1 | 0.00206 | -1.8105 | 0.000138 |
| SLC27A3 | 3.37E-06 | -1.8065 | 8.31E-08 |
| ITGAM | 0.0002 | -1.798 | 8.86E-06 |
| MS4A1 | 0.00015 | -1.7954 | 6.54E-06 |
| KLRC1 | 8.95E-06 | -1.775 | 2.54E-07 |
| HIST1H2AK | 0.00851 | -1.7739 | 0.000776 |
| GALNT3 | 3.10E-06 | -1.7693 | 7.46E-08 |
| MATK | 3.65E-09 | -1.7562 | 4.58E-11 |
| ADRBK2 | 0.00307 | -1.7529 | 0.000224 |
| NEXN | 0.00325 | -1.744 | 0.00024 |
| CD247 | 6.30E-25 | -1.7345 | 8.38E-28 |
| GOLIM4 | 0.00996 | -1.7304 | 0.000938 |
| PLEKHO2 | 0.00025 | -1.7178 | 1.19E-05 |
| SLC4A4 | 0.00168 | -1.7171 | 0.000108 |
| MAP3K8 | 0.00042 | -1.7098 | 2.13E-05 |
| CD79A | 0.00074 | -1.7096 | 4.07E-05 |
| TBKBP1 | 0.00636 | -1.7078 | 0.000543 |
| BCL2L1 | 2.53E-07 | -1.7028 | 4.45E-09 |
| ARHGEF12 | 4.78E-05 | -1.6973 | 1.75E-06 |
| DYRK1B | 0.00978 | -1.6894 | 0.000917 |
| TIGIT | 8.01E-06 | -1.6829 | 2.23E-07 |
| PYHIN1 | 5.11E-20 | -1.6689 | 1.31E-22 |
| PTCH2 | 2.28E-05 | -1.6509 | 7.31E-07 |
| TMCC3 | 0.00931 | -1.6354 | 0.000865 |
| CCDC50 | 0.00097 | -1.6347 | 5.63E-05 |
| IRAK2 | 0.00867 | -1.6344 | 0.000796 |
| GAPT | 0.00998 | -1.6336 | 0.000941 |

| | | | |
|---|---|---|---|
| NOTCH1 | 4.93E-08 | -1.6315 | 7.77E-10 |
| HVCN1 | 0.00059 | -1.6209 | 3.17E-05 |
| EHD3 | 0.00658 | -1.6115 | 0.000566 |
| ZBTB16 | 0.00082 | -1.6067 | 4.62E-05 |
| CRIM1 | 7.54E-05 | -1.6025 | 2.90E-06 |
| PLEKHF1 | 5.70E-06 | -1.6007 | 1.49E-07 |
| HLA-DOA | 0.00509 | -1.597 | 0.000418 |
| HOXB4 | 0.00159 | -1.5953 | 0.000101 |
| FAM102B | 1.48E-05 | -1.5947 | 4.47E-07 |
| LIMK1 | 9.84E-06 | -1.5915 | 2.82E-07 |
| HLA-DOB | 0.00179 | -1.5906 | 0.000117 |
| HLA-DMA | 0.00681 | -1.5905 | 0.000591 |
| RAB27B | 0.00645 | -1.5858 | 0.000552 |
| ADRB2 | 0.0064 | -1.5746 | 0.000547 |
| HIST1H2AM | 0.00136 | -1.5604 | 8.36E-05 |
| NPC1 | 2.93E-06 | -1.5538 | 6.94E-08 |
| PRR11 | 0.0003 | -1.5482 | 1.42E-05 |
| SYNE1 | 1.77E-18 | -1.5422 | 5.55E-21 |
| THEMIS2 | 0.00018 | -1.5294 | 7.96E-06 |
| GRAMD4 | 0.00541 | -1.5259 | 0.00045 |
| GNPTAB | 3.80E-11 | -1.5253 | 3.40E-13 |
| TFEB | 0.0048 | -1.5241 | 0.000387 |
| IL18RAP | 0.00119 | -1.514 | 7.19E-05 |
| CHST12 | 3.66E-12 | -1.5069 | 2.96E-14 |
| CACNA2D2 | 0.00358 | -1.5014 | 0.000271 |
| HOPX | 6.86E-10 | -1.5012 | 7.70E-12 |
| PON2 | 8.23E-05 | -1.4995 | 3.20E-06 |
| LLGL2 | 4.04E-14 | -1.491 | 2.46E-16 |
| SSX2IP | 0.00048 | -1.4868 | 2.46E-05 |
| GZMM | 6.13E-08 | -1.4789 | 9.86E-10 |
| ENPP4 | 0.00087 | -1.4786 | 4.96E-05 |
| ARHGAP18 | 0.00025 | -1.4671 | 1.15E-05 |
| RDH10 | 0.00751 | -1.4669 | 0.000662 |
| SCLT1 | 3.60E-06 | -1.4607 | 8.97E-08 |
| PTPRJ | 1.72E-12 | -1.4583 | 1.31E-14 |
| PIK3AP1 | 0.00106 | -1.4539 | 6.27E-05 |
| LYST | 2.85E-16 | -1.448 | 1.33E-18 |
| G6PD | 6.64E-08 | -1.4208 | 1.07E-09 |
| JAZF1 | 4.98E-06 | -1.4147 | 1.27E-07 |
| NRROS | 0.00308 | -1.4119 | 0.000225 |
| RAP1GAP2 | 1.69E-10 | -1.4115 | 1.68E-12 |
| FUT11 | 0.00395 | -1.4111 | 0.000303 |
| SASH3 | 2.45E-10 | -1.4079 | 2.50E-12 |
| PLAC8 | 5.15E-07 | -1.3871 | 9.99E-09 |
| SIRT2 | 0.00029 | -1.3856 | 1.38E-05 |
| TPST2 | 6.86E-08 | -1.3809 | 1.12E-09 |
| HIRA | 0.00077 | -1.3797 | 4.26E-05 |
| HLA-DPB1 | 0.00102 | -1.3692 | 5.99E-05 |
| CASZ1 | 0.00104 | -1.3685 | 6.17E-05 |
| TMEM2 | 3.67E-06 | -1.3628 | 9.20E-08 |
| TGFBR3 | 8.84E-13 | -1.358 | 6.64E-15 |
| MAN1A1 | 0.00037 | -1.3573 | 1.82E-05 |
| HSH2D | 0.00052 | -1.3545 | 2.73E-05 |
| LDLR | 0.00338 | -1.3434 | 0.000252 |

| | | | |
|---|---|---|---|
| APOL1 | 0.00244 | -1.3419 | 0.000169 |
| MPST | 0.00194 | -1.3335 | 0.000128 |
| STX7 | 0.00061 | -1.3335 | 3.26E-05 |
| CTSW | 9.67E-07 | -1.3317 | 1.94E-08 |
| SAMD3 | 2.15E-07 | -1.3293 | 3.72E-09 |
| C12orf75 | 1.37E-06 | -1.3231 | 2.95E-08 |
| C1orf174 | 1.16E-06 | -1.3205 | 2.44E-08 |
| IRF5 | 0.00275 | -1.3183 | 0.000195 |
| TFDP2 | 2.22E-11 | -1.3088 | 1.94E-13 |
| PTPRE | 1.48E-05 | -1.3079 | 4.49E-07 |
| LMNB1 | 0.0026 | -1.3056 | 0.000182 |
| ZNF600 | 2.16E-05 | -1.3043 | 6.87E-07 |
| PRR5L | 0.00192 | -1.2998 | 0.000126 |
| PAK1 | 0.00017 | -1.298 | 7.09E-06 |
| SLC25A20 | 0.00116 | -1.2954 | 7.00E-05 |
| SH2D2A | 6.13E-08 | -1.2921 | 9.80E-10 |
| LIMD1 | 0.00302 | -1.2861 | 0.000219 |
| APOBEC3G | 4.70E-08 | -1.2812 | 7.37E-10 |
| INPP1 | 0.00917 | -1.2804 | 0.00085 |
| EIF4G3 | 3.98E-08 | -1.2803 | 6.05E-10 |
| GNG2 | 3.61E-10 | -1.277 | 3.74E-12 |
| PPP2R2B | 3.12E-05 | -1.2729 | 1.08E-06 |
| HIST1H2BK | 0.00146 | -1.2683 | 9.09E-05 |
| GSE1 | 2.62E-06 | -1.2644 | 6.08E-08 |
| SBK1 | 3.64E-10 | -1.2603 | 3.81E-12 |
| CEP126 | 0.00525 | -1.2513 | 0.000435 |
| ACAA2 | 2.30E-05 | -1.2484 | 7.42E-07 |
| ADA | 8.24E-05 | -1.2402 | 3.21E-06 |
| ZNF117 | 0.00768 | -1.2369 | 0.00068 |
| RAB37 | 6.85E-06 | -1.2356 | 1.83E-07 |
| PDE4A | 0.0009 | -1.2166 | 5.14E-05 |
| ERMP1 | 0.00426 | -1.2102 | 0.000335 |
| NCALD | 0.00628 | -1.2076 | 0.000534 |
| HIST1H1E | 0.00014 | -1.2059 | 5.99E-06 |
| PAM | 0.00376 | -1.2014 | 0.000287 |
| CTNNBIP1 | 8.39E-05 | -1.1996 | 3.29E-06 |
| RRAS2 | 0.00762 | -1.1981 | 0.000673 |
| ADAMTS10 | 0.00035 | -1.1959 | 1.72E-05 |
| GAB3 | 5.98E-06 | -1.1946 | 1.58E-07 |
| HIPK2 | 2.93E-11 | -1.192 | 2.59E-13 |
| KLRK1 | 1.48E-05 | -1.1905 | 4.49E-07 |
| EXTL3 | 0.00573 | -1.1897 | 0.000482 |
| SH3BP2 | 1.14E-05 | -1.1742 | 3.35E-07 |
| RHOQ | 2.46E-05 | -1.1714 | 8.07E-07 |
| RALGPS2 | 0.00865 | -1.1682 | 0.000792 |
| ABHD17A | 7.53E-07 | -1.1528 | 1.49E-08 |
| LAIR1 | 0.00099 | -1.1412 | 5.76E-05 |
| HEG1 | 0.00203 | -1.1399 | 0.000136 |
| CTSC | 1.58E-05 | -1.1331 | 4.83E-07 |
| DGKQ | 0.00514 | -1.1301 | 0.000424 |
| STARD3NL | 0.00118 | -1.1239 | 7.10E-05 |
| AGK | 0.00084 | -1.1224 | 4.80E-05 |
| MVD | 0.00121 | -1.1171 | 7.31E-05 |
| IGF2R | 3.58E-09 | -1.113 | 4.43E-11 |

| | | | |
|---|---|---|---|
| LASP1 | 2.80E-07 | -1.1093 | 4.96E-09 |
| USB1 | 1.13E-05 | -1.1073 | 3.31E-07 |
| AGTRAP | 0.00693 | -1.1066 | 0.000603 |
| CASP3 | 0.00332 | -1.1024 | 0.000247 |
| XKR8 | 0.00427 | -1.087 | 0.000336 |
| LRMP | 1.69E-05 | -1.0864 | 5.26E-07 |
| CTSD | 1.34E-05 | -1.0835 | 3.96E-07 |
| FAM49B | 9.09E-11 | -1.0805 | 8.55E-13 |
| ABHD2 | 2.46E-05 | -1.0615 | 8.04E-07 |
| APLP2 | 0.0006 | -1.0561 | 3.20E-05 |
| ASL | 0.0067 | -1.0557 | 0.00058 |
| NCOA4 | 0.0002 | -1.054 | 9.16E-06 |
| MTURN | 0.00988 | -1.0468 | 0.000929 |
| NAA40 | 0.00391 | -1.042 | 0.0003 |
| CD63 | 0.00023 | -1.0265 | 1.06E-05 |
| ADCY7 | 2.73E-09 | -1.0246 | 3.30E-11 |
| TMEM107 | 0.00545 | -1.021 | 0.000456 |
| LGALS9 | 0.00019 | -1.018 | 8.35E-06 |
| STMN1 | 0.00498 | -1.0174 | 0.000406 |
| CCL5 | 7.36E-07 | -1.01 | 1.45E-08 |
| MAPK1 | 7.00E-15 | -1.0093 | 3.86E-17 |
| SKAP2 | 0.00289 | -1.0093 | 0.000208 |
| HENMT1 | 0.00339 | -1.009 | 0.000253 |
| PTPN18 | 4.16E-05 | -1.0077 | 1.50E-06 |
| IKZF3 | 3.15E-06 | -1.0065 | 7.72E-08 |
| RUNX2 | 0.00474 | 1.00987 | 0.000382 |
| FOXO1 | 5.70E-06 | 1.01292 | 1.50E-07 |
| SLC39A8 | 0.00116 | 1.01332 | 6.97E-05 |
| FXYD5 | 4.70E-17 | 1.0154 | 2.01E-19 |
| FLJ44635 | 0.00985 | 1.01749 | 0.000925 |
| PPA1 | 0.00048 | 1.02262 | 2.49E-05 |
| RLTPR | 0.00843 | 1.0304 | 0.000762 |
| LEPROTL1 | 7.34E-06 | 1.03066 | 1.99E-07 |
| ARHGAP5 | 2.31E-06 | 1.03107 | 5.26E-08 |
| HIVEP2 | 0.00093 | 1.03513 | 5.33E-05 |
| DGKA | 3.24E-05 | 1.03711 | 1.13E-06 |
| CD3E | 0.00075 | 1.04071 | 4.17E-05 |
| BEX2 | 0.00032 | 1.05205 | 1.57E-05 |
| DNAJB1 | 0.00036 | 1.06652 | 1.78E-05 |
| ATP6V0E2 | 0.00048 | 1.07157 | 2.50E-05 |
| SVIL | 5.03E-05 | 1.07718 | 1.86E-06 |
| SLC26A11 | 0.00702 | 1.07976 | 0.000615 |
| TTC39C | 4.53E-05 | 1.0882 | 1.66E-06 |
| REEP3 | 0.00267 | 1.09618 | 0.000188 |
| RAB3GAP1 | 0.00473 | 1.09949 | 0.000379 |
| NFKB1 | 4.76E-07 | 1.10322 | 9.01E-09 |
| BCL11B | 1.22E-05 | 1.10685 | 3.58E-07 |
| PDE8A | 0.00851 | 1.10792 | 0.000772 |
| SAMSN1 | 2.73E-05 | 1.11056 | 9.17E-07 |
| CASK | 0.00043 | 1.127 | 2.17E-05 |
| KLHL6 | 0.00433 | 1.13914 | 0.000343 |
| SIRPG | 0.00332 | 1.14004 | 0.000247 |
| ITGA6 | 1.90E-07 | 1.14387 | 3.27E-09 |
| SC5D | 0.00215 | 1.14726 | 0.000146 |

| | | | |
|---|---|---|---|
| CREBL2 | 0.00014 | 1.14878 | 5.99E-06 |
| MAP9 | 0.00513 | 1.14965 | 0.000422 |
| BIRC3 | 0.00011 | 1.15286 | 4.39E-06 |
| KDSR | 0.00285 | 1.1626 | 0.000205 |
| PLP2 | 1.11E-09 | 1.16448 | 1.28E-11 |
| ARHGAP15 | 2.94E-08 | 1.17007 | 4.25E-10 |
| STMN3 | 3.91E-09 | 1.1721 | 4.94E-11 |
| SYPL1 | 0.00013 | 1.17635 | 5.07E-06 |
| ZBTB25 | 0.00012 | 1.18282 | 4.99E-06 |
| LDHB | 1.75E-11 | 1.18321 | 1.51E-13 |
| DYNLT3 | 2.39E-05 | 1.21413 | 7.75E-07 |
| SPEF2 | 0.00739 | 1.21714 | 0.00065 |
| TBC1D4 | 0.00193 | 1.23092 | 0.000127 |
| MPZL3 | 0.00659 | 1.23422 | 0.000568 |
| CD82 | 0.00016 | 1.23728 | 6.76E-06 |
| CMSS1 | 0.00491 | 1.23841 | 0.000399 |
| ANTXR2 | 3.78E-05 | 1.24659 | 1.34E-06 |
| GRAP | 0.00341 | 1.24878 | 0.000255 |
| MINA | 0.00321 | 1.25301 | 0.000237 |
| ENO2 | 0.00407 | 1.25368 | 0.000315 |
| MDFIC | 1.59E-05 | 1.26434 | 4.87E-07 |
| EPHA4 | 0.00025 | 1.26666 | 1.14E-05 |
| USP53 | 0.00652 | 1.26702 | 0.000559 |
| PIK3IP1 | 1.19E-06 | 1.26808 | 2.52E-08 |
| MTM1 | 0.00602 | 1.27235 | 0.000509 |
| LINS1 | 0.00022 | 1.28157 | 1.00E-05 |
| CD27 | 0.00049 | 1.28562 | 2.57E-05 |
| DNPH1 | 2.46E-05 | 1.29006 | 8.06E-07 |
| ATP8B2 | 9.82E-09 | 1.29235 | 1.33E-10 |
| TTC9 | 0.00611 | 1.29284 | 0.000519 |
| FKBP5 | 0.00407 | 1.29372 | 0.000314 |
| THEM4 | 1.72E-05 | 1.29844 | 5.39E-07 |
| BNIP3 | 0.00027 | 1.29862 | 1.29E-05 |
| MAST4 | 0.00075 | 1.30109 | 4.17E-05 |
| ACVR1 | 0.00867 | 1.30611 | 0.000796 |
| MAML2 | 0.00016 | 1.32824 | 6.77E-06 |
| S1PR1 | 0.0007 | 1.32827 | 3.83E-05 |
| SVIP | 3.10E-07 | 1.33961 | 5.63E-09 |
| PBX4 | 0.001 | 1.34469 | 5.81E-05 |
| DISC1 | 0.00489 | 1.34843 | 0.000396 |
| CHD7 | 4.17E-09 | 1.34911 | 5.36E-11 |
| GPR171 | 0.00046 | 1.35765 | 2.37E-05 |
| FHL1 | 0.00775 | 1.35837 | 0.00069 |
| MFHAS1 | 2.45E-06 | 1.36722 | 5.63E-08 |
| CD6 | 0.00185 | 1.37135 | 0.000121 |
| GATA3 | 3.98E-09 | 1.37652 | 5.07E-11 |
| IL4R | 1.40E-10 | 1.37778 | 1.37E-12 |
| MTFP1 | 0.00021 | 1.37887 | 9.49E-06 |
| GPRASP1 | 0.00222 | 1.37956 | 0.000152 |
| LRRC8C | 0.00021 | 1.38838 | 9.65E-06 |
| PAG1 | 1.21E-06 | 1.39053 | 2.56E-08 |
| SLC34A2 | 0.00816 | 1.39256 | 0.000733 |
| MFGE8 | 0.00234 | 1.40303 | 0.000161 |
| IL6ST | 7.09E-09 | 1.4052 | 9.43E-11 |

| | | | |
|---|---|---|---|
| DOCK9 | 0.00056 | 1.41183 | 2.92E-05 |
| RTKN2 | 0.00502 | 1.41902 | 0.000411 |
| TMEM123 | 8.72E-18 | 1.42772 | 3.23E-20 |
| USP46 | 0.00032 | 1.43213 | 1.53E-05 |
| TCF7 | 1.78E-08 | 1.43986 | 2.54E-10 |
| CDR2 | 0.00043 | 1.44117 | 2.20E-05 |
| PRKCA | 0.00018 | 1.44192 | 7.87E-06 |
| GPR183 | 0.00015 | 1.45379 | 6.36E-06 |
| CFAP36 | 1.80E-12 | 1.46792 | 1.39E-14 |
| FAM84B | 0.00036 | 1.47558 | 1.79E-05 |
| TOB1 | 1.16E-06 | 1.47885 | 2.43E-08 |
| A1BG | 0.00145 | 1.48854 | 8.98E-05 |
| TSHZ2 | 0.00851 | 1.4911 | 0.000775 |
| PLK3 | 0.00026 | 1.49125 | 1.20E-05 |
| PCNXL2 | 2.93E-06 | 1.5018 | 6.91E-08 |
| NOSIP | 2.01E-09 | 1.50362 | 2.39E-11 |
| PDE3B | 1.42E-05 | 1.5043 | 4.27E-07 |
| TNFAIP3 | 0.00019 | 1.50879 | 8.65E-06 |
| LEF1 | 0.00025 | 1.51393 | 1.18E-05 |
| ANO9 | 4.36E-06 | 1.52652 | 1.10E-07 |
| PDE4B | 4.03E-08 | 1.52971 | 6.20E-10 |
| PLAG1 | 0.00833 | 1.53721 | 0.00075 |
| SUPT3H | 0.00631 | 1.54111 | 0.000537 |
| FLT3LG | 5.44E-16 | 1.54359 | 2.69E-18 |
| CD5 | 0.00117 | 1.54467 | 7.04E-05 |
| TRADD | 1.13E-13 | 1.54762 | 7.33E-16 |
| FAM102A | 5.06E-06 | 1.55696 | 1.29E-07 |
| SORBS3 | 7.77E-06 | 1.56391 | 2.15E-07 |
| ADAM19 | 3.01E-06 | 1.57073 | 7.19E-08 |
| LRIG1 | 1.57E-05 | 1.57167 | 4.79E-07 |
| LTB | 1.38E-09 | 1.5879 | 1.61E-11 |
| AP3M2 | 1.69E-05 | 1.59441 | 5.26E-07 |
| TNFRSF10A | 1.63E-06 | 1.59653 | 3.57E-08 |
| TESPA1 | 6.38E-10 | 1.59994 | 7.10E-12 |
| OXNAD1 | 2.16E-07 | 1.60537 | 3.76E-09 |
| NSG1 | 3.81E-05 | 1.61218 | 1.35E-06 |
| SESN3 | 0.00116 | 1.61912 | 6.98E-05 |
| NR1D1 | 0.00159 | 1.62603 | 0.000101 |
| CYB561 | 1.53E-06 | 1.64324 | 3.31E-08 |
| FHIT | 0.00033 | 1.66075 | 1.59E-05 |
| BAIAP3 | 0.00109 | 1.68074 | 6.49E-05 |
| TNFRSF25 | 7.07E-15 | 1.69034 | 3.96E-17 |
| CDC14A | 2.35E-14 | 1.70428 | 1.41E-16 |
| IL6R | 8.13E-06 | 1.71159 | 2.29E-07 |
| UBASH3A | 0.00407 | 1.72382 | 0.000314 |
| EPHX2 | 0.00032 | 1.74822 | 1.51E-05 |
| CITED4 | 9.72E-05 | 1.74899 | 3.82E-06 |
| GDPD5 | 5.55E-06 | 1.75262 | 1.44E-07 |
| SERINC5 | 1.29E-06 | 1.76488 | 2.75E-08 |
| PKIA | 0.00013 | 1.76689 | 5.14E-06 |
| TIAM1 | 2.27E-06 | 1.77862 | 5.13E-08 |
| SOCS3 | 1.84E-09 | 1.78585 | 2.17E-11 |
| MEGF6 | 5.40E-05 | 1.80019 | 2.01E-06 |
| PASK | 3.07E-07 | 1.80294 | 5.51E-09 |

| | | | |
|---|---|---|---|
| C1orf228 | 0.00022 | 1.80655 | 1.01E-05 |
| NPDC1 | 0.0009 | 1.80831 | 5.13E-05 |
| BCL2 | 9.20E-14 | 1.81135 | 5.86E-16 |
| LDOC1 | 0.00042 | 1.82396 | 2.12E-05 |
| ANK3 | 2.94E-06 | 1.83549 | 7.00E-08 |
| IFI44L | 2.65E-05 | 1.84654 | 8.79E-07 |
| CD28 | 4.04E-08 | 1.85209 | 6.26E-10 |
| MLLT4 | 0.00136 | 1.86293 | 8.35E-05 |
| SH2D3A | 1.34E-06 | 1.86304 | 2.86E-08 |
| TSPAN18 | 3.58E-09 | 1.87057 | 4.46E-11 |
| MAN1C1 | 0.00022 | 1.87743 | 9.93E-06 |
| FAM134B | 3.88E-07 | 1.89162 | 7.19E-09 |
| FBXL16 | 6.78E-05 | 1.904 | 2.57E-06 |
| RNF157 | 5.50E-09 | 1.90668 | 7.16E-11 |
| RCAN3 | 7.13E-13 | 1.91546 | 5.22E-15 |
| RHPN1 | 0.00219 | 1.92939 | 0.00015 |
| HSF4 | 0.002 | 1.92972 | 0.000133 |
| TRAT1 | 3.10E-06 | 1.95331 | 7.51E-08 |
| ZC3H12D | 1.05E-05 | 1.95762 | 3.03E-07 |
| TNFSF8 | 7.86E-13 | 1.97937 | 5.83E-15 |
| SUSD3 | 2.39E-06 | 1.99167 | 5.47E-08 |
| DCHS1 | 0.00106 | 2.01656 | 6.29E-05 |
| HAPLN3 | 9.62E-11 | 2.01891 | 9.14E-13 |
| RIC3 | 0.00495 | 2.03856 | 0.000402 |
| CAMK4 | 3.09E-07 | 2.07368 | 5.59E-09 |
| MYC | 0.00025 | 2.07632 | 1.16E-05 |
| DUSP16 | 5.88E-10 | 2.08058 | 6.43E-12 |
| PFN2 | 0.00558 | 2.08215 | 0.000468 |
| VIPR1 | 0.00164 | 2.10094 | 0.000104 |
| CLN5 | 0.00456 | 2.11681 | 0.000364 |
| SLAMF1 | 0.00045 | 2.15538 | 2.29E-05 |
| AQP3 | 1.73E-07 | 2.17165 | 2.95E-09 |
| KIF5C | 8.05E-06 | 2.18337 | 2.26E-07 |
| RASGRF2 | 0.00482 | 2.1978 | 0.00039 |
| INADL | 1.01E-11 | 2.20538 | 8.68E-14 |
| ACSL6 | 2.16E-06 | 2.20962 | 4.85E-08 |
| LTK | 0.00496 | 2.23318 | 0.000404 |
| LMTK3 | 0.00799 | 2.24851 | 0.000715 |
| IGFBP3 | 0.00163 | 2.26246 | 0.000104 |
| CCR7 | 3.27E-08 | 2.27012 | 4.85E-10 |
| CRIP2 | 3.44E-07 | 2.28562 | 6.34E-09 |
| PLEKHB1 | 7.77E-06 | 2.2888 | 2.16E-07 |
| RNF207 | 0.00767 | 2.31212 | 0.000679 |
| LSR | 4.18E-12 | 2.31633 | 3.41E-14 |
| FXYD7 | 2.84E-17 | 2.31882 | 1.19E-19 |
| RGMB | 0.00367 | 2.33056 | 0.000279 |
| INPP4B | 1.28E-09 | 2.33411 | 1.48E-11 |
| MAL | 9.29E-12 | 2.35897 | 7.77E-14 |
| CD40LG | 1.03E-08 | 2.41292 | 1.41E-10 |
| NT5E | 0.00645 | 2.4538 | 0.000553 |
| NR3C2 | 9.07E-06 | 2.45733 | 2.59E-07 |
| FXYD1 | 0.00851 | 2.47802 | 0.000776 |
| IL7R | 2.03E-22 | 2.52712 | 3.85E-25 |
| SUSD4 | 0.00163 | 2.55347 | 0.000104 |

| | | | |
|---|---|---|---|
| TRABD2A | 7.04E-21 | 2.55465 | 1.67E-23 |
| ANKRD18A | 7.51E-05 | 2.56974 | 2.89E-06 |
| DPP4 | 2.01E-13 | 2.58455 | 1.34E-15 |
| CISH | 0.0096 | 2.58462 | 0.000895 |
| EEPD1 | 0.0042 | 2.60773 | 0.00033 |
| KLF5 | 0.00101 | 2.64464 | 5.89E-05 |
| HOOK1 | 1.71E-06 | 2.64738 | 3.78E-08 |
| NELL2 | 4.42E-10 | 2.64812 | 4.71E-12 |
| VSIG1 | 1.50E-05 | 2.67446 | 4.55E-07 |
| TLCD2 | 0.00058 | 2.68881 | 3.04E-05 |
| MCF2L | 0.00327 | 2.69041 | 0.000242 |
| CMTM8 | 0.00025 | 2.71021 | 1.16E-05 |
| GREM2 | 0.00232 | 2.71039 | 0.00016 |
| CNTNAP1 | 3.42E-05 | 2.72796 | 1.20E-06 |
| ACVR1C | 0.00545 | 2.77706 | 0.000456 |
| SEC14L2 | 0.00207 | 2.80671 | 0.00014 |
| TCEA3 | 6.15E-11 | 2.81928 | 5.67E-13 |
| IL23A | 3.15E-06 | 2.82492 | 7.73E-08 |
| C10orf35 | 0.00893 | 2.83738 | 0.000823 |
| CCR6 | 8.01E-07 | 2.87541 | 1.60E-08 |
| LRRN3 | 0.00288 | 2.93989 | 0.000207 |
| AXIN2 | 1.03E-09 | 2.99886 | 1.18E-11 |
| PLEKHG5 | 0.00151 | 3.06271 | 9.44E-05 |
| TMEM45B | 0.00252 | 3.14204 | 0.000175 |
| TMEM30B | 0.00102 | 3.17131 | 5.96E-05 |
| WNT7A | 5.43E-07 | 3.18476 | 1.06E-08 |
| CFH | 1.11E-08 | 3.19139 | 1.52E-10 |
| HLF | 0.0002 | 3.19395 | 8.77E-06 |
| CACNA1I | 1.69E-06 | 3.23724 | 3.73E-08 |
| HPCAL4 | 0.00163 | 3.25958 | 0.000103 |
| RORC | 0.00505 | 3.27927 | 0.000414 |
| FGF9 | 0.00058 | 3.28866 | 3.09E-05 |
| SEMA5A | 0.00027 | 3.36094 | 1.26E-05 |
| HDAC11 | 0.00321 | 3.36153 | 0.000237 |
| TRIM2 | 3.83E-05 | 3.39958 | 1.37E-06 |
| PDE9A | 0.00893 | 3.55719 | 0.000824 |
| RFPL2 | 0.00866 | 3.56363 | 0.000794 |
| KRT1 | 0.00262 | 3.58154 | 0.000184 |
| PTGDR2 | 0.00865 | 3.66971 | 0.000791 |
| HID1 | 0.00327 | 3.76304 | 0.000243 |
| TMEM200A | 0.00107 | 3.80581 | 6.35E-05 |
| IL4I1 | 0.00334 | 3.88601 | 0.000249 |
| C14orf132 | 0.00199 | 4.04366 | 0.000132 |
| GAP43 | 0.00259 | 4.1698 | 0.000182 |
| ACE | 1.51E-08 | 4.19368 | 2.12E-10 |
| ST6GALNAC1 | 0.00262 | 4.20983 | 0.000184 |
| MB21D2 | 4.09E-05 | 4.31855 | 1.47E-06 |
| NEFL | 1.07E-07 | 4.40005 | 1.77E-09 |
| ADAM23 | 1.06E-06 | 4.56689 | 2.16E-08 |
| RBPMS | 0.00174 | 4.92728 | 0.000112 |

**CD127- reactome analysis**

| Reactome pathways | # | # | expected | Fold Enrichment | +/- | raw P value | FDR |
|---|---|---|---|---|---|---|---|
| Sema4D induced cell migration and growth-cone collapse | 20 | 5 | 0.39 | 12.87 | + | 9.50E-05 | 1.97E-02 |
| Sema4D in semaphorin signaling | 24 | 5 | 0.47 | 10.73 | + | 2.00E-04 | 3.80E-02 |
| DAP12 signaling | 25 | 5 | 0.49 | 10.3 | + | 2.36E-04 | 3.85E-02 |
| DAP12 interactions | 39 | 6 | 0.76 | 7.92 | + | 2.00E-04 | 3.52E-02 |
| Immunoregulatory interactions between a Lymphoid and a non-Lymphoid cell | 199 | 20 | 3.87 | 5.17 | + | 8.42E-09 | 9.61E-06 |
| Platelet degranulation | 127 | 10 | 2.47 | 4.05 | + | 3.02E-04 | 4.31E-02 |
| Platelet activation, signaling and aggregatio | 259 | 19 | 5.03 | 3.78 | + | 1.81E-06 | 6.89E-04 |
| Neutrophil degranulation | 478 | 28 | 9.28 | 3.02 | + | 5.32E-07 | 4.05E-04 |
| Signaling by Interleukins | 447 | 24 | 8.68 | 2.76 | + | 1.39E-05 | 3.53E-03 |
| Hemostasis | 670 | 33 | 13.01 | 2.54 | + | 3.11E-06 | 1.01E-03 |
| Adaptive Immune System | 820 | 39 | 15.93 | 2.45 | + | 5.82E-07 | 3.32E-04 |
| Immune System | 2159 | 95 | 41.94 | 2.27 | + | 8.36E-14 | 1.91E-10 |
| Innate Immune System | 1105 | 47 | 21.46 | 2.19 | + | 1.07E-06 | 4.87E-04 |
| Cytokine Signaling in Immune system | 823 | 32 | 15.99 | 2 | + | 2.86E-04 | 4.35E-02 |
| Unclassified | 10195 | 157 | 198.02 | 0.79 | - | 5.66E-05 | 1.29E-02 |
| Metabolism of RNA | 661 | 0 | 12.84 | < 0.01 | - | 3.98E-06 | 1.14E-03 |
| **CD127+ reactome analysis** | | | | | | | |
| Reactome pathways | # | # | expected | Fold Enrichment | +/- | raw P value | FDR |
| Binding of TCF/LEF:CTNNB1 to target gene promoters | 8 | 4 | 0.09 | 42.9 | + | 7.91E-06 | 6.02E-03 |
| Interleukin-4 and Interleukin-13 signaling | 111 | 10 | 1.29 | 7.73 | + | 1.41E-06 | 1.62E-03 |
| Cytokine Signaling in Immune system | 823 | 28 | 9.59 | 2.92 | + | 6.29E-07 | 1.44E-03 |
| Death Receptor Signalling | 141 | 9 | 1.64 | 5.48 | + | 6.25E-05 | 2.85E-02 |

| Supplemental Table 3 \| DEgenes from paired comparison of Lin$^-$ populations and reactome analysis | | | | | | | | | | | |
|---|---|---|---|---|---|---|---|---|---|---|---|
| DEgenes | | | | | | | | | | | |
| CD56$^{neg}$NK vs ILC | | | | CD56$^{dim}$NK vs ILC | | | | CD56$^{hi}$NK vs ILC | | | |
| ID | log2FC | pvalue | padj | ID | log2FC | pvalue | padj | ID | log2FC | pvalue | padj |
| CX3CR1 | 9.7843433 | 2.56E-19 | 5.00E-17 | CX3CR1 | 9.6650869 | 1.67E-21 | 3.23E-19 | NCAM1 | 9.1479289 | 3.31E-76 | 1.06E-72 |
| KIR2DL3 | 9.5546208 | 4.06E-23 | 1.06E-20 | KIR2DL3 | 9.4724334 | 2.52E-14 | 1.93E-12 | PACSIN1 | 8.1239984 | 7.97E-20 | 2.54E-17 |
| S1PR5 | 8.9185066 | 6.40E-48 | 7.68E-45 | DAB2 | 8.9368757 | 1.56E-26 | 4.51E-24 | IL12RB2 | 7.7684498 | 3.00E-19 | 7.98E-17 |
| SPON2 | 8.7650315 | 1.79E-121 | 7.53E-118 | SPON2 | 8.914952 | 1.91E-135 | 1.77E-131 | EOMES | 7.6256249 | 7.66E-34 | 1.05E-30 |
| DAB2 | 8.6634417 | 5.53E-23 | 1.41E-20 | S1PR5 | 8.899882 | 3.29E-51 | 3.04E-48 | KIR2DL3 | 7.5978329 | 3.43E-13 | 4.64E-11 |
| EOMES | 7.8042566 | 1.58E-32 | 8.28E-30 | RASSF4 | 8.4120152 | 3.60E-23 | 9.01E-21 | ZMAT4 | 7.4953002 | 4.09E-19 | 1.06E-16 |
| CCL3 | 7.6300074 | 1.35E-13 | 1.12E-11 | LGR6 | 8.3187948 | 7.82E-23 | 1.86E-20 | CCR1 | 7.3223937 | 1.06E-15 | 1.99E-13 |
| CCL4L2 | 7.3702392 | 3.92E-11 | 2.07E-09 | FGFBP2 | 7.8061475 | 6.35E-17 | 7.26E-15 | STYK1 | 7.161679 | 3.61E-14 | 5.59E-12 |
| FGFBP2 | 7.2552058 | 2.52E-14 | 2.46E-12 | NCAM1 | 7.7584731 | 5.19E-53 | 5.34E-50 | CDHR1 | 6.8600585 | 2.49E-19 | 6.81E-17 |
| RASSF4 | 7.1944812 | 6.39E-13 | 4.83E-11 | CCL3 | 7.7295036 | 5.17E-18 | 6.46E-16 | KLRF1 | 6.6325475 | 3.63E-84 | 1.74E-80 |
| GPR56 | 7.1533249 | 1.58E-51 | 2.22E-48 | FEZ1 | 7.5576093 | 5.16E-17 | 5.98E-15 | ASF1B | 6.4934936 | 1.25E-09 | 9.69E-08 |
| FCGR3A | 6.9748785 | 4.28E-17 | 6.42E-15 | EOMES | 7.5473543 | 3.95E-26 | 1.11E-23 | C16orf70 | 6.4169375 | 2.53E-08 | 1.42E-06 |
| FEZ1 | 6.9372466 | 2.02E-11 | 1.15E-09 | FCGR3A | 7.1726895 | 5.42E-18 | 6.69E-16 | SPTSSB | 6.3859365 | 1.39E-11 | 1.45E-09 |
| CD38 | 6.8150885 | 2.41E-10 | 1.08E-08 | GPR56 | 7.1545212 | 5.25E-56 | 6.07E-53 | GPR97 | 6.195343 | 2.40E-09 | 1.64E-07 |
| FCRL6 | 6.7926635 | 3.19E-13 | 2.57E-11 | DTHD1 | 7.1419512 | 9.22E-14 | 6.37E-12 | GNLY | 6.1934255 | 2.25E-101 | 2.16E-97 |
| LAIR2 | 6.6859514 | 1.02E-14 | 1.08E-12 | CXCR2 | 6.9962547 | 9.28E-18 | 1.12E-15 | LDB2 | 6.0721342 | 8.86E-10 | 7.09E-08 |
| GZMH | 6.6688996 | 2.68E-21 | 5.78E-19 | PRSS23 | 6.929696 | 5.43E-42 | 3.35E-39 | S1PR5 | 6.0686949 | 1.16E-12 | 1.44E-10 |
| GZMA | 6.584581 | 1.97E-65 | 4.14E-62 | CCL4L2 | 6.886574 | 1.89E-10 | 7.92E-09 | KRT86 | 5.9463457 | 1.28E-10 | 1.20E-08 |
| CCL4 | 6.5467991 | 7.35E-15 | 8.12E-13 | LINGO2 | 6.8282918 | 6.50E-12 | 3.48E-10 | KLRD1 | 5.9416855 | 2.87E-15 | 5.09E-13 |
| PIGB | 6.5426906 | 3.20E-08 | 9.69E-07 | FCRL6 | 6.8140763 | 3.04E-14 | 2.27E-12 | KIR2DL4 | 5.7402289 | 3.65E-09 | 2.42E-07 |
| MYOM2 | 6.4717036 | 8.82E-08 | 2.43E-06 | GZMA | 6.7238509 | 2.71E-70 | 5.01E-67 | KLRC4 | 5.6875836 | 1.39E-09 | 1.04E-07 |
| GZMB | 6.379399 | 1.55E-09 | 5.90E-08 | LAIR2 | 6.6656761 | 4.73E-15 | 4.09E-13 | PPIL2 | 5.6567993 | 5.17E-06 | 0.0001829 |
| BCL2A1 | 6.2872725 | 7.18E-08 | 2.02E-06 | GZMB | 6.6505648 | 2.73E-10 | 1.10E-08 | CATSPER1 | 5.6350779 | 1.95E-06 | 7.59E-05 |
| CXCR2 | 6.2402429 | 1.87E-10 | 8.40E-09 | B3GAT1 | 6.61237 | 6.08E-11 | 2.83E-09 | GPR114 | 5.5723366 | 1.51E-09 | 1.12E-07 |
| PYHIN1 | 6.1932504 | 2.47E-25 | 7.39E-23 | CMKLR1 | 6.581545 | 1.63E-18 | 2.25E-16 | RASSF4 | 5.5561368 | 9.33E-07 | 3.86E-05 |
| PRSS23 | 6.1765624 | 5.10E-30 | 2.25E-27 | ASCL2 | 6.5771225 | 7.27E-11 | 3.35E-09 | DTHD1 | 5.514018 | 7.40E-06 | 0.0002508 |
| ASCL2 | 6.1474048 | 3.17E-07 | 8.04E-06 | KLRF1 | 6.5600449 | 8.56E-94 | 2.64E-90 | GZMA | 5.4705133 | 2.71E-29 | 1.73E-26 |
| KLRF1 | 6.1324357 | 4.29E-71 | 1.20E-67 | COL13A1 | 6.410747 | 6.86E-08 | 1.81E-06 | XIRP2 | 5.4171113 | 9.21E-08 | 4.60E-06 |
| GPR114 | 6.1136223 | 2.78E-11 | 1.53E-09 | GZMH | 6.3910887 | 7.45E-22 | 1.57E-19 | HMMR | 5.3462285 | 5.24E-06 | 0.0001841 |
| KIR2DS4 | 6.0634048 | 1.23E-10 | 6.07E-09 | NLRP7 | 6.3744647 | 6.36E-09 | 2.03E-07 | CD38 | 5.142899 | 4.31E-05 | 0.0011251 |
| PDGFRB | 5.8220054 | 2.71E-06 | 5.77E-05 | CCL4 | 6.3728608 | 9.88E-15 | 8.17E-13 | QPRT | 5.0996477 | 0.0001453 | 0.0032893 |
| KIR3DL1 | 5.808989 | 5.85E-13 | 4.47E-11 | PDGFRB | 6.3500218 | 1.39E-09 | 5.05E-08 | ROR1 | 5.089324 | 1.31E-05 | 0.0004017 |
| DTHD1 | 5.7468713 | 1.48E-05 | 0.0002683 | MYOM2 | 6.3396057 | 6.33E-10 | 2.44E-08 | RAD51 | 5.0097053 | 9.05E-07 | 3.76E-05 |
| KLRD1 | 5.6072194 | 2.30E-12 | 1.55E-10 | KIR3DL1 | 6.2384163 | 1.72E-08 | 4.98E-07 | OSBPL5 | 5.0007885 | 1.73E-17 | 4.05E-15 |
| MMP23A | 5.5570153 | 1.86E-08 | 5.85E-07 | SLAMF7 | 6.1991478 | 1.62E-08 | 4.71E-07 | GMPR | 4.9925227 | 1.32E-06 | 5.30E-05 |
| CST7 | 5.5455674 | 2.00E-47 | 2.10E-44 | KIR2DS4 | 6.1364354 | 4.23E-16 | 4.21E-14 | NDC80 | 4.9589625 | 5.96E-06 | 0.0002055 |
| PRF1 | 5.5317832 | 6.90E-38 | 5.79E-35 | PCDH1 | 5.9091935 | 5.37E-07 | 1.20E-05 | CABLES1 | 4.9108142 | 2.76E-09 | 1.88E-07 |
| CD244 | 5.4621082 | 6.91E-17 | 9.84E-15 | CD244 | 5.8973871 | 1.03E-27 | 3.19E-25 | NCR1 | 4.9033716 | 0.0004377 | 0.0082307 |
| NKG7 | 5.4207343 | 9.30E-43 | 8.67E-40 | GPR114 | 5.8574718 | 2.59E-12 | 1.48E-10 | NKG7 | 4.8928123 | 6.81E-32 | 5.02E-29 |
| SLAMF7 | 5.419671 | 1.98E-06 | 4.36E-05 | XIRP2 | 5.8274554 | 1.60E-07 | 3.93E-06 | NR1H3 | 4.8652824 | 0.0002197 | 0.0046629 |
| LOC100130 | 5.388469 | 1.29E-16 | 1.78E-14 | CST7 | 5.8064546 | 1.10E-61 | 1.45E-58 | RAD51AP1 | 4.8630876 | 2.17E-05 | 0.0006212 |
| LGALS9C | 5.3279329 | 8.79E-15 | 9.58E-13 | MIR181A2HG | 5.8026678 | 3.85E-06 | 6.94E-05 | CD244 | 4.8526246 | 8.62E-14 | 1.25E-11 |
| ENC1 | 5.3255938 | 2.83E-05 | 0.0004832 | PYHIN1 | 5.7885789 | 2.88E-10 | 1.16E-08 | SLC27A2 | 4.8501228 | 0.0002634 | 0.0054445 |
| HIC1 | 5.2444597 | 3.95E-06 | 8.08E-05 | CD38 | 5.6858239 | 2.06E-06 | 3.95E-05 | TMEM241 | 4.8198156 | 3.75E-08 | 2.01E-06 |
| HIST1H3D | 5.2345839 | 3.94E-07 | 9.90E-06 | PRF1 | 5.6774152 | 1.48E-42 | 1.06E-39 | ZAK | 4.800202 | 5.11E-07 | 2.26E-05 |
| ATP9A | 5.2314866 | 1.07E-09 | 4.24E-08 | CERCAM | 5.6512876 | 2.35E-06 | 4.45E-05 | GZMB | 4.7916718 | 6.26E-06 | 0.0002137 |
| BAAT | 5.2087371 | 0.0002094 | 0.0027822 | RBPMS2 | 5.6391011 | 1.20E-06 | 2.46E-05 | CX3CR1 | 4.790556 | 6.63E-07 | 2.83E-05 |
| TIGIT | 5.1111197 | 7.38E-07 | 1.78E-05 | DGKK | 5.6165807 | 1.26E-10 | 5.50E-09 | LGALS9C | 4.766653 | 3.05E-12 | 3.65E-10 |
| HSPA7 | 5.1023668 | 7.60E-09 | 2.61E-07 | KLRD1 | 5.5841223 | 9.08E-13 | 5.61E-11 | SLAMF7 | 4.7085086 | 2.12E-05 | 0.0006119 |
| OSBPL5 | 5.0668246 | 4.20E-16 | 5.34E-14 | NKG7 | 5.5815292 | 2.94E-46 | 2.27E-43 | SPON2 | 4.7034295 | 1.40E-13 | 2.00E-11 |

| | | | | | | | | | | | |
|---|---|---|---|---|---|---|---|---|---|---|---|
| LINC00437 | 5.0618245 | 0.0006303 | 0.0073414 | NME8 | 5.568825 | 2.76E-11 | 1.35E-09 | CST7 | 4.6736719 | 3.99E-32 | 3.19E-29 |
| NME8 | 5.0337109 | 6.27E-08 | 1.79E-06 | PIF1 | 5.4195798 | 1.17E-05 | 0.0001944 | CMC1 | 4.6646848 | 1.04E-25 | 4.97E-23 |
| CMKLR1 | 4.9963613 | 1.25E-06 | 2.88E-05 | KRT86 | 5.4193077 | 6.56E-08 | 1.74E-06 | DGAT1 | 4.6287293 | 0.0007476 | 0.0128291 |
| MKI67 | 4.9908465 | 8.44E-07 | 2.01E-05 | MELK | 5.4185803 | 2.30E-05 | 0.0003606 | B3GNT7 | 4.4998584 | 4.45E-35 | 8.53E-32 |
| ABI3 | 4.9685542 | 1.15E-10 | 5.72E-09 | HIC1 | 5.382263 | 8.29E-07 | 1.80E-05 | ENO3 | 4.4965736 | 0.0002428 | 0.0050516 |
| KIF11 | 4.9596052 | 2.87E-05 | 0.0004873 | HMOX1 | 5.377093 | 5.46E-05 | 0.0007774 | CKAP2L | 4.4866717 | 0.0001598 | 0.0035642 |
| KIR2DL1 | 4.9509493 | 1.01E-14 | 1.08E-12 | TMEM97 | 5.3603913 | 3.53E-05 | 0.0005315 | SH3BP4 | 4.4662147 | 2.90E-05 | 0.0008022 |
| PLEK | 4.918992 | 1.39E-09 | 5.37E-08 | GNAL | 5.3479812 | 9.41E-12 | 4.92E-10 | ATP9A | 4.4487988 | 2.21E-09 | 1.53E-07 |
| GNAL | 4.8948445 | 1.32E-08 | 4.29E-07 | GNLY | 5.3471319 | 1.76E-65 | 2.71E-62 | TBX21 | 4.3657619 | 1.79E-24 | 7.80E-22 |
| TTC38 | 4.8053037 | 6.94E-15 | 7.77E-13 | CCDC28B | 5.3433422 | 7.15E-08 | 1.89E-06 | GZMK | 4.352003 | 8.20E-20 | 2.54E-17 |
| HAVCR2 | 4.7871964 | 1.05E-10 | 5.28E-09 | OSBPL5 | 5.3355922 | 5.40E-20 | 9.62E-18 | FLJ20021 | 4.2807677 | 1.27E-05 | 0.0003902 |
| KIR3DX1 | 4.7693649 | 0.0002729 | 0.003504 | CATSPER1 | 5.2924569 | 1.31E-05 | 0.0002159 | SPRY2 | 4.2528894 | 0.0010354 | 0.0166616 |
| TBX21 | 4.708798 | 4.27E-28 | 1.71E-25 | LGALS9C | 5.2758882 | 2.78E-15 | 2.52E-13 | IRF8 | 4.2405911 | 4.96E-06 | 0.000177 |
| USP28 | 4.6983362 | 1.17E-15 | 1.39E-13 | KIR2DL1 | 5.2737683 | 3.06E-19 | 4.72E-17 | FEZ1 | 4.2330183 | 0.0029729 | 0.0366488 |
| ADRB2 | 4.6762296 | 4.97E-12 | 3.09E-10 | CXCR1 | 5.1769727 | 7.38E-05 | 0.0010204 | KIR2DS4 | 4.2102052 | 0.0001148 | 0.0026607 |
| BRI3BP | 4.6021074 | 0.0002026 | 0.0027217 | KIR3DX1 | 5.0720578 | 2.43E-05 | 0.0003779 | ZNF418 | 4.1528822 | 0.002495 | 0.0325128 |
| PAFAH2 | 4.5830599 | 2.71E-14 | 2.56E-12 | HSPA7 | 5.0610922 | 1.41E-10 | 6.12E-09 | TRMT61B | 4.1370846 | 8.75E-06 | 0.0002865 |
| SFT2D2 | 4.5631832 | 1.05E-05 | 0.0001988 | LOC1001308 | 5.0570867 | 4.59E-15 | 4.01E-13 | DCLRE1A | 4.0949475 | 0.0050407 | 0.0559556 |
| B3GNT7 | 4.5611498 | 7.99E-10 | 3.27E-08 | FAM216A | 5.0487531 | 1.84E-06 | 3.62E-05 | FGFBP2 | 4.0941473 | 8.08E-07 | 3.38E-05 |
| TMCC3 | 4.5402617 | 5.21E-06 | 0.0001044 | LIPC | 5.0463142 | 9.62E-05 | 0.0012795 | KIF11 | 4.0761118 | 0.0009668 | 0.0158782 |
| AKR1C3 | 4.535182 | 9.46E-12 | 5.67E-10 | TTC38 | 5.0300529 | 1.19E-08 | 3.58E-07 | CD2 | 4.0228757 | 3.51E-29 | 2.10E-26 |
| PDGFD | 4.533633 | 2.57E-12 | 1.70E-10 | PLEK | 4.9498072 | 1.21E-09 | 4.42E-08 | GZMH | 4.016802 | 2.19E-05 | 0.0006241 |
| ATP6V1E2 | 4.483721 | 0.0002796 | 0.0035788 | MGAM | 4.9269237 | 6.77E-06 | 0.0001171 | AQP11 | 4.001874 | 0.0099065 | 0.0915121 |
| FGR | 4.472404 | 5.17E-37 | 3.95E-34 | DGAT1 | 4.915056 | 0.0002483 | 0.0029461 | HVCN1 | 3.9953819 | 1.73E-24 | 7.80E-22 |
| GNLY | 4.4116509 | 1.70E-36 | 1.09E-33 | ABI3 | 4.861948 | 1.22E-10 | 5.35E-09 | CXCR3 | 3.9940218 | 1.95E-11 | 1.99E-09 |
| HIST1H2BB | 4.4080827 | 0.0013545 | 0.0136883 | LIM2 | 4.7882197 | 0.0004941 | 0.0053428 | FCGR3A | 3.9891459 | 5.29E-05 | 0.0013324 |
| NCAM1 | 4.3774335 | 1.29E-06 | 2.96E-05 | SMKR1 | 4.7782611 | 0.0003823 | 0.0042792 | FCRL6 | 3.9806692 | 1.37E-08 | 8.03E-07 |
| CHST2 | 4.3614356 | 2.20E-26 | 7.38E-24 | ENC1 | 4.7603479 | 0.0002635 | 0.0030878 | NAT1 | 3.9788622 | 0.0008996 | 0.0149919 |
| MYO5B | 4.3203167 | 0.0001342 | 0.0019068 | COL15A1 | 4.6775338 | 0.0005849 | 0.0061949 | TMEM163 | 3.9648612 | 0.0080277 | 0.0784849 |
| SLCO4C1 | 4.3151709 | 1.35E-10 | 6.49E-09 | TBX21 | 4.6749475 | 1.80E-39 | 8.77E-37 | LOC10192 | 3.9628311 | 0.000282 | 0.0057186 |
| ERBB2 | 4.2625966 | 2.37E-06 | 5.11E-05 | SMAGP | 4.6705432 | 1.04E-09 | 3.81E-08 | IFNG-AS1 | 3.9601046 | 3.22E-06 | 0.0001222 |
| GZMM | 4.2066121 | 1.73E-11 | 1.01E-09 | SYNGR1 | 4.6458261 | 2.57E-05 | 0.0003963 | XCL2 | 3.9511453 | 1.10E-33 | 1.26E-30 |
| ZEB2 | 4.1821605 | 1.55E-26 | 5.43E-24 | FICD | 4.5980394 | 0.0009129 | 0.0090282 | LOC10192 | 3.9440879 | 3.48E-11 | 3.37E-09 |
| MLC1 | 4.1638678 | 4.69E-10 | 2.02E-08 | MMP23A | 4.5476733 | 2.77E-05 | 0.0004253 | DICER1-A | 3.9312905 | 0.0082274 | 0.080111 |
| LOC101928 | 4.0965387 | 7.18E-09 | 2.50E-07 | ZAK | 4.541755 | 3.48E-06 | 6.33E-05 | IL2RB | 3.9261578 | 1.12E-21 | 4.12E-19 |
| BRMS1L | 4.0911867 | 0.0004899 | 0.005877 | USP28 | 4.5391002 | 4.15E-16 | 4.17E-14 | ITGAX | 3.9124837 | 2.27E-32 | 1.98E-29 |
| CEP78 | 4.0750885 | 9.67E-10 | 3.89E-08 | MYO5B | 4.517041 | 8.30E-06 | 0.0001417 | PDGFD | 3.8856256 | 1.59E-09 | 1.15E-07 |
| IGFBP7 | 4.0681023 | 5.89E-06 | 0.0001171 | HS3ST3B1 | 4.5058845 | 1.15E-06 | 2.38E-05 | CCNJL | 3.8783136 | 2.35E-05 | 0.0006682 |
| GFOD1 | 4.0532062 | 0.0018812 | 0.0178713 | PDGFD | 4.4844096 | 1.63E-12 | 9.65E-11 | SMAGP | 3.8667312 | 5.14E-06 | 0.0001826 |
| UNC45B | 3.9875454 | 0.0030861 | 0.0266914 | FGR | 4.4599867 | 2.78E-42 | 1.84E-39 | GFOD1 | 3.8577166 | 0.0036174 | 0.0430991 |
| CD160 | 3.9661214 | 1.27E-13 | 1.07E-11 | TIGIT | 4.4520378 | 9.59E-12 | 4.99E-10 | GPR56 | 3.8570666 | 9.94E-09 | 6.04E-07 |
| LPCAT1 | 3.9610399 | 9.94E-36 | 5.96E-33 | RGS3 | 4.4395643 | 1.49E-07 | 3.68E-06 | FGR | 3.8303575 | 1.29E-26 | 6.89E-24 |
| PTPRJ | 3.9551812 | 4.29E-16 | 5.38E-14 | SLCO4C1 | 4.4069712 | 1.13E-15 | 1.08E-13 | BAALC | 3.8029257 | 0.0004215 | 0.0080052 |
| TTC16 | 3.9373161 | 0.0048028 | 0.0383766 | AKR1C3 | 4.3967637 | 7.25E-15 | 6.10E-13 | FAM173B | 3.801355 | 0.0003737 | 0.0072706 |
| CCL5 | 3.9272458 | 3.37E-26 | 1.09E-23 | NAT1 | 4.3647318 | 0.0001468 | 0.0018611 | MKI67 | 3.7639507 | 4.73E-07 | 2.10E-05 |
| TBKBP1 | 3.9239836 | 0.002173 | 0.0202318 | PTGDS | 4.3637295 | 2.66E-07 | 6.29E-06 | SGSM1 | 3.6979435 | 0.0045313 | 0.051371 |
| CEBPD | 3.904847 | 3.80E-06 | 7.81E-05 | IRF8 | 4.3485743 | 5.37E-06 | 9.38E-05 | XCL1 | 3.6975579 | 6.52E-26 | 3.29E-23 |
| BAALC | 3.8949631 | 0.0027151 | 0.0240268 | HAVCR2 | 4.3107548 | 5.79E-10 | 2.25E-08 | PYHIN1 | 3.6805598 | 9.93E-07 | 4.09E-05 |
| TBC1D7 | 3.8844444 | 7.00E-05 | 0.0010558 | RNF217 | 4.308275 | 0.0015834 | 0.014455 | TTC8 | 3.6696108 | 0.0007401 | 0.0127893 |
| ENPP5 | 3.8514938 | 0.0006966 | 0.0079378 | ADRB2 | 4.2874039 | 3.18E-10 | 1.27E-08 | KLHL23 | 3.64195 | 0.0002045 | 0.0043781 |
| TMEM62 | 3.8419732 | 4.17E-05 | 0.0006753 | GZMM | 4.2826827 | 9.57E-14 | 6.56E-12 | LOC72873 | 3.6403517 | 0.0051039 | 0.0563961 |
| LMNB1 | 3.7924434 | 4.27E-11 | 2.23E-09 | C7orf31 | 4.2623572 | 0.000845 | 0.0084569 | PRSS23 | 3.624622 | 2.84E-08 | 1.57E-06 |
| CBX3P2 | 3.7627749 | 0.0001393 | 0.0019634 | SETBP1 | 4.2459003 | 4.84E-11 | 2.30E-09 | NOD2 | 3.6231324 | 8.88E-06 | 0.0002896 |
| EFHD2 | 3.7414063 | 3.15E-27 | 1.20E-24 | IGFBP7 | 4.2155563 | 5.16E-07 | 1.16E-05 | KLRC2 | 3.6229397 | 5.14E-13 | 6.57E-11 |

| | | | | | | | | | | | |
|---|---|---|---|---|---|---|---|---|---|---|---|
| TMLHE | 3.7325968 | 0.0007435 | 0.0083698 | TMCC3 | 4.2147054 | 6.22E-06 | 0.000108 | TMEM62 | 3.6037357 | 2.15E-05 | 0.0006171 |
| PXYLP1 | 3.7273392 | 0.0002914 | 0.0036977 | GFOD1 | 4.1946877 | 0.0020668 | 0.0181341 | HAVCR2 | 3.5959336 | 3.24E-06 | 0.0001224 |
| LOC653160 | 3.7084076 | 0.0005393 | 0.0064247 | NUDT11 | 4.1567529 | 8.18E-05 | 0.0011157 | MICAL2 | 3.591392 | 3.54E-08 | 1.92E-06 |
| AGTRAP | 3.7033653 | 0.0009165 | 0.009931 | IFNG | 4.148498 | 2.83E-09 | 9.73E-08 | IL18RAP | 3.5498102 | 2.35E-11 | 2.35E-09 |
| CMC1 | 3.6926305 | 3.67E-11 | 1.95E-09 | PAM | 4.1134866 | 7.24E-09 | 2.29E-07 | DCAF13P3 | 3.5417868 | 0.0024947 | 0.0325128 |
| RGS3 | 3.6025282 | 0.0013045 | 0.0132627 | PAFAH2 | 4.1101528 | 1.59E-14 | 1.25E-12 | COTL1 | 3.5309812 | 8.05E-35 | 1.29E-31 |
| SYNE1 | 3.5712709 | 1.47E-30 | 7.26E-28 | HIST1H3D | 4.1013517 | 0.0005843 | 0.0061949 | DPH3P1 | 3.4909718 | 0.002556 | 0.0330799 |
| TGFBR3 | 3.5512135 | 4.94E-27 | 1.80E-24 | ZEB2 | 4.0966668 | 5.53E-29 | 1.90E-26 | USP28 | 3.4572616 | 8.85E-08 | 4.45E-06 |
| IRF8 | 3.5415573 | 0.0013996 | 0.0140427 | HOXB4 | 4.0880093 | 9.31E-06 | 0.000157 | TIGIT | 3.4433769 | 1.66E-06 | 6.56E-05 |
| LYN | 3.5038959 | 1.99E-20 | 4.07E-18 | SPTB | 4.0669454 | 1.40E-07 | 3.49E-06 | PAFAH2 | 3.4131575 | 3.66E-09 | 2.42E-07 |
| C1orf21 | 3.5032692 | 0.001456 | 0.0144706 | KIFC3 | 4.0611988 | 6.08E-08 | 1.63E-06 | S100A13 | 3.378113 | 0.0002059 | 0.0043973 |
| KIR3DL2 | 3.4028284 | 8.60E-05 | 0.0012777 | ARVCF | 4.0572685 | 7.70E-05 | 0.0010581 | KIR3DL1 | 3.3588362 | 0.0026269 | 0.0335923 |
| CHN2 | 3.3670222 | 0.0002924 | 0.0036977 | CEBPD | 4.0329924 | 1.06E-07 | 2.72E-06 | NEIL1 | 3.3566399 | 4.06E-06 | 0.0001496 |
| KIAA1841 | 3.3420329 | 0.0001304 | 0.0018595 | LOC440461 | 4.0186812 | 0.009877 | 0.0666407 | PLEK | 3.3563388 | 3.01E-10 | 2.72E-08 |
| SETBP1 | 3.3359685 | 0.0001498 | 0.0020799 | CA13 | 4.0152123 | 0.000121 | 0.0015663 | CD300A | 3.3434573 | 4.12E-09 | 2.70E-07 |
| SLC14A1 | 3.3310684 | 0.0074577 | 0.0541778 | CCL5 | 3.9769364 | 2.19E-41 | 1.20E-38 | KLRC3 | 3.330743 | 8.22E-14 | 1.21E-11 |
| ASTN2 | 3.3165786 | 0.0007637 | 0.0085629 | CHST2 | 3.9597809 | 9.00E-22 | 1.85E-19 | WDSUB1 | 3.3279166 | 0.0001233 | 0.002828 |
| AGK | 3.299258 | 1.30E-10 | 6.30E-09 | CEP78 | 3.9354437 | 5.43E-05 | 0.0007771 | ZNF831 | 3.3256024 | 2.35E-08 | 1.33E-06 |
| LPAR5 | 3.2977384 | 0.0030958 | 0.0267055 | LPCAT1 | 3.9159325 | 8.08E-42 | 4.68E-39 | UEVLD | 3.3209867 | 0.0033109 | 0.0400324 |
| HES6 | 3.2928013 | 0.0005687 | 0.006703 | NDC80 | 3.9138813 | 0.0031536 | 0.0259265 | ITPKC | 3.3010783 | 0.0059671 | 0.0633085 |
| RAB27B | 3.2676843 | 6.95E-07 | 1.68E-05 | B3GNT7 | 3.8952938 | 1.39E-15 | 1.31E-13 | HOXA5 | 3.2949792 | 1.86E-05 | 0.0005429 |
| ZNF322 | 3.2632382 | 0.0013042 | 0.0132627 | ANKRD20A1 | 3.8793092 | 0.0016222 | 0.0147369 | LOC10192 | 3.2882267 | 0.0009588 | 0.0157737 |
| TKTL1 | 3.2529722 | 1.83E-07 | 4.86E-06 | C20orf197 | 3.8761526 | 0.0003013 | 0.0034645 | C1QTNF6 | 3.2881505 | 0.0044683 | 0.0510187 |
| SIGLEC7 | 3.2474607 | 0.0001866 | 0.0025194 | TKTL1 | 3.8658045 | 2.88E-05 | 0.0004399 | PAM | 3.2653121 | 1.31E-05 | 0.0004017 |
| WDSUB1 | 3.2444516 | 0.0022896 | 0.0210605 | ENPP4 | 3.8358939 | 1.00E-09 | 3.70E-08 | PRF1 | 3.2650199 | 3.22E-12 | 3.82E-10 |
| IL2RB | 3.242332 | 8.63E-16 | 1.03E-13 | TMEM241 | 3.8350218 | 0.0008115 | 0.0081563 | CA13 | 3.2279764 | 0.0050643 | 0.0560873 |
| ULK4P1 | 3.1493931 | 0.0004104 | 0.0050169 | SIGLEC7 | 3.8288843 | 6.42E-07 | 1.42E-05 | ENG | 3.2232509 | 0.0007689 | 0.0131686 |
| PAM | 3.1249327 | 0.0002863 | 0.003649 | WDSUB1 | 3.8247766 | 7.50E-05 | 0.0010349 | SYK | 3.1962549 | 1.60E-13 | 2.22E-11 |
| TNFRSF1B | 3.1135844 | 1.35E-22 | 3.24E-20 | FABP3 | 3.8200936 | 0.0074556 | 0.0528902 | PPFIBP2 | 3.1823882 | 0.0077238 | 0.0765273 |
| SPTB | 3.0936363 | 0.0026722 | 0.0237727 | EFHD2 | 3.8136109 | 4.81E-34 | 2.02E-31 | ENC1 | 3.1801281 | 0.0009856 | 0.0160771 |
| FAM49A | 3.0756573 | 3.94E-05 | 0.0006429 | SYNE1 | 3.7946005 | 9.17E-35 | 4.24E-32 | CEBPD | 3.1654043 | 0.0002811 | 0.0057111 |
| APMAP | 3.0415284 | 6.58E-17 | 9.53E-15 | FBXO44 | 3.7633853 | 1.41E-08 | 4.19E-07 | IGFBP4 | 3.1461717 | 1.93E-06 | 7.57E-05 |
| PLEKHF1 | 3.0244136 | 2.97E-06 | 6.23E-05 | CBX3P2 | 3.7367585 | 2.91E-05 | 0.0004438 | RGS3 | 3.1427264 | 0.0019529 | 0.0272245 |
| SH2D1B | 3.0182312 | 4.69E-12 | 2.96E-10 | CCNJL | 3.6971232 | 0.0002574 | 0.0030282 | DFNB31 | 3.1347569 | 0.0068235 | 0.0700811 |
| HENMT1 | 3.0150034 | 0.0004897 | 0.005877 | MLC1 | 3.6959122 | 5.01E-08 | 1.36E-06 | TYROBP | 3.1329005 | 1.45E-23 | 6.05E-21 |
| MED20 | 3.014813 | 0.0060671 | 0.0460177 | ZNF831 | 3.6889184 | 1.10E-10 | 4.86E-09 | ADAMTS1 | 3.1310342 | 0.0015568 | 0.0230783 |
| CORO1C | 3.0070052 | 0.0010265 | 0.0108984 | BAALC | 3.6182334 | 0.005434 | 0.0413332 | MLC1 | 3.118352 | 8.56E-06 | 0.0002812 |
| FAM179A | 3.0021127 | 1.41E-05 | 0.0002592 | TTC16 | 3.5844566 | 0.0066336 | 0.0485049 | DNM3 | 3.1165231 | 0.0097077 | 0.0902145 |
| RSPRY1 | 3.0012032 | 0.0021915 | 0.0203807 | TMEM62 | 3.5842985 | 5.43E-05 | 0.0007771 | IGFBP7 | 3.1153353 | 2.38E-07 | 1.10E-05 |
| SYK | 2.9939955 | 2.63E-11 | 1.47E-09 | FAM49A | 3.5584042 | 7.77E-08 | 2.04E-06 | EFNA5 | 3.114134 | 0.0001973 | 0.0042516 |
| ADAM28 | 2.9856155 | 0.0014531 | 0.0144706 | LOC1019285 | 3.5554405 | 0.0011415 | 0.0108821 | CEP78 | 3.101137 | 1.41E-05 | 0.0004253 |
| CLIC3 | 2.9807545 | 0.0092051 | 0.0636778 | UEVLD | 3.5488334 | 0.0024359 | 0.0208984 | FAM161B | 3.085837 | 0.0026758 | 0.0339576 |
| CHAF1A | 2.9628 | 0.0002909 | 0.0036977 | APMAP | 3.5297166 | 4.94E-32 | 1.99E-29 | SPTB | 3.0506209 | 0.0003721 | 0.0072545 |
| NTNG2 | 2.9413393 | 0.0038373 | 0.0319128 | ADAMTS1 | 3.5186309 | 0.0090506 | 0.0621585 | TBC1D1 | 3.0178759 | 1.15E-08 | 6.92E-07 |
| GK5 | 2.9373636 | 6.09E-18 | 9.84E-16 | PTPRJ | 3.5173967 | 9.77E-13 | 5.95E-11 | PLCG2 | 3.0165776 | 6.81E-19 | 1.72E-16 |
| CHST12 | 2.9138898 | 2.60E-06 | 5.56E-05 | SLC14A1 | 3.5064155 | 0.0008597 | 0.0085755 | ZNF620 | 3.009427 | 0.0043757 | 0.0501998 |
| MIR646HG | 2.8887962 | 0.0071899 | 0.0526251 | LOC1019281 | 3.4770423 | 1.50E-06 | 3.01E-05 | AGK | 2.9973134 | 3.21E-11 | 3.14E-09 |
| ADAMTS1 | 2.8745269 | 0.0063104 | 0.0475292 | TGFBR3 | 3.4426646 | 6.09E-28 | 1.94E-25 | TIE1 | 2.9797045 | 0.0030262 | 0.0371156 |
| TMTC4 | 2.8681564 | 1.11E-07 | 3.00E-06 | ZNF28 | 3.4414853 | 0.0004865 | 0.0052797 | PVRIG | 2.9740062 | 2.55E-11 | 2.52E-09 |
| RNASEH2A | 2.8615395 | 0.0021143 | 0.0197727 | FCGR3B | 3.4257689 | 0.0035474 | 0.0287553 | ZNF322 | 2.9671241 | 0.0012712 | 0.0196324 |
| MICAL2 | 2.8506489 | 0.0002284 | 0.0030155 | ERBB2 | 3.4109193 | 0.0002876 | 0.0033317 | FAM49A | 2.9212501 | 2.83E-05 | 0.0007933 |
| ENAH | 2.8504639 | 0.0048918 | 0.0387193 | MYPOP | 3.4088868 | 0.0031958 | 0.0262494 | ZNF329 | 2.9211367 | 0.0036402 | 0.0433169 |
| LBH | 2.8488731 | 2.77E-14 | 2.58E-12 | PTCH1 | 3.3817991 | 1.07E-07 | 2.74E-06 | SIGLEC7 | 2.911885 | 0.0004236 | 0.0080289 |
| MRPL10 | 2.8445678 | 1.77E-08 | 5.63E-07 | CACNA2D2 | 3.3737739 | 1.05E-07 | 2.72E-06 | FUT7 | 2.8987185 | 0.0003044 | 0.0061416 |

| | | | | | | | | | | | |
|---|---|---|---|---|---|---|---|---|---|---|---|
| ENPP4 | 2.8392125 | 9.96E-05 | 0.0014572 | F2R | 3.343397 | 0.0004143 | 0.0045879 | PLA2G6 | 2.8867393 | 3.24E-08 | 1.78E-06 |
| HIST1H2AC | 2.8333365 | 1.22E-05 | 0.0002254 | CD160 | 3.3336945 | 7.56E-10 | 2.86E-08 | LINC0099 | 2.8859983 | 0.0024802 | 0.0324086 |
| AREL1 | 2.8279596 | 6.88E-07 | 1.67E-05 | LYN | 3.3305847 | 3.99E-16 | 4.06E-14 | MYOM2 | 2.8852137 | 0.0023924 | 0.0316495 |
| TTYH3 | 2.8182005 | 0.0080734 | 0.0578012 | IL18RAP | 3.3269884 | 6.90E-10 | 2.65E-08 | ULK4P1 | 2.8823921 | 0.0009813 | 0.0160606 |
| GBP5 | 2.8012083 | 4.42E-06 | 8.99E-05 | RRN3P2 | 3.3265159 | 0.0001308 | 0.0016816 | RFC3 | 2.8767357 | 0.0070404 | 0.0715297 |
| FBXO44 | 2.7654516 | 0.0032068 | 0.0273963 | TADA2A | 3.3113456 | 0.0026505 | 0.0225097 | ADAM28 | 2.8752632 | 0.0011963 | 0.0186571 |
| CD247 | 2.7572826 | 3.55E-21 | 7.46E-19 | NTNG2 | 3.3076182 | 0.0001847 | 0.0022798 | GZMM | 2.8629485 | 3.71E-05 | 0.0009995 |
| BNC2 | 2.749738 | 2.30E-06 | 4.97E-05 | TBKBP1 | 3.2412889 | 0.0096227 | 0.0652107 | NLRP2 | 2.8479854 | 5.99E-06 | 0.000206 |
| FCER1G | 2.7355518 | 3.50E-07 | 8.86E-06 | GNPTAB | 3.2196943 | 4.84E-06 | 8.53E-05 | MLYCD | 2.824806 | 0.0048339 | 0.0541607 |
| ZNF831 | 2.7325572 | 0.0001634 | 0.0022424 | GNGT2 | 3.2148147 | 0.0021619 | 0.018862 | GLIDR | 2.7972273 | 0.0027132 | 0.0343308 |
| KRT72 | 2.7172167 | 0.0008446 | 0.0092957 | FBXL6 | 3.2127634 | 0.0050864 | 0.0390421 | FCER1G | 2.7953168 | 0.0003514 | 0.0069058 |
| SLC15A4 | 2.6695171 | 1.83E-11 | 1.05E-09 | CMC1 | 3.1683945 | 7.72E-09 | 2.42E-07 | VANGL1 | 2.7925342 | 4.89E-05 | 0.0012443 |
| NLRP2 | 2.6557158 | 5.86E-05 | 0.0009023 | SERPINH1 | 3.1645127 | 0.0035414 | 0.0287312 | FOXC1 | 2.7635578 | 0.0045079 | 0.0512263 |
| ZNF786 | 2.6538673 | 0.0089976 | 0.0627072 | IL2RB | 3.1568578 | 5.79E-15 | 4.96E-13 | GAB1 | 2.7625044 | 1.61E-05 | 0.0004793 |
| ARHGEF12 | 2.6517961 | 0.0037419 | 0.0313613 | PXYLP1 | 3.124753 | 0.002517 | 0.0214956 | LYN | 2.7411843 | 4.13E-12 | 4.82E-10 |
| CRYZ | 2.6511014 | 0.0093005 | 0.064126 | HENMT1 | 3.0996414 | 5.12E-06 | 9.01E-05 | IL18 | 2.7311092 | 0.0072095 | 0.0727853 |
| SLC16A6 | 2.6446726 | 7.50E-06 | 0.0001458 | FHL3 | 3.0907471 | 3.68E-05 | 0.000551 | C6orf25 | 2.7253436 | 0.0032766 | 0.0397202 |
| TMEM229B | 2.6445957 | 0.0026766 | 0.0237867 | GBP5 | 3.0807726 | 1.10E-08 | 3.37E-07 | SYNE1 | 2.7061149 | 7.07E-14 | 1.08E-11 |
| GNPTAB | 2.6374514 | 3.05E-07 | 7.77E-06 | C1orf21 | 3.0514876 | 0.0065484 | 0.0479958 | HLA-DMA | 2.6951835 | 0.0017262 | 0.0250088 |
| AOAH | 2.6353277 | 0.0011339 | 0.0117706 | CYP4F29P | 3.0453628 | 0.0008714 | 0.0086832 | CHN2 | 2.6831215 | 0.0028601 | 0.0356255 |
| APOBEC3H | 2.6208238 | 0.0020696 | 0.0193979 | TNFRSF1B | 3.0327589 | 2.53E-18 | 3.35E-16 | TBC1D7 | 2.6814668 | 0.0099421 | 0.0916868 |
| SEC14L1P1 | 2.5925537 | 0.0007234 | 0.0081876 | KIR3DL2 | 2.9940221 | 7.97E-05 | 0.001093 | STAG3L5P | 2.6771731 | 0.0013294 | 0.0203029 |
| TMEM2 | 2.5861495 | 3.59E-16 | 4.64E-14 | ZNF335 | 2.9895486 | 0.000558 | 0.0059369 | GSN | 2.6756494 | 9.70E-10 | 7.69E-08 |
| GOLGA8J | 2.5779117 | 0.0068329 | 0.0506469 | AGAP1 | 2.9781268 | 3.71E-06 | 6.72E-05 | ZNF600 | 2.6729336 | 1.13E-07 | 5.51E-06 |
| XKR8 | 2.5718553 | 0.007408 | 0.0538632 | RAB27B | 2.9620595 | 1.21E-06 | 2.48E-05 | RRN3P2 | 2.671762 | 0.0025661 | 0.0330799 |
| NHLRC3 | 2.5627979 | 0.001821 | 0.0174376 | PIK3R3 | 2.9470625 | 0.0002538 | 0.0029933 | UGGT2 | 2.6608296 | 0.0061509 | 0.0649706 |
| LLGL2 | 2.5601056 | 1.06E-05 | 0.0002002 | SYK | 2.9371061 | 2.85E-12 | 1.60E-10 | LRFN1 | 2.6505282 | 0.0063137 | 0.0661802 |
| TYROBP | 2.5498867 | 1.83E-16 | 2.48E-14 | ARHGEF12 | 2.9304823 | 2.20E-05 | 0.0003469 | CNNM2 | 2.648238 | 0.0022306 | 0.0298799 |
| YES1 | 2.546544 | 3.92E-08 | 1.16E-06 | TMEM229B | 2.9212117 | 0.000145 | 0.0018443 | MLX | 2.6438458 | 0.0057219 | 0.0614044 |
| MPP1 | 2.5423241 | 0.0091452 | 0.0633155 | TMEM91 | 2.9101413 | 0.0003403 | 0.0038702 | ST20 | 2.6432414 | 0.0061891 | 0.0653021 |
| MRPL15 | 2.539865 | 0.0057766 | 0.0441419 | IGF2R | 2.8976744 | 7.00E-14 | 4.91E-12 | TTC38 | 2.6298163 | 0.0003121 | 0.0062631 |
| VPS37B | 2.5391375 | 0.000156 | 0.0021516 | PPP2R2B | 2.8944817 | 9.44E-07 | 1.99E-05 | VAV3 | 2.6113235 | 1.93E-08 | 1.11E-06 |
| MCTP2 | 2.5359113 | 5.11E-12 | 3.16E-10 | BNC2 | 2.8919849 | 4.09E-08 | 1.13E-06 | FGD2 | 2.6031572 | 0.00244 | 0.0320581 |
| APBA2 | 2.5319785 | 0.0021304 | 0.0199008 | DOK2 | 2.8874777 | 2.38E-13 | 1.58E-11 | HIST1H2A | 2.5975653 | 2.40E-05 | 0.0006805 |
| PTPN12 | 2.5167917 | 5.22E-15 | 5.93E-13 | C6orf25 | 2.8808948 | 0.0034444 | 0.0280183 | LYST | 2.5953615 | 8.68E-17 | 1.81E-14 |
| FEM1A | 2.5145528 | 0.0060779 | 0.0460253 | AGTRAP | 2.8769461 | 0.0043664 | 0.0341382 | PRCC | 2.5787578 | 0.00078 | 0.0133209 |
| IGF2R | 2.5116586 | 1.52E-08 | 4.86E-07 | ZNF322 | 2.8643804 | 0.0012473 | 0.0117464 | CHST2 | 2.5712189 | 2.03E-08 | 1.17E-06 |
| ITGAL | 2.507017 | 3.75E-15 | 4.32E-13 | TYROBP | 2.8373452 | 3.20E-22 | 7.05E-20 | SLC9B2 | 2.5657383 | 0.0021939 | 0.0294703 |
| PRKCH | 2.5053387 | 9.23E-18 | 1.46E-15 | MCTP2 | 2.8274897 | 2.52E-18 | 3.35E-16 | SLFN12 | 2.5650075 | 0.0090911 | 0.08582 |
| IL18RAP | 2.5018367 | 3.69E-05 | 0.0006081 | CD300A | 2.8214918 | 4.57E-06 | 8.10E-05 | APMAP | 2.5607579 | 1.03E-15 | 1.97E-13 |
| ACAA2 | 2.4942926 | 0.0048683 | 0.0386065 | GOLGA8J | 2.8125433 | 0.0005147 | 0.0055406 | HLA-DPA1 | 2.5561547 | 2.05E-12 | 2.49E-10 |
| PHLDB2 | 2.4839599 | 1.15E-05 | 0.000214 | PHLDB2 | 2.792365 | 1.12E-08 | 3.41E-07 | CD300C | 2.551652 | 0.0093066 | 0.0870828 |
| PTGDR | 2.4822218 | 6.03E-07 | 1.47E-05 | BRCA1 | 2.7851186 | 0.0007134 | 0.0073133 | TCIRG1 | 2.5515245 | 1.37E-11 | 1.44E-09 |
| IFIT2 | 2.4821885 | 0.0006039 | 0.0070541 | SH2D1B | 2.7785894 | 2.01E-10 | 8.33E-09 | CXorf21 | 2.5387895 | 0.0009342 | 0.0154476 |
| DOK2 | 2.4731134 | 7.12E-08 | 2.01E-06 | ACAA2 | 2.7451102 | 0.0002206 | 0.0026622 | ATP8B4 | 2.5366747 | 8.36E-08 | 4.24E-06 |
| HDDC2 | 2.4683786 | 7.67E-07 | 1.84E-05 | CACNA1A | 2.7416248 | 0.0038785 | 0.0308828 | HLA-DRB1 | 2.5265552 | 0.0042128 | 0.0488813 |
| HEATR1 | 2.4557502 | 0.0023583 | 0.0216447 | HIST1H2AC | 2.737495 | 0.0004231 | 0.0046621 | TEC | 2.5183284 | 0.0002829 | 0.0057244 |
| GNG2 | 2.4536215 | 3.38E-24 | 9.79E-22 | RPL39L | 2.7320767 | 0.0004286 | 0.0047067 | CCL5 | 2.499258 | 5.57E-10 | 4.65E-08 |
| F2R | 2.4462886 | 0.0007404 | 0.0083465 | LMNB1 | 2.7214459 | 2.32E-05 | 0.0003624 | SCD | 2.4962435 | 0.0003468 | 0.0068305 |
| PLCG2 | 2.4391835 | 3.63E-08 | 1.09E-06 | PLEKHF1 | 2.7113958 | 5.26E-06 | 9.23E-05 | PYROXD2 | 2.4667852 | 0.0001593 | 0.0035617 |
| PRDM1 | 2.4366818 | 4.23E-10 | 1.83E-08 | RHOC | 2.7104914 | 2.41E-06 | 4.54E-05 | NHLRC3 | 2.4543308 | 0.0006551 | 0.0115664 |
| SLC4A4 | 2.4243147 | 3.02E-05 | 0.0005097 | DBNDD2 | 2.7042948 | 8.40E-05 | 0.0011401 | ID2 | 2.4463502 | 6.53E-21 | 2.32E-18 |
| HIRA | 2.4241757 | 2.97E-06 | 6.23E-05 | AGK | 2.659667 | 1.11E-08 | 3.41E-07 | LPCAT1 | 2.4458142 | 4.46E-12 | 5.16E-10 |
| MYBL1 | 2.4134011 | 2.93E-12 | 1.90E-10 | GK5 | 2.6594675 | 2.42E-15 | 2.22E-13 | PTPRJ | 2.4361274 | 4.77E-06 | 0.0001706 |

| | | | | | | | | | | | |
|---|---|---|---|---|---|---|---|---|---|---|---|
| SH2D2A | 2.4118126 | 1.03E-06 | 2.42E-05 | MYBL1 | 2.6542305 | 1.33E-18 | 1.87E-16 | C12orf79 | 2.435866 | 0.0097166 | 0.0902145 |
| THEMIS2 | 2.4082669 | 1.95E-06 | 4.31E-05 | PRDM1 | 2.639978 | 2.98E-16 | 3.10E-14 | CIDEB | 2.4197786 | 3.61E-08 | 1.95E-06 |
| RNF14 | 2.3997827 | 0.0010699 | 0.0112873 | ITGAL | 2.6289413 | 6.44E-19 | 9.31E-17 | AGAP5 | 2.4019586 | 0.0070388 | 0.0715297 |
| SBK1 | 2.3971644 | 1.77E-07 | 4.74E-06 | SLFN12 | 2.6022133 | 0.0099105 | 0.0668182 | HENMT1 | 2.4011428 | 0.0023752 | 0.0315078 |
| LINC01138 | 2.3959515 | 0.0040213 | 0.0331617 | HES6 | 2.5819118 | 0.0052477 | 0.040081 | CHST12 | 2.3977673 | 2.88E-05 | 0.0007988 |
| CENPF | 2.3944883 | 5.89E-08 | 1.69E-06 | MLLT11 | 2.5791804 | 0.0051218 | 0.0392811 | CPNE8 | 2.3902748 | 1.30E-06 | 5.25E-05 |
| SSH1 | 2.3941508 | 0.0002835 | 0.0036213 | FCER1G | 2.5738166 | 0.000896 | 0.0088707 | CARKD | 2.3742661 | 0.005513 | 0.0598138 |
| CD300A | 2.3834212 | 0.0008159 | 0.0090391 | PVRIG | 2.5726038 | 1.32E-07 | 3.32E-06 | COMMD4 | 2.3731223 | 0.0045486 | 0.0514453 |
| G6PD | 2.3747081 | 3.58E-13 | 2.84E-11 | KIR2DL4 | 2.5643757 | 0.005696 | 0.0430429 | DACH1 | 2.3720086 | 0.0032223 | 0.0391703 |
| TENM1 | 2.3743066 | 1.06E-07 | 2.90E-06 | ADAM28 | 2.5636027 | 0.0073259 | 0.0521521 | KLRK1 | 2.3632057 | 7.10E-09 | 4.54E-07 |
| GBP3 | 2.3651359 | 0.000737 | 0.0083195 | KLRC2 | 2.5591481 | 0.000101 | 0.0013353 | GPR68 | 2.3524554 | 0.0008576 | 0.0144815 |
| CMTR2 | 2.3499886 | 1.16E-05 | 0.0002164 | SLC15A4 | 2.5371745 | 6.46E-12 | 3.48E-10 | CYTH3 | 2.345374 | 0.002123 | 0.0287455 |
| GSDMB | 2.3451764 | 0.0070858 | 0.0520276 | ADHFE1 | 2.5366613 | 2.85E-05 | 0.0004362 | TMEM229 | 2.3439113 | 0.0037023 | 0.0439466 |
| HPS5 | 2.3293039 | 0.0009538 | 0.0102826 | LBH | 2.5278089 | 6.98E-11 | 3.23E-09 | HLA-DPB1 | 2.3330963 | 0.0002761 | 0.0056465 |
| HOPX | 2.323766 | 1.43E-09 | 5.48E-08 | CAPN5 | 2.5164782 | 0.005356 | 0.0408073 | IER3 | 2.3185793 | 1.84E-05 | 0.0005399 |
| KLHDC4 | 2.3141461 | 0.0048155 | 0.0383766 | THEMIS2 | 2.5021255 | 4.51E-07 | 1.03E-05 | ARHGEF12 | 2.3179957 | 0.0021282 | 0.0287492 |
| UBE2F | 2.3009712 | 3.69E-06 | 7.61E-05 | LLGL2 | 2.4972553 | 1.57E-05 | 0.0002543 | WHAMMP | 2.3151331 | 0.0003145 | 0.0062971 |
| APOL1 | 2.2942046 | 0.0005554 | 0.006579 | CD2 | 2.490055 | 4.52E-09 | 1.49E-07 | MCTP2 | 2.313953 | 7.05E-12 | 7.95E-10 |
| AUTS2 | 2.2922318 | 1.47E-05 | 0.0002682 | PTPN12 | 2.4891988 | 1.92E-17 | 2.27E-15 | TCEAL8 | 2.3118769 | 0.002125 | 0.0287455 |
| DDX28 | 2.270343 | 0.0056222 | 0.043282 | SBK1 | 2.4649186 | 1.76E-09 | 6.33E-08 | EFHD2 | 2.3069639 | 4.29E-10 | 3.77E-08 |
| B4GALT6 | 2.2581372 | 0.0022848 | 0.0210395 | NLRP2 | 2.4641682 | 0.000175 | 0.0021751 | SLC17A9 | 2.2977201 | 0.0001144 | 0.0026564 |
| PEAK1 | 2.2492484 | 0.0034121 | 0.0289155 | MIAT | 2.4617387 | 0.0013663 | 0.0127631 | MCC | 2.2929171 | 0.0005147 | 0.0094925 |
| ADHFE1 | 2.2412383 | 0.0012999 | 0.0132627 | YES1 | 2.4580813 | 7.74E-09 | 2.42E-07 | ACSL6 | 2.28276 | 0.0012854 | 0.0197881 |
| ZNF682 | 2.2394079 | 0.0062293 | 0.0470449 | GM2A | 2.4530503 | 0.0007572 | 0.0076943 | APEX2 | 2.2584095 | 0.0003048 | 0.0061416 |
| ZNF710 | 2.2388493 | 0.0063937 | 0.0479839 | SYNE2 | 2.4502993 | 1.27E-16 | 1.35E-14 | MIB2 | 2.25193 | 0.0087344 | 0.0835213 |
| SCARNA9 | 2.2363306 | 3.26E-05 | 0.0005413 | IFIT2 | 2.4399256 | 0.0002911 | 0.0033686 | FAM133B | 2.2515876 | 0.0007966 | 0.013546 |
| HLA-DPB1 | 2.2355031 | 0.0008223 | 0.0090984 | CD247 | 2.4352501 | 1.08E-16 | 1.17E-14 | TENM1 | 2.2427041 | 7.14E-07 | 3.03E-05 |
| TTC13 | 2.2347688 | 0.003759 | 0.031442 | ZNF600 | 2.4279066 | 8.85E-06 | 0.0001501 | PTGDR | 2.2221765 | 1.13E-05 | 0.0003553 |
| XCL2 | 2.233584 | 8.14E-07 | 1.95E-05 | PTGDR | 2.4264363 | 2.04E-06 | 3.93E-05 | STYXL1 | 2.1978339 | 0.0010559 | 0.0167947 |
| TFDP2 | 2.2296464 | 1.55E-10 | 7.39E-09 | MYO6 | 2.4072782 | 3.65E-05 | 0.000548 | ABHD2 | 2.1963451 | 1.62E-10 | 1.50E-08 |
| PVRIG | 2.227173 | 2.71E-05 | 0.0004666 | TENM1 | 2.4048178 | 4.98E-09 | 1.64E-07 | BCL6 | 2.1908215 | 0.0001716 | 0.0037832 |
| RHOC | 2.2163535 | 0.001741 | 0.016896 | PRCC | 2.4040434 | 0.0061904 | 0.0458807 | DOK2 | 2.1656605 | 3.87E-07 | 1.73E-05 |
| ZNF600 | 2.2104636 | 0.0001864 | 0.0025194 | ANXA4 | 2.3966111 | 2.50E-05 | 0.0003861 | SLC20A1 | 2.1641531 | 1.37E-09 | 1.04E-07 |
| SAYSD1 | 2.2102632 | 0.007563 | 0.054848 | FBXO32 | 2.3955654 | 2.26E-05 | 0.0003553 | ACAA2 | 2.1623617 | 0.003961 | 0.046499 |
| GSAP | 2.2041053 | 2.69E-07 | 6.90E-06 | GOLM1 | 2.3949449 | 0.0003664 | 0.004136 | TMEM11 | 2.1583763 | 0.0077356 | 0.0765273 |
| CD2 | 2.1929562 | 9.13E-06 | 0.0001746 | VAV3 | 2.389996 | 5.93E-07 | 1.32E-05 | GAS7 | 2.1523717 | 5.58E-08 | 2.91E-06 |
| TEX264 | 2.1883514 | 0.0040202 | 0.0331617 | TRIP4 | 2.3892153 | 0.002732 | 0.0230959 | GRAMD4 | 2.1465356 | 0.0008652 | 0.0145502 |
| FKTN | 2.1770869 | 0.0051165 | 0.040245 | PLCG2 | 2.3720516 | 3.58E-09 | 1.21E-07 | PSTPIP1 | 2.1399374 | 0.0001933 | 0.004176 |
| VNN2 | 2.152124 | 0.0001484 | 0.0020661 | TDRD7 | 2.3716837 | 0.0062042 | 0.0459455 | CTNNBIP1 | 2.1396755 | 5.55E-07 | 2.41E-05 |
| NAP1L5 | 2.1480913 | 0.008268 | 0.0589426 | MRPL15 | 2.3686465 | 0.0058447 | 0.0438093 | PPP1R9A | 2.1371454 | 2.59E-06 | 9.90E-05 |
| ABCC5 | 2.1400468 | 0.0009506 | 0.0102612 | GOLGA8N | 2.3645528 | 3.38E-06 | 6.19E-05 | SLFN13 | 2.1159998 | 1.33E-09 | 1.02E-07 |
| ITPRIP | 2.1381254 | 4.28E-05 | 0.0006927 | L3MBTL4-AS | 2.3537178 | 0.0014028 | 0.0130506 | GOLGA8N | 2.1154623 | 6.67E-05 | 0.0016319 |
| LYST | 2.1307997 | 3.13E-09 | 1.16E-07 | SCARNA17 | 2.3517069 | 0.0067659 | 0.0493198 | SIGLEC17F | 2.1154542 | 0.0012112 | 0.0188587 |
| PPP2R2B | 2.122686 | 0.0038075 | 0.0317531 | BZRAP1 | 2.3508163 | 1.33E-05 | 0.0002192 | RPE | 2.1067086 | 0.0010013 | 0.0162766 |
| S100PBP | 2.1100093 | 2.13E-05 | 0.0003771 | CTNNA1 | 2.3468705 | 0.0010259 | 0.0099757 | ENPP1 | 2.1066306 | 0.0018821 | 0.0265823 |
| SPIN4 | 2.1042854 | 0.0001108 | 0.0016127 | SMAD7 | 2.3343267 | 0.0002166 | 0.0026212 | LLGL2 | 2.0922701 | 0.000168 | 0.0037182 |
| NUP88 | 2.0989304 | 0.0040238 | 0.0331617 | B4GALT6 | 2.3252558 | 0.0006713 | 0.0069587 | B4GALT6 | 2.0899622 | 0.0019805 | 0.0274497 |
| IDH2 | 2.0973208 | 2.73E-05 | 0.0004696 | VPS37B | 2.2955352 | 0.0002317 | 0.0027786 | LINC0042 | 2.0877586 | 0.0003983 | 0.0076396 |
| C12orf75 | 2.0913962 | 1.47E-08 | 4.76E-07 | RGS9 | 2.2909501 | 0.0036791 | 0.0296152 | SEC14L1P1 | 2.0875662 | 0.001589 | 0.0233743 |
| IKZF3 | 2.0907281 | 1.73E-10 | 8.03E-09 | LINC01138 | 2.2903447 | 0.0043603 | 0.0341192 | TGFBR3 | 2.0779704 | 7.37E-10 | 5.99E-08 |
| CACNA2D2 | 2.0874557 | 0.0064644 | 0.0484284 | ZNF490 | 2.2760529 | 0.0018109 | 0.0161806 | SH2D1B | 2.0758799 | 4.62E-06 | 0.0001679 |
| COMT | 2.0733349 | 0.0018103 | 0.0173945 | MKI67 | 2.2727069 | 0.0074136 | 0.0527092 | HOPX | 2.0721679 | 3.34E-08 | 1.83E-06 |
| SYCP2 | 2.0689659 | 0.0022561 | 0.0208546 | PRKCH | 2.270295 | 5.19E-16 | 5.11E-14 | MAP4K3 | 2.0719732 | 0.0017155 | 0.024914 |
| ANXA4 | 2.0658974 | 0.0010484 | 0.0111032 | CHST12 | 2.2672185 | 0.0001205 | 0.0015628 | MATK | 2.0539751 | 3.90E-08 | 2.06E-06 |

| | | | | | | | | | | | |
|---|---|---|---|---|---|---|---|---|---|---|---|
| DTX3 | 2.063532 | 0.0006064 | 0.0070727 | HLA-DPB1 | 2.2574224 | 0.0002461 | 0.0029317 | ARAP2 | 2.0514176 | 5.20E-07 | 2.28E-05 |
| OAS2 | 2.060558 | 4.53E-05 | 0.0007227 | PRSS30P | 2.2560515 | 0.0083278 | 0.0582256 | PISD | 2.0325671 | 0.0037177 | 0.0440207 |
| HVCN1 | 2.0589244 | 0.0024828 | 0.0224923 | CD8A | 2.2476685 | 0.0022495 | 0.0195157 | NOTCH1 | 2.0304343 | 5.65E-06 | 0.0001969 |
| ABHD2 | 2.029975 | 9.29E-10 | 3.75E-08 | SH2D2A | 2.2048115 | 7.46E-07 | 1.63E-05 | CTSW | 1.9951995 | 5.54E-10 | 4.65E-08 |
| VPS26B | 2.0227206 | 2.88E-07 | 7.36E-06 | GAB3 | 2.2020682 | 0.0002081 | 0.0025317 | BZRAP1 | 1.9813957 | 0.0006153 | 0.0109891 |
| KLRC2 | 2.0226948 | 0.0083296 | 0.0592813 | ARHGEF9 | 2.1964224 | 1.02E-05 | 0.0001704 | ARHGEF9 | 1.9650527 | 5.98E-05 | 0.0014781 |
| ITGAX | 2.0202579 | 6.00E-08 | 1.72E-06 | RHBDF2 | 2.1955385 | 0.0007251 | 0.0074164 | TTC13 | 1.9646691 | 0.002132 | 0.028759 |
| STX12 | 2.0161054 | 0.0030308 | 0.0263213 | HIPK2 | 2.186006 | 3.24E-16 | 3.34E-14 | STK17A | 1.9626181 | 1.80E-14 | 2.88E-12 |
| ADA | 2.0134267 | 0.0001141 | 0.0016487 | AREL1 | 2.1852053 | 5.39E-05 | 0.0007734 | C4orf33 | 1.9510526 | 0.0098837 | 0.0914293 |
| BPGM | 1.9953132 | 0.0055929 | 0.0432497 | APOBR | 2.1556157 | 1.37E-06 | 2.78E-05 | PTPN12 | 1.9498687 | 8.87E-10 | 7.09E-08 |
| STX8 | 1.9935169 | 0.0079554 | 0.0572492 | ADCY9 | 2.1540684 | 0.0070877 | 0.0510195 | UACA | 1.9494263 | 0.0094827 | 0.0882997 |
| HIPK2 | 1.9930169 | 1.09E-12 | 7.82E-11 | MRPL10 | 2.1508563 | 8.65E-06 | 0.0001472 | APOL1 | 1.9409086 | 0.0017319 | 0.0250166 |
| MXRA7 | 1.9842942 | 0.0066758 | 0.0496572 | UBE2F | 2.1461969 | 1.56E-06 | 3.12E-05 | FES | 1.9387447 | 0.0042806 | 0.0492856 |
| AKAP2 | 1.9796356 | 0.0052543 | 0.0411364 | AUTS2 | 2.1364649 | 1.89E-05 | 0.0003027 | IFITM3 | 1.9357726 | 3.84E-08 | 2.03E-06 |
| CTBP2 | 1.9776818 | 8.00E-08 | 2.22E-06 | RAB9A | 2.1333377 | 0.0005164 | 0.0055515 | MAFF | 1.9257071 | 1.25E-06 | 5.09E-05 |
| FBXO32 | 1.9735126 | 0.0013862 | 0.0139414 | HOPX | 2.1271855 | 2.38E-08 | 6.77E-07 | LITAF | 1.9220559 | 1.35E-06 | 5.40E-05 |
| SYNE2 | 1.9557406 | 2.81E-11 | 1.53E-09 | ALOX5AP | 2.1193337 | 1.14E-08 | 3.47E-07 | GPR82 | 1.9197291 | 0.0049047 | 0.0548268 |
| SCARNA17 | 1.9526161 | 0.0023778 | 0.0217764 | LITAF | 2.1048157 | 8.72E-09 | 2.71E-07 | GBP5 | 1.9191214 | 0.001055 | 0.0167947 |
| SH2D1A | 1.9422459 | 2.48E-06 | 5.33E-05 | STX8 | 2.1022862 | 0.0008876 | 0.0088061 | RUNX3 | 1.9165982 | 1.09E-11 | 1.18E-09 |
| METRNL | 1.9417978 | 4.48E-05 | 0.0007188 | LINC00504 | 2.0984672 | 0.0003276 | 0.0037396 | ZEB2 | 1.9013332 | 2.86E-05 | 0.0007966 |
| STMN1 | 1.9381784 | 1.76E-05 | 0.0003171 | SLC16A6 | 2.0960325 | 0.0001448 | 0.0018443 | FMNL3 | 1.8930737 | 0.0051443 | 0.0566468 |
| RASA2 | 1.9337803 | 0.001084 | 0.0113798 | KIAA1377 | 2.0864505 | 0.0001711 | 0.0021374 | STARD9 | 1.8712132 | 0.001754 | 0.0251999 |
| KLRK1 | 1.9219343 | 1.75E-06 | 3.93E-05 | SSH1 | 2.0775815 | 0.0009221 | 0.0090902 | TMEM120 | 1.8646527 | 0.0090028 | 0.0853117 |
| PTPRE | 1.920187 | 2.83E-10 | 1.25E-08 | NPC1 | 2.0674984 | 0.0009241 | 0.0091001 | CRTAM | 1.8487237 | 0.0023116 | 0.0308776 |
| VAV3 | 1.9171601 | 0.0002132 | 0.0028244 | DLG3 | 2.0647951 | 0.0009187 | 0.009067 | ARID5A | 1.8438515 | 7.46E-05 | 0.0018025 |
| ARHGAP9 | 1.906382 | 1.11E-06 | 2.59E-05 | XCL2 | 2.0526357 | 8.68E-07 | 1.85E-05 | LTB4R | 1.843764 | 0.000953 | 0.0157046 |
| SASH3 | 1.9047266 | 0.0002312 | 0.0030427 | FAM179A | 2.0464852 | 0.0061794 | 0.0458725 | PPP2R2B | 1.8304492 | 0.0053326 | 0.0581851 |
| TRAPPC10 | 1.9027098 | 1.11E-07 | 3.00E-06 | SLA | 2.0463089 | 1.11E-11 | 5.72E-10 | ANO9 | 1.824642 | 1.74E-05 | 0.0005144 |
| PPP2R2D | 1.9018906 | 0.0024671 | 0.0223747 | IKZF3 | 2.0301211 | 2.23E-09 | 7.83E-08 | THAP3 | 1.8234729 | 0.0020875 | 0.0283585 |
| CORO1A | 1.8975219 | 0.0003616 | 0.0044986 | GNG2 | 2.0260092 | 3.93E-18 | 5.05E-16 | MNAT1 | 1.8148493 | 0.007657 | 0.0761807 |
| GAB3 | 1.8971396 | 0.0033259 | 0.0282989 | G6PD | 2.0215463 | 1.48E-09 | 5.35E-08 | METRNL | 1.806379 | 4.77E-06 | 0.0001706 |
| ARHGAP26 | 1.8879358 | 3.14E-05 | 0.0005251 | CPD | 2.0195485 | 3.89E-05 | 0.0005777 | G6PD | 1.805282 | 1.89E-08 | 1.10E-06 |
| RAB9A | 1.8858425 | 0.0085115 | 0.0603124 | LINC00861 | 2.0173751 | 9.15E-06 | 0.0001548 | SPDYE8P | 1.7922076 | 0.0089888 | 0.0852735 |
| ITGB2 | 1.8820088 | 8.24E-14 | 7.06E-12 | ADA | 2.0137885 | 1.52E-05 | 0.0002474 | IFIT2 | 1.7915616 | 0.0085321 | 0.0822422 |
| IFI16 | 1.8806004 | 1.67E-10 | 7.93E-09 | APOL1 | 2.0102875 | 0.0017015 | 0.0153512 | PDE4A | 1.7859265 | 3.17E-05 | 0.0008657 |
| APOBEC3C | 1.8698728 | 0.0001583 | 0.0021759 | PALLD | 1.9992728 | 0.0078645 | 0.0553203 | SCLY | 1.7804374 | 0.0004247 | 0.0080332 |
| NMUR1 | 1.8611503 | 0.0002087 | 0.0027782 | HDDC2 | 1.9925141 | 0.0001486 | 0.0018793 | NR4A3 | 1.777611 | 0.0077397 | 0.0765273 |
| ARPC5L | 1.8591975 | 0.0004045 | 0.0049516 | LAX1 | 1.9644003 | 0.0029551 | 0.0245117 | AGAP6 | 1.7717438 | 0.004251 | 0.0490587 |
| LINC00861 | 1.8553086 | 0.0008331 | 0.0091938 | CORO1A | 1.962111 | 3.91E-06 | 7.03E-05 | VPS37B | 1.7703328 | 0.0069546 | 0.0709589 |
| LOC102723 | 1.8551295 | 3.16E-05 | 0.000528 | GSAP | 1.9559528 | 4.69E-08 | 1.28E-06 | CD74 | 1.768475 | 3.81E-08 | 2.03E-06 |
| GOLGA8N | 1.8521445 | 0.0019482 | 0.0183624 | CMTR2 | 1.9500143 | 0.0002787 | 0.0032407 | BHLHE40 | 1.7666983 | 3.86E-05 | 0.0010294 |
| EIF4G3 | 1.8440566 | 7.16E-09 | 2.50E-07 | PTPN18 | 1.9389591 | 1.50E-07 | 3.71E-06 | B4GALT1 | 1.7623334 | 1.21E-08 | 7.18E-07 |
| STK35 | 1.8423687 | 0.0045368 | 0.0366347 | HECTD3 | 1.9329323 | 0.0073194 | 0.0521521 | DPF3 | 1.7591028 | 0.0065742 | 0.0681809 |
| SLAMF6 | 1.8370818 | 5.14E-05 | 0.0008065 | ABHD2 | 1.930713 | 4.43E-09 | 1.47E-07 | HIP1 | 1.7558698 | 0.0038993 | 0.0459434 |
| ATP6V0D1 | 1.8321587 | 0.0070585 | 0.0519973 | AMOT | 1.9237386 | 0.0087171 | 0.060445 | NCALD | 1.7513358 | 0.0042491 | 0.0490587 |
| RUNX3 | 1.8256334 | 6.59E-13 | 4.94E-11 | LNPEP | 1.923483 | 0.0018814 | 0.0167138 | ARPC5L | 1.7485281 | 9.32E-05 | 0.0022083 |
| LITAF | 1.8234641 | 2.12E-07 | 5.53E-06 | CTBP2 | 1.9231504 | 1.51E-08 | 4.43E-07 | SLA | 1.7356743 | 4.31E-06 | 0.0001577 |
| MICB | 1.8177708 | 0.0017085 | 0.0166061 | APOBEC3C | 1.922721 | 3.02E-08 | 8.51E-07 | UBE2F | 1.7296309 | 0.0011099 | 0.0174797 |
| TCIRG1 | 1.8173652 | 0.0001144 | 0.00165 | APOBEC3F | 1.9144377 | 0.0071755 | 0.051511 | RHOC | 1.7157714 | 0.0067936 | 0.0699113 |
| SLC20A1 | 1.8083902 | 1.41E-05 | 0.0002593 | ARHGAP26 | 1.9120277 | 1.91E-06 | 3.73E-05 | ADHFE1 | 1.7051743 | 0.0079851 | 0.0781645 |
| PATL2 | 1.8042956 | 0.0026381 | 0.0236092 | KLRK1 | 1.9096437 | 2.24E-06 | 4.25E-05 | CD63 | 1.6904433 | 8.05E-12 | 8.77E-10 |
| KIAA1377 | 1.7978917 | 0.0058071 | 0.0442946 | SLC35A4 | 1.9087241 | 0.0015288 | 0.0140401 | RAI1 | 1.6871339 | 0.001717 | 0.024914 |
| SLA2 | 1.7976852 | 0.0011094 | 0.0115741 | PEAK1 | 1.8984118 | 0.0097162 | 0.0657479 | AREG | 1.6798813 | 0.0003899 | 0.0075084 |
| HLA-DPA1 | 1.7969231 | 0.0002087 | 0.0027782 | TRAPPC10 | 1.8907013 | 8.46E-10 | 3.16E-08 | SAMD3 | 1.6755395 | 1.84E-05 | 0.0005404 |

| | | | | | | | | | | | |
|---|---|---|---|---|---|---|---|---|---|---|---|
| STOM | 1.7964439 | 3.75E-07 | 9.47E-06 | DTX3 | 1.8809941 | 0.0022708 | 0.0196638 | HPS5 | 1.6525093 | 0.0092598 | 0.0868989 |
| KLRC3 | 1.7933435 | 0.0018292 | 0.0174765 | TTC13 | 1.8749254 | 0.0057457 | 0.0431947 | S100PBP | 1.6478656 | 0.0002326 | 0.0048675 |
| SPN | 1.7863811 | 8.07E-06 | 0.0001562 | BPGM | 1.8741891 | 0.0029322 | 0.0243755 | ARHGAP2( | 1.645788 | 0.0002021 | 0.0043365 |
| TGFBR1 | 1.7863775 | 6.53E-05 | 0.0009953 | CIDEB | 1.8716078 | 0.0038231 | 0.0305368 | ABCB1 | 1.6233402 | 0.0027304 | 0.0345024 |
| TMEM181 | 1.7849188 | 1.28E-05 | 0.0002361 | ITGB2 | 1.8709014 | 2.11E-15 | 1.95E-13 | HSPA4 | 1.6184391 | 0.0097839 | 0.0906645 |
| ZAP70 | 1.7775875 | 2.19E-05 | 0.0003865 | TRAFD1 | 1.8699301 | 0.0012802 | 0.0120189 | ARHGAP9 | 1.6168558 | 9.56E-08 | 4.75E-06 |
| RAB8A | 1.7745856 | 0.0002061 | 0.0027516 | RNF14 | 1.869718 | 0.0068185 | 0.0496608 | ACSS1 | 1.6097422 | 4.64E-05 | 0.0011907 |
| KCTD10 | 1.7732242 | 7.98E-06 | 0.0001549 | PLCB1 | 1.8693285 | 0.0015492 | 0.0141988 | PLCB1 | 1.5985869 | 0.0067148 | 0.0692492 |
| CREM | 1.7706361 | 0.0022172 | 0.0205463 | PTPRE | 1.8673316 | 1.79E-11 | 9.07E-10 | ALDH3A2 | 1.5985745 | 0.0001524 | 0.0034226 |
| APLP2 | 1.7692658 | 2.87E-07 | 7.34E-06 | CYTH4 | 1.8628248 | 0.0018218 | 0.0162566 | SACM1L | 1.5972594 | 2.11E-08 | 1.20E-06 |
| ARHGEF3 | 1.7665255 | 0.000463 | 0.0055791 | SLC20A1 | 1.8628227 | 4.92E-07 | 1.11E-05 | SERPINB9 | 1.5865104 | 2.33E-07 | 1.09E-05 |
| PEF1 | 1.759051 | 0.0040669 | 0.0334515 | FNDC3B | 1.8516742 | 1.18E-08 | 3.56E-07 | MADD | 1.5706954 | 0.0001818 | 0.0039728 |
| STK26 | 1.7590265 | 2.08E-06 | 4.56E-05 | EIF4G3 | 1.8513362 | 4.44E-07 | 1.02E-05 | QSER1 | 1.5683336 | 0.0064882 | 0.0675655 |
| PTPN4 | 1.7586935 | 2.16E-08 | 6.69E-07 | PEF1 | 1.8503577 | 9.41E-05 | 0.0012587 | DUSP2 | 1.5645222 | 0.0002266 | 0.0047869 |
| MATK | 1.7580868 | 0.000171 | 0.0023316 | ITPRIP | 1.8490013 | 0.0002596 | 0.0030452 | CORO1A | 1.5595761 | 0.0003665 | 0.0071587 |
| PTPN18 | 1.757112 | 0.0001389 | 0.0019603 | PATL2 | 1.8488258 | 0.0002025 | 0.0024669 | NARF | 1.5591143 | 0.0039157 | 0.0460241 |
| SLA | 1.7518559 | 5.34E-06 | 0.0001066 | TGFBR1 | 1.8459052 | 2.28E-06 | 4.32E-05 | DHRS1 | 1.5590631 | 0.0002156 | 0.0045854 |
| SERPINB9 | 1.7509707 | 7.19E-06 | 0.0001414 | SIGLEC17P | 1.8444426 | 0.0084872 | 0.0591164 | PLSCR1 | 1.5556708 | 0.0027381 | 0.0345536 |
| COTL1 | 1.7481013 | 2.98E-06 | 6.23E-05 | SPPL2A | 1.8409336 | 0.0061411 | 0.0456952 | OTULIN | 1.5535794 | 0.0012423 | 0.0192357 |
| ALOX5AP | 1.7465716 | 2.42E-05 | 0.0004234 | ZAP70 | 1.8303286 | 1.18E-06 | 2.43E-05 | TPST2 | 1.551227 | 0.0003252 | 0.0064707 |
| GALNT3 | 1.7450201 | 0.0008407 | 0.0092655 | ABCC5 | 1.823022 | 0.0011566 | 0.0109855 | ITGAM | 1.5401526 | 7.70E-09 | 4.86E-07 |
| CROCCP2 | 1.7369389 | 0.0064243 | 0.0481707 | RUNX3 | 1.8159695 | 8.70E-14 | 6.06E-12 | FBXO18 | 1.5388114 | 0.0065181 | 0.0678043 |
| NFATC2 | 1.7364527 | 3.40E-10 | 1.48E-08 | TCIRG1 | 1.8153407 | 3.44E-05 | 0.000519 | BID | 1.5341935 | 0.0044299 | 0.0507613 |
| DENND2D | 1.7356137 | 0.0001524 | 0.0021084 | FGL2 | 1.8125081 | 0.0013493 | 0.0126163 | TNFRSF1B | 1.5337917 | 0.0001076 | 0.0025058 |
| CPD | 1.7320116 | 0.0009826 | 0.0105386 | ITGAX | 1.8115081 | 2.95E-07 | 6.91E-06 | VNN2 | 1.5330524 | 0.0026219 | 0.0335739 |
| FRMD8 | 1.727015 | 6.96E-05 | 0.0010539 | PTPN4 | 1.811015 | 2.33E-11 | 1.16E-09 | CD97 | 1.52404 | 1.15E-07 | 5.57E-06 |
| DIP2A | 1.7256221 | 3.09E-06 | 6.43E-05 | EPS15L1 | 1.8044298 | 0.0037132 | 0.029838 | PAPD5 | 1.510611 | 8.08E-06 | 0.0002689 |
| ASB1 | 1.71454 | 0.0039801 | 0.032931 | PRR5L | 1.8010285 | 5.10E-05 | 0.0007405 | HCST | 1.5065272 | 2.36E-07 | 1.10E-05 |
| ARHGEF9 | 1.7127512 | 0.0041101 | 0.0337737 | STOM | 1.7999617 | 3.32E-08 | 9.25E-07 | ITGAL | 1.504863 | 1.42E-06 | 5.67E-05 |
| C1orf174 | 1.7116377 | 0.0010607 | 0.0112044 | ZNF595 | 1.7927043 | 0.0019083 | 0.0169209 | CAMK2D | 1.5045211 | 0.0016655 | 0.0243139 |
| PDE4A | 1.7077054 | 0.0006723 | 0.0077022 | IFI16 | 1.791643 | 1.04E-10 | 4.67E-09 | CROCCP2 | 1.4999681 | 0.0070613 | 0.0715908 |
| RIPK2 | 1.7048503 | 0.0039939 | 0.0330128 | SYCP2 | 1.7876481 | 0.0047356 | 0.0367452 | FURIN | 1.4972554 | 9.45E-05 | 0.0022317 |
| C20orf24 | 1.701234 | 4.07E-05 | 0.0006608 | RAP1GAP2 | 1.7852734 | 1.23E-08 | 3.68E-07 | PIAS2 | 1.4957107 | 0.0003659 | 0.0071587 |
| RAP1GAP2 | 1.6940634 | 2.11E-08 | 6.57E-07 | MAN1A1 | 1.770427 | 2.36E-06 | 4.45E-05 | PRKCD | 1.4807743 | 0.0020126 | 0.0276946 |
| STK17A | 1.6938422 | 9.68E-07 | 2.30E-05 | NMUR1 | 1.7681335 | 0.0003067 | 0.0035177 | STK26 | 1.4762288 | 7.24E-06 | 0.0002464 |
| FAM49B | 1.6934837 | 1.21E-09 | 4.76E-08 | TFDP2 | 1.7626134 | 5.78E-08 | 1.56E-06 | RAB8A | 1.475575 | 0.000973 | 0.0159523 |
| PRKCB | 1.6922644 | 4.78E-08 | 1.40E-06 | ZNF117 | 1.7552052 | 0.0046865 | 0.0363951 | ZC3H18 | 1.4724002 | 0.0012435 | 0.0192357 |
| PTPN6 | 1.6915829 | 0.0092583 | 0.06394 | GSN | 1.7533167 | 0.0002228 | 0.0026824 | FYN | 1.4635717 | 1.99E-05 | 0.0005777 |
| GBP4 | 1.6885519 | 6.69E-07 | 1.62E-05 | JAZF1 | 1.751146 | 1.42E-06 | 2.87E-05 | CPD | 1.4631468 | 0.0020445 | 0.0280508 |
| ABHD17A | 1.6872787 | 6.04E-05 | 0.0009287 | S100PBP | 1.7497639 | 5.51E-05 | 0.000783 | DAPP1 | 1.4616418 | 0.0029556 | 0.0366247 |
| TRG-AS1 | 1.6849298 | 0.0055892 | 0.0432497 | VNN2 | 1.739136 | 0.0006053 | 0.0063961 | ADA | 1.4548979 | 0.0026659 | 0.0339552 |
| SAMD3 | 1.678368 | 2.18E-08 | 6.72E-07 | LYST | 1.737153 | 3.78E-07 | 8.72E-06 | LOC10013 | 1.4469941 | 0.0076886 | 0.0763969 |
| MAN1A1 | 1.6738797 | 0.0014014 | 0.0140442 | SLA2 | 1.7198394 | 0.0011653 | 0.0110414 | SH2D2A | 1.4440972 | 0.0027667 | 0.0348229 |
| PPP2R5A | 1.6705802 | 0.0001466 | 0.002048 | ARRB1 | 1.7148554 | 0.0003668 | 0.004136 | PEF1 | 1.4440508 | 0.0026958 | 0.0341546 |
| ASCC1 | 1.6703006 | 0.0090886 | 0.0631315 | C5orf56 | 1.7107208 | 1.94E-07 | 4.66E-06 | SH2D1A | 1.4387341 | 0.0006501 | 0.0115036 |
| FXR2 | 1.6686649 | 0.0013881 | 0.0139441 | VPS16 | 1.7070261 | 0.0039131 | 0.0310403 | MAP3K8 | 1.4373178 | 2.38E-07 | 1.10E-05 |
| DENND1C | 1.6557626 | 0.0005585 | 0.0066064 | ACTR1A | 1.7041402 | 0.0091058 | 0.0623924 | YARS | 1.4231081 | 0.0013787 | 0.0208891 |
| CDKN2C | 1.6432032 | 0.0081797 | 0.0584122 | ARAP2 | 1.7013285 | 8.79E-06 | 0.0001493 | GATAD1 | 1.4230798 | 0.000809 | 0.0137324 |
| SUSD1 | 1.636898 | 0.0001498 | 0.0020799 | IDH2 | 1.6971201 | 0.0003961 | 0.0044128 | UBE2Q2P1 | 1.420437 | 0.0058656 | 0.0626468 |
| TRAPPC2 | 1.632566 | 5.25E-06 | 0.000105 | DIP2A | 1.6955235 | 1.23E-07 | 3.12E-06 | GBP4 | 1.4159416 | 2.74E-05 | 0.0007711 |
| ARRB1 | 1.630711 | 0.001568 | 0.0154369 | ARHGEF3 | 1.6948938 | 6.45E-05 | 0.0009062 | CXorf40B | 1.4105838 | 0.0025414 | 0.0329831 |
| PSMB9 | 1.6298535 | 0.0002122 | 0.0028148 | MATK | 1.6941341 | 2.16E-05 | 0.0003407 | ACAA1 | 1.4022936 | 0.0064338 | 0.067213 |
| ARID5A | 1.6268695 | 0.0069595 | 0.0514944 | FYN | 1.6930229 | 8.78E-10 | 3.27E-08 | ZNF853 | 1.4005397 | 0.0073328 | 0.0734892 |
| USB1 | 1.6246643 | 0.0011926 | 0.012304 | C12orf75 | 1.6740122 | 1.30E-07 | 3.29E-06 | CALCOCO | 1.3826634 | 0.0009953 | 0.0162071 |

| | | | | | | | | | | | |
|---|---|---|---|---|---|---|---|---|---|---|---|
| YPEL1 | 1.6240689 | 0.0003973 | 0.0048855 | CBR4 | 1.6656238 | 0.0002426 | 0.002894 | AGPAT4 | 1.3805873 | 0.0005846 | 0.0105385 |
| MAPRE2 | 1.6081671 | 8.89E-05 | 0.0013185 | KLRC3 | 1.6530164 | 0.0016614 | 0.015034 | CLSTN3 | 1.378183 | 0.0046875 | 0.0527674 |
| RFTN1 | 1.6059634 | 0.000633 | 0.0073596 | CD2BP2 | 1.6522412 | 0.0080464 | 0.0563432 | LDLRAP1 | 1.3755476 | 0.0007477 | 0.0128291 |
| PPP1CA | 1.5995074 | 0.0003332 | 0.0041768 | GNS | 1.6462732 | 0.0001061 | 0.0013947 | GPRIN3 | 1.3694124 | 0.0080642 | 0.0786815 |
| ZNF136 | 1.5993049 | 0.0044023 | 0.0356515 | SLAMF6 | 1.6407574 | 0.0017461 | 0.0156476 | DNAJC1 | 1.3680814 | 0.0017546 | 0.0251999 |
| AMZ2 | 1.5833735 | 3.07E-05 | 0.0005149 | YPEL1 | 1.640549 | 8.14E-05 | 0.0011124 | RFTN1 | 1.3658122 | 0.0017551 | 0.0251999 |
| PLEKHA1 | 1.5831901 | 3.83E-05 | 0.0006275 | METRNL | 1.6362977 | 5.66E-05 | 0.0008005 | HSH2D | 1.3609649 | 3.40E-05 | 0.0009208 |
| GCH1 | 1.581688 | 8.17E-06 | 0.0001577 | HLA-DPA1 | 1.6321014 | 0.0001788 | 0.0022118 | SLC44A1 | 1.3547444 | 0.0001658 | 0.0036804 |
| TOX | 1.5725236 | 1.73E-05 | 0.0003113 | SASH3 | 1.6314482 | 0.0006095 | 0.006426 | PPP2R2D | 1.3442954 | 0.0036954 | 0.043919 |
| DGKQ | 1.5673356 | 0.0002502 | 0.0032683 | PSTPIP1 | 1.6302191 | 0.0050426 | 0.0387701 | LINC01278 | 1.3380795 | 0.0094753 | 0.0882997 |
| IFNGR1 | 1.566346 | 0.0002606 | 0.0033773 | ARPC5L | 1.6256591 | 0.0006426 | 0.0067368 | JAK1 | 1.3369973 | 1.48E-05 | 0.0004476 |
| C5orf56 | 1.5613032 | 1.72E-05 | 0.0003106 | NFIL3 | 1.6252023 | 2.38E-05 | 0.0003709 | CLDND1 | 1.3187002 | 0.000202 | 0.0043365 |
| RNF213 | 1.5535505 | 2.79E-09 | 1.04E-07 | ZNF266 | 1.622557 | 0.0090515 | 0.0621585 | CTBP2 | 1.3110121 | 0.0002329 | 0.0048675 |
| MRPS5 | 1.551331 | 0.0072605 | 0.0529121 | NFATC2 | 1.6223785 | 5.52E-10 | 2.16E-08 | PSMB9 | 1.303771 | 0.0005715 | 0.0103614 |
| RAB29 | 1.5445837 | 1.98E-05 | 0.0003537 | MTSS1 | 1.6102247 | 0.000118 | 0.0015405 | AKNA | 1.3000298 | 1.10E-05 | 0.0003493 |
| PAPD5 | 1.532076 | 6.98E-05 | 0.0010548 | UBFD1 | 1.607131 | 1.68E-05 | 0.0002704 | GK5 | 1.2948992 | 0.0003319 | 0.0065913 |
| JAZF1 | 1.5287516 | 0.0002044 | 0.0027416 | KCNAB2 | 1.6037166 | 4.96E-05 | 0.0007229 | TRAPPC10 | 1.2851341 | 5.40E-05 | 0.0013531 |
| TMIGD2 | 1.5284854 | 0.0034984 | 0.0295867 | TXK | 1.6029987 | 0.0009638 | 0.0094415 | SIPA1 | 1.2817642 | 0.0040663 | 0.047619 |
| PRR5L | 1.5254256 | 0.0019039 | 0.0180408 | GBP4 | 1.5961841 | 6.36E-07 | 1.41E-05 | PRKCH | 1.2766841 | 1.09E-05 | 0.0003457 |
| ITCH | 1.5237426 | 0.0038248 | 0.0318658 | SPN | 1.582446 | 4.57E-05 | 0.0006679 | STX4 | 1.2645247 | 0.0068247 | 0.0700811 |
| WHAMMP2 | 1.5191835 | 0.0007135 | 0.0080974 | SEC23IP | 1.5791928 | 0.0037767 | 0.0302429 | SELL | 1.2574824 | 0.0005828 | 0.0105267 |
| CDK19 | 1.5181553 | 0.0025332 | 0.022875 | ARHGAP25 | 1.5759448 | 0.0003005 | 0.0034643 | GSTP1 | 1.2462637 | 0.0055086 | 0.0598138 |
| NFIL3 | 1.5165923 | 0.0017802 | 0.0171639 | SYTL2 | 1.5716891 | 3.69E-05 | 0.0005522 | NDUFB7 | 1.2376913 | 0.0068574 | 0.0702186 |
| APOL6 | 1.5157212 | 1.68E-06 | 3.80E-05 | GPR65 | 1.5695263 | 5.09E-05 | 0.0007401 | ATHL1 | 1.2316627 | 0.005186 | 0.05691 |
| TAP1 | 1.5150019 | 0.0027278 | 0.0240882 | TBC1D1 | 1.5533498 | 0.0083935 | 0.0585934 | WHAMMP | 1.2247953 | 0.0026048 | 0.0334436 |
| LGALS9 | 1.5097264 | 0.0004556 | 0.0055127 | NOTCH1 | 1.5526227 | 0.0007331 | 0.0074906 | DIP2A | 1.2176451 | 0.0005424 | 0.0099284 |
| TOPBP1 | 1.4942991 | 0.0011252 | 0.0117094 | ABCB1 | 1.5460883 | 0.0089449 | 0.061614 | CBLB | 1.2152574 | 0.0052848 | 0.0577954 |
| TXNRD1 | 1.4843931 | 0.00305 | 0.0264329 | TRAPPC2 | 1.5193833 | 2.41E-06 | 4.54E-05 | FLNB | 1.2135847 | 0.0091173 | 0.0859825 |
| LASP1 | 1.4839405 | 2.41E-07 | 6.24E-06 | PRKCB | 1.5164928 | 2.69E-07 | 6.35E-06 | ZMAT2 | 1.2100166 | 0.000336 | 0.0066447 |
| STAM2 | 1.4802119 | 0.0075542 | 0.0548319 | SNX18 | 1.5162956 | 0.0001379 | 0.0017658 | FUT8 | 1.2088945 | 0.0031587 | 0.0384449 |
| ATP2B4 | 1.4772536 | 4.29E-07 | 1.07E-05 | DIAPH2 | 1.5139857 | 0.0017131 | 0.0154415 | YES1 | 1.2032756 | 0.0086303 | 0.0828424 |
| PCNT | 1.4770865 | 1.24E-05 | 0.000229 | KCTD10 | 1.5122054 | 9.42E-05 | 0.0012587 | CDYL | 1.1978823 | 0.0036075 | 0.0430343 |
| AKNA | 1.4568967 | 1.10E-07 | 2.98E-06 | PIN4 | 1.5085619 | 0.00809 | 0.0566055 | SRGN | 1.1879087 | 1.25E-05 | 0.0003857 |
| SCP2 | 1.4564915 | 0.0002487 | 0.0032532 | PTPN22 | 1.5040333 | 1.89E-06 | 3.71E-05 | ARHGAP2! | 1.1876375 | 0.0056862 | 0.0611396 |
| ACSL5 | 1.451237 | 0.0018461 | 0.0175884 | MAST3 | 1.498523 | 0.0023154 | 0.0200311 | ZAP70 | 1.1870263 | 0.0050482 | 0.0559737 |
| NOTCH1 | 1.4498757 | 0.0036367 | 0.0306325 | TMEM2 | 1.495887 | 2.09E-07 | 5.01E-06 | ATP6V0D1 | 1.1731083 | 0.0065757 | 0.0681809 |
| VAV1 | 1.4458721 | 0.0047746 | 0.0381875 | FRMD8 | 1.4942257 | 7.60E-05 | 0.0010468 | CTSD | 1.1619818 | 0.0003812 | 0.0073714 |
| ARHGAP25 | 1.4391209 | 0.001284 | 0.0131342 | SLC25A42 | 1.4938515 | 0.0026713 | 0.0226657 | ITGB2 | 1.1516702 | 9.93E-06 | 0.0003207 |
| IRF2 | 1.4386765 | 0.0055702 | 0.0431932 | ARHGAP9 | 1.490129 | 1.60E-06 | 3.17E-05 | KCNAB2 | 1.1458145 | 0.0049236 | 0.0549732 |
| SLFN13 | 1.4345951 | 0.0001813 | 0.0024594 | SAP30L-AS1 | 1.4890744 | 0.0010834 | 0.0104249 | RPAP3 | 1.1366604 | 0.0053144 | 0.0580531 |
| LOC388692 | 1.4327392 | 0.0020959 | 0.0196227 | EIF2B2 | 1.4887553 | 0.0097059 | 0.0657264 | PRKCB | 1.1240747 | 0.0005769 | 0.0104406 |
| FYN | 1.4239837 | 4.51E-06 | 9.15E-05 | ZNF410 | 1.4855509 | 0.0036678 | 0.0295501 | APOBEC3( | 1.1162751 | 0.0031295 | 0.0381784 |
| ATP1B3 | 1.4236922 | 0.0002837 | 0.0036213 | AKNA | 1.4815474 | 2.58E-08 | 7.30E-07 | SP140 | 1.1138007 | 0.0011575 | 0.0181402 |
| CAMK2D | 1.4231012 | 0.0056229 | 0.043282 | ABHD17A | 1.4766878 | 0.0004437 | 0.0048662 | PLAC8 | 1.1136592 | 0.001948 | 0.0271948 |
| CBR4 | 1.4174691 | 0.0020673 | 0.0193979 | PIAS2 | 1.4740178 | 0.0024859 | 0.0212905 | PILRB | 1.1119411 | 0.0010522 | 0.0167947 |
| VASP | 1.4159022 | 0.0052394 | 0.0410832 | GLTP | 1.4696182 | 3.08E-07 | 7.19E-06 | MBP | 1.1079531 | 3.51E-05 | 0.0009492 |
| YARS | 1.4146352 | 0.0085662 | 0.0605546 | CASP1 | 1.4590594 | 0.0001236 | 0.0015982 | GLTP | 1.105906 | 8.74E-05 | 0.0020778 |
| TROVE2 | 1.4117794 | 0.0009227 | 0.0099861 | LINC01278 | 1.4566542 | 0.0031378 | 0.0258425 | BACH2 | 1.1045603 | 5.08E-05 | 0.0012848 |
| CASP1 | 1.4117348 | 0.0071526 | 0.0524608 | CYBA | 1.4547806 | 0.0021518 | 0.0187921 | RAB1B | 1.0964504 | 0.0001683 | 0.0037182 |
| BSG | 1.4089047 | 0.0088916 | 0.0621823 | FAM49B | 1.4526152 | 1.43E-08 | 4.23E-07 | UTRN | 1.0873492 | 0.0008273 | 0.0139938 |
| CTSC | 1.4048101 | 0.0011861 | 0.012252 | PIK3R2 | 1.4523572 | 0.003262 | 0.0267226 | KMT2E-AS | 1.0829839 | 0.0029991 | 0.0369245 |
| BTN3A2 | 1.4040903 | 0.0001286 | 0.0018367 | TMEM181 | 1.4425557 | 0.0001139 | 0.0014907 | CHD9 | 1.0760836 | 0.0006179 | 0.0110147 |
| DGKZ | 1.4019258 | 0.0097101 | 0.0665678 | TPST2 | 1.4423079 | 0.0009752 | 0.0095429 | ZFHX3 | 1.0752037 | 0.0054426 | 0.059184 |
| CLPTM1L | 1.3975332 | 0.0056792 | 0.0435161 | GALNT2 | 1.4418303 | 0.0071519 | 0.0514015 | PDCD7 | 1.0570425 | 0.0019637 | 0.0272951 |

| | | | | | | | | | | | |
|---|---|---|---|---|---|---|---|---|---|---|---|
| ARAP2 | 1.3937233 | 0.0006685 | 0.0076796 | SERPINB9 | 1.4370971 | 6.65E-05 | 0.0009287 | AGPAT5 | 1.0531058 | 0.0063052 | 0.0661632 |
| GPR65 | 1.3895584 | 0.0011831 | 0.0122362 | SH2D1A | 1.4357895 | 0.0008369 | 0.0084023 | CD247 | 1.0506559 | 0.0007178 | 0.0124722 |
| ATAD5 | 1.3860371 | 0.0027213 | 0.0240562 | STAM2 | 1.4335753 | 0.0023381 | 0.0202088 | CTAGE5 | 1.0477305 | 0.0034313 | 0.0413434 |
| DNAJC1 | 1.3837219 | 0.0058984 | 0.0449093 | DHRS1 | 1.4306201 | 0.0029202 | 0.0243096 | NDUFV1 | 1.0462086 | 0.0066846 | 0.0690113 |
| SP140 | 1.3631003 | 0.0002593 | 0.0033654 | ETAA1 | 1.4286149 | 0.0090877 | 0.0623608 | NLRC5 | 1.0401526 | 0.0018381 | 0.0261541 |
| DNM2 | 1.3615093 | 0.0066035 | 0.0492948 | PPP1CA | 1.4274257 | 0.0006452 | 0.0067492 | RAB29 | 1.0345492 | 0.001908 | 0.0267612 |
| COX8A | 1.3551221 | 0.0026398 | 0.0236092 | TBC1D31 | 1.4183222 | 0.0076341 | 0.0539456 | CPNE1 | 1.0195356 | 0.0008138 | 0.0137901 |
| DBI | 1.3532465 | 0.0005931 | 0.0069372 | HCST | 1.4152276 | 5.31E-06 | 9.29E-05 | STIM2 | 1.0148174 | 0.0010502 | 0.0167947 |
| APOBEC3G | 1.3471046 | 0.0052559 | 0.0411364 | MBP | 1.4144105 | 2.26E-07 | 5.38E-06 | ESYT2 | 1.005918 | 0.0014987 | 0.0224589 |
| CTBP1 | 1.3457785 | 3.95E-06 | 8.08E-05 | CLIP4 | 1.413351 | 0.0014272 | 0.0132384 | YBX1 | -1.000172 | 0.0001065 | 0.002485 |
| COPZ1 | 1.3413144 | 0.0014549 | 0.0144706 | MAPRE2 | 1.4089517 | 0.0001579 | 0.0019892 | RSBN1 | -1.005241 | 0.0089423 | 0.0850007 |
| JAK1 | 1.3386798 | 7.04E-06 | 0.0001392 | UTRN | 1.405852 | 1.41E-07 | 3.49E-06 | SOD1 | -1.00669 | 0.0013071 | 0.0199947 |
| SRPK2 | 1.3332116 | 8.74E-06 | 0.0001675 | DENND1C | 1.4006933 | 0.0023419 | 0.0202226 | KLF12 | -1.007554 | 0.0063025 | 0.0661632 |
| VOPP1 | 1.3297049 | 1.94E-05 | 0.0003471 | PPP2R5A | 1.3979305 | 0.0011 | 0.0105412 | EPM2AIP1 | -1.009033 | 0.0042149 | 0.0488813 |
| KCNAB2 | 1.3242809 | 0.0024493 | 0.0222368 | SP140 | 1.3958271 | 0.0002016 | 0.0024585 | TOMM20 | -1.010382 | 0.0013616 | 0.0207285 |
| FAM8A1 | 1.324163 | 0.0015282 | 0.0151525 | LINC00623 | 1.3937109 | 0.0092367 | 0.0631026 | RASGRP2 | -1.013181 | 0.0015525 | 0.0230496 |
| CWC22 | 1.3223858 | 0.0028092 | 0.0247034 | RPA2 | 1.3927326 | 0.0006298 | 0.0066249 | CBX7 | -1.014473 | 0.0042465 | 0.0490587 |
| ALDH9A1 | 1.3194661 | 0.0026553 | 0.0236919 | ZBTB38 | 1.3914264 | 0.001251 | 0.0117691 | KLF6 | -1.015732 | 0.0013565 | 0.0206845 |
| PIAS2 | 1.3045829 | 0.0088927 | 0.0621823 | PHPT1 | 1.371071 | 0.0050529 | 0.0388168 | RHOH | -1.022719 | 0.0035317 | 0.0422878 |
| PPP1R18 | 1.3035577 | 1.24E-06 | 2.86E-05 | MRE11A | 1.3681535 | 0.0001861 | 0.002291 | HSD17B11 | -1.024655 | 0.0042083 | 0.0488813 |
| MBP | 1.2980325 | 1.03E-06 | 2.42E-05 | SAMD3 | 1.3675161 | 5.80E-05 | 0.0008168 | UBE2H | -1.025375 | 0.0001211 | 0.002792 |
| VAMP1 | 1.2975939 | 0.0029496 | 0.0256957 | CBX1 | 1.3651735 | 0.0018566 | 0.01651 | FAM84B | -1.027043 | 0.0085052 | 0.0821481 |
| RAP1B | 1.2972374 | 2.07E-06 | 4.53E-05 | RNF213 | 1.3645571 | 4.33E-09 | 1.44E-07 | HMGCS1 | -1.030014 | 0.0034918 | 0.0419144 |
| ID2 | 1.2969669 | 1.31E-06 | 3.00E-05 | STK10 | 1.3615755 | 2.58E-05 | 0.0003977 | GIMAP1 | -1.031545 | 0.0024496 | 0.0321121 |
| DGKD | 1.2938271 | 0.0041444 | 0.0339556 | ATP6V0D1 | 1.3552382 | 0.0064538 | 0.0474147 | GNG2 | -1.035692 | 0.0046336 | 0.0523451 |
| MYO1F | 1.2903388 | 7.55E-05 | 0.0011319 | DNAJC1 | 1.3536372 | 0.0015918 | 0.0144891 | ABCE1 | -1.036839 | 0.0008662 | 0.0145502 |
| PLAC8 | 1.2902733 | 4.30E-05 | 0.0006927 | SCP2 | 1.3496518 | 0.0001197 | 0.0015542 | ASPH | -1.040161 | 0.001529 | 0.0227364 |
| TM9SF2 | 1.2864828 | 0.0024139 | 0.0220395 | CTSW | 1.3467364 | 0.000236 | 0.0028259 | CKAP2 | -1.042726 | 0.0079413 | 0.0781645 |
| NCKAP1L | 1.283995 | 0.0018841 | 0.0178786 | BTN3A2 | 1.3438234 | 6.51E-05 | 0.0009131 | NOL9 | -1.045683 | 0.0009089 | 0.0151086 |
| IQGAP1 | 1.2810505 | 1.12E-05 | 0.0002102 | HEG1 | 1.3403003 | 0.0026969 | 0.022841 | ARRDC3 | -1.045722 | 2.99E-05 | 0.0008252 |
| PSMB10 | 1.2789124 | 0.0004171 | 0.0050773 | loc4618 | 1.3334742 | 1.41E-05 | 0.0002304 | ARNTL | -1.048612 | 0.0060433 | 0.0639751 |
| SERPINB1 | 1.2775567 | 0.0004886 | 0.005877 | DENND2D | 1.3317733 | 0.0035392 | 0.0287312 | ASXL1 | -1.054067 | 0.0083503 | 0.0809781 |
| ARHGAP27 | 1.2709157 | 0.0051229 | 0.040245 | VAV1 | 1.3265637 | 0.0065471 | 0.0479958 | DNAJB1 | -1.054885 | 0.0007467 | 0.0128291 |
| HTT | 1.2634355 | 0.008429 | 0.0598872 | VPS26B | 1.3251705 | 0.0007432 | 0.0075682 | TXNIP | -1.064015 | 0.0002757 | 0.0056465 |
| NCOA1 | 1.2597833 | 2.32E-06 | 5.00E-05 | MAP2K4 | 1.3248048 | 0.0013459 | 0.0125976 | PABPC4 | -1.067721 | 0.0037717 | 0.0446044 |
| UTRN | 1.2573304 | 2.23E-06 | 4.85E-05 | APOBEC3G | 1.3168444 | 0.0036302 | 0.0293491 | NCOA6 | -1.075693 | 0.007304 | 0.0733411 |
| RNF113A | 1.2380083 | 0.0081634 | 0.0583457 | HIVEP3 | 1.3120084 | 0.000692 | 0.007149 | LMAN1 | -1.07594 | 0.0004551 | 0.008525 |
| ARPC5 | 1.2274067 | 0.0031637 | 0.0271106 | CDK2AP2 | 1.3118808 | 0.0078887 | 0.0554486 | SARAF | -1.079894 | 0.000188 | 0.0040804 |
| CTSD | 1.2239844 | 0.0005067 | 0.0060698 | CD97 | 1.3073629 | 4.54E-06 | 8.08E-05 | ARHGAP1 | -1.082559 | 0.0005467 | 0.0099873 |
| RAC2 | 1.2196196 | 4.93E-08 | 1.44E-06 | ATP2B4 | 1.3016039 | 1.12E-06 | 2.33E-05 | SLC30A7 | -1.083984 | 0.0035733 | 0.0427292 |
| BRK1 | 1.2125018 | 0.0017558 | 0.0169876 | DGKD | 1.3005469 | 0.0024274 | 0.020847 | RHOG | -1.084436 | 0.0086174 | 0.0828344 |
| SLC35A3 | 1.2107207 | 0.0040659 | 0.0334515 | MMP24-AS1 | 1.2945581 | 0.0083604 | 0.0584093 | SLCO3A1 | -1.090453 | 0.0063681 | 0.0666778 |
| STARD3NL | 1.2098519 | 0.0015312 | 0.0151635 | LASP1 | 1.2927032 | 3.39E-06 | 6.19E-05 | ATXN2 | -1.093052 | 0.0010439 | 0.0167702 |
| ARPC4 | 1.2014334 | 8.41E-06 | 0.0001621 | MAPK1 | 1.2916769 | 3.78E-07 | 8.72E-06 | TNK2 | -1.097564 | 0.0004853 | 0.0090087 |
| ADD1 | 1.2009079 | 5.40E-05 | 0.0008422 | SLFN13 | 1.2913154 | 0.0011522 | 0.0109621 | FAM199X | -1.099378 | 0.0050844 | 0.0562447 |
| TERF1 | 1.1959621 | 0.0002409 | 0.0031606 | STX4 | 1.2866217 | 0.0070021 | 0.0505605 | SNRPE | -1.123093 | 0.0019576 | 0.0272503 |
| LRMP | 1.1958398 | 0.0043757 | 0.0355046 | BTBD9 | 1.2862734 | 0.0007908 | 0.0079901 | HIPK3 | -1.123884 | 1.50E-05 | 0.0004519 |
| MTSS1 | 1.1899424 | 0.0091183 | 0.0632333 | RFTN1 | 1.2842353 | 0.0059912 | 0.0447989 | PRKCQ | -1.12579 | 0.0014373 | 0.0217425 |
| IDS | 1.1803101 | 0.0002022 | 0.0027217 | IFNGR1 | 1.2824002 | 0.0014547 | 0.0134645 | WWP1 | -1.125881 | 0.0001854 | 0.0040329 |
| ZMAT2 | 1.1750999 | 0.0001766 | 0.0024038 | RAB8A | 1.2807101 | 0.0080271 | 0.0562508 | SPIN1 | -1.126139 | 0.005553 | 0.060179 |
| SACM1L | 1.1658235 | 0.0001442 | 0.0020223 | STK17A | 1.2786547 | 9.17E-06 | 0.000155 | DUSP1 | -1.128002 | 0.0003371 | 0.0066535 |
| ATP8A1 | 1.16328 | 0.0020012 | 0.0188336 | CAMK2D | 1.2776507 | 0.0073064 | 0.0521493 | ZFP36L1 | -1.136504 | 2.51E-05 | 0.0007114 |
| RAB1B | 1.1587447 | 0.0013252 | 0.0134404 | CDC42EP3 | 1.2768744 | 0.0017716 | 0.0158604 | LAT | -1.139445 | 0.0017844 | 0.0255462 |
| ITGAM | 1.1584112 | 3.53E-05 | 0.0005833 | GATAD1 | 1.2756846 | 0.0044773 | 0.0348876 | TUBA1A | -1.141005 | 2.15E-05 | 0.0006171 |

| | | | | | | | | | | | |
|---|---|---|---|---|---|---|---|---|---|---|---|
| GNS | 1.1553337 | 0.0058772 | 0.044788 | CD63 | 1.2739036 | 1.56E-06 | 3.12E-05 | ZC3HAV1 | -1.145232 | 2.69E-05 | 0.0007584 |
| ACTR2 | 1.1508266 | 1.94E-06 | 4.29E-05 | PLEKHA1 | 1.2726783 | 0.0015597 | 0.0142807 | SLC16A7 | -1.145723 | 0.0005685 | 0.0103274 |
| CD81 | 1.1475354 | 0.0001692 | 0.0023108 | PAPD5 | 1.2684739 | 0.0006071 | 0.0064077 | HMG20A | -1.147457 | 0.0062459 | 0.0657565 |
| WDR11 | 1.14122 | 0.0016281 | 0.0159542 | ZNF746 | 1.2604351 | 0.0025656 | 0.0218895 | DTX3L | -1.151506 | 0.0099136 | 0.0915121 |
| loc4618 | 1.1404735 | 0.0056201 | 0.043282 | DNAJC5 | 1.2589304 | 0.0031127 | 0.0256583 | AGL | -1.157732 | 0.0057237 | 0.0614044 |
| GLTP | 1.1383285 | 0.0016632 | 0.0162414 | CTSD | 1.2556401 | 0.0001014 | 0.0013385 | CYP51A1 | -1.159139 | 0.0044835 | 0.0510994 |
| TRIM26 | 1.1319779 | 0.0070869 | 0.0520276 | ID2 | 1.254352 | 9.62E-07 | 2.02E-05 | JUND | -1.161315 | 5.25E-05 | 0.0013263 |
| SAMD9L | 1.1270998 | 0.0008997 | 0.0097994 | GSTP1 | 1.2455332 | 0.0071783 | 0.051511 | SLC33A1 | -1.166149 | 0.0057547 | 0.0615992 |
| STIM2 | 1.1270047 | 0.0027074 | 0.0240091 | BTN3A1 | 1.2408785 | 9.50E-05 | 0.0012675 | FAM129A | -1.168059 | 0.0039154 | 0.0460241 |
| ADAM8 | 1.126715 | 0.0006336 | 0.0073596 | STK26 | 1.239473 | 0.0003033 | 0.0034831 | LIN7C | -1.168155 | 0.0025661 | 0.0330799 |
| STK10 | 1.1252766 | 0.000914 | 0.0099294 | NCOA1 | 1.2390598 | 2.89E-06 | 5.41E-05 | ANXA5 | -1.184118 | 0.0019945 | 0.0275245 |
| HELZ | 1.1174602 | 0.0001651 | 0.0022619 | RGS2 | 1.2338056 | 0.0091533 | 0.0626719 | TTC39C | -1.184152 | 0.0003856 | 0.0074421 |
| EAF1 | 1.1170697 | 0.0029546 | 0.0257124 | RAB29 | 1.2241421 | 0.0001854 | 0.0022856 | TSPYL1 | -1.184215 | 0.0012414 | 0.0192357 |
| SRGN | 1.108018 | 6.21E-05 | 0.0009513 | SRGN | 1.2230828 | 1.95E-06 | 3.80E-05 | ATP11B | -1.185384 | 0.0040375 | 0.0473398 |
| ZMAT3 | 1.1043983 | 0.0080627 | 0.0577735 | APLP2 | 1.2071002 | 0.0004165 | 0.0046014 | USP53 | -1.185468 | 0.0011308 | 0.0177802 |
| SRSF1 | 1.1015286 | 0.0005709 | 0.0067149 | C20orf24 | 1.2026046 | 0.0075706 | 0.0535789 | SESN1 | -1.18654 | 0.0017316 | 0.0250166 |
| BTN3A1 | 1.1001487 | 0.0077874 | 0.0562324 | TRIM56 | 1.196003 | 0.0041026 | 0.0323489 | CRKL | -1.191244 | 0.0057493 | 0.0615992 |
| NADK | 1.0967805 | 0.0070338 | 0.0519069 | STIM2 | 1.1908755 | 5.46E-05 | 0.0007774 | KLF7 | -1.192156 | 0.0037152 | 0.0440207 |
| ESYT2 | 1.0965369 | 0.0031759 | 0.0271854 | ITGAM | 1.1896782 | 3.26E-06 | 6.02E-05 | GPX4 | -1.196251 | 0.0088488 | 0.0845308 |
| CD97 | 1.0957956 | 0.000546 | 0.0064947 | APOL6 | 1.1841927 | 1.82E-05 | 0.0002918 | BCAS2 | -1.197796 | 0.0072176 | 0.0727908 |
| ZBTB7A | 1.089158 | 4.31E-05 | 0.0006927 | HSPB11 | 1.1820285 | 0.0046227 | 0.03593 | FAM78A | -1.198409 | 0.005881 | 0.0627331 |
| NFAT5 | 1.0861899 | 2.36E-05 | 0.000413 | CTSC | 1.1724201 | 0.0060399 | 0.045031 | HERC1 | -1.198577 | 0.0019085 | 0.0267612 |
| SELPLG | 1.0828019 | 0.0025323 | 0.022875 | SACM1L | 1.1710982 | 8.62E-05 | 0.0011644 | RAB4A | -1.199871 | 0.003371 | 0.0406682 |
| ELK4 | 1.0801167 | 0.0031278 | 0.0269129 | TAP1 | 1.1691074 | 0.0038589 | 0.0307944 | FDFT1 | -1.201816 | 0.0007283 | 0.0126318 |
| ARL4C | 1.064701 | 1.13E-05 | 0.0002123 | RAB27A | 1.1647354 | 0.0004269 | 0.0046929 | TAB2 | -1.207712 | 0.0006739 | 0.0118377 |
| PTPN22 | 1.0644403 | 0.0024436 | 0.0222091 | TBC1D10C | 1.158771 | 9.69E-05 | 0.0012863 | MTX3 | -1.210384 | 0.0047338 | 0.0532261 |
| CTSS | 1.0621558 | 0.001404 | 0.0140531 | DNAJC8 | 1.1564661 | 0.0006777 | 0.0070169 | HMGXB4 | -1.210427 | 0.0079828 | 0.0781645 |
| CD164 | 1.0571656 | 0.0011044 | 0.0115738 | AP4B1 | 1.1563894 | 0.0013245 | 0.01241 | JUNB | -1.213222 | 0.0001005 | 0.0023516 |
| CTSW | 1.0562127 | 0.0051209 | 0.040245 | PTK2B | 1.1516886 | 7.70E-05 | 0.0010581 | PDE7A | -1.219118 | 4.01E-05 | 0.0010584 |
| HMGN3 | 1.0519497 | 0.0034264 | 0.0290071 | RAC2 | 1.1481373 | 1.55E-06 | 3.10E-05 | NDFIP1 | -1.226453 | 1.04E-05 | 0.0003335 |
| WIPF1 | 1.0509948 | 0.0007127 | 0.0080974 | LUZP6 | 1.1413051 | 0.0002001 | 0.0024432 | EZH1 | -1.227304 | 0.0031328 | 0.0381784 |
| HCST | 1.0485969 | 0.0059501 | 0.0452617 | RAB22A | 1.1380267 | 0.0004846 | 0.005265 | NGFRAP1 | -1.234713 | 0.0077733 | 0.0767799 |
| FNDC3B | 1.0422449 | 0.0066671 | 0.0496369 | ZMAT2 | 1.1372613 | 0.0001901 | 0.0023375 | MMGT1 | -1.241557 | 0.003057 | 0.0373971 |
| PRKACB | 1.0345891 | 0.0013618 | 0.0137459 | MYO1F | 1.1266932 | 0.0015811 | 0.0144487 | MAP3K1 | -1.245871 | 4.58E-05 | 0.0011771 |
| BUB3 | 1.0326721 | 0.0016788 | 0.0163748 | MEGF9 | 1.1199093 | 0.0069844 | 0.0504717 | LRRC58 | -1.251433 | 0.0078482 | 0.0773605 |
| LUZP6 | 1.0305375 | 0.0009641 | 0.0103666 | ADCY7 | 1.1179905 | 2.00E-06 | 3.88E-05 | RGS1 | -1.251673 | 0.000862 | 0.0145303 |
| BZRAP1-AS | 1.02941 | 0.0031075 | 0.0267663 | ARF3 | 1.1108094 | 0.0057482 | 0.0431947 | AHR | -1.255931 | 0.0006121 | 0.010953 |
| MSN | 1.0268699 | 4.74E-05 | 0.0007514 | TET2 | 1.1096361 | 0.000693 | 0.0071519 | KLF10 | -1.257907 | 0.00226 | 0.0302316 |
| ADCY7 | 1.0254146 | 5.75E-05 | 0.0008884 | ADD1 | 1.1090221 | 3.80E-05 | 0.0005668 | ARL8A | -1.261988 | 0.0007297 | 0.0126331 |
| SSBP3 | 1.0129659 | 0.0078404 | 0.056518 | IL12RB1 | 1.0884687 | 0.0094754 | 0.064401 | TMBIM1 | -1.262419 | 0.0001638 | 0.0036443 |
| 7-Sep | 1.0103732 | 9.64E-06 | 0.0001839 | ARPC5 | 1.088435 | 0.0026226 | 0.0222936 | ANXA2 | -1.268093 | 0.0051968 | 0.0569631 |
| MAPK1 | 1.0054759 | 0.0002183 | 0.0028875 | CCDC88C | 1.0873564 | 0.0003149 | 0.0036033 | EDEM1 | -1.276199 | 6.25E-05 | 0.0015375 |
| SARAF | -1.006754 | 0.0010812 | 0.0113798 | ARL4C | 1.0824059 | 1.16E-06 | 2.39E-05 | PDP1 | -1.282747 | 0.0022008 | 0.029521 |
| FOXP1 | -1.008219 | 0.0004541 | 0.005503 | ZBTB7A | 1.0779216 | 2.11E-05 | 0.0003351 | KIAA1586 | -1.291668 | 0.0094386 | 0.0880599 |
| LMAN1 | -1.014616 | 0.0023696 | 0.0217249 | SPATA13 | 1.055772 | 0.0006517 | 0.0067879 | COPB2 | -1.302667 | 0.0043709 | 0.0501998 |
| KDM7A | -1.016694 | 0.0080323 | 0.0576543 | JAK1 | 1.0519595 | 0.0001531 | 0.0019304 | FAM117B | -1.30303 | 0.0068679 | 0.0702237 |
| PFDN5 | -1.028749 | 0.0022598 | 0.0208546 | TRIM22 | 1.0364391 | 0.0001782 | 0.0022108 | KIF3A | -1.303042 | 0.0066443 | 0.0687433 |
| CDC25B | -1.030875 | 0.0056399 | 0.0433209 | ZMAT3 | 1.0361242 | 0.0062927 | 0.0464841 | NDUFAF2 | -1.304151 | 0.0085968 | 0.0827832 |
| KLRB1 | -1.031422 | 0.0013767 | 0.0138708 | PPP1R18 | 1.0335151 | 5.80E-05 | 0.0008168 | TMOD3 | -1.305536 | 1.55E-05 | 0.0004656 |
| BCL7B | -1.049457 | 0.0096655 | 0.066316 | DBI | 1.0286552 | 0.0092785 | 0.063295 | BCL7B | -1.314845 | 0.0015881 | 0.0233743 |
| KLF12 | -1.050778 | 0.0005728 | 0.0067276 | IQGAP1 | 1.0275566 | 6.74E-05 | 0.0009388 | SVIL | -1.317371 | 3.10E-05 | 0.0008506 |
| BTF3 | -1.056567 | 0.0016559 | 0.0161893 | ADAM10 | 1.024903 | 0.0003799 | 0.0042571 | PCNX | -1.318107 | 4.87E-05 | 0.0012443 |
| CXCR4 | -1.060341 | 0.0067153 | 0.0498192 | SLC44A2 | 1.0218813 | 0.0040593 | 0.0321173 | SAMSN1 | -1.322711 | 5.76E-05 | 0.0014331 |
| TOMM20 | -1.062529 | 0.0029739 | 0.0258542 | ADAM8 | 1.0213621 | 0.0012591 | 0.0118331 | DNAJB9 | -1.324917 | 0.0088631 | 0.0845833 |

| | | | | | | | | | | | |
|---|---|---|---|---|---|---|---|---|---|---|---|
| C6orf48 | -1.065412 | 0.0024386 | 0.0221877 | DIDO1 | 1.0169522 | 0.0016147 | 0.0146832 | P2RY8 | -1.3254 | 0.0001798 | 0.0039371 |
| ELOVL5 | -1.073592 | 0.0031612 | 0.0271106 | RAP1B | 1.0098909 | 1.89E-06 | 3.71E-05 | FOSB | -1.330368 | 4.46E-05 | 0.0011573 |
| SNHG16 | -1.079756 | 0.0009969 | 0.0106647 | PRMT2 | 1.009565 | 0.0032855 | 0.0268914 | TP53RK | -1.333109 | 0.0085023 | 0.0821481 |
| RPL7 | -1.082189 | 9.79E-07 | 2.32E-05 | KARS | -1.001642 | 0.0043833 | 0.0342255 | CAPN1 | -1.333349 | 0.0092382 | 0.0868661 |
| HECA | -1.084006 | 0.0025495 | 0.0229726 | RUFY3 | -1.004311 | 0.0040756 | 0.0321913 | OSBPL3 | -1.333432 | 0.0009204 | 0.0152656 |
| MLXIP | -1.084149 | 0.0003837 | 0.0047457 | TIMP1 | -1.004589 | 0.001083 | 0.0104249 | PASK | -1.336382 | 0.0052129 | 0.0570745 |
| BCL11B | -1.084413 | 0.0013026 | 0.0132627 | STT3B | -1.006868 | 0.0001588 | 0.0019951 | FAM53B | -1.337374 | 0.0015233 | 0.0227219 |
| SPTBN1 | -1.085338 | 0.0001174 | 0.0016916 | BTF3 | -1.009864 | 6.87E-05 | 0.0009546 | SSFA2 | -1.344572 | 0.0090298 | 0.0854092 |
| RPLP1 | -1.086805 | 3.46E-05 | 0.000573 | RPS27 | -1.010261 | 4.29E-06 | 7.70E-05 | MEI1 | -1.346987 | 0.0020082 | 0.0276732 |
| RPL23 | -1.08766 | 1.86E-06 | 4.15E-05 | CREB1 | -1.012181 | 0.0011765 | 0.0111355 | TAF1B | -1.355286 | 0.0073619 | 0.0737036 |
| TTC39C | -1.093993 | 0.0056434 | 0.0433209 | ELP2 | -1.012395 | 0.0098593 | 0.0665997 | EIF4E3 | -1.356533 | 0.0001454 | 0.0032893 |
| SIK3 | -1.097082 | 0.0027142 | 0.0240268 | NOP58 | -1.013232 | 0.0057447 | 0.0431947 | GMPR2 | -1.358648 | 0.0040799 | 0.0477195 |
| NSA2 | -1.100019 | 0.0003988 | 0.0048968 | ERP29 | -1.018137 | 0.0012445 | 0.0117436 | TRIB2 | -1.359075 | 0.0004096 | 0.0078251 |
| STT3B | -1.107668 | 0.0001502 | 0.0020809 | MKNK2 | -1.021573 | 0.002755 | 0.0232694 | SLC2A4RG | -1.35951 | 0.00328 | 0.0397202 |
| RPL24 | -1.108948 | 0.00101 | 0.010791 | NSA2 | -1.023395 | 0.0008736 | 0.0086958 | LEPROTL1 | -1.360511 | 0.0003354 | 0.0066447 |
| EXOSC6 | -1.112853 | 0.0056011 | 0.0432736 | RPL31 | -1.024839 | 1.92E-06 | 3.75E-05 | PCYOX1L | -1.3615 | 0.0025885 | 0.0332787 |
| EIF3D | -1.11376 | 0.0013075 | 0.0132779 | TSNAX | -1.028752 | 0.0027806 | 0.0234287 | HK1 | -1.370096 | 0.0012763 | 0.0196808 |
| RPS3 | -1.119007 | 1.06E-05 | 0.0002003 | RPL7A | -1.028867 | 3.96E-05 | 0.000588 | ATP6V1B2 | -1.370526 | 0.0046684 | 0.0526144 |
| ARHGAP15 | -1.122307 | 0.0013542 | 0.0136883 | RPS11 | -1.034271 | 8.65E-07 | 1.85E-05 | PLSCR3 | -1.373658 | 3.61E-06 | 0.0001351 |
| IER2 | -1.123033 | 7.35E-05 | 0.001104 | DUT | -1.043872 | 0.0065971 | 0.0483142 | FKBP5 | -1.376649 | 0.0010262 | 0.0165618 |
| IFRD1 | -1.130009 | 0.0047061 | 0.0377474 | DNMT3A | -1.044555 | 0.0057209 | 0.043161 | SSBP3 | -1.379078 | 0.0006893 | 0.0120634 |
| RPS4X | -1.131156 | 0.0002625 | 0.0033917 | CD48 | -1.046435 | 8.11E-06 | 0.0001387 | ANKRD28 | -1.380716 | 1.32E-05 | 0.0004017 |
| RPL37 | -1.132482 | 1.14E-05 | 0.000214 | PPP1CB | -1.048262 | 2.36E-05 | 0.0003677 | CNST | -1.381372 | 0.0001184 | 0.0027357 |
| EDEM1 | -1.134759 | 0.0018339 | 0.0175015 | HNRNPA0 | -1.048699 | 0.000139 | 0.0017776 | TNFAIP8 | -1.381502 | 0.001297 | 0.0199038 |
| RPL7A | -1.134947 | 4.64E-06 | 9.38E-05 | SATB1 | -1.052724 | 2.32E-05 | 0.000362 | KDM6B | -1.383004 | 0.0079782 | 0.0781645 |
| BTF3L4 | -1.13627 | 0.0045676 | 0.0368478 | RPL27A | -1.053067 | 1.81E-07 | 4.37E-06 | ZMPSTE24 | -1.384483 | 0.0028405 | 0.0354466 |
| ZBTB20 | -1.139123 | 2.08E-05 | 0.0003686 | CCT2 | -1.057076 | 0.0004038 | 0.0044822 | SMAD3 | -1.392177 | 0.00897 | 0.0851793 |
| IRF2BP2 | -1.140595 | 2.86E-06 | 6.05E-05 | MIF | -1.059432 | 0.0029334 | 0.0243755 | UXS1 | -1.406801 | 0.0030234 | 0.0371156 |
| ZFP36L1 | -1.14661 | 0.0004044 | 0.0049516 | TFG | -1.061408 | 0.0074562 | 0.0528902 | P2RY10 | -1.40971 | 0.0071174 | 0.0719404 |
| ABCA5 | -1.152359 | 0.009786 | 0.0670334 | TYW3 | -1.061889 | 0.0060289 | 0.0450075 | BTG2 | -1.412223 | 0.0015918 | 0.02338 |
| RSL1D1 | -1.153561 | 0.0003058 | 0.0038623 | SPOCK2 | -1.067198 | 2.49E-05 | 0.0003861 | EI24 | -1.412566 | 0.0004608 | 0.0085982 |
| S100A10 | -1.160513 | 0.0018285 | 0.0174765 | NSMCE1 | -1.080691 | 0.0097434 | 0.0658837 | SOD2 | -1.412864 | 0.0072369 | 0.0729085 |
| RPL27 | -1.168875 | 2.86E-06 | 6.05E-05 | HSD17B11 | -1.08303 | 0.0057805 | 0.0433981 | MAPK8 | -1.414528 | 6.13E-05 | 0.0015115 |
| EEF2 | -1.16959 | 7.21E-06 | 0.0001415 | GSPT1 | -1.083839 | 0.0025987 | 0.0221311 | STAT5B | -1.416004 | 0.0010556 | 0.0167947 |
| RPL31 | -1.172103 | 1.07E-07 | 2.93E-06 | ASAP1 | -1.084129 | 0.000362 | 0.0040913 | ERCC6L2 | -1.416681 | 0.0049731 | 0.0553977 |
| PDCD4 | -1.173584 | 3.29E-05 | 0.0005458 | STX17 | -1.08581 | 0.0003282 | 0.003741 | ATP6V0A2 | -1.426871 | 0.007818 | 0.0771425 |
| NPM1 | -1.176336 | 1.72E-06 | 3.89E-05 | TMEM167A | -1.088576 | 0.00388 | 0.0308828 | SLC30A5 | -1.429261 | 0.0089089 | 0.0849168 |
| EDEM3 | -1.177758 | 0.0092578 | 0.06394 | JAK3 | -1.094804 | 0.0095658 | 0.0648723 | ERO1L | -1.430168 | 1.05E-05 | 0.0003335 |
| KLHL5 | -1.179608 | 0.0038593 | 0.0320577 | RPL24 | -1.095232 | 0.0006208 | 0.0065377 | RNASET2 | -1.430532 | 0.0004856 | 0.0090087 |
| ITGB1 | -1.179664 | 6.77E-05 | 0.001028 | TAGLN2 | -1.096276 | 8.92E-05 | 0.0012021 | GATA3 | -1.432397 | 1.92E-06 | 7.54E-05 |
| CD52 | -1.183043 | 0.0006536 | 0.0075291 | RPL29 | -1.097342 | 3.31E-06 | 6.09E-05 | SOX4 | -1.433768 | 1.32E-05 | 0.0004025 |
| RPS11 | -1.187563 | 1.91E-08 | 5.97E-07 | CD52 | -1.098265 | 0.0026168 | 0.0222641 | TANC2 | -1.436319 | 0.0048169 | 0.0540343 |
| EIF3E | -1.191272 | 6.96E-05 | 0.0010539 | TC2N | -1.100901 | 0.0064027 | 0.0471514 | ATP2A3 | -1.438158 | 0.0026509 | 0.0338092 |
| NUDT4 | -1.192538 | 2.30E-05 | 0.0004049 | ZBTB10 | -1.101859 | 0.0007712 | 0.0078107 | CXCR4 | -1.440726 | 7.09E-05 | 0.0017259 |
| SLC25A6 | -1.192664 | 4.76E-06 | 9.60E-05 | MALT1 | -1.102216 | 0.0007216 | 0.0073896 | CERK | -1.44953 | 0.0003083 | 0.0061998 |
| ANXA7 | -1.194724 | 0.0090482 | 0.0629554 | ASXL1 | -1.103911 | 0.0006407 | 0.0067248 | CORO2A | -1.455437 | 0.0035261 | 0.0422738 |
| SELL | -1.204362 | 0.0001053 | 0.001536 | RPL37A | -1.104008 | 1.04E-06 | 2.17E-05 | IRF2BP2 | -1.457859 | 9.73E-09 | 6.01E-07 |
| LZTFL1 | -1.20927 | 0.0063004 | 0.047496 | HNRNPA1 | -1.105152 | 9.88E-08 | 2.56E-06 | PIM2 | -1.458035 | 0.0025818 | 0.033237 |
| HSP90AB1 | -1.209373 | 1.75E-06 | 3.92E-05 | TNFAIP3 | -1.105294 | 0.0069554 | 0.0503854 | CISD2 | -1.463059 | 0.0009216 | 0.0152656 |
| RPL15 | -1.213597 | 2.69E-08 | 8.20E-07 | UXT | -1.107245 | 0.0077584 | 0.0546632 | MCMBP | -1.464972 | 0.0024984 | 0.0325131 |
| SNRPN | -1.21966 | 0.0014377 | 0.014339 | PLSCR3 | -1.112125 | 0.0002751 | 0.0032107 | NUDT4 | -1.465554 | 1.13E-06 | 4.60E-05 |
| GNB2L1 | -1.227133 | 7.27E-06 | 0.0001419 | GLTSCR2 | -1.115443 | 8.51E-07 | 1.84E-05 | IRS2 | -1.467002 | 8.42E-05 | 0.0020093 |
| RPS15A | -1.22904 | 1.17E-07 | 3.15E-06 | NME2 | -1.117888 | 4.47E-05 | 0.000656 | RABL2B | -1.472711 | 0.0058867 | 0.0627331 |
| MBNL2 | -1.23501 | 0.008996 | 0.0627072 | KRTCAP2 | -1.118501 | 0.0005513 | 0.0058859 | ZBED5 | -1.477024 | 0.0018668 | 0.026447 |

| | | | | | | | | | | | |
|---|---|---|---|---|---|---|---|---|---|---|---|
| PHACTR2 | -1.235994 | 3.09E-05 | 0.0005181 | NOL7 | -1.11903 | 0.004462 | 0.0347975 | TGFBR2 | -1.480644 | 0.0073001 | 0.0733411 |
| CYLD | -1.238871 | 0.0002756 | 0.0035341 | EZR | -1.122118 | 9.10E-05 | 0.0012231 | ZXDB | -1.484888 | 0.0024259 | 0.0319603 |
| RPL26 | -1.24234 | 1.82E-08 | 5.76E-07 | RPL41 | -1.131713 | 2.99E-06 | 5.54E-05 | FAM222B | -1.485714 | 0.0007885 | 0.0134324 |
| SNHG6 | -1.249721 | 2.84E-05 | 0.0004851 | RSL1D1 | -1.132569 | 0.0003971 | 0.0044183 | PRKCQ-AS | -1.486336 | 0.0017358 | 0.025035 |
| HSD17B11 | -1.249977 | 0.0030598 | 0.0264912 | NDRG1 | -1.134045 | 0.0061457 | 0.0456952 | DAG1 | -1.49228 | 0.0080573 | 0.0786815 |
| NME2 | -1.25069 | 9.82E-05 | 0.0014413 | HIVEP1 | -1.136406 | 0.0027815 | 0.0234287 | ITGB1 | -1.493158 | 3.01E-07 | 1.37E-05 |
| CD48 | -1.252968 | 5.85E-07 | 1.43E-05 | LZTFL1 | -1.138404 | 0.004974 | 0.0383705 | PIK3IP1 | -1.494976 | 4.22E-05 | 0.001105 |
| TAGLN2 | -1.253606 | 4.91E-05 | 0.0007735 | RPL35A | -1.140994 | 1.60E-07 | 3.93E-06 | USP31 | -1.51031 | 0.0083872 | 0.0812539 |
| RPL35 | -1.253728 | 1.27E-06 | 2.92E-05 | SLC30A7 | -1.14139 | 0.0028586 | 0.0239672 | ETHE1 | -1.517605 | 0.0021568 | 0.0290536 |
| ST3GAL1 | -1.258032 | 0.0001576 | 0.0021691 | CD69 | -1.14321 | 2.95E-06 | 5.48E-05 | RAB3GAP1 | -1.522912 | 0.004486 | 0.0510994 |
| RPL3 | -1.259618 | 1.86E-07 | 4.92E-06 | RPL17 | -1.144995 | 4.15E-07 | 9.53E-06 | TRAF3IP2 | -1.527917 | 0.0018869 | 0.0265823 |
| RPS29 | -1.265307 | 3.94E-09 | 1.43E-07 | ST3GAL1 | -1.146041 | 0.0013199 | 0.0123792 | SNX30 | -1.531021 | 0.007441 | 0.0744174 |
| ANKRD28 | -1.270903 | 4.90E-05 | 0.0007735 | EIF3D | -1.14722 | 5.34E-05 | 0.0007698 | ICAM2 | -1.535098 | 9.97E-05 | 0.0023389 |
| RCSD1 | -1.27438 | 0.0063417 | 0.0477219 | RPS19 | -1.149707 | 7.88E-07 | 1.72E-05 | HECA | -1.540525 | 5.17E-10 | 4.39E-08 |
| TOMM7 | -1.280639 | 2.76E-06 | 5.86E-05 | RPS4X | -1.155095 | 1.04E-05 | 0.000174 | APOOL | -1.540612 | 0.0051604 | 0.056693 |
| RPS20 | -1.287643 | 4.73E-09 | 1.68E-07 | RPS24 | -1.157492 | 1.31E-07 | 3.29E-06 | GRPEL2 | -1.545085 | 0.001897 | 0.0266783 |
| RPL6 | -1.290169 | 3.31E-09 | 1.21E-07 | SESN1 | -1.161065 | 0.0005434 | 0.0058155 | HERC3 | -1.550363 | 0.0069975 | 0.0713214 |
| STK17B | -1.291405 | 2.22E-07 | 5.78E-06 | PDE7A | -1.16172 | 0.0002248 | 0.0027029 | SEC11C | -1.551565 | 3.10E-05 | 0.0008506 |
| ANKRD26 | -1.291512 | 0.0003772 | 0.0046797 | EDEM1 | -1.165741 | 0.0009572 | 0.0093963 | SC5D | -1.557631 | 0.0001377 | 0.0031454 |
| KIF3A | -1.293742 | 0.00357 | 0.0301313 | BCL11B | -1.166637 | 0.0002765 | 0.0032199 | RPS6KA3 | -1.559151 | 7.49E-09 | 4.76E-07 |
| ITGA6 | -1.294514 | 0.0012972 | 0.0132532 | PABPC1 | -1.168631 | 8.16E-05 | 0.0011144 | SPINT2 | -1.568728 | 0.0050235 | 0.0558292 |
| IL17RA | -1.295456 | 0.0053029 | 0.0414269 | GNB2L1 | -1.180974 | 8.21E-07 | 1.78E-05 | CDR2 | -1.573623 | 2.87E-05 | 0.0007966 |
| OCIAD2 | -1.295994 | 0.0001284 | 0.0018367 | ZFAS1 | -1.181567 | 0.000449 | 0.0049193 | SLC25A33 | -1.574714 | 0.0032303 | 0.0392176 |
| UBE2I | -1.297222 | 5.56E-05 | 0.0008628 | SNRPE | -1.182066 | 0.0014649 | 0.0135334 | ARHGAP5 | -1.577493 | 1.02E-05 | 0.0003285 |
| RPL30 | -1.297996 | 2.09E-07 | 5.47E-06 | RPL15 | -1.187379 | 2.25E-09 | 7.84E-08 | S100A11 | -1.577693 | 0.0002288 | 0.0048234 |
| RPL27A | -1.300788 | 1.15E-08 | 3.80E-07 | PRPSAP2 | -1.1909 | 0.0024862 | 0.0212905 | RCAN3 | -1.578327 | 0.001978 | 0.0274497 |
| RPS6KA3 | -1.30771 | 4.19E-06 | 8.54E-05 | HMGCS1 | -1.192417 | 0.007084 | 0.0510195 | CDKN1B | -1.580148 | 5.59E-07 | 2.41E-05 |
| CHD7 | -1.308147 | 0.0005691 | 0.006703 | KLHL5 | -1.193936 | 0.0013977 | 0.0130294 | IBA57 | -1.582159 | 0.0056703 | 0.0610369 |
| SNHG1 | -1.308711 | 0.00081 | 0.0089858 | RPL3 | -1.194132 | 1.15E-08 | 3.48E-07 | INTS2 | -1.585088 | 0.0087227 | 0.083492 |
| RPS7 | -1.318663 | 1.29E-10 | 6.29E-09 | DAAM1 | -1.194165 | 0.0008794 | 0.0087436 | PPT1 | -1.58714 | 0.000173 | 0.0038054 |
| S100A6 | -1.320312 | 1.33E-05 | 0.0002455 | PAFAH1B2 | -1.197449 | 0.0014305 | 0.0132555 | MAP4K2 | -1.588797 | 0.0091862 | 0.0864619 |
| VWA8 | -1.320485 | 0.006216 | 0.0469866 | OCIAD2 | -1.197822 | 0.0020218 | 0.017825 | CYSLTR1 | -1.593367 | 1.19E-05 | 0.0003697 |
| RPL41 | -1.32393 | 2.47E-07 | 6.36E-06 | TMEM33 | -1.198437 | 0.0057347 | 0.0431947 | NDRG1 | -1.594363 | 0.0027524 | 0.0346885 |
| TAB2 | -1.324631 | 0.0003153 | 0.0039699 | RPS15A | -1.204096 | 1.91E-07 | 4.60E-06 | SPOCK2 | -1.594627 | 3.16E-10 | 2.83E-08 |
| DDX21 | -1.324871 | 1.99E-06 | 4.38E-05 | ABLIM1 | -1.204653 | 0.0002385 | 0.0028525 | INPP4B | -1.595564 | 3.28E-05 | 0.0008918 |
| TFRC | -1.333042 | 0.0026575 | 0.0236919 | DDX21 | -1.207432 | 1.39E-06 | 2.82E-05 | LSR | -1.596217 | 0.0053685 | 0.0585108 |
| RNASET2 | -1.336591 | 0.0010837 | 0.0113798 | DPYD | -1.21267 | 0.0073181 | 0.0521521 | SPECC1L | -1.59741 | 0.0092485 | 0.0868775 |
| ZFP36L2 | -1.344025 | 2.56E-06 | 5.49E-05 | NUP54 | -1.217104 | 0.0030848 | 0.025451 | SNX2 | -1.597899 | 0.000247 | 0.0051267 |
| RPS24 | -1.35065 | 4.63E-09 | 1.65E-07 | TOMM7 | -1.217228 | 1.08E-05 | 0.0001795 | PDLIM5 | -1.600821 | 0.0010146 | 0.0164098 |
| MAP3K1 | -1.351593 | 2.21E-05 | 0.0003897 | ZFP36L1 | -1.220031 | 4.44E-06 | 7.92E-05 | TRADD | -1.602357 | 0.0018875 | 0.0265823 |
| RPL29 | -1.353711 | 1.21E-06 | 2.81E-05 | RPL28 | -1.225768 | 5.37E-07 | 1.20E-05 | DCK | -1.606293 | 6.67E-05 | 0.0016319 |
| STARD4 | -1.359905 | 0.0001658 | 0.0022678 | RPL6 | -1.228361 | 5.20E-09 | 1.69E-07 | CCND2 | -1.607346 | 1.36E-08 | 8.00E-07 |
| RPSAP58 | -1.359951 | 0.0036835 | 0.0309342 | RALA | -1.233225 | 0.0060417 | 0.045031 | RPS26P11 | -1.613282 | 0.0042557 | 0.0490587 |
| RPS27 | -1.361584 | 6.38E-07 | 1.55E-05 | RPL14 | -1.238328 | 1.05E-08 | 3.26E-07 | ZYG11B | -1.613764 | 0.0002571 | 0.0053264 |
| NFIC | -1.363091 | 0.0007812 | 0.0087127 | RPS20 | -1.245501 | 2.49E-10 | 1.01E-08 | NDST2 | -1.617409 | 0.0005506 | 0.010039 |
| WWP1 | -1.365425 | 0.0001879 | 0.0025323 | LSM5 | -1.245838 | 0.0086027 | 0.0598309 | LDHB | -1.630167 | 1.78E-10 | 1.63E-08 |
| EEF1A1 | -1.365715 | 1.71E-10 | 7.98E-09 | RPL36A | -1.246691 | 1.43E-08 | 4.23E-07 | MBNL2 | -1.63145 | 3.99E-05 | 0.0010565 |
| SNHG15 | -1.371715 | 0.0019055 | 0.0180408 | STK17B | -1.247905 | 1.17E-06 | 2.41E-05 | CCSAP | -1.633966 | 9.80E-05 | 0.0023096 |
| RPL19 | -1.372067 | 1.71E-10 | 7.98E-09 | H2AFY | -1.249122 | 1.14E-06 | 2.36E-05 | ZBTB21 | -1.636424 | 0.0059149 | 0.0629185 |
| HERC5 | -1.37286 | 0.0070088 | 0.0518132 | RPS7 | -1.251111 | 6.32E-09 | 2.02E-07 | YPEL2 | -1.636663 | 5.78E-06 | 0.0002001 |
| AIM1 | -1.373861 | 0.000653 | 0.0075291 | CXCR4 | -1.255458 | 0.0014188 | 0.013173 | IDE | -1.639934 | 0.002817 | 0.0353015 |
| MAF | -1.374627 | 0.0019332 | 0.0182414 | DDT | -1.256303 | 0.0015895 | 0.0144823 | IL7R | -1.643672 | 6.25E-07 | 2.69E-05 |
| LMO4 | -1.376409 | 6.67E-05 | 0.0010154 | CRYL1 | -1.256945 | 0.0052176 | 0.0399263 | TOB1 | -1.64588 | 4.74E-06 | 0.0001706 |
| ESD | -1.381414 | 0.0037912 | 0.03168 | RPS27A | -1.257346 | 6.38E-08 | 1.70E-06 | TIMP1 | -1.649982 | 5.74E-08 | 2.97E-06 |

| | | | | | | | | | | | |
|---|---|---|---|---|---|---|---|---|---|---|---|
| TXNDC12 | -1.382226 | 0.0060634 | 0.0460177 | RPL10 | -1.259821 | 5.70E-09 | 1.83E-07 | VIM | -1.650761 | 1.48E-10 | 1.38E-08 |
| RPL28 | -1.38669 | 1.21E-08 | 3.94E-07 | EIF3H | -1.265489 | 0.0001248 | 0.0016091 | KCNA3 | -1.657036 | 7.74E-05 | 0.0018598 |
| JUNB | -1.389141 | 1.98E-05 | 0.0003535 | NRIP1 | -1.266096 | 9.73E-05 | 0.0012909 | ZNF445 | -1.673414 | 0.0086375 | 0.0828424 |
| RPS27A | -1.389236 | 5.87E-08 | 1.69E-06 | RPS25 | -1.266707 | 2.14E-09 | 7.53E-08 | RGCC | -1.674352 | 0.0015758 | 0.0232627 |
| MAPK6 | -1.389433 | 0.0085265 | 0.0603251 | CBWD5 | -1.267196 | 0.0057179 | 0.043161 | LEF1 | -1.677836 | 4.51E-06 | 0.0001646 |
| ERCC6L2 | -1.392724 | 0.0046325 | 0.0372937 | SIK1 | -1.267246 | 3.81E-05 | 0.0005676 | RUNX1 | -1.680169 | 0.0002111 | 0.0044994 |
| RPL14 | -1.39727 | 2.30E-09 | 8.64E-08 | CDKN1B | -1.268199 | 2.19E-06 | 4.18E-05 | CCDC167 | -1.685812 | 0.0051346 | 0.0566064 |
| RPL37A | -1.398319 | 3.96E-08 | 1.17E-06 | RPL18 | -1.268792 | 1.63E-07 | 3.96E-06 | TRIM4 | -1.692181 | 0.0044393 | 0.0508085 |
| RPSA | -1.401325 | 1.09E-07 | 2.98E-06 | CDK6 | -1.268834 | 0.000301 | 0.0034645 | RARG | -1.727848 | 0.0028171 | 0.0353015 |
| RPS14 | -1.402112 | 7.01E-08 | 1.98E-06 | ELOVL5 | -1.274259 | 6.73E-05 | 0.0009383 | SZT2 | -1.742996 | 0.0098855 | 0.0914293 |
| RPL10 | -1.40513 | 5.10E-10 | 2.18E-08 | NDUFV2 | -1.275131 | 0.0009309 | 0.0091581 | ASNS | -1.744172 | 0.0011835 | 0.0184869 |
| SNHG5 | -1.405842 | 0.0006886 | 0.0078574 | SEC11C | -1.275489 | 2.10E-05 | 0.0003342 | CRYBG3 | -1.746378 | 0.0026135 | 0.0335108 |
| CLUHP3 | -1.407989 | 0.007929 | 0.0571077 | CD59 | -1.27606 | 0.0011315 | 0.0107985 | THAP5 | -1.748772 | 3.77E-05 | 0.0010101 |
| NAP1L1 | -1.415658 | 2.43E-07 | 6.27E-06 | RPLP2 | -1.27649 | 1.12E-09 | 4.08E-08 | SYNJ2 | -1.751617 | 4.04E-06 | 0.0001495 |
| ZMIZ1 | -1.419591 | 0.0003873 | 0.0047759 | TLK1 | -1.279208 | 4.54E-05 | 0.0006644 | PHLDA1 | -1.752949 | 0.0094261 | 0.0880288 |
| RPL36A | -1.424886 | 3.15E-09 | 1.16E-07 | PAPSS1 | -1.279956 | 0.0071999 | 0.051626 | EDEM3 | -1.7606 | 2.56E-08 | 1.43E-06 |
| RPL5 | -1.427727 | 2.68E-11 | 1.48E-09 | NPM1 | -1.283031 | 8.31E-07 | 1.80E-05 | SNX9 | -1.773005 | 0.0011756 | 0.0183938 |
| YPEL2 | -1.430224 | 0.0001478 | 0.0020614 | IRF2BPL | -1.283657 | 0.0029547 | 0.0245117 | LINC0015 | -1.775515 | 0.000656 | 0.0115664 |
| RPS14P3 | -1.430783 | 0.002684 | 0.0238265 | MTA1 | -1.285217 | 0.0069561 | 0.0503854 | ELOVL5 | -1.780574 | 3.94E-05 | 0.0010468 |
| RPS23 | -1.43222 | 9.00E-10 | 3.65E-08 | ITGA6 | -1.285274 | 0.0028796 | 0.0241013 | OSGEP | -1.784115 | 0.0092979 | 0.0870828 |
| RPL38 | -1.434048 | 2.68E-08 | 8.20E-07 | RPL34 | -1.285684 | 1.74E-07 | 4.22E-06 | CREBL2 | -1.785308 | 4.41E-05 | 0.0011487 |
| RPL39 | -1.435284 | 2.42E-08 | 7.44E-07 | EIF3L | -1.289395 | 8.61E-07 | 1.85E-05 | TFRC | -1.796473 | 0.0003757 | 0.0072941 |
| FOXJ2 | -1.436444 | 0.0036178 | 0.030504 | FAM60A | -1.290655 | 4.18E-05 | 0.0006149 | TP53INP1 | -1.804853 | 4.15E-09 | 2.71E-07 |
| BEX2 | -1.43704 | 0.0032551 | 0.0277806 | NSUN6 | -1.292774 | 0.0054398 | 0.0413435 | EVI2B | -1.807635 | 6.46E-07 | 2.77E-05 |
| TMEM14B | -1.441615 | 0.0023861 | 0.0218285 | RPL38 | -1.292945 | 1.05E-07 | 2.72E-06 | SLC2A3 | -1.817055 | 5.25E-07 | 2.29E-05 |
| LRRC75A-A | -1.443172 | 3.19E-06 | 6.61E-05 | SARAF | -1.295021 | 3.30E-06 | 6.08E-05 | AMMECR1 | -1.817691 | 8.21E-06 | 0.0002723 |
| RPL23A | -1.446729 | 3.72E-08 | 1.11E-06 | RPS14 | -1.296324 | 2.08E-09 | 7.35E-08 | PIEZO1 | -1.828476 | 4.89E-05 | 0.0012443 |
| PAICS | -1.447857 | 0.0031789 | 0.0271854 | AIMP1 | -1.297751 | 0.0007859 | 0.0079509 | NAP1L2 | -1.82885 | 0.0023845 | 0.0315885 |
| STYX | -1.449625 | 0.0028986 | 0.0253566 | PTGER4 | -1.299359 | 0.0023655 | 0.0203885 | NAMPT | -1.831859 | 7.81E-06 | 0.0002613 |
| RPL22 | -1.456854 | 2.77E-10 | 1.23E-08 | EPM2AIP1 | -1.300657 | 0.0004876 | 0.0052856 | RNF144A | -1.832284 | 0.000394 | 0.0075735 |
| NFKBIZ | -1.462295 | 2.67E-05 | 0.0004608 | CCSAP | -1.305571 | 0.0012356 | 0.0116712 | FAM102A | -1.834473 | 9.78E-06 | 0.0003169 |
| STAT5A | -1.46315 | 0.000383 | 0.0047437 | KAT2A | -1.307676 | 0.0088194 | 0.0609718 | IL6ST | -1.834974 | 4.52E-05 | 0.0011649 |
| RPS28 | -1.464268 | 5.89E-08 | 1.69E-06 | ZMAT1 | -1.308635 | 0.0003073 | 0.0035203 | TMEM167 | -1.840311 | 7.82E-06 | 0.0002613 |
| CDKN1B | -1.465412 | 2.86E-05 | 0.0004865 | TTC39C | -1.310358 | 0.0001524 | 0.0019251 | PCBD2 | -1.851052 | 0.0082267 | 0.080111 |
| STK11 | -1.467374 | 0.0098536 | 0.067353 | RPS16 | -1.312501 | 4.53E-11 | 2.16E-09 | FOCAD | -1.857634 | 0.0076264 | 0.0759553 |
| LPHN1 | -1.473701 | 0.0048183 | 0.0383766 | RPSA | -1.314009 | 1.08E-07 | 2.75E-06 | KLRB1 | -1.867241 | 2.10E-09 | 1.48E-07 |
| RPL35A | -1.474905 | 4.28E-11 | 2.23E-09 | RPS17 | -1.314223 | 8.25E-09 | 2.57E-07 | ZNF516 | -1.868575 | 0.0007033 | 0.0122644 |
| RPL34 | -1.474946 | 8.43E-09 | 2.89E-07 | RPL27 | -1.314626 | 1.31E-08 | 3.91E-07 | NBEA | -1.868829 | 0.0021249 | 0.0287455 |
| 11-Sep | -1.475219 | 1.77E-06 | 3.96E-05 | DARS | -1.318127 | 0.000267 | 0.0031253 | GLIPR1 | -1.879291 | 2.50E-07 | 1.15E-05 |
| SLC7A6 | -1.478795 | 0.0012086 | 0.0124539 | S100A10 | -1.318214 | 6.92E-06 | 0.0001195 | PTGER2 | -1.883215 | 3.76E-05 | 0.0010091 |
| PABPC4 | -1.480841 | 0.0010264 | 0.0108984 | DUSP10 | -1.319036 | 0.0072074 | 0.0516397 | ZNF101 | -1.8854 | 0.0004597 | 0.0085953 |
| C10orf128 | -1.48104 | 0.0070873 | 0.0520276 | NABP1 | -1.322111 | 0.0066086 | 0.0483606 | MSRB3 | -1.887764 | 0.0019939 | 0.0275245 |
| PEBP1 | -1.481523 | 0.0010605 | 0.0112044 | EEF1A1 | -1.327826 | 4.71E-12 | 2.60E-10 | VCL | -1.889969 | 1.53E-09 | 1.13E-07 |
| EIF3L | -1.481957 | 5.09E-07 | 1.26E-05 | EEF2 | -1.331731 | 4.08E-08 | 1.13E-06 | KDM7A | -1.89879 | 2.15E-06 | 8.33E-05 |
| LTA4H | -1.487236 | 0.0004172 | 0.0050773 | SLC25A6 | -1.334149 | 5.01E-08 | 1.36E-06 | DGKA | -1.902737 | 1.75E-09 | 1.25E-07 |
| RPL17 | -1.489791 | 1.92E-09 | 7.31E-08 | SETD1B | -1.3348 | 0.0003573 | 0.0040486 | RHOBTB3 | -1.910211 | 0.0089235 | 0.0849168 |
| RPL12 | -1.491869 | 1.00E-09 | 4.02E-08 | RPL37 | -1.337282 | 1.63E-08 | 4.74E-07 | PLIN2 | -1.927229 | 1.04E-05 | 0.0003335 |
| FAM65A | -1.492894 | 0.0088764 | 0.0621718 | RPS18 | -1.342301 | 2.30E-09 | 8.01E-08 | FAM134B | -1.930163 | 0.0013748 | 0.0208629 |
| RPS6 | -1.494064 | 3.27E-11 | 1.76E-09 | TSPYL1 | -1.34272 | 2.93E-05 | 0.0004459 | NR3C2 | -1.936645 | 0.0089246 | 0.0849168 |
| GPX4 | -1.49529 | 0.0043278 | 0.0351839 | RPL13A | -1.34374 | 3.32E-11 | 1.62E-09 | KCNMB3 | -1.945118 | 0.0069026 | 0.0705041 |
| CCNG2 | -1.495349 | 3.03E-05 | 0.0005098 | ARHGAP15 | -1.347411 | 1.57E-05 | 0.0002541 | PLP2 | -1.947085 | 1.06E-07 | 5.20E-06 |
| KLF6 | -1.498872 | 4.01E-09 | 1.45E-07 | TTN | -1.351821 | 9.08E-05 | 0.0012223 | RASA4 | -1.948297 | 1.64E-05 | 0.0004871 |
| RPS3A | -1.501647 | 4.16E-13 | 3.23E-11 | RPL23 | -1.35222 | 7.36E-11 | 3.37E-09 | CRYL1 | -1.951907 | 0.0004104 | 0.0078253 |
| RPS9 | -1.503575 | 1.76E-10 | 8.03E-09 | RPL23A | -1.353009 | 6.03E-08 | 1.62E-06 | PLXDC2 | -1.961793 | 3.78E-06 | 0.0001412 |

| | | | | | | | | | | | |
|---|---|---|---|---|---|---|---|---|---|---|---|
| NRIP1 | -1.504506 | 9.95E-05 | 0.0014572 | RPS9 | -1.353236 | 2.95E-10 | 1.18E-08 | GRASP | -1.964475 | 0.0026361 | 0.0336654 |
| ACTN1 | -1.505304 | 0.0042255 | 0.0344525 | CCDC88A | -1.355739 | 0.0001359 | 0.0017454 | RORA | -1.965079 | 1.02E-10 | 9.70E-09 |
| CCNJ | -1.507646 | 0.0075824 | 0.0549411 | PSMD12 | -1.356944 | 0.0055461 | 0.0420474 | SOCS3 | -1.982613 | 1.14E-05 | 0.0003572 |
| ZFHX3 | -1.511123 | 0.0072595 | 0.0529121 | ZFP36L2 | -1.361485 | 1.57E-06 | 3.12E-05 | MGAT5 | -1.987921 | 2.11E-09 | 1.48E-07 |
| GNAQ | -1.515048 | 1.64E-08 | 5.24E-07 | RPSAP58 | -1.361709 | 0.0005089 | 0.0054838 | SLC35A2 | -1.99368 | 0.0015263 | 0.0227311 |
| TC2N | -1.517329 | 0.0014121 | 0.0141177 | PDE4B | -1.363209 | 0.0094987 | 0.0645117 | MDFIC | -1.999266 | 1.44E-14 | 2.35E-12 |
| RPL10A | -1.519904 | 8.45E-08 | 2.33E-06 | TMEM147 | -1.364276 | 0.0036431 | 0.0294281 | DPP4 | -1.999366 | 3.97E-07 | 1.77E-05 |
| SPOCK2 | -1.527498 | 5.79E-09 | 2.05E-07 | C16orf54 | -1.366866 | 5.14E-05 | 0.0007459 | PRNP | -2.021913 | 4.61E-13 | 5.98E-11 |
| RPL4 | -1.528772 | 3.76E-12 | 2.41E-10 | IFNAR1 | -1.368217 | 0.0034347 | 0.0279886 | RAPGEF2 | -2.022557 | 5.74E-10 | 4.75E-08 |
| HINT1 | -1.531301 | 1.81E-10 | 8.20E-09 | RPL19 | -1.368381 | 1.95E-10 | 8.15E-09 | REEP3 | -2.027539 | 3.98E-06 | 0.0001481 |
| CCDC88A | -1.537053 | 3.91E-05 | 0.0006385 | RPS21 | -1.369095 | 3.38E-09 | 1.15E-07 | HMGN5 | -2.034828 | 0.0023688 | 0.0314671 |
| TSPYL4 | -1.542605 | 0.000452 | 0.0054855 | PDCD4 | -1.370257 | 3.14E-08 | 8.80E-07 | ETS2 | -2.038303 | 0.0029636 | 0.0366283 |
| RPL13A | -1.542922 | 3.83E-13 | 3.01E-11 | PTP4A1 | -1.37144 | 2.21E-06 | 4.22E-05 | S100A4 | -2.040842 | 8.06E-07 | 3.38E-05 |
| ARHGAP21 | -1.544949 | 0.0025279 | 0.022875 | TXNDC15 | -1.371606 | 0.0033187 | 0.0271153 | A2M | -2.046289 | 7.03E-05 | 0.0017162 |
| SEL1L3 | -1.556346 | 0.0009468 | 0.0102331 | ATIC | -1.371836 | 0.0018229 | 0.0162566 | STYX | -2.053157 | 0.0002781 | 0.005674 |
| RPL21 | -1.558079 | 1.13E-13 | 9.62E-12 | GADD45A | -1.373789 | 0.0001082 | 0.001421 | SDCBP | -2.058798 | 3.85E-05 | 0.0010294 |
| ITM2A | -1.558467 | 0.007402 | 0.0538632 | HSF1 | -1.37381 | 0.0007113 | 0.0073004 | LOC10272 | -2.061032 | 0.002417 | 0.0319299 |
| RPS16 | -1.560235 | 8.61E-13 | 6.29E-11 | USP36 | -1.386624 | 0.0025171 | 0.0214956 | ARHGEF18 | -2.065714 | 1.14E-07 | 5.56E-06 |
| RPS17 | -1.561946 | 5.28E-10 | 2.25E-08 | CYLD | -1.390347 | 2.56E-06 | 4.81E-05 | GLRX | -2.066035 | 1.55E-07 | 7.37E-06 |
| SKI | -1.566129 | 3.80E-05 | 0.0006237 | WDR75 | -1.391038 | 0.0038885 | 0.0308973 | SOS1 | -2.068851 | 9.89E-09 | 6.04E-07 |
| TSC22D3 | -1.566318 | 1.08E-08 | 3.62E-07 | SKI | -1.391374 | 8.47E-06 | 0.0001444 | PCSK5 | -2.069043 | 0.0041089 | 0.0479425 |
| SPG20 | -1.572749 | 0.0012803 | 0.0131125 | NXT1 | -1.393635 | 0.0024277 | 0.020847 | PRKCA | -2.069196 | 0.0010966 | 0.017327 |
| EEF1B2 | -1.574704 | 3.13E-10 | 1.37E-08 | KLRB1 | -1.39445 | 1.07E-06 | 2.23E-05 | SLC6A6 | -2.114258 | 0.0002339 | 0.0048759 |
| MIEF1 | -1.574771 | 5.51E-05 | 0.0008592 | RNASET2 | -1.395492 | 0.0004555 | 0.0049728 | NR4A1 | -2.138966 | 0.0018434 | 0.0261541 |
| RPS18 | -1.581874 | 1.95E-11 | 1.12E-09 | IER2 | -1.397897 | 4.35E-09 | 1.44E-07 | MAL | -2.15088 | 0.0018054 | 0.0258054 |
| RPL18 | -1.582369 | 8.87E-09 | 3.02E-07 | PPP1R2 | -1.402717 | 3.69E-06 | 6.70E-05 | ITGA6 | -2.163371 | 1.59E-05 | 0.0004765 |
| EEF1G | -1.583539 | 2.50E-11 | 1.41E-09 | RPL12 | -1.40305 | 4.64E-10 | 1.83E-08 | ATF7IP2 | -2.163387 | 4.48E-05 | 0.0011573 |
| RPLP2 | -1.583597 | 3.27E-09 | 1.20E-07 | IER5 | -1.411334 | 1.01E-05 | 0.0001696 | MARCKSL1 | -2.167735 | 7.22E-05 | 0.0017525 |
| GAS5 | -1.587199 | 5.66E-07 | 1.39E-05 | PABPC4 | -1.411913 | 0.0002126 | 0.0025828 | CENPV | -2.177015 | 0.0001496 | 0.0033689 |
| RPS10 | -1.588343 | 3.47E-09 | 1.26E-07 | RPS10 | -1.413537 | 5.26E-09 | 1.71E-07 | ANTXR2 | -2.177212 | 7.77E-06 | 0.0002613 |
| STX17 | -1.592788 | 0.0008524 | 0.0093577 | CBX7 | -1.415514 | 0.0004036 | 0.0044822 | KLRG1 | -2.179987 | 1.92E-05 | 0.0005566 |
| RPS25 | -1.593295 | 9.82E-13 | 7.11E-11 | RPS23 | -1.418212 | 8.63E-12 | 4.56E-10 | FAM118A | -2.182082 | 0.0005056 | 0.0093436 |
| TPT1 | -1.598696 | 6.85E-13 | 5.09E-11 | THEM4 | -1.418437 | 0.0017299 | 0.0155773 | AHCY | -2.185622 | 0.0054068 | 0.0588612 |
| RPS19 | -1.599677 | 2.32E-09 | 8.66E-08 | CKAP2 | -1.41904 | 0.0010974 | 0.0105275 | LIMA1 | -2.189881 | 0.000189 | 0.0040911 |
| JUND | -1.631656 | 7.73E-08 | 2.15E-06 | 11-Sep | -1.420141 | 1.63E-06 | 3.23E-05 | PSMB5 | -2.191122 | 0.0083998 | 0.0812942 |
| ZNF101 | -1.632092 | 0.0036702 | 0.0308535 | RPS3A | -1.421254 | 1.43E-10 | 6.20E-09 | CHD7 | -2.191352 | 7.35E-12 | 8.11E-10 |
| MXI1 | -1.640628 | 0.0041524 | 0.033988 | KCTD18 | -1.425798 | 0.0007561 | 0.0076916 | PDE4D | -2.199208 | 2.37E-09 | 1.64E-07 |
| FAM222B | -1.644648 | 0.0033718 | 0.028605 | LINC00936 | -1.430514 | 0.002915 | 0.0243096 | TRABD2A | -2.205896 | 0.0010546 | 0.0167947 |
| LRRC8C | -1.651724 | 0.0066858 | 0.049688 | RPL22 | -1.432555 | 3.54E-12 | 1.96E-10 | CDC14A | -2.22703 | 3.00E-09 | 2.03E-07 |
| PRKCI | -1.652846 | 0.0054682 | 0.0425205 | TRIM4 | -1.433147 | 0.0087512 | 0.0605456 | TTC33 | -2.237809 | 0.000883 | 0.0147793 |
| SCOC | -1.654645 | 0.0098567 | 0.067353 | SGSM3 | -1.434905 | 0.0020813 | 0.0182278 | LOC10192 | -2.238048 | 0.0058021 | 0.0620373 |
| RALGPS2 | -1.656959 | 0.0025912 | 0.0232988 | THAP5 | -1.435458 | 0.0004549 | 0.0049711 | ZNF204P | -2.243885 | 0.0016592 | 0.0242576 |
| RPS13 | -1.657765 | 2.84E-11 | 1.54E-09 | NFIC | -1.435704 | 4.07E-05 | 0.0006009 | SOCS2 | -2.260267 | 0.0024343 | 0.0320268 |
| RPS21 | -1.670898 | 4.98E-11 | 2.57E-09 | HECA | -1.435939 | 1.99E-06 | 3.85E-05 | S100A6 | -2.261595 | 1.01E-06 | 4.12E-05 |
| WDR6 | -1.681786 | 0.0017795 | 0.0171639 | CEP170 | -1.438046 | 0.0006865 | 0.007101 | POLR3C | -2.265585 | 0.0009504 | 0.0156887 |
| RPL36 | -1.683919 | 7.60E-11 | 3.84E-09 | RPS28 | -1.438085 | 1.46E-10 | 6.30E-09 | FLT3LG | -2.265662 | 9.75E-09 | 6.01E-07 |
| AGO2 | -1.690015 | 0.0060714 | 0.0460177 | MRPL3 | -1.440172 | 0.0003832 | 0.0042839 | PHTF2 | -2.266018 | 5.39E-12 | 6.16E-10 |
| RPS2 | -1.698564 | 1.19E-08 | 3.91E-07 | RPL7 | -1.44513 | 2.22E-12 | 1.28E-10 | CCDC86 | -2.273329 | 0.0010124 | 0.0164021 |
| SOS1 | -1.698713 | 6.37E-06 | 0.0001264 | SSFA2 | -1.446901 | 0.0089453 | 0.061614 | MAGEF1 | -2.28128 | 7.56E-06 | 0.0002552 |
| FAM213B | -1.711513 | 0.0009596 | 0.0103318 | PPP1R15A | -1.447119 | 0.0002984 | 0.0034444 | SPPL2B | -2.283949 | 0.0056551 | 0.0610097 |
| TNFSF10 | -1.719673 | 0.0007708 | 0.008631 | PIEZO1 | -1.448626 | 0.0016271 | 0.0147665 | TBC1D4 | -2.296363 | 7.25E-07 | 3.06E-05 |
| REEP3 | -1.720945 | 0.0026351 | 0.0236092 | ERO1L | -1.44974 | 2.58E-06 | 4.84E-05 | ADAM22 | -2.311078 | 0.0006458 | 0.0114494 |
| NFKB1 | -1.721879 | 0.0001344 | 0.0019068 | TNFSF10 | -1.453749 | 0.0002389 | 0.0028531 | TNFRSF10 | -2.321804 | 0.0092887 | 0.0870828 |
| RPL13 | -1.725929 | 4.07E-14 | 3.64E-12 | VWA8 | -1.455783 | 0.0025019 | 0.0214045 | ID3 | -2.331514 | 0.0091527 | 0.0862315 |

| | | | | | | | | | | | |
|---|---|---|---|---|---|---|---|---|---|---|---|
| TUBB4B | -1.728648 | 0.0071938 | 0.0526251 | RPL36 | -1.456408 | 7.38E-10 | 2.80E-08 | PPP2R3A | -2.332975 | 0.0029688 | 0.0366461 |
| GUCY1A3 | -1.736536 | 0.0004619 | 0.0055731 | KDM5B | -1.46271 | 0.0032903 | 0.0269068 | TGIF1 | -2.335015 | 4.19E-06 | 0.000154 |
| RPL9 | -1.7373 | 4.40E-13 | 3.39E-11 | KLF6 | -1.464635 | 1.78E-09 | 6.35E-08 | MORC2-A | -2.348833 | 0.0004959 | 0.0091814 |
| IER5 | -1.752114 | 2.19E-06 | 4.77E-05 | SLC7A6 | -1.470035 | 0.0002744 | 0.0032069 | EBLN2 | -2.368815 | 0.0015214 | 0.0227219 |
| RPL32 | -1.757193 | 2.53E-12 | 1.69E-10 | SNX30 | -1.473764 | 0.0048726 | 0.0376819 | BCL2 | -2.376 | 8.39E-06 | 0.0002773 |
| FKBP5 | -1.76193 | 8.05E-05 | 0.0012006 | HSP90AB1 | -1.474235 | 8.30E-10 | 3.12E-08 | PTGER4 | -2.397711 | 4.35E-10 | 3.79E-08 |
| RPLP0 | -1.767848 | 5.45E-10 | 2.31E-08 | RPS6KA5 | -1.477823 | 0.0014014 | 0.0130506 | GRAP2 | -2.406162 | 5.78E-05 | 0.0014331 |
| ASPH | -1.771616 | 0.0087437 | 0.0616017 | RPL30 | -1.479906 | 1.60E-13 | 1.08E-11 | TESPA1 | -2.413103 | 1.38E-09 | 1.04E-07 |
| CRTAP | -1.779128 | 1.51E-05 | 0.000273 | AMMECR1 | -1.480113 | 4.00E-05 | 0.0005918 | MTFP1 | -2.442295 | 0.0016232 | 0.0238045 |
| MAP7D3 | -1.796557 | 0.0047124 | 0.0377625 | GPX4 | -1.480472 | 8.48E-05 | 0.0011497 | GADD45G | -2.450084 | 7.54E-05 | 0.001816 |
| COG2 | -1.797826 | 0.0054322 | 0.0423189 | RPL9 | -1.482577 | 9.97E-11 | 4.48E-09 | TTC9 | -2.460812 | 5.72E-05 | 0.001428 |
| CD59 | -1.801001 | 0.0019309 | 0.0182404 | SDCCAG3 | -1.482677 | 0.0029779 | 0.0246793 | BIRC3 | -2.468092 | 8.75E-21 | 3.00E-18 |
| PBXIP1 | -1.804994 | 9.52E-07 | 2.27E-05 | RPS2 | -1.484889 | 2.66E-07 | 6.29E-06 | KIT | -2.473385 | 1.88E-09 | 1.34E-07 |
| GPR155 | -1.808747 | 0.0042029 | 0.0343013 | NT5DC1 | -1.48857 | 0.0016287 | 0.014767 | CSGALNA | -2.479946 | 2.88E-06 | 0.0001095 |
| RPL11 | -1.809872 | 2.72E-14 | 2.56E-12 | RPL32 | -1.493843 | 1.54E-10 | 6.62E-09 | SLC48A1 | -2.485496 | 0.0019283 | 0.0269724 |
| RPS12 | -1.811169 | 3.47E-19 | 6.33E-17 | NUDT4 | -1.495745 | 3.50E-07 | 8.13E-06 | NCKAP1 | -2.488998 | 0.0023627 | 0.03143 |
| PDE4DIP | -1.812406 | 0.0087427 | 0.0616017 | RPLP1 | -1.497057 | 1.08E-10 | 4.81E-09 | FAM24B | -2.491417 | 0.0023284 | 0.0310592 |
| RPS5 | -1.815045 | 8.58E-12 | 5.18E-10 | NAP1L2 | -1.507185 | 0.0090077 | 0.0619499 | BCL7A | -2.495899 | 0.0055943 | 0.0605587 |
| ZNF445 | -1.81507 | 0.0096595 | 0.066316 | SPTBN1 | -1.508531 | 3.89E-08 | 1.08E-06 | TNFRSF10 | -2.529241 | 7.46E-05 | 0.0018025 |
| GLTSCR2 | -1.815307 | 3.26E-13 | 2.60E-11 | RPL39 | -1.50926 | 1.60E-10 | 6.81E-09 | AK5 | -2.558826 | 0.0002308 | 0.0048534 |
| CCDC86 | -1.825317 | 0.0044816 | 0.0362588 | STAM | -1.512917 | 0.001386 | 0.0129339 | CLUHP3 | -2.568076 | 2.27E-06 | 8.75E-05 |
| RPL8 | -1.826608 | 1.33E-13 | 1.11E-11 | IRF2BP2 | -1.513929 | 7.05E-09 | 2.24E-07 | VCAN | -2.571604 | 0.005155 | 0.056693 |
| ZFAND1 | -1.832186 | 0.0054636 | 0.0425205 | EEF1G | -1.521339 | 5.17E-12 | 2.81E-10 | ATP8B2 | -2.57852 | 8.46E-11 | 8.12E-09 |
| THAP5 | -1.840371 | 0.0020347 | 0.0191131 | TBC1D15 | -1.525183 | 0.0006521 | 0.0067879 | RHOBTB1 | -2.580014 | 0.008306 | 0.0807124 |
| NABP1 | -1.843423 | 0.0067143 | 0.0498192 | NAMPT | -1.526866 | 0.0036668 | 0.0295501 | ZBED6CL | -2.59189 | 1.21E-05 | 0.0003743 |
| BCL3 | -1.845536 | 0.0007534 | 0.0084698 | SVIP | -1.527802 | 0.0023959 | 0.0206137 | LDLR | -2.603711 | 5.39E-05 | 0.0013531 |
| TUBA1A | -1.846758 | 3.77E-08 | 1.12E-06 | PAIP1 | -1.53422 | 0.0016473 | 0.0149204 | PAG1 | -2.608713 | 1.21E-19 | 3.61E-17 |
| SPIDR | -1.852056 | 0.0083823 | 0.0596058 | LRRC75A-AS | -1.54047 | 3.97E-07 | 9.13E-06 | BCAS1 | -2.629445 | 8.75E-05 | 0.0020778 |
| RPL18A | -1.860338 | 4.67E-14 | 4.08E-12 | EEF1B2 | -1.542864 | 2.38E-11 | 1.18E-09 | SGK1 | -2.635913 | 0.0013737 | 0.0208629 |
| RPS8 | -1.862093 | 2.89E-19 | 5.39E-17 | RHOB | -1.547453 | 0.001683 | 0.0152148 | IL32 | -2.651049 | 2.69E-17 | 5.86E-15 |
| GKAP1 | -1.866477 | 0.0056394 | 0.0433209 | RPL5 | -1.547801 | 3.07E-14 | 2.27E-12 | PLAGL1 | -2.658324 | 0.0018421 | 0.0261541 |
| APOL3 | -1.869424 | 0.001559 | 0.0153816 | TPT1 | -1.548538 | 1.70E-13 | 1.14E-11 | ARL4A | -2.667448 | 7.17E-08 | 3.68E-06 |
| GTPBP10 | -1.885935 | 0.009587 | 0.0658849 | SLC39A8 | -1.550698 | 0.0020396 | 0.0179473 | MID1IP1 | -2.671526 | 5.67E-06 | 0.0001972 |
| CBX7 | -1.89866 | 1.99E-05 | 0.0003548 | MMGT1 | -1.55084 | 0.0002516 | 0.0029709 | FAM92A1 | -2.707624 | 0.004516 | 0.0512574 |
| ANXA5 | -1.905642 | 2.10E-05 | 0.0003721 | PEBP1 | -1.552314 | 3.36E-06 | 6.16E-05 | HK2 | -2.713655 | 3.26E-05 | 0.000889 |
| ICAM3 | -1.931258 | 0.0003866 | 0.0047751 | SNHG12 | -1.552588 | 0.0088283 | 0.0609877 | PAFAH1B3 | -2.748291 | 0.0001224 | 0.0028151 |
| CFL2 | -1.938412 | 0.0039103 | 0.0324493 | ARHGAP21 | -1.555648 | 0.0008092 | 0.0081422 | CADM1 | -2.76473 | 0.0003772 | 0.0073089 |
| PPP1R15A | -1.946987 | 1.49E-05 | 0.0002703 | RPL11 | -1.56569 | 2.96E-14 | 2.23E-12 | DIP2C | -2.782101 | 0.0011068 | 0.01746 |
| ANAPC13 | -1.953955 | 0.0030973 | 0.0267055 | TOB1 | -1.577155 | 0.0002486 | 0.0029461 | ABCA1 | -2.787034 | 0.0001793 | 0.0039348 |
| SYPL1 | -1.955317 | 0.0033721 | 0.028605 | SOS1 | -1.582472 | 2.27E-05 | 0.0003557 | CYP2E1 | -2.80192 | 2.13E-06 | 8.26E-05 |
| HIF1A-AS2 | -1.967656 | 2.45E-05 | 0.0004265 | MAP9 | -1.586557 | 0.0074975 | 0.0531023 | N4BP3 | -2.802647 | 0.0007477 | 0.0128291 |
| R3HDM4 | -1.968793 | 0.0042768 | 0.034803 | NFKBIZ | -1.594864 | 1.67E-06 | 3.31E-05 | LYL1 | -2.844859 | 0.0004129 | 0.0078582 |
| TGIF1 | -1.979672 | 0.0029048 | 0.0253842 | YPEL2 | -1.595944 | 6.51E-07 | 1.44E-05 | ZFYVE28 | -2.858149 | 0.0001427 | 0.0032443 |
| FAM210B | -1.987535 | 0.0015892 | 0.0156274 | PRNP | -1.596614 | 4.65E-06 | 8.21E-05 | E2F2 | -2.866599 | 0.006854 | 0.0702186 |
| WWC2 | -1.988691 | 0.0050333 | 0.0397271 | DEGS1 | -1.597079 | 0.0009839 | 0.0096179 | BACE2 | -2.870898 | 1.12E-05 | 0.0003525 |
| A2M | -1.989079 | 4.96E-05 | 0.0007802 | GTPBP8 | -1.598166 | 0.0057996 | 0.0435063 | CD84 | -2.872522 | 3.93E-15 | 6.73E-13 |
| SLC25A33 | -1.990866 | 0.007027 | 0.0519023 | TSPYL4 | -1.6077 | 2.00E-05 | 0.0003183 | INPP5A | -2.899022 | 0.0003226 | 0.0064329 |
| A2M-AS1 | -1.994314 | 0.0008874 | 0.009691 | MAF | -1.611103 | 1.63E-05 | 0.0002627 | GRAP | -2.913544 | 6.03E-06 | 0.0002066 |
| DLL1 | -1.998264 | 0.009136 | 0.0633037 | TUBA1A | -1.613022 | 3.75E-09 | 1.26E-07 | MAST4 | -2.915123 | 7.50E-14 | 1.12E-11 |
| ZMAT1 | -2.005049 | 7.19E-06 | 0.0001414 | RPS6 | -1.614528 | 1.83E-14 | 1.43E-12 | ACTN1 | -2.933002 | 8.44E-06 | 0.0002783 |
| BCL2 | -2.010276 | 0.000334 | 0.0041798 | SS18L2 | -1.614858 | 0.0039306 | 0.0311518 | LTB | -2.941794 | 1.67E-35 | 4.00E-32 |
| NADK2 | -2.013363 | 0.0009785 | 0.0105083 | HSPBAP1 | -1.62097 | 0.0049355 | 0.0381046 | DGKH | -2.944914 | 2.04E-07 | 9.62E-06 |
| MINOS1P1 | -2.016167 | 0.0088108 | 0.0618546 | RPS12 | -1.621399 | 1.99E-15 | 1.86E-13 | TPST1 | -2.94556 | 0.0038745 | 0.0457077 |
| ACVR2A | -2.016616 | 0.0086622 | 0.0611307 | BCL3 | -1.621472 | 0.0027643 | 0.0233263 | BCDIN3D | -2.965225 | 0.0005626 | 0.0102388 |

| | | | | | | | | | | | |
|---|---|---|---|---|---|---|---|---|---|---|---|
| RPS26P11 | -2.017357 | 0.0012334 | 0.0126622 | HK1 | -1.625937 | 7.12E-05 | 0.0009883 | IL2RA | -2.974783 | 3.96E-13 | 5.20E-11 |
| SAMSN1 | -2.020938 | 4.86E-06 | 9.79E-05 | RBM7 | -1.626044 | 0.006831 | 0.0497129 | LINC00892 | -3.027948 | 0.0074818 | 0.0746456 |
| PRNP | -2.024683 | 1.07E-06 | 2.49E-05 | NME9 | -1.626544 | 0.0033993 | 0.0277249 | LOC10050 | -3.033518 | 4.47E-05 | 0.0011573 |
| MPP7 | -2.026034 | 0.00486 | 0.0385771 | TNRC6C-AS1 | -1.62754 | 0.002919 | 0.0243096 | FRY | -3.042722 | 1.90E-17 | 4.30E-15 |
| JUN | -2.028338 | 1.77E-13 | 1.45E-11 | RPS13 | -1.6281 | 7.45E-11 | 3.40E-09 | LRP12 | -3.043746 | 1.69E-05 | 0.000501 |
| TRADD | -2.035669 | 0.0057295 | 0.0438219 | ACTN1 | -1.634649 | 0.0074236 | 0.0527398 | TNFRSF25 | -3.060335 | 6.98E-10 | 5.72E-08 |
| MDFIC | -2.036314 | 6.70E-08 | 1.90E-06 | PPAPDC1B | -1.638628 | 0.0012462 | 0.0117464 | IL17RB | -3.075381 | 0.0028201 | 0.0353015 |
| SC5D | -2.03736 | 0.0002664 | 0.0034361 | RPS14P3 | -1.640726 | 0.0004219 | 0.0046554 | COL4A3 | -3.087489 | 0.0005857 | 0.0105397 |
| IL6ST | -2.039352 | 4.87E-05 | 0.0007707 | PIK3IP1 | -1.649591 | 6.80E-07 | 1.50E-05 | CFAP36 | -3.104628 | 2.03E-12 | 2.49E-10 |
| FAM129A | -2.039464 | 2.63E-05 | 0.0004557 | MXI1 | -1.654815 | 0.0010662 | 0.0103025 | PTP4A3 | -3.161394 | 8.24E-05 | 0.0019715 |
| MRPL41 | -2.041233 | 0.0088161 | 0.0618546 | HEBP2 | -1.65964 | 8.30E-05 | 0.0011302 | STARD4 | -3.192929 | 1.33E-11 | 1.42E-09 |
| SOCS5 | -2.046013 | 0.0088164 | 0.0618546 | IFRD1 | -1.660347 | 1.34E-05 | 0.0002197 | IFIT3 | -3.223939 | 0.0004396 | 0.0082511 |
| PLXDC2 | -2.049425 | 1.14E-05 | 0.000214 | HINT1 | -1.662314 | 3.15E-12 | 1.76E-10 | TUBB2A | -3.289417 | 0.0003188 | 0.0063705 |
| TRAF3IP2 | -2.054339 | 0.0007174 | 0.0081308 | ACYP1 | -1.663415 | 0.0052188 | 0.0399263 | ENO2 | -3.329322 | 9.91E-05 | 0.0023299 |
| THEM4 | -2.055919 | 0.0043898 | 0.0355845 | FAM213B | -1.667668 | 4.10E-05 | 0.0006041 | ERN1 | -3.3312 | 0.0002223 | 0.0047071 |
| CSRNP1 | -2.065983 | 5.66E-05 | 0.0008765 | KLF10 | -1.668469 | 0.0008945 | 0.0088652 | CRIP2 | -3.337027 | 0.0014972 | 0.0224589 |
| YBX3 | -2.075296 | 0.0037582 | 0.031442 | RPS5 | -1.669604 | 3.17E-13 | 2.08E-11 | LOC10192 | -3.34295 | 0.0025072 | 0.032583 |
| LINS | -2.076841 | 0.0001121 | 0.0016228 | PLEKHA5 | -1.670228 | 0.0077593 | 0.0546632 | CD6 | -3.358854 | 0.0012406 | 0.0192357 |
| EPB41L2 | -2.105065 | 0.0050644 | 0.0399353 | EDEM3 | -1.674635 | 9.37E-06 | 0.0001578 | CHRM3-A | -3.413379 | 0.0010695 | 0.016983 |
| COMMD10 | -2.105701 | 0.0079742 | 0.0573353 | SIK3 | -1.680789 | 1.41E-07 | 3.49E-06 | BCORL1 | -3.447029 | 0.001841 | 0.0261541 |
| EML4 | -2.106095 | 1.04E-14 | 1.08E-12 | LACTB | -1.681482 | 0.0002839 | 0.003297 | LINC00324 | -3.463051 | 0.0002725 | 0.0056089 |
| CD44 | -2.107099 | 8.86E-10 | 3.61E-08 | MPP7 | -1.684789 | 0.0002133 | 0.0025878 | AQP3 | -3.478543 | 9.71E-06 | 0.0003156 |
| ZXDB | -2.10943 | 0.0011082 | 0.0115741 | TRADD | -1.686296 | 0.0004814 | 0.0052426 | CD3G | -3.491915 | 9.75E-08 | 4.82E-06 |
| ARL4A | -2.110133 | 1.69E-06 | 3.83E-05 | SOX12 | -1.686483 | 0.0042659 | 0.0334088 | LZTFL1 | -3.515663 | 5.16E-07 | 2.27E-05 |
| KDM6B | -2.110582 | 0.0008488 | 0.0093307 | SUN1 | -1.688753 | 0.0026827 | 0.0227412 | PKIA | -3.596375 | 2.40E-06 | 9.23E-05 |
| IL32 | -2.119654 | 1.21E-07 | 3.25E-06 | ABCE1 | -1.69096 | 3.16E-08 | 8.84E-07 | STMN3 | -3.601367 | 2.57E-14 | 4.03E-12 |
| LACTB | -2.125778 | 0.0017834 | 0.0171756 | JUNB | -1.691528 | 7.34E-10 | 2.80E-08 | DUSP16 | -3.671523 | 4.01E-17 | 8.55E-15 |
| STRBP | -2.136909 | 5.36E-05 | 0.0008389 | HIVEP2 | -1.705343 | 0.0011111 | 0.0106254 | SLC25A15 | -3.732459 | 3.80E-07 | 1.71E-05 |
| PIK3IP1 | -2.14147 | 5.43E-07 | 1.34E-05 | AHI1 | -1.705514 | 0.0014979 | 0.0138106 | SLC8A1 | -3.75074 | 0.0015593 | 0.0230794 |
| FAM162A | -2.145968 | 0.0058017 | 0.0442935 | USP53 | -1.706857 | 0.0056711 | 0.04289 | MRPL42P5 | -3.764732 | 4.09E-05 | 0.0010771 |
| SDCCAG3 | -2.146613 | 0.0003183 | 0.0040014 | SRM | -1.707054 | 0.0015201 | 0.0139738 | MIATNB | -3.777897 | 1.15E-05 | 0.0003583 |
| BTG2 | -2.151952 | 1.73E-06 | 3.90E-05 | MYADM | -1.707084 | 0.0001741 | 0.0021685 | C12orf66 | -3.806478 | 4.15E-05 | 0.0010898 |
| ZEB1 | -2.156047 | 7.30E-10 | 3.02E-08 | MEF2A | -1.710724 | 0.0003367 | 0.0038337 | PHLDB3 | -3.882696 | 0.0002699 | 0.0055674 |
| METTL16 | -2.156883 | 0.0048211 | 0.0383766 | NAP1L1 | -1.710955 | 8.62E-11 | 3.91E-09 | IL23A | -3.892008 | 3.00E-07 | 1.37E-05 |
| KLF7 | -2.158401 | 9.51E-09 | 3.22E-07 | RHOH | -1.718582 | 0.0002166 | 0.0026212 | SERINC5 | -3.918155 | 9.21E-19 | 2.26E-16 |
| IMPACT | -2.164255 | 0.0015997 | 0.0156946 | IRS2 | -1.719105 | 2.03E-06 | 3.92E-05 | NT5E | -3.934307 | 0.000355 | 0.0069623 |
| ATP8B2 | -2.167307 | 1.47E-05 | 0.0002682 | AGO2 | -1.71984 | 0.0041892 | 0.0329196 | P2RY1 | -3.944936 | 0.0006084 | 0.0109076 |
| IFI44L | -2.183151 | 0.0022582 | 0.0208546 | SC5D | -1.720501 | 1.63E-05 | 0.0002629 | CNTN3 | -3.959854 | 0.0002743 | 0.0056328 |
| C9orf89 | -2.193881 | 0.0002696 | 0.0034674 | RPL13 | -1.723364 | 6.10E-14 | 4.38E-12 | NRIP3 | -3.980683 | 8.89E-07 | 3.71E-05 |
| FXYD5 | -2.201728 | 9.92E-12 | 5.91E-10 | IFT57 | -1.724351 | 7.39E-06 | 0.0001271 | KLF5 | -4.01866 | 2.12E-08 | 1.20E-06 |
| CEP68 | -2.218486 | 0.0006516 | 0.0075273 | TAB2 | -1.727754 | 1.93E-06 | 3.77E-05 | TRAT1 | -4.05739 | 8.67E-08 | 4.38E-06 |
| GRASP | -2.227151 | 0.0095844 | 0.0658849 | MTFMT | -1.739356 | 0.0053896 | 0.0410292 | RTKN2 | -4.064434 | 1.55E-09 | 1.14E-07 |
| LONRF1 | -2.237142 | 0.0023068 | 0.0211949 | TUBE1 | -1.745399 | 0.0062869 | 0.0464838 | KIF5C | -4.082013 | 3.46E-09 | 2.32E-07 |
| DGKA | -2.245567 | 1.75E-10 | 8.03E-09 | AIM1 | -1.746367 | 6.36E-06 | 0.0001103 | SCN1B | -4.096704 | 1.41E-05 | 0.0004253 |
| RASA4 | -2.246947 | 0.0003936 | 0.004847 | RPL18A | -1.749588 | 1.52E-13 | 1.03E-11 | BCL11B | -4.115531 | 9.17E-17 | 1.87E-14 |
| CHCHD10 | -2.250382 | 0.0037351 | 0.0313361 | SOCS5 | -1.755565 | 0.0038868 | 0.0308973 | MAF | -4.261265 | 1.65E-23 | 6.58E-21 |
| BBIP1 | -2.251865 | 0.0015134 | 0.0150228 | RPL4 | -1.757873 | 1.01E-16 | 1.11E-14 | GRK5 | -4.26575 | 3.38E-08 | 1.84E-06 |
| CD84 | -2.279791 | 1.76E-10 | 8.03E-09 | LTA4H | -1.759133 | 3.09E-08 | 8.69E-07 | FGFR1 | -4.27328 | 1.30E-07 | 6.25E-06 |
| TNFRSF10B | -2.281292 | 0.0055223 | 0.0429016 | SNHG16 | -1.761056 | 2.71E-07 | 6.37E-06 | 8-Sep | -4.27514 | 4.64E-06 | 0.0001679 |
| USP53 | -2.281665 | 4.05E-05 | 0.000659 | ITGB1 | -1.761788 | 5.83E-11 | 2.73E-09 | LRP6 | -4.297406 | 2.84E-05 | 0.0007935 |
| RAB31 | -2.282864 | 0.0094471 | 0.0650836 | PBXIP1 | -1.763592 | 1.07E-08 | 3.29E-07 | SPDYA | -4.316516 | 7.04E-08 | 3.63E-06 |
| PCYOX1L | -2.284905 | 0.0039322 | 0.0325664 | GKAP1 | -1.766179 | 0.0041658 | 0.0327913 | KIAA0226L | -4.345785 | 0.0004281 | 0.0080827 |
| LDHB | -2.286106 | 2.27E-14 | 2.25E-12 | CCT6P1 | -1.769533 | 0.001835 | 0.0163488 | CFH | -4.379241 | 1.48E-07 | 7.06E-06 |
| AHCY | -2.296936 | 0.0025833 | 0.0232522 | TSC22D3 | -1.77058 | 2.32E-10 | 9.45E-09 | HLF | -4.390345 | 4.76E-10 | 4.12E-08 |

| | | | | | | | | | | | |
|---|---|---|---|---|---|---|---|---|---|---|---|
| OXNAD1 | -2.303335 | 0.0030919 | 0.0267055 | ZMIZ1 | -1.771363 | 7.71E-08 | 2.03E-06 | TNFSF8 | -4.409603 | 1.58E-09 | 1.15E-07 |
| EPHA4 | -2.305665 | 7.25E-06 | 0.0001419 | ANK3 | -1.772073 | 0.0003887 | 0.0043409 | PSD3 | -4.596284 | 4.96E-10 | 4.25E-08 |
| FLT3LG | -2.307682 | 0.000135 | 0.0019114 | ATF7IP2 | -1.773586 | 0.0020011 | 0.0176758 | ROGDI | -4.691861 | 1.20E-09 | 9.39E-08 |
| SVIL | -2.30839 | 1.45E-05 | 0.0002645 | PPA1 | -1.790377 | 0.0011571 | 0.0109855 | MBOAT2 | -4.735789 | 0.0001418 | 0.0032301 |
| WASL | -2.310766 | 0.0026055 | 0.0234023 | RPL8 | -1.794625 | 7.67E-17 | 8.66E-15 | EMR2 | -4.749823 | 5.22E-06 | 0.0001839 |
| FRY | -2.319331 | 9.14E-05 | 0.0013512 | NPHP3 | -1.795336 | 0.0001747 | 0.0021733 | SCN2A | -4.836458 | 7.30E-08 | 3.73E-06 |
| PHTF2 | -2.385712 | 4.49E-09 | 1.61E-07 | PAFAH1B3 | -1.797583 | 0.009304 | 0.0633756 | TNFRSF19 | -4.839114 | 5.45E-06 | 0.0001909 |
| CASK | -2.392089 | 0.0031476 | 0.0270554 | FKBP5 | -1.799949 | 4.47E-05 | 0.000656 | PHLDA2 | -4.855379 | 0.0001246 | 0.0028518 |
| SNX30 | -2.397653 | 0.0093 | 0.064126 | CYSLTR1 | -1.804112 | 3.75E-06 | 6.78E-05 | TLE2 | -4.898821 | 8.01E-05 | 0.0019202 |
| IL4R | -2.400107 | 0.0002341 | 0.0030768 | EML4 | -1.804709 | 3.59E-11 | 1.74E-09 | RGS18 | -5.013882 | 1.74E-06 | 6.86E-05 |
| RNF144A | -2.412203 | 0.0012258 | 0.012615 | NUP43 | -1.80494 | 0.0034418 | 0.0280183 | IL6R | -5.058671 | 1.56E-19 | 4.53E-17 |
| VIM | -2.429524 | 1.10E-16 | 1.54E-14 | KLF7 | -1.805625 | 1.06E-07 | 2.72E-06 | HTR1F | -5.066316 | 7.33E-12 | 8.11E-10 |
| SOX4 | -2.435468 | 1.87E-07 | 4.95E-06 | SELL | -1.818659 | 7.14E-07 | 1.57E-05 | THEMIS | -5.111496 | 9.68E-09 | 6.01E-07 |
| RAPGEF2 | -2.436143 | 0.0001315 | 0.0018724 | IRGQ | -1.819483 | 0.0041122 | 0.0323971 | HPGD | -5.176032 | 1.64E-29 | 1.12E-26 |
| TRAF3 | -2.446084 | 0.0015482 | 0.0152963 | MAPK6 | -1.82757 | 8.67E-05 | 0.0011706 | FSCN1 | -5.241288 | 3.44E-06 | 0.0001292 |
| SGSM3 | -2.463459 | 0.0004151 | 0.0050667 | FAM84B | -1.834624 | 1.60E-07 | 3.93E-06 | ZC3H12D | -5.25548 | 2.23E-07 | 1.05E-05 |
| FLNB | -2.475303 | 0.0027704 | 0.0243877 | UNC93B1 | -1.83632 | 0.004856 | 0.037617 | TCEA3 | -5.333783 | 4.09E-08 | 2.14E-06 |
| FAM210A | -2.497774 | 0.0001114 | 0.0016184 | IFI44 | -1.836807 | 0.0045474 | 0.0353739 | SETD7 | -5.417696 | 1.84E-11 | 1.90E-09 |
| DUSP1 | -2.501474 | 1.32E-11 | 7.82E-10 | PPT1 | -1.840156 | 3.55E-05 | 0.0005341 | ANTXRLP1 | -5.532857 | 9.77E-09 | 6.01E-07 |
| SPRYD7 | -2.50191 | 0.0052967 | 0.0414165 | IFI44L | -1.840554 | 0.0048601 | 0.0376173 | MYRF | -5.550998 | 4.31E-09 | 2.79E-07 |
| TP53INP1 | -2.509952 | 7.96E-13 | 5.87E-11 | KCNA3 | -1.844184 | 2.60E-11 | 1.28E-09 | CR2 | -5.607805 | 1.90E-05 | 0.000555 |
| TCEAL4 | -2.522364 | 0.0007726 | 0.00864 | KLHL15 | -1.861708 | 0.0008054 | 0.0081129 | IL1R1 | -5.656374 | 3.98E-16 | 7.79E-14 |
| FAM115C | -2.525615 | 0.000367 | 0.0045587 | SAMD12 | -1.862575 | 0.008699 | 0.06041 | GCSAM | -5.663606 | 2.21E-11 | 2.23E-09 |
| CYSLTR1 | -2.531292 | 3.80E-06 | 7.81E-05 | CSRNP1 | -1.866141 | 9.37E-07 | 1.98E-05 | C10orf128 | -5.748311 | 1.65E-06 | 6.56E-05 |
| PIM2 | -2.539346 | 7.43E-08 | 2.08E-06 | PDCD5 | -1.866287 | 0.0060186 | 0.0449673 | C6orf10 | -5.888162 | 5.18E-09 | 3.33E-07 |
| MTFP1 | -2.544007 | 0.0076881 | 0.0556113 | PPP1R9A | -1.885495 | 0.0006438 | 0.0067415 | GAB2 | -6.1113 | 1.18E-08 | 7.01E-07 |
| FOSB | -2.547683 | 0.0084399 | 0.0599139 | RPL10A | -1.886923 | 4.78E-14 | 3.51E-12 | CLIC4 | -6.171744 | 2.26E-13 | 3.09E-11 |
| ARHGAP5 | -2.548155 | 2.02E-12 | 1.37E-10 | MORC2-AS1 | -1.889287 | 0.0068449 | 0.0497745 | ALS2CL | -6.433869 | 3.13E-07 | 1.42E-05 |
| URB1 | -2.584945 | 0.0056873 | 0.043539 | CNST | -1.889826 | 7.40E-07 | 1.62E-05 | GATA3-AS | -6.770723 | 1.07E-16 | 2.14E-14 |
| SNHG8 | -2.599694 | 1.24E-10 | 6.07E-09 | EIF3E | -1.89388 | 2.12E-10 | 8.71E-09 | GCNT4 | -6.787775 | 1.01E-08 | 6.08E-07 |
| TESPA1 | -2.611286 | 6.78E-09 | 2.39E-07 | FAM129A | -1.894244 | 9.58E-06 | 0.0001609 | CD28 | -6.975289 | 1.65E-22 | 6.34E-20 |
| SMPD3 | -2.614421 | 0.0002573 | 0.0033453 | CRYBG3 | -1.909462 | 0.0010745 | 0.0103607 | GAPT | -6.986329 | 1.20E-09 | 9.39E-08 |
| TUBE1 | -2.631861 | 0.0015605 | 0.0153816 | NOSIP | -1.909498 | 0.0001588 | 0.0019951 | NSG1 | -7.10343 | 3.48E-13 | 4.64E-11 |
| NXT1 | -2.633099 | 2.48E-05 | 0.0004314 | NADK2 | -1.917145 | 0.0010016 | 0.0097493 | GAP43 | -7.129039 | 1.18E-33 | 1.26E-30 |
| PAG1 | -2.643991 | 2.72E-18 | 4.48E-16 | JUN | -1.917215 | 3.37E-13 | 2.20E-11 | PKIB | -7.154968 | 1.76E-19 | 4.96E-17 |
| PIM3 | -2.655902 | 7.09E-05 | 0.0010665 | MAML2 | -1.918819 | 0.0006991 | 0.0072061 | KCTD12 | -7.159776 | 1.23E-17 | 2.96E-15 |
| LST1 | -2.660067 | 0.0001452 | 0.0020318 | MAP3K1 | -1.928873 | 2.11E-11 | 1.06E-09 | FERMT2 | -7.209531 | 3.18E-10 | 2.83E-08 |
| RELB | -2.664346 | 0.0005897 | 0.0069075 | FXYD5 | -1.932649 | 2.80E-09 | 9.66E-08 | ADAM19 | -7.301496 | 1.26E-07 | 6.05E-06 |
| ETS2 | -2.668166 | 0.0008696 | 0.0095212 | CADM1 | -1.934666 | 0.0085389 | 0.0594319 | FAAH2 | -7.367155 | 1.93E-17 | 4.30E-15 |
| MGAT5 | -2.682457 | 6.62E-11 | 3.39E-09 | RPLP0 | -1.943758 | 1.52E-12 | 9.07E-11 | CD5 | -7.371774 | 1.45E-15 | 2.63E-13 |
| ISPD | -2.68309 | 0.0030399 | 0.0263728 | VIM | -1.944895 | 4.49E-10 | 1.78E-08 | MEGF6 | -7.442101 | 8.12E-15 | 1.37E-12 |
| MID1IP1 | -2.68371 | 0.0026617 | 0.0237039 | CCNG2 | -1.945934 | 3.26E-09 | 1.12E-07 | CD40LG | -7.711911 | 9.94E-13 | 1.25E-10 |
| TNFRSF11A | -2.706359 | 0.0018145 | 0.0174034 | BTG2 | -1.950004 | 4.71E-07 | 1.07E-05 | ANK1 | -7.715101 | 1.56E-13 | 2.20E-11 |
| LOC100506 | -2.723339 | 0.0011083 | 0.0115741 | ESD | -1.958001 | 0.0001032 | 0.0013589 | FSTL4 | -7.786685 | 1.38E-14 | 2.29E-12 |
| TNFRSF25 | -2.747712 | 2.98E-06 | 6.23E-05 | RPS6KA3 | -1.961929 | 1.56E-14 | 1.24E-12 | SLAMF1 | -7.85768 | 1.32E-15 | 2.43E-13 |
| COQ2 | -2.749088 | 0.0072645 | 0.0529121 | RPS8 | -1.969564 | 1.80E-22 | 4.14E-20 | MAL2 | -8.063223 | 2.98E-15 | 5.20E-13 |
| NFKBIA | -2.749409 | 1.73E-30 | 8.08E-28 | SAMSN1 | -2.006845 | 5.45E-05 | 0.0007774 | KRT1 | -8.286655 | 1.11E-20 | 3.66E-18 |
| FAM161A | -2.750769 | 0.0024245 | 0.0221071 | TGIF1 | -2.008368 | 1.71E-05 | 0.0002736 | NPDC1 | -8.758226 | 1.21E-26 | 6.84E-24 |
| MAST4 | -2.753949 | 1.21E-08 | 3.94E-07 | MAFG | -2.011399 | 0.0023567 | 0.020332 | HPGDS | -9.100707 | 1.17E-32 | 1.12E-29 |
| NPIPA3 | -2.769749 | 0.0089663 | 0.0625924 | RARG | -2.012061 | 0.0008545 | 0.0085331 | | | | |
| RASSF2 | -2.795956 | 5.53E-05 | 0.0008607 | ZBTB25 | -2.012844 | 0.000525 | 0.0056315 | | | | |
| C19orf60 | -2.79679 | 0.000123 | 0.0017625 | ADK | -2.017851 | 0.002212 | 0.0192447 | | | | |
| MAML3 | -2.798341 | 0.0046563 | 0.0374195 | AGPAT5 | -2.017935 | 0.0002479 | 0.0029461 | | | | |
| PDE8A | -2.801886 | 0.0001228 | 0.0017625 | NFKB1 | -2.021023 | 7.70E-09 | 2.42E-07 | | | | |

| | | | | | | | |
|---|---|---|---|---|---|---|---|
| DOCK9 | -2.829753 | 8.25E-05 | 0.0012281 | LEPROTL1 | -2.033779 | 2.03E-06 | 3.92E-05 |
| GRAP | -2.834538 | 0.0007943 | 0.0088349 | CD44 | -2.040034 | 7.34E-15 | 6.12E-13 |
| CYB5R1 | -2.836015 | 0.0090135 | 0.0627654 | RNF144A | -2.045019 | 0.0003742 | 0.004199 |
| CFAP36 | -2.848825 | 1.21E-09 | 4.76E-08 | GOLGA6L9 | -2.049639 | 0.0009903 | 0.0096595 |
| IMPDH2 | -2.852065 | 7.31E-06 | 0.0001425 | JUND | -2.052418 | 6.83E-14 | 4.83E-12 |
| CDC14A | -2.863278 | 7.28E-16 | 8.85E-14 | DGKA | -2.058491 | 4.32E-08 | 1.18E-06 |
| SERINC5 | -2.870434 | 0.0038381 | 0.0319128 | PLAGL1 | -2.075765 | 0.0087401 | 0.0605142 |
| TNFRSF18 | -2.877437 | 2.87E-06 | 6.05E-05 | C10orf128 | -2.076799 | 2.15E-06 | 4.10E-05 |
| PASK | -2.879692 | 4.74E-05 | 0.0007514 | CC2D2A | -2.079552 | 0.0032424 | 0.0266087 |
| RUNX2 | -2.925279 | 1.85E-08 | 5.84E-07 | EPHA4 | -2.083877 | 3.41E-05 | 0.0005161 |
| HIBADH | -2.929296 | 0.002277 | 0.0209903 | ZNF226 | -2.085408 | 0.0020154 | 0.0177855 |
| APP | -2.944887 | 0.001157 | 0.0119959 | ATP8B2 | -2.0874 | 3.24E-06 | 6.01E-05 |
| LRP12 | -2.945797 | 0.0043383 | 0.0352348 | PCYOX1L | -2.089668 | 3.52E-05 | 0.0005305 |
| S1PR1 | -2.947577 | 5.72E-08 | 1.66E-06 | ARHGAP5 | -2.09022 | 3.12E-07 | 7.27E-06 |
| FHL1 | -2.953731 | 0.0007564 | 0.0084922 | GPR155 | -2.105485 | 0.0001926 | 0.0023612 |
| PPP1R14B | -2.97087 | 0.0039151 | 0.0324569 | ANKRD28 | -2.113357 | 1.90E-07 | 4.59E-06 |
| FBXW8 | -2.997493 | 0.0029481 | 0.0256957 | RNF157 | -2.116528 | 0.0049962 | 0.0384775 |
| INPP5A | -3.005836 | 0.0006379 | 0.0073998 | PAICS | -2.116979 | 9.08E-07 | 1.93E-05 |
| FOS | -3.006169 | 3.63E-11 | 1.94E-09 | IL6ST | -2.118845 | 7.63E-06 | 0.0001308 |
| MPZL3 | -3.017342 | 1.41E-06 | 3.21E-05 | AMPD2 | -2.126285 | 0.0021465 | 0.0187628 |
| ZNF669 | -3.019387 | 0.0025438 | 0.0229462 | STARD4 | -2.141582 | 5.42E-08 | 1.47E-06 |
| E2F2 | -3.020693 | 0.0079822 | 0.0573432 | CORO2A | -2.14281 | 0.0006591 | 0.0068476 |
| TMEM123 | -3.028394 | 1.89E-17 | 2.89E-15 | RPL23P8 | -2.143505 | 0.0029109 | 0.0242979 |
| AKAP1 | -3.035143 | 0.0007781 | 0.0086891 | TFRC | -2.145537 | 0.000179 | 0.0022118 |
| JAG1 | -3.042292 | 0.001598 | 0.0156946 | JAG1 | -2.149537 | 0.0037462 | 0.030051 |
| CENPV | -3.055784 | 0.0002051 | 0.0027465 | CORO7 | -2.149612 | 0.0050014 | 0.0384855 |
| TOB1 | -3.065627 | 4.13E-07 | 1.03E-05 | YBX3 | -2.149665 | 0.0028609 | 0.0239672 |
| SCMH1 | -3.072306 | 0.0045966 | 0.0370464 | BCL2 | -2.153841 | 1.73E-07 | 4.20E-06 |
| NFKB2 | -3.073896 | 0.0033533 | 0.0285033 | LRRC8D | -2.154457 | 0.0001186 | 0.0015462 |
| TPST1 | -3.08253 | 0.000886 | 0.0096879 | LINC00341 | -2.157512 | 0.0007078 | 0.0072722 |
| TCF7 | -3.084178 | 1.07E-21 | 2.36E-19 | TNFRSF18 | -2.158284 | 2.88E-05 | 0.0004399 |
| SCML1 | -3.09421 | 7.70E-05 | 0.0011507 | PAG1 | -2.159054 | 7.82E-21 | 1.48E-18 |
| SPPL2B | -3.118784 | 0.0021997 | 0.0204351 | IMPDH2 | -2.16629 | 0.0015194 | 0.0139738 |
| DIP2C | -3.119487 | 0.0056097 | 0.043282 | TAF9B | -2.175961 | 6.01E-06 | 0.0001048 |
| AHR | -3.122624 | 6.76E-06 | 0.000134 | FAM117B | -2.186577 | 0.0004654 | 0.0050748 |
| NT5E | -3.122665 | 0.0071663 | 0.0525154 | C9orf89 | -2.188531 | 0.0010285 | 0.0099907 |
| NSUN4 | -3.167816 | 0.0021686 | 0.0202131 | STRBP | -2.190396 | 3.58E-07 | 8.31E-06 |
| ADCY3 | -3.16836 | 0.0055568 | 0.0431292 | KIAA1586 | -2.199633 | 0.0007365 | 0.0075166 |
| DGKH | -3.175403 | 3.47E-06 | 7.17E-05 | HABP4 | -2.233395 | 0.0011032 | 0.0105612 |
| LMO7 | -3.177636 | 0.0003068 | 0.0038681 | PCBD2 | -2.235063 | 0.001456 | 0.0134645 |
| GPM6A | -3.185768 | 0.0024144 | 0.0220395 | ANXA5 | -2.242159 | 9.36E-07 | 1.98E-05 |
| TMEM14C | -3.19547 | 0.0005113 | 0.0061168 | SYPL1 | -2.245871 | 5.28E-05 | 0.0007624 |
| GOLGA6L9 | -3.209626 | 8.93E-05 | 0.0013228 | RASSF2 | -2.273866 | 0.0015171 | 0.0139738 |
| LYL1 | -3.218788 | 0.0048279 | 0.0383942 | NFKBIA | -2.287127 | 3.37E-23 | 8.67E-21 |
| FAM24B | -3.234393 | 0.0027353 | 0.0241035 | SLC25A33 | -2.291359 | 0.0001466 | 0.0018611 |
| MIATNB | -3.239823 | 0.0009944 | 0.0106521 | MDFIC | -2.29159 | 1.61E-09 | 5.80E-08 |
| LINC00341 | -3.254366 | 0.0002059 | 0.0027516 | RARA | -2.292145 | 5.26E-05 | 0.0007605 |
| HNRNPLL | -3.27325 | 4.30E-05 | 0.0006927 | LYRM4 | -2.326256 | 0.0025968 | 0.0221311 |
| CYBRD1 | -3.325684 | 0.0015352 | 0.015186 | ZFAND1 | -2.329372 | 0.0005482 | 0.0058595 |
| GCSAM | -3.352024 | 0.0005871 | 0.0068856 | IL32 | -2.333399 | 2.08E-10 | 8.59E-09 |
| RARG | -3.366999 | 2.90E-05 | 0.0004914 | SNX9 | -2.335278 | 0.0002513 | 0.0029708 |
| GATA3 | -3.426042 | 1.94E-25 | 6.03E-23 | P2RY10 | -2.335704 | 0.0002979 | 0.003443 |
| KCNA3 | -3.443288 | 2.88E-16 | 3.84E-14 | RNF214 | -2.337161 | 0.0001166 | 0.0015248 |
| RTKN2 | -3.455773 | 1.28E-09 | 5.00E-08 | FAM118A | -2.378254 | 0.0088652 | 0.0611972 |

| | | | | | | | |
|---|---|---|---|---|---|---|---|
| C4orf32 | -3.5032 | 0.001351 | 0.0136864 | REEP3 | -2.384258 | 2.14E-05 | 0.000339 |
| SGK1 | -3.505665 | 0.0026549 | 0.0236919 | GALT | -2.396442 | 0.0016952 | 0.0153101 |
| PVT1 | -3.525202 | 0.0001421 | 0.0019953 | EPB41L2 | -2.396996 | 0.0004841 | 0.005265 |
| CNTN3 | -3.526059 | 0.0041254 | 0.0338489 | ZEB1 | -2.405647 | 2.81E-12 | 1.59E-10 |
| RNF207 | -3.554194 | 0.0011741 | 0.0121583 | CASK | -2.411969 | 1.80E-09 | 6.40E-08 |
| SSBP2 | -3.560995 | 8.68E-09 | 2.96E-07 | SNHG8 | -2.415869 | 8.97E-12 | 4.72E-10 |
| FARP1 | -3.58086 | 0.0005154 | 0.0061571 | SLC25A15 | -2.416644 | 0.0007705 | 0.0078107 |
| BIRC3 | -3.623255 | 6.32E-37 | 4.43E-34 | FLNB | -2.424139 | 1.32E-05 | 0.0002178 |
| TAF4B | -3.626787 | 0.0006789 | 0.007757 | TMEM123 | -2.445266 | 4.03E-13 | 2.57E-11 |
| NBEA | -3.634853 | 0.0001782 | 0.0024209 | SPPL2B | -2.461138 | 0.0036593 | 0.0295331 |
| ANTXR2 | -3.646757 | 1.62E-12 | 1.10E-10 | CSRNP2 | -2.467756 | 0.0004246 | 0.0046732 |
| GPR183 | -3.656975 | 7.48E-09 | 2.59E-07 | SVIL | -2.473758 | 9.53E-13 | 5.84E-11 |
| A2MP1 | -3.664144 | 0.0027307 | 0.0240888 | RAPGEF2 | -2.475304 | 6.63E-14 | 4.72E-12 |
| MTRNR2L9 | -3.678034 | 0.0054305 | 0.0423189 | DUSP1 | -2.505366 | 3.50E-15 | 3.11E-13 |
| SLC40A1 | -3.685944 | 0.0041273 | 0.0338489 | SLC35F2 | -2.50628 | 0.0010707 | 0.0103348 |
| ANO9 | -3.697036 | 0.0006446 | 0.0074569 | GUCY1A3 | -2.507621 | 1.15E-07 | 2.92E-06 |
| FOXP1-AS1 | -3.699379 | 0.0006414 | 0.0074297 | NUDCD1 | -2.51068 | 0.0098637 | 0.0665997 |
| CAPG | -3.711267 | 0.0001017 | 0.0014849 | BCAS4 | -2.517637 | 0.0066525 | 0.0486051 |
| HPGD | -3.740885 | 0.000136 | 0.0019222 | LMO7 | -2.517646 | 0.0006471 | 0.0067613 |
| NR4A1 | -3.748741 | 1.87E-06 | 4.16E-05 | NREP | -2.536857 | 8.00E-05 | 0.0010954 |
| SNX9 | -3.751194 | 9.94E-09 | 3.35E-07 | PPP2R3A | -2.537748 | 0.0028265 | 0.0237647 |
| ZNF546 | -3.764824 | 0.0009156 | 0.009931 | GAS5 | -2.541466 | 3.76E-15 | 3.31E-13 |
| PLP2 | -3.768695 | 1.41E-12 | 9.69E-11 | KCNMB3 | -2.554258 | 0.0063894 | 0.0470911 |
| CORO2A | -3.771396 | 3.30E-08 | 9.97E-07 | DENND5A | -2.55647 | 2.64E-05 | 0.0004052 |
| TNFRSF9 | -3.811082 | 3.18E-06 | 6.60E-05 | MZF1-AS1 | -2.562534 | 0.0020687 | 0.0181341 |
| NRIP3 | -3.833433 | 0.0010147 | 0.010814 | GRIP1 | -2.567747 | 0.0012232 | 0.011566 |
| CHRM3-AS | -3.844169 | 0.0008022 | 0.0089113 | MFGE8 | -2.582483 | 0.0030445 | 0.0251406 |
| ENO2 | -3.852259 | 5.15E-05 | 0.0008072 | PIM2 | -2.584916 | 2.34E-08 | 6.69E-07 |
| CADM1 | -3.868681 | 7.66E-05 | 0.001146 | RGPD6 | -2.590644 | 0.0004939 | 0.0053428 |
| TNFSF8 | -3.87107 | 3.15E-08 | 9.57E-07 | RASA4 | -2.596997 | 0.0001477 | 0.0018701 |
| CD40LG | -3.887107 | 0.000244 | 0.0031964 | SUFU | -2.613067 | 0.0051589 | 0.0395334 |
| NCKAP1 | -3.888467 | 4.60E-05 | 0.0007309 | SPDYA | -2.619467 | 0.0006494 | 0.0067777 |
| ACVR1B | -3.960839 | 9.19E-05 | 0.0013566 | CHCHD10 | -2.626527 | 9.13E-05 | 0.0012245 |
| SETD7 | -3.982579 | 2.14E-06 | 4.66E-05 | NT5DC3 | -2.641841 | 0.0083994 | 0.0585934 |
| PBX1 | -3.991189 | 3.20E-05 | 0.0005331 | TP53INP1 | -2.646086 | 6.65E-20 | 1.14E-17 |
| DUSP16 | -4.006126 | 6.32E-10 | 2.64E-08 | CENPV | -2.663078 | 0.000406 | 0.0045005 |
| SV2A | -4.01397 | 6.11E-05 | 0.0009382 | RGS1 | -2.664784 | 4.65E-06 | 8.21E-05 |
| RAB3GAP1 | -4.053846 | 1.97E-07 | 5.17E-06 | FRY | -2.681066 | 1.24E-06 | 2.52E-05 |
| CCDC167 | -4.054535 | 2.08E-09 | 7.84E-08 | MID1IP1 | -2.687706 | 0.0015782 | 0.0144363 |
| ROGDI | -4.071745 | 2.95E-05 | 0.0005003 | PRDM8 | -2.693924 | 0.0032527 | 0.0266701 |
| LOC100996 | -4.083017 | 0.0011053 | 0.0115738 | PASK | -2.704376 | 2.92E-06 | 5.44E-05 |
| 8-Sep | -4.087014 | 4.47E-05 | 0.0007185 | KCTD12 | -2.705979 | 0.0009606 | 0.0094194 |
| CKAP4 | -4.104948 | 0.0005483 | 0.0065128 | PHTF2 | -2.71643 | 1.35E-20 | 2.49E-18 |
| SCN2A | -4.108441 | 2.43E-05 | 0.0004239 | ASPH | -2.73745 | 0.0076889 | 0.0542503 |
| LINC00324 | -4.16487 | 0.0002307 | 0.0030415 | FMN1 | -2.740133 | 0.0042045 | 0.0330121 |
| ZNF460 | -4.183057 | 0.0011139 | 0.0116062 | PLXDC2 | -2.743891 | 7.03E-10 | 2.69E-08 |
| TLE2 | -4.198189 | 0.0024308 | 0.0221409 | AHCY | -2.762668 | 0.0003191 | 0.0036471 |
| NR1D1 | -4.209271 | 0.0002923 | 0.0036977 | FAM161A | -2.776168 | 0.000124 | 0.0016011 |
| ANTXRLP1 | -4.210488 | 1.90E-05 | 0.0003405 | LDHB | -2.782927 | 2.79E-14 | 2.12E-12 |
| AXIN2 | -4.224654 | 4.58E-05 | 0.0007294 | INADL | -2.787579 | 8.88E-10 | 3.29E-08 |
| TIAM1 | -4.251849 | 1.08E-08 | 3.62E-07 | KANK3 | -2.793852 | 0.0019157 | 0.0169702 |
| SMYD5 | -4.260044 | 0.0010138 | 0.010814 | TLCD2 | -2.799598 | 0.0010582 | 0.0102364 |
| STMN3 | -4.298639 | 5.14E-16 | 6.35E-14 | CDC14A | -2.801047 | 2.15E-20 | 3.90E-18 |
| HLF | -4.321729 | 1.33E-08 | 4.30E-07 | DLL1 | -2.804548 | 1.01E-06 | 2.13E-05 |

| | | | | | | | |
|---|---|---|---|---|---|---|---|
| NBPF15 | -4.334826 | 0.0062635 | 0.0472605 | EGR1 | -2.806981 | 0.0001436 | 0.0018314 |
| ZNF510 | -4.354841 | 0.0001405 | 0.0019762 | FAM24B | -2.817871 | 0.0005057 | 0.0054556 |
| TNFRSF10A | -4.371159 | 0.0003272 | 0.0041078 | DGKH | -2.82195 | 2.50E-08 | 7.09E-07 |
| ZC3H12D | -4.401175 | 0.000156 | 0.0021516 | PGAP1 | -2.831544 | 0.000171 | 0.0021374 |
| MBOAT2 | -4.404317 | 0.0017015 | 0.016558 | DISC1 | -2.836533 | 0.0003591 | 0.0040642 |
| PGBD1 | -4.428427 | 0.000112 | 0.0016228 | RAB33B | -2.841771 | 0.0011432 | 0.0108875 |
| CHPT1 | -4.431629 | 6.42E-05 | 0.0009799 | TNFRSF10B | -2.842104 | 5.35E-05 | 0.0007699 |
| PHLDA2 | -4.458455 | 0.0029085 | 0.0253902 | RAB3GAP1 | -2.844766 | 0.0002553 | 0.0030068 |
| BCL7A | -4.513122 | 1.01E-05 | 0.0001923 | FAM92A1 | -2.856267 | 0.0073296 | 0.0521521 |
| LEF1 | -4.566367 | 2.62E-18 | 4.41E-16 | FLT3LG | -2.870285 | 2.17E-11 | 1.09E-09 |
| ZNF347 | -4.592178 | 1.19E-05 | 0.0002203 | BMP3 | -2.870635 | 0.0070227 | 0.0506693 |
| PRDM8 | -4.609865 | 8.58E-06 | 0.0001649 | PLK3 | -2.874141 | 0.0017348 | 0.0155959 |
| PAWR | -4.653841 | 2.67E-05 | 0.0004608 | ANTXR2 | -2.889184 | 9.24E-07 | 1.96E-05 |
| TBC1D4 | -4.70538 | 3.71E-08 | 1.11E-06 | ROGDI | -2.901393 | 0.0003565 | 0.0040448 |
| IL17RB | -4.708988 | 1.49E-08 | 4.80E-07 | FAR2 | -2.909707 | 0.0079798 | 0.0560036 |
| SPINK2 | -4.718381 | 0.0003542 | 0.0044197 | MGAT5 | -2.913331 | 2.33E-18 | 3.17E-16 |
| CR2 | -4.735675 | 0.0018472 | 0.0175884 | PIM3 | -2.917801 | 4.42E-07 | 1.01E-05 |
| MSRB3 | -4.790241 | 0.0004515 | 0.0054855 | AHR | -2.926946 | 6.98E-13 | 4.33E-11 |
| KIAA0226L | -4.796228 | 0.0002685 | 0.0034579 | INPP5E | -2.930304 | 3.33E-06 | 6.11E-05 |
| P2RY1 | -4.822412 | 3.71E-05 | 0.0006112 | FOS | -2.931609 | 1.64E-12 | 9.65E-11 |
| GAB2 | -4.82759 | 4.51E-05 | 0.0007208 | RPS6KA6 | -2.940861 | 0.0059725 | 0.0446944 |
| DPP4 | -4.834099 | 0.0004618 | 0.0055731 | ARL4D | -2.952892 | 0.002902 | 0.0242453 |
| FSCN1 | -4.897617 | 6.24E-05 | 0.0009542 | PVT1 | -2.963245 | 0.0049939 | 0.0384775 |
| RPS6KA6 | -4.905203 | 2.82E-05 | 0.0004832 | TCF7 | -2.97423 | 2.49E-14 | 1.92E-12 |
| LOC101927 | -4.907584 | 0.000357 | 0.0044483 | CDR2 | -2.984734 | 1.18E-12 | 7.08E-11 |
| SIRPG | -4.940179 | 3.95E-07 | 9.90E-06 | C4orf32 | -3.007746 | 0.0015895 | 0.0144823 |
| P2RX1 | -5.009818 | 0.0002625 | 0.0033917 | SSBP2 | -3.013061 | 4.33E-06 | 7.76E-05 |
| ALS2CL | -5.025252 | 0.0003373 | 0.0042154 | IL4R | -3.032377 | 2.21E-12 | 1.28E-10 |
| EGR1 | -5.026988 | 1.03E-09 | 4.11E-08 | FOSB | -3.066008 | 6.72E-13 | 4.20E-11 |
| MBOAT1 | -5.089231 | 1.96E-08 | 6.13E-07 | PDE3B | -3.066798 | 0.0001909 | 0.0023432 |
| COL4A3 | -5.117779 | 1.04E-05 | 0.0001967 | CFAP36 | -3.069382 | 9.79E-20 | 1.62E-17 |
| TCEA3 | -5.187175 | 7.86E-10 | 3.24E-08 | BCAS1 | -3.073243 | 2.49E-05 | 0.0003861 |
| A1BG | -5.195935 | 0.0001861 | 0.0025194 | TIAM1 | -3.092701 | 3.75E-13 | 2.42E-11 |
| CRYBG3 | -5.212258 | 1.28E-09 | 5.00E-08 | DUSP16 | -3.134988 | 1.36E-16 | 1.43E-14 |
| SOCS3 | -5.260603 | 2.66E-14 | 2.56E-12 | MTRNR2L9 | -3.147055 | 0.0052263 | 0.03995 |
| FAM102A | -5.299588 | 2.87E-19 | 5.39E-17 | ETS2 | -3.150102 | 0.0002754 | 0.0032107 |
| CCR7 | -5.305331 | 1.69E-10 | 7.97E-09 | TMEM14C | -3.175217 | 2.64E-07 | 6.28E-06 |
| PAQR7 | -5.32469 | 1.07E-06 | 2.49E-05 | MAML3 | -3.176022 | 0.0028412 | 0.0238452 |
| SHE | -5.356795 | 0.0001217 | 0.0017505 | GPR183 | -3.1762 | 1.99E-09 | 7.05E-08 |
| ZNF10 | -5.432733 | 9.51E-05 | 0.0013995 | GATA3 | -3.20753 | 1.91E-29 | 6.78E-27 |
| NSG1 | -5.508524 | 4.78E-07 | 1.19E-05 | NBEA | -3.209339 | 7.77E-07 | 1.70E-05 |
| ZNF540 | -5.526681 | 2.97E-05 | 0.0005023 | PLP2 | -3.240108 | 5.40E-09 | 1.75E-07 |
| GADD45G | -5.551293 | 1.25E-12 | 8.82E-11 | KIF5C | -3.260836 | 1.09E-06 | 2.27E-05 |
| ABCA1 | -5.690531 | 1.75E-11 | 1.02E-09 | NGFRAP1 | -3.270132 | 0.0015323 | 0.0140582 |
| FOXO3B | -5.709041 | 2.04E-07 | 5.35E-06 | TESPA1 | -3.271133 | 5.20E-14 | 3.79E-12 |
| SPINT2 | -5.7412 | 4.01E-12 | 2.55E-10 | N4BP3 | -3.283458 | 0.0005613 | 0.0059653 |
| ZNF204P | -5.767731 | 7.72E-08 | 2.15E-06 | TCEAL4 | -3.287024 | 6.44E-05 | 0.000906 |
| CDCP1 | -5.790737 | 1.65E-06 | 3.76E-05 | SLC48A1 | -3.290745 | 6.57E-05 | 0.0009183 |
| PKIA | -5.791355 | 2.14E-14 | 2.14E-12 | GRAP | -3.31111 | 5.27E-06 | 9.23E-05 |
| HPGDS | -5.825961 | 4.91E-06 | 9.87E-05 | CD40LG | -3.342102 | 0.0028894 | 0.0241619 |
| KRT1 | -5.828313 | 1.31E-10 | 6.34E-09 | GNAQ | -3.363377 | 8.13E-17 | 9.07E-15 |
| PKIB | -5.862121 | 1.07E-10 | 5.36E-09 | MAST4 | -3.369485 | 1.31E-08 | 3.91E-07 |
| NKX3-1 | -5.883097 | 1.02E-06 | 2.41E-05 | NCDN | -3.385821 | 0.0019928 | 0.0176194 |
| VSIG1 | -5.90201 | 6.46E-08 | 1.84E-06 | SERINC5 | -3.399736 | 9.02E-20 | 1.52E-17 |

| | | | | | | | |
|---|---|---|---|---|---|---|---|
| NELL2 | -5.967084 | 6.44E-10 | 2.68E-08 | MPZL3 | -3.400978 | 2.22E-10 | 9.11E-09 |
| RGCC | -6.057354 | 2.03E-15 | 2.37E-13 | BACE2 | -3.410498 | 2.04E-05 | 0.0003237 |
| SCN1B | -6.101189 | 1.79E-07 | 4.77E-06 | TNFRSF9 | -3.433162 | 0.0004515 | 0.0049408 |
| FXYD7 | -6.106394 | 3.41E-14 | 3.08E-12 | EMR2 | -3.44261 | 1.38E-05 | 0.0002267 |
| TLCD2 | -6.151317 | 2.86E-12 | 1.86E-10 | 8-Sep | -3.449152 | 4.57E-06 | 8.10E-05 |
| LTB | -6.162951 | 4.89E-150 | 4.11E-146 | PHLDB3 | -3.457597 | 0.0042304 | 0.0331592 |
| COL9A2 | -6.209419 | 6.86E-09 | 2.41E-07 | C8orf82 | -3.463548 | 0.002995 | 0.0247867 |
| CSGALNAC | -6.221076 | 5.32E-07 | 1.31E-05 | AFF3 | -3.467639 | 0.0005575 | 0.0059369 |
| DNER | -6.221237 | 4.50E-08 | 1.32E-06 | NFIX | -3.472339 | 0.0009451 | 0.0092872 |
| LSR | -6.237735 | 1.30E-12 | 9.07E-11 | TBC1D4 | -3.479799 | 8.58E-07 | 1.85E-05 |
| LINC00892 | -6.359508 | 5.53E-08 | 1.61E-06 | FARP1 | -3.494316 | 0.0002493 | 0.0029507 |
| FSTL4 | -6.390947 | 7.41E-09 | 2.57E-07 | IFNGR2 | -3.504068 | 1.42E-05 | 0.0002314 |
| WDR86-AS | -6.408479 | 1.12E-08 | 3.73E-07 | A2MP1 | -3.505825 | 0.002266 | 0.0196408 |
| SLAMF1 | -6.458216 | 2.08E-09 | 7.84E-08 | STK36 | -3.517007 | 0.0001934 | 0.0023649 |
| HAPLN3 | -6.476789 | 2.70E-12 | 1.77E-10 | FZD6 | -3.526109 | 5.63E-05 | 0.0007972 |
| IL1RL1 | -6.536811 | 6.59E-12 | 4.01E-10 | ARL4A | -3.531194 | 1.64E-21 | 3.23E-19 |
| GATA3-AS1 | -6.594082 | 6.15E-10 | 2.60E-08 | UBE2Q2P2 | -3.554342 | 0.0020585 | 0.0180965 |
| ANK1 | -6.627571 | 1.15E-08 | 3.80E-07 | SOCS3 | -3.555204 | 1.70E-12 | 9.95E-11 |
| MAL2 | -6.667491 | 1.31E-09 | 5.08E-08 | TNFRSF25 | -3.568423 | 2.57E-09 | 8.91E-08 |
| PSD3 | -6.743195 | 1.22E-12 | 8.66E-11 | MTM1 | -3.58665 | 3.37E-05 | 0.000511 |
| RPS6KA2 | -6.863703 | 1.39E-14 | 1.40E-12 | LEF1 | -3.602775 | 1.07E-12 | 6.48E-11 |
| CD5 | -6.866762 | 6.31E-10 | 2.64E-08 | INPP5A | -3.606705 | 1.40E-05 | 0.0002301 |
| MEGF6 | -6.893583 | 3.06E-10 | 1.34E-08 | LSR | -3.611202 | 8.03E-08 | 2.10E-06 |
| IL6R | -6.914222 | 6.74E-24 | 1.89E-21 | HPGD | -3.619543 | 0.000761 | 0.007724 |
| KLF5 | -6.918254 | 4.31E-11 | 2.23E-09 | SCML1 | -3.619874 | 0.0076321 | 0.0539456 |
| MAL | -6.964931 | 7.57E-11 | 3.84E-09 | MRPL42P5 | -3.668897 | 0.0001716 | 0.0021408 |
| AQP3 | -6.989069 | 4.87E-12 | 3.05E-10 | FOXP1-AS1 | -3.687902 | 0.0009991 | 0.0097351 |
| BACE2 | -7.034063 | 6.33E-23 | 1.56E-20 | CYLC2 | -3.700909 | 0.000573 | 0.0060831 |
| FAAH2 | -7.058275 | 1.64E-11 | 9.65E-10 | TUBB2A | -3.717656 | 3.01E-05 | 0.000457 |
| TSPAN18 | -7.07871 | 2.82E-14 | 2.60E-12 | BCL7A | -3.728558 | 0.000128 | 0.0016478 |
| PLEKHN1 | -7.101676 | 1.04E-14 | 1.08E-12 | MTL5 | -3.759243 | 0.0022375 | 0.0194298 |
| LMNA | -7.123078 | 5.26E-14 | 4.55E-12 | CCDC86 | -3.795427 | 5.58E-11 | 2.64E-09 |
| NPDC1 | -7.157082 | 1.43E-17 | 2.22E-15 | DNER | -3.811172 | 0.0003911 | 0.004362 |
| TNFRSF19 | -7.268798 | 6.25E-12 | 3.83E-10 | LINC00324 | -3.849217 | 0.0003735 | 0.004196 |
| N4BP3 | -7.328897 | 4.42E-14 | 3.91E-12 | FAM134B | -3.857157 | 5.24E-07 | 1.17E-05 |
| CYP2E1 | -7.33878 | 1.35E-12 | 9.34E-11 | TPST1 | -3.867074 | 5.58E-05 | 0.0007917 |
| KIAA0125 | -7.453114 | 2.55E-11 | 1.43E-09 | PLEKHN1 | -3.867675 | 1.47E-05 | 0.0002388 |
| IL1R1 | -7.468221 | 1.18E-18 | 2.02E-16 | RUNX2 | -3.874429 | 8.39E-10 | 3.15E-08 |
| RCAN3 | -7.571585 | 5.40E-17 | 7.95E-15 | CUL7 | -3.886399 | 0.0005335 | 0.0057157 |
| IL23A | -7.590864 | 3.36E-14 | 3.07E-12 | GRASP | -3.888801 | 3.44E-06 | 6.27E-05 |
| IL7R | -7.685174 | 1.83E-10 | 8.25E-09 | SGK1 | -3.89188 | 1.98E-05 | 0.000316 |
| CD27 | -7.70286 | 1.12E-14 | 1.14E-12 | FXYD7 | -3.933541 | 1.89E-10 | 7.92E-09 |
| HTR1F | -7.889526 | 3.19E-16 | 4.19E-14 | RNF207 | -3.941196 | 4.03E-05 | 0.0005961 |
| CCR6 | -8.193609 | 2.87E-23 | 7.77E-21 | AQP3 | -3.951551 | 3.01E-05 | 0.000457 |
| CD28 | -8.208232 | 9.15E-19 | 1.64E-16 | CD84 | -3.953582 | 9.77E-16 | 9.42E-14 |
| KIT | -8.213345 | 9.48E-19 | 1.66E-16 | DIP2C | -3.95796 | 2.21E-05 | 0.0003475 |
| ITM2C | -8.233504 | 4.19E-28 | 1.71E-25 | NR4A1 | -3.998344 | 5.59E-05 | 0.0007917 |
| INPP4B | -8.29208 | 3.63E-34 | 2.03E-31 | PLD6 | -4.008686 | 4.64E-05 | 0.000678 |
| IL2RA | -8.397689 | 3.09E-22 | 7.02E-20 | SGTB | -4.088038 | 5.13E-09 | 1.68E-07 |
| GAP43 | -8.598172 | 3.05E-22 | 7.02E-20 | FAM102A | -4.09116 | 6.46E-13 | 4.07E-11 |
| CRIP2 | -8.794469 | 2.74E-20 | 5.47E-18 | APP | -4.11917 | 4.43E-06 | 7.91E-05 |
| ADAM19 | -9.717975 | 8.86E-55 | 1.49E-51 | KLF5 | -4.147806 | 4.91E-07 | 1.11E-05 |
| | | | | TNFSF8 | -4.155638 | 6.94E-12 | 3.69E-10 |
| | | | | LRP6 | -4.223327 | 0.0001394 | 0.0017794 |

| | | | | | | | |
|---|---|---|---|---|---|---|---|
| | | | | HDHD3 | -4.254771 | 9.58E-05 | 0.0012762 |
| | | | | SCN2A | -4.260041 | 7.50E-06 | 0.0001288 |
| | | | | MBOAT2 | -4.285003 | 0.0038038 | 0.0304333 |
| | | | | SETD7 | -4.299767 | 0.0006601 | 0.0068508 |
| | | | | SOX4 | -4.306897 | 7.31E-19 | 1.04E-16 |
| | | | | HK2 | -4.316839 | 7.20E-06 | 0.0001241 |
| | | | | PAWR | -4.357792 | 3.82E-05 | 0.0005687 |
| | | | | BIRC3 | -4.386341 | 1.79E-47 | 1.50E-44 |
| | | | | GCSAM | -4.399485 | 1.54E-05 | 0.0002498 |
| | | | | CFH | -4.452984 | 2.09E-06 | 4.00E-05 |
| | | | | STMN3 | -4.490404 | 7.39E-05 | 0.0010204 |
| | | | | HTR1F | -4.550853 | 5.57E-08 | 1.50E-06 |
| | | | | IL17RB | -4.560626 | 9.55E-08 | 2.48E-06 |
| | | | | ADCY3 | -4.575601 | 9.04E-07 | 1.93E-05 |
| | | | | NELL2 | -4.587518 | 1.35E-07 | 3.38E-06 |
| | | | | TRAT1 | -4.663026 | 3.64E-09 | 1.22E-07 |
| | | | | RGCC | -4.664145 | 1.14E-08 | 3.47E-07 |
| | | | | LINC01215 | -4.666638 | 1.15E-10 | 5.08E-09 |
| | | | | TNFRSF10A | -4.668642 | 9.84E-06 | 0.000165 |
| | | | | MYC | -4.684684 | 1.30E-06 | 2.65E-05 |
| | | | | SHE | -4.712878 | 9.80E-05 | 0.0012982 |
| | | | | TNFRSF11A | -4.746062 | 4.21E-11 | 2.02E-09 |
| | | | | COL9A2 | -4.781165 | 4.68E-07 | 1.07E-05 |
| | | | | CAPG | -4.781893 | 7.08E-09 | 2.24E-07 |
| | | | | ITM2C | -4.793534 | 0.0011256 | 0.0107532 |
| | | | | FSCN1 | -4.835573 | 1.12E-05 | 0.000185 |
| | | | | E2F2 | -4.849997 | 2.90E-06 | 5.42E-05 |
| | | | | CCR6 | -4.879278 | 0.0001639 | 0.0020554 |
| | | | | GPR132 | -4.895548 | 2.08E-08 | 6.00E-07 |
| | | | | CRIP2 | -4.920341 | 0.0001784 | 0.0022108 |
| | | | | A1BG | -5.003357 | 1.64E-05 | 0.0002644 |
| | | | | MBOAT1 | -5.026883 | 3.31E-09 | 1.13E-07 |
| | | | | THEMIS | -5.052624 | 6.84E-08 | 1.81E-06 |
| | | | | NRIP3 | -5.075753 | 1.61E-07 | 3.94E-06 |
| | | | | IL1RL1 | -5.087146 | 9.97E-11 | 4.48E-09 |
| | | | | GATA3-AS1 | -5.155315 | 5.66E-09 | 1.83E-07 |
| | | | | RTKN2 | -5.231141 | 1.27E-16 | 1.35E-14 |
| | | | | MEGF6 | -5.237968 | 8.45E-08 | 2.20E-06 |
| | | | | SV2A | -5.265041 | 5.48E-10 | 2.15E-08 |
| | | | | LMNA | -5.2751 | 5.05E-12 | 2.77E-10 |
| | | | | ZC3H12D | -5.27964 | 1.13E-06 | 2.34E-05 |
| | | | | CR2 | -5.334956 | 8.36E-05 | 0.001137 |
| | | | | PSD3 | -5.361235 | 3.34E-10 | 1.33E-08 |
| | | | | GADD45G | -5.365761 | 5.87E-10 | 2.28E-08 |
| | | | | PKIB | -5.388727 | 2.30E-08 | 6.59E-07 |
| | | | | CSGALNACT1 | -5.427718 | 5.24E-05 | 0.0007592 |
| | | | | KIAA0226L | -5.443779 | 6.19E-06 | 0.0001077 |
| | | | | VSIG1 | -5.448493 | 3.93E-09 | 1.31E-07 |
| | | | | RCAN3 | -5.517755 | 6.36E-18 | 7.74E-16 |
| | | | | SPINK2 | -5.543735 | 1.70E-06 | 3.36E-05 |
| | | | | TRABD2A | -5.721168 | 1.43E-11 | 7.29E-10 |
| | | | | LTB | -5.867817 | 2.76E-97 | 1.28E-93 |
| | | | | LOC1009962 | -5.897722 | 1.40E-06 | 2.83E-05 |
| | | | | ALS2CL | -5.919936 | 3.84E-06 | 6.94E-05 |

| | | | | | | | |
|---|---|---|---|---|---|---|---|
| | | | | ZNF667-AS1 | -5.934773 | 1.06E-07 | 2.72E-06 |
| | | | | HAPLN3 | -5.95995 | 1.53E-14 | 1.23E-12 |
| | | | | NT5E | -6.085549 | 4.25E-08 | 1.17E-06 |
| | | | | IL6R | -6.170638 | 7.59E-25 | 2.07E-22 |
| | | | | IL2RA | -6.187872 | 1.58E-23 | 4.18E-21 |
| | | | | GCNT4 | -6.250922 | 2.16E-07 | 5.15E-06 |
| | | | | IL7R | -6.260636 | 3.86E-89 | 8.94E-86 |
| | | | | DPP4 | -6.277795 | 5.21E-07 | 1.17E-05 |
| | | | | FERMT2 | -6.312234 | 1.58E-08 | 4.60E-07 |
| | | | | COL4A3 | -6.314059 | 3.64E-09 | 1.22E-07 |
| | | | | PKIA | -6.346284 | 1.16E-14 | 9.47E-13 |
| | | | | GAP43 | -6.364745 | 7.58E-23 | 1.85E-20 |
| | | | | RHOBTB1 | -6.526148 | 4.22E-08 | 1.16E-06 |
| | | | | P2RY1 | -6.534517 | 2.28E-08 | 6.56E-07 |
| | | | | IL1R1 | -6.543093 | 4.33E-18 | 5.50E-16 |
| | | | | KIT | -6.622989 | 4.84E-41 | 2.49E-38 |
| | | | | ABCA1 | -6.649653 | 1.17E-21 | 2.35E-19 |
| | | | | CD28 | -6.678824 | 3.27E-19 | 4.96E-17 |
| | | | | SCN1B | -6.918915 | 1.61E-10 | 6.85E-09 |
| | | | | MSRB3 | -6.931866 | 1.44E-08 | 4.24E-07 |
| | | | | INPP4B | -6.951916 | 7.85E-32 | 3.03E-29 |
| | | | | TOX2 | -6.994566 | 1.80E-10 | 7.63E-09 |
| | | | | SPINT2 | -7.024288 | 1.84E-22 | 4.14E-20 |
| | | | | AXIN2 | -7.045332 | 4.04E-11 | 1.95E-09 |
| | | | | CCR7 | -7.158078 | 5.34E-19 | 7.97E-17 |
| | | | | SRGAP3 | -7.176054 | 1.54E-11 | 7.83E-10 |
| | | | | LINC00892 | -7.176421 | 5.75E-11 | 2.70E-09 |
| | | | | FSTL4 | -7.194955 | 2.46E-12 | 1.41E-10 |
| | | | | WDR86-AS1 | -7.21156 | 5.20E-12 | 2.81E-10 |
| | | | | SLAMF1 | -7.256838 | 3.90E-13 | 2.51E-11 |
| | | | | GAB2 | -7.325827 | 1.48E-30 | 5.50E-28 |
| | | | | PTP4A3 | -7.407663 | 2.66E-12 | 1.51E-10 |
| | | | | ANK1 | -7.456137 | 1.11E-11 | 5.72E-10 |
| | | | | MAL2 | -7.473766 | 4.76E-13 | 3.02E-11 |
| | | | | RPS6KA2 | -7.508391 | 2.71E-19 | 4.24E-17 |
| | | | | CYP2E1 | -7.522182 | 5.15E-17 | 5.98E-15 |
| | | | | CD5 | -7.698975 | 2.90E-13 | 1.92E-11 |
| | | | | TNFRSF19 | -7.72637 | 5.29E-16 | 5.15E-14 |
| | | | | TSPAN18 | -7.77396 | 2.69E-19 | 4.24E-17 |
| | | | | SIRPG | -7.775703 | 5.88E-14 | 4.25E-12 |
| | | | | MAL | -7.785625 | 1.84E-14 | 1.43E-12 |
| | | | | FAAH2 | -7.881482 | 2.91E-15 | 2.61E-13 |
| | | | | HLF | -7.980051 | 1.95E-34 | 8.60E-32 |
| | | | | CD27 | -8.182757 | 1.88E-19 | 3.05E-17 |
| | | | | TCEA3 | -8.197722 | 5.83E-19 | 8.57E-17 |
| | | | | KIAA0125 | -8.26463 | 1.45E-14 | 1.18E-12 |
| | | | | KRT1 | -8.289733 | 6.38E-20 | 1.11E-17 |
| | | | | IL23A | -8.383312 | 2.99E-18 | 3.90E-16 |
| | | | | ANTXRLP1 | -8.498836 | 6.32E-15 | 5.37E-13 |
| | | | | NPDC1 | -8.507668 | 5.26E-22 | 1.13E-19 |
| | | | | HPGDS | -8.579528 | 1.08E-27 | 3.22E-25 |
| | | | | ADAM19 | -9.175876 | 1.46E-28 | 4.82E-26 |

| CD56negNK vs ILC | | | | | | | |
|---|---|---|---|---|---|---|---|
| Reactome pathways | # | # | expected | Fold Enrichment | +/- | raw P value | FDR |
| PECAM1 interactions | 11 | 4 | 0.27 | 14.84 | + | 3.58E-04 | 3.03E-02 |
| DAP12 signaling | 25 | 8 | 0.61 | 13.06 | + | 8.32E-07 | 1.58E-04 |
| Regulation of KIT signaling | 13 | 4 | 0.32 | 12.56 | + | 6.01E-04 | 4.29E-02 |
| GPVI-mediated activation cascade | 35 | 9 | 0.86 | 10.49 | + | 8.05E-07 | 1.67E-04 |
| Regulation of signaling by CBL | 21 | 5 | 0.51 | 9.72 | + | 3.33E-04 | 3.17E-02 |
| Interleukin-37 signaling | 21 | 5 | 0.51 | 9.72 | + | 3.33E-04 | 3.04E-02 |
| DAP12 interactions | 39 | 9 | 0.96 | 9.42 | + | 1.75E-06 | 2.85E-04 |
| RHO GTPases Activate WASPs and WAVEs | 35 | 7 | 0.86 | 8.16 | + | 5.62E-05 | 6.41E-03 |
| Interleukin-3, Interleukin-5 and GM-CSF signaling | 47 | 8 | 1.15 | 6.95 | + | 4.60E-05 | 5.53E-03 |
| Interferon gamma signaling | 91 | 14 | 2.23 | 6.28 | + | 2.11E-07 | 4.81E-05 |
| Generation of second messenger molecules | 40 | 6 | 0.98 | 6.12 | + | 7.57E-04 | 4.67E-02 |
| Costimulation by the CD28 family | 72 | 8 | 1.76 | 4.53 | + | 6.51E-04 | 4.37E-02 |
| Immunoregulatory interactions between a Lymphoid and a non-Lymphoid cell | 199 | 22 | 4.88 | 4.51 | + | 1.91E-08 | 5.45E-06 |
| Fcgamma receptor (FCGR) dependent phagocytosis | 149 | 16 | 3.65 | 4.38 | + | 2.34E-06 | 3.57E-04 |
| RAB GEFs exchange GTP for GDP on RABs | 89 | 9 | 2.18 | 4.13 | + | 5.71E-04 | 4.21E-02 |
| Platelet activation, signaling and aggregation | 259 | 26 | 6.35 | 4.1 | + | 6.18E-09 | 2.02E-06 |
| Rho GTPase cycle | 140 | 13 | 3.43 | 3.79 | + | 8.21E-05 | 8.53E-03 |
| TCR signaling | 124 | 11 | 3.04 | 3.62 | + | 4.10E-04 | 3.34E-02 |
| Regulation of actin dynamics for phagocytic cup formation | 125 | 11 | 3.06 | 3.59 | + | 4.36E-04 | 3.32E-02 |
| Neutrophil degranulation | 478 | 42 | 11.71 | 3.59 | + | 6.54E-12 | 3.73E-09 |
| Interferon Signaling | 196 | 17 | 4.8 | 3.54 | + | 1.58E-05 | 2.00E-03 |
| Cell surface interactions at the vascular wall | 199 | 16 | 4.88 | 3.28 | + | 6.48E-05 | 7.05E-03 |
| Adaptive Immune System | 820 | 61 | 20.1 | 3.04 | + | 8.56E-14 | 6.52E-11 |
| RAF/MAP kinase cascade | 232 | 16 | 5.69 | 2.81 | + | 3.35E-04 | 2.95E-02 |
| MAPK1/MAPK3 signaling | 237 | 16 | 5.81 | 2.75 | + | 4.18E-04 | 3.29E-02 |
| Hemostasis | 670 | 45 | 16.42 | 2.74 | + | 3.89E-09 | 1.48E-06 |
| Signaling by Interleukins | 447 | 30 | 10.95 | 2.74 | + | 1.68E-06 | 2.95E-04 |
| Innate Immune System | 1105 | 73 | 27.08 | 2.7 | + | 6.87E-14 | 7.84E-11 |
| Signaling by Rho GTPases | 417 | 27 | 10.22 | 2.64 | + | 1.02E-05 | 1.37E-03 |
| FLT3 Signaling | 250 | 16 | 6.13 | 2.61 | + | 7.20E-04 | 4.70E-02 |
| Immune System | 2159 | 138 | 52.91 | 2.61 | + | 2.72E-25 | 6.20E-22 |
| Signaling by Receptor Tyrosine Kinases | 457 | 29 | 11.2 | 2.59 | + | 6.93E-06 | 9.89E-04 |
| Cytokine Signaling in Immune system | 823 | 52 | 20.17 | 2.58 | + | 2.03E-09 | 9.29E-07 |
| MAPK family signaling cascades | 276 | 17 | 6.76 | 2.51 | + | 7.45E-04 | 4.73E-02 |
| Signal Transduction | 2730 | 99 | 66.9 | 1.48 | + | 8.99E-05 | 8.92E-03 |
| Unclassified | 10195 | 187 | 249.85 | 0.75 | - | 4.02E-08 | 1.02E-05 |
| Olfactory Signaling Pathway | 432 | 1 | 10.59 | 0.09 | - | 6.34E-04 | 4.39E-02 |
| CD56dimNK vs ILC | | | | | | | |
| Reactome pathways | # | # | expected | Fold Enrichment | +/- | raw P value | FDR |
| DAP12 signaling | 25 | 9 | 0.63 | 14.27 | + | 9.36E-08 | 2.67E-05 |
| Interleukin-2 signaling | 12 | 4 | 0.3 | 13.21 | + | 5.21E-04 | 4.76E-02 |
| Regulation of signaling by CBL | 21 | 7 | 0.53 | 13.21 | + | 3.97E-06 | 6.47E-04 |
| GPVI-mediated activation cascade | 35 | 10 | 0.88 | 11.33 | + | 1.10E-07 | 2.79E-05 |
| DAP12 interactions | 39 | 10 | 0.98 | 10.16 | + | 2.59E-07 | 5.38E-05 |
| CD28 co-stimulation | 31 | 6 | 0.78 | 7.67 | + | 2.62E-04 | 2.85E-02 |
| Interleukin-3, Interleukin-5 and GM-CSF signaling | 47 | 9 | 1.19 | 7.59 | + | 8.36E-06 | 1.27E-03 |
| Interleukin-2 family signaling | 44 | 7 | 1.11 | 6.31 | + | 2.38E-04 | 2.72E-02 |
| Costimulation by the CD28 family | 72 | 9 | 1.82 | 4.96 | + | 1.67E-04 | 2.01E-02 |

| | | | | | | | |
|---|---|---|---|---|---|---|---|
| Interferon gamma signaling | 91 | 11 | 2.3 | 4.79 | + | 4.26E-05 | 6.09E-03 |
| Immunoregulatory interactions between a Lymphoid and a non-Lymphoid cell | 199 | 22 | 5.02 | 4.38 | + | 3.15E-08 | 1.20E-05 |
| Platelet activation, signaling and aggregation | 259 | 28 | 6.53 | 4.29 | + | 6.57E-10 | 3.00E-07 |
| Neutrophil degranulation | 478 | 47 | 12.06 | 3.9 | + | 2.19E-14 | 2.50E-11 |
| Rho GTPase cycle | 140 | 12 | 3.53 | 3.4 | + | 3.91E-04 | 4.06E-02 |
| Signaling by Interleukins | 447 | 32 | 11.28 | 2.84 | + | 3.72E-07 | 7.09E-05 |
| Adaptive Immune System | 820 | 57 | 20.69 | 2.76 | + | 2.41E-11 | 1.38E-08 |
| Innate Immune System | 1105 | 73 | 27.88 | 2.62 | + | 3.67E-13 | 2.79E-10 |
| Immune System | 2159 | 139 | 54.46 | 2.55 | + | 1.88E-24 | 4.29E-21 |
| Hemostasis | 670 | 41 | 16.9 | 2.43 | + | 7.42E-07 | 1.30E-04 |
| Cytokine Signaling in Immune system | 823 | 50 | 20.76 | 2.41 | + | 3.40E-08 | 1.11E-05 |
| Signaling by Receptor Tyrosine Kinases | 457 | 27 | 11.53 | 2.34 | + | 7.97E-05 | 1.01E-02 |
| Membrane Trafficking | 625 | 34 | 15.77 | 2.16 | + | 5.34E-05 | 7.18E-03 |
| Vesicle-mediated transport | 726 | 35 | 18.31 | 1.91 | + | 4.54E-04 | 4.32E-02 |
| Unclassified | 10195 | 196 | 257.19 | 0.76 | - | 1.32E-07 | 3.01E-05 |
| Olfactory Signaling Pathway | 432 | 1 | 10.9 | 0.09 | - | 4.28E-04 | 4.25E-02 |
| **CD56hiNK vs ILC** | | | | | | | |
| Reactome pathways | # | # | expected | Fold Enrichment | +/- | raw P value | FDR |
| DAP12 signaling | 25 | 8 | 0.5 | 16 | + | 1.85E-07 | 6.02E-05 |
| DAP12 interactions | 39 | 9 | 0.78 | 11.54 | + | 3.35E-07 | 9.56E-05 |
| GPVI-mediated activation cascade | 35 | 6 | 0.7 | 8.57 | + | 1.38E-04 | 2.10E-02 |
| Interleukin-3, Interleukin-5 and GM-CSF signaling | 47 | 7 | 0.94 | 7.45 | + | 8.47E-05 | 1.49E-02 |
| Interferon gamma signaling | 91 | 10 | 1.82 | 5.49 | + | 2.96E-05 | 6.14E-03 |
| Immunoregulatory interactions between a Lymphoid and a non-Lymphoid cell | 199 | 21 | 3.98 | 5.28 | + | 2.66E-09 | 1.52E-06 |
| Cell surface interactions at the vascular wall | 199 | 14 | 3.98 | 3.52 | + | 8.54E-05 | 1.39E-02 |
| Neutrophil degranulation | 478 | 33 | 9.56 | 3.45 | + | 2.38E-09 | 1.81E-06 |
| Interferon Signaling | 196 | 13 | 3.92 | 3.32 | + | 2.61E-04 | 3.73E-02 |
| Adaptive Immune System | 820 | 45 | 16.4 | 2.74 | + | 2.75E-09 | 1.26E-06 |
| Signaling by Interleukins | 447 | 24 | 8.94 | 2.68 | + | 2.22E-05 | 5.07E-03 |
| Innate Immune System | 1105 | 55 | 22.1 | 2.49 | + | 1.25E-09 | 1.43E-06 |
| Immune System | 2159 | 105 | 43.18 | 2.43 | + | 2.04E-17 | 4.66E-14 |
| Cytokine Signaling in Immune system | 823 | 40 | 16.46 | 2.43 | + | 4.96E-07 | 1.26E-04 |
| Hemostasis | 670 | 30 | 13.4 | 2.24 | + | 6.93E-05 | 1.32E-02 |
| Unclassified | 10195 | 150 | 203.89 | 0.74 | - | 1.78E-07 | 6.76E-05 |
| Olfactory Signaling Pathway | 432 | 0 | 8.64 | < 0.01 | - | 3.15E-04 | 4.23E-02 |
| **CD56negNK vs ILC** | | | | | | | |
| Reactome pathways | # | # | expected | Fold Enrichment | +/- | raw P value | FDR |
| Peptide chain elongation | 89 | 77 | 2.44 | 31.54 | + | 1.78E-76 | 2.04E-73 |
| Viral mRNA Translation | 89 | 75 | 2.44 | 30.72 | + | 6.99E-74 | 3.99E-71 |
| Eukaryotic Translation Elongation | 94 | 79 | 2.58 | 30.64 | + | 8.68E-78 | 1.98E-74 |
| Selenocysteine synthesis | 93 | 75 | 2.55 | 29.4 | + | 6.98E-73 | 3.19E-70 |
| Eukaryotic Translation Termination | 93 | 75 | 2.55 | 29.4 | + | 6.98E-73 | 2.66E-70 |
| Nonsense Mediated Decay (NMD) independent of the Exon Junction Complex (EJC) | 95 | 75 | 2.61 | 28.78 | + | 2.14E-72 | 6.12E-70 |
| Formation of a pool of free 40S subunits | 101 | 78 | 2.77 | 28.15 | + | 8.75E-75 | 6.66E-72 |

| | | | | | | | |
|---|---|---|---|---|---|---|---|
| Response of EIF2AK4 (GCN2) to amino acid deficiency | 101 | 76 | 2.77 | 27.43 | + | 3.03E-72 | 6.93E-70 |
| L13a-mediated translational silencing of Ceruloplasmin expression | 111 | 78 | 3.05 | 25.62 | + | 1.69E-72 | 5.52E-70 |
| GTP hydrolysis and joining of the 60S ribosomal subunit | 112 | 78 | 3.07 | 25.39 | + | 2.80E-72 | 7.12E-70 |
| SRP-dependent cotranslational protein targeting to membrane | 112 | 75 | 3.07 | 24.41 | + | 1.49E-68 | 2.43E-66 |
| Eukaryotic Translation Initiation | 119 | 78 | 3.26 | 23.89 | + | 8.73E-71 | 1.81E-68 |
| Cap-dependent Translation Initiation | 119 | 78 | 3.26 | 23.89 | + | 8.73E-71 | 1.66E-68 |
| Selenoamino acid metabolism | 116 | 76 | 3.18 | 23.88 | + | 6.02E-69 | 1.06E-66 |
| Nonsense-Mediated Decay (NMD) | 115 | 75 | 3.15 | 23.77 | + | 6.33E-68 | 9.64E-66 |
| Nonsense Mediated Decay (NMD) enhanced by the Exon Junction Complex (EJC) | 115 | 75 | 3.15 | 23.77 | + | 6.33E-68 | 9.04E-66 |
| Formation of the ternary complex, and subsequently, the 43S complex | 51 | 32 | 1.4 | 22.87 | + | 5.14E-29 | 3.56E-27 |
| Influenza Viral RNA Transcription and Replication | 132 | 75 | 3.62 | 20.71 | + | 1.33E-64 | 1.79E-62 |
| Ribosomal scanning and start codon recognition | 58 | 32 | 1.59 | 20.11 | + | 1.07E-27 | 7.21E-26 |
| Translation initiation complex formation | 58 | 32 | 1.59 | 20.11 | + | 1.07E-27 | 7.01E-26 |
| Activation of the mRNA upon binding of the cap-binding complex and eIFs, and subsequent binding to 43S | 59 | 32 | 1.62 | 19.77 | + | 1.62E-27 | 9.97E-26 |
| Influenza Life Cycle | 141 | 75 | 3.87 | 19.39 | + | 5.50E-63 | 6.98E-61 |
| STAT5 Activation | 6 | 3 | 0.16 | 18.23 | + | 1.41E-03 | 3.88E-02 |
| Influenza Infection | 152 | 76 | 4.17 | 18.23 | + | 2.79E-62 | 3.35E-60 |
| Regulation of expression of SLITs and ROBOs | 169 | 75 | 4.64 | 16.18 | + | 1.78E-58 | 1.94E-56 |
| Major pathway of rRNA processing in the nucleolus and cytosol | 182 | 77 | 4.99 | 15.42 | + | 8.34E-59 | 9.53E-57 |
| rRNA processing in the nucleus and cytosol | 192 | 77 | 5.27 | 14.62 | + | 2.06E-57 | 2.14E-55 |
| Regulation by c-FLIP | 10 | 4 | 0.27 | 14.58 | + | 4.07E-04 | 1.29E-02 |
| Dimerization of procaspase-8 | 10 | 4 | 0.27 | 14.58 | + | 4.07E-04 | 1.27E-02 |
| CASP8 activity is inhibited | 10 | 4 | 0.27 | 14.58 | + | 4.07E-04 | 1.26E-02 |
| rRNA processing | 202 | 78 | 5.54 | 14.08 | + | 3.69E-57 | 3.66E-55 |
| Signaling by ROBO receptors | 215 | 78 | 5.9 | 13.22 | + | 1.68E-55 | 1.60E-53 |
| Regulation of necroptotic cell death | 17 | 6 | 0.47 | 12.87 | + | 2.42E-05 | 1.13E-03 |
| Regulated Necrosis | 19 | 6 | 0.52 | 11.51 | + | 4.05E-05 | 1.65E-03 |
| RIPK1-mediated regulated necrosis | 19 | 6 | 0.52 | 11.51 | + | 4.05E-05 | 1.62E-03 |
| TNF receptor superfamily (TNFSF) members mediating non-canonical NF-kB pathway | 16 | 5 | 0.44 | 11.39 | + | 1.90E-04 | 6.49E-03 |
| Translation | 293 | 83 | 8.04 | 10.33 | + | 5.12E-52 | 4.68E-50 |
| Caspase activation via Death Receptors in the presence of ligand | 15 | 4 | 0.41 | 9.72 | + | 1.42E-03 | 3.85E-02 |
| The NLRP3 inflammasome | 15 | 4 | 0.41 | 9.72 | + | 1.42E-03 | 3.81E-02 |
| p75NTR signals via NF-kB | 16 | 4 | 0.44 | 9.11 | + | 1.73E-03 | 4.60E-02 |
| Inflammasomes | 20 | 5 | 0.55 | 9.11 | + | 4.55E-04 | 1.39E-02 |
| Metabolism of amino acids and derivatives | 367 | 79 | 10.07 | 7.85 | + | 5.61E-42 | 4.57E-40 |
| Diseases of Immune System | 24 | 5 | 0.66 | 7.59 | + | 9.31E-04 | 2.69E-02 |
| TRAF6 mediated NF-kB activation | 24 | 5 | 0.66 | 7.59 | + | 9.31E-04 | 2.66E-02 |
| Diseases associated with the TLR signaling cascade | 24 | 5 | 0.66 | 7.59 | + | 9.31E-04 | 2.63E-02 |
| Infectious disease | 465 | 88 | 12.76 | 6.9 | + | 6.73E-43 | 5.70E-41 |

| | | | | | | | |
|---|---|---|---|---|---|---|---|
| TAK1 activates NFkB by phosphorylation and activation of IKKs complex | 32 | 6 | 0.88 | 6.83 | + | 4.71E-04 | 1.42E-02 |
| Axon guidance | 548 | 97 | 15.03 | 6.45 | + | 3.83E-45 | 3.36E-43 |
| Cellular responses to stress | 548 | 92 | 15.03 | 6.12 | + | 4.23E-41 | 3.33E-39 |
| TNF signaling | 42 | 7 | 1.15 | 6.08 | + | 3.03E-04 | 9.74E-03 |
| Nucleotide-binding domain, leucine rich repeat containing receptor (NLR) signaling pathways | 54 | 9 | 1.48 | 6.08 | + | 4.24E-05 | 1.67E-03 |
| Cellular responses to external stimuli | 562 | 92 | 15.42 | 5.97 | + | 2.61E-40 | 1.99E-38 |
| TNFR2 non-canonical NF-kB pathway | 99 | 13 | 2.72 | 4.79 | + | 9.28E-06 | 4.71E-04 |
| MyD88 cascade initiated on plasma membrane | 84 | 11 | 2.3 | 4.77 | + | 4.59E-05 | 1.78E-03 |
| Toll Like Receptor 5 (TLR5) Cascade | 84 | 11 | 2.3 | 4.77 | + | 4.59E-05 | 1.75E-03 |
| Toll Like Receptor 10 (TLR10) Cascade | 84 | 11 | 2.3 | 4.77 | + | 4.59E-05 | 1.72E-03 |
| Toll Like Receptor 3 (TLR3) Cascade | 92 | 12 | 2.52 | 4.75 | + | 2.17E-05 | 1.05E-03 |
| MyD88:MAL(TIRAP) cascade initiated on plasma membrane | 94 | 12 | 2.58 | 4.65 | + | 2.63E-05 | 1.20E-03 |
| Toll Like Receptor TLR6:TLR2 Cascade | 94 | 12 | 2.58 | 4.65 | + | 2.63E-05 | 1.18E-03 |
| Interleukin-17 signaling | 71 | 9 | 1.95 | 4.62 | + | 2.78E-04 | 9.08E-03 |
| Interleukin-4 and Interleukin-13 signaling | 111 | 14 | 3.05 | 4.6 | + | 6.45E-06 | 3.35E-04 |
| TRIF(TICAM1)-mediated TLR4 signaling | 96 | 12 | 2.63 | 4.56 | + | 3.17E-05 | 1.39E-03 |
| MyD88-independent TLR4 cascade | 96 | 12 | 2.63 | 4.56 | + | 3.17E-05 | 1.37E-03 |
| Toll Like Receptor TLR1:TLR2 Cascade | 97 | 12 | 2.66 | 4.51 | + | 3.48E-05 | 1.47E-03 |
| Toll Like Receptor 2 (TLR2) Cascade | 97 | 12 | 2.66 | 4.51 | + | 3.48E-05 | 1.45E-03 |
| Metabolism of RNA | 661 | 81 | 18.13 | 4.47 | + | 1.28E-27 | 8.10E-26 |
| TRAF6 mediated induction of NFkB and MAP kinases upon TLR7/8 or 9 activation | 90 | 11 | 2.47 | 4.46 | + | 8.09E-05 | 2.98E-03 |
| Toll Like Receptor 7/8 (TLR7/8) Cascade | 91 | 11 | 2.5 | 4.41 | + | 8.86E-05 | 3.21E-03 |
| MyD88 dependent cascade initiated on endosome | 91 | 11 | 2.5 | 4.41 | + | 8.86E-05 | 3.16E-03 |
| DDX58/IFIH1-mediated induction of interferon-alpha/beta | 77 | 9 | 2.11 | 4.26 | + | 4.79E-04 | 1.42E-02 |
| Toll Like Receptor 9 (TLR9) Cascade | 95 | 11 | 2.61 | 4.22 | + | 1.26E-04 | 4.41E-03 |
| Toll Like Receptor 4 (TLR4) Cascade | 126 | 14 | 3.46 | 4.05 | + | 2.38E-05 | 1.13E-03 |
| Death Receptor Signalling | 141 | 15 | 3.87 | 3.88 | + | 1.96E-05 | 9.75E-04 |
| Developmental Biology | 1056 | 109 | 28.97 | 3.76 | + | 2.57E-31 | 1.89E-29 |
| Disease | 1127 | 111 | 30.92 | 3.59 | + | 2.85E-30 | 2.03E-28 |
| L1CAM interactions | 118 | 11 | 3.24 | 3.4 | + | 6.94E-04 | 2.03E-02 |
| Toll-like Receptor Cascades | 152 | 14 | 4.17 | 3.36 | + | 1.53E-04 | 5.30E-03 |
| Signaling by Interleukins | 447 | 36 | 12.26 | 2.94 | + | 3.41E-08 | 1.85E-06 |
| Cytokine Signaling in Immune system | 823 | 60 | 22.58 | 2.66 | + | 3.30E-11 | 1.84E-09 |
| Metabolism of proteins | 1978 | 125 | 54.26 | 2.3 | + | 5.13E-18 | 3.08E-16 |
| Metabolism | 2079 | 122 | 57.03 | 2.14 | + | 4.11E-15 | 2.41E-13 |
| Hemostasis | 670 | 33 | 18.38 | 1.8 | + | 1.78E-03 | 4.68E-02 |
| Immune System | 2159 | 97 | 59.23 | 1.64 | + | 2.24E-06 | 1.19E-04 |
| Signal Transduction | 2730 | 103 | 74.89 | 1.38 | + | 1.10E-03 | 3.06E-02 |
| Unclassified | 10195 | 198 | 279.68 | 0.71 | - | 1.19E-11 | 6.77E-10 |
| G alpha (s) signalling events | 575 | 3 | 15.77 | 0.19 | - | 2.03E-04 | 6.81E-03 |
| Olfactory Signaling Pathway | 432 | 1 | 11.85 | 0.08 | - | 2.09E-04 | 6.93E-03 |

**CD56dimNK vs ILC**

| Reactome pathways | # | # | expected | Fold Enrichment | +/- | raw P value | FDR |
|---|---|---|---|---|---|---|---|
| Peptide chain elongation | 89 | 72 | 2.88 | 25.03 | + | 6.66E-65 | 7.61E-62 |
| Eukaryotic Translation Elongation | 94 | 74 | 3.04 | 24.35 | + | 4.29E-66 | 9.79E-63 |
| Viral mRNA Translation | 89 | 70 | 2.88 | 24.33 | + | 1.66E-62 | 5.41E-60 |
| Eukaryotic Translation Termination | 93 | 71 | 3.01 | 23.62 | + | 9.39E-63 | 3.57E-60 |

| | | | | | | | |
|---|---|---|---|---|---|---|---|
| Nonsense Mediated Decay (NMD) independent of the Exon Junction Complex (EJC) | 95 | 72 | 3.07 | 23.45 | + | 1.78E-63 | 1.02E-60 |
| Selenocysteine synthesis | 93 | 70 | 3.01 | 23.29 | + | 1.44E-61 | 2.74E-59 |
| Formation of a pool of free 40S subunits | 101 | 74 | 3.26 | 22.67 | + | 1.81E-64 | 1.38E-61 |
| Response of EIF2AK4 (GCN2) to amino acid deficiency | 101 | 70 | 3.26 | 21.44 | + | 8.71E-60 | 1.53E-57 |
| L13a-mediated translational silencing of Ceruloplasmin expression | 111 | 75 | 3.59 | 20.9 | + | 1.86E-63 | 8.52E-61 |
| GTP hydrolysis and joining of the 60S ribosomal subunit | 112 | 74 | 3.62 | 20.44 | + | 4.32E-62 | 1.10E-59 |
| Selenoamino acid metabolism | 116 | 75 | 3.75 | 20 | + | 2.03E-62 | 5.80E-60 |
| SRP-dependent cotranslational protein targeting to membrane | 112 | 71 | 3.62 | 19.61 | + | 1.18E-58 | 1.69E-56 |
| Eukaryotic Translation Initiation | 119 | 75 | 3.85 | 19.5 | + | 8.16E-62 | 1.86E-59 |
| Cap-dependent Translation Initiation | 119 | 75 | 3.85 | 19.5 | + | 8.16E-62 | 1.70E-59 |
| Nonsense-Mediated Decay (NMD) | 115 | 72 | 3.72 | 19.37 | + | 3.41E-59 | 5.57E-57 |
| Nonsense Mediated Decay (NMD) enhanced by the Exon Junction Complex (EJC) | 115 | 72 | 3.72 | 19.37 | + | 3.41E-59 | 5.20E-57 |
| Formation of the ternary complex, and subsequently, the 43S complex | 51 | 31 | 1.65 | 18.8 | + | 9.82E-26 | 6.60E-24 |
| CREB phosphorylation | 7 | 4 | 0.23 | 17.68 | + | 2.64E-04 | 8.37E-03 |
| Translation initiation complex formation | 58 | 32 | 1.87 | 17.07 | + | 1.54E-25 | 1.01E-23 |
| Influenza Viral RNA Transcription and Replication | 132 | 72 | 4.27 | 16.87 | + | 5.11E-56 | 6.87E-54 |
| Activation of the mRNA upon binding of the cap-binding complex and eIFs, and subsequent binding to 43S | 59 | 32 | 1.91 | 16.78 | + | 2.31E-25 | 1.47E-23 |
| Ribosomal scanning and start codon recognition | 58 | 31 | 1.87 | 16.53 | + | 1.83E-24 | 1.13E-22 |
| Influenza Life Cycle | 141 | 72 | 4.56 | 15.8 | + | 1.78E-54 | 2.25E-52 |
| Binding of TCF/LEF:CTNNB1 to target gene promoters | 8 | 4 | 0.26 | 15.47 | + | 3.86E-04 | 1.21E-02 |
| Influenza Infection | 152 | 73 | 4.91 | 14.86 | + | 9.10E-54 | 1.09E-51 |
| Regulation of expression of SLITs and ROBOs | 169 | 73 | 5.46 | 13.36 | + | 3.26E-51 | 3.72E-49 |
| Major pathway of rRNA processing in the nucleolus and cytosol | 182 | 73 | 5.88 | 12.41 | + | 2.06E-49 | 2.14E-47 |
| Regulation by c-FLIP | 10 | 4 | 0.32 | 12.37 | + | 7.42E-04 | 2.20E-02 |
| Dimerization of procaspase-8 | 10 | 4 | 0.32 | 12.37 | + | 7.42E-04 | 2.17E-02 |
| CASP8 activity is inhibited | 10 | 4 | 0.32 | 12.37 | + | 7.42E-04 | 2.15E-02 |
| rRNA processing in the nucleus and cytosol | 192 | 73 | 6.21 | 11.76 | + | 4.16E-48 | 3.96E-46 |
| rRNA processing | 202 | 73 | 6.53 | 11.18 | + | 7.29E-47 | 6.66E-45 |
| Signaling by ROBO receptors | 215 | 77 | 6.95 | 11.08 | + | 3.44E-49 | 3.41E-47 |
| Regulation of necroptotic cell death | 17 | 6 | 0.55 | 10.92 | + | 5.90E-05 | 2.17E-03 |
| Regulated Necrosis | 19 | 6 | 0.61 | 9.77 | + | 9.81E-05 | 3.34E-03 |
| RIPK1-mediated regulated necrosis | 19 | 6 | 0.61 | 9.77 | + | 9.81E-05 | 3.30E-03 |
| Translation | 293 | 87 | 9.47 | 9.19 | + | 4.39E-50 | 4.78E-48 |
| TP53 regulates transcription of additional cell cycle genes whose exact role in the p53 pathway remain uncertain | 21 | 5 | 0.68 | 7.37 | + | 1.13E-03 | 3.11E-02 |
| MAPK targets/ Nuclear events mediated by MAP kinases | 31 | 7 | 1 | 6.99 | + | 1.55E-04 | 4.98E-03 |
| Metabolism of amino acids and derivatives | 367 | 80 | 11.86 | 6.74 | + | 1.11E-37 | 9.36E-36 |
| TNFR1-induced NFkappaB signaling pathway | 30 | 6 | 0.97 | 6.19 | + | 8.05E-04 | 2.27E-02 |
| Infectious disease | 465 | 88 | 15.03 | 5.85 | + | 2.77E-37 | 2.26E-35 |
| Regulation of TNFR1 signaling | 32 | 6 | 1.03 | 5.8 | + | 1.08E-03 | 3.02E-02 |
| Axon guidance | 548 | 102 | 17.71 | 5.76 | + | 1.02E-42 | 9.00E-41 |
| Signaling by NOTCH2 | 33 | 6 | 1.07 | 5.62 | + | 1.25E-03 | 3.39E-02 |
| Cellular responses to stress | 548 | 92 | 17.71 | 5.19 | + | 2.46E-35 | 1.94E-33 |

| | | | | | | | |
|---|---|---|---|---|---|---|---|
| TNF signaling | 42 | 7 | 1.36 | 5.16 | + | 7.83E-04 | 2.24E-02 |
| Nucleotide-binding domain, leucine rich repeat containing receptor (NLR) signaling pathways | 54 | 9 | 1.75 | 5.16 | + | 1.43E-04 | 4.68E-03 |
| Cellular responses to external stimuli | 562 | 92 | 18.17 | 5.06 | + | 1.42E-34 | 1.08E-32 |
| MAP kinase activation | 63 | 10 | 2.04 | 4.91 | + | 8.89E-05 | 3.08E-03 |
| Interleukin-17 signaling | 71 | 11 | 2.3 | 4.79 | + | 4.86E-05 | 1.82E-03 |
| MyD88 cascade initiated on plasma membrane | 84 | 13 | 2.72 | 4.79 | + | 1.03E-05 | 4.81E-04 |
| Toll Like Receptor 5 (TLR5) Cascade | 84 | 13 | 2.72 | 4.79 | + | 1.03E-05 | 4.71E-04 |
| Toll Like Receptor 10 (TLR10) Cascade | 84 | 13 | 2.72 | 4.79 | + | 1.03E-05 | 4.62E-04 |
| MyD88:MAL(TIRAP) cascade initiated on plasma membrane | 94 | 14 | 3.04 | 4.61 | + | 7.04E-06 | 3.58E-04 |
| Toll Like Receptor TLR6:TLR2 Cascade | 94 | 14 | 3.04 | 4.61 | + | 7.04E-06 | 3.50E-04 |
| Recycling pathway of L1 | 47 | 7 | 1.52 | 4.61 | + | 1.41E-03 | 3.75E-02 |
| TRAF6 mediated induction of NFkB and MAP kinases upon TLR7/8 or 9 activation | 90 | 13 | 2.91 | 4.47 | + | 1.99E-05 | 8.76E-04 |
| Toll Like Receptor TLR1:TLR2 Cascade | 97 | 14 | 3.14 | 4.47 | + | 9.72E-06 | 4.72E-04 |
| Toll Like Receptor 2 (TLR2) Cascade | 97 | 14 | 3.14 | 4.47 | + | 9.72E-06 | 4.63E-04 |
| Interleukin-4 and Interleukin-13 signaling | 111 | 16 | 3.59 | 4.46 | + | 2.32E-06 | 1.23E-04 |
| Toll Like Receptor 7/8 (TLR7/8) Cascade | 91 | 13 | 2.94 | 4.42 | + | 2.22E-05 | 9.55E-04 |
| MyD88 dependent cascade initiated on endosome | 91 | 13 | 2.94 | 4.42 | + | 2.22E-05 | 9.37E-04 |
| TP53 Regulates Transcription of Cell Cycle Genes | 49 | 7 | 1.58 | 4.42 | + | 1.75E-03 | 4.60E-02 |
| Toll Like Receptor 3 (TLR3) Cascade | 92 | 13 | 2.97 | 4.37 | + | 2.46E-05 | 1.02E-03 |
| Toll Like Receptor 9 (TLR9) Cascade | 95 | 13 | 3.07 | 4.23 | + | 3.33E-05 | 1.33E-03 |
| TRIF(TICAM1)-mediated TLR4 signaling | 96 | 13 | 3.1 | 4.19 | + | 3.67E-05 | 1.45E-03 |
| MyD88-independent TLR4 cascade | 96 | 13 | 3.1 | 4.19 | + | 3.67E-05 | 1.42E-03 |
| Metabolism of RNA | 661 | 87 | 21.37 | 4.07 | + | 1.37E-26 | 9.47E-25 |
| Toll Like Receptor 4 (TLR4) Cascade | 126 | 15 | 4.07 | 3.68 | + | 3.72E-05 | 1.42E-03 |
| L1CAM interactions | 118 | 14 | 3.81 | 3.67 | + | 6.93E-05 | 2.51E-03 |
| TNFR2 non-canonical NF-kB pathway | 99 | 11 | 3.2 | 3.44 | + | 6.73E-04 | 2.02E-02 |
| Developmental Biology | 1056 | 116 | 34.13 | 3.4 | + | 4.80E-29 | 3.54E-27 |
| Death Receptor Signalling | 141 | 15 | 4.56 | 3.29 | + | 1.19E-04 | 3.94E-03 |
| Toll-like Receptor Cascades | 152 | 16 | 4.91 | 3.26 | + | 8.00E-05 | 2.85E-03 |
| Disease | 1127 | 116 | 36.43 | 3.18 | + | 8.85E-27 | 6.32E-25 |
| Signaling by Interleukins | 447 | 42 | 14.45 | 2.91 | + | 3.93E-09 | 2.14E-07 |
| Cytokine Signaling in Immune system | 823 | 70 | 26.6 | 2.63 | + | 1.78E-12 | 9.93E-11 |
| MAPK family signaling cascades | 276 | 21 | 8.92 | 2.35 | + | 5.47E-04 | 1.66E-02 |
| Metabolism of proteins | 1978 | 143 | 63.94 | 2.24 | + | 3.84E-19 | 2.31E-17 |
| Metabolism | 2079 | 138 | 67.2 | 2.05 | + | 2.01E-15 | 1.18E-13 |
| Immune System | 2159 | 110 | 69.79 | 1.58 | + | 2.92E-06 | 1.52E-04 |
| Signal Transduction | 2730 | 125 | 88.25 | 1.42 | + | 8.31E-05 | 2.92E-03 |
| Unclassified | 10195 | 232 | 329.55 | 0.7 | - | 9.59E-14 | 5.47E-12 |
| G alpha (s) signalling events | 575 | 5 | 18.59 | 0.27 | - | 3.95E-04 | 1.22E-02 |
| DNA Repair | 309 | 1 | 9.99 | 0.1 | - | 1.36E-03 | 3.67E-02 |
| Olfactory Signaling Pathway | 432 | 1 | 13.96 | 0.07 | - | 3.24E-05 | 1.32E-03 |

**CD56hiNK vs ILC**

| Reactome pathways | # | # | expected | Fold Enrichment | +/- | raw P value | FDR |
|---|---|---|---|---|---|---|---|
| Signaling by SCF-KIT | 39 | 7 | 0.85 | 8.19 | + | 5.18E-05 | 1.69E-02 |
| Interleukin-4 and Interleukin-13 signaling | 111 | 11 | 2.43 | 4.52 | + | 6.49E-05 | 1.85E-02 |
| Platelet activation, signaling and aggregation | 259 | 18 | 5.68 | 3.17 | + | 3.32E-05 | 1.26E-02 |
| Signaling by Interleukins | 447 | 31 | 9.8 | 3.16 | + | 5.30E-08 | 4.04E-05 |
| Cytokine Signaling in Immune system | 823 | 46 | 18.04 | 2.55 | + | 2.21E-08 | 2.52E-05 |

| | | | | | | | |
|---|---|---|---|---|---|---|---|
| Hemostasis | 670 | 32 | 14.68 | 2.18 | + | 8.35E-05 | 2.12E-02 |
| Immune System | 2159 | 90 | 47.32 | 1.9 | + | 4.78E-09 | 1.09E-05 |
| Signal Transduction | 2730 | 94 | 59.83 | 1.57 | + | 1.34E-05 | 6.11E-03 |
| Unclassified | 10195 | 171 | 223.45 | 0.77 | - | 1.27E-06 | 7.26E-04 |

| Supplemental Table 4 \| mouse gene expression | | | | | | | |
|---|---|---|---|---|---|---|---|
| | | | | Normalized counts | | | |
| FileID | Cell Type | GEO number | Tissue | Areg | Gata3 | Rorc | Tbx21 |
| GSM2056340_sample_49_RNA_LV_ILC1_rpt1 | ILC1 | GSE77695 | Liver | 2.96 | 788.27 | 33.77 | 2437.81 |
| GSM2056341_sample_50_RNA_LV_ILC1_rpt2 | ILC1 | GSE77695 | Liver | 3.51 | 763.16 | 22.34 | 2746.61 |
| GSM3208792_SNW7_ILC1GSM3208804_WT_1 | ILC1 | GSE116092 | Small_Intestine | 450.20 | 845.29 | 3023.69 | 2607.32 |
| GSM3208793_SNW8_ILC1GSM3208804_WT_2 | ILC1 | GSE116092 | Small_Intestine | 733.38 | 955.68 | 2472.37 | 2057.07 |
| GSM3208794_SNW9_ILC1GSM3208804_WT_3 | ILC1 | GSE116092 | Small_Intestine | 487.08 | 452.87 | 2724.12 | 940.29 |
| GSM3208820_11_GGACCC_ILC1_SILP_3 | ILC1 | GSE116092 | Small_Intestine | 1212.38 | 781.12 | 182.04 | 1518.78 |
| GSM3208824_15_GGATGT_ILC1_SILP_4 | ILC1 | GSE116092 | Small_Intestine | 877.24 | 572.30 | 377.95 | 1557.07 |
| GSM2056342_sample_51_RNA_GT_ILC2_rpt1 | ILC2 | GSE77695 | SI_lamina_propria | 5644.13 | 15088.12 | 148.94 | 17.06 |
| GSM2056343_sample_52_RNA_GT_ILC2_rpt2 | ILC2 | GSE77695 | SI_lamina_propria | 7671.94 | 13926.24 | 170.35 | 21.02 |
| GSM2056350_sample_59_RNA_GT_ILC2_IL33_IL25_rpt1 | ILC2 | GSE77695 | SI_lamina_propria | 1052.80 | 5886.53 | 19.24 | 9.59 |
| GSM2056351_sample_60_RNA_GT_ILC2_IL33_IL25_rpt2 | ILC2 | GSE77695 | SI_lamina_propria | 2423.93 | 6187.70 | 18.40 | 10.57 |
| GSM2932539_ILC2.SI_1_RNA-seq | ILC2 | GSE109125 | Small_Intestine | 4620.95 | 25104.32 | 178.28 | 34.33 |
| GSM2932540_ILC2.SI_2_RNA-seq | ILC2 | GSE109125 | Small_Intestine | 3534.37 | 24923.85 | 186.24 | 9.99 |
| GSM2932541_ILC2.ST2neg.SI_2_RNA-seq | ILC2 | GSE109125 | Small_Intestine | 1286.27 | 27956.01 | 390.89 | 11.43 |
| GSM2932542_ILC2.ST2neg.SI_1_RNA-seq | ILC2 | GSE109125 | Small_Intestine | 1159.72 | 29206.48 | 229.00 | 49.26 |
| GSM3208810_1_AAGGGA_ILC2_SILP_2 | ILC2 | GSE116092 | Small_Intestine | 4532.85 | 10484.19 | 29.79 | 62.65 |
| GSM3208813_4_TTCAGC_ILC2_LILP_1 | ILC2 | GSE116092 | Large_Intestine | 7988.15 | 6620.74 | 48.20 | 0.00 |
| GSM3208826_SL39396_ILC2s_LI_SS | ILC2 | GSE116092 | Large_Intestine | 4286.55 | 8985.73 | 105.45 | 1.01 |
| GSM3208829_SL39392_ILC2s_SI_SS | ILC2 | GSE116092 | Small_Intestine | 2993.93 | 14147.27 | 105.61 | 20.90 |
| GSM2056354_sample_63_RNA_BM_ILC2p_rpt1 | ILC2p | GSE77695 | Bone_marrow | 57.81 | 7499.92 | 270.24 | 3.58 |
| GSM2056355_sample_64_RNA_BM_ILC2p_rpt2 | ILC2p | GSE77695 | Bone_marrow | 40.45 | 6188.91 | 373.70 | 8.90 |
| GSM2932543_ILC3.CCR6pos.SI_1_RNA-seq | ILC3 | GSE109125 | Lamina_Propria | 46.29 | 3790.17 | 11013.79 | 44.68 |
| GSM2932544_ILC3.CCR6pos.SI_2_RNA-seq | ILC3 | GSE109125 | Lamina_Propria | 11.26 | 4025.81 | 15582.24 | 21.18 |
| GSM2932545_ILC3.NKp46negCCR6neg.SI_2_RNA-seq | ILC3 | GSE109125 | Lamina_Propria | 4.11 | 2939.14 | 11968.31 | 454.23 |
| GSM2932546_ILC3.NKp46pos.SI_1_RNA-seq | ILC3 | GSE109125 | Lamina_Propria | 30.35 | 3027.23 | 7929.88 | 1840.31 |
| GSM2932547_ILC3.NKp46pos.SI_2_RNA-seq | ILC3 | GSE109125 | Lamina_Propria | 15.82 | 2489.72 | 9478.99 | 1603.77 |
| GSM3208798_SNW13_ILC3GSM3208804_CCR6neg_WT_1 | ILC3 | GSE116092 | Small_Intestine | 684.20 | 412.27 | 9746.00 | 587.01 |
| GSM3208799_SNW14_ILC3GSM3208804_CCR6neg_WT_2 | ILC3 | GSE116092 | Small_Intestine | 985.51 | 457.78 | 6808.20 | 411.92 |
| GSM3208800_SNW15_ILC3GSM3208804_CCR6neg_WT_3 | ILC3 | GSE116092 | Small_Intestine | 1027.21 | 523.24 | 8567.42 | 228.94 |
| GSM3208804_SNW19_ILC3GSM3208804_CCR6pos_WT_1 | ILC3 | GSE116092 | Small_Intestine | 0.89 | 794.70 | 10053.42 | 0.94 |
| GSM3208805_SNW20_ILC3_SI_CCR6pos_WT_2 | ILC3 | GSE116092 | Small_Intestine | 165.33 | 536.87 | 8394.08 | 0.00 |
| GSM3208806_SNW21_ILC3_SI_CCR6pos_WT_3 | ILC3 | GSE116092 | Small_Intestine | 104.49 | 525.44 | 7875.30 | 46.53 |
| GSM3208811_2_CCTTCA_ILC3_CCR6p_SILP_2 | ILC3 | GSE116092 | Small_Intestine | 162.50 | 502.36 | 4705.19 | 17.84 |
| GSM3208812_3_GGACCC_ILC3_CCR6n_SILP_2 | ILC3 | GSE116092 | Small_Intestine | 804.77 | 691.09 | 4312.99 | 162.47 |
| GSM3208814_5_AAGACG_ILC3_CCR6p_LILP_1 | ILC3 | GSE116092 | Large_Intestine | 290.67 | 591.09 | 3242.17 | 7.54 |
| GSM3208815_6_CCTCGG_ILC3_CCR6n_LILP_1 | ILC3 | GSE116092 | Large_Intestine | 745.63 | 1040.18 | 2757.25 | 237.00 |
| GSM3208821_12_TTCAGC_ILC3_CCR6n_SILP_3 | ILC3 | GSE116092 | Small_Intestine | 2153.08 | 1411.60 | 1951.89 | 136.81 |
| GSM3208822_13_AAGACG_ILC3_CCR6p_SILP_3 | ILC3 | GSE116092 | Small_Intestine | 824.73 | 1198.20 | 1478.35 | 6.08 |
| GSM3208827_SL39395_CCR6negILC3s_SI_SS | ILC3 | GSE116092 | Small_Intestine | 547.86 | 1256.41 | 8018.74 | 189.90 |
| GSM3208828_SL39394_CCR6posILC3s_SI_SS | ILC3 | GSE116092 | Small_Intestine | 90.54 | 842.39 | 8283.87 | 16.24 |
| GSM2056338_sample_47_RNA_Sp_NK_rpt1 | NK | GSE77695 | Spleen | 0.00 | 598.50 | 5.14 | 3783.91 |
| GSM2056339_sample_48_RNA_Sp_NK_rpt2 | NK | GSE77695 | Spleen | 0.00 | 620.87 | 0.00 | 3437.77 |
| GSM2056348_sample_57_RNA_Sp_NK_IL2_IL12_rpt1 | NK | GSE77695 | Spleen | 0.51 | 285.66 | 0.29 | 3086.20 |
| GSM2056349_sample_58_RNA_Sp_NK_IL2_IL12_rpt2 | NK | GSE77695 | Spleen | 0.26 | 429.66 | 1.17 | 2810.07 |
| GSM2056356_sample_65_RNA_BM_NKp_rpt1 | NK | GSE77695 | Bone_marrow | 1.37 | 1732.51 | 642.48 | 552.51 |
| GSM2056357_sample_66_RNA_BM_NKp_rpt2 | NK | GSE77695 | Bone_marrow | 0.00 | 1872.84 | 657.52 | 777.77 |
| GSM2056358_sample_67_RNA_BM_iNK_rpt1 | NK | GSE77695 | Bone_marrow | 25.63 | 565.73 | 10.98 | 2258.76 |
| GSM2056359_sample_68_RNA_BM_iNK_rpt2 | NK | GSE77695 | Bone_marrow | 0.00 | 613.93 | 7.04 | 2149.21 |
| GSM2932569_NK.27neg11bpos.BM_1_RNA-seq | NK | GSE109125 | Bone_marrow | 3.99 | 2245.07 | 0.00 | 6904.97 |
| GSM2932570_NK.27neg11bpos.BM_2_RNA-seq | NK | GSE109125 | Bone_marrow | 0.00 | 1627.92 | 3.16 | 5612.25 |
| GSM2932571_NK.27neg11bpos.Sp_2_RNA-seq | NK | GSE109125 | Spleen | 0.00 | 1425.28 | 2.75 | 6356.32 |

| | | | | | | | |
|---|---|---|---|---|---|---|---|
| GSM2932572_NK.27pos11bneg.BM_1_RNA-seq | NK | GSE109125 | Bone_marrow | 0.00 | 1601.62 | 6.44 | 5273.34 |
| GSM2932573_NK.27pos11bneg.BM_2_RNA-seq | NK | GSE109125 | Bone_marrow | 0.00 | 1578.18 | 0.00 | 4054.36 |
| GSM2932574_NK.27pos11bneg.Sp_1_RNA-seq | NK | GSE109125 | Spleen | 0.00 | 1719.40 | 3.67 | 4957.17 |
| GSM2932575_NK.27pos11bneg.Sp_2_RNA-seq | NK | GSE109125 | Spleen | 0.00 | 1243.08 | 0.00 | 3713.91 |
| GSM2932576_NK.27pos11bpos.BM_1_RNA-seq | NK | GSE109125 | Bone_marrow | 0.00 | 2007.78 | 0.00 | 6806.63 |
| GSM2932577_NK.27pos11bpos.BM_2_RNA-seq | NK | GSE109125 | Bone_marrow | 0.00 | 1816.18 | 0.00 | 4922.10 |
| GSM2932578_NK.27pos11bpos.Sp_1_RNA-seq | NK | GSE109125 | Spleen | 0.00 | 1834.09 | 0.00 | 6931.00 |
| GSM2932579_NK.27pos11bpos.Sp_2_RNA-seq | NK | GSE109125 | Spleen | 0.00 | 1692.63 | 1.69 | 5478.77 |
| GSM3208786_SNW1_NKGSM3208804_WT_1 | NK | GSE116092 | Small_Intestine | 177.77 | 870.91 | 30.25 | 2788.88 |
| GSM3208787_SNW2_NKGSM3208804_WT_2 | NK | GSE116092 | Small_Intestine | 164.45 | 868.10 | 0.00 | 2657.50 |
| GSM3208788_SNW3_NKGSM3208804_WT_3 | NK | GSE116092 | Small_Intestine | 235.27 | 283.68 | 0.00 | 2064.01 |
| GSM3208819_10_CCTTCA_NK_SILP_3 | NK | GSE116092 | Small_Intestine | 555.50 | 670.54 | 3.50 | 701.45 |

| Supplemental Table 5 \| Unique genes for cluster 1, 2, 3, 4 | | | | | | |
|---|---|---|---|---|---|---|
| Higher expressed by cluster 4 | | | | | | |
| gene | logFC(4vs1) | padj(4vs1) | logFC(4vs3) | padj(4vs3) | logFC(4vs2) | padj(4vs2) |
| ABHD17A | 1.02849232 | 2.86E-72 | 0.6864923 | 3.42E-30 | 0.99940212 | 6.10E-56 |
| ABI3 | 0.95219825 | 5.45E-72 | 0.8399836 | 3.51E-42 | 0.96942436 | 7.34E-48 |
| ACTB | 0.72911056 | 2.99E-178 | 0.9823637 | 6.94E-226 | 0.85326567 | 1.11E-165 |
| ADGRG1 | 0.96602874 | 1.75E-75 | 0.8683214 | 1.25E-46 | 0.97926226 | 7.40E-49 |
| AGTRAP | 0.73098967 | 2.25E-44 | 0.5005441 | 5.54E-14 | 0.76591071 | 5.80E-33 |
| AKR1C3 | 0.96162791 | 2.06E-58 | 1.0531379 | 7.55E-62 | 1.05145104 | 3.51E-53 |
| ALOX5AP | 1.06131601 | 9.02E-76 | 1.5199977 | 6.75E-125 | 1.53767239 | 7.18E-107 |
| AOAH | 0.90568074 | 1.13E-58 | 0.5157773 | 2.42E-13 | 0.93172119 | 3.22E-44 |
| APMAP | 1.51790527 | 1.52E-175 | 0.7036985 | 2.00E-45 | 1.43151939 | 6.76E-115 |
| ARPC2 | 0.71475272 | 1.58E-111 | 0.7629433 | 3.23E-107 | 0.71542608 | 9.83E-78 |
| ATP6V0E1 | 0.72163546 | 1.74E-34 | 0.5939859 | 3.98E-25 | 0.59216311 | 1.88E-26 |
| C12orf75 | 1.00759159 | 2.59E-79 | 0.7331798 | 2.22E-42 | 0.63112591 | 1.82E-31 |
| C1orf21 | 0.97441215 | 5.61E-74 | 0.5952669 | 6.37E-21 | 0.95654972 | 1.73E-47 |
| CARD16 | 0.68952626 | 2.42E-25 | 0.7336526 | 6.57E-35 | 0.78392269 | 1.21E-36 |
| CCL3 | 1.4334961 | 2.38E-77 | 1.3783067 | 1.05E-56 | 1.44316923 | 5.32E-52 |
| CCL3L1 | 0.57101388 | 7.48E-32 | 0.5641168 | 8.80E-24 | 0.5693876 | 1.55E-19 |
| CCL4 | 1.65134083 | 8.91E-175 | 1.5466787 | 6.70E-131 | 1.66867916 | 4.48E-116 |
| CCL5 | 3.07275369 | 0 | 1.252792 | 1.32E-90 | 3.10614817 | 1.45E-217 |
| CD160 | 1.16475985 | 2.40E-86 | 0.9331294 | 5.18E-48 | 1.16881442 | 1.30E-59 |
| CD247 | 1.38358915 | 1.33E-179 | 1.0473561 | 6.93E-111 | 1.63954382 | 1.61E-160 |
| CD99 | 0.82155643 | 1.52E-72 | 0.9661463 | 7.65E-90 | 0.78164461 | 1.51E-55 |
| CEP78 | 1.09722663 | 6.74E-85 | 0.5406132 | 1.18E-14 | 1.06523169 | 3.34E-55 |
| CFL1 | 0.69398318 | 6.80E-139 | 0.5619778 | 1.13E-84 | 0.56736036 | 1.15E-70 |
| CHST2 | 0.91970265 | 1.74E-69 | 0.5052267 | 5.34E-14 | 0.92844574 | 1.12E-45 |
| CLIC3 | 1.98218008 | 3.86E-246 | 1.4732553 | 3.46E-144 | 1.31407252 | 5.48E-108 |
| CST7 | 2.63257715 | 0 | 1.1291199 | 2.62E-182 | 2.63864781 | 0 |
| CTSC | 0.88966288 | 1.91E-50 | 0.5161959 | 2.06E-15 | 0.70999038 | 1.76E-29 |
| CYBA | 1.03239818 | 1.21E-227 | 0.8236316 | 1.83E-141 | 1.18849344 | 1.29E-184 |
| DBI | 0.86204751 | 1.63E-62 | 0.7944628 | 2.96E-54 | 0.93524577 | 4.01E-61 |
| DOK2 | 1.01149759 | 1.63E-66 | 0.6146814 | 1.09E-22 | 0.95981805 | 3.06E-51 |
| EFHD2 | 1.19916731 | 5.90E-104 | 0.7321075 | 1.76E-40 | 1.32344546 | 1.76E-90 |
| FAM49B | 1.04610292 | 1.45E-95 | 0.5194944 | 2.80E-27 | 1.07359727 | 1.20E-80 |
| FCER1G | 2.02758112 | 0 | 0.5691294 | 2.39E-57 | 0.6131419 | 1.18E-55 |
| FCGR3A | 2.00583204 | 1.10E-262 | 1.6695591 | 9.11E-174 | 2.06120189 | 3.88E-181 |
| FCRL6 | 1.01754266 | 3.66E-78 | 0.880116 | 1.55E-45 | 1.02870904 | 8.93E-52 |
| FGFBP2 | 2.46733102 | 0 | 2.2943263 | 8.41E-256 | 2.52584942 | 1.39E-232 |
| FGR | 1.15340195 | 6.75E-100 | 0.5442589 | 1.31E-16 | 0.99670629 | 1.50E-52 |
| GNG2 | 0.84285833 | 2.58E-43 | 1.1114421 | 4.35E-74 | 1.11973882 | 1.87E-63 |
| GNPTAB | 0.8108165 | 5.67E-43 | 0.5610346 | 3.81E-16 | 0.67676117 | 2.21E-24 |
| GZMA | 2.8329271 | 0 | 1.000146 | 1.09E-163 | 2.82142782 | 0 |
| GZMB | 2.91585881 | 0 | 1.536282 | 1.97E-240 | 3.02180769 | 0 |
| GZMH | 2.02082038 | 8.42E-216 | 1.6468591 | 4.90E-131 | 2.04639117 | 2.03E-145 |
| GZMM | 1.89794634 | 4.93E-250 | 1.1740812 | 2.30E-114 | 1.95367047 | 3.58E-173 |
| HIST1H1E | 0.87665431 | 1.55E-36 | 0.5907124 | 6.89E-16 | 0.71554599 | 3.18E-26 |
| HMOX2 | 0.52907566 | 6.05E-11 | 0.51296 | 8.85E-16 | 0.53120131 | 2.64E-16 |
| IFI27L2 | 0.56902218 | 1.37E-15 | 0.5473756 | 1.81E-16 | 0.80907445 | 2.68E-35 |
| IFNG | 0.81604308 | 2.09E-46 | 0.658839 | 1.01E-21 | 0.77997647 | 2.12E-27 |
| IGFBP7 | 1.44058263 | 2.30E-133 | 0.6694827 | 5.08E-26 | 0.96813151 | 2.80E-52 |
| ITGB2 | 1.38983646 | 1.73E-230 | 0.6007945 | 1.14E-66 | 1.13190403 | 5.58E-133 |
| KIR2DL3 | 0.63065307 | 4.52E-35 | 0.5542488 | 1.65E-21 | 0.63951092 | 6.90E-23 |
| LBH | 0.98209773 | 7.01E-70 | 0.6809207 | 9.83E-27 | 1.01089528 | 7.13E-52 |

| | | | | | | |
|---|---|---|---|---|---|---|
| LYAR | 0.97415497 | 1.67E-55 | 0.6999873 | 2.01E-24 | 0.97819158 | 7.47E-42 |
| MAPK1 | 0.63764706 | 3.25E-24 | 0.520755 | 1.15E-14 | 0.62238848 | 1.23E-22 |
| MRPL10 | 0.66384543 | 7.63E-29 | 0.5371727 | 7.95E-16 | 0.62812944 | 4.53E-22 |
| MYBL1 | 0.72282351 | 1.88E-34 | 0.5407381 | 1.58E-14 | 0.78934527 | 1.07E-32 |
| MYL12A | 0.75533085 | 4.85E-161 | 0.7927915 | 5.76E-145 | 0.6347121 | 2.87E-85 |
| MYOM2 | 0.6220542 | 3.48E-23 | 0.6009644 | 4.52E-16 | 0.60615749 | 4.23E-14 |
| NKG7 | 3.4572828 | 0 | 0.9808386 | 1.62E-293 | 3.50669399 | 0 |
| OSTF1 | 0.905565 | 3.58E-64 | 0.5023685 | 2.64E-21 | 0.60594616 | 2.34E-29 |
| PFN1 | 0.84661855 | 7.42E-252 | 0.5981358 | 3.69E-130 | 0.747473 | 2.15E-151 |
| PLEK | 1.38157231 | 7.12E-126 | 0.8832824 | 3.30E-47 | 1.41824238 | 3.26E-89 |
| PPP1CA | 0.734661 | 1.06E-47 | 0.6706098 | 2.27E-41 | 0.70357677 | 2.28E-40 |
| PPP1R18 | 0.822038 | 3.90E-39 | 0.8017507 | 4.23E-39 | 0.68634409 | 9.37E-27 |
| PPP2R5C | 0.55598457 | 4.14E-13 | 1.0143849 | 1.10E-60 | 0.98806796 | 6.37E-55 |
| PRF1 | 2.63888225 | 0 | 1.5648576 | 3.31E-220 | 2.28294598 | 8.89E-257 |
| PRKCH | 1.12712317 | 4.76E-92 | 0.5480798 | 5.58E-20 | 0.7679906 | 2.18E-35 |
| PRSS23 | 1.14839654 | 5.61E-93 | 1.0370398 | 4.26E-60 | 1.17559557 | 5.09E-63 |
| PSAP | 0.6582682 | 9.95E-25 | 0.6013618 | 1.19E-26 | 0.52245592 | 1.66E-18 |
| PTGDS | 0.9809975 | 3.58E-10 | 1.0332344 | 1.32E-15 | 1.0073765 | 5.24E-11 |
| PYHIN1 | 1.82351723 | 4.07E-225 | 1.2216877 | 7.32E-109 | 1.84672003 | 4.07E-153 |
| RARRES3 | 0.55055156 | 2.92E-25 | 0.7762643 | 2.62E-53 | 1.09351577 | 5.26E-78 |
| RHOC | 1.33505253 | 4.94E-127 | 0.6772014 | 1.24E-34 | 1.10918686 | 4.61E-73 |
| S1PR5 | 1.40671079 | 1.56E-139 | 1.2817479 | 1.18E-94 | 1.41623508 | 2.19E-92 |
| SCP2 | 0.6118367 | 3.43E-22 | 0.5492352 | 5.43E-22 | 0.65998804 | 1.29E-31 |
| SPON2 | 2.48905021 | 0 | 2.2711884 | 5.47E-265 | 2.51193235 | 3.29E-242 |
| SYNE2 | 0.91740489 | 5.24E-56 | 1.123552 | 6.79E-75 | 1.28205879 | 2.52E-78 |
| TRBC1 | 0.64469567 | 2.07E-18 | 0.746251 | 1.15E-35 | 0.9083837 | 3.20E-46 |
| TTC38 | 1.16824056 | 3.76E-94 | 0.9703404 | 6.56E-55 | 1.00410571 | 3.22E-51 |
| ZEB2 | 1.0466087 | 2.97E-64 | 0.8556956 | 3.83E-42 | 1.2492348 | 3.89E-72 |
| **Higher expressed by cluster 1** | | | | | | |
| gene | logFC(1vs4) | padj(1vs4) | logFC(1vs3) | padj(1vs3) | logFC(1vs2) | padj(1vs2) |
| GADD45G | 0.84015272 | 1.23E-87 | 0.6317183 | 1.13E-40 | 0.70923687 | 2.62E-45 |
| GAP43 | 0.78036078 | 3.04E-63 | 0.7709391 | 3.09E-46 | 0.76196007 | 1.06E-36 |
| GATA3-AS | 0.66035526 | 4.20E-76 | 0.6554248 | 1.29E-56 | 0.63913429 | 2.90E-45 |
| HPGD | 1.03448995 | 1.71E-141 | 1.0801706 | 1.24E-116 | 0.82661819 | 1.27E-73 |
| HPGDS | 0.9119051 | 1.29E-105 | 0.9207918 | 2.09E-81 | 0.90756862 | 7.88E-66 |
| IL32 | 1.19229597 | 2.44E-268 | 1.4699825 | 1.05E-285 | 0.86201041 | 4.02E-146 |
| KLRB1 | 0.928484 | 2.40E-262 | 1.7728424 | 0 | 0.78799747 | 8.15E-157 |
| KRT1 | 0.78892441 | 1.59E-81 | 0.8060853 | 8.54E-64 | 0.80529956 | 9.13E-52 |
| NPDC1 | 0.79134721 | 9.61E-93 | 0.7868058 | 1.05E-68 | 0.54510614 | 7.15E-35 |
| PHTF2 | 0.79164857 | 1.35E-93 | 0.7287094 | 1.15E-61 | 0.55858572 | 1.28E-39 |
| PKIB | 0.67327527 | 3.24E-63 | 0.6735674 | 9.92E-48 | 0.67398523 | 1.21E-38 |
| RPL18A | 1.04327689 | 0 | 0.5490508 | 2.90E-245 | 0.50237434 | 7.02E-182 |
| RPLP0 | 1.33690649 | 0 | 0.7348578 | 1.90E-174 | 0.83244842 | 2.54E-169 |
| **Higher expressed by cluster 3** | | | | | | |
| gene | logFC(3vs4) | padj(3vs4) | logFC(3vs1) | padj(3vs1) | logFC(3vs2) | padj(3vs2) |
| CD2 | 0.84526937 | 4.41E-65 | 1.6661952 | 1.13E-192 | 1.48638139 | 3.87E-112 |
| CD74 | 0.68370123 | 3.43E-50 | 1.3185242 | 3.79E-136 | 0.55177468 | 8.62E-29 |
| CLDND1 | 0.53018511 | 1.98E-23 | 0.6568885 | 4.46E-26 | 0.67253419 | 3.93E-29 |
| CMC1 | 0.62842728 | 1.04E-78 | 2.9672114 | 0 | 2.80756487 | 2.23E-296 |
| COTL1 | 1.5862131 | 2.25E-185 | 1.6230941 | 5.31E-181 | 1.57808899 | 2.14E-126 |

| | | | | | | |
|---|---|---|---|---|---|---|
| DUSP2 | 0.81211709 | 1.56E-71 | 1.5162993 | 8.35E-161 | 1.29118748 | 9.46E-99 |
| GZMK | 2.22665774 | 0 | 2.4402401 | 0 | 2.42437356 | 7.55E-230 |
| ITGAX | 0.60061093 | 4.44E-36 | 0.8434358 | 1.50E-66 | 0.6464044 | 1.42E-26 |
| KIR2DL4 | 0.62896739 | 6.16E-38 | 0.7351497 | 1.01E-51 | 0.73113343 | 2.75E-31 |
| KLRC1 | 0.75183305 | 7.35E-79 | 2.1188614 | 8.42E-295 | 1.4483877 | 4.39E-135 |
| KLRC2 | 0.60632576 | 1.95E-21 | 0.9177363 | 4.56E-53 | 0.84925804 | 9.84E-29 |
| LINC0187 | 0.58031742 | 2.68E-33 | 1.3803044 | 4.49E-142 | 1.38345503 | 4.87E-96 |
| NCAM1 | 0.58756401 | 6.35E-34 | 0.8307709 | 4.34E-68 | 0.82131143 | 3.22E-42 |
| SPTSSB | 0.88210477 | 2.49E-74 | 0.9030137 | 4.31E-78 | 0.89914379 | 1.86E-48 |
| XCL1 | 1.532881 | 6.59E-176 | 1.8186694 | 1.05E-217 | 0.55631506 | 2.38E-34 |
| **Higher expressed by cluster 2** | | | | | | |
| gene | logFC(2vs4) | padj(2vs4) | logFC(2vs1) | padj(2vs1) | logFC(2vs3) | padj(2vs3) |
| AC245014 | 1.07272144 | 9.63E-80 | 0.9031628 | 3.29E-46 | 0.89703273 | 5.03E-45 |
| CAMK1 | 0.62419456 | 9.51E-30 | 0.599569 | 8.00E-18 | 0.61323255 | 2.26E-19 |
| CARD19 | 1.00944654 | 1.54E-61 | 0.7064022 | 9.17E-14 | 0.68580185 | 6.50E-19 |
| CST3 | 0.8611375 | 1.37E-42 | 0.9474414 | 1.80E-40 | 0.67067902 | 5.89E-16 |
| DDIT4 | 1.00533385 | 3.21E-61 | 0.6708241 | 2.72E-19 | 1.07670244 | 8.25E-56 |
| FAM30A | 0.88717763 | 1.76E-56 | 0.5758637 | 7.10E-09 | 0.72295257 | 5.10E-24 |
| FOS | 1.56189782 | 4.20E-181 | 0.5738614 | 3.94E-41 | 0.77628194 | 4.54E-61 |
| FXYD5 | 1.53437198 | 1.37E-248 | 0.6980536 | 8.60E-100 | 0.95644318 | 2.40E-135 |
| FXYD7 | 1.45038221 | 5.66E-116 | 0.8543447 | 5.77E-24 | 1.04722188 | 2.88E-44 |
| IER2 | 0.82679872 | 9.42E-107 | 0.6327746 | 2.05E-70 | 0.69461521 | 6.74E-72 |
| ITM2C | 1.2146067 | 1.08E-97 | 0.8798511 | 4.53E-35 | 0.95680307 | 5.30E-45 |
| JUN | 1.08841167 | 9.70E-141 | 0.6369185 | 1.47E-58 | 0.77518706 | 1.57E-76 |
| JUNB | 1.17771355 | 3.15E-159 | 0.7392858 | 1.94E-81 | 0.92143809 | 8.04E-102 |
| LST1 | 1.67002486 | 9.90E-126 | 0.9688903 | 5.51E-31 | 1.22555567 | 2.65E-55 |
| LTA4H | 1.18618186 | 6.61E-74 | 1.1867526 | 3.90E-58 | 0.68510606 | 5.04E-17 |
| PPP1R15A | 1.06859196 | 5.16E-87 | 0.6577667 | 1.07E-26 | 0.62668642 | 2.25E-26 |
| PTH2 | 0.62133309 | 5.14E-31 | 0.5950461 | 1.61E-26 | 0.61393202 | 1.81E-21 |
| RRBP1 | 1.03434821 | 3.49E-68 | 1.1123472 | 6.54E-65 | 0.9241851 | 1.29E-43 |
| TIMP1 | 1.28805495 | 3.07E-127 | 0.742736 | 1.63E-41 | 1.36483725 | 3.94E-112 |
| TMEM107 | 0.78878625 | 3.43E-28 | 0.9971618 | 1.43E-31 | 0.96335131 | 1.49E-35 |
| TMEM123 | 1.62386572 | 2.05E-188 | 0.604006 | 3.38E-34 | 0.53552525 | 3.20E-29 |
| TNFRSF18 | 1.52449699 | 9.87E-156 | 1.1446407 | 6.36E-83 | 0.93551993 | 6.57E-59 |
| TNFRSF4 | 1.44521006 | 9.24E-128 | 1.2906058 | 8.48E-89 | 1.27348087 | 9.88E-80 |
| TRAC | 0.83331443 | 9.04E-38 | 0.7149963 | 1.20E-15 | 0.90007765 | 9.34E-35 |
| ZFP36 | 0.55732339 | 5.74E-23 | 0.5278366 | 1.04E-12 | 0.5890367 | 1.14E-21 |
| **Less expressed by cluster 4** | | | | | | |
| gene | logFC(4vs1) | padj(4vs1) | logFC(4vs3) | padj(4vs3) | logFC(4vs2) | padj(4vs2) |
| CAPG | -0.8969185 | 6.89E-109 | -1.067645 | 4.43E-99 | -0.9434378 | 6.08E-57 |
| CD44 | -0.6473298 | 1.12E-75 | -1.269704 | 1.08E-151 | -1.1301286 | 4.15E-93 |
| CD52 | -0.9715955 | 1.13E-204 | -0.591307 | 7.43E-68 | -1.061707 | 1.82E-146 |
| EEF1A1 | -1.1947321 | 0 | -0.6555 | 4.24E-208 | -0.7910367 | 1.81E-213 |
| FOS | -0.9880364 | 5.78E-112 | -0.785616 | 8.04E-62 | -1.5618978 | 4.20E-181 |
| FXYD5 | -0.8363184 | 1.16E-153 | -0.577929 | 1.10E-48 | -1.534372 | 1.37E-248 |
| GATA3 | -1.3435341 | 1.53E-217 | -0.718867 | 1.51E-47 | -1.0302307 | 2.13E-58 |
| GPR183 | -0.6895004 | 1.29E-69 | -1.005777 | 2.06E-87 | -0.9474408 | 1.65E-61 |
| IL7R | -1.8914315 | 0 | -1.295211 | 4.73E-128 | -1.8986904 | 1.30E-200 |
| LTB | -2.5888772 | 0 | -1.227885 | 5.15E-104 | -2.6704619 | 4.35E-289 |
| NFKBIA | -1.0263772 | 7.87E-151 | -0.866689 | 1.79E-86 | -1.5192322 | 5.72E-180 |

| | | | | | | |
|---|---|---|---|---|---|---|
| NOP53 | -0.6699371 | 9.89E-90 | -0.515395 | 2.55E-30 | -0.653723 | 1.26E-34 |
| RPL10A | -0.90894 | 0 | -0.628403 | 8.48E-140 | -0.5376451 | 1.76E-80 |
| RPL11 | -0.8333591 | 0 | -0.544323 | 3.89E-149 | -0.5763142 | 9.34E-136 |
| RPL13 | -1.0298704 | 0 | -0.605578 | 1.07E-235 | -0.7007852 | 2.06E-224 |
| RPL13A | -0.7910354 | 0 | -0.641412 | 5.14E-258 | -0.7030105 | 6.42E-227 |
| RPL21 | -0.6634309 | 0 | -0.514023 | 7.73E-192 | -0.5436096 | 1.02E-167 |
| RPL32 | -0.9476081 | 0 | -0.593791 | 3.68E-202 | -0.6093497 | 2.29E-160 |
| RPL34 | -0.9352903 | 0 | -0.645245 | 1.72E-273 | -0.7170408 | 1.63E-259 |
| RPL36 | -0.8461866 | 0 | -0.54499 | 2.27E-131 | -0.6611573 | 5.93E-149 |
| RPL8 | -0.9977533 | 0 | -0.653628 | 1.41E-146 | -0.8039695 | 1.44E-167 |
| RPL9 | -0.7476072 | 4.73E-246 | -0.557651 | 4.16E-110 | -0.5158043 | 3.84E-74 |
| RPLP0 | -1.3369065 | 0 | -0.602049 | 1.30E-67 | -0.5044581 | 1.07E-34 |
| RPS12 | -0.9120804 | 0 | -0.652312 | 8.28E-176 | -0.5856169 | 2.14E-115 |
| RPS14 | -0.6724964 | 0 | -0.5324 | 8.87E-202 | -0.5417616 | 8.15E-163 |
| RPS19 | -0.7115064 | 0 | -0.59972 | 1.02E-232 | -0.5394401 | 1.32E-135 |
| RPS23 | -0.7998303 | 4.15E-301 | -0.629707 | 5.17E-161 | -0.623643 | 5.42E-120 |
| RPS25 | -0.781948 | 3.81E-292 | -0.528181 | 4.02E-115 | -0.6196309 | 6.54E-116 |
| RPS27 | -0.5802804 | 0 | -0.565789 | 8.93E-277 | -0.6354948 | 2.37E-261 |
| RPS27A | -0.6707433 | 0 | -0.50376 | 3.96E-179 | -0.5831807 | 7.01E-169 |
| RPS5 | -1.0669339 | 0 | -0.584673 | 2.97E-95 | -0.5447889 | 1.78E-59 |
| RPS6 | -0.9523814 | 0 | -0.592589 | 1.67E-202 | -0.577453 | 1.19E-154 |
| RPS8 | -1.2039157 | 0 | -0.588919 | 2.73E-134 | -0.7134152 | 4.36E-142 |
| RPSA | -0.8151757 | 2.20E-206 | -0.613978 | 1.06E-87 | -0.5075834 | 7.88E-44 |
| SNHG8 | -0.6807797 | 3.05E-82 | -0.577262 | 1.60E-32 | -0.6007691 | 5.74E-20 |
| SYPL1 | -0.5000602 | 1.16E-50 | -0.502926 | 3.52E-30 | -0.7090409 | 1.68E-33 |
| TMEM123 | -1.0198597 | 1.56E-165 | -1.08834 | 1.70E-106 | -1.6238657 | 2.05E-188 |
| TPT1 | -0.6625549 | 3.64E-182 | -0.581109 | 1.22E-98 | -0.5811386 | 2.40E-78 |
| **Less expressed by cluster 1** | | | | | | |
| gene | logFC(1vs4) | padj(1vs4) | logFC(1vs3) | padj(1vs3) | logFC(1vs2) | padj(1vs2) |
| AC007952 | -0.7003976 | 2.62E-25 | -0.548936 | 4.13E-20 | -0.8864825 | 8.10E-29 |
| CD63 | -0.6968817 | 1.43E-48 | -0.67556 | 4.59E-57 | -0.6495242 | 1.87E-27 |
| CD7 | -0.8625055 | 4.62E-134 | -0.831741 | 2.51E-90 | -1.0416055 | 4.91E-86 |
| CD74 | -0.634823 | 2.59E-17 | -1.318524 | 3.79E-136 | -0.7667495 | 7.46E-19 |
| CLIC3 | -1.9821801 | 3.86E-246 | -0.508925 | 3.08E-14 | -0.6681076 | 7.92E-14 |
| CTSW | -1.2866429 | 2.01E-270 | -1.472816 | 3.97E-294 | -1.1084031 | 9.12E-122 |
| FCER1G | -2.0275811 | 0 | -1.458452 | 3.18E-191 | -1.4144392 | 2.08E-126 |
| GNLY | -3.4117249 | 0 | -3.71212 | 0 | -1.0584355 | 3.54E-25 |
| GSN | -0.5526072 | 1.48E-26 | -0.651963 | 1.35E-44 | -0.5640818 | 6.66E-22 |
| HOPX | -1.3581677 | 2.36E-157 | -1.217764 | 9.01E-144 | -0.7618046 | 3.00E-24 |
| HSH2D | -0.8695527 | 5.07E-40 | -0.977372 | 5.98E-74 | -0.9731517 | 1.32E-38 |
| ID2 | -0.882743 | 1.17E-69 | -1.45381 | 3.01E-201 | -1.1805834 | 9.42E-71 |
| IFITM2 | -0.7428166 | 1.84E-102 | -0.792875 | 5.51E-119 | -0.9306615 | 2.59E-111 |
| KLRC1 | -1.3670283 | 1.01E-97 | -2.118861 | 8.42E-295 | -0.6704737 | 4.52E-22 |
| MAP3K8 | -0.9136255 | 2.56E-44 | -1.205517 | 5.50E-113 | -1.0339965 | 7.37E-43 |
| SUN2 | -0.7307716 | 6.07E-37 | -0.59637 | 9.10E-24 | -0.5680384 | 5.67E-07 |
| TPST2 | -1.1239019 | 5.07E-86 | -0.998135 | 2.02E-83 | -0.7372623 | 2.20E-17 |
| TYROBP | -2.3739202 | 0 | -2.345082 | 0 | -1.9640156 | 3.48E-243 |
| WDR74 | -0.6692673 | 1.97E-12 | -0.73899 | 4.15E-26 | -0.9388984 | 1.07E-28 |
| XCL2 | -0.8058121 | 3.18E-44 | -1.652623 | 3.21E-186 | -1.2025338 | 5.18E-75 |
| **Less expressed by cluster 3** | | | | | | |

| gene | logFC(3vs4) | padj(3vs4) | logFC(3vs1) | padj(3vs1) | logFC(3vs2) | padj(3vs2) |
| --- | --- | --- | --- | --- | --- | --- |
| KLRB1 | -0.8443584 | 2.60E-121 | -1.772842 | 0 | -0.9848449 | 1.17E-102 |
| LGALS1 | -1.0831479 | 5.23E-63 | -0.53824 | 1.68E-17 | -1.1819617 | 7.18E-69 |
| S100A4 | -0.8854237 | 1.53E-129 | -1.148179 | 2.00E-231 | -0.9941064 | 4.45E-129 |
| S100A6 | -0.6653582 | 2.31E-48 | -1.206061 | 6.57E-213 | -1.2125185 | 1.53E-124 |

| Supplemental Table 6 \| The effect of HIV-1 infection on NK cell subsets and ILCs | | | | | | | | | | | |
|---|---|---|---|---|---|---|---|---|---|---|---|
| DEgenes of CD56$^{hi}$NK cells | | | | | | | | | | | |
| HIV-1$^-$ vs HIV-1$^+$ viremic | | | | HIV-1$^-$ vs HIV-1$^+$ART | | | | HIV-1$^-$ vs HIV-1$^+$controller | | | |
| ID | log2FC | padj | type | ID | log2FC | padj | type | ID | log2FC | padj | type |
| CCL4L2 | -4.823 | 0.022762 | DN | LYAR | -1.206 | 0.03761 | DN | CXCL8 | -4.6909 | 0.009463 | DN |
| IFI44L | -3.331 | 1.92E-09 | DN | | | | | CSTB | -1.6534 | 0.004483 | DN |
| ODF3B | -2.558 | 0.0108075 | DN | | | | | MT-TS1 | -1.5669 | 0.007025 | DN |
| LINC02446 | -2.476 | 0.0428916 | DN | | | | | PARVG | -1.1431 | 0.002834 | DN |
| IFI6 | -2.351 | 1.54E-07 | DN | | | | | MYDGF | -1.0304 | 0.009974 | DN |
| IFI44 | -2.223 | 4.54E-05 | DN | | | | | NPIPB5 | 1.01876 | 0.037643 | UP |
| MX1 | -2.107 | 4.54E-05 | DN | | | | | CHPT1 | 1.11063 | 0.022431 | UP |
| RGS1 | -1.851 | 0.0182225 | DN | | | | | MYSM1 | 1.15636 | 2.54E-05 | UP |
| MT2A | -1.806 | 0.0119618 | DN | | | | | AC090398.2 | 1.21877 | 0.038815 | UP |
| ISG15 | -1.779 | 0.0045199 | DN | | | | | AC097532.3 | 1.22976 | 0.038944 | UP |
| EPSTI1 | -1.655 | 0.0053832 | DN | | | | | FANCM | 1.24395 | 0.038369 | UP |
| IRF7 | -1.648 | 0.000276 | DN | | | | | LINC00621 | 1.25512 | 0.015239 | UP |
| PRAL | -1.222 | 0.0300718 | DN | | | | | AC068418.3 | 1.28488 | 0.029121 | UP |
| SAT1 | -1.156 | 0.0182225 | DN | | | | | LRRK1 | 1.28942 | 0.038369 | UP |
| ISG20 | -1.053 | 0.0031996 | DN | | | | | PDE5A | 1.29729 | 0.049248 | UP |
| MSRB3 | 1.9256 | 0.0350385 | UP | | | | | SPATA6L | 1.29984 | 0.016635 | UP |
| A2MP1 | 2.1708 | 0.0392407 | UP | | | | | FAM241A | 1.30161 | 0.036776 | UP |
| | | | | | | | | TMEM45A | 1.36993 | 0.005479 | UP |
| | | | | | | | | TP53BP1 | 1.39629 | 0.018075 | UP |
| | | | | | | | | HUNK | 1.4325 | 0.049273 | UP |
| | | | | | | | | AC105233.4 | 1.50717 | 0.041895 | UP |
| | | | | | | | | L3MBTL4-AS1 | 1.50806 | 0.029121 | UP |
| | | | | | | | | PPAT | 1.53946 | 0.035214 | UP |
| | | | | | | | | CXorf56 | 1.55704 | 0.015239 | UP |
| | | | | | | | | Z82195.3 | 1.57065 | 0.008737 | UP |
| | | | | | | | | MCF2L2 | 1.61564 | 0.027608 | UP |
| | | | | | | | | AL109809.4 | 1.68263 | 0.040129 | UP |
| | | | | | | | | MYT1L | 1.69935 | 0.040283 | UP |
| | | | | | | | | WARS2-AS1 | 1.72143 | 0.015064 | UP |
| | | | | | | | | FSIP2 | 1.86617 | 0.005859 | UP |
| | | | | | | | | AC124290.1 | 1.91129 | 0.036045 | UP |
| | | | | | | | | AL117378.1 | 1.92478 | 0.028803 | UP |
| | | | | | | | | AC079949.1 | 1.95172 | 0.014264 | UP |
| | | | | | | | | AL390800.1 | 2.00253 | 0.038024 | UP |
| | | | | | | | | AC004801.7 | 2.02903 | 0.01847 | UP |
| | | | | | | | | LINC01550 | 2.12189 | 0.029121 | UP |
| | | | | | | | | PRKRA | 2.21844 | 0.028665 | UP |
| | | | | | | | | AL355312.2 | 2.24238 | 0.034336 | UP |
| | | | | | | | | AC234782.4 | 2.27261 | 0.011175 | UP |
| | | | | | | | | XKR9 | 2.43337 | 0.038369 | UP |
| | | | | | | | | AL021368.5 | 2.45903 | 0.046599 | UP |
| | | | | | | | | AC063950.1 | 2.53898 | 0.021037 | UP |
| | | | | | | | | LINC02196 | 2.60095 | 0.029121 | UP |
| | | | | | | | | AC018678.1 | 2.62945 | 0.041426 | UP |
| | | | | | | | | CHKB-DT | 2.70648 | 0.046599 | UP |
| | | | | | | | | AC116158.1 | 2.78554 | 0.022667 | UP |
| | | | | | | | | AC019257.9 | 2.83835 | 0.015918 | UP |
| | | | | | | | | AC093725.2 | 2.86119 | 0.01106 | UP |
| | | | | | | | | AC132938.6 | 2.98303 | 0.02471 | UP |
| | | | | | | | | PDE10A | 3.7179 | 0.032393 | UP |
| | | | | | | | | B3GALT2 | 4.13755 | 0.007025 | UP |
| | | | | | | | | MFSD1P1 | 4.32401 | 0.036776 | UP |
| DEgenes of CD56$^{dim}$NK cells | | | | | | | | | | | |
| HIV-1$^-$ vs HIV-1$^+$ viremic | | | | HIV-1$^-$ vs HIV-1$^+$ART | | | | HIV-1$^-$ vs HIV-1$^+$controller | | | |
| ID | log2FC | padj | type | ID | log2FC | padj | type | ID | log2FC | padj | type |
| IFI44L | -4.414 | 5.00E-08 | DN | No DEgenes | | | | ADGRG4 | 2.79888 | 0.005509 | UP |
| IFI44 | -3.498 | 1.73E-07 | DN | | | | | | | | |
| EGR2 | -2.678 | 0.0019053 | DN | | | | | | | | |
| LAG3 | -2.416 | 0.03233 | DN | | | | | | | | |
| IFIT1 | -2.407 | 0.0274472 | DN | | | | | | | | |
| IFIT3 | -2.375 | 0.0081911 | DN | | | | | | | | |
| IFI6 | -2.351 | 9.47E-12 | DN | | | | | | | | |
| MX1 | -2.22 | 1.29E-07 | DN | | | | | | | | |
| USP18 | -2.202 | 0.0043247 | DN | | | | | | | | |
| EPSTI1 | -2.139 | 1.91E-08 | DN | | | | | | | | |
| CMPK2 | -2.135 | 0.0032907 | DN | | | | | | | | |
| OAS3 | -2.029 | 0.0006523 | DN | | | | | | | | |
| OR10G3 | -1.839 | 0.0040054 | DN | | | | | | | | |
| XAF1 | -1.83 | 0.0203535 | DN | | | | | | | | |
| IRF7 | -1.814 | 6.06E-05 | DN | | | | | | | | |
| RSAD2 | -1.798 | 0.0474467 | DN | | | | | | | | |

| | | | | | | | | | | | |
|---|---|---|---|---|---|---|---|---|---|---|---|
| KIR3DX1 | -1.797 | 0.013072 | DN | | | | | | | | |
| MCOLN2 | -1.668 | 0.0331103 | DN | | | | | | | | |
| AHR | -1.598 | 0.0033068 | DN | | | | | | | | |
| HERC6 | -1.506 | 0.0100101 | DN | | | | | | | | |
| PARP9 | -1.485 | 2.21E-06 | DN | | | | | | | | |
| DUSP5 | -1.481 | 0.0327784 | DN | | | | | | | | |
| CDCA4 | -1.448 | 0.0001096 | DN | | | | | | | | |
| IFITM3 | -1.418 | 0.0401335 | DN | | | | | | | | |
| MX2 | -1.407 | 8.45E-05 | DN | | | | | | | | |
| LGALS3 | -1.333 | 0.0346109 | DN | | | | | | | | |
| HERC5 | -1.292 | 0.0008699 | DN | | | | | | | | |
| AC073264.3 | -1.256 | 0.0277054 | DN | | | | | | | | |
| DAPK2 | -1.222 | 0.0483201 | DN | | | | | | | | |
| HLA-DPA1 | -1.216 | 0.0157484 | DN | | | | | | | | |
| LGALS9 | -1.195 | 0.0006523 | DN | | | | | | | | |
| HELZ2 | -1.172 | 0.0008699 | DN | | | | | | | | |
| PRAL | -1.155 | 0.0012748 | DN | | | | | | | | |
| EHD4 | -1.121 | 0.0287293 | DN | | | | | | | | |
| GPR174 | -1.118 | 0.0218106 | DN | | | | | | | | |
| HLA-DPB1 | -1.11 | 0.0301533 | DN | | | | | | | | |
| SLC30A1 | -1.099 | 0.0049141 | DN | | | | | | | | |
| REC8 | -1.077 | 0.0234842 | DN | | | | | | | | |
| SAT1 | -1.056 | 0.0134206 | DN | | | | | | | | |
| RTCA | -1.033 | 0.0018512 | DN | | | | | | | | |
| LY6E | -1.013 | 0.0114896 | DN | | | | | | | | |
| AL137779.2 | 1.1168 | 0.0161268 | UP | | | | | | | | |
| CFAP298-TCP10L | 1.1374 | 0.0016388 | UP | | | | | | | | |
| IL18RAP | 1.2001 | 0.0483201 | UP | | | | | | | | |
| AC116158.3 | 1.2549 | 0.0457666 | UP | | | | | | | | |
| PLCB1 | 1.3283 | 0.0006033 | UP | | | | | | | | |
| AC090377.1 | 1.3455 | 0.029711 | UP | | | | | | | | |
| KIF13A | 1.3693 | 0.0133072 | UP | | | | | | | | |
| MYCBP2-AS1 | 1.4723 | 0.0492571 | UP | | | | | | | | |
| PRDM11 | 1.5001 | 0.031201 | UP | | | | | | | | |
| ITGA6 | 1.5304 | 0.0483201 | UP | | | | | | | | |
| A2M | 1.5648 | 0.0100101 | UP | | | | | | | | |
| NEIL1 | 1.5742 | 0.0049127 | UP | | | | | | | | |
| IGF1R | 1.602 | 0.0031895 | UP | | | | | | | | |
| AC063977.6 | 1.6154 | 0.0241633 | UP | | | | | | | | |
| ZNF595 | 1.6475 | 0.001023 | UP | | | | | | | | |
| ZNF718 | 1.684 | 0.0007949 | UP | | | | | | | | |
| AC118658.1 | 1.7705 | 0.0108068 | UP | | | | | | | | |
| PZP | 1.8543 | 0.0169264 | UP | | | | | | | | |
| CENPJ | 1.8805 | 0.0007949 | UP | | | | | | | | |
| MGAM | 2.0508 | 0.0152928 | UP | | | | | | | | |
| SIGLEC17P | 2.0911 | 0.0006523 | UP | | | | | | | | |
| A2MP1 | 2.2723 | 0.0016388 | UP | | | | | | | | |
| PFN2 | 2.2869 | 0.0492571 | UP | | | | | | | | |
| LINC00987 | 2.4723 | 0.0029037 | UP | | | | | | | | |
| ADGRG4 | 2.4822 | 0.0016388 | UP | | | | | | | | |
| COL13A1 | 2.7349 | 0.010237 | UP | | | | | | | | |
| AC011246.1 | 2.7525 | 0.0452351 | UP | | | | | | | | |
| MTND2P28 | 3.3671 | 0.029711 | UP | | | | | | | | |
| DEgenes of CD56$^{neg}$NK cells | | | | | | | | | | | |
| HIV-1$^-$ vs HIV-1$^+$ viremic | | | | HIV-1$^-$ vs HIV-1$^+$ART | | | | HIV-1$^-$ vs HIV-1$^+$controller | | | |
| ID | log2FC | padj | type | ID | log2FC | padj | type | ID | log2FC | padj | |
| LINC02446 | -4.115 | 0.000199 | DN | AC138907.8 | 4.566 | 0.01445 | UP | CYRIA | -1.3393 | 0.025132 | DN |
| MLF1 | -3.843 | 0.0487144 | DN | | | | | FRK | 1.28651 | 0.039409 | UP |
| IFI44L | -3.509 | 0.0005924 | DN | | | | | ZSCAN30 | 1.60053 | 0.017253 | UP |
| IFI44 | -3.478 | 3.47E-05 | DN | | | | | AC135983.3 | 1.60286 | 0.003325 | UP |
| KLRA1P | -2.979 | 0.0014181 | DN | | | | | AL589743.5 | 1.79344 | 0.015642 | UP |
| RSAD2 | -2.921 | 0.0054769 | DN | | | | | AC092802.2 | 1.98358 | 0.035034 | UP |
| GPR171 | -2.624 | 0.0054257 | DN | | | | | LINC01776 | 2.55579 | 0.015642 | UP |
| RHOB | -2.367 | 0.0033297 | DN | | | | | AC020910.7 | 3.33615 | 0.017253 | UP |
| TTN | -2.124 | 3.41E-06 | DN | | | | | | | | |
| MCOLN2 | -2.024 | 0.0069824 | DN | | | | | | | | |
| EPSTI1 | -1.944 | 0.0385931 | DN | | | | | | | | |
| IFI6 | -1.902 | 3.57E-05 | DN | | | | | | | | |
| MX1 | -1.813 | 0.0347771 | DN | | | | | | | | |
| GBP1 | -1.639 | 0.0151676 | DN | | | | | | | | |
| TNFRSF9 | -1.468 | 0.0399124 | DN | | | | | | | | |
| EOMES | -1.458 | 0.0032228 | DN | | | | | | | | |
| H1-10 | -1.414 | 0.0302705 | DN | | | | | | | | |
| ITM2A | -1.377 | 0.0492475 | DN | | | | | | | | |
| ISG20 | -1.376 | 0.0030535 | DN | | | | | | | | |

| | | | | | | | | | | | |
|---|---|---|---|---|---|---|---|---|---|---|---|
| VMP1 | -1.375 | 0.0130034 | DN | | | | | | | | |
| PPP1R16B | -1.321 | 0.0073477 | DN | | | | | | | | |
| AC058791.1 | -1.282 | 0.0069824 | DN | | | | | | | | |
| ARID5B | -1.239 | 0.0030535 | DN | | | | | | | | |
| ADAMTS2 | -1.227 | 0.0492475 | DN | | | | | | | | |
| TRAPPC13 | -1.216 | 0.0391446 | DN | | | | | | | | |
| CMC1 | -1.213 | 0.0067486 | DN | | | | | | | | |
| EZR | -1.149 | 0.0062903 | DN | | | | | | | | |
| EIF2AK2 | -1.127 | 0.0073825 | DN | | | | | | | | |
| B4GALT5 | -1.106 | 0.041828 | DN | | | | | | | | |
| KLF2 | -1.079 | 0.0055422 | DN | | | | | | | | |
| KRAS | -1.028 | 0.0160396 | DN | | | | | | | | |
| PREX1 | 1.0893 | 0.010829 | UP | | | | | | | | |
| IL2RB | 1.0956 | 0.0040217 | UP | | | | | | | | |
| HIPK2 | 1.302 | 0.0421387 | UP | | | | | | | | |
| SPOCK2 | 1.3796 | 0.021069 | UP | | | | | | | | |
| PEAK1 | 1.3926 | 0.0465665 | UP | | | | | | | | |
| SPON2 | 1.4836 | 0.0464834 | UP | | | | | | | | |
| CIDEB | 1.6256 | 0.0160396 | UP | | | | | | | | |
| ITGAM | 1.7053 | 4.96E-11 | UP | | | | | | | | |
| SYK | 1.7686 | 0.0073477 | UP | | | | | | | | |
| FCER1G | 1.8872 | 0.0145193 | UP | | | | | | | | |
| AC234782.4 | 1.9462 | 0.0489997 | UP | | | | | | | | |
| TANC2 | 1.9581 | 0.0218284 | UP | | | | | | | | |
| IL18RAP | 2.0254 | 0.0401695 | UP | | | | | | | | |
| KLRB1 | 2.0354 | 0.0004664 | UP | | | | | | | | |
| IRS2 | 2.0964 | 0.015646 | UP | | | | | | | | |
| RNF165 | 2.1307 | 0.0067486 | UP | | | | | | | | |
| LINC00294 | 2.2006 | 0.0048287 | UP | | | | | | | | |
| PLCB1 | 2.252 | 0.0069824 | UP | | | | | | | | |
| ZBTB16 | 2.3103 | 0.0003968 | UP | | | | | | | | |
| C1orf162 | 2.3805 | 0.0004664 | UP | | | | | | | | |
| AKR1C3 | 2.5819 | 0.0097332 | UP | | | | | | | | |
| AC008870.1 | 2.5828 | 0.0348354 | UP | | | | | | | | |
| IGF1R | 2.6304 | 0.0391446 | UP | | | | | | | | |
| FADS1 | 2.7356 | 0.0273885 | UP | | | | | | | | |
| TKTL1 | 2.965 | 0.0455554 | UP | | | | | | | | |
| SYNJ2 | 3.0076 | 0.0430589 | UP | | | | | | | | |
| LGALS9B | 3.2067 | 0.0073825 | UP | | | | | | | | |
| MYOM2 | 3.3616 | 0.0185163 | UP | | | | | | | | |
| SCRN1 | 4.0402 | 0.0030535 | UP | | | | | | | | |
| AC099494.2 | 4.7536 | 0.0243918 | UP | | | | | | | | |
| WIPF3 | 6.5278 | 0.0049415 | UP | | | | | | | | |
| DEgenes of ILCs | | | | | | | | | | | |
| HIV-1$^-$ vs HIV-1$^+$ viremic | | | | HIV-1$^-$ vs HIV-1$^+$ART | | | | HIV-1$^-$ vs HIV-1$^+$controller | | | |
| ID | log2FC | padj | type | ID | log2FC | padj | type | ID | log2FC | padj | type |
| IFI44L | -3.537 | 0.0003189 | DN | No DEgenes | | | | TTN | 1.78481 | 0.037471 | UP |
| TRDN-AS1 | -3.492 | 0.0273885 | DN | | | | | | | | |
| IFI44 | -2.902 | 0.0002956 | DN | | | | | | | | |
| PRPF40A | -1.003 | 0.0318127 | DN | | | | | | | | |
| DEgenes of ILCs (based on pvalue) | | | | | | | | | | | |
| HIV-1$^-$ vs HIV-1$^+$ viremic | | | | HIV-1$^-$ vs HIV-1$^+$ART | | | | HIV-1$^-$ vs HIV-1$^+$controller | | | |
| ID | log2FC | pvalue | type | ID | log2FC | pvalue | type | ID | log2FC | pvalue | type |
| RN7SL279P | -5.004 | 0.0029986 | DN | AC004982.2 | -3.171 | 0.00362 | DN | G0S2 | -4.3706 | 0.000147 | DN |
| SWT1 | -4.587 | 0.0115557 | DN | AL008638.6 | -2.979 | 0.00019 | DN | CKB | -4.2378 | 0.039931 | DN |
| LRRC59 | -4.316 | 0.0001652 | DN | AL354793.1 | -2.979 | 0.03406 | DN | AC093462.1 | -4.1777 | 0.001899 | DN |
| G0S2 | -4.009 | 0.0005862 | DN | VDR | -2.902 | 0.01622 | DN | AC005225.4 | -4.0639 | 0.000163 | DN |
| AC245128.3 | -3.93 | 0.0049713 | DN | APCDD1L-DT | -2.897 | 0.00614 | DN | CXCL8 | -3.983 | 0.00132 | DN |
| IGF2BP1 | -3.897 | 0.0008693 | DN | UBR5-AS1 | -2.84 | 0.00078 | DN | SERPINA1 | -3.4651 | 0.004516 | DN |
| OR6C4 | -3.852 | 0.0009683 | DN | CIT | -2.745 | 0.03053 | DN | ADORA2A-AS1 | -3.4649 | 0.012165 | DN |
| CXCL8 | -3.836 | 0.0021935 | DN | PAQR8 | -2.635 | 0.03466 | DN | AC004982.2 | -3.4175 | 0.002045 | DN |
| APOBEC3A | -3.813 | 0.0003773 | DN | ZSCAN25 | -2.58 | 0.01026 | DN | AKAP6 | -3.1896 | 0.004287 | DN |
| MPP1 | -3.802 | 0.0031641 | DN | FAM155A | -2.516 | 0.03379 | DN | LINC02160 | -3.1621 | 0.032155 | DN |
| VDR | -3.729 | 0.00234 | DN | BICD1 | -2.464 | 0.04526 | DN | IER3 | -3.0958 | 0.003352 | DN |
| ADAP1 | -3.713 | 0.0084878 | DN | SLC9B2 | -2.431 | 0.00564 | DN | AC105345.1 | -3.0195 | 0.00247 | DN |
| MAFB | -3.607 | 0.0030942 | DN | EPB41L4A | -2.41 | 0.00491 | DN | AC007106.2 | -2.9959 | 0.028353 | DN |
| IFI44L | -3.537 | 8.47E-08 | DN | LINC01926 | -2.361 | 0.04069 | DN | BEX3 | -2.9842 | 0.000105 | DN |
| NOD2 | -3.523 | 0.0004864 | DN | LINC02274 | -2.338 | 0.00636 | DN | CD36 | -2.9833 | 0.002928 | DN |
| TRDN-AS1 | -3.492 | 1.45E-05 | DN | HLA-J | -2.238 | 0.01021 | DN | AC132219.2 | -2.9329 | 0.007542 | DN |
| AC018678.1 | -3.43 | 0.0010081 | DN | METTL25 | -2.23 | 0.03379 | DN | MAFB | -2.9325 | 0.015907 | DN |
| AC016747.3 | -3.4 | 0.006786 | DN | ZNF529-AS1 | -2.176 | 0.00269 | DN | ANKRD9 | -2.9303 | 0.01511 | DN |
| IFIT2 | -3.397 | 0.0078606 | DN | PNKD | -2.089 | 0.00587 | DN | AL022394.1 | -2.924 | 0.045259 | DN |
| LGALS3BP | -3.37 | 0.0012351 | DN | AC243654.3 | -2.077 | 0.03164 | DN | NOTCH2NLC | -2.9212 | 9.79E-05 | DN |

| | | | | | | | | | | | |
|---|---|---|---|---|---|---|---|---|---|---|---|
| MED7 | -3.253 | 0.0178483 | DN | AC007191.1 | -2.077 | 0.0431 | DN | NECTIN3 | -2.9207 | 0.003177 | DN |
| SERPINA1 | -3.23 | 0.0084345 | DN | SLC39A1 | -2.072 | 0.01129 | DN | AL589862.1 | -2.8931 | 0.008486 | DN |
| FCGR2A | -3.219 | 0.0134429 | DN | PLEKHD1 | -2.071 | 0.02422 | DN | ZNF462 | -2.8332 | 0.022834 | DN |
| SRRM4 | -3.203 | 0.0076649 | DN | RN7SL624P | -2.013 | 0.0347 | DN | LINC01926 | -2.8144 | 0.015852 | DN |
| RTL8A | -3.201 | 0.0160547 | DN | H2BC15 | -2.005 | 0.04174 | DN | FBXO44 | -2.7841 | 0.00127 | DN |
| AC004982.2 | -3.173 | 0.0046926 | DN | SCGB3A2 | -1.994 | 0.0431 | DN | MINDY4 | -2.7519 | 0.018516 | DN |
| LINC01020 | -3.13 | 0.0388779 | DN | AC009090.6 | -1.984 | 0.04763 | DN | EML6 | -2.747 | 0.030076 | DN |
| TCEANC | -3.117 | 0.0008952 | DN | AC008403.3 | -1.95 | 0.04539 | DN | AC010889.2 | -2.7432 | 0.008334 | DN |
| AC010677.2 | -3.112 | 0.0223939 | DN | KIR3DL2 | -1.937 | 0.03802 | DN | MTFR1 | -2.7122 | 0.001686 | DN |
| CCR1 | -3.074 | 0.0191133 | DN | AL590627.1 | -1.869 | 0.04624 | DN | FPR1 | -2.6856 | 0.009398 | DN |
| LINC02384 | -3.035 | 0.0029287 | DN | FAM156A | -1.853 | 0.01194 | DN | ZNF438 | -2.6834 | 0.041727 | DN |
| LINC00520 | -3.022 | 0.0087628 | DN | METTL15 | -1.846 | 0.00021 | DN | CD14 | -2.6806 | 0.01041 | DN |
| EML6 | -3.012 | 0.0178022 | DN | AC097717.1 | -1.841 | 0.03379 | DN | HCK | -2.6693 | 0.007332 | DN |
| EBF1 | -3.004 | 0.0311753 | DN | AC100827.3 | -1.833 | 0.03487 | DN | AC005261.3 | -2.6336 | 0.007984 | DN |
| CD22 | -2.994 | 9.25E-05 | DN | FOXH1 | -1.829 | 0.01378 | DN | SLC4A4 | -2.6059 | 0.036198 | DN |
| AC005225.4 | -2.984 | 0.0068821 | DN | FBXO44 | -1.806 | 0.03586 | DN | IGF2BP1 | -2.582 | 0.029271 | DN |
| SHARPIN | -2.977 | 0.0043496 | DN | FCRL6 | -1.802 | 0.04446 | DN | CMC1 | -2.5754 | 0.003423 | DN |
| PLOD2 | -2.965 | 0.031857 | DN | GFM2 | -1.778 | 0.02279 | DN | AOAH | -2.57 | 0.000913 | DN |
| NKAIN3-IT1 | -2.964 | 0.0335624 | DN | LINC00578 | -1.776 | 0.03503 | DN | IRAK2 | -2.5285 | 0.0173 | DN |
| HINT2 | -2.955 | 0.0088395 | DN | TEP1 | -1.775 | 0.00688 | DN | AC108751.4 | -2.5249 | 0.027823 | DN |
| MAN1B1-DT | -2.952 | 0.0134989 | DN | AL157895.2 | -1.772 | 0.03803 | DN | PLOD1 | -2.5177 | 0.023697 | DN |
| AC126121.3 | -2.937 | 0.0012894 | DN | TCTE1 | -1.766 | 0.0255 | DN | SPIN4 | -2.5004 | 0.013176 | DN |
| MOB3B | -2.923 | 0.0107758 | DN | ZBTB2 | -1.729 | 0.01631 | DN | SFR1 | -2.4885 | 0.021894 | DN |
| AC022784.5 | -2.91 | 0.0238675 | DN | PARVG | -1.689 | 0.00287 | DN | BMERB1 | -2.4324 | 0.029988 | DN |
| IFI44 | -2.902 | 3.92E-08 | DN | EMILIN2 | -1.681 | 0.01983 | DN | PPBP | -2.4225 | 0.044195 | DN |
| PLOD1 | -2.883 | 0.0097283 | DN | GALNT16 | -1.68 | 0.02848 | DN | EIF2B3 | -2.4206 | 0.017115 | DN |
| SKA2 | -2.876 | 0.0004597 | DN | CENPE | -1.659 | 0.02402 | DN | AC005034.4 | -2.4178 | 0.018852 | DN |
| ZBED5-AS1 | -2.863 | 0.0464007 | DN | MUTYH | -1.645 | 0.02346 | DN | UBR5-AS1 | -2.4171 | 0.005607 | DN |
| ATP6V0E2 | -2.806 | 0.0011323 | DN | CUTC | -1.643 | 0.03645 | DN | TRUB1 | -2.4156 | 0.015613 | DN |
| PEX5 | -2.801 | 0.0168632 | DN | LINC00200 | -1.639 | 0.0144 | DN | MC2R | -2.4147 | 0.027739 | DN |
| ALPK2 | -2.8 | 0.0366791 | DN | AC110614.1 | -1.628 | 0.03515 | DN | GET1 | -2.3771 | 0.018314 | DN |
| AC007613.1 | -2.788 | 0.015933 | DN | MVB12B | -1.627 | 0.04885 | DN | HHEX | -2.3695 | 0.043435 | DN |
| MTCO1P12 | -2.786 | 0.0002051 | DN | NOTCH2NLC | -1.599 | 0.03292 | DN | SPART-AS1 | -2.363 | 0.029466 | DN |
| PON2 | -2.761 | 0.0010815 | DN | TIMM8B | -1.597 | 0.02169 | DN | ZNF385A | -2.3537 | 0.02137 | DN |
| AC022498.2 | -2.748 | 0.020021 | DN | SMIM11A | -1.593 | 0.00445 | DN | TREML2 | -2.3497 | 0.018 | DN |
| LRRC25 | -2.74 | 0.0018666 | DN | GSDMB | -1.581 | 0.01318 | DN | THBS1 | -2.3435 | 0.008222 | DN |
| TMEM170B | -2.705 | 0.016033 | DN | AC131649.1 | -1.581 | 0.01731 | DN | MPP3 | -2.3168 | 0.024009 | DN |
| SIN3B | -2.703 | 0.0004943 | DN | GZMA | -1.577 | 0.02744 | DN | ATP6AP1L | -2.2899 | 0.040289 | DN |
| RN7SL743P | -2.703 | 0.0098402 | DN | CD22 | -1.534 | 0.04481 | DN | S100A8 | -2.2561 | 0.020374 | DN |
| AC089998.4 | -2.701 | 0.0208381 | DN | SMIM15 | -1.514 | 0.04981 | DN | AC125494.1 | -2.2525 | 0.028801 | DN |
| RGS13 | -2.698 | 0.0371552 | DN | GBP5 | -1.511 | 0.00822 | DN | GPR160P2 | -2.2518 | 0.01453 | DN |
| AC022148.1 | -2.694 | 0.0357649 | DN | CD3G | -1.482 | 0.01048 | DN | OSM | -2.247 | 0.003562 | DN |
| TCF21 | -2.689 | 0.0122774 | DN | ZCCHC10 | -1.48 | 0.01311 | DN | PARP10 | -2.231 | 0.030381 | DN |
| LZTS3 | -2.685 | 0.0203343 | DN | TMEM267 | -1.478 | 0.04462 | DN | AC090241.3 | -2.2072 | 0.024123 | DN |
| RRAGD | -2.671 | 0.0264235 | DN | TBPL1 | -1.446 | 0.02235 | DN | HLA-J | -2.2051 | 0.013879 | DN |
| AC009951.4 | -2.626 | 0.022624 | DN | SYMPK | -1.445 | 0.03298 | DN | STX1B | -2.1683 | 0.021623 | DN |
| KLHL7-DT | -2.625 | 0.010676 | DN | DCK | -1.444 | 0.01171 | DN | LINC02482 | -2.1677 | 0.037946 | DN |
| MEG3 | -2.619 | 0.0325113 | DN | USP40 | -1.392 | 0.03708 | DN | GZMA | -2.1562 | 0.002791 | DN |
| TNFSF13B | -2.609 | 0.0118985 | DN | IL21R | -1.381 | 0.03648 | DN | NFIL3 | -2.1554 | 0.010947 | DN |
| C5orf66 | -2.602 | 0.0016764 | DN | BLOC1S1 | -1.343 | 0.02372 | DN | AC083799.1 | -2.1542 | 0.003475 | DN |
| CSF3R | -2.591 | 0.0018896 | DN | MIR4435-2HG | -1.321 | 0.03263 | DN | COMMD10 | -2.1394 | 0.042993 | DN |
| LINC01289 | -2.589 | 0.0496742 | DN | ESS2 | -1.298 | 0.04578 | DN | RNF34 | -2.1223 | 0.012332 | DN |
| TUFT1 | -2.589 | 0.0264863 | DN | TRAC | -1.296 | 0.00836 | DN | SMIM25 | -2.1166 | 0.024352 | DN |
| LITAFD | -2.586 | 0.0372919 | DN | GALR1 | -1.283 | 0.0435 | DN | POFUT2 | -2.1131 | 0.004575 | DN |
| ASB4 | -2.566 | 0.0287997 | DN | KIAA2013 | -1.264 | 0.01032 | DN | ZCWPW1 | -2.1049 | 0.033375 | DN |
| AC131280.1 | -2.555 | 0.0373077 | DN | PLXDC1 | -1.263 | 0.04735 | DN | AL359878.1 | -2.0982 | 0.012503 | DN |
| FPR1 | -2.555 | 0.0147526 | DN | PPFIBP1 | -1.259 | 0.03949 | DN | S100A9 | -2.0953 | 0.039181 | DN |
| AC073611.1 | -2.554 | 0.0310696 | DN | RGPD6 | -1.241 | 0.0097 | DN | AC020656.1 | -2.095 | 0.044639 | DN |
| ZNF385A | -2.547 | 0.0135254 | DN | BNIP3 | -1.23 | 0.03516 | DN | MOV10 | -2.0884 | 0.016741 | DN |
| CACNA1G | -2.544 | 0.0211032 | DN | CD3D | -1.228 | 0.04676 | DN | AC130456.5 | -2.0826 | 0.015233 | DN |
| AL161640.4 | -2.539 | 0.0278304 | DN | AL034351.3 | -1.226 | 0.04109 | DN | PRDX3 | -2.076 | 0.012726 | DN |
| RN7SL200P | -2.537 | 0.0160293 | DN | TNFRSF13C | -1.199 | 0.04793 | DN | GRN | -2.075 | 0.020938 | DN |
| PDLIM1 | -2.519 | 0.0232368 | DN | PTPRJ | -1.184 | 0.02997 | DN | EOLA2 | -2.0704 | 0.009733 | DN |
| PER2 | -2.516 | 0.0149762 | DN | SLC25A42 | -1.176 | 0.03673 | DN | PLBD2 | -2.064 | 0.019132 | DN |
| AC097374.1 | -2.511 | 0.0232009 | DN | TRAF1 | -1.175 | 0.01499 | DN | GORASP1 | -2.0509 | 0.02861 | DN |
| AC009779.4 | -2.511 | 0.0115856 | DN | C2orf92 | -1.168 | 0.03284 | DN | PACC1 | -2.039 | 0.039746 | DN |
| AC016042.1 | -2.508 | 0.0297964 | DN | GGA3 | -1.165 | 0.01686 | DN | KIR3DL2 | -2.0365 | 0.030689 | DN |
| ITPRIPL1 | -2.495 | 0.0033133 | DN | TRAF3IP2-AS1 | -1.161 | 0.04565 | DN | DANCR | -2.0216 | 0.005811 | DN |
| AP001525.1 | -2.48 | 0.0085512 | DN | HTR3B | -1.156 | 0.02693 | DN | AC233280.1 | -2.0059 | 0.004454 | DN |
| AC005034.4 | -2.476 | 0.017298 | DN | MIAT | -1.156 | 0.0106 | DN | AL133373.2 | -1.9819 | 0.049879 | DN |
| TPD52L1 | -2.474 | 0.0054195 | DN | MTATP6P1 | -1.155 | 0.00962 | DN | ADCY2 | -1.9702 | 0.029779 | DN |
| LINC00887 | -2.474 | 0.0487894 | DN | TAC4 | -1.127 | 0.03027 | DN | RELT | -1.9699 | 0.000863 | DN |
| AC108693.1 | -2.472 | 0.0239199 | DN | POLR2A | -1.124 | 0.00808 | DN | PLXNB2 | -1.9673 | 0.022034 | DN |
| AL358472.4 | -2.467 | 0.0312709 | DN | BISPR | -1.111 | 0.03251 | DN | SS18L1 | -1.9401 | 0.012526 | DN |
| IFI35 | -2.455 | 0.0183016 | DN | AC008013.2 | -1.102 | 0.03473 | DN | NPIPB3 | -1.9386 | 0.046304 | DN |
| AC116348.5 | -2.436 | 0.0485578 | DN | TENT5C | -1.1 | 0.03111 | DN | RER1 | -1.9345 | 0.011295 | DN |
| ZNF286B | -2.427 | 0.0212289 | DN | AC234782.4 | -1.079 | 0.04153 | DN | FRMD4B | -1.9305 | 0.0068 | DN |
| C5AR1 | -2.418 | 0.0005956 | DN | TBCE | -1.061 | 0.02052 | DN | CFP | -1.9158 | 0.015631 | DN |

| | | | | | | | | | | | |
|---|---|---|---|---|---|---|---|---|---|---|---|
| ZNF593 | -2.417 | 0.0238104 | DN | TPD52 | -1.045 | 0.04625 | DN | ETS2 | -1.9155 | 0.011341 | DN |
| AC092979.1 | -2.408 | 0.0497087 | DN | TRIM39 | -1.044 | 0.03248 | DN | TLR1 | -1.8992 | 0.043241 | DN |
| HEATR3 | -2.408 | 0.0063086 | DN | MEA1 | -1.028 | 0.04053 | DN | C5AR1 | -1.8875 | 0.006918 | DN |
| IFITM3 | -2.408 | 0.0036779 | DN | TUBA4A | -1.023 | 0.00142 | DN | LYZ | -1.8864 | 0.014187 | DN |
| HCK | -2.4 | 0.0177471 | DN | POLR2K | -1.02 | 0.01836 | DN | FCN1 | -1.8773 | 0.02018 | DN |
| DZANK1 | -2.389 | 0.0136687 | DN | LINC01588 | -1.016 | 0.02007 | DN | CDKN1A | -1.873 | 0.001386 | DN |
| OAS3 | -2.385 | 0.0002365 | DN | ODR4 | -1.015 | 0.04528 | DN | USP19 | -1.8671 | 0.024792 | DN |
| ADAMTSL4 | -2.384 | 0.049152 | DN | ARL17B | -1.013 | 0.02838 | DN | IFI30 | -1.8538 | 0.020043 | DN |
| XAF1 | -2.376 | 0.0293796 | DN | CD3E | -1.004 | 0.02572 | DN | PLXDC1 | -1.8415 | 0.004123 | DN |
| ADCY1 | -2.363 | 0.0488698 | DN | USPL1 | 1.0001 | 0.02509 | UP | SLC35E2B | -1.8375 | 0.005183 | DN |
| CPOX | -2.361 | 0.0260733 | DN | ZBTB16 | 1.0077 | 0.00528 | UP | AJ009632.2 | -1.8365 | 0.032286 | DN |
| IFI6 | -2.346 | 0.0001544 | DN | SYTL2 | 1.0151 | 0.03403 | UP | RBM4B | -1.8234 | 0.040355 | DN |
| RELT | -2.345 | 8.07E-05 | DN | ATP6V1F | 1.0186 | 0.03213 | UP | GALNT2 | -1.8129 | 0.006853 | DN |
| C6orf226 | -2.336 | 0.0094752 | DN | SIVA1 | 1.0287 | 0.0244 | UP | EREG | -1.8091 | 0.020826 | DN |
| DMXL2 | -2.333 | 0.0041618 | DN | ZKSCAN1 | 1.0289 | 0.02304 | UP | STMND1 | -1.8056 | 0.035232 | DN |
| GREB1 | -2.32 | 0.0362134 | DN | TUBB | 1.0348 | 0.00763 | UP | AC008629.1 | -1.7805 | 0.04393 | DN |
| AL031670.1 | -2.318 | 0.0138305 | DN | TMEM51-AS1 | 1.0474 | 0.04372 | UP | MCTP1 | -1.7766 | 0.032203 | DN |
| SYK | -2.316 | 0.0402733 | DN | TLE3 | 1.0533 | 0.03065 | UP | LINC00200 | -1.7731 | 0.00987 | DN |
| TRAPPC3L | -2.313 | 0.0129647 | DN | PHLPP1 | 1.0752 | 0.0218 | UP | ZNF574 | -1.7531 | 0.033646 | DN |
| NEDD4L | -2.31 | 0.0106637 | DN | AC068418.3 | 1.0828 | 0.01534 | UP | URB1 | -1.7482 | 0.039604 | DN |
| RPS4XP5 | -2.307 | 0.0164617 | DN | CCNG2 | 1.0886 | 0.0199 | UP | PLEK | -1.7464 | 0.031234 | DN |
| ERLIN1 | -2.305 | 0.0354306 | DN | CHAF1B | 1.0904 | 0.03378 | UP | RIT1 | -1.7311 | 0.002229 | DN |
| LINC01170 | -2.3 | 0.0497832 | DN | SNRPC | 1.0951 | 0.01616 | UP | PRPF8 | -1.7308 | 0.003441 | DN |
| FGGY | -2.3 | 0.0127136 | DN | CFH | 1.0998 | 0.04417 | UP | AC008915.2 | -1.7212 | 0.036797 | DN |
| SH3GL1 | -2.285 | 0.0009033 | DN | POLR2J | 1.0999 | 0.04254 | UP | SLC8B1 | -1.7138 | 0.025317 | DN |
| GZMM | -2.28 | 0.0190392 | DN | NRSN2-AS1 | 1.1005 | 0.0173 | UP | OR4F17 | -1.701 | 0.028008 | DN |
| NOTCH2NLC | -2.276 | 0.0029752 | DN | RNF166 | 1.1014 | 0.01815 | UP | CST3 | -1.7006 | 0.023712 | DN |
| FCN1 | -2.27 | 0.0051115 | DN | PUSL1 | 1.1058 | 0.04823 | UP | VGLL4 | -1.6994 | 0.01941 | DN |
| AC074286.1 | -2.246 | 0.037478 | DN | THUMPD2 | 1.1115 | 0.04996 | UP | CARD16 | -1.6975 | 0.024318 | DN |
| AL139002.1 | -2.237 | 0.044028 | DN | GSN | 1.1136 | 0.04824 | UP | SGMS1 | -1.6932 | 0.018292 | DN |
| PDE1B | -2.229 | 0.0072934 | DN | NSMCE2 | 1.1228 | 0.04321 | UP | TMEM135 | -1.6704 | 0.018411 | DN |
| EYA3 | -2.224 | 0.0045073 | DN | SUCLG2 | 1.1296 | 0.01863 | UP | TRMT44 | -1.6563 | 0.008673 | DN |
| LINC02246 | -2.221 | 0.0132566 | DN | TCP11L2 | 1.1308 | 0.04821 | UP | KLHL5 | -1.6514 | 0.01826 | DN |
| MRPS17 | -2.216 | 0.0166305 | DN | KIT | 1.1356 | 0.04818 | UP | SP2 | -1.6421 | 0.005755 | DN |
| KRCC1 | -2.216 | 0.0139077 | DN | LIN7C | 1.139 | 0.02912 | UP | UBE2M | -1.6359 | 0.010142 | DN |
| GSTM3 | -2.215 | 0.0171202 | DN | ZNF525 | 1.1462 | 0.03057 | UP | HERC6 | -1.6282 | 0.006193 | DN |
| IFI30 | -2.199 | 0.0061962 | DN | ABRAXAS1 | 1.1656 | 0.04476 | UP | MARCKS | -1.6255 | 0.046665 | DN |
| NBPF10 | -2.193 | 0.0087072 | DN | ZFHX2-AS1 | 1.1705 | 0.0461 | UP | C9orf85 | -1.6191 | 0.038729 | DN |
| THBS1 | -2.192 | 0.014047 | DN | STAG3L3 | 1.1788 | 0.0241 | UP | UBE2A | -1.6083 | 0.001283 | DN |
| CPEB4 | -2.192 | 0.0006687 | DN | PZP | 1.1804 | 0.04363 | UP | LINC01148 | -1.586 | 0.049587 | DN |
| CLDN12 | -2.186 | 0.0044172 | DN | DUSP16 | 1.1918 | 0.01408 | UP | NBEAL1 | -1.5763 | 0.022354 | DN |
| E2F5 | -2.186 | 0.0147003 | DN | PLXND1 | 1.2139 | 0.03178 | UP | PARVG | -1.5699 | 0.006564 | DN |
| PPP1CA | -2.183 | 0.0042506 | DN | RRP1 | 1.2141 | 0.04969 | UP | FAM156A | -1.5545 | 0.038865 | DN |
| DNAJC15 | -2.182 | 0.0024147 | DN | RNF126 | 1.2191 | 0.00126 | UP | TEX9 | -1.5515 | 0.009766 | DN |
| SASS6 | -2.163 | 0.0045162 | DN | RASSF8-AS1 | 1.2197 | 0.0314 | UP | AC123912.4 | -1.5256 | 0.011125 | DN |
| AC008533.1 | -2.134 | 0.0050191 | DN | EIF2B4 | 1.2508 | 0.02487 | UP | KIAA1328 | -1.5209 | 0.005842 | DN |
| SPOCD1 | -2.132 | 0.0325013 | DN | EXOC4 | 1.2527 | 0.01769 | UP | ACSL1 | -1.5114 | 0.022409 | DN |
| H2AW | -2.111 | 0.0466193 | DN | ATP5ME | 1.2712 | 0.04252 | UP | ACSM2B | -1.5086 | 0.037358 | DN |
| SLC30A1 | -2.102 | 0.0364793 | DN | COMMD3 | 1.2713 | 0.01076 | UP | CASS4 | -1.4947 | 0.012437 | DN |
| TNFAIP2 | -2.097 | 0.0033556 | DN | MYO18A | 1.2728 | 0.03452 | UP | MXD3 | -1.4915 | 0.049736 | DN |
| MMP25 | -2.094 | 0.005095 | DN | LINC00467 | 1.2779 | 0.009 | UP | GORAB | -1.4901 | 0.032151 | DN |
| ZNRD2 | -2.088 | 0.013246 | DN | MSRB3 | 1.2819 | 0.03553 | UP | SGK1 | -1.4817 | 0.006346 | DN |
| AC091959.3 | -2.088 | 0.0432913 | DN | ASPH | 1.2859 | 0.02644 | UP | IFI44L | -1.4808 | 0.026044 | DN |
| CFP | -2.08 | 0.0095653 | DN | COLCA1 | 1.2931 | 0.04294 | UP | GBP4 | -1.479 | 0.000479 | DN |
| PINX1 | -2.078 | 0.0468279 | DN | COLEC12 | 1.2968 | 0.01458 | UP | OAS3 | -1.4704 | 0.023132 | DN |
| FCER1G | -2.076 | 0.0211159 | DN | PPP4R1 | 1.3017 | 0.01554 | UP | PPFIA1 | -1.4559 | 0.013411 | DN |
| OASL | -2.074 | 0.0277001 | DN | ACBD7 | 1.3062 | 0.04649 | UP | LINC00174 | -1.4515 | 0.042961 | DN |
| HLA-DQB1 | -2.071 | 0.0254275 | DN | PLA2G12A | 1.3067 | 0.00992 | UP | KANK2 | -1.4363 | 0.048041 | DN |
| CD36 | -2.054 | 0.0444561 | DN | FXYD7 | 1.3178 | 0.04616 | UP | KATNA1 | -1.4117 | 0.025613 | DN |
| GOLGA2P5 | -2.05 | 0.0008118 | DN | GBA2 | 1.329 | 0.04551 | UP | AL109837.3 | -1.4072 | 0.030588 | DN |
| PRKCE | -2.046 | 0.0074086 | DN | MMS22L | 1.347 | 0.0063 | UP | UBE2W | -1.4038 | 0.045662 | DN |
| FKBP4 | -2.045 | 0.0157105 | DN | BDH1 | 1.3479 | 0.04034 | UP | GGA3 | -1.397 | 0.004755 | DN |
| EPSTI1 | -2.043 | 0.0081605 | DN | TYMP | 1.3479 | 0.01296 | UP | RBM42 | -1.3872 | 0.045197 | DN |
| AIF1 | -2.039 | 0.0336493 | DN | IL2RA | 1.3657 | 0.02719 | UP | SCYL3 | -1.3817 | 0.02813 | DN |
| PMAIP1 | -2.032 | 0.0009083 | DN | LINC00665 | 1.3775 | 0.00483 | UP | TRGC2 | -1.3738 | 0.015419 | DN |
| AP002812.3 | -2.029 | 0.0490541 | DN | SPTSSA | 1.3949 | 0.04594 | UP | ENO2 | -1.3679 | 0.020816 | DN |
| AC103858.3 | -2.021 | 0.0324918 | DN | ZBTB8A | 1.3959 | 0.04626 | UP | CCNK | -1.3653 | 0.001959 | DN |
| AC019080.1 | -2.017 | 0.0080458 | DN | BID | 1.399 | 0.02234 | UP | CHRNA5 | -1.365 | 0.031141 | DN |
| RETSAT | -2.016 | 0.0214555 | DN | MIR22HG | 1.4018 | 0.04515 | UP | SYNGR2 | -1.3642 | 0.007502 | DN |
| IRF7 | -2.014 | 0.0035109 | DN | MRPL52 | 1.407 | 0.00624 | UP | ZNF76 | -1.363 | 0.013594 | DN |
| GNLY | -2.011 | 0.0024213 | DN | EMC1 | 1.4073 | 0.02084 | UP | ZNF136 | -1.3602 | 0.011022 | DN |
| ITGAX | -2.01 | 0.0191218 | DN | AP3S2 | 1.4127 | 0.02357 | UP | DCK | -1.3518 | 0.021833 | DN |
| MARCKS | -1.995 | 0.0156456 | DN | ZNF397 | 1.4288 | 0.03897 | UP | ZACN | -1.3511 | 0.020216 | DN |
| PDE6B | -1.99 | 0.0076951 | DN | DPEP2 | 1.4309 | 0.04542 | UP | WDR91 | -1.3502 | 0.047693 | DN |
| CST3 | -1.978 | 0.0093114 | DN | SWAP70 | 1.4311 | 0.01102 | UP | MFSD1 | -1.3352 | 0.028776 | DN |
| STT3A | -1.97 | 0.0024467 | DN | TOR1A | 1.4343 | 0.03806 | UP | CD3G | -1.3332 | 0.022815 | DN |
| ZNF707 | -1.944 | 0.0045036 | DN | NUDT22 | 1.4706 | 0.02651 | UP | ZDHHC21 | -1.3247 | 0.034418 | DN |
| AL513497.1 | -1.942 | 0.0455801 | DN | PCAT19 | 1.4729 | 0.047 | UP | AC004047.1 | -1.315 | 0.012237 | DN |

| | | | | | | | | | | | |
|---|---|---|---|---|---|---|---|---|---|---|---|
| AL031595.2 | -1.939 | 8.15E-05 | DN | ASB8 | 1.4744 | 0.02895 | UP | MMUT | -1.3048 | 0.040932 | DN |
| RHD | -1.938 | 0.008299 | DN | PLEKHF1 | 1.4913 | 0.03595 | UP | PRORP | -1.2962 | 0.004404 | DN |
| AC098591.3 | -1.931 | 0.0446705 | DN | BET1 | 1.5122 | 0.04801 | UP | NPIPB5 | -1.2937 | 0.007983 | DN |
| EPB41L4A | -1.922 | 0.0317594 | DN | PPP4C | 1.5179 | 0.02803 | UP | RIN3 | -1.2906 | 0.030794 | DN |
| LGALS3 | -1.914 | 0.0023046 | DN | AP005482.4 | 1.531 | 0.04337 | UP | ATP6V1B2 | -1.2885 | 0.045114 | DN |
| FBXO44 | -1.914 | 0.0321648 | DN | CRCP | 1.5346 | 0.00617 | UP | F2RL2 | -1.2857 | 0.02037 | DN |
| HLA-DRA | -1.906 | 0.0188659 | DN | AC096645.1 | 1.5355 | 0.04999 | UP | AL034351.3 | -1.2843 | 0.038007 | DN |
| AL035587.1 | -1.906 | 0.0078119 | DN | OR5H14 | 1.5511 | 0.02826 | UP | RNU1-46P | -1.2817 | 0.042048 | DN |
| POP4 | -1.903 | 0.041787 | DN | NIT1 | 1.5657 | 0.01418 | UP | RASSF3 | -1.2668 | 0.01143 | DN |
| OR5AU1 | -1.899 | 0.0188808 | DN | EIF2D | 1.5677 | 0.00054 | UP | B4GALT5 | -1.2537 | 0.009065 | DN |
| RIT1 | -1.897 | 0.0009327 | DN | STK39 | 1.573 | 0.01174 | UP | GLUL | -1.2512 | 0.006485 | DN |
| ASAP1-IT2 | -1.895 | 0.0456205 | DN | AC127035.1 | 1.5732 | 0.03964 | UP | ATP11A | -1.2416 | 0.037404 | DN |
| HLA-DMA | -1.893 | 0.0352025 | DN | DDR1-DT | 1.5736 | 0.04102 | UP | TRIM69 | -1.2229 | 0.020266 | DN |
| HDAC10 | -1.892 | 0.0185809 | DN | EEF1E1 | 1.5749 | 0.03703 | UP | TANK | -1.2176 | 0.020886 | DN |
| PNKD | -1.891 | 0.0176036 | DN | COG8 | 1.5787 | 0.00404 | UP | LONRF1 | -1.2148 | 0.042818 | DN |
| RASAL2 | -1.887 | 0.0375139 | DN | AC090515.6 | 1.5822 | 0.04658 | UP | AC006064.4 | -1.2117 | 0.000126 | DN |
| MRPS33 | -1.882 | 0.0008272 | DN | DCXR | 1.5929 | 0.00992 | UP | COQ10A | -1.2044 | 0.03479 | DN |
| AC091152.4 | -1.88 | 0.0272871 | DN | NCK1-DT | 1.5952 | 0.0341 | UP | COPS3 | -1.2039 | 0.035175 | DN |
| ZNF696 | -1.874 | 0.0330609 | DN | MYO15B | 1.6 | 0.03219 | UP | PIM2 | -1.2029 | 0.014566 | DN |
| NFIL3 | -1.871 | 0.0296466 | DN | TCF4 | 1.6126 | 0.0166 | UP | STK38 | -1.2001 | 0.003926 | DN |
| AC034238.2 | -1.86 | 0.0072254 | DN | AC008982.2 | 1.6143 | 0.02615 | UP | JADE2 | -1.197 | 0.017716 | DN |
| RN7SL105P | -1.86 | 0.0270049 | DN | PHKA2 | 1.6224 | 0.0268 | UP | RHCE | -1.1877 | 0.025452 | DN |
| GRN | -1.857 | 0.0402623 | DN | AL359641.1 | 1.6273 | 0.04964 | UP | ANXA2 | -1.1872 | 0.002333 | DN |
| LINC01563 | -1.847 | 0.0062407 | DN | AC025580.2 | 1.6277 | 0.01789 | UP | TRAPPC10 | -1.1819 | 0.037639 | DN |
| TNFRSF13C | -1.847 | 0.0031109 | DN | ZNF765 | 1.6294 | 0.00013 | UP | DHRS1 | -1.1732 | 0.030266 | DN |
| NAPB | -1.844 | 0.015855 | DN | MT-TC | 1.6297 | 0.02428 | UP | DPH3 | -1.1697 | 0.032931 | DN |
| CHRM5 | -1.842 | 0.0291366 | DN | AC008537.3 | 1.6371 | 0.0262 | UP | AL713852.1 | -1.1556 | 0.047148 | DN |
| TAF4B | -1.834 | 0.0168692 | DN | LRP12 | 1.6398 | 0.00742 | UP | UBE4A | -1.1504 | 0.010314 | DN |
| GBP5 | -1.821 | 0.0020712 | DN | MTRNR2L8 | 1.6472 | 0.002 | UP | PSMB7 | -1.149 | 0.009283 | DN |
| RN7SL525P | -1.82 | 0.0223245 | DN | GUF1 | 1.6537 | 0.01683 | UP | AC106845.1 | -1.142 | 0.018234 | DN |
| FILIP1L | -1.812 | 0.0367416 | DN | CKAP2 | 1.6542 | 0.03334 | UP | NBDY | -1.1405 | 0.024218 | DN |
| AC090952.2 | -1.81 | 0.0395838 | DN | UBE2Q2P2 | 1.6618 | 0.03063 | UP | MAGED2 | -1.1345 | 0.02458 | DN |
| CC2D2A | -1.808 | 0.0109641 | DN | SV2A | 1.6655 | 0.01472 | UP | ST8SIA5 | -1.1204 | 0.04306 | DN |
| LZTR1 | -1.807 | 0.0194565 | DN | PWRN1 | 1.6675 | 0.03479 | UP | YTHDF3 | -1.1059 | 0.009307 | DN |
| AC005841.1 | -1.806 | 0.0367598 | DN | SH3PXD2A | 1.6694 | 0.01896 | UP | NINJ1 | -1.1025 | 0.017251 | DN |
| HSF2 | -1.802 | 0.0052577 | DN | PPIB | 1.6708 | 0.01366 | UP | INF2 | -1.1003 | 0.022124 | DN |
| ZHX2 | -1.8 | 0.0212359 | DN | TMEM67 | 1.6772 | 0.00428 | UP | WDR83OS | -1.0994 | 0.006452 | DN |
| OSBPL1A | -1.796 | 0.047031 | DN | SMG5 | 1.6865 | 0.00752 | UP | PIM3 | -1.0964 | 0.014063 | DN |
| TMEM62 | -1.795 | 0.034025 | DN | AC010325.1 | 1.692 | 0.03557 | UP | HIPK2 | -1.0826 | 0.029885 | DN |
| FXN | -1.783 | 0.0194046 | DN | UHRF1BP1L | 1.6939 | 0.01447 | UP | SERPINB9 | -1.0825 | 0.022878 | DN |
| AF001548.3 | -1.781 | 0.0317983 | DN | AC005487.1 | 1.699 | 0.01024 | UP | SHKBP1 | -1.0811 | 0.008891 | DN |
| MT2A | -1.766 | 0.0079491 | DN | ZNF500 | 1.7217 | 0.01657 | UP | TUBA4A | -1.0765 | 0.001033 | DN |
| RAB3B | -1.765 | 0.0294491 | DN | LINC00884 | 1.7231 | 0.00669 | UP | ACSS1 | -1.0619 | 0.049556 | DN |
| C18orf25 | -1.762 | 0.0041914 | DN | DNAJC14 | 1.732 | 0.01235 | UP | TSPAN3 | -1.0569 | 0.044054 | DN |
| ATP6V0E1 | -1.753 | 0.008954 | DN | AC020978.9 | 1.741 | 0.03665 | UP | MIDN | -1.0514 | 0.022635 | DN |
| PIGG | -1.744 | 0.0158951 | DN | EAF1-AS1 | 1.7445 | 0.02374 | UP | NBPF12 | -1.0514 | 0.036045 | DN |
| SGK1 | -1.737 | 0.0016053 | DN | DCAF1 | 1.7463 | 0.02479 | UP | ATG13 | -1.0354 | 0.02947 | DN |
| STK33 | -1.732 | 0.0413551 | DN | MTRF1L | 1.7518 | 0.03523 | UP | ZNF654 | -1.017 | 0.030818 | DN |
| RCBTB1 | -1.724 | 0.013583 | DN | PPIEL | 1.7553 | 0.04623 | UP | CD74 | -1.0166 | 0.025382 | DN |
| ZNF529-AS1 | -1.723 | 0.0242864 | DN | CLU | 1.7658 | 0.03751 | UP | AC068759.1 | -1.0119 | 0.039874 | DN |
| IDH2-DT | -1.72 | 0.0182893 | DN | RFC2 | 1.7831 | 0.03966 | UP | PRKCH | -1.0048 | 0.004636 | DN |
| AC098934.2 | -1.715 | 0.0250257 | DN | AC025268.1 | 1.7859 | 0.03966 | UP | SMAP2 | -1.0006 | 0.002657 | DN |
| TSPY26P | -1.715 | 0.0361187 | DN | RMI2 | 1.7875 | 0.008 | UP | LINC01609 | 1.00087 | 0.008087 | UP |
| EAF1 | -1.709 | 0.0147389 | DN | DENND2A | 1.7877 | 0.04329 | UP | UBXN2A | 1.00249 | 0.02586 | UP |
| NCOA6 | -1.701 | 0.0278852 | DN | PLAAT3 | 1.7884 | 0.00701 | UP | AC073225.1 | 1.00803 | 0.035834 | UP |
| NOMO3 | -1.696 | 0.0163187 | DN | MCCC1 | 1.7892 | 0.04922 | UP | HLA-F | 1.01615 | 0.0198 | UP |
| FZD10-AS1 | -1.684 | 0.0374611 | DN | LINC02334 | 1.7934 | 0.00561 | UP | TMED2-DT | 1.02175 | 0.025509 | UP |
| AL049776.1 | -1.677 | 0.0046816 | DN | HERC2P3 | 1.7949 | 0.04427 | UP | AL138899.1 | 1.02205 | 0.045832 | UP |
| KIF22 | -1.671 | 0.0212922 | DN | CREB5 | 1.8055 | 0.03745 | UP | ARID3B | 1.02233 | 0.036079 | UP |
| DNASE2 | -1.668 | 0.0147395 | DN | MAPT-AS1 | 1.8071 | 0.00829 | UP | POLK | 1.03073 | 0.027992 | UP |
| HERC6 | -1.667 | 0.0058364 | DN | CEACAM22P | 1.8124 | 0.0302 | UP | SH2D3A | 1.03101 | 0.027105 | UP |
| PTCD2 | -1.66 | 0.0336433 | DN | TMEM132A | 1.815 | 0.03202 | UP | LINC00836 | 1.03522 | 0.045288 | UP |
| CYP27C1 | -1.654 | 0.0134006 | DN | CEP76 | 1.8248 | 0.02796 | UP | WDFY2 | 1.03882 | 0.020654 | UP |
| PCID2 | -1.649 | 0.0017072 | DN | EIF2B2 | 1.8293 | 0.004 | UP | PRKCA | 1.04168 | 0.041079 | UP |
| MRTO4 | -1.647 | 0.0128556 | DN | MIR3936HG | 1.8303 | 0.02053 | UP | BCL7A | 1.04538 | 0.03581 | UP |
| ZNF622 | -1.635 | 0.0177972 | DN | ASB1 | 1.8385 | 0.0132 | UP | RNF125 | 1.05051 | 0.004901 | UP |
| ARL11 | -1.625 | 0.0466081 | DN | A2ML1 | 1.8443 | 0.03193 | UP | PRXL2C | 1.05067 | 0.037163 | UP |
| PPEF2 | -1.625 | 0.0107284 | DN | PDIK1L | 1.8508 | 0.04218 | UP | AL132799.1 | 1.05067 | 0.010362 | UP |
| HELZ2 | -1.623 | 0.0389732 | DN | AC012313.8 | 1.8518 | 0.02147 | UP | CIAO1 | 1.05271 | 0.014864 | UP |
| LYZ | -1.62 | 0.035786 | DN | RMND1 | 1.8523 | 0.00572 | UP | AC010970.1 | 1.05512 | 0.012302 | UP |
| STOML2 | -1.619 | 0.0283962 | DN | SDC3 | 1.8605 | 0.03556 | UP | SOX4 | 1.05516 | 0.016065 | UP |
| PPIF | -1.617 | 0.0191483 | DN | DTD2 | 1.8689 | 0.04573 | UP | DESI2 | 1.05815 | 0.043739 | UP |
| CENPE | -1.615 | 0.0351792 | DN | ELMOD2 | 1.8709 | 0.02363 | UP | PEX1 | 1.05875 | 0.048411 | UP |
| KIAA2013 | -1.613 | 0.0015811 | DN | FOCAD | 1.8728 | 0.043 | UP | AC011447.3 | 1.06539 | 0.036059 | UP |
| GFM2 | -1.606 | 0.0498667 | DN | CFLAR-AS1 | 1.877 | 0.02929 | UP | MYSM1 | 1.07049 | 0.009189 | UP |
| BLOC1S3 | -1.606 | 0.0152638 | DN | PLXDC2 | 1.8812 | 0.00028 | UP | CCDC144A | 1.07211 | 0.014343 | UP |
| TIMP1 | -1.605 | 0.0014067 | DN | NUDT17 | 1.8832 | 0.0092 | UP | RC3H1 | 1.07799 | 0.04046 | UP |
| CEBPB | -1.602 | 0.0056116 | DN | GLDC | 1.8869 | 0.04298 | UP | ZNF274 | 1.07866 | 0.007567 | UP |

| | | | | | | | | | | | |
|---|---|---|---|---|---|---|---|---|---|---|---|
| AC141586.1 | -1.602 | 0.0274441 | DN | AC011287.1 | 1.8881 | 0.04365 | UP | BORCS7 | 1.07998 | 0.033404 | UP |
| KANK2 | -1.587 | 0.0311636 | DN | BLCAP | 1.8906 | 0.00875 | UP | PPP1R16B | 1.08914 | 0.017762 | UP |
| STRN4 | -1.58 | 0.0055646 | DN | FYB2 | 1.8917 | 0.04047 | UP | AC121333.1 | 1.09865 | 0.014872 | UP |
| NOP2 | -1.556 | 0.0069349 | DN | NOCT | 1.8968 | 0.02558 | UP | MFSD4B | 1.10591 | 0.01122 | UP |
| FAM92A | -1.555 | 0.0431119 | DN | USP54 | 1.9051 | 0.00903 | UP | OARD1 | 1.10791 | 0.030906 | UP |
| RBX1 | -1.555 | 0.0138973 | DN | CEP85 | 1.9138 | 0.00714 | UP | COPS5 | 1.11041 | 0.035115 | UP |
| GZF1 | -1.554 | 0.0022698 | DN | FAM110D | 1.9165 | 0.04123 | UP | ORMDL3 | 1.12347 | 0.039878 | UP |
| CLEC7A | -1.551 | 0.0350635 | DN | SMIM29 | 1.9167 | 0.03019 | UP | WDR3 | 1.12349 | 0.034728 | UP |
| PPARGC1B | -1.54 | 0.0452472 | DN | SMPD1 | 1.9175 | 0.01198 | UP | NKIRAS2 | 1.12406 | 0.046014 | UP |
| ATP6V0B | -1.54 | 0.0090029 | DN | CIPC | 1.9237 | 0.0239 | UP | ARL5B | 1.12728 | 0.024199 | UP |
| HSF1 | -1.533 | 0.0049665 | DN | LCAT | 1.9307 | 0.03376 | UP | RGPD5 | 1.1278 | 0.006775 | UP |
| MBD1 | -1.532 | 0.023993 | DN | ANKRD44-IT1 | 1.9307 | 0.04642 | UP | AP005901.5 | 1.13327 | 0.028608 | UP |
| ARID3A | -1.529 | 0.0474745 | DN | FO681492.1 | 1.9713 | 0.0497 | UP | CASC19 | 1.1525 | 0.018452 | UP |
| C21orf91 | -1.518 | 0.0100896 | DN | AJ009632.2 | 1.9714 | 0.04135 | UP | SLC25A32 | 1.15452 | 0.033116 | UP |
| RNH1 | -1.509 | 0.0014931 | DN | AL390071.1 | 1.9752 | 0.00981 | UP | FARSB | 1.15597 | 0.044802 | UP |
| AC018628.3 | -1.506 | 0.0093412 | DN | SDAD1P1 | 1.9758 | 0.02547 | UP | TNFRSF9 | 1.15771 | 0.021628 | UP |
| DACT3 | -1.504 | 0.0001672 | DN | LINC01237 | 2.0324 | 0.02319 | UP | ALYREF | 1.1668 | 0.034102 | UP |
| ZNF708 | -1.502 | 0.0241626 | DN | CATSPER2P1 | 2.0615 | 0.0421 | UP | AC092802.2 | 1.17068 | 0.034318 | UP |
| CCDC50 | -1.499 | 0.0054644 | DN | FAM43A | 2.0706 | 0.01984 | UP | SBDS | 1.17283 | 0.023615 | UP |
| KYAT3 | -1.496 | 0.0247018 | DN | SERPINH1 | 2.0734 | 0.02954 | UP | CCDC125 | 1.17286 | 0.02046 | UP |
| RBM42 | -1.485 | 0.034869 | DN | PRRG3 | 2.0752 | 0.01579 | UP | ERGIC2 | 1.19422 | 0.006708 | UP |
| FAM117A | -1.484 | 0.0468337 | DN | IRS1 | 2.0963 | 0.03918 | UP | RBMS3-AS3 | 1.20339 | 0.023207 | UP |
| LIN54 | -1.484 | 0.0068251 | DN | FSD1L | 2.0965 | 0.01539 | UP | AL136441.1 | 1.2038 | 0.04823 | UP |
| MPP6 | -1.478 | 0.04429 | DN | CCDC86 | 2.0967 | 0.02569 | UP | NGRN | 1.20679 | 0.024034 | UP |
| ACSM2B | -1.478 | 0.0451299 | DN | PJA1 | 2.1096 | 0.02147 | UP | AC090114.2 | 1.21314 | 0.018986 | UP |
| MIR4500HG | -1.477 | 0.0291276 | DN | GNRH1 | 2.1291 | 0.01951 | UP | USP39 | 1.21501 | 0.031789 | UP |
| SCD5 | -1.47 | 0.0469924 | DN | ARRDC1 | 2.1293 | 0.01507 | UP | STAG3L3 | 1.24016 | 0.021782 | UP |
| LINC00702 | -1.468 | 0.0216759 | DN | SH3TC1 | 2.1387 | 0.02547 | UP | LINC00882 | 1.24451 | 0.025961 | UP |
| RILPL2 | -1.463 | 0.0086997 | DN | IGSF9B | 2.1477 | 0.00925 | UP | AC135983.3 | 1.2595 | 0.010799 | UP |
| PINK1 | -1.462 | 0.040517 | DN | TMEM191C | 2.1734 | 0.01185 | UP | WDR61 | 1.25967 | 0.040598 | UP |
| KDELR1 | -1.462 | 0.0111012 | DN | DPF1 | 2.205 | 0.02599 | UP | RN7SL474P | 1.26023 | 0.047315 | UP |
| SERPINA10 | -1.462 | 0.0408194 | DN | TLE1 | 2.2058 | 0.01962 | UP | LTB4R | 1.26029 | 0.041601 | UP |
| GLUL | -1.461 | 0.0016103 | DN | OTUD7A | 2.2074 | 0.0299 | UP | ZKSCAN1 | 1.2637 | 0.006479 | UP |
| RRP12 | -1.458 | 0.0397063 | DN | SRSF9 | 2.2172 | 0.02979 | UP | RCOR1 | 1.27751 | 0.031143 | UP |
| UTP15 | -1.454 | 0.0270143 | DN | AL355312.6 | 2.2189 | 0.02569 | UP | ZNF557 | 1.28389 | 0.00677 | UP |
| AL365295.1 | -1.447 | 0.0037596 | DN | FAM71D | 2.2325 | 0.00367 | UP | EIF5A2 | 1.28603 | 0.043723 | UP |
| METTL15 | -1.447 | 0.0059681 | DN | SLC5A6 | 2.2456 | 0.03476 | UP | ASPH | 1.29049 | 0.028773 | UP |
| AC007216.3 | -1.435 | 0.0131913 | DN | AC109460.1 | 2.2482 | 0.02654 | UP | GTF2IRD2B | 1.29252 | 0.024418 | UP |
| DHTKD1 | -1.435 | 0.0396909 | DN | AC022140.1 | 2.2524 | 0.02833 | UP | AP003072.3 | 1.29562 | 0.007218 | UP |
| TET3 | -1.434 | 0.0191147 | DN | NEK2 | 2.257 | 0.03854 | UP | KBTBD6 | 1.30758 | 0.047591 | UP |
| IER5 | -1.431 | 0.0022284 | DN | XIST | 2.263 | 0.0285 | UP | LYPLA2 | 1.31166 | 0.016124 | UP |
| HMGN4 | -1.421 | 0.0104707 | DN | FERMT2 | 2.2643 | 0.00774 | UP | ANKRD10-IT1 | 1.31503 | 0.047478 | UP |
| MRPL10 | -1.419 | 0.0272945 | DN | COQ7 | 2.2687 | 0.04678 | UP | IL12RB1 | 1.32333 | 0.01985 | UP |
| MX1 | -1.41 | 0.013118 | DN | PDC-AS1 | 2.2892 | 0.01579 | UP | DDX60 | 1.32705 | 0.035202 | UP |
| RGS1 | -1.406 | 0.0085834 | DN | GOLIM4 | 2.2895 | 0.00161 | UP | ZNF555 | 1.32855 | 0.027629 | UP |
| CPQ | -1.406 | 0.0453797 | DN | AC036214.2 | 2.2904 | 0.03733 | UP | FLT1 | 1.34342 | 0.01313 | UP |
| PIM1 | -1.404 | 0.0060897 | DN | IFIT3 | 2.3026 | 0.00751 | UP | NOC2L | 1.34617 | 0.013118 | UP |
| NBPF20 | -1.403 | 0.0151876 | DN | FAR2P1 | 2.308 | 0.00579 | UP | RASSF8-AS1 | 1.34837 | 0.023012 | UP |
| BCL6 | -1.401 | 0.0304038 | DN | ZNF213 | 2.3125 | 0.03816 | UP | MRPL19 | 1.35579 | 0.028481 | UP |
| MIR4435-2HG | -1.4 | 0.0293118 | DN | AC007823.1 | 2.3256 | 0.03137 | UP | FBXO48 | 1.35647 | 0.034539 | UP |
| BLVRA | -1.396 | 0.0490332 | DN | MSMO1 | 2.334 | 0.00143 | UP | UBE2J2 | 1.36157 | 0.00943 | UP |
| ATG2A | -1.394 | 0.0026061 | DN | TTC28 | 2.3456 | 0.02516 | UP | LRRC40 | 1.36295 | 0.021132 | UP |
| VTI1B | -1.388 | 0.0330267 | DN | ADAMTS7P4 | 2.3468 | 0.0368 | UP | CLN8 | 1.36364 | 0.019362 | UP |
| COA5 | -1.382 | 0.0456738 | DN | LINC00674 | 2.3563 | 0.04382 | UP | ELP4 | 1.36734 | 0.040716 | UP |
| TAB3 | -1.382 | 0.0198438 | DN | PGAM1P8 | 2.359 | 0.01191 | UP | FXYD7 | 1.37376 | 0.039916 | UP |
| EMC8 | -1.381 | 0.0283456 | DN | BACE2 | 2.3632 | 0.0202 | UP | SCLT1 | 1.38456 | 0.00257 | UP |
| CDKN1A | -1.373 | 0.0214145 | DN | AC009563.1 | 2.364 | 0.04598 | UP | PZP | 1.38762 | 0.023414 | UP |
| WDR91 | -1.366 | 0.0489629 | DN | ZNF552 | 2.3701 | 0.0031 | UP | CHAF1B | 1.38799 | 0.010544 | UP |
| MOB1B | -1.364 | 0.0075803 | DN | ADGRG4 | 2.3737 | 0.00139 | UP | AC018647.2 | 1.38803 | 0.009629 | UP |
| TMEM70 | -1.358 | 0.0478597 | DN | LINC00888 | 2.379 | 0.0352 | UP | AC108676.1 | 1.39646 | 0.031414 | UP |
| SUCLG1 | -1.349 | 0.0050181 | DN | HNF1A-AS1 | 2.3934 | 0.00357 | UP | ASB8 | 1.39921 | 0.043037 | UP |
| PECAM1 | -1.347 | 0.0181744 | DN | SLC48A1 | 2.4081 | 0.01193 | UP | MT-ND4L | 1.39957 | 0.00313 | UP |
| E2F3 | -1.342 | 0.0122719 | DN | AC103858.3 | 2.41 | 0.02676 | UP | FBXL2 | 1.40861 | 0.024579 | UP |
| CFL2 | -1.339 | 0.0302402 | DN | SEPTIN3 | 2.4302 | 0.01237 | UP | TTPAL | 1.41125 | 0.004186 | UP |
| SCAMP4 | -1.336 | 0.0170002 | DN | CCDC85C | 2.4403 | 0.00435 | UP | FAM219B | 1.41989 | 0.029417 | UP |
| POMT1 | -1.335 | 0.0408747 | DN | WARS2 | 2.4539 | 0.02501 | UP | WEE1 | 1.42273 | 0.039531 | UP |
| FCGRT | -1.331 | 0.0402217 | DN | FP671120.8 | 2.4785 | 0.03121 | UP | ZNF677 | 1.43698 | 0.037699 | UP |
| ZNF432 | -1.319 | 0.0490047 | DN | RRAS | 2.4885 | 0.03333 | UP | AC079949.1 | 1.44259 | 0.036556 | UP |
| AF111167.1 | -1.318 | 0.0283822 | DN | AC008691.1 | 2.495 | 0.04554 | UP | EXOSC2 | 1.44318 | 0.031144 | UP |
| TAGLN | -1.316 | 0.0293906 | DN | FP325332.1 | 2.4987 | 0.02973 | UP | LINC01006 | 1.44465 | 0.037822 | UP |
| RNASEK-C17orf4 | -1.314 | 0.0124987 | DN | NDST1 | 2.5055 | 0.02752 | UP | LRP12 | 1.44623 | 0.021698 | UP |
| HLA-DPB1 | -1.312 | 0.017838 | DN | AC024560.4 | 2.5066 | 0.00945 | UP | SLC7A1 | 1.46266 | 0.02635 | UP |
| PRAL | -1.311 | 0.0023599 | DN | AC012467.2 | 2.5119 | 0.03092 | UP | LINC01524 | 1.4648 | 0.016331 | UP |
| MIDN | -1.298 | 0.0055105 | DN | AL596087.3 | 2.5294 | 0.01924 | UP | GUSB | 1.47229 | 0.001373 | UP |
| LYN | -1.296 | 0.0204639 | DN | AP001330.1 | 2.5321 | 0.02907 | UP | NDUFAB1 | 1.47278 | 0.008909 | UP |
| DPH3 | -1.296 | 0.0202884 | DN | MACC1 | 2.5377 | 0.00106 | UP | LINC01547 | 1.50317 | 0.040872 | UP |
| MED27 | -1.295 | 0.0333236 | DN | KANSL1L-AS1 | 2.5466 | 0.02523 | UP | AL354809.1 | 1.50557 | 0.002286 | UP |
| TULP4 | -1.29 | 0.0247129 | DN | GFER | 2.5483 | 0.01819 | UP | NPFFR1 | 1.50897 | 0.023457 | UP |

| | | | | | | | | | | | |
|---|---|---|---|---|---|---|---|---|---|---|---|
| MRPS15 | -1.288 | 0.0261768 | DN | PCGF1 | 2.5545 | 0.03271 | UP | ZFHX3 | 1.5132 | 0.026014 | UP |
| RNF111 | -1.28 | 0.0209047 | DN | CSRP1 | 2.5596 | 0.00176 | UP | SOCS2-AS1 | 1.51486 | 0.002923 | UP |
| TPMT | -1.28 | 0.0296224 | DN | AL359715.1 | 2.5724 | 0.04496 | UP | AL365203.2 | 1.52212 | 0.02349 | UP |
| RORA-AS1 | -1.273 | 0.0451333 | DN | DRAXIN | 2.607 | 0.01519 | UP | SPTSSA | 1.52359 | 0.034326 | UP |
| NCBP2AS2 | -1.269 | 0.0065575 | DN | AL157938.3 | 2.6117 | 0.0096 | UP | LRRC8A | 1.52842 | 0.036398 | UP |
| BOLA2 | -1.265 | 0.0170958 | DN | TCEAL8 | 2.6187 | 0.01031 | UP | PLEKHF1 | 1.53208 | 0.037797 | UP |
| RIN3 | -1.258 | 0.0393621 | DN | LINC01204 | 2.6198 | 0.00824 | UP | AC067817.2 | 1.53409 | 0.035831 | UP |
| PCGF3 | -1.254 | 0.0350427 | DN | LINC01840 | 2.644 | 0.02049 | UP | SCN2A | 1.53994 | 0.007958 | UP |
| RABEPK | -1.252 | 0.0258491 | DN | SORBS1 | 2.6445 | 0.01511 | UP | EIF2B2 | 1.54193 | 0.018141 | UP |
| AC060780.1 | -1.249 | 0.0214928 | DN | FARSA | 2.6471 | 0.02652 | UP | FCRL1 | 1.54258 | 0.015984 | UP |
| SARNP | -1.237 | 0.0263745 | DN | AC005632.3 | 2.6514 | 0.02694 | UP | AC007342.3 | 1.546 | 0.042804 | UP |
| ULK3 | -1.236 | 0.0129563 | DN | UNG | 2.6599 | 0.03236 | UP | YJU2 | 1.55317 | 0.036007 | UP |
| PPP6C | -1.222 | 0.0040661 | DN | AC073255.1 | 2.6609 | 0.03171 | UP | MAPK9 | 1.55523 | 0.027635 | UP |
| PHF10 | -1.218 | 0.0109731 | DN | AC009961.1 | 2.6716 | 0.03985 | UP | NRSN2-AS1 | 1.55563 | 0.001527 | UP |
| MTATP6P1 | -1.218 | 0.0067627 | DN | ADAMTSL4 | 2.6881 | 0.04455 | UP | ITPRID2 | 1.576 | 0.011098 | UP |
| CD74 | -1.209 | 0.0080528 | DN | LINC02511 | 2.6886 | 0.01861 | UP | LINC02068 | 1.57786 | 0.017424 | UP |
| MYL12B | -1.205 | 0.0173822 | DN | MPPED2 | 2.7565 | 0.02184 | UP | ZFYVE16 | 1.6019 | 0.03375 | UP |
| TAFA2 | -1.205 | 0.0179124 | DN | OR10A2 | 2.7696 | 0.03638 | UP | CCDC25 | 1.60475 | 0.012244 | UP |
| GABPB1 | -1.204 | 0.0320024 | DN | TCTA | 2.7836 | 0.00653 | UP | GTF2H3 | 1.60855 | 0.014026 | UP |
| PPM1K | -1.204 | 0.0014557 | DN | ZKSCAN2-DT | 2.7941 | 0.00813 | UP | LINC00299 | 1.61083 | 0.032643 | UP |
| SYT11 | -1.201 | 0.0075826 | DN | RPS6KA2 | 2.7955 | 0.03538 | UP | CSRP1 | 1.61224 | 0.049561 | UP |
| TNFRSF1B | -1.2 | 0.0030541 | DN | AC073365.1 | 2.8058 | 0.01182 | UP | AP3S2 | 1.62068 | 0.013194 | UP |
| AC010615.2 | -1.197 | 0.0158586 | DN | LINC00839 | 2.8127 | 0.03878 | UP | ANTXR2 | 1.62133 | 0.012243 | UP |
| ADNP2 | -1.196 | 0.003081 | DN | BEGAIN | 2.8227 | 0.01962 | UP | AC087359.1 | 1.62275 | 0.036319 | UP |
| ZNF654 | -1.196 | 0.0122984 | DN | IFFO2 | 2.8454 | 3.17E-05 | UP | AC016831.1 | 1.62333 | 0.040429 | UP |
| CRYM-AS1 | -1.194 | 0.0375258 | DN | CYP4F26P | 2.8576 | 0.0182 | UP | MT-TC | 1.62382 | 0.029401 | UP |
| HERC5 | -1.19 | 0.0281627 | DN | SLC5A4-AS1 | 2.8686 | 0.03193 | UP | PMS1 | 1.63187 | 0.017349 | UP |
| CCNK | -1.188 | 0.0081102 | DN | BRF2 | 2.8901 | 0.02358 | UP | AL133330.1 | 1.64548 | 0.047142 | UP |
| CUTA | -1.183 | 0.0053093 | DN | AC013460.1 | 2.912 | 0.0257 | UP | ATAD5 | 1.64725 | 0.017213 | UP |
| ASAH1 | -1.182 | 0.0113008 | DN | TMEM115 | 2.9407 | 0.03652 | UP | PWRN1 | 1.65192 | 0.039418 | UP |
| BISPR | -1.182 | 0.0288412 | DN | CXADR | 2.9423 | 0.01559 | UP | RINL | 1.65332 | 0.027287 | UP |
| TADA2B | -1.181 | 0.0468272 | DN | CEACAM1 | 2.946 | 0.00715 | UP | AP5S1 | 1.66411 | 0.018386 | UP |
| TMEM50A | -1.179 | 0.0088857 | DN | TRIM21 | 2.9547 | 0.03291 | UP | DIXDC1 | 1.66641 | 0.045386 | UP |
| PICALM | -1.178 | 0.0018756 | DN | BCDIN3D | 2.955 | 0.00575 | UP | ABI3BP | 1.66773 | 0.029953 | UP |
| SFXN2 | -1.177 | 0.0161055 | DN | SLC35G1 | 2.9678 | 0.00157 | UP | PDE5A | 1.66987 | 0.013094 | UP |
| ABR | -1.176 | 0.0379951 | DN | GAL3ST4 | 2.9727 | 0.03585 | UP | LINC01205 | 1.6734 | 0.007464 | UP |
| NEAT1 | -1.168 | 0.0002257 | DN | LRP11 | 2.9741 | 0.03878 | UP | AC008459.1 | 1.68851 | 0.042249 | UP |
| SHFL | -1.164 | 0.0141744 | DN | AC079921.1 | 2.9863 | 0.0029 | UP | GABRD | 1.69303 | 0.034971 | UP |
| MADD | -1.163 | 0.042205 | DN | ADD2 | 3.0073 | 0.02558 | UP | AL162391.1 | 1.69321 | 0.011174 | UP |
| HIF1A | -1.162 | 0.0029697 | DN | MRVI1 | 3.0119 | 0.02026 | UP | LINC01322 | 1.69355 | 0.003889 | UP |
| BHLHE40 | -1.157 | 0.0394261 | DN | AC125603.4 | 3.0179 | 0.00848 | UP | CCBE1 | 1.69706 | 0.04613 | UP |
| DENND3 | -1.156 | 0.0083983 | DN | AL355312.2 | 3.0208 | 0.01329 | UP | GHET1 | 1.70084 | 0.011106 | UP |
| MX2 | -1.155 | 0.0017585 | DN | BAIAP2L1 | 3.0605 | 0.00639 | UP | AC005858.3 | 1.71758 | 0.021479 | UP |
| RBMS2 | -1.149 | 0.0092825 | DN | AC090015.1 | 3.0635 | 0.0016 | UP | PHKA2 | 1.74086 | 0.02133 | UP |
| UQCRFS1 | -1.146 | 0.0214885 | DN | MGLL | 3.0773 | 0.00016 | UP | ZFY | 1.75041 | 0.005043 | UP |
| PSMF1 | -1.144 | 0.0447088 | DN | AC073655.2 | 3.0882 | 0.04628 | UP | GNPNAT1 | 1.75747 | 0.007261 | UP |
| NAMPT | -1.144 | 0.0156342 | DN | DCUN1D3 | 3.0957 | 9.36E-05 | UP | TACC2 | 1.77828 | 0.017774 | UP |
| TOM1 | -1.144 | 0.0228279 | DN | AC004540.1 | 3.1114 | 0.02996 | UP | GEMIN2 | 1.78205 | 0.031437 | UP |
| PPHLN1 | -1.142 | 0.0018974 | DN | AP001528.2 | 3.1148 | 0.003 | UP | RPS6KC1 | 1.78255 | 0.025714 | UP |
| RAB8B | -1.139 | 0.0065539 | DN | NKAPP1 | 3.1192 | 0.00416 | UP | TTN | 1.78481 | 2.43E-06 | UP |
| B4GALT5 | -1.138 | 0.0198804 | DN | CFAP45 | 3.1224 | 0.01426 | UP | FAM200B | 1.78549 | 0.017296 | UP |
| TMEM183A | -1.137 | 0.0187794 | DN | CDRT1 | 3.1251 | 0.04756 | UP | DNAJC14 | 1.79063 | 0.011544 | UP |
| CTSB | -1.119 | 0.0366081 | DN | SLFN12 | 3.1276 | 0.04596 | UP | LINC00649 | 1.7911 | 0.03867 | UP |
| JADE2 | -1.119 | 0.0300014 | DN | LINC02649 | 3.1679 | 0.00927 | UP | LINC00271 | 1.80128 | 0.017773 | UP |
| UBQLN1 | -1.117 | 0.0017931 | DN | RPS4XP2 | 3.1764 | 0.01868 | UP | ELP3 | 1.80413 | 0.040339 | UP |
| GTF2F1 | -1.115 | 0.0457266 | DN | H2AC6 | 3.1939 | 0.00937 | UP | AC107884.1 | 1.82065 | 0.03547 | UP |
| SENP5 | -1.114 | 0.0328581 | DN | LINC02642 | 3.1966 | 0.01535 | UP | ZNF718 | 1.82258 | 0.031948 | UP |
| AC018629.1 | -1.113 | 0.0388444 | DN | NOL3 | 3.2287 | 0.02247 | UP | AC022929.2 | 1.82897 | 0.010918 | UP |
| VCPKMT | -1.107 | 0.0259865 | DN | ZNF282 | 3.2446 | 0.01371 | UP | DCUN1D3 | 1.83916 | 0.018719 | UP |
| PPP2R2D | -1.106 | 0.0179824 | DN | SPINK2 | 3.2591 | 0.04575 | UP | TMEM107 | 1.87363 | 0.012111 | UP |
| BRI3 | -1.104 | 0.0131625 | DN | HYOU1 | 3.2767 | 0.03533 | UP | UQCC3 | 1.8748 | 0.044642 | UP |
| MRPS21 | -1.096 | 0.0312318 | DN | BNC2 | 3.2837 | 0.03022 | UP | TANC1 | 1.89176 | 0.034136 | UP |
| EDEM3 | -1.095 | 0.0487335 | DN | GRIK1 | 3.3031 | 0.02412 | UP | ARHGAP29 | 1.89583 | 0.03922 | UP |
| PARP9 | -1.089 | 0.0166609 | DN | AC079943.2 | 3.3106 | 0.01815 | UP | AP005262.1 | 1.90746 | 0.024326 | UP |
| NFKB2 | -1.089 | 0.0282078 | DN | NRIR | 3.3196 | 0.02892 | UP | MAP3K9 | 1.90802 | 0.012266 | UP |
| STX12 | -1.088 | 0.028365 | DN | ALAS1 | 3.3368 | 0.00078 | UP | U2AF1L4 | 1.90985 | 0.00023 | UP |
| GRK5 | -1.087 | 0.021675 | DN | BMP2 | 3.3495 | 0.01139 | UP | SLC6A1 | 1.92612 | 0.041174 | UP |
| CAPN15 | -1.086 | 0.0234117 | DN | STXBP5-AS1 | 3.356 | 0.01326 | UP | HNF1A-AS1 | 1.93099 | 0.019192 | UP |
| PIM3 | -1.084 | 0.016689 | DN | BAHCC1 | 3.3705 | 0.00116 | UP | ZNF461 | 1.94551 | 0.011683 | UP |
| CBX4 | -1.078 | 0.0001961 | DN | LINC01003 | 3.3736 | 0.03848 | UP | AC009135.1 | 1.94763 | 0.040032 | UP |
| GRSF1 | -1.078 | 0.0167348 | DN | KCNA7 | 3.3891 | 0.00184 | UP | ZNF829 | 1.94785 | 0.003992 | UP |
| WDR83OS | -1.075 | 0.0093564 | DN | AC108125.1 | 3.3989 | 0.02712 | UP | GUF1 | 1.94804 | 0.006937 | UP |
| CNOT11 | -1.074 | 0.0033912 | DN | RN7SL505P | 3.4017 | 0.02805 | UP | AL009031.1 | 1.95007 | 0.048013 | UP |
| MSL2 | -1.073 | 0.0382183 | DN | ICAM1 | 3.402 | 0.03122 | UP | MTRNR2L12 | 1.96119 | 0.005384 | UP |
| SPATA17 | -1.073 | 0.0284607 | DN | AC121761.1 | 3.4104 | 0.02218 | UP | AC073332.1 | 1.96569 | 0.002548 | UP |
| SAT1 | -1.07 | 0.0065134 | DN | AC093297.2 | 3.4272 | 0.01305 | UP | CORO1B | 1.9747 | 0.012517 | UP |
| GAPVD1 | -1.067 | 0.0012835 | DN | AL157932.1 | 3.4368 | 0.01066 | UP | NDUFAF4 | 1.99816 | 0.016534 | UP |
| LMF2 | -1.061 | 0.0376874 | DN | AC009087.1 | 3.439 | 0.00644 | UP | BAIAP3 | 2.0049 | 0.001549 | UP |

| | | | | | | | | | | | |
|---|---|---|---|---|---|---|---|---|---|---|---|
| NIP7 | -1.06 | 0.0406799 | DN | LANCL3 | 3.443 | 0.00965 | UP | LRRC37A4P | 2.0068 | 0.043242 | UP |
| BFAR | -1.06 | 0.0176338 | DN | AC007382.1 | 3.4477 | 0.02167 | UP | RNF24 | 2.01051 | 0.003703 | UP |
| GNAI2 | -1.058 | 0.0018695 | DN | LINC02798 | 3.4536 | 0.03902 | UP | LINC00884 | 2.01926 | 0.002499 | UP |
| PRORP | -1.054 | 0.0234765 | DN | HLA-DRB6 | 3.4692 | 0.00293 | UP | SEMA3F-AS1 | 2.02459 | 0.022973 | UP |
| FDX1 | -1.051 | 0.040757 | DN | AL109946.1 | 3.4924 | 0.00089 | UP | AC021683.3 | 2.02484 | 0.015423 | UP |
| PSMD2 | -1.048 | 0.0384928 | DN | BDKRB1 | 3.5127 | 0.00487 | UP | SCNN1G | 2.02887 | 0.032262 | UP |
| LSM7 | -1.046 | 0.028271 | DN | AC011472.1 | 3.536 | 0.04557 | UP | NUBP2 | 2.02973 | 0.0065 | UP |
| EIF2AK2 | -1.045 | 0.0043774 | DN | RN7SL159P | 3.6994 | 0.04698 | UP | MFSD11 | 2.03857 | 0.032954 | UP |
| COTL1 | -1.043 | 0.0181867 | DN | ZNF189 | 3.7346 | 0.01822 | UP | AC009303.4 | 2.03975 | 0.045628 | UP |
| NFKBIZ | -1.041 | 0.0067415 | DN | COQ9 | 3.742 | 0.00098 | UP | AGBL2 | 2.04719 | 0.041211 | UP |
| YARS1 | -1.041 | 0.0056075 | DN | RN7SL242P | 3.7815 | 0.04782 | UP | PPIB | 2.05252 | 0.003822 | UP |
| YTHDF2 | -1.038 | 0.0209457 | DN | AC016526.1 | 3.7888 | 0.00915 | UP | PRRG3 | 2.05466 | 0.019867 | UP |
| YWHAE | -1.034 | 0.0128107 | DN | STX1A | 3.7932 | 0.04697 | UP | HERC2P3 | 2.05964 | 0.024218 | UP |
| AC006064.4 | -1.032 | 0.0014054 | DN | AL157871.1 | 3.8079 | 0.01187 | UP | FAM30A | 2.06336 | 0.044099 | UP |
| MYDGF | -1.031 | 0.0460118 | DN | BASP1 | 3.8088 | 0.00394 | UP | USP32P1 | 2.08765 | 0.020108 | UP |
| HBP1 | -1.024 | 0.0352164 | DN | PDZD11 | 3.8099 | 0.03314 | UP | AL034550.2 | 2.09137 | 0.014824 | UP |
| GRIPAP1 | -1.021 | 0.0460246 | DN | AC010333.1 | 3.8359 | 0.01061 | UP | AL929472.2 | 2.0919 | 0.012341 | UP |
| ECE1 | -1.019 | 0.0268825 | DN | SLC9A3R2 | 3.8588 | 0.00245 | UP | PLPP1 | 2.10395 | 0.0406 | UP |
| MIAT | -1.006 | 0.0337708 | DN | SLC35E4 | 3.8638 | 4.35E-05 | UP | ZNF283 | 2.10477 | 0.031649 | UP |
| SNW1 | -1.005 | 0.0041957 | DN | OR4K13 | 3.9776 | 0.00112 | UP | COL4A4 | 2.10626 | 0.034508 | UP |
| PRPF40A | -1.003 | 2.11E-05 | DN | KIAA0895L | 3.9825 | 0.00125 | UP | TCFL5 | 2.11121 | 0.013941 | UP |
| KLRG1 | 1.006 | 0.0344093 | UP | INTS9 | 4.0229 | 0.00242 | UP | AC068620.3 | 2.11572 | 0.015767 | UP |
| LINC02605 | 1.0131 | 0.0485888 | UP | AC068722.1 | 4.1289 | 0.00491 | UP | AC008731.1 | 2.12703 | 0.025691 | UP |
| ST3GAL3 | 1.0138 | 0.0470925 | UP | ALG14-AS1 | 4.178 | 0.03217 | UP | SLC12A2 | 2.13155 | 0.00188 | UP |
| AC009403.2 | 1.0263 | 0.0113129 | UP | AP003168.2 | 4.2131 | 0.02297 | UP | AL136526.1 | 2.13675 | 0.022198 | UP |
| SNX1 | 1.0274 | 0.0145683 | UP | TCF19 | 4.2243 | 0.0053 | UP | AL589642.2 | 2.14319 | 0.006412 | UP |
| RB1 | 1.0286 | 0.0161482 | UP | NPHP4 | 4.2724 | 0.02358 | UP | TLCD2 | 2.15013 | 0.00262 | UP |
| GMEB1 | 1.0302 | 0.0371734 | UP | EPB41L3 | 4.2739 | 0.00139 | UP | LINC00598 | 2.1618 | 0.006645 | UP |
| TAF1 | 1.0467 | 0.0145085 | UP | CLDN7 | 4.2856 | 0.02864 | UP | TMEM25 | 2.17115 | 0.017642 | UP |
| TERF1 | 1.0482 | 0.0162174 | UP | C2orf88 | 4.3421 | 0.00685 | UP | MIER2 | 2.18202 | 0.032664 | UP |
| WASHC4 | 1.0486 | 0.0254622 | UP | CTXND1 | 4.3454 | 0.00473 | UP | ZNF839 | 2.19634 | 0.010295 | UP |
| SCAPER | 1.0499 | 0.0120613 | UP | IFITM10 | 4.4184 | 0.01437 | UP | LINC02611 | 2.20087 | 0.038286 | UP |
| CBX5 | 1.0502 | 0.0441643 | UP | UPF3AP3 | 4.4867 | 0.02043 | UP | AL034427.1 | 2.20375 | 0.019443 | UP |
| DIS3L | 1.0521 | 0.0363672 | UP | CCDC173 | 4.6566 | 0.02889 | UP | AC138207.4 | 2.20793 | 0.016116 | UP |
| HLA-C | 1.0523 | 0.0480541 | UP | PCOLCE | 5.1189 | 0.00581 | UP | EYS | 2.21192 | 0.012617 | UP |
| SSBP2 | 1.0559 | 0.0150992 | UP | AL442663.2 | 6.492 | 0.01582 | UP | AC005487.1 | 2.21918 | 0.001501 | UP |
| WAC-AS1 | 1.056 | 0.0123586 | UP | AL445235.1 | 6.9369 | 0.00099 | UP | LPCAT3 | 2.25125 | 0.030936 | UP |
| SCAMP1 | 1.0574 | 0.0411993 | UP | | | | | JPH2 | 2.25171 | 0.007836 | UP |
| A2M | 1.0664 | 0.0022882 | UP | | | | | CEP76 | 2.25186 | 0.009307 | UP |
| AGO1 | 1.0665 | 0.0447683 | UP | | | | | SYDE2 | 2.25738 | 0.01614 | UP |
| ZNF638 | 1.075 | 0.0070867 | UP | | | | | AL450992.2 | 2.26594 | 0.002303 | UP |
| OSGEP | 1.0798 | 0.0133098 | UP | | | | | FRG1-DT | 2.30599 | 0.040352 | UP |
| EIF1B-AS1 | 1.082 | 0.0093903 | UP | | | | | AC074032.1 | 2.33453 | 0.016654 | UP |
| USP32P3 | 1.0918 | 0.0017447 | UP | | | | | UTP11 | 2.34764 | 0.015048 | UP |
| COMMD3 | 1.0932 | 0.0362619 | UP | | | | | GPM6B | 2.361 | 0.008238 | UP |
| ZNF557 | 1.0933 | 0.02406 | UP | | | | | BEGAIN | 2.4036 | 0.045267 | UP |
| IBA57 | 1.1012 | 0.0494776 | UP | | | | | ITIH5 | 2.41834 | 0.025073 | UP |
| SNX13 | 1.1015 | 0.0022805 | UP | | | | | BX284613.2 | 2.42288 | 0.04083 | UP |
| MGAT5 | 1.1047 | 0.0427231 | UP | | | | | IFNW1 | 2.43917 | 0.014497 | UP |
| NGRN | 1.1153 | 0.0409525 | UP | | | | | CLMN | 2.44757 | 0.002788 | UP |
| MTM1 | 1.119 | 0.0380002 | UP | | | | | AC246817.1 | 2.45061 | 0.027403 | UP |
| ZNF429 | 1.1192 | 0.0463243 | UP | | | | | SLC9A9 | 2.45207 | 0.026874 | UP |
| POLR2J | 1.1292 | 0.0485927 | UP | | | | | NTRK3 | 2.45347 | 0.011893 | UP |
| CLCN3 | 1.1299 | 0.0404069 | UP | | | | | CEP85 | 2.4937 | 0.00083 | UP |
| BIVM | 1.1346 | 0.0372145 | UP | | | | | AL449363.2 | 2.49731 | 0.02422 | UP |
| RFX3 | 1.1379 | 0.0117975 | UP | | | | | AC079921.1 | 2.5 | 0.011469 | UP |
| ALAD | 1.139 | 0.0490117 | UP | | | | | AC105415.1 | 2.50293 | 0.047575 | UP |
| ZNF75A | 1.1408 | 0.0299802 | UP | | | | | AC099518.5 | 2.5096 | 0.013795 | UP |
| RNF125 | 1.1417 | 0.0028244 | UP | | | | | ZNF362 | 2.53679 | 0.000185 | UP |
| ESF1 | 1.1438 | 0.0168941 | UP | | | | | IQANK1 | 2.55569 | 0.037663 | UP |
| EXOC2 | 1.1523 | 0.0293347 | UP | | | | | PALB2 | 2.5714 | 0.028922 | UP |
| RRP15 | 1.1579 | 0.0236724 | UP | | | | | ATAD3C | 2.58012 | 0.004779 | UP |
| AP000704.2 | 1.1589 | 0.0029346 | UP | | | | | PMS2CL | 2.58559 | 0.031282 | UP |
| LINC00670 | 1.1627 | 0.0347187 | UP | | | | | AGAP5 | 2.61687 | 0.008228 | UP |
| ZCCHC7 | 1.1634 | 0.0352835 | UP | | | | | OR4K13 | 2.64511 | 0.023017 | UP |
| LINC02754 | 1.1726 | 0.0240142 | UP | | | | | NAALAD2 | 2.66553 | 0.042189 | UP |
| TPGS2 | 1.1735 | 0.0337158 | UP | | | | | ATP11A-AS1 | 2.72472 | 0.04813 | UP |
| CSNK1G1 | 1.175 | 0.0249482 | UP | | | | | AC087286.4 | 2.73246 | 0.018705 | UP |
| MEMO1 | 1.1813 | 0.040769 | UP | | | | | BAHCC1 | 2.75512 | 0.006348 | UP |
| AC114546.3 | 1.1846 | 0.040677 | UP | | | | | PALM2AKAP2 | 2.76394 | 0.009126 | UP |
| RAD51-AS1 | 1.1856 | 0.0467219 | UP | | | | | ADGRG4 | 2.76951 | 0.000376 | UP |
| NSMCE2 | 1.1869 | 0.0459797 | UP | | | | | AC008798.3 | 2.78255 | 0.005371 | UP |
| LSG1 | 1.1883 | 0.0132679 | UP | | | | | AC010300.1 | 2.81381 | 0.020687 | UP |
| LINC01358 | 1.1886 | 0.0341921 | UP | | | | | PAPSS2 | 2.81511 | 0.032732 | UP |
| PPP1R3E | 1.19 | 0.0353296 | UP | | | | | GPR22 | 2.83473 | 0.025619 | UP |
| DCP2 | 1.1924 | 0.0093802 | UP | | | | | C12orf60 | 2.84226 | 0.034997 | UP |
| SEPHS2 | 1.1998 | 0.0260497 | UP | | | | | PPP3R1 | 2.84507 | 0.007372 | UP |
| ARHGAP25 | 1.2012 | 0.0252473 | UP | | | | | SLC35E4 | 2.84637 | 0.001428 | UP |

| | | | | | | | | | | | |
|---|---|---|---|---|---|---|---|---|---|---|---|
| SH3YL1 | 1.2045 | 0.013898 | UP | | | | | FAM180B | 2.90796 | 0.032001 | UP |
| ZBTB38 | 1.2089 | 0.0344379 | UP | | | | | N4BP2L2-IT2 | 2.92043 | 0.004916 | UP |
| SCN2A | 1.2136 | 0.0387112 | UP | | | | | MYLK4 | 2.92902 | 0.008923 | UP |
| MIR646HG | 1.2272 | 0.0349669 | UP | | | | | CD38 | 2.94524 | 0.009574 | UP |
| AL035071.1 | 1.2275 | 0.0416691 | UP | | | | | SLC9A3R2 | 2.95426 | 0.018066 | UP |
| MSH2 | 1.2315 | 0.0222491 | UP | | | | | AC009141.1 | 2.97664 | 0.016363 | UP |
| ATP11C | 1.2321 | 0.0284831 | UP | | | | | AC024270.5 | 2.98331 | 0.015438 | UP |
| PSMG3 | 1.2355 | 0.0208898 | UP | | | | | LY86-AS1 | 2.99192 | 0.012357 | UP |
| CARD8 | 1.24 | 0.0009362 | UP | | | | | CYSLTR2 | 3.007 | 0.025166 | UP |
| AC018647.2 | 1.24 | 0.0252476 | UP | | | | | AC234031.1 | 3.05353 | 0.001148 | UP |
| SLCO3A1 | 1.2435 | 0.0007735 | UP | | | | | ANKRD44-IT1 | 3.08763 | 0.002554 | UP |
| LINC01205 | 1.2445 | 0.0495885 | UP | | | | | LINC01572 | 3.11096 | 0.008766 | UP |
| SRSF8 | 1.2483 | 0.0006782 | UP | | | | | AC021683.5 | 3.11583 | 0.037453 | UP |
| MAGED1 | 1.2526 | 0.0115422 | UP | | | | | AGAP1 | 3.14661 | 0.015229 | UP |
| ANAPC4 | 1.2559 | 0.035867 | UP | | | | | INTS9 | 3.15141 | 0.014903 | UP |
| NCK2 | 1.2704 | 0.0100026 | UP | | | | | AC007406.5 | 3.15295 | 0.003152 | UP |
| RDH14 | 1.2727 | 0.0363265 | UP | | | | | AL023882.1 | 3.15684 | 0.000679 | UP |
| ZNF770 | 1.2758 | 0.0236207 | UP | | | | | AC008056.3 | 3.17467 | 0.034729 | UP |
| GNRHR2 | 1.2799 | 0.0166318 | UP | | | | | AGAP6 | 3.17883 | 0.040106 | UP |
| OSTM1 | 1.2818 | 0.0140012 | UP | | | | | AC034111.1 | 3.29544 | 0.033599 | UP |
| ITPKB | 1.2877 | 0.0021498 | UP | | | | | CEACAM1 | 3.3408 | 0.002978 | UP |
| SLC35E2A | 1.2922 | 0.0421806 | UP | | | | | LINC00547 | 3.44142 | 0.008319 | UP |
| ERVK13-1 | 1.2923 | 0.0427073 | UP | | | | | NRIR | 3.51968 | 0.019577 | UP |
| PPP2R1A | 1.2991 | 0.0084536 | UP | | | | | AJUBA | 3.5681 | 9.40E-05 | UP |
| BCLAF3 | 1.3075 | 0.0117748 | UP | | | | | AC004076.2 | 3.59206 | 0.007147 | UP |
| NDUFB11 | 1.3092 | 0.015897 | UP | | | | | LONRF3 | 3.61792 | 0.049636 | UP |
| PLEKHH2 | 1.313 | 0.045623 | UP | | | | | AL356417.1 | 3.77715 | 0.008378 | UP |
| G6PC | 1.3162 | 0.0276175 | UP | | | | | AP000919.1 | 4.05824 | 0.013528 | UP |
| OSBPL10 | 1.3226 | 0.0165731 | UP | | | | | ZNF184 | 4.09041 | 0.000122 | UP |
| NDUFB5 | 1.3262 | 0.0058811 | UP | | | | | STX1A | 4.11061 | 0.03257 | UP |
| STK11 | 1.3277 | 0.0490857 | UP | | | | | AL442663.2 | 5.50706 | 0.039712 | UP |
| MT-ND4L | 1.331 | 0.0053726 | UP | | | | | | | | |
| AC008543.1 | 1.3351 | 0.029012 | UP | | | | | | | | |
| STK39 | 1.3379 | 0.0402016 | UP | | | | | | | | |
| PRKCQ-AS1 | 1.3484 | 0.0059875 | UP | | | | | | | | |
| AC012557.1 | 1.3492 | 0.0368029 | UP | | | | | | | | |
| CYP1B1-AS1 | 1.3562 | 0.031869 | UP | | | | | | | | |
| TCAF1 | 1.3617 | 0.0303762 | UP | | | | | | | | |
| RAD17 | 1.3708 | 0.0323956 | UP | | | | | | | | |
| FAM217B | 1.3713 | 0.0192574 | UP | | | | | | | | |
| GTF2E2 | 1.3714 | 0.0302632 | UP | | | | | | | | |
| PRPSAP2 | 1.3748 | 0.0224091 | UP | | | | | | | | |
| PCYOX1 | 1.3776 | 0.0214681 | UP | | | | | | | | |
| DUSP22 | 1.3832 | 0.045501 | UP | | | | | | | | |
| AP001992.1 | 1.3842 | 0.0431791 | UP | | | | | | | | |
| TMEM184C | 1.3843 | 0.009481 | UP | | | | | | | | |
| STAG3L3 | 1.3867 | 0.0128277 | UP | | | | | | | | |
| PRKX-AS1 | 1.3922 | 0.0434467 | UP | | | | | | | | |
| GADD45A | 1.3955 | 0.0075172 | UP | | | | | | | | |
| SETD3 | 1.3971 | 0.0394926 | UP | | | | | | | | |
| LY75 | 1.4063 | 0.0386916 | UP | | | | | | | | |
| ZNF789 | 1.4152 | 0.0103929 | UP | | | | | | | | |
| MMS22L | 1.4157 | 0.0066703 | UP | | | | | | | | |
| CACNB2 | 1.4181 | 0.0177832 | UP | | | | | | | | |
| NNT | 1.4202 | 0.0013057 | UP | | | | | | | | |
| ZFY | 1.4249 | 0.0239538 | UP | | | | | | | | |
| TOMM5 | 1.4277 | 0.0313068 | UP | | | | | | | | |
| FAM102B | 1.4294 | 0.0446706 | UP | | | | | | | | |
| TXLNA | 1.4337 | 0.0280819 | UP | | | | | | | | |
| PPP2R3C | 1.4357 | 0.0229079 | UP | | | | | | | | |
| SMIM19 | 1.4401 | 0.0245272 | UP | | | | | | | | |
| AC009487.3 | 1.4403 | 0.0286746 | UP | | | | | | | | |
| KIAA1841 | 1.4472 | 0.0270335 | UP | | | | | | | | |
| FANCL | 1.4513 | 0.0274053 | UP | | | | | | | | |
| CASC19 | 1.4526 | 0.0041908 | UP | | | | | | | | |
| AC007262.2 | 1.4609 | 0.0298551 | UP | | | | | | | | |
| LINC00882 | 1.462 | 0.0117206 | UP | | | | | | | | |
| GHET1 | 1.4768 | 0.0294555 | UP | | | | | | | | |
| OSBPL3 | 1.4872 | 0.034662 | UP | | | | | | | | |
| FSIP2 | 1.4914 | 0.0078493 | UP | | | | | | | | |
| ELK3 | 1.4964 | 0.0038487 | UP | | | | | | | | |
| IGSF8 | 1.4965 | 0.0365228 | UP | | | | | | | | |
| IRF3 | 1.4991 | 0.0143283 | UP | | | | | | | | |
| EMC1 | 1.5003 | 0.019255 | UP | | | | | | | | |
| ZNF738 | 1.5019 | 0.0392411 | UP | | | | | | | | |
| DGKH | 1.5023 | 6.11E-05 | UP | | | | | | | | |
| S100PBP | 1.504 | 0.0062882 | UP | | | | | | | | |
| WBP1L | 1.5085 | 0.0085252 | UP | | | | | | | | |

| | | | |
|---|---|---|---|
| SPECC1L | 1.5123 | 0.0323602 | UP |
| AC008555.2 | 1.5237 | 0.0079345 | UP |
| LMO7 | 1.5239 | 0.0291436 | UP |
| TSPAN18 | 1.524 | 0.0008786 | UP |
| ANO6 | 1.5249 | 0.0227015 | UP |
| CATSPER2 | 1.5358 | 0.0063 | UP |
| AC090627.1 | 1.5384 | 0.0009425 | UP |
| PGM2L1 | 1.5396 | 0.0042148 | UP |
| GALK2 | 1.5396 | 0.0158277 | UP |
| TCP11L2 | 1.5555 | 0.012028 | UP |
| AC008467.1 | 1.57 | 0.0030195 | UP |
| GDF11 | 1.5718 | 0.0450995 | UP |
| NRF1 | 1.574 | 0.0237669 | UP |
| HPS5 | 1.5754 | 0.0299061 | UP |
| CROT | 1.5777 | 0.0185806 | UP |
| AC131571.1 | 1.5815 | 0.0365637 | UP |
| PUF60 | 1.5965 | 0.0165651 | UP |
| TTC13 | 1.5971 | 0.0494411 | UP |
| SS18L2 | 1.5999 | 0.0151864 | UP |
| CASP6 | 1.601 | 0.0349786 | UP |
| PWWP3A | 1.6092 | 0.0471163 | UP |
| CDC23 | 1.6274 | 0.019987 | UP |
| KCNB1 | 1.629 | 0.0298293 | UP |
| FAM210B | 1.6319 | 0.0112853 | UP |
| RAD50 | 1.635 | 0.0028654 | UP |
| RINL | 1.6421 | 0.0328843 | UP |
| TRMT13 | 1.6427 | 0.0331263 | UP |
| ZNF354A | 1.6567 | 0.0339215 | UP |
| CAVIN1 | 1.666 | 0.024824 | UP |
| DCBLD2 | 1.6684 | 0.0478112 | UP |
| LCLAT1 | 1.6917 | 0.000386 | UP |
| MTRF1L | 1.6959 | 0.0484262 | UP |
| LINC00299 | 1.7121 | 0.026309 | UP |
| ZNF146 | 1.7125 | 0.0114566 | UP |
| SPRTN | 1.7159 | 0.0377889 | UP |
| AC104791.2 | 1.7215 | 0.0294523 | UP |
| THAP5 | 1.7363 | 0.0069742 | UP |
| ACSL5 | 1.7372 | 0.019277 | UP |
| ZDBF2 | 1.7396 | 0.0151446 | UP |
| GVINP1 | 1.7399 | 0.0092117 | UP |
| AARS2 | 1.7412 | 0.0287814 | UP |
| OGFOD1 | 1.7473 | 0.0283377 | UP |
| CEP76 | 1.7527 | 0.0438789 | UP |
| FOXL2NB | 1.7533 | 0.0419396 | UP |
| TPM2 | 1.7536 | 0.0148741 | UP |
| TNFRSF14-AS1 | 1.7542 | 0.0401933 | UP |
| TRAM2 | 1.7587 | 0.0124669 | UP |
| PYM1 | 1.7608 | 0.0220735 | UP |
| STAU2 | 1.7672 | 0.0010364 | UP |
| AC025580.2 | 1.7687 | 0.0163189 | UP |
| AL139424.2 | 1.7731 | 0.0212718 | UP |
| CLASP1 | 1.7739 | 0.0039865 | UP |
| LRRC8A | 1.7747 | 0.0184491 | UP |
| OR2T12 | 1.784 | 0.0439301 | UP |
| USP54 | 1.7916 | 0.0190183 | UP |
| FCRL3 | 1.7918 | 0.0323873 | UP |
| AC105233.4 | 1.7932 | 0.0322153 | UP |
| CHIC1 | 1.7975 | 0.045362 | UP |
| TXNDC16 | 1.8025 | 0.0450041 | UP |
| CACNA1C-AS2 | 1.8106 | 0.0166959 | UP |
| LRIG1 | 1.8179 | 0.0333428 | UP |
| SMYD4 | 1.8241 | 0.0079125 | UP |
| EBAG9 | 1.8243 | 0.0011485 | UP |
| SMIM14 | 1.8307 | 0.0003675 | UP |
| PIGC | 1.8311 | 0.0166377 | UP |
| KLHDC1 | 1.8327 | 0.0402759 | UP |
| AC139792.1 | 1.8366 | 0.0297466 | UP |
| MMP25-AS1 | 1.8429 | 0.000908 | UP |
| ZNF354B | 1.8455 | 0.0107701 | UP |
| ZNF518A | 1.8467 | 0.0005444 | UP |
| ZFHX2-AS1 | 1.8485 | 0.0032383 | UP |
| CUBN | 1.8589 | 0.045748 | UP |
| PZP | 1.8721 | 0.0035839 | UP |
| DNAJC17 | 1.887 | 0.0222467 | UP |
| ITPA | 1.8887 | 0.0103748 | UP |
| ACTR3B | 1.8966 | 0.0410882 | UP |
| JPH2 | 1.906 | 0.0268035 | UP |
| MRPL53 | 1.9128 | 0.010647 | UP |

| | | | |
|---|---|---|---|
| CYP4V2 | 1.9154 | 0.0054329 | UP |
| PTCSC3 | 1.9278 | 0.0109549 | UP |
| FAM157C | 1.9337 | 0.0318229 | UP |
| MCC | 1.9572 | 0.0188803 | UP |
| AC068620.3 | 1.9811 | 0.0270842 | UP |
| PIP4K2B | 1.9897 | 0.0015141 | UP |
| SFXN3 | 1.9962 | 0.0080731 | UP |
| NUDT17 | 1.9964 | 0.0088472 | UP |
| AC027237.4 | 2.0199 | 0.0409612 | UP |
| CYB5R1 | 2.0205 | 0.0451798 | UP |
| MEGF6 | 2.0229 | 0.0072949 | UP |
| AC243967.3 | 2.0238 | 0.0424947 | UP |
| ARLNC1 | 2.0477 | 0.0426003 | UP |
| HERC2P3 | 2.0655 | 0.02864 | UP |
| AC024610.2 | 2.0863 | 0.0445717 | UP |
| SLX4IP | 2.0997 | 0.0032919 | UP |
| LMOD3 | 2.1018 | 0.0415352 | UP |
| AC108676.1 | 2.1089 | 0.0028805 | UP |
| CCBE1 | 2.1093 | 0.0171356 | UP |
| C9orf64 | 2.1221 | 0.0319953 | UP |
| RPAP1 | 2.1245 | 0.0318784 | UP |
| ZNF718 | 2.129 | 0.0171924 | UP |
| BLCAP | 2.1443 | 0.0051731 | UP |
| AC022150.4 | 2.1451 | 0.0482214 | UP |
| FDXACB1 | 2.1461 | 0.0239502 | UP |
| ATP8B3 | 2.1506 | 0.0457939 | UP |
| AC007823.1 | 2.156 | 0.0415963 | UP |
| LINC00921 | 2.1647 | 0.029241 | UP |
| LINC01006 | 2.1691 | 0.0038576 | UP |
| TLCD2 | 2.2038 | 0.0025238 | UP |
| PACRGL | 2.2076 | 0.0201001 | UP |
| SPDYE3 | 2.2118 | 0.0426633 | UP |
| SIRT5 | 2.2402 | 0.033767 | UP |
| AC139256.2 | 2.2449 | 0.0193505 | UP |
| SPRYD3 | 2.2451 | 0.0293089 | UP |
| AP001372.2 | 2.248 | 0.0123188 | UP |
| HNF1A-AS1 | 2.262 | 0.0082988 | UP |
| DISC1-IT1 | 2.2634 | 0.04399 | UP |
| AC103874.1 | 2.2648 | 0.0334364 | UP |
| TNFRSF17 | 2.2795 | 0.0463908 | UP |
| SYDE2 | 2.2819 | 0.0185149 | UP |
| PCTP | 2.2992 | 0.041473 | UP |
| AL109946.1 | 2.3029 | 0.028594 | UP |
| EPB41L2 | 2.3267 | 0.0189372 | UP |
| AC104623.2 | 2.3297 | 0.0208063 | UP |
| LINC00649 | 2.3317 | 0.0089324 | UP |
| AC007406.5 | 2.333 | 0.0282438 | UP |
| ZNF688 | 2.334 | 0.003777 | UP |
| ARRDC5 | 2.3357 | 0.0494353 | UP |
| ZNF532 | 2.3383 | 0.0062251 | UP |
| SERPINH1 | 2.3433 | 0.0188453 | UP |
| AC005972.3 | 2.3439 | 0.0159979 | UP |
| LINC00884 | 2.3646 | 0.0007215 | UP |
| PRRG3 | 2.3826 | 0.0090432 | UP |
| SEPTIN8 | 2.3978 | 0.026877 | UP |
| TOX2 | 2.4153 | 0.0255167 | UP |
| ACE | 2.4192 | 0.0419526 | UP |
| AL929472.2 | 2.4214 | 0.0060068 | UP |
| AC138150.1 | 2.4238 | 0.0214013 | UP |
| GEMIN5 | 2.4409 | 0.0294764 | UP |
| ERVH-1 | 2.4552 | 0.0461758 | UP |
| SH3D19 | 2.4718 | 0.0064852 | UP |
| ELP3 | 2.4738 | 0.0066969 | UP |
| CEP162 | 2.4903 | 0.0061169 | UP |
| DCUN1D3 | 2.4991 | 0.002132 | UP |
| AL157886.1 | 2.5241 | 0.0014748 | UP |
| AC092910.3 | 2.5287 | 0.0121146 | UP |
| AC053545.1 | 2.5431 | 0.0097142 | UP |
| MECR | 2.566 | 0.0161904 | UP |
| DTX2P1-UPK3BP1 | 2.5795 | 0.0030527 | UP |
| PDK2 | 2.6026 | 0.0240876 | UP |
| GPR22 | 2.6443 | 0.0383409 | UP |
| MED31 | 2.6458 | 0.029158 | UP |
| CLMN | 2.663 | 0.0019716 | UP |
| BAHCC1 | 2.6716 | 0.0096217 | UP |
| BMP8B | 2.6798 | 0.0157646 | UP |
| AC005899.9 | 2.7082 | 0.008443 | UP |
| LINC02273 | 2.7118 | 0.0036674 | UP |

| | | | |
|---|---|---|---|
| AL391121.1 | 2.7536 | 0.0484863 | UP |
| COG2 | 2.7828 | 0.0018208 | UP |
| FNBP1L | 2.8073 | 0.0449528 | UP |
| ZNF184 | 2.828 | 0.0068023 | UP |
| TMEM128 | 2.8441 | 0.0104004 | UP |
| ARMCX5 | 2.8497 | 0.0283542 | UP |
| AC068282.1 | 2.8741 | 0.0154788 | UP |
| MOSPD2 | 2.882 | 0.0075706 | UP |
| LINC02604 | 2.9046 | 0.0177313 | UP |
| DYNC1I1 | 2.9086 | 0.0403901 | UP |
| RUSC1 | 2.955 | 0.0153139 | UP |
| APOL2 | 2.9971 | 0.0016915 | UP |
| NPIPP1 | 3.0213 | 0.0031077 | UP |
| MAFTRR | 3.039 | 0.0060809 | UP |
| IPO13 | 3.1269 | 0.0222368 | UP |
| SLC9A9 | 3.1375 | 0.0060192 | UP |
| GPM6B | 3.1383 | 0.000835 | UP |
| AC009299.2 | 3.1411 | 0.0338638 | UP |
| AC087286.4 | 3.1667 | 0.0082004 | UP |
| CEACAM1 | 3.1914 | 0.0051536 | UP |
| ZNF354C | 3.228 | 0.0016115 | UP |
| AP000919.1 | 3.2964 | 0.0443216 | UP |
| RASGRF2 | 3.3004 | 0.0002466 | UP |
| MAL2 | 3.399 | 0.0249221 | UP |
| ZNF81 | 3.4035 | 0.0140484 | UP |
| INTS9 | 3.4705 | 0.0087965 | UP |
| C11orf65 | 3.4778 | 0.0114508 | UP |
| AL356417.1 | 3.5255 | 0.0144683 | UP |
| AC040975.2 | 3.7168 | 0.000711 | UP |
| NPTXR | 4.1068 | 0.0007554 | UP |
| ADGRG4 | 4.1984 | 1.08E-06 | UP |

| Supplemental Table 7 \| PBMCs and plasma from University of California, San Francisco SCOPE Cohort | | | | | | | |
|---|---|---|---|---|---|---|---|
| Donors of PBMCs or plasma for flow cytometry or cytokine detection | | | | | | | |
| **Sample ID#** | **Group** | **CD4** | **CD8** | **CD4/CD8** | **CD4 Nadir** | **Viral load** | **Related Figure** |
| 1824 | HIV- | 875 | 553 | 1.582278 | 791 | N/A | Figure 6, 7, 8 |
| 1836 | HIV- | 700 | 618 | 1.132686 | 700 | N/A | Figure 6, 7, 8 |
| 1838 | HIV- | 898 | 1108 | 0.810469 | 898 | N/A | Figure 6, 7, 8 |
| 1877 | HIV- | 678 | 290 | 2.337931 | 604 | N/A | Figure 6, 7, 8 |
| 1888 | HIV- | 1112 | 148 | 7.513514 | 1112 | N/A | Figure 6, 7, 8 |
| 1922 | HIV- | 1434 | 1011 | 1.418398 | 1004 | N/A | Figure 6, 7, 8 |
| 1938 | HIV- | 553 | 1006 | 0.549702 | 408 | N/A | Figure 6, 7, 8 |
| 1940 | HIV- | 1290 | 555 | 2.324324 | 718 | N/A | Figure 6, 7, 8 |
| 1948 | HIV- | 742 | 342 | 2.169591 | 576 | N/A | Figure 6, 7, 8 |
| 1977 | HIV- | 734 | 393 | 1.867684 | 734 | N/A | Figure 6, 7, 8 |
| 1986 | HIV- | 993 | 460 | 2.158696 | 864 | N/A | Figure 6, 7, 8 |
| 6512 | HIV- | 752 | 605 | 1.242975 | 626 | N/A | Figure 6, 7, 8 |
| 6526 | HIV- | 1198 | 241 | 4.970954 | 1198 | N/A | Figure 6, 7, 8 |
| 6529 | HIV- | 1052 | 405 | 2.597531 | 1052 | N/A | Figure 6, 7, 8 |
| 6573 | HIV- | 527 | 545 | 0.966972 | 527 | N/A | Figure 6, 7, 8 |
| 6578 | HIV- | 1140 | 443 | 2.573363 | 1140 | N/A | Figure 6, 7, 8 |
| 6581 | HIV- | 444 | 354 | 1.254237 | 444 | N/A | Figure 6, 7, 8 |
| 6608 | HIV- | 832 | 497 | 1.674044 | 832 | N/A | Figure 6, 7, 8 |
| 6609 | HIV- | 1191 | 368 | 3.236413 | 1191 | N/A | Figure 6, 7, 8 |
| 6612 | HIV- | 725 | 279 | 2.598566 | 725 | N/A | Figure 6, 7, 8 |
| 1811 | HIV- | 332 | 467 | 0.710921 | 332 | N/A | Figure 6, 7, 8 |
| 1822 | HIV- | 638 | 383 | 1.665796 | 606 | N/A | Figure 6, 7 |
| 1833 | HIV- | 956 | 349 | 2.739255 | 729 | N/A | Figure 6, 7 |
| 1891 | HIV- | 829 | 281 | 2.950178 | 722 | N/A | Figure 6, 7 |
| 1930 | HIV- | 1064 | 604 | 1.761589 | 774 | N/A | Figure 6, 7 |
| 1943 | HIV- | 1729 | 707 | 2.445545 | 1543 | N/A | Figure 6, 7 |
| 1956 | HIV- | 897 | 554 | 1.619134 | 897 | N/A | Figure 6, 7 |
| 1961 | HIV- | 750 | 274 | 2.737226 | 712 | N/A | Figure 6, 7 |
| 3722 | HIV- | 509 | 198 | 2.570707 | 509 | N/A | Figure 6, 7 |
| 6513 | HIV- | 883 | 604 | 1.461921 | 676 | N/A | Figure 6, 7 |
| 6532 | HIV- | 1131 | 397 | 2.848866 | 656 | N/A | Figure 6, 7 |
| 6549 | HIV- | 1178 | 365 | 3.227397 | 884 | N/A | Figure 6, 7 |
| 6558 | HIV- | 1809 | 1186 | 1.525295 | 1809 | N/A | Figure 6, 7 |
| 6575 | HIV- | 1703 | 791 | 2.152971 | 1703 | N/A | Figure 6, 7 |
| 6576 | HIV- | 1757 | 427 | 4.114754 | 1757 | N/A | Figure 6, 7 |
| 6579 | HIV- | 1079 | 667 | 1.617691 | 1062 | N/A | Figure 6, 7 |
| 6587 | HIV- | 951 | 237 | 4.012658 | 951 | N/A | Figure 6, 7 |
| 6613 | HIV- | 1337 | 939 | 1.423855 | 1337 | N/A | Figure 6, 7 |
| 6622 | HIV- | 661 | 562 | 1.176157 | 661 | N/A | Figure 6, 7 |
| 8040 | HIV- | 550 | 664 | 0.828313 | 550 | N/A | Figure 6, 7 |
| 1038 | viremic | 178 | 819 | 0.217338 | 168 | 27846 | Figure 6, 7 |
| 1183 | viremic | 473 | 1269 | 0.372734 | 355 | 10744 | Figure 6, 7 |
| 1582 | viremic | 491 | 816 | 0.601716 | 418 | 57600 | Figure 6, 7, 8 |
| 1588 | viremic | 787 | 2306 | 0.341284 | 461 | 39200 | Figure 6, 7, 8 |
| 1602 | viremic | 601 | 412 | 1.458738 | 523 | 20684 | Figure 6, 7, 8 |
| 1628 | viremic | 432 | 475 | 0.909474 | 329 | 17944 | Figure 6, 7, 8 |
| 1636 | viremic | 914 | 1852 | 0.493521 | 500 | 65030 | Figure 6, 7, 8 |
| 1705 | viremic | 356 | 1008 | 0.353175 | 356 | 26574 | Figure 6, 7, 8 |
| 1795 | viremic | 350 | 544 | 0.643382 | 333 | 48131 | Figure 6, 7, 8 |

| | | | | | | | |
|---|---|---|---|---|---|---|---|
| 1245 | viremic | 333 | 893 | 0.3729 | 293 | 11058 | Figure 6, 7 |
| 1292 | viremic | 587 | 1887 | 0.311076 | 513 | 25725 | Figure 6, 7 |
| 3630 | viremic | 36 | 647 | 0.055641 | 36 | 145163 | Figure 6, 7, 8 |
| 3673 | viremic | 594 | 1306 | 0.454824 | 594 | 23503 | Figure 6, 7, 8 |
| 3691 | viremic | 362 | 1046 | 0.34608 | 362 | 21972 | Figure 6, 7, 8 |
| 1293 | viremic | 622 | 543 | 1.145488 | 516 | 10668 | Figure 6, 7 |
| 3731 | viremic | 195 | 608 | 0.320724 | 191 | 18261 | Figure 6, 7, 8 |
| 3734 | viremic | 60 | 560 | 0.107143 | 60 | 17179 | Figure 6, 7, 8 |
| 1324 | viremic | 588 | 874 | 0.672769 | 410 | 36954 | Figure 6, 7 |
| 1404 | viremic | 245 | 651 | 0.376344 | 191 | 23839 | Figure 6, 7 |
| 1441 | viremic | 251 | 1150 | 0.218261 | 251 | 54222 | Figure 6, 7 |
| 1053 | viremic | 308 | 1635 | 0.188379 | 308 | 180915 | Figure 6, 7 |
| 1469 | viremic | 385 | 1596 | 0.241228 | 385 | 19667 | Figure 6, 7 |
| 1614 | viremic | 606 | 1421 | 0.42646 | 606 | 122415 | Figure 6, 7 |
| 1107 | viremic | 458 | 923 | 0.496208 | 396 | 37604 | Figure 6, 7 |
| 3626 | viremic | 178 | 1059 | 0.168083 | 178 | 86380 | Figure 6, 7 |
| 3639 | viremic | 479 | 1266 | 0.378357 | 192 | 12990 | Figure 6, 7 |
| 3662 | viremic | 144 | 251 | 0.573705 | 144 | 97014 | Figure 6, 7 |
| 3675 | viremic | 310 | 1426 | 0.217391 | 228 | 104081 | Figure 6, 7 |
| 3698 | viremic | 230 | 1376 | 0.167151 | 230 | 1607407 | Figure 6, 7 |
| 4007 | viremic | 916 | 2463 | 0.371904 | 400 | 22233 | Figure 6, 7 |
| 4016 | viremic | 867 | 1902 | 0.455836 | 867 | 13600 | Figure 6, 7 |
| 1027 | viremic | 483 | 719 | 0.671766 | 388 | 89075 | Figure 6, 7, 8 |
| 1047 | viremic | 161 | 779 | 0.206675 | 161 | 168856 | Figure 6, 7, 8 |
| 1049 | viremic | 316 | 3474 | 0.090961 | 289 | 85902 | Figure 6, 7, 8 |
| 1138 | viremic | 413 | 1052 | 0.392586 | 413 | 30178 | Figure 6, 7, 8 |
| 1208 | viremic | 1227 | 2861 | 0.428871 | 877 | 19649 | Figure 6, 7, 8 |
| 1226 | viremic | 728 | 1886 | 0.386002 | 623 | 53962 | Figure 6, 7, 8 |
| 1182 | viremic | 447 | 1248 | 0.358173 | 447 | 61300 | Figure 6, 7 |
| 1375 | viremic | 685 | 1083 | 0.632502 | 617 | 33528 | Figure 6, 7, 8 |
| 1562 | viremic | 581 | 1706 | 0.340563 | 399 | 12000 | Figure 6, 7, 8 |
| 1240 | ART | 1125 | 934 | 1.204497 | 212 | <40 | Figure 6, 7 |
| 2013 | ART | 656 | 479 | 1.36952 | 13 | <40 | Figure 6, 7, 8 |
| 2032 | ART | 912 | 1726 | 0.528389 | 93 | <50 | Figure 6, 7, 8 |
| 2046 | ART | 587 | 810 | 0.724691 | 10 | <40 | Figure 6, 7, 8 |
| 2056 | ART | 956 | 875 | 1.092571 | 423 | <40 | Figure 6, 7, 8 |
| 2060 | ART | 796 | 1034 | 0.769826 | 398 | <75 | Figure 6, 7, 8 |
| 2114 | ART | 727 | 1301 | 0.558801 | 639 | <40 | Figure 6, 7, 8 |
| 2147 | ART | 478 | 927 | 0.515642 | 292 | <40 | Figure 6, 7, 8 |
| 2151 | ART | 1013 | 448 | 2.261161 | 733 | <40 | Figure 6, 7, 8 |
| 2193 | ART | 607 | 469 | 1.294243 | 606 | <40 | Figure 6, 7, 8 |
| 2616 | ART | 2139 | 1396 | 1.532235 | 1177 | <40 | Figure 6, 7, 8 |
| 2669 | ART | 802 | 1570 | 0.510828 | 205 | <40 | Figure 6, 7, 8 |
| 2676 | ART | 551 | 526 | 1.047529 | 320 | <40 | Figure 6, 7, 8 |
| 3132 | ART | 786 | 1492 | 0.52681 | 1 | <40 | Figure 6, 7, 8 |
| 3169 | ART | 605 | 2684 | 0.22541 | 26 | <40 | Figure 6, 7, 8 |
| 3223 | ART | 485 | 1175 | 0.412766 | 264 | <40 | Figure 6, 7, 8 |
| 1723 | ART | 880 | 755 | 1.165563 | 435 | <40 | Figure 6, 7 |
| 1762 | ART | 913 | 730 | 1.250685 | 193 | <40 | Figure 6, 7 |
| 1126 | ART | 1063 | 1396 | 0.761461 | 227 | <40 | Figure 6, 7 |
| 2024 | ART | 731 | 916 | 0.798035 | 102 | <40 | Figure 6, 7 |
| 2155 | ART | 1081 | 1532 | 0.705614 | 213 | <40 | Figure 6, 7 |

| | | | | | | | |
|---|---|---|---|---|---|---|---|
| 2167 | ART | 509 | 521 | 0.976967 | 241 | <40 | Figure 6, 7 |
| 2303 | ART | 853 | 700 | 1.218571 | 478 | <40 | Figure 6, 7 |
| 2459 | ART | 415 | 1234 | 0.336305 | 113 | <40 | Figure 6, 7 |
| 2472 | ART | 758 | 867 | 0.874279 | 150 | <40 | Figure 6, 7 |
| 2523 | ART | 899 | 906 | 0.992274 | 490 | <40 | Figure 6, 7 |
| 2576 | ART | 534 | 515 | 1.036893 | 276 | <40 | Figure 6, 7 |
| 2722 | ART | 1051 | 443 | 2.37246 | 143 | <40 | Figure 6, 7 |
| 3025 | ART | 368 | 1091 | 0.337305 | 39 | <40 | Figure 6, 7 |
| 3135 | ART | 837 | 543 | 1.541436 | 101 | <40 | Figure 6, 7 |
| 3641 | ART | 1342 | 553 | 2.426763 | 666 | <40 | Figure 6, 7 |
| 3672 | ART | 466 | 638 | 0.730408 | 323 | <40 | Figure 6, 7 |
| 3706 | ART | 1213 | 942 | 1.287686 | 1039 | <40 | Figure 6, 7 |
| 1151 | ART | 998 | 887 | 1.125141 | 502 | <40 | Figure 6, 7 |
| 8019 | ART | 596 | 542 | 1.099631 | 518 | <40 | Figure 6, 7 |
| 1181 | ART | 599 | 811 | 0.738594 | 211 | <40 | Figure 6, 7, 8 |
| 1228 | ART | 412 | 321 | 1.283489 | 136 | <40 | Figure 6, 7, 8 |
| 1408 | ART | 708 | 628 | 1.127389 | 357 | <40 | Figure 6, 7, 8 |
| 1440 | ART | 659 | 636 | 1.036164 | 455 | <40 | Figure 6, 7, 8 |
| 1570 | ART | 479 | 706 | 0.67847 | 149 | <40 | Figure 6, 7, 8 |
| 1635 | controller | 1433 | 1950 | 0.734872 | 900 | 388 | Figure 6, 7, 8 |
| 1637 | controller | 909 | 1082 | 0.840111 | 434 | 676 | Figure 6, 7, 8 |
| 1640 | controller | 794 | 1043 | 0.761266 | 673 | 1314 | Figure 6, 7, 8 |
| 1726 | controller | 325 | 850 | 0.382353 | 325 | 1077 | Figure 6, 7, 8 |
| 1740 | controller | 340 | 1126 | 0.301954 | 340 | 1545 | Figure 6, 7, 8 |
| 1761 | controller | 911 | 885 | 1.029379 | 911 | 1445 | Figure 6, 7, 8 |
| 1788 | controller | 768 | 710 | 1.08169 | 459 | 715 | Figure 6, 7, 8 |
| 1242 | controller | 1317 | 2443 | 0.539091 | 815 | 158 | Figure 6, 7 |
| 3604 | controller | 1273 | 1105 | 1.152036 | 738 | <40 | Figure 6, 7, 8 |
| 3679 | controller | 843 | 510 | 1.652941 | 843 | 131 | Figure 6, 7, 8 |
| 3708 | controller | 555 | 1098 | 0.505464 | 555 | 242 | Figure 6, 7, 8 |
| 3724 | controller | 375 | 341 | 1.099707 | 375 | <40 | Figure 6, 7, 8 |
| 3725 | controller | 576 | 1324 | 0.435045 | 576 | <40 | Figure 6, 7, 8 |
| 3736 | controller | 771 | 383 | 2.013055 | 771 | <40 | Figure 6, 7, 8 |
| 8002 | controller | 514 | 756 | 0.679894 | 514 | 1755 | Figure 6, 7, 8 |
| 1465 | controller | 553 | 2004 | 0.275948 | 553 | 624 | Figure 6, 7 |
| 1471 | controller | 2025 | 479 | 4.227557 | 1423 | <40 | Figure 6, 7 |
| 1508 | controller | 421 | 1195 | 0.352301 | 255 | 347 | Figure 6, 7 |
| 1509 | controller | 785 | 665 | 1.180451 | 758 | 246 | Figure 6, 7 |
| 1531 | controller | 698 | 2319 | 0.300992 | 273 | 1497 | Figure 6, 7 |
| 1545 | controller | 625 | 527 | 1.185958 | 536 | 182 | Figure 6, 7 |
| 1550 | controller | 1043 | 1140 | 0.914912 | 917 | 1149 | Figure 6, 7 |
| 1581 | controller | 648 | 671 | 0.965723 | 625 | <40 | Figure 6, 7 |
| 1620 | controller | 602 | 1229 | 0.489829 | 579 | 87 | Figure 6, 7 |
| 1731 | controller | 791 | 705 | 1.121986 | 768 | 1136 | Figure 6, 7 |
| 1748 | controller | 560 | 991 | 0.565086 | 236 | 1759 | Figure 6, 7 |
| 1749 | controller | 712 | 1807 | 0.394023 | 286 | 986 | Figure 6, 7 |
| 1775 | controller | 924 | 831 | 1.111913 | 924 | 879 | Figure 6, 7 |
| 1128 | controller | 648 | 300 | 2.16 | 133 | <40 | Figure 6, 7 |
| 3602 | controller | 734 | 546 | 1.344322 | 524 | <40 | Figure 6, 7 |
| 1145 | controller | 578 | 792 | 0.729798 | 420 | 1500 | Figure 6, 7 |
| 3664 | controller | 1751 | 911 | 1.922064 | 1274 | <40 | Figure 6, 7 |
| 3713 | controller | 666 | 597 | 1.115578 | 424 | 632 | Figure 6, 7 |

| | | | | | | | |
|---|---|---|---|---|---|---|---|
| 1161 | controller | 1149 | 1259 | 0.912629 | 694 | 1516 | Figure 6, 7 |
| 1247 | controller | 821 | 3762 | 0.218235 | 782 | 1010 | Figure 6, 7, 8 |
| 1268 | controller | 804 | 878 | 0.915718 | 804 | 1065 | Figure 6, 7, 8 |
| 1403 | controller | 654 | 1055 | 0.619905 | 493 | 776 | Figure 6, 7, 8 |
| 1527 | controller | 813 | 748 | 1.086898 | 454 | 98 | Figure 6, 7, 8 |
| 1541 | controller | 1174 | 1040 | 1.128846 | 881 | 1589 | Figure 6, 7, 8 |
| 1549 | controller | 1043 | 986 | 1.057809 | 663 | 99 | Figure 6, 7, 8 |
| N/A: Not Applicable; CD4, CD4 nadir and CD8: counts/mm3; viral load:copies/ml | | | | | | | |

| Donors of PBMCs for RNA-Seq | | | | | |
|---|---|---|---|---|---|
| **Sample ID#** | **Group** | **CD4** | **CD4 Nadi** | **Viral load** | **Related Figure** |
| 1822 | HIV- | 638 | 606 | N/A | Figure 4 |
| 1956 | HIV- | 897 | 897 | N/A | Figure 4 |
| 1986 | HIV- | 903 | 792 | N/A | Figure 4 |
| 6526 | HIV- | 1647 | 1647 | N/A | Figure 4 |
| 6532 | HIV- | 983 | 656 | N/A | Figure 4 |
| 6561 | HIV- | 664 | 453 | N/A | Figure 4 |
| 6575 | HIV- | 1703 | 1703 | N/A | Figure 4 |
| 6587 | HIV- | 951 | 951 | N/A | Figure 4 |
| 6613 | HIV- | 1337 | 1337 | N/A | Figure 4 |
| 6622 | HIV- | 661 | 661 | N/A | Figure 4 |
| 1151 | ART | 998 | 502 | <40 | Figure 4 |
| 2024 | ART | 581 | 102 | <40 | Figure 4 |
| 2032 | ART | 1259 | 93 | <50 | Figure 4 |
| 2046 | ART | 1082 | 10 | <40 | Figure 4 |
| 2151 | ART | 955 | 733 | <40 | Figure 4 |
| 2472 | ART | 758 | 150 | <40 | Figure 4 |
| 2523 | ART | 1341 | 490 | <40 | Figure 4 |
| 2576 | ART | 479 | 276 | <40 | Figure 4 |
| 2669 | ART | 906 | 205 | <40 | Figure 4 |
| 3706 | ART | 1199 | 1039 | <40 | Figure 4 |
| 1226 | viremic | 178 | 178 | 166463 | Figure 4 |
| 1531 | viremic | 607 | 273 | 3100 | Figure 4 |
| 1705 | viremic | 421 | 384 | 16733 | Figure 4 |
| 3102 | viremic | 290 | 64 | 15067 | Figure 4 |
| 3610 | viremic | 266 | 266 | 166381 | Figure 4 |
| 3626 | viremic | 151 | 151 | 86380 | Figure 4 |
| 3639 | viremic | 403 | 192 | 28085 | Figure 4 |
| 3675 | viremic | 310 | 228 | 104081 | Figure 4 |
| 4007 | viremic | 778 | 400 | >100000 | Figure 4 |
| 6007 | viremic | 586 | 221 | 27671 | Figure 4 |
| 1204 | elite controller | 701 | 701 | <40 | Figure 4 |
| 1471 | elite controller | 2025 | 1423 | <40 | Figure 4 |
| 1517 | elite controller | 1237 | 509 | <40 | Figure 4 |
| 1559 | elite controller | 599 | 557 | <40 | Figure 4 |
| 1581 | elite controller | 926 | 625 | <40 | Figure 4 |
| 1612 | elite controller | 629 | 328 | <40 | Figure 4 |
| 1783 | elite controller | 677 | 648 | <40 | Figure 4 |
| 3602 | elite controller | 734 | 524 | <40 | Figure 4 |
| 3664 | elite controller | 1751 | 1274 | <40 | Figure 4 |
| 4014 | elite controller | 1054 | 665 | <40 | Figure 4 |

N/A: Not Applicable; CD4 and CD4 nadir: counts/mm3; viral load:copies/ml

| Supplemental Table 8 \| DEgenes CD56dim vs CD56neg with or without stimulaiton | | | |
|---|---|---|---|
| CD56dim vs CD56neg stimulated | | | |
| ID | log2FC | pval | padj |
| NCOA2 | -4.574373019 | 0.000461582 | 0.070251322 |
| KATNAL1 | -4.19998043 | 7.29E-07 | 0.001889407 |
| WTH3DI | -4.151765439 | 0.000889018 | 0.094721863 |
| DEAF1 | -4.088787515 | 0.004400772 | 0.194857443 |
| PRICKLE2-AS3 | -4.048325066 | 0.001637741 | 0.118679127 |
| TSHZ2 | -4.019250593 | 3.54E-06 | 0.006886975 |
| ELMOD1 | -3.979173375 | 0.008629571 | 0.260225033 |
| TMEM65 | -3.816975569 | 0.002835059 | 0.150552867 |
| LINC01410 | -3.737321632 | 0.000260694 | 0.061925081 |
| ZCCHC4 | -3.730195868 | 0.000772605 | 0.094721863 |
| ABCB4 | -3.689496574 | 0.006182077 | 0.222669258 |
| LINC00550 | -3.686081383 | 0.002567612 | 0.149641995 |
| SRRM4 | -3.640156595 | 0.004408086 | 0.194857443 |
| MAGI3 | -3.628152916 | 0.001261697 | 0.109066697 |
| MRPL1 | -3.616290149 | 3.38E-05 | 0.026296754 |
| ZNF345 | -3.546695813 | 0.000227228 | 0.059814599 |
| MED12L | -3.537484037 | 0.001653697 | 0.118679127 |
| EBF1 | -3.502291385 | 0.001838267 | 0.12545364 |
| SLX1A | -3.449399236 | 0.000830278 | 0.094721863 |
| TBC1D32 | -3.423401698 | 0.00846731 | 0.258335956 |
| KCNAB2 | -3.41725889 | 0.000618269 | 0.084388295 |
| TUBGCP2 | -3.416571549 | 0.000401408 | 0.06644581 |
| RAD54B | -3.391780288 | 0.001150132 | 0.107977767 |
| PDE4C | -3.355387213 | 0.002577381 | 0.149641995 |
| SLC9B1 | -3.342667788 | 0.000238336 | 0.059814599 |
| ENPP1 | -3.337733308 | 0.005886207 | 0.221285493 |
| DCAF17 | -3.289380223 | 0.007267851 | 0.243533745 |
| MPP5 | -3.27841103 | 0.007711854 | 0.250730739 |
| MAP3K5 | -3.222879585 | 0.000465545 | 0.070251322 |
| CWC27 | -3.216540676 | 0.001192304 | 0.108056659 |
| PKD2 | -3.20821115 | 0.00036155 | 0.063432511 |
| CENPN | -3.15476919 | 0.005171176 | 0.210798674 |
| NFIB | -3.137551334 | 0.001232012 | 0.108056659 |
| C5orf51 | -3.136376113 | 0.009249708 | 0.271708846 |
| HMG20A | -3.125895129 | 0.001509056 | 0.118679127 |
| PTS | -3.125145562 | 0.000114309 | 0.042348826 |
| NUDT16 | -3.115152919 | 0.002650029 | 0.150552867 |
| TMEM38B | -3.101791212 | 0.001803353 | 0.125268656 |
| MCF2 | -3.095841857 | 0.006954737 | 0.239415288 |
| SLC1A2 | -3.089584472 | 0.002275257 | 0.142754057 |
| NOTCH2NL | -3.073731646 | 0.00199718 | 0.130571923 |
| HADH | -3.064868075 | 0.000316002 | 0.063361991 |
| ICE2 | -3.06326013 | 0.000994639 | 0.099208835 |
| CLVS2 | -3.042581607 | 0.008027182 | 0.250806592 |
| TSC2 | -3.026014825 | 0.004908098 | 0.20731555 |
| DCAF12 | -3.022396661 | 0.002770637 | 0.150552867 |
| CERK | -2.983090908 | 0.001151948 | 0.107977767 |

| | | | |
|---|---|---|---|
| IFT80 | -2.983052051 | 0.006623287 | 0.234223515 |
| CITED2 | -2.966572256 | 0.001985161 | 0.130571923 |
| HIC2 | -2.955884211 | 7.09E-05 | 0.034461692 |
| PDE6A | -2.951039678 | 0.001773839 | 0.124328514 |
| TBC1D1 | -2.944946584 | 0.002670126 | 0.150552867 |
| ATMIN | -2.908322875 | 0.000266994 | 0.061925081 |
| OR4F29 | -2.905585846 | 0.003200628 | 0.157600561 |
| CEP250 | -2.905560662 | 0.008931429 | 0.267255827 |
| ITGB8 | -2.903082722 | 0.007293491 | 0.243533745 |
| ZNRD1 | -2.87819771 | 0.004232744 | 0.19145783 |
| SETD1B | -2.857959968 | 0.00143048 | 0.1176218 |
| HTATSF1P2 | -2.824005866 | 0.007909036 | 0.250806592 |
| LINC01531 | -2.817961085 | 0.007986161 | 0.250806592 |
| SLC25A1 | -2.799892339 | 0.004547842 | 0.198776486 |
| P2RX5-TAX1BP3 | -2.789635875 | 0.002702319 | 0.150552867 |
| CSMD3 | -2.77600624 | 0.003739624 | 0.175266717 |
| MED15 | -2.762570098 | 0.00966822 | 0.277092077 |
| NTRK2 | -2.747296704 | 0.00576229 | 0.220840476 |
| LMNB1 | -2.742272744 | 0.001348271 | 0.11526972 |
| EPHA1-AS1 | -2.741482967 | 0.001236124 | 0.108056659 |
| PPM1D | -2.723387714 | 0.001611498 | 0.118679127 |
| PTBP2 | -2.722799133 | 0.007865334 | 0.250806592 |
| PPP4R1 | -2.722754508 | 0.00522426 | 0.210798674 |
| SPATS2L | -2.71872555 | 0.005082726 | 0.210798674 |
| CENPF | -2.711735738 | 0.006173032 | 0.222669258 |
| GHRLOS | -2.709691443 | 0.007624088 | 0.249224399 |
| CCNG2 | -2.709572288 | 0.002390075 | 0.146415644 |
| MTIF2 | -2.693448756 | 0.000829767 | 0.094721863 |
| MRPS27 | -2.686262206 | 0.005418989 | 0.210798674 |
| BAZ2B | -2.683396193 | 0.001996118 | 0.130571923 |
| BRCA2 | -2.680797758 | 0.005899592 | 0.221285493 |
| CWC22 | -2.679905137 | 0.005120609 | 0.210798674 |
| UBTF | -2.67592143 | 0.000615124 | 0.084388295 |
| METTL2B | -2.660518943 | 0.006166979 | 0.222669258 |
| BTBD1 | -2.652259547 | 0.003264117 | 0.15971591 |
| ENTPD4 | -2.63506121 | 0.001662728 | 0.118679127 |
| NRXN3 | -2.610281224 | 0.001209565 | 0.108056659 |
| MGAT4A | -2.606223833 | 0.000704739 | 0.091381213 |
| TMEM167A | -2.585150195 | 0.003126471 | 0.155922708 |
| WDR76 | -2.57487898 | 0.002553536 | 0.149641995 |
| C2CD5 | -2.559785563 | 0.001219284 | 0.108056659 |
| DIS3L | -2.510038534 | 0.002737606 | 0.150552867 |
| AFF1 | -2.503481262 | 0.001772647 | 0.124328514 |
| TTC7A | -2.50006042 | 0.00743689 | 0.247260694 |
| KCTD7 | -2.48804307 | 0.007961204 | 0.250806592 |
| CEP57 | -2.483108849 | 0.008029363 | 0.250806592 |
| CTPS1 | -2.482361191 | 0.003050842 | 0.155922708 |
| KATNBL1 | -2.468367076 | 0.002355714 | 0.14545598 |
| EBPL | -2.446301556 | 0.008750153 | 0.262842446 |
| AAAS | -2.430071069 | 0.002342768 | 0.14545598 |

| | | | |
|---|---|---|---|
| BID | -2.421542932 | 0.005335122 | 0.210798674 |
| CHD7 | -2.413002706 | 0.00183059 | 0.12545364 |
| CCDC132 | -2.400267562 | 0.000423267 | 0.068604496 |
| INPPL1 | -2.398737528 | 0.002778556 | 0.150552867 |
| NCSTN | -2.392392717 | 0.009705268 | 0.277092077 |
| KLHL6 | -2.376117498 | 0.004303972 | 0.193012594 |
| DDX55 | -2.361428268 | 0.001442232 | 0.1176218 |
| NDUFV1 | -2.327681809 | 0.005916116 | 0.221285493 |
| BEX4 | -2.322536406 | 0.008015501 | 0.250806592 |
| AATF | -2.293896996 | 0.009630234 | 0.277092077 |
| REPIN1 | -2.28928804 | 0.006546417 | 0.232562196 |
| ARL17A | -2.26416954 | 0.00522185 | 0.210798674 |
| FAM200B | -2.243692448 | 0.007129348 | 0.242624506 |
| MGRN1 | -2.223026352 | 0.003116657 | 0.155922708 |
| TMEM106B | -2.190352732 | 0.006948799 | 0.239415288 |
| PHTF2 | -2.183680895 | 5.82E-05 | 0.034074952 |
| PAPOLG | -2.182843771 | 0.009289788 | 0.271708846 |
| HAUS6 | -2.159142594 | 0.009164795 | 0.271110658 |
| ZCCHC7 | -2.155356908 | 0.004603497 | 0.199239722 |
| ECHDC1 | -2.127712545 | 0.009717736 | 0.277092077 |
| C10orf32 | -2.117867539 | 0.001553634 | 0.118679127 |
| R3HDM1 | -2.085192014 | 0.001379105 | 0.116624356 |
| KAT2B | -2.080492621 | 0.003533143 | 0.167608869 |
| TARBP1 | -2.057957723 | 0.005517559 | 0.213565203 |
| QTRTD1 | -2.057310068 | 0.00986171 | 0.280014976 |
| KIAA2013 | -2.038049476 | 0.002674278 | 0.150552867 |
| PFDN4 | -2.036117631 | 0.007607554 | 0.249224399 |
| BRI3 | -2.034051095 | 0.0008912 | 0.094721863 |
| ACOT13 | -2.006495234 | 0.003013474 | 0.155263752 |
| NCOA7 | -1.991945845 | 0.008589392 | 0.260225033 |
| ADRM1 | -1.982842453 | 0.001610033 | 0.118679127 |
| FBXO28 | -1.967519769 | 0.007734624 | 0.250730739 |
| MIDN | -1.93268293 | 0.006945622 | 0.239415288 |
| GPS2 | -1.924053697 | 0.007515819 | 0.248821575 |
| EXOC1 | -1.902493698 | 0.002510334 | 0.149087 |
| SEC14L1 | -1.898467317 | 0.006129434 | 0.222669258 |
| LIX1L | -1.893748157 | 0.007183148 | 0.242977806 |
| MSH2 | -1.863574253 | 0.005569042 | 0.214490816 |
| ZNHIT1 | -1.863244418 | 0.006388505 | 0.227993423 |
| FAM122B | -1.862509408 | 0.000954067 | 0.097110642 |
| FAM114A2 | -1.838835206 | 0.005367283 | 0.210798674 |
| GOLPH3 | -1.785596537 | 0.009487413 | 0.276449716 |
| ZMAT5 | -1.770028532 | 0.006663886 | 0.234592926 |
| GLUD1 | -1.754828702 | 0.009589609 | 0.277092077 |
| FDX1 | -1.71345281 | 0.007237362 | 0.243533745 |
| RAB1B | -1.695927743 | 0.002872728 | 0.150552867 |
| STAU1 | -1.63127619 | 0.005311828 | 0.210798674 |
| CLDND1 | -1.514086011 | 0.002774542 | 0.150552867 |
| DDX3X | -1.397380034 | 0.005915853 | 0.221285493 |
| SNRPE | -1.340107634 | 0.009119708 | 0.270806601 |

| | | | |
|---|---|---|---|
| BTF3L4 | 1.763754561 | 0.005403412 | 0.210798674 |
| CD82 | 1.99983897 | 0.000900953 | 0.094721863 |
| SNX9 | 2.209788024 | 0.006109007 | 0.222669258 |
| PIN4 | 2.218230951 | 0.00927053 | 0.271708846 |
| S1PR1 | 2.230113516 | 0.007993316 | 0.250806592 |
| PHPT1 | 2.23028656 | 0.006950476 | 0.239415288 |
| SLC35B3 | 2.337138809 | 0.006777972 | 0.237534334 |
| GSDMB | 2.378940351 | 0.004752044 | 0.203136812 |
| HMGXB4 | 2.447554157 | 0.009723154 | 0.277092077 |
| MADD | 2.660586618 | 0.005300022 | 0.210798674 |
| MINPP1 | 2.745288208 | 0.003949629 | 0.181823174 |
| STRA13 | 2.751334727 | 0.002466367 | 0.148755298 |
| PPP2R2B | 2.824257179 | 0.000550285 | 0.0778403 |
| ACAD8 | 2.864303451 | 0.000961121 | 0.097110642 |
| UHRF1BP1 | 2.9044382 | 0.006121808 | 0.222669258 |
| COQ4 | 2.90451587 | 0.004200084 | 0.191091533 |
| CKAP5 | 2.937707729 | 0.003835547 | 0.178182263 |
| LONRF3 | 2.940194209 | 0.008195718 | 0.25302652 |
| CSF2 | 2.943483613 | 0.004316734 | 0.193012594 |
| TRMT13 | 2.977152765 | 0.002857404 | 0.150552867 |
| TFB1M | 3.014793903 | 0.000877893 | 0.094721863 |
| CDK16 | 3.018932069 | 0.008015958 | 0.250806592 |
| DEXI | 3.022171611 | 0.000469546 | 0.070251322 |
| EOGT | 3.035644714 | 0.003160454 | 0.156613604 |
| ZNF32 | 3.069840028 | 0.003104903 | 0.155922708 |
| INPP5E | 3.070315489 | 0.005389096 | 0.210798674 |
| NCAM1 | 3.09697015 | 0.008234337 | 0.253213989 |
| SERPINB6 | 3.098043583 | 0.002861787 | 0.150552867 |
| GOLIM4 | 3.098136414 | 0.003428812 | 0.164667648 |
| GPM6B | 3.143729739 | 0.004664779 | 0.200508198 |
| GOLGA8B | 3.183232382 | 0.004023084 | 0.184115256 |
| RAB39B | 3.184043826 | 0.004865141 | 0.206834957 |
| N4BP3 | 3.277880735 | 0.001426511 | 0.1176218 |
| LOC101927827 | 3.280502628 | 0.003070599 | 0.155922708 |
| ATG4A | 3.331200356 | 0.002883339 | 0.150552867 |
| CXCR6 | 3.335972793 | 0.004437738 | 0.195059914 |
| SLC35A4 | 3.350752243 | 0.005310294 | 0.210798674 |
| ZNF780B | 3.360745903 | 0.000505045 | 0.074136827 |
| C10orf54 | 3.400274418 | 0.00514353 | 0.210798674 |
| ACTA2 | 3.495563389 | 0.001940421 | 0.130571923 |
| OBFC1 | 3.526080037 | 0.003365795 | 0.162645228 |
| RASSF2 | 3.546484985 | 0.002921681 | 0.151537842 |
| CTSO | 3.572996245 | 0.000854283 | 0.094721863 |
| FKTN | 3.575845975 | 0.000221541 | 0.059814599 |
| MRPS28 | 3.603762019 | 0.006221485 | 0.223056017 |
| CRNKL1 | 3.660771967 | 0.000270624 | 0.061925081 |
| LRRTM2 | 3.670169254 | 0.000321669 | 0.063361991 |
| RAB9B | 3.679644866 | 0.002499924 | 0.149087 |
| KCTD17 | 3.693339452 | 0.001563698 | 0.118679127 |
| RNF103 | 3.700188012 | 0.000527646 | 0.076020157 |

| | | | |
|---|---|---|---|
| GPATCH3 | 3.73162744 | 0.000817046 | 0.094721863 |
| KAT7 | 3.762111374 | 0.000205082 | 0.059814599 |
| KCNC2 | 3.795755253 | 0.001056715 | 0.102240823 |
| NAB2 | 3.831742796 | 0.000398788 | 0.06644581 |
| SLC35F1 | 3.836403683 | 0.007585731 | 0.249224399 |
| SS18L1 | 3.844066955 | 0.001580584 | 0.118679127 |
| PDPR | 3.867764694 | 0.003847638 | 0.178182263 |
| OTUD3 | 3.879867184 | 0.003335546 | 0.162190944 |
| LOC102724612 | 3.892669831 | 0.007018323 | 0.24053989 |
| ATP7A | 3.907208251 | 0.008605911 | 0.260225033 |
| GRIP1 | 3.927677169 | 0.000366898 | 0.063432511 |
| LINC01204 | 3.945890107 | 0.006115514 | 0.222669258 |
| TRAPPC12 | 3.990449478 | 0.000686061 | 0.090466987 |
| DHRS7B | 4.01691458 | 0.009040578 | 0.269485427 |
| TMEM140 | 4.053558606 | 0.002205939 | 0.140673842 |
| COX11 | 4.059747953 | 0.001201706 | 0.108056659 |
| TPST2 | 4.085296465 | 0.007141518 | 0.242624506 |
| ZUFSP | 4.088054778 | 0.005833758 | 0.221285493 |
| TTN-AS1 | 4.111399994 | 0.001451374 | 0.1176218 |
| ATOX1 | 4.143170278 | 0.002466508 | 0.148755298 |
| MTERF4 | 4.150396189 | 0.005999369 | 0.222669258 |
| RBKS | 4.161199388 | 0.008059338 | 0.250806592 |
| HEBP1 | 4.167774118 | 0.00811427 | 0.251510029 |
| OTUD7B | 4.236720579 | 0.001560218 | 0.118679127 |
| APP | 4.240153852 | 0.000879551 | 0.094721863 |
| LOC283299 | 4.243335974 | 0.004609659 | 0.199239722 |
| SRSF12 | 4.245978065 | 0.001064461 | 0.102240823 |
| TICAM2 | 4.282174354 | 0.00373584 | 0.175266717 |
| PPIL3 | 4.304507918 | 0.000451726 | 0.070251322 |
| F13A1 | 4.323398062 | 0.00846647 | 0.258335956 |
| C9orf85 | 4.36133184 | 0.003478244 | 0.166016792 |
| APIP | 4.388507911 | 0.000325769 | 0.063361991 |
| MX2 | 4.416913185 | 0.002136431 | 0.137367204 |
| ESR2 | 4.419153257 | 0.00492974 | 0.20731555 |
| ZSCAN26 | 4.458658412 | 0.000307498 | 0.063361991 |
| MRPL48 | 4.484873857 | 1.32E-05 | 0.017115225 |
| FLJ12825 | 4.499664036 | 0.001947907 | 0.130571923 |
| RCAN3 | 4.579627757 | 0.002229943 | 0.141048451 |
| HOTTIP | 4.579902893 | 0.005276742 | 0.210798674 |
| LDLRAP1 | 4.625830282 | 0.002048477 | 0.132809621 |
| BBS7 | 4.62921038 | 0.00285271 | 0.150552867 |
| CTNNBIP1 | 4.635779833 | 1.60E-05 | 0.01779942 |
| KAT2A | 4.679354386 | 0.000848102 | 0.094721863 |
| SLC30A4 | 4.68132514 | 0.000861767 | 0.094721863 |
| HOXB3 | 4.68730844 | 0.001519347 | 0.118679127 |
| CEP85 | 4.710936419 | 0.001045515 | 0.102240823 |
| ANKRD13B | 4.719842418 | 0.001636482 | 0.118679127 |
| IFT46 | 4.748927131 | 0.000720311 | 0.091869175 |
| KLRF1 | 4.799955815 | 5.50E-07 | 0.001889407 |
| NUAK1 | 4.802265258 | 0.002736208 | 0.150552867 |

| | | | |
|---|---|---|---|
| SPIRE1 | 4.825794069 | 0.00030899 | 0.063361991 |
| PPARGC1B | 4.845886863 | 0.000356035 | 0.063432511 |
| ANKRD26P3 | 4.875017854 | 0.000740147 | 0.092876495 |
| ZNF663P | 4.893018186 | 0.001623629 | 0.118679127 |
| LAPTM4B | 4.929501194 | 0.000113979 | 0.042348826 |
| SEC31B | 4.930352325 | 0.000345568 | 0.063432511 |
| LOC101927237 | 5.024391197 | 0.000343936 | 0.063432511 |
| DFNB31 | 5.100947692 | 0.000953456 | 0.097110642 |
| USP30 | 5.121253435 | 0.000230796 | 0.059814599 |
| CDKL1 | 5.18420668 | 0.000323342 | 0.063361991 |
| LOC101927580 | 5.203032727 | 0.000149438 | 0.052846834 |
| DUSP28 | 5.250029205 | 6.78E-05 | 0.034461692 |
| TSPAN17 | 5.258425824 | 0.00022357 | 0.059814599 |
| ST6GALNAC3 | 5.289366199 | 5.02E-05 | 0.032546001 |
| ZSCAN25 | 5.303573812 | 0.000642857 | 0.086231519 |
| CENPV | 5.31120672 | 0.000103973 | 0.042348826 |
| FAM213B | 5.346772102 | 0.000174983 | 0.05919007 |
| C9orf40 | 5.399803692 | 6.13E-05 | 0.034074952 |
| AMY2B | 5.401743364 | 0.000213631 | 0.059814599 |
| TMEM71 | 5.454462454 | 8.07E-05 | 0.036925505 |
| TMCC3 | 5.505814796 | 0.00018884 | 0.059814599 |
| GPX7 | 5.549545099 | 9.42E-05 | 0.040701525 |
| NDUFAF4P1 | 5.620158125 | 2.78E-05 | 0.024002376 |
| CDKN2C | 5.623684009 | 2.03E-05 | 0.0197352 |
| LOC653513 | 5.70639893 | 7.42E-06 | 0.011547905 |
| CDADC1 | 5.918475322 | 3.92E-05 | 0.027702229 |
| DDX19B | 6.606921338 | 2.43E-08 | 0.000189313 |
| **CD56dim vs CD56neg unstimulated** | | | |
| ID | log2FC | pval | padj |
| MKI67 | -2.745796458 | 3.14E-19 | 1.38E-15 |
| PRMT6 | -2.546009896 | 0.000942331 | 0.172564331 |
| LOC101929147 | -2.474680707 | 0.009710497 | 0.457372214 |
| TOP2A | -2.381053853 | 0.00024091 | 0.076672877 |
| MED20 | -2.352282857 | 0.002762235 | 0.282758131 |
| GGCT | -2.3081131 | 2.29E-06 | 0.00251406 |
| NDUFAF5 | -2.179486502 | 0.002736584 | 0.282758131 |
| COPS7A | -2.177094774 | 0.001141737 | 0.1858494 |
| PSMD5-AS1 | -2.103210092 | 1.20E-05 | 0.008618905 |
| IQCE | -2.09985949 | 0.002144515 | 0.261809565 |
| NTRK2 | -2.087086604 | 0.008332584 | 0.421282021 |
| ASCC1 | -1.990855656 | 3.84E-05 | 0.018771761 |
| ERCC4 | -1.989816734 | 0.003904205 | 0.308364563 |
| MIR646HG | -1.981664959 | 0.003775751 | 0.308364563 |
| TMEM65 | -1.977733661 | 0.006633048 | 0.383582209 |
| USP30 | -1.975826237 | 0.002403475 | 0.274370672 |
| ACSL1 | -1.950404296 | 0.007895115 | 0.421282021 |
| UCHL3 | -1.886184914 | 0.002464642 | 0.27774624 |
| SOX4 | -1.867001734 | 0.00323663 | 0.292036316 |

| | | | |
|---|---|---|---|
| GNAQ | -1.851776766 | 1.52E-05 | 0.009551648 |
| ASPM | -1.848661728 | 0.000376813 | 0.100369256 |
| RGS1 | -1.834923536 | 0.004110654 | 0.31148836 |
| SYNJ1 | -1.816433454 | 0.001137028 | 0.1858494 |
| RPRD1A | -1.805058287 | 0.002497561 | 0.277893223 |
| LRRC8D | -1.775651804 | 0.000797468 | 0.164202887 |
| WDR13 | -1.765317434 | 0.004824803 | 0.332582646 |
| KDM1B | -1.755293278 | 0.002260938 | 0.268562746 |
| ZBTB46 | -1.753143414 | 0.008224341 | 0.421282021 |
| XPOT | -1.742790301 | 0.004068706 | 0.31099066 |
| SCARNA9 | -1.733057804 | 9.08E-06 | 0.007981604 |
| STMN1 | -1.731697245 | 7.37E-05 | 0.029438851 |
| AGPAT5 | -1.695295525 | 0.008415882 | 0.421282021 |
| CD84 | -1.678610125 | 0.004051348 | 0.31099066 |
| ELK4 | -1.674939123 | 8.16E-07 | 0.001024543 |
| PBX2 | -1.646519158 | 0.003846883 | 0.308364563 |
| ALG2 | -1.632025672 | 0.005378858 | 0.34714397 |
| SNF8 | -1.60120578 | 0.004751989 | 0.332582646 |
| COMT | -1.565674425 | 7.08E-05 | 0.029438851 |
| ARPC1A | -1.56562915 | 0.001900861 | 0.253160157 |
| TMEM200B | -1.519385814 | 0.007373319 | 0.407177618 |
| BDH2 | -1.499835746 | 0.009676938 | 0.457372214 |
| TRG-AS1 | -1.491625687 | 0.002946685 | 0.284630375 |
| SPIN4 | -1.455389056 | 0.000815181 | 0.164202887 |
| GAR1 | -1.454031684 | 0.000289385 | 0.079490507 |
| OAS2 | -1.441400367 | 0.002722745 | 0.282758131 |
| TMTC4 | -1.431686414 | 0.003922656 | 0.308364563 |
| ARL4A | -1.427032447 | 0.008435226 | 0.421282021 |
| DCAF16 | -1.418015381 | 0.006304234 | 0.371630586 |
| STX12 | -1.416285202 | 0.000625197 | 0.14090986 |
| CHKA | -1.415806376 | 0.001655249 | 0.242493972 |
| HPS5 | -1.414632375 | 0.008160823 | 0.421282021 |
| JAGN1 | -1.38146601 | 0.009295465 | 0.456162251 |
| WEE1 | -1.35202467 | 0.005410549 | 0.34714397 |
| SF3A3 | -1.335384601 | 0.004286835 | 0.316649387 |
| CENPE | -1.304281237 | 0.007855216 | 0.421282021 |
| ARHGAP17 | -1.29579279 | 0.00320036 | 0.292036316 |
| CERS6 | -1.292141494 | 0.00306186 | 0.284686247 |
| MCM5 | -1.290219807 | 0.008355145 | 0.421282021 |
| COTL1 | -1.2547852 | 0.005983761 | 0.365258763 |
| CCDC50 | -1.251488726 | 0.004880906 | 0.332582646 |
| PAGR1 | -1.231416886 | 0.003892814 | 0.308364563 |
| MICB | -1.231144244 | 0.009412571 | 0.457107748 |
| NR2C2 | -1.163343718 | 0.007159656 | 0.402586145 |
| LGALS9 | -1.155115852 | 0.001962687 | 0.257492754 |
| TRIM8 | -1.135837986 | 0.002067247 | 0.261028582 |
| MMS19 | -1.112219023 | 0.007966997 | 0.421282021 |
| DGKQ | -1.107937145 | 0.006355779 | 0.371630586 |
| GCH1 | -1.107116637 | 0.000843799 | 0.164202887 |
| TMEM2 | -1.10390285 | 0.000260173 | 0.076672877 |

| | | | |
|---|---|---|---|
| RB1 | -1.093489518 | 0.001869973 | 0.252877936 |
| SS18 | -1.078432547 | 0.00338748 | 0.294811386 |
| COPZ1 | -1.074153346 | 0.006255884 | 0.371630586 |
| LYRM2 | -1.07304506 | 0.002343623 | 0.271058541 |
| TCF4 | -1.049941706 | 0.006384098 | 0.371630586 |
| DCTD | -1.037265949 | 0.002844323 | 0.282758131 |
| LAPTM4A | -1.000135673 | 0.002537787 | 0.278839323 |
| RASSF4 | 1.125426441 | 0.002862966 | 0.282758131 |
| IARS | 1.139514735 | 0.009612988 | 0.457372214 |
| RGS2 | 1.174842614 | 0.005039713 | 0.338160894 |
| TXNDC12 | 1.21521305 | 0.00951072 | 0.457372214 |
| AGPAT6 | 1.216702263 | 0.006778306 | 0.386891622 |
| CACNA2D2 | 1.271684771 | 0.001567267 | 0.237522011 |
| SHKBP1 | 1.322431072 | 0.002201651 | 0.265102872 |
| BZRAP1 | 1.351795032 | 0.008309828 | 0.421282021 |
| A2M | 1.368504179 | 0.0017749 | 0.246543176 |
| LNPEP | 1.393029539 | 0.005611957 | 0.354885619 |
| GPR107 | 1.502213886 | 0.001404249 | 0.220416889 |
| ICAM3 | 1.579611094 | 0.000668407 | 0.14688245 |
| CMKLR1 | 1.592415631 | 0.00103836 | 0.178964369 |
| LRRC8C | 1.595959845 | 0.003605104 | 0.307658893 |
| CEP162 | 1.596267285 | 0.001750274 | 0.246543176 |
| SPIDR | 1.602292623 | 0.004621331 | 0.331036256 |
| KCNA3 | 1.60791333 | 0.000855054 | 0.164202887 |
| LSM2 | 1.684040835 | 0.007003016 | 0.397138769 |
| MED15 | 1.702008894 | 0.009341207 | 0.456162251 |
| S1PR1 | 1.771266796 | 0.006744893 | 0.386891622 |
| IRAK2 | 1.786784898 | 0.008936023 | 0.441278871 |
| EPB41L4A | 1.828779425 | 0.007190674 | 0.402586145 |
| FAM210A | 1.837784786 | 0.008206478 | 0.421282021 |
| FAM115C | 1.944467246 | 0.004853155 | 0.332582646 |
| NFIA | 1.945533281 | 0.002108422 | 0.261028582 |
| DYNC2H1 | 1.981639478 | 0.000261682 | 0.076672877 |
| TRAF3 | 2.025352171 | 0.007543933 | 0.409328208 |
| GM2A | 2.11571981 | 0.005540353 | 0.352896396 |
| CD6 | 2.12079735 | 0.005794169 | 0.361210982 |
| METTL16 | 2.161879777 | 0.000183738 | 0.062117567 |
| ZNF626 | 2.259008559 | 0.007411652 | 0.407177618 |
| LOC100506990 | 2.263724423 | 0.004241821 | 0.316649387 |
| AMOT | 2.309982367 | 0.001480383 | 0.2282906 |
| TCP11L1 | 2.341453481 | 0.006313196 | 0.371630586 |
| AASS | 2.394645068 | 0.005354589 | 0.34714397 |
| EXPH5 | 2.402182062 | 0.00800524 | 0.421282021 |
| GPM6B | 2.438193431 | 0.005758616 | 0.361210982 |
| PTGDS | 2.468742667 | 0.000597343 | 0.138174838 |
| DGKK | 2.557598091 | 0.000275224 | 0.078039278 |
| ARAP3 | 2.619739427 | 0.003255923 | 0.292036316 |
| NUAK1 | 2.705949152 | 0.003766022 | 0.308364563 |
| ZNF578 | 2.708424969 | 0.009832386 | 0.459716333 |
| NSUN4 | 2.83955819 | 0.003893197 | 0.308364563 |

| | | | |
|---|---|---|---|
| PPM1N | 2.882753846 | 0.005934938 | 0.364811948 |
| DOCK9 | 2.889576559 | 6.98E-06 | 0.006821498 |
| SIL1 | 2.904558814 | 0.003054293 | 0.284686247 |
| LOC100132057 | 2.943804152 | 0.000522577 | 0.127595949 |
| SH3GLB2 | 2.988367733 | 0.000498178 | 0.125113872 |
| ZNF818P | 3.000852796 | 0.004669909 | 0.331036256 |
| CCDC167 | 3.023328086 | 2.19E-05 | 0.012849043 |
| AGAP1 | 3.146288558 | 1.15E-08 | 2.53E-05 |
| NCAM1 | 3.249905095 | 1.84E-21 | 1.62E-17 |
| MREG | 3.311287171 | 0.00498621 | 0.337144501 |
| ADCY9 | 3.372734955 | 0.000585913 | 0.138174838 |
| CRYBG3 | 3.377763739 | 0.00209258 | 0.261028582 |
| ARMCX4 | 3.496557335 | 6.57E-05 | 0.028855502 |
| CATSPER1 | 3.610164539 | 0.000455413 | 0.117737584 |
| PFKM | 3.630638913 | 0.002762004 | 0.282758131 |
| B3GAT1 | 3.681548385 | 3.35E-05 | 0.017345727 |
| SPIRE1 | 3.776632429 | 0.000248234 | 0.076672877 |
| MRPL24 | 3.802021798 | 4.38E-07 | 0.000641196 |
| NCKAP1 | 3.918304814 | 3.35E-05 | 0.017345727 |
| LOC100288798 | 3.994242518 | 0.003076814 | 0.284686247 |
| FABP3 | 4.094853074 | 0.006187251 | 0.371630586 |
| GORASP1 | 4.122306718 | 4.99E-05 | 0.023077839 |
| SORBS3 | 4.212948965 | 0.000938354 | 0.172564331 |
| TADA2A | 4.244993285 | 0.000163865 | 0.057986884 |
| PIGL | 4.264578425 | 1.06E-05 | 0.00848699 |
| PCBP4 | 4.451000019 | 0.001795081 | 0.246543176 |
| ZNF823 | 4.505587258 | 0.001630857 | 0.242493972 |
| ZNF540 | 4.918442571 | 0.000164923 | 0.057986884 |
| PLEKHB1 | 5.462203049 | 1.27E-05 | 0.008618905 |
| ACAT2 | 5.964026462 | 4.80E-08 | 8.44E-05 |
| TXNDC5 | 6.137288149 | 3.98E-09 | 1.17E-05 |
| **CD56$^{dim}$NK stimulated vs CD56$^{dim}$NK unstimulated** | | | |
| ID | log2FC | pval | padj |
| FCRL6 | -11.27544069 | 2.47E-36 | 1.27E-33 |
| CD160 | -9.533270642 | 4.68E-25 | 1.43E-22 |
| SIGLEC17P | -8.809108734 | 4.09E-18 | 5.14E-16 |
| RAB37 | -8.272345653 | 1.52E-22 | 3.32E-20 |
| ZNF831 | -8.258293664 | 7.08E-22 | 1.42E-19 |
| CIDEB | -7.895346162 | 5.45E-12 | 2.80E-10 |
| KRT72 | -7.786967872 | 1.45E-10 | 5.77E-09 |
| HVCN1 | -7.757739929 | 1.62E-13 | 1.09E-11 |
| TLR1 | -7.73321033 | 2.45E-13 | 1.59E-11 |
| CMKLR1 | -7.579873205 | 1.03E-15 | 9.76E-14 |
| MYOM2 | -7.561562314 | 4.54E-16 | 4.46E-14 |
| LGR6 | -7.483695257 | 2.61E-20 | 4.20E-18 |
| SCARNA9L | -7.466825422 | 1.70E-11 | 8.23E-10 |
| DBNDD2 | -7.273750855 | 3.63E-11 | 1.65E-09 |
| A2M-AS1 | -7.135284365 | 5.49E-11 | 2.45E-09 |

| | | | |
|---|---|---|---|
| GPR89B | -7.122726585 | 2.97E-11 | 1.37E-09 |
| SPTB | -7.106960105 | 1.33E-10 | 5.37E-09 |
| HIST1H2AM | -7.068069064 | 7.03E-10 | 2.47E-08 |
| RRN3P2 | -7.041066155 | 7.16E-09 | 2.07E-07 |
| NMRAL1 | -6.993907701 | 2.80E-10 | 1.08E-08 |
| RPL39L | -6.898231355 | 3.83E-09 | 1.15E-07 |
| FAIM3 | -6.843061401 | 9.54E-56 | 1.34E-52 |
| AMT | -6.839515478 | 2.98E-10 | 1.14E-08 |
| GNGT2 | -6.801692962 | 1.56E-08 | 4.20E-07 |
| PITPNM2 | -6.766698085 | 4.89E-10 | 1.78E-08 |
| TKTL1 | -6.736250366 | 1.04E-07 | 2.35E-06 |
| ZNF618 | -6.618490564 | 1.19E-10 | 4.89E-09 |
| CAPN5 | -6.609411572 | 1.79E-08 | 4.74E-07 |
| CD302 | -6.566812166 | 5.78E-08 | 1.36E-06 |
| NEIL1 | -6.551625155 | 1.69E-08 | 4.53E-07 |
| SLC39A11 | -6.551213804 | 2.33E-08 | 6.11E-07 |
| COL13A1 | -6.490271478 | 1.36E-06 | 2.25E-05 |
| ZNF570 | -6.484179049 | 2.72E-07 | 5.47E-06 |
| DFFB | -6.456661273 | 8.44E-11 | 3.65E-09 |
| PFN2 | -6.380401369 | 1.90E-06 | 3.07E-05 |
| PECAM1 | -6.353633119 | 1.02E-15 | 9.72E-14 |
| DPEP2 | -6.353203649 | 8.44E-07 | 1.49E-05 |
| KPNA5 | -6.349510026 | 3.37E-12 | 1.80E-10 |
| CTSF | -6.34696041 | 1.24E-07 | 2.72E-06 |
| ZNF296 | -6.285112174 | 3.97E-08 | 9.78E-07 |
| LOC100506123 | -6.261793308 | 7.11E-07 | 1.28E-05 |
| CACNA2D4 | -6.2477362 | 3.90E-06 | 5.70E-05 |
| GM2A | -6.203839867 | 2.78E-09 | 8.70E-08 |
| LINC01184 | -6.197835627 | 2.58E-06 | 3.99E-05 |
| PVRIG | -6.181674321 | 2.83E-17 | 3.19E-15 |
| CACNA2D2 | -6.127552305 | 1.34E-16 | 1.40E-14 |
| GPRIN3 | -6.089837582 | 6.21E-15 | 5.16E-13 |
| ANKMY2 | -6.077419194 | 1.17E-09 | 3.91E-08 |
| ZNF624 | -6.023284016 | 7.67E-06 | 0.000103499 |
| DLL1 | -5.98857786 | 6.37E-06 | 8.82E-05 |
| CEMP1 | -5.975323758 | 1.15E-06 | 1.94E-05 |
| LGALS9C | -5.967537503 | 1.26E-13 | 8.67E-12 |
| TET1 | -5.887677359 | 2.76E-07 | 5.54E-06 |
| IL18BP | -5.882001756 | 7.30E-06 | 9.94E-05 |
| SYNGR1 | -5.881042856 | 1.04E-05 | 0.000136569 |
| TK1 | -5.866723656 | 1.85E-05 | 0.000224323 |
| PDGFD | -5.859623412 | 3.25E-12 | 1.74E-10 |
| RASGRP2 | -5.834786816 | 1.42E-27 | 4.49E-25 |
| C1orf162 | -5.817207801 | 6.77E-25 | 2.01E-22 |
| GNPDA1 | -5.791608848 | 1.56E-09 | 5.15E-08 |
| USP3-AS1 | -5.770552134 | 1.24E-05 | 0.000158992 |
| METTL25 | -5.755465347 | 1.41E-05 | 0.000178114 |
| TIAF1 | -5.735006885 | 1.86E-05 | 0.000224892 |
| MBLAC2 | -5.718381684 | 3.39E-05 | 0.000381247 |
| LHPP | -5.701443789 | 3.92E-05 | 0.000433403 |

| | | | |
|---|---|---|---|
| PSMD6-AS2 | -5.63816937 | 3.47E-05 | 0.000388593 |
| AMOT | -5.631870667 | 6.39E-09 | 1.86E-07 |
| SERGEF | -5.628478977 | 1.05E-05 | 0.000137153 |
| MFSD1 | -5.609742048 | 1.50E-12 | 8.52E-11 |
| ZNF318 | -5.592952037 | 5.52E-12 | 2.82E-10 |
| LINC00426 | -5.557010833 | 8.85E-05 | 0.000880166 |
| PDE6G | -5.554460406 | 2.98E-05 | 0.000343309 |
| NAT1 | -5.520272317 | 4.82E-05 | 0.000518009 |
| BNIP3 | -5.514791452 | 7.92E-07 | 1.41E-05 |
| PLXND1 | -5.500778806 | 4.34E-08 | 1.06E-06 |
| LINC00852 | -5.492993064 | 3.93E-05 | 0.000433403 |
| UVRAG | -5.49090705 | 2.23E-13 | 1.48E-11 |
| ACSS1 | -5.479198354 | 5.08E-16 | 4.94E-14 |
| PDGFRB | -5.471324641 | 4.90E-08 | 1.18E-06 |
| NREP | -5.396545419 | 0.000178724 | 0.001616097 |
| PVRL1 | -5.392760403 | 5.21E-05 | 0.000554832 |
| TNRC6C-AS1 | -5.388535896 | 3.14E-06 | 4.70E-05 |
| CATSPER1 | -5.383411567 | 3.54E-05 | 0.000395612 |
| LOC101928517 | -5.351486938 | 7.52E-06 | 0.000101573 |
| ABCA7 | -5.324767048 | 7.21E-06 | 9.84E-05 |
| LRSAM1 | -5.316540018 | 0.000159787 | 0.001468518 |
| RGS9 | -5.297242241 | 1.88E-11 | 9.00E-10 |
| CXCR1 | -5.295762265 | 0.000204076 | 0.001800532 |
| ZNF300 | -5.287888279 | 0.000103299 | 0.001004428 |
| CCDC57 | -5.275928892 | 9.94E-05 | 0.000972547 |
| DTX3 | -5.265875472 | 0.00012716 | 0.001200738 |
| SGSH | -5.258573726 | 0.000201719 | 0.001787082 |
| PZP | -5.244171071 | 0.000202974 | 0.001794032 |
| TTLL1 | -5.237965294 | 0.000330041 | 0.002698361 |
| SLC36A1 | -5.23730097 | 0.001089461 | 0.007222098 |
| ANO9 | -5.234066771 | 0.000516905 | 0.003980659 |
| NUAK2 | -5.208692579 | 1.21E-06 | 2.02E-05 |
| MAGEE1 | -5.185330293 | 0.000263514 | 0.002236449 |
| PLTP | -5.177398487 | 3.76E-06 | 5.54E-05 |
| ABHD11 | -5.159892067 | 8.01E-05 | 0.000805529 |
| PGLS | -5.155157369 | 2.12E-05 | 0.000253703 |
| SNHG11 | -5.144101379 | 0.00022933 | 0.001977117 |
| C14orf93 | -5.091715621 | 0.000218935 | 0.001904227 |
| TP53TG1 | -5.090525892 | 0.000321697 | 0.002644515 |
| RASSF7 | -5.077392093 | 0.000889935 | 0.006131427 |
| SULT1B1 | -5.069497506 | 0.000556045 | 0.004215885 |
| CYP4F29P | -5.065829058 | 8.33E-08 | 1.91E-06 |
| RAB27B | -5.064523115 | 3.32E-09 | 1.03E-07 |
| LST1 | -5.053904811 | 0.000145132 | 0.001354798 |
| NDRG2 | -5.013860334 | 0.000694891 | 0.005046317 |
| PLXDC1 | -5.003921775 | 0.000446864 | 0.003515781 |
| TPCN2 | -4.96464551 | 0.000950536 | 0.006453779 |
| ZNF627 | -4.951669941 | 0.000440316 | 0.003481015 |
| SORBS3 | -4.924246764 | 0.000517986 | 0.003985848 |
| ZNF74 | -4.915165423 | 4.66E-05 | 0.000503496 |

| | | | |
|---|---|---|---|
| PHOSPHO2 | -4.914741755 | 0.001076121 | 0.007155652 |
| ACN9 | -4.907934691 | 0.000588142 | 0.004401418 |
| DUSP1 | -4.868832449 | 4.46E-09 | 1.33E-07 |
| C14orf159 | -4.856680059 | 6.64E-13 | 4.00E-11 |
| A2M | -4.843346213 | 5.94E-15 | 4.98E-13 |
| SNX21 | -4.832641826 | 1.18E-05 | 0.000151957 |
| ARSG | -4.824529993 | 0.001596098 | 0.009780963 |
| ZNF823 | -4.815710761 | 0.001817378 | 0.010845679 |
| SNAI3 | -4.809876267 | 0.001478169 | 0.009201372 |
| GMDS-AS1 | -4.805059223 | 0.001161632 | 0.007562554 |
| SLC22A18 | -4.800533926 | 0.000223494 | 0.001937017 |
| SBK1 | -4.788114032 | 0.000374515 | 0.003024167 |
| HERC2P9 | -4.768093267 | 2.84E-08 | 7.28E-07 |
| ECSIT | -4.742488025 | 0.001091965 | 0.007233808 |
| GALNT12 | -4.74016673 | 0.001600901 | 0.009804271 |
| COMTD1 | -4.730774667 | 0.001454601 | 0.009098882 |
| GSTM1 | -4.728191538 | 5.65E-07 | 1.05E-05 |
| DIAPH2 | -4.724606375 | 4.40E-10 | 1.63E-08 |
| SLC25A14 | -4.724529718 | 0.002921891 | 0.016113924 |
| FOS | -4.710057281 | 1.44E-14 | 1.15E-12 |
| TMEM86B | -4.707206531 | 0.001778959 | 0.010661802 |
| ARMCX4 | -4.700705874 | 6.61E-06 | 9.04E-05 |
| LOC100288798 | -4.680407932 | 0.001318671 | 0.008406423 |
| ARMC7 | -4.663889993 | 0.002020041 | 0.011867441 |
| BTBD11 | -4.663729048 | 3.02E-05 | 0.000346064 |
| ZNF34 | -4.649915928 | 0.000196601 | 0.001747152 |
| MANBA | -4.647257535 | 9.06E-07 | 1.59E-05 |
| PLEKHG4 | -4.646180126 | 0.002069777 | 0.012080059 |
| CD300A | -4.643506452 | 1.15E-13 | 8.02E-12 |
| CCDC85C | -4.637947607 | 0.000431842 | 0.003422297 |
| PER1 | -4.598018279 | 1.14E-06 | 1.94E-05 |
| ANO8 | -4.59651517 | 1.77E-05 | 0.00021626 |
| PATL2 | -4.591599707 | 3.67E-07 | 7.16E-06 |
| FRMD4A | -4.590923068 | 0.001839203 | 0.010942644 |
| LOC101928020 | -4.586203014 | 2.18E-05 | 0.000260359 |
| VNN2 | -4.583617252 | 1.07E-06 | 1.84E-05 |
| MYO18A | -4.577025136 | 0.004174353 | 0.02136117 |
| HIST1H2BN | -4.566141301 | 0.003883621 | 0.020117322 |
| TEF | -4.557131176 | 0.003164463 | 0.017077306 |
| RCCD1 | -4.556418392 | 0.002853697 | 0.015787213 |
| HIST1H1A | -4.52509237 | 0.006639257 | 0.03063864 |
| PLEKHM1P | -4.515556356 | 0.001520177 | 0.009380159 |
| LRFN1 | -4.51153622 | 0.00352813 | 0.018621004 |
| RAD9A | -4.489385012 | 0.000153157 | 0.001418908 |
| MIR646HG | -4.481631209 | 0.001146182 | 0.007496791 |
| SPON2 | -4.477495911 | 1.57E-06 | 2.57E-05 |
| FAM8A1 | -4.467517783 | 1.60E-09 | 5.25E-08 |
| RNVU1-19 | -4.460231389 | 0.000152426 | 0.001414805 |
| PTPRN2 | -4.458962938 | 0.004052134 | 0.020858072 |
| CLEC11A | -4.447271371 | 0.005041587 | 0.024719147 |

| | | | |
|---|---|---|---|
| ITGB2 | -4.432853645 | 2.76E-36 | 1.29E-33 |
| FBXW4P1 | -4.432643823 | 0.000168722 | 0.001538409 |
| TPM2 | -4.387715751 | 0.000101841 | 0.000993207 |
| PLEKHG3 | -4.386986177 | 0.000374331 | 0.003024167 |
| CCDC136 | -4.367100962 | 0.009970443 | 0.042049136 |
| LINC00339 | -4.346035891 | 0.005820001 | 0.027660249 |
| RASSF1 | -4.336917178 | 2.48E-11 | 1.16E-09 |
| PCOLCE | -4.33166409 | 0.00560808 | 0.026813289 |
| ZCWPW1 | -4.327908272 | 0.002595634 | 0.014610307 |
| ZNF577 | -4.312947698 | 2.65E-05 | 0.000309925 |
| CXCR2 | -4.305581155 | 6.90E-08 | 1.61E-06 |
| LOC100130872 | -4.289969407 | 8.69E-11 | 3.74E-09 |
| C12orf66 | -4.28085743 | 0.001306338 | 0.008344062 |
| PTPRCAP | -4.262164619 | 4.20E-13 | 2.61E-11 |
| IFITM3 | -4.256090478 | 4.58E-12 | 2.39E-10 |
| APOBR | -4.245143654 | 9.64E-21 | 1.66E-18 |
| OSBPL5 | -4.234660593 | 1.09E-10 | 4.52E-09 |
| PSKH1 | -4.214671451 | 0.000209937 | 0.001839011 |
| FMN1 | -4.204817947 | 0.002154102 | 0.012431701 |
| ASB14 | -4.19585647 | 0.001036466 | 0.006936406 |
| HIBCH | -4.194265677 | 9.68E-06 | 0.000128345 |
| DECR2 | -4.187765718 | 0.009951779 | 0.042012438 |
| NRROS | -4.18584701 | 7.82E-05 | 0.000789704 |
| NTHL1 | -4.18480802 | 0.009690806 | 0.041140848 |
| APBA2 | -4.182721049 | 0.000122179 | 0.001159286 |
| UBA7 | -4.17852724 | 2.16E-07 | 4.49E-06 |
| TTLL3 | -4.173576841 | 6.64E-07 | 1.21E-05 |
| USP43 | -4.165186863 | 0.003074729 | 0.016740379 |
| PIGL | -4.16388901 | 2.54E-05 | 0.000299128 |
| RNF166 | -4.15295979 | 2.86E-09 | 8.92E-08 |
| ZNF329 | -4.1332098 | 0.002572576 | 0.014505487 |
| MYOM1 | -4.128800771 | 0.00215805 | 0.012447163 |
| KCNAB2 | -4.117766447 | 1.62E-14 | 1.28E-12 |
| RHOBTB2 | -4.113538933 | 0.000706796 | 0.005113845 |
| ATXN7L1 | -4.11024728 | 1.69E-05 | 0.000207511 |
| CX3CR1 | -4.107922909 | 7.17E-13 | 4.27E-11 |
| FGR | -4.096741713 | 1.74E-33 | 7.77E-31 |
| TBC1D10C | -4.092988539 | 1.65E-13 | 1.10E-11 |
| ZNF711 | -4.091152043 | 9.10E-05 | 0.000900449 |
| TCEAL1 | -4.085595704 | 2.38E-05 | 0.000280923 |
| TMC8 | -4.083738363 | 2.94E-37 | 1.70E-34 |
| LINC00263 | -4.066717225 | 0.000213483 | 0.001865076 |
| SLC25A53 | -4.062980933 | 0.000627421 | 0.004624811 |
| CD52 | -4.058981432 | 6.58E-12 | 3.33E-10 |
| TSPAN32 | -4.050509033 | 0.004473506 | 0.022484407 |
| CTBP1-AS2 | -4.049660901 | 1.68E-05 | 0.000205755 |
| ZBTB46 | -4.03180519 | 0.001066659 | 0.00710454 |
| MROH1 | -4.026452361 | 0.002169971 | 0.01249389 |
| L3HYPDH | -4.019103119 | 0.000324154 | 0.002659094 |
| MAP4K2 | -4.016749392 | 0.000433557 | 0.003430898 |

| | | | |
|---|---|---|---|
| KIR3DL1 | -4.016665595 | 2.95E-08 | 7.45E-07 |
| C3orf35 | -4.007584861 | 0.001292836 | 0.008273982 |
| PCED1A | -3.996531598 | 1.08E-06 | 1.85E-05 |
| KLF2 | -3.995631163 | 0.002182349 | 0.012557789 |
| EFHC1 | -3.978060849 | 3.66E-06 | 5.42E-05 |
| DOCK5 | -3.975797952 | 2.92E-06 | 4.43E-05 |
| TTC16 | -3.973689354 | 0.006696423 | 0.030815482 |
| VPS26B | -3.97199384 | 1.41E-08 | 3.80E-07 |
| CPQ | -3.964231554 | 2.84E-07 | 5.67E-06 |
| GSN-AS1 | -3.926512122 | 0.005525162 | 0.026507267 |
| LY75 | -3.925031747 | 8.34E-13 | 4.87E-11 |
| NME8 | -3.921189487 | 0.000177517 | 0.001609627 |
| NAPB | -3.889000626 | 0.00020712 | 0.00182084 |
| SPTBN1 | -3.880444218 | 7.25E-07 | 1.30E-05 |
| TRG-AS1 | -3.870462126 | 1.23E-07 | 2.72E-06 |
| HIST1H1D | -3.860684822 | 9.98E-07 | 1.73E-05 |
| LRGUK | -3.852191898 | 0.000375471 | 0.003026905 |
| AKT1S1 | -3.852114254 | 0.007929731 | 0.035330877 |
| SAMHD1 | -3.849934317 | 2.87E-05 | 0.000333341 |
| CEP89 | -3.846893431 | 0.001366879 | 0.008635192 |
| TMEM229B | -3.845816198 | 0.000182946 | 0.001639167 |
| HOTAIRM1 | -3.844441403 | 2.18E-06 | 3.44E-05 |
| PET117 | -3.841798316 | 0.000179808 | 0.001624396 |
| TRIB3 | -3.835177039 | 0.003806017 | 0.019872717 |
| ADCK3 | -3.830935724 | 5.72E-06 | 8.06E-05 |
| HSDL1 | -3.820911774 | 0.000608649 | 0.004506756 |
| CHI3L2 | -3.80573296 | 0.00469178 | 0.023293977 |
| CECR1 | -3.789275377 | 4.34E-07 | 8.26E-06 |
| FAN1 | -3.779339209 | 0.006225406 | 0.029070663 |
| SLC25A45 | -3.773090016 | 2.59E-06 | 4.00E-05 |
| ARHGAP12 | -3.770831281 | 1.40E-07 | 3.03E-06 |
| RASA3 | -3.766726665 | 7.60E-11 | 3.34E-09 |
| KRT73 | -3.755607261 | 0.005700801 | 0.027190356 |
| PXYLP1 | -3.746811143 | 0.001270098 | 0.008144399 |
| LRIG2 | -3.740923188 | 5.47E-05 | 0.000579495 |
| RAB11FIP4 | -3.730937387 | 1.92E-06 | 3.09E-05 |
| ZNF668 | -3.713282269 | 0.003299696 | 0.01770969 |
| ZNF682 | -3.705172655 | 0.000449579 | 0.003534309 |
| CARNS1 | -3.701348178 | 6.71E-05 | 0.000688805 |
| CD300C | -3.689146977 | 0.007174466 | 0.032699084 |
| CDKN1C | -3.680579479 | 0.005059435 | 0.024757279 |
| MGAM | -3.65943228 | 0.003189938 | 0.017186427 |
| RNU4ATAC | -3.64891939 | 0.005862285 | 0.027811837 |
| GRK5 | -3.643639282 | 8.46E-07 | 1.49E-05 |
| RNASET2 | -3.640445015 | 1.77E-05 | 0.00021626 |
| VPS8 | -3.63693466 | 1.26E-05 | 0.000161136 |
| THTPA | -3.621022045 | 0.009494817 | 0.040607519 |
| ADAMTS10 | -3.617284431 | 4.80E-06 | 6.87E-05 |
| DLG3 | -3.61431864 | 5.61E-05 | 0.000589578 |
| ZNF276 | -3.607345536 | 1.40E-20 | 2.33E-18 |

| | | | |
|---|---|---|---|
| CXCR5 | -3.607144565 | 0.001171847 | 0.007608863 |
| LOC440434 | -3.596354064 | 3.31E-07 | 6.51E-06 |
| TBC1D10A | -3.592257994 | 0.001035751 | 0.006936349 |
| CMC4 | -3.588934312 | 0.001223335 | 0.00790134 |
| FAM179A | -3.584022569 | 0.00052795 | 0.00404034 |
| TSPAN2 | -3.581643542 | 1.47E-05 | 0.000183151 |
| SYTL1 | -3.578074757 | 4.73E-15 | 4.07E-13 |
| ANXA2R | -3.577715033 | 6.05E-07 | 1.11E-05 |
| TMEM107 | -3.558617828 | 4.82E-06 | 6.88E-05 |
| DDX58 | -3.554193641 | 7.48E-06 | 0.000101416 |
| MZF1 | -3.553175473 | 0.009610866 | 0.04096148 |
| ZBP1 | -3.550508604 | 7.42E-07 | 1.33E-05 |
| ITGB2-AS1 | -3.544027398 | 0.001230536 | 0.007937399 |
| PPM1N | -3.509120756 | 0.001358715 | 0.00860023 |
| ICAM3 | -3.508784958 | 1.20E-05 | 0.000153994 |
| PLA2G4B | -3.507223254 | 0.001158017 | 0.007549042 |
| LOC728392 | -3.503200321 | 0.000112784 | 0.001082707 |
| CLCN6 | -3.483033299 | 0.001301666 | 0.008319641 |
| AHNAK | -3.481318598 | 1.92E-39 | 1.71E-36 |
| LOC653160 | -3.437787882 | 0.009638086 | 0.041019615 |
| PDZK1 | -3.437753082 | 0.002080189 | 0.012119201 |
| ARHGAP4 | -3.429829732 | 7.82E-07 | 1.39E-05 |
| ZAP70 | -3.40490839 | 2.45E-08 | 6.40E-07 |
| EDC3 | -3.402640829 | 0.006838193 | 0.031364896 |
| RASSF4 | -3.401144946 | 2.75E-11 | 1.28E-09 |
| MELK | -3.400168011 | 0.003392165 | 0.018092388 |
| KIR3DL2 | -3.398806568 | 0.001457795 | 0.009101105 |
| IFITM1 | -3.398380922 | 1.77E-18 | 2.38E-16 |
| MRPL1 | -3.396751807 | 5.67E-10 | 2.04E-08 |
| TNFSF13B | -3.39253993 | 0.003245842 | 0.017439737 |
| TTC38 | -3.38875964 | 2.31E-06 | 3.62E-05 |
| PLCH2 | -3.385983513 | 0.004188401 | 0.021382472 |
| KDM2B | -3.383307917 | 2.26E-05 | 0.000267939 |
| MKL1 | -3.376677333 | 0.001789161 | 0.010709856 |
| SIGLEC7 | -3.37304881 | 1.04E-05 | 0.000136569 |
| CEP192 | -3.372296006 | 1.30E-05 | 0.000165371 |
| NBPF3 | -3.368331597 | 0.001945614 | 0.011471407 |
| AGAP1 | -3.352156276 | 0.000461185 | 0.003616859 |
| MIB2 | -3.349264364 | 0.000833722 | 0.005834838 |
| ULK4P1 | -3.346942703 | 0.006013703 | 0.028351965 |
| PYCARD-AS1 | -3.331484608 | 0.008337336 | 0.036713466 |
| HCST | -3.331037407 | 3.31E-23 | 7.92E-21 |
| TSHZ1 | -3.327930089 | 3.97E-07 | 7.63E-06 |
| SPN | -3.314254691 | 2.12E-14 | 1.59E-12 |
| MAST3 | -3.302167159 | 3.00E-10 | 1.14E-08 |
| RASGEF1A | -3.282749721 | 0.004119913 | 0.021162547 |
| FAM175A | -3.280361732 | 0.002271335 | 0.012993627 |
| CITED2 | -3.27266285 | 2.94E-08 | 7.44E-07 |
| MKI67 | -3.266798687 | 0.000579526 | 0.004343568 |
| CHST2 | -3.265574548 | 3.25E-14 | 2.40E-12 |

| | | | |
|---|---|---|---|
| NDUFAF7 | -3.26319628 | 0.004201141 | 0.02142276 |
| FGL2 | -3.256584258 | 1.55E-05 | 0.000193045 |
| RASA2 | -3.248986157 | 2.86E-07 | 5.69E-06 |
| UBTF | -3.240542037 | 1.07E-14 | 8.70E-13 |
| IFITM2 | -3.238047357 | 6.68E-32 | 2.73E-29 |
| HSPA7 | -3.234169329 | 0.001746268 | 0.010517275 |
| STAP1 | -3.22656129 | 0.002668573 | 0.014921456 |
| PYHIN1 | -3.226104486 | 3.34E-14 | 2.45E-12 |
| LINC01420 | -3.214710963 | 0.00028556 | 0.002392514 |
| GSDMD | -3.211642014 | 0.000224971 | 0.001944664 |
| ALDOC | -3.210525368 | 0.002034853 | 0.011947304 |
| SIGIRR | -3.199807085 | 3.30E-07 | 6.51E-06 |
| CDK5R1 | -3.191284225 | 0.005142884 | 0.02506549 |
| IL16 | -3.189690274 | 9.51E-10 | 3.26E-08 |
| PLAC8 | -3.180448707 | 5.83E-15 | 4.93E-13 |
| LOC101927780 | -3.174724587 | 0.00513678 | 0.025048187 |
| L3MBTL4-AS1 | -3.174643935 | 0.001219215 | 0.007879918 |
| PIP4K2C | -3.170213544 | 8.09E-05 | 0.000812797 |
| EP400NL | -3.17009004 | 0.006195434 | 0.028985887 |
| GRHL2 | -3.152164638 | 0.00592978 | 0.028064191 |
| FAM78A | -3.144292951 | 2.31E-07 | 4.75E-06 |
| KLRF1 | -3.13792939 | 7.70E-21 | 1.35E-18 |
| SLC8B1 | -3.137849748 | 0.003465147 | 0.018366591 |
| RCSD1 | -3.135000137 | 0.003007503 | 0.016453594 |
| TENM1 | -3.132501523 | 2.12E-06 | 3.36E-05 |
| UCP2 | -3.125884283 | 3.71E-18 | 4.73E-16 |
| KRBOX4 | -3.124230876 | 0.004557375 | 0.022835755 |
| PLA2G6 | -3.118904436 | 0.001422997 | 0.008937917 |
| BSCL2 | -3.105091769 | 0.007894653 | 0.035270694 |
| SLC2A12 | -3.092848536 | 0.005924825 | 0.028054273 |
| TYROBP | -3.090730378 | 2.65E-12 | 1.44E-10 |
| ERBB2 | -3.085436208 | 0.002678403 | 0.014956071 |
| DYM | -3.068181809 | 0.001441402 | 0.009041941 |
| FCGR3A | -3.064592843 | 7.68E-31 | 2.90E-28 |
| GPRASP1 | -3.062187637 | 0.005375916 | 0.025994632 |
| RNF165 | -3.051700581 | 6.35E-05 | 0.000654982 |
| TMEM173 | -3.049606086 | 1.65E-05 | 0.0002038 |
| DLG5 | -3.038466266 | 0.000209569 | 0.001837426 |
| SCML4 | -3.033388766 | 0.005300523 | 0.025693393 |
| SLC2A4RG | -3.026309908 | 0.000226419 | 0.001953734 |
| TTC39B | -3.018191972 | 0.001235048 | 0.007956043 |
| PXN | -3.014939971 | 6.89E-19 | 9.52E-17 |
| ABHD14A | -3.012403516 | 0.004675162 | 0.023255211 |
| TSEN54 | -3.009918743 | 7.38E-09 | 2.12E-07 |
| CTSW | -3.007225678 | 6.65E-13 | 4.00E-11 |
| PTGDR | -3.007044462 | 1.28E-10 | 5.20E-09 |
| GZMM | -3.004772632 | 5.25E-12 | 2.71E-10 |
| SLC9A3R1 | -3.002240095 | 7.28E-06 | 9.92E-05 |
| NSUN5P1 | -3.002123645 | 0.003615875 | 0.018991086 |
| PIK3IP1 | -2.99967236 | 2.48E-07 | 5.06E-06 |

| | | | |
|---|---|---|---|
| GMIP | -2.992154811 | 0.005648339 | 0.026979482 |
| RGS2 | -2.9810041 | 2.45E-07 | 5.02E-06 |
| CRIP1 | -2.975284528 | 9.59E-07 | 1.67E-05 |
| DOK2 | -2.969167544 | 2.16E-11 | 1.02E-09 |
| LOC285074 | -2.96686444 | 0.000841545 | 0.005880625 |
| GRK6 | -2.965756832 | 4.97E-05 | 0.000532191 |
| MED15 | -2.953958214 | 2.03E-05 | 0.000243344 |
| ARVCF | -2.951736198 | 0.006221685 | 0.029067121 |
| GPR56 | -2.94931319 | 1.25E-07 | 2.73E-06 |
| CCDC7 | -2.944661145 | 0.007287871 | 0.033071833 |
| NCOA2 | -2.940013202 | 5.88E-05 | 0.000614912 |
| TRAF3IP3 | -2.938345579 | 2.04E-14 | 1.55E-12 |
| LOC100131564 | -2.93334771 | 0.000184151 | 0.001648458 |
| NUDT16 | -2.919456935 | 8.34E-05 | 0.000834964 |
| CAT | -2.907133038 | 0.003334865 | 0.017869122 |
| PMS2P5 | -2.902393741 | 0.008893111 | 0.038538125 |
| PARVG | -2.900124661 | 0.000180981 | 0.001633491 |
| LOC100506076 | -2.898983189 | 0.000601755 | 0.004478784 |
| COTL1 | -2.891524934 | 0.000982217 | 0.006623046 |
| UVSSA | -2.890705991 | 0.002286129 | 0.013070636 |
| ABHD15 | -2.880662387 | 0.000575449 | 0.004322921 |
| ADHFE1 | -2.879263934 | 0.00044571 | 0.003512336 |
| SLC12A7 | -2.87340584 | 0.006214154 | 0.029045767 |
| USP53 | -2.867841209 | 0.00271405 | 0.015094983 |
| PIK3R3 | -2.864959817 | 0.000490086 | 0.003810012 |
| AREG | -2.862153203 | 0.007699785 | 0.034525866 |
| MMP24-AS1 | -2.856658388 | 6.56E-10 | 2.32E-08 |
| MYLIP | -2.856593695 | 8.22E-09 | 2.34E-07 |
| NOL4L | -2.856235718 | 2.87E-18 | 3.70E-16 |
| ITGAM | -2.852833063 | 4.88E-08 | 1.18E-06 |
| ESYT2 | -2.851883795 | 1.59E-05 | 0.00019686 |
| SYNE1 | -2.851489295 | 4.32E-14 | 3.10E-12 |
| PTGDS | -2.850024003 | 0.000474867 | 0.003712289 |
| CYTH4 | -2.846786556 | 8.42E-06 | 0.000112707 |
| SAMD3 | -2.844907274 | 3.50E-08 | 8.70E-07 |
| RMDN2 | -2.843275455 | 0.008608373 | 0.037636699 |
| MTFR1L | -2.842725199 | 0.009152592 | 0.039418823 |
| AK1 | -2.840793181 | 0.001492765 | 0.009263451 |
| BTBD2 | -2.837407097 | 0.007021173 | 0.03206924 |
| FAM98C | -2.834144433 | 0.000603463 | 0.004485281 |
| LAT | -2.828720158 | 8.92E-06 | 0.000118913 |
| DKFZP586I1420 | -2.827199389 | 0.000590849 | 0.00441831 |
| ZNF431 | -2.819316248 | 0.000602132 | 0.004478784 |
| DENND1C | -2.817206925 | 1.13E-05 | 0.000146169 |
| CCND3 | -2.817100507 | 1.33E-10 | 5.35E-09 |
| TMEM55B | -2.816588174 | 0.001353349 | 0.008582875 |
| ADD3 | -2.813099163 | 1.44E-10 | 5.77E-09 |
| SPECC1 | -2.808681761 | 0.00065068 | 0.004764045 |
| CCDC107 | -2.80632244 | 3.29E-05 | 0.000371636 |
| LGALS9 | -2.802007572 | 0.000745846 | 0.005345141 |

| | | | |
|---|---|---|---|
| SELPLG | -2.800309935 | 4.05E-19 | 5.68E-17 |
| LSP1 | -2.796812834 | 3.81E-08 | 9.42E-07 |
| CLSTN3 | -2.794094106 | 3.16E-08 | 7.95E-07 |
| BIN2 | -2.787897662 | 3.99E-16 | 4.04E-14 |
| VAMP2 | -2.78053424 | 6.23E-09 | 1.82E-07 |
| CEP128 | -2.774580243 | 0.001145577 | 0.007496791 |
| DENND2D | -2.772525962 | 0.000150113 | 0.001395976 |
| CASP10 | -2.770487436 | 3.36E-07 | 6.61E-06 |
| RCBTB1 | -2.758581105 | 0.000154426 | 0.001426704 |
| TMEM71 | -2.757881313 | 3.46E-07 | 6.77E-06 |
| GLTSCR2 | -2.756995632 | 4.25E-21 | 7.88E-19 |
| CYBA | -2.756805969 | 8.00E-08 | 1.85E-06 |
| ZNF317 | -2.747247549 | 0.000675585 | 0.004924343 |
| VAV3 | -2.746024329 | 2.96E-06 | 4.46E-05 |
| SUN2 | -2.74208235 | 3.97E-10 | 1.48E-08 |
| IL10RA | -2.738556441 | 2.59E-07 | 5.26E-06 |
| H1FX | -2.728980744 | 0.000295124 | 0.002455861 |
| PIK3R1 | -2.724291765 | 6.97E-20 | 1.04E-17 |
| MAD1L1 | -2.720848597 | 0.005813352 | 0.027646533 |
| SFXN3 | -2.720418501 | 0.000430369 | 0.003416144 |
| DYNLL2 | -2.7200133 | 0.000845656 | 0.005900945 |
| TMEM91 | -2.717922271 | 0.002256615 | 0.012929183 |
| RABGAP1L | -2.715798802 | 1.40E-06 | 2.32E-05 |
| HKR1 | -2.71427102 | 0.001508999 | 0.009334674 |
| CAPRIN2 | -2.714010023 | 0.000265285 | 0.002248366 |
| EVI2B | -2.713216098 | 1.78E-16 | 1.84E-14 |
| CHAF1A | -2.706479528 | 0.007360745 | 0.03330176 |
| TSC22D3 | -2.706011607 | 4.77E-10 | 1.75E-08 |
| ABHD17A | -2.701618248 | 1.86E-08 | 4.91E-07 |
| MTRNR2L1 | -2.692169043 | 0.00010649 | 0.001031364 |
| CCDC15 | -2.691457124 | 0.007820917 | 0.035019199 |
| TRANK1 | -2.689796469 | 0.000521236 | 0.004002072 |
| CNN2 | -2.689151069 | 1.90E-11 | 9.04E-10 |
| CASD1 | -2.688628233 | 0.001010926 | 0.006802604 |
| EPB41 | -2.685002621 | 1.42E-07 | 3.07E-06 |
| HSH2D | -2.684595032 | 3.76E-06 | 5.54E-05 |
| MAP4K3 | -2.677943701 | 0.006248685 | 0.029137759 |
| MXD4 | -2.675727921 | 1.49E-09 | 4.97E-08 |
| METTL7A | -2.674840352 | 5.13E-06 | 7.30E-05 |
| TRAPPC6A | -2.667854596 | 0.003911882 | 0.020242341 |
| NCAPD2 | -2.666048326 | 0.004530872 | 0.022729149 |
| KLRG1 | -2.663437527 | 9.17E-07 | 1.60E-05 |
| IPP | -2.655518787 | 0.000470221 | 0.003678897 |
| CEP78 | -2.648191937 | 1.32E-06 | 2.19E-05 |
| CRTAP | -2.646304908 | 0.00172547 | 0.010411186 |
| SH3BGRL | -2.644164844 | 1.31E-06 | 2.17E-05 |
| FBXO32 | -2.638122877 | 0.000172562 | 0.00157196 |
| OTUD1 | -2.634734345 | 0.00052449 | 0.004020132 |
| RNF44 | -2.628100433 | 0.000489046 | 0.003804939 |
| SMPD1 | -2.62773971 | 0.001372197 | 0.008657638 |

| | | | |
|---|---|---|---|
| ZBTB18 | -2.621108186 | 0.003892191 | 0.020151072 |
| CCDC88C | -2.62069032 | 1.95E-12 | 1.07E-10 |
| C16orf54 | -2.618246091 | 2.72E-07 | 5.47E-06 |
| RBM38 | -2.612626671 | 0.000118126 | 0.001130671 |
| ARHGAP9 | -2.61261432 | 2.59E-13 | 1.65E-11 |
| SSX2IP | -2.603551945 | 0.001626171 | 0.00992804 |
| PRR5 | -2.59421306 | 0.00450605 | 0.0226248 |
| TCIRG1 | -2.592318916 | 2.52E-05 | 0.000297262 |
| PTPN22 | -2.589777008 | 3.84E-09 | 1.15E-07 |
| PEX1 | -2.587397617 | 0.000702855 | 0.005096604 |
| ILF3-AS1 | -2.587171343 | 0.000631783 | 0.004644193 |
| PQLC3 | -2.584486863 | 0.002549636 | 0.014392682 |
| NR1D2 | -2.57645174 | 1.06E-09 | 3.56E-08 |
| EHBP1L1 | -2.571928426 | 0.004278623 | 0.021688746 |
| GIMAP4 | -2.570591465 | 1.31E-07 | 2.85E-06 |
| LNPEP | -2.57028384 | 0.006510679 | 0.030132117 |
| ARRB1 | -2.562055823 | 0.002653031 | 0.014878564 |
| MYO1F | -2.558652926 | 3.64E-09 | 1.10E-07 |
| SLC30A9 | -2.549885775 | 0.002712658 | 0.015094983 |
| RNF126 | -2.549876597 | 2.72E-05 | 0.000316683 |
| PHF21A | -2.549243591 | 0.00249624 | 0.014107495 |
| FAM134A | -2.546835112 | 4.18E-07 | 7.96E-06 |
| NMUR1 | -2.540649562 | 6.49E-06 | 8.92E-05 |
| CYTH1 | -2.529666182 | 1.16E-05 | 0.00014988 |
| BAZ2B | -2.52950276 | 8.66E-09 | 2.45E-07 |
| SLFN12L | -2.523756701 | 0.000929219 | 0.006353008 |
| SORL1 | -2.520354988 | 1.10E-06 | 1.88E-05 |
| LMF2 | -2.519878302 | 0.005947127 | 0.028132722 |
| FAM117A | -2.51969113 | 1.54E-06 | 2.53E-05 |
| ADAM28 | -2.516959803 | 0.00948036 | 0.040563372 |
| CCDC69 | -2.516587238 | 8.95E-08 | 2.05E-06 |
| FYB | -2.514547583 | 5.40E-14 | 3.84E-12 |
| SCARNA9 | -2.514462301 | 0.006531025 | 0.030210225 |
| TRMT10B | -2.513917176 | 0.00189822 | 0.011246036 |
| SIPA1 | -2.509654015 | 0.000770542 | 0.005482078 |
| ENC1 | -2.507321985 | 0.000605043 | 0.004493327 |
| SH3BP2 | -2.504896585 | 9.81E-06 | 0.000129654 |
| SIK1 | -2.50109834 | 0.008550678 | 0.037483702 |
| MATK | -2.498872789 | 3.02E-05 | 0.000346064 |
| PRSS23 | -2.498531782 | 2.69E-08 | 6.95E-07 |
| MYO1G | -2.49683869 | 1.02E-05 | 0.000134847 |
| NAAA | -2.492752817 | 1.61E-07 | 3.44E-06 |
| LOC283070 | -2.489875411 | 0.003851879 | 0.020004554 |
| TPST2 | -2.489789946 | 9.96E-08 | 2.26E-06 |
| CDC14A | -2.489738864 | 0.000553334 | 0.004201824 |
| BTN3A1 | -2.482826885 | 2.57E-07 | 5.23E-06 |
| GLTSCR1L | -2.479727659 | 2.34E-05 | 0.000277335 |
| TBC1D5 | -2.477922839 | 0.001752119 | 0.010539573 |
| VPS16 | -2.476353824 | 0.000350385 | 0.002848077 |
| CTBP1 | -2.467057599 | 1.10E-13 | 7.71E-12 |

| | | | |
|---|---|---|---|
| DGKZ | -2.46498916 | 4.44E-05 | 0.000481469 |
| STK10 | -2.4591105 | 6.40E-08 | 1.50E-06 |
| SUMF1 | -2.458139698 | 0.006723973 | 0.030913262 |
| LLGL2 | -2.44594785 | 0.000281573 | 0.00236112 |
| ATP2A3 | -2.445456963 | 0.000196126 | 0.001744506 |
| CERK | -2.44346374 | 4.83E-05 | 0.000519461 |
| VPS37B | -2.438103138 | 0.000710869 | 0.005135741 |
| CA5B | -2.437689771 | 0.000165398 | 0.001512325 |
| DOCK11 | -2.43693835 | 3.72E-14 | 2.71E-12 |
| ADCY7 | -2.414853822 | 6.13E-16 | 5.89E-14 |
| AP1G2 | -2.414065006 | 6.00E-07 | 1.10E-05 |
| ACVR2A | -2.413444923 | 0.002162189 | 0.012463711 |
| CBX4 | -2.409858546 | 7.32E-09 | 2.11E-07 |
| SNX20 | -2.401939402 | 0.006168875 | 0.028889178 |
| SYNJ2BP | -2.400875687 | 0.004260852 | 0.021614904 |
| ADRB2 | -2.398713275 | 0.005538727 | 0.02655936 |
| MRS2P2 | -2.395056359 | 0.004395594 | 0.022189572 |
| PAPD5 | -2.392953375 | 2.66E-07 | 5.37E-06 |
| BNIP3L | -2.390210302 | 2.26E-05 | 0.000268167 |
| KATNAL1 | -2.386847826 | 0.00096383 | 0.006525973 |
| ATM | -2.382233332 | 2.77E-08 | 7.12E-07 |
| TWF1 | -2.380530105 | 4.52E-05 | 0.000490492 |
| ITPKB | -2.380149174 | 1.36E-08 | 3.71E-07 |
| PTOV1-AS2 | -2.376051661 | 0.003833829 | 0.019943635 |
| MGAT4A | -2.368994288 | 1.27E-06 | 2.11E-05 |
| MBNL3 | -2.368954212 | 0.002417073 | 0.013699538 |
| TRMT2B | -2.366215277 | 0.008717616 | 0.037978921 |
| SLC44A2 | -2.364919319 | 4.96E-07 | 9.33E-06 |
| ITGA6 | -2.364720694 | 2.65E-05 | 0.000309925 |
| EFHD2 | -2.357303732 | 5.00E-13 | 3.07E-11 |
| FAM111A | -2.353281554 | 0.001481227 | 0.009209329 |
| BIN1 | -2.348714949 | 0.000278898 | 0.002343685 |
| EVL | -2.344033751 | 1.81E-14 | 1.42E-12 |
| FLOT2 | -2.343752519 | 1.10E-05 | 0.000142159 |
| GYPC | -2.343434157 | 2.56E-13 | 1.65E-11 |
| SLAMF6 | -2.342871827 | 0.00020649 | 0.001816924 |
| FOSL2 | -2.342792727 | 0.001170469 | 0.007608863 |
| PSTPIP1 | -2.333908278 | 0.008109023 | 0.035901455 |
| S100A4 | -2.326322364 | 1.05E-08 | 2.92E-07 |
| NME3 | -2.322562376 | 0.00527398 | 0.025608994 |
| CCDC186 | -2.322371374 | 3.73E-10 | 1.40E-08 |
| PTPN18 | -2.314016864 | 0.000310739 | 0.002570536 |
| TUBGCP2 | -2.307471429 | 0.00140926 | 0.008862983 |
| CXCR4 | -2.30711602 | 0.00010548 | 0.001023602 |
| ZBTB16 | -2.301702538 | 0.000324856 | 0.002662622 |
| NCBP2-AS2 | -2.30039611 | 3.16E-11 | 1.45E-09 |
| NCOR2 | -2.29667608 | 6.29E-09 | 1.84E-07 |
| LAPTM5 | -2.296195654 | 2.97E-13 | 1.88E-11 |
| TC2N | -2.29362753 | 0.000271438 | 0.002289834 |
| BEX2 | -2.292597594 | 0.001103239 | 0.00728881 |

| | | | |
|---|---|---|---|
| SMARCA2 | -2.286408364 | 1.89E-06 | 3.05E-05 |
| FGFBP2 | -2.285488137 | 3.59E-10 | 1.36E-08 |
| CDKN2D | -2.278774884 | 8.26E-06 | 0.000110822 |
| SUSD1 | -2.275423138 | 0.008975211 | 0.038842431 |
| TCEAL8 | -2.273284508 | 0.001643893 | 0.00999271 |
| GSAP | -2.272227049 | 6.54E-06 | 8.96E-05 |
| FGD3 | -2.2688971 | 3.27E-05 | 0.000369283 |
| GAB3 | -2.268855897 | 5.30E-09 | 1.56E-07 |
| ESYT1 | -2.267830738 | 0.004666771 | 0.023250876 |
| TXNIP | -2.264587462 | 7.19E-17 | 7.66E-15 |
| SPOCK2 | -2.263311372 | 5.80E-07 | 1.07E-05 |
| PRAF2 | -2.262290603 | 0.006716828 | 0.03089489 |
| KAT6B | -2.260801361 | 5.99E-11 | 2.66E-09 |
| CCNG2 | -2.255521342 | 0.000311069 | 0.002571104 |
| UBXN7 | -2.244322654 | 0.000425858 | 0.003385812 |
| TRIB2 | -2.241067745 | 0.000354834 | 0.002881849 |
| CWC27 | -2.238405639 | 0.000421273 | 0.003357526 |
| GTPBP1 | -2.237418616 | 0.004621065 | 0.023107682 |
| PCM1 | -2.235502428 | 4.48E-10 | 1.65E-08 |
| AFF1 | -2.234500416 | 6.74E-10 | 2.38E-08 |
| TXK | -2.233479755 | 5.58E-08 | 1.32E-06 |
| DENND6A | -2.232224775 | 1.92E-05 | 0.000231666 |
| MAN2B2 | -2.231307717 | 2.79E-05 | 0.000324254 |
| SKAP1 | -2.228217573 | 0.000570611 | 0.004296442 |
| B3GNT7 | -2.227811978 | 2.20E-05 | 0.000261992 |
| S1PR5 | -2.225520655 | 1.98E-14 | 1.52E-12 |
| SAMD9 | -2.22500124 | 2.13E-09 | 6.83E-08 |
| MAP4K1 | -2.224249921 | 0.006181437 | 0.028934197 |
| LINC00861 | -2.223875949 | 7.48E-06 | 0.000101416 |
| TNFSF12 | -2.220391162 | 0.009672027 | 0.041096689 |
| PHF3 | -2.217341987 | 8.81E-11 | 3.76E-09 |
| ATAD2 | -2.216200169 | 0.002668375 | 0.014921456 |
| PTPRA | -2.206024607 | 2.67E-08 | 6.90E-07 |
| TECR | -2.202301074 | 0.002082346 | 0.012119716 |
| MTSS1 | -2.200369335 | 0.000173077 | 0.001575191 |
| C12orf75 | -2.179949214 | 8.78E-09 | 2.47E-07 |
| UTRN | -2.178976984 | 4.65E-09 | 1.38E-07 |
| SHKBP1 | -2.178160367 | 0.000306017 | 0.002537901 |
| LRRCC1 | -2.177250953 | 0.004224096 | 0.021495127 |
| DENND4C | -2.177045589 | 3.91E-06 | 5.71E-05 |
| OSBPL2 | -2.176052307 | 1.03E-05 | 0.000136012 |
| HAUS3 | -2.175743716 | 8.59E-07 | 1.51E-05 |
| SPG7 | -2.175282971 | 0.000488823 | 0.003804939 |
| PRKACB | -2.173868861 | 1.51E-05 | 0.000188115 |
| KLF3 | -2.158795924 | 7.11E-10 | 2.49E-08 |
| APBB1IP | -2.157700751 | 7.90E-10 | 2.75E-08 |
| HLA-DPB1 | -2.155514821 | 0.002111803 | 0.012245214 |
| KLHDC3 | -2.154467896 | 3.46E-05 | 0.00038801 |
| FAM53B | -2.149348735 | 0.000121301 | 0.001152589 |
| AGTPBP1 | -2.144264423 | 1.94E-06 | 3.11E-05 |

| | | | |
|---|---|---|---|
| JMJD1C | -2.143030065 | 3.64E-10 | 1.37E-08 |
| SYK | -2.136287299 | 1.18E-06 | 1.98E-05 |
| ARHGDIB | -2.134647079 | 7.48E-13 | 4.39E-11 |
| RNF139 | -2.131756241 | 1.07E-05 | 0.000139249 |
| ZFP36 | -2.126968956 | 0.003860522 | 0.020018805 |
| PGPEP1 | -2.125883665 | 4.80E-06 | 6.87E-05 |
| FAM21A | -2.121912298 | 0.009918862 | 0.041907427 |
| CNOT6L | -2.120454871 | 3.42E-12 | 1.81E-10 |
| LAIR1 | -2.114116891 | 0.003309102 | 0.017750462 |
| AGPAT4 | -2.083772569 | 0.004344775 | 0.02196115 |
| DTHD1 | -2.079987242 | 0.005425965 | 0.026172145 |
| ARRDC2 | -2.077222038 | 0.000286029 | 0.002394396 |
| IDH2 | -2.077181085 | 2.73E-06 | 4.17E-05 |
| USP28 | -2.072661544 | 1.67E-05 | 0.000204794 |
| NCALD | -2.071501564 | 1.37E-05 | 0.000173249 |
| ZNF394 | -2.070104406 | 0.000978151 | 0.006602998 |
| BTN3A3 | -2.06609766 | 0.000446095 | 0.003512547 |
| PLSCR3 | -2.063036509 | 0.000480388 | 0.003752454 |
| VRK3 | -2.058492402 | 3.42E-05 | 0.000384862 |
| ERMP1 | -2.050094753 | 0.002005634 | 0.01179693 |
| IFT74 | -2.04883696 | 0.006555459 | 0.030308957 |
| DAB2 | -2.034408636 | 0.000174027 | 0.001582372 |
| GZMA | -2.025842408 | 4.65E-15 | 4.04E-13 |
| YPEL5 | -2.017609908 | 1.37E-08 | 3.71E-07 |
| IQSEC1 | -2.017305509 | 0.001707237 | 0.010320211 |
| FCER1G | -2.017198913 | 1.17E-05 | 0.00015047 |
| FAM122B | -2.015569219 | 1.83E-09 | 5.90E-08 |
| AUTS2 | -2.010635663 | 0.001408443 | 0.008862983 |
| SNX29 | -2.008635847 | 0.004440012 | 0.022341713 |
| ZMYND8 | -2.007399181 | 6.50E-07 | 1.18E-05 |
| ZFP36L2 | -2.00553074 | 8.75E-11 | 3.75E-09 |
| SUPT6H | -2.00414142 | 0.001094687 | 0.007246945 |
| INPP5D | -1.996048919 | 1.09E-08 | 3.01E-07 |
| NMT2 | -1.992901872 | 0.003087952 | 0.016767407 |
| AES | -1.992392155 | 2.52E-08 | 6.56E-07 |
| MOV10 | -1.986608076 | 0.006692346 | 0.030812076 |
| CEP250 | -1.984160033 | 0.003669274 | 0.019230367 |
| NKTR | -1.98031621 | 1.52E-11 | 7.37E-10 |
| FAM214A | -1.974935944 | 1.86E-08 | 4.91E-07 |
| PLCL2 | -1.971551738 | 0.002667518 | 0.014921456 |
| AP4B1 | -1.968721268 | 2.65E-05 | 0.000309925 |
| NCAM1 | -1.967764629 | 6.14E-05 | 0.000637676 |
| RBM33 | -1.96771944 | 3.44E-06 | 5.11E-05 |
| GOLGA8N | -1.964486984 | 0.000844254 | 0.005895357 |
| ERCC5 | -1.958461259 | 0.000876205 | 0.006058105 |
| CWC22 | -1.95696676 | 0.000323828 | 0.002658639 |
| GTF3A | -1.955132084 | 0.000442079 | 0.003492135 |
| ITGB7 | -1.954415071 | 6.00E-06 | 8.38E-05 |
| ATP8A1 | -1.95125939 | 0.00015458 | 0.001426704 |
| TAPBPL | -1.943963891 | 0.000629878 | 0.00463596 |

| | | | |
|---|---|---|---|
| YPEL3 | -1.943766431 | 0.001142585 | 0.007490553 |
| SNRK | -1.942803089 | 0.000103864 | 0.00100892 |
| AGO1 | -1.942312333 | 0.002142724 | 0.012395205 |
| KLF12 | -1.942165491 | 6.85E-05 | 0.000699573 |
| GLIPR2 | -1.939241729 | 1.69E-07 | 3.59E-06 |
| PREX1 | -1.93119698 | 0.000657285 | 0.004805234 |
| LPCAT1 | -1.927535119 | 9.33E-10 | 3.21E-08 |
| DIP2A | -1.922864774 | 2.70E-08 | 6.96E-07 |
| YPEL2 | -1.92112687 | 0.007904385 | 0.035290092 |
| RASAL3 | -1.920005202 | 0.000128544 | 0.001212639 |
| CDK5RAP2 | -1.919828305 | 0.001761855 | 0.010591639 |
| ADA | -1.916192453 | 0.000225471 | 0.001947271 |
| JUN | -1.910959009 | 1.12E-08 | 3.08E-07 |
| CTSS | -1.901098092 | 0.00014122 | 0.001320794 |
| ARHGAP1 | -1.894811623 | 0.006637727 | 0.03063864 |
| FRAT1 | -1.888944905 | 0.005725394 | 0.027281126 |
| C5orf56 | -1.885195923 | 4.60E-07 | 8.69E-06 |
| CREBRF | -1.883815703 | 5.53E-08 | 1.31E-06 |
| SH3BP1 | -1.878985144 | 0.000678253 | 0.004940122 |
| CPT1A | -1.874844438 | 0.000136214 | 0.001276405 |
| PARP8 | -1.873419954 | 0.000846658 | 0.005903737 |
| GNS | -1.872852594 | 0.002318905 | 0.013234891 |
| LINC01278 | -1.870230264 | 0.001320056 | 0.008409785 |
| RARRES3 | -1.868106193 | 1.85E-06 | 2.99E-05 |
| SLX1A | -1.867985607 | 0.002960565 | 0.016272329 |
| LYN | -1.865166145 | 8.11E-05 | 0.000813689 |
| BPTF | -1.855283632 | 2.21E-10 | 8.66E-09 |
| ECHDC2 | -1.853405969 | 0.003374694 | 0.018043117 |
| MAPKAP1 | -1.852234635 | 1.21E-05 | 0.000155116 |
| MIDN | -1.85209646 | 5.51E-05 | 0.000581913 |
| PRKCH | -1.847889762 | 1.25E-08 | 3.42E-07 |
| NUP210 | -1.837633513 | 0.00191072 | 0.011292817 |
| CYFIP2 | -1.832891314 | 9.85E-10 | 3.37E-08 |
| GAS7 | -1.82957595 | 0.006081934 | 0.028584217 |
| LOC100132352 | -1.822260436 | 6.82E-05 | 0.00069662 |
| MGEA5 | -1.820844946 | 1.28E-10 | 5.20E-09 |
| ARL4C | -1.820720365 | 1.38E-13 | 9.40E-12 |
| DTNBP1 | -1.804037961 | 0.000749507 | 0.00535963 |
| NKG7 | -1.80350195 | 5.37E-10 | 1.94E-08 |
| FXYD5 | -1.801897638 | 0.003000709 | 0.016437722 |
| FAM199X | -1.801638592 | 0.002983402 | 0.016370336 |
| APEH | -1.794846587 | 0.003653234 | 0.019166782 |
| NRD1 | -1.79395359 | 2.62E-06 | 4.04E-05 |
| FBXW5 | -1.792534131 | 0.006099811 | 0.028634089 |
| ANKRD13A | -1.790701147 | 9.86E-05 | 0.000967391 |
| NAA38 | -1.786762452 | 0.007512298 | 0.033808787 |
| CLN8 | -1.766827232 | 0.003994795 | 0.020584526 |
| EIF2AK2 | -1.760297769 | 0.005437319 | 0.026204196 |
| VPS41 | -1.759982875 | 0.004763548 | 0.023591707 |
| INPP4A | -1.752573742 | 4.32E-08 | 1.05E-06 |

| | | | |
|---|---|---|---|
| MEI1 | -1.752362516 | 0.003404636 | 0.018134032 |
| PHF12 | -1.752214573 | 0.003623333 | 0.019020077 |
| MED29 | -1.750668822 | 0.003963228 | 0.020475633 |
| GMFG | -1.748070802 | 2.60E-08 | 6.75E-07 |
| DPH7 | -1.746485029 | 0.003499407 | 0.018518167 |
| ARHGEF1 | -1.742689522 | 0.005515006 | 0.026471489 |
| RPS6KA1 | -1.739687351 | 0.004790303 | 0.023682896 |
| PIKFYVE | -1.736647506 | 0.006860695 | 0.031453399 |
| C19orf66 | -1.724887878 | 4.14E-05 | 0.00045247 |
| ITGAL | -1.720284473 | 1.21E-08 | 3.31E-07 |
| FBXW7 | -1.719990551 | 1.07E-05 | 0.000139249 |
| TMEM259 | -1.719458462 | 0.001455116 | 0.009098882 |
| TNRC6C | -1.718473638 | 0.000221156 | 0.001918442 |
| PIK3R5 | -1.714907276 | 8.18E-07 | 1.45E-05 |
| NIPBL | -1.71307765 | 2.20E-07 | 4.56E-06 |
| SESN1 | -1.697750861 | 0.001826955 | 0.010896204 |
| TFDP2 | -1.69716827 | 6.13E-05 | 0.000636734 |
| RAP2B | -1.696149535 | 6.18E-09 | 1.82E-07 |
| BEX4 | -1.693743275 | 5.76E-05 | 0.000603228 |
| WBP1 | -1.693468897 | 0.009472806 | 0.040563372 |
| ZRANB1 | -1.692447887 | 0.000348799 | 0.002839888 |
| EZH1 | -1.689217574 | 0.001239223 | 0.007977703 |
| CST7 | -1.688604973 | 3.45E-09 | 1.06E-07 |
| ARHGAP30 | -1.685881895 | 3.14E-06 | 4.70E-05 |
| TPRG1L | -1.676856968 | 0.005446059 | 0.026230381 |
| RBL2 | -1.675835418 | 0.000120705 | 0.001148629 |
| SSH2 | -1.6722302 | 5.53E-05 | 0.000581913 |
| PFKL | -1.666059669 | 0.006354116 | 0.029545133 |
| TRAPPC2 | -1.665937018 | 0.000376237 | 0.003030588 |
| DNAJC13 | -1.664188008 | 0.000119029 | 0.001135705 |
| CUX1 | -1.664109378 | 3.18E-05 | 0.000362204 |
| PALLD | -1.661800614 | 0.008571161 | 0.037524168 |
| WNK1 | -1.660910888 | 6.27E-06 | 8.73E-05 |
| LIMD2 | -1.658290626 | 0.005281356 | 0.025625808 |
| SLC25A42 | -1.65733939 | 0.002962603 | 0.016274414 |
| ZBTB24 | -1.657031778 | 0.002082748 | 0.012119716 |
| CTDSP1 | -1.650243217 | 3.21E-06 | 4.79E-05 |
| GDE1 | -1.650073504 | 0.004911983 | 0.024131931 |
| HMGN3 | -1.647889553 | 1.04E-05 | 0.000136948 |
| PAN3 | -1.646288919 | 4.58E-06 | 6.60E-05 |
| PRKCB | -1.642446711 | 4.09E-05 | 0.000448629 |
| NPRL2 | -1.640751182 | 0.008555658 | 0.037483702 |
| LITAF | -1.637571067 | 1.28E-07 | 2.79E-06 |
| TIMP1 | -1.636731556 | 0.007688822 | 0.034494987 |
| ARHGAP15 | -1.635710013 | 0.002349501 | 0.013393932 |
| KIAA1109 | -1.630181159 | 5.16E-08 | 1.23E-06 |
| MEF2D | -1.628422207 | 3.04E-06 | 4.57E-05 |
| CEP85L | -1.620256895 | 1.58E-07 | 3.38E-06 |
| PACS1 | -1.619249632 | 0.003010779 | 0.016456127 |
| ERBB2IP | -1.616677491 | 4.05E-06 | 5.90E-05 |

| | | | |
|---|---|---|---|
| ARHGEF18 | -1.614860346 | 0.009014252 | 0.038977006 |
| TRABD | -1.599253774 | 0.00569436 | 0.027172845 |
| MKRN1 | -1.599184055 | 8.66E-05 | 0.000864012 |
| RAB11B | -1.598980378 | 0.002095363 | 0.012164568 |
| KIAA2013 | -1.596879557 | 0.00081791 | 0.005745267 |
| PIK3C2A | -1.594984557 | 0.000224931 | 0.001944664 |
| YPEL1 | -1.594425028 | 0.003091732 | 0.016767407 |
| ERC1 | -1.593345204 | 0.000937083 | 0.006388969 |
| CRBN | -1.592552345 | 1.38E-05 | 0.00017518 |
| R3HDM1 | -1.591552139 | 0.000741872 | 0.005322187 |
| DDX3X | -1.591232703 | 1.05E-09 | 3.56E-08 |
| GOLGA8B | -1.586191621 | 8.96E-06 | 0.000119269 |
| CD7 | -1.583363891 | 0.000302672 | 0.002514406 |
| FAM134C | -1.582352915 | 0.001561077 | 0.009590313 |
| FLNA | -1.574002128 | 5.78E-08 | 1.36E-06 |
| LY6E | -1.573343184 | 0.009213779 | 0.039647537 |
| HERC1 | -1.573121597 | 1.60E-06 | 2.62E-05 |
| PIGK | -1.571399106 | 0.008131098 | 0.035966728 |
| DGKD | -1.570040737 | 4.91E-05 | 0.00052752 |
| STK4 | -1.569196769 | 1.09E-10 | 4.53E-09 |
| SRSF5 | -1.567200352 | 3.56E-06 | 5.27E-05 |
| APMAP | -1.565945029 | 7.65E-07 | 1.36E-05 |
| CSNK1G2 | -1.543523325 | 1.44E-05 | 0.000180649 |
| STK38 | -1.537475445 | 6.22E-05 | 0.000643065 |
| LOC606724 | -1.537225912 | 1.43E-05 | 0.000179494 |
| ABHD4 | -1.537096714 | 0.006238978 | 0.029120177 |
| GVINP1 | -1.53469345 | 0.000620867 | 0.004586843 |
| TRAPPC1 | -1.533192847 | 0.00997333 | 0.042049136 |
| ABCC5 | -1.531640187 | 0.003826026 | 0.019913602 |
| ESCO1 | -1.531472684 | 4.78E-06 | 6.86E-05 |
| LRBA | -1.522486672 | 0.007864356 | 0.035151342 |
| ALOX5AP | -1.521140814 | 0.007909945 | 0.035290802 |
| PPP6R1 | -1.515412938 | 8.73E-05 | 0.000869107 |
| NLRC5 | -1.514787735 | 0.000396568 | 0.003173515 |
| MACF1 | -1.510340682 | 7.52E-07 | 1.35E-05 |
| PRMT2 | -1.510327523 | 0.00057609 | 0.004324421 |
| SHISA5 | -1.508542523 | 0.000202622 | 0.001792539 |
| CCDC144B | -1.508228935 | 0.009479053 | 0.040563372 |
| NDUFV1 | -1.506621615 | 0.001668785 | 0.010125201 |
| ARID4A | -1.502584192 | 0.000124867 | 0.001180222 |
| THAP11 | -1.501358751 | 0.007313514 | 0.033172853 |
| AXIN1 | -1.501324075 | 3.22E-05 | 0.000365389 |
| KAT2B | -1.500485631 | 0.000752223 | 0.005375134 |
| ZFC3H1 | -1.500181123 | 0.000739511 | 0.00531138 |
| DPP8 | -1.498609451 | 0.007465964 | 0.033615684 |
| P2RY8 | -1.498041848 | 0.001307294 | 0.00834474 |
| ZNF148 | -1.496156274 | 2.27E-05 | 0.000268921 |
| KLRB1 | -1.495676883 | 2.63E-06 | 4.05E-05 |
| ABLIM1 | -1.489679299 | 0.009788631 | 0.041471535 |
| C19orf12 | -1.488371987 | 0.003771282 | 0.019733356 |

| | | | |
|---|---|---|---|
| SYNE2 | -1.488244738 | 1.20E-07 | 2.66E-06 |
| CUTA | -1.487439999 | 0.006796637 | 0.031232695 |
| FAM65B | -1.486322273 | 4.08E-07 | 7.81E-06 |
| XPC | -1.485453949 | 0.006416554 | 0.029793094 |
| TMC6 | -1.483525099 | 0.008768476 | 0.03809899 |
| KLRK1 | -1.479784989 | 6.77E-07 | 1.23E-05 |
| ADD1 | -1.477422854 | 1.61E-05 | 0.000199491 |
| SCAMP1 | -1.476818961 | 0.002377661 | 0.013507371 |
| ZNF621 | -1.474346374 | 0.004325571 | 0.021875348 |
| ARHGAP25 | -1.470776894 | 0.003385804 | 0.018079034 |
| SCARNA17 | -1.465958007 | 8.73E-05 | 0.000869107 |
| SNRPN | -1.461935947 | 0.001314012 | 0.008382169 |
| ABR | -1.461692241 | 0.008302149 | 0.036623884 |
| JMY | -1.457768343 | 0.002299438 | 0.013139074 |
| DDX60L | -1.456248631 | 0.007389479 | 0.033363173 |
| ATF7IP | -1.455845808 | 5.53E-05 | 0.000581913 |
| CRIM1 | -1.455294949 | 0.009464988 | 0.040550656 |
| IFFO1 | -1.454455274 | 0.004223437 | 0.021495127 |
| TMX4 | -1.453036007 | 0.000597017 | 0.004454244 |
| ICAM2 | -1.451879856 | 0.006905523 | 0.0316146 |
| PCNT | -1.445900688 | 0.001524063 | 0.009398231 |
| TRIM38 | -1.442041372 | 0.005495466 | 0.026403534 |
| GATAD2A | -1.440747133 | 0.000631943 | 0.004644193 |
| CORO1A | -1.437156234 | 0.004417414 | 0.022282389 |
| SCLT1 | -1.436157577 | 0.002054586 | 0.012034356 |
| HMHA1 | -1.435497499 | 2.95E-06 | 4.46E-05 |
| BRWD1 | -1.435308714 | 5.11E-06 | 7.29E-05 |
| CSK | -1.431241127 | 1.87E-05 | 0.000225206 |
| CXXC5 | -1.431012515 | 0.001326349 | 0.008444392 |
| LINC00342 | -1.429787007 | 0.002327267 | 0.013274894 |
| ABI3 | -1.428341751 | 0.00868705 | 0.03789624 |
| NUB1 | -1.427537969 | 0.000521318 | 0.004002072 |
| TAOK3 | -1.424240174 | 2.27E-07 | 4.68E-06 |
| MAGED1 | -1.424059688 | 0.002413992 | 0.013689986 |
| MYBL1 | -1.423190901 | 0.000218916 | 0.001904227 |
| BNC2 | -1.420056107 | 0.00486793 | 0.023988976 |
| MTIF2 | -1.418749009 | 0.004176203 | 0.02136117 |
| MYO9A | -1.416244274 | 0.004188881 | 0.021382472 |
| OSBPL8 | -1.415996155 | 0.00049331 | 0.003831035 |
| TRIM44 | -1.409052571 | 0.001536683 | 0.009464154 |
| CAST | -1.407281459 | 5.43E-07 | 1.01E-05 |
| ARHGEF9 | -1.405095653 | 0.000872915 | 0.006043875 |
| ATRX | -1.396729418 | 3.05E-08 | 7.70E-07 |
| C6orf106 | -1.392702466 | 3.27E-05 | 0.000369283 |
| GPCPD1 | -1.387131294 | 0.000134418 | 0.001261717 |
| RAB1B | -1.386208074 | 0.000529868 | 0.004048701 |
| WHAMM | -1.383241411 | 0.008752565 | 0.038097346 |
| MBNL1 | -1.382463483 | 9.80E-09 | 2.73E-07 |
| TCF7 | -1.380260107 | 0.008051499 | 0.035711236 |
| KIAA1551 | -1.379597497 | 0.000301127 | 0.002503691 |

| | | | |
|---|---|---|---|
| JAKMIP2 | -1.379097287 | 0.000859769 | 0.005978169 |
| JADE2 | -1.377711489 | 0.002261045 | 0.012942305 |
| CARD11 | -1.377657087 | 0.005884047 | 0.027888108 |
| EMB | -1.373812249 | 0.000512064 | 0.003952681 |
| MBP | -1.372432781 | 0.000108053 | 0.001042387 |
| VPS13C | -1.365983659 | 2.75E-07 | 5.53E-06 |
| CCNL2 | -1.36293675 | 0.000562418 | 0.004257627 |
| LCK | -1.356938967 | 0.000862905 | 0.005987246 |
| TTC3 | -1.356330875 | 2.38E-06 | 3.72E-05 |
| NLRC3 | -1.355670251 | 0.001672485 | 0.010135114 |
| ITPRIP | -1.354304489 | 0.003115283 | 0.016876886 |
| MALAT1 | -1.348716109 | 4.41E-10 | 1.63E-08 |
| SLCO4C1 | -1.345362968 | 0.00028896 | 0.002414808 |
| MSL2 | -1.343695039 | 0.00424623 | 0.021574191 |
| SYTL2 | -1.342890753 | 0.000868065 | 0.00601454 |
| NOTCH2NL | -1.338376874 | 0.009891703 | 0.041830818 |
| TPP1 | -1.33506349 | 0.000242531 | 0.002074518 |
| STK17A | -1.334788247 | 0.001193649 | 0.007735069 |
| FAM192A | -1.332573879 | 0.004600576 | 0.023028701 |
| ANKRD36 | -1.332484669 | 0.000212819 | 0.00186093 |
| AAK1 | -1.32969343 | 0.000178101 | 0.001613434 |
| PLEKHA2 | -1.32885065 | 0.001789066 | 0.010709856 |
| MYSM1 | -1.326989963 | 0.000864817 | 0.005996269 |
| DMTF1 | -1.323780435 | 0.001232267 | 0.007943347 |
| SMC5 | -1.318296252 | 0.002410487 | 0.013678016 |
| RPL3 | -1.313311271 | 2.08E-06 | 3.30E-05 |
| GPATCH8 | -1.31003402 | 6.44E-06 | 8.89E-05 |
| GON4L | -1.299775121 | 0.001098828 | 0.00726456 |
| ASCC2 | -1.295431369 | 0.008016932 | 0.035622335 |
| SCAPER | -1.293240982 | 0.000787235 | 0.005588687 |
| KLHL24 | -1.292253992 | 0.003124021 | 0.016905557 |
| TET2 | -1.290563257 | 0.000736403 | 0.005292933 |
| EIF4B | -1.289941082 | 0.000265701 | 0.002249176 |
| CPNE3 | -1.289235587 | 0.001287751 | 0.008246817 |
| CAPNS1 | -1.287486896 | 2.78E-06 | 4.25E-05 |
| KIAA0922 | -1.285838287 | 0.00207929 | 0.012119201 |
| USP25 | -1.280867755 | 0.000502209 | 0.003891918 |
| MYCBP2 | -1.277692801 | 0.000147552 | 0.001374773 |
| CERKL | -1.277060664 | 0.008536456 | 0.037441182 |
| CALM1 | -1.275551188 | 4.25E-06 | 6.14E-05 |
| PTPRC | -1.275153472 | 2.18E-07 | 4.53E-06 |
| SEPHS2 | -1.274824207 | 8.11E-05 | 0.000813689 |
| ATG16L2 | -1.274601568 | 0.001157329 | 0.007549042 |
| ZNF33A | -1.273366065 | 0.003851893 | 0.020004554 |
| PNN | -1.273346513 | 2.26E-07 | 4.68E-06 |
| RNF20 | -1.267167431 | 0.008279644 | 0.036557869 |
| KIF21B | -1.266602013 | 0.006052066 | 0.028491755 |
| CREBBP | -1.263254498 | 0.000749391 | 0.00535963 |
| CTSD | -1.262232819 | 0.000466046 | 0.003652057 |
| TOPORS | -1.262105829 | 0.005032389 | 0.024686383 |

| | | | |
|---|---|---|---|
| FAM208A | -1.26006271 | 0.001444989 | 0.009052866 |
| HELZ | -1.258773002 | 0.000690629 | 0.00501908 |
| WHSC1L1 | -1.24573796 | 0.000216804 | 0.001889046 |
| PCSK7 | -1.244979892 | 0.000610788 | 0.004519185 |
| ZNF652 | -1.243877668 | 0.00258445 | 0.014564065 |
| GNAI2 | -1.243374117 | 0.000231963 | 0.001994555 |
| PPP3CC | -1.23104283 | 0.003512355 | 0.018566657 |
| BAG5 | -1.227435449 | 0.006610496 | 0.030534639 |
| KIF2A | -1.22713848 | 0.007355955 | 0.03330176 |
| SLC25A36 | -1.22648413 | 0.004650608 | 0.023210775 |
| ZBTB44 | -1.225916218 | 4.00E-05 | 0.00044048 |
| PAPD4 | -1.224560873 | 0.0014823 | 0.009210163 |
| RBM3 | -1.221916109 | 6.45E-05 | 0.000665211 |
| IDS | -1.221044469 | 1.51E-06 | 2.49E-05 |
| PDCD4 | -1.213387942 | 5.75E-06 | 8.09E-05 |
| IGBP1 | -1.211685069 | 0.006432387 | 0.029852481 |
| PSD4 | -1.207400743 | 0.005237755 | 0.025452013 |
| DCAF8 | -1.205874203 | 0.000807083 | 0.005700716 |
| TOMM20 | -1.201804948 | 0.002139666 | 0.012384818 |
| STAT6 | -1.20152926 | 2.23E-05 | 0.000265758 |
| RIOK3 | -1.193627214 | 0.001011786 | 0.006803722 |
| WIPF1 | -1.191936426 | 0.000519932 | 0.003997688 |
| PRDM2 | -1.186564679 | 0.000503574 | 0.003896837 |
| GLTP | -1.184776594 | 0.000989972 | 0.00667075 |
| PTPN12 | -1.180952218 | 4.34E-05 | 0.000471332 |
| PLCG2 | -1.174986252 | 0.009789835 | 0.041471535 |
| EEF1D | -1.174925909 | 0.001259597 | 0.008092931 |
| GK5 | -1.173117703 | 0.002661906 | 0.014914883 |
| CD74 | -1.173018847 | 0.001282215 | 0.00821673 |
| SUGP2 | -1.172893536 | 0.007394071 | 0.033368553 |
| BCLAF1 | -1.170121139 | 2.10E-06 | 3.32E-05 |
| ZCCHC6 | -1.169311826 | 0.005721117 | 0.027273993 |
| CD99 | -1.168307819 | 3.32E-05 | 0.000374282 |
| GIMAP2 | -1.166079405 | 0.005643666 | 0.02697029 |
| XRN1 | -1.162459613 | 0.001039817 | 0.006950139 |
| KIAA2026 | -1.161707244 | 0.00023665 | 0.002029523 |
| SLC2A3 | -1.160103623 | 0.001525071 | 0.009398539 |
| HIPK1 | -1.149943174 | 0.004889857 | 0.024045841 |
| JAK1 | -1.148573827 | 6.99E-05 | 0.000712622 |
| FOXO3 | -1.148358826 | 0.00113277 | 0.007443806 |
| ZC3H13 | -1.144200726 | 0.000209093 | 0.001834893 |
| CIRBP | -1.143717811 | 7.10E-05 | 0.000722211 |
| CPNE1 | -1.143178105 | 0.003655229 | 0.019166995 |
| BTN3A2 | -1.140309617 | 0.000182339 | 0.001636279 |
| PHIP | -1.138821831 | 5.04E-05 | 0.000539096 |
| KANSL2 | -1.138709501 | 0.008992085 | 0.038898299 |
| NBPF8 | -1.137403792 | 0.001181613 | 0.007667197 |
| CLASP1 | -1.13737482 | 0.005790748 | 0.02756576 |
| GLG1 | -1.130661754 | 0.000212616 | 0.001860817 |
| THEMIS2 | -1.1270195 | 0.007607471 | 0.034174405 |

| | | | |
|---|---|---|---|
| VPS13B | -1.126816582 | 0.000624946 | 0.004610221 |
| ITGB1 | -1.112023212 | 0.00015279 | 0.001416848 |
| INPPL1 | -1.108493831 | 0.003855735 | 0.020004554 |
| PTPN4 | -1.106949319 | 0.000756872 | 0.00540049 |
| GAK | -1.104302493 | 0.009306218 | 0.039957684 |
| SNTB2 | -1.102399379 | 0.008697414 | 0.037924591 |
| ELF1 | -1.10164462 | 0.00147901 | 0.009201372 |
| OPTN | -1.097026644 | 0.000493571 | 0.003831035 |
| RPL13A | -1.096307346 | 9.52E-07 | 1.66E-05 |
| SP100 | -1.096306788 | 0.001130935 | 0.007436734 |
| FBXO9 | -1.092685133 | 0.005059458 | 0.024757279 |
| CHST12 | -1.091456602 | 0.003818873 | 0.019913602 |
| CD81 | -1.089970594 | 0.005854816 | 0.02778984 |
| CNOT2 | -1.088247455 | 0.003957478 | 0.020456702 |
| GSK3B | -1.085147648 | 0.002918985 | 0.016106951 |
| NAA16 | -1.082478461 | 0.008669853 | 0.0378486 |
| CNTRL | -1.079744562 | 0.001157107 | 0.007549042 |
| DHRS7 | -1.076342912 | 0.000368322 | 0.002983982 |
| OFD1 | -1.072006404 | 0.000181434 | 0.001636072 |
| TMEM181 | -1.066400638 | 0.004608652 | 0.023057363 |
| HLA-E | -1.06541516 | 2.61E-05 | 0.00030617 |
| ZNF800 | -1.061814675 | 0.00344339 | 0.018280899 |
| ITGA4 | -1.055996493 | 0.002068156 | 0.012080059 |
| RBM39 | -1.053810661 | 0.000522052 | 0.004004577 |
| ANKRD17 | -1.053710848 | 0.005275281 | 0.025608994 |
| RABGAP1 | -1.051065676 | 0.003777234 | 0.019753967 |
| ANKRD10 | -1.046768458 | 0.004885349 | 0.024045841 |
| MYO5A | -1.042364101 | 0.00222106 | 0.012750627 |
| EID1 | -1.042191259 | 0.00427983 | 0.021688746 |
| SAMD9L | -1.03736805 | 0.005959661 | 0.028164851 |
| ZBTB4 | -1.03335688 | 0.005392516 | 0.026051541 |
| ATMIN | -1.03031099 | 0.008500496 | 0.037314706 |
| RAB7A | -1.027058807 | 0.002460898 | 0.013923799 |
| GCC2 | -1.023224887 | 6.56E-05 | 0.000674438 |
| FBXO33 | -1.019227954 | 0.002952698 | 0.016238181 |
| ZC3H6 | -1.016551595 | 0.007414034 | 0.033412537 |
| ZNF292 | -1.012452734 | 0.004178176 | 0.02136117 |
| NIN | -1.010527651 | 0.008107478 | 0.035901455 |
| HNRNPUL2 | -1.00850095 | 0.001085267 | 0.007204031 |
| TES | -1.007438686 | 0.002640499 | 0.014823267 |
| TAF10 | -1.005823394 | 0.004286511 | 0.021711387 |
| MAN1A2 | -1.005579203 | 1.32E-05 | 0.000167604 |
| TNRC6B | -1.004091483 | 0.003032176 | 0.016554635 |
| SRSF6 | 1.003480865 | 0.001567553 | 0.009618053 |
| ANP32A | 1.004533303 | 0.000961644 | 0.006515668 |
| FOXN2 | 1.00704408 | 0.009621681 | 0.040971491 |
| RWDD4 | 1.009471932 | 0.00882476 | 0.038275738 |
| SSBP1 | 1.014958705 | 0.000375073 | 0.003026187 |
| POLR2L | 1.015908402 | 0.003057398 | 0.016664519 |
| UBR4 | 1.019244691 | 0.001902783 | 0.011261968 |

| | | | |
|---|---|---|---|
| SREBF2 | 1.023093995 | 0.008356447 | 0.036781115 |
| DHX9 | 1.025348874 | 0.005512604 | 0.026471489 |
| SATB1 | 1.028941839 | 0.000201823 | 0.001787082 |
| HSP90B1 | 1.034990897 | 3.19E-05 | 0.000362634 |
| MRPL33 | 1.039388536 | 0.000564949 | 0.004273311 |
| BCL2 | 1.040632157 | 0.007405234 | 0.033389951 |
| ZC3H15 | 1.043003838 | 0.000732605 | 0.005269494 |
| ZNF706 | 1.05613269 | 0.008761227 | 0.03809899 |
| RHEB | 1.056577594 | 0.002190952 | 0.012592518 |
| GOLGA7 | 1.056641144 | 0.002369388 | 0.013475982 |
| PTGER2 | 1.060576244 | 0.001885508 | 0.01118423 |
| PIK3CD | 1.062115798 | 0.000966947 | 0.006542565 |
| COX5A | 1.063640923 | 0.003454405 | 0.018319551 |
| FAM60A | 1.063649906 | 0.004476376 | 0.022487316 |
| TRMT112 | 1.067095631 | 0.000811052 | 0.005716401 |
| MIF | 1.068502363 | 0.008024665 | 0.035640558 |
| RDH11 | 1.06885581 | 0.003034601 | 0.016558658 |
| EIF4E | 1.070485211 | 0.001506325 | 0.009329893 |
| DARS | 1.074331231 | 0.004124996 | 0.021177568 |
| DDIT4 | 1.078028522 | 0.000543736 | 0.004135344 |
| NUP153 | 1.07842935 | 0.004676715 | 0.023255211 |
| IL21R | 1.080555328 | 0.003214146 | 0.017288369 |
| FAM96B | 1.080738222 | 0.00487066 | 0.023988976 |
| NAP1L1 | 1.084765727 | 0.000886448 | 0.006111693 |
| ACSL4 | 1.086954746 | 0.005450883 | 0.026240731 |
| BRK1 | 1.087673298 | 0.001751987 | 0.010539573 |
| DYNLRB1 | 1.089707212 | 0.00016413 | 0.001502127 |
| E2F4 | 1.08995393 | 0.001638719 | 0.009979811 |
| NME2 | 1.09094491 | 0.000879568 | 0.006072795 |
| API5 | 1.095521476 | 0.009712594 | 0.041215509 |
| PSMD8 | 1.099073646 | 0.006067275 | 0.028549658 |
| ANKLE2 | 1.099910371 | 0.002372483 | 0.013485763 |
| PRR5L | 1.099920148 | 0.004155402 | 0.021300236 |
| HAX1 | 1.101399691 | 0.006466123 | 0.029980687 |
| ELAVL1 | 1.105617635 | 0.002774641 | 0.015388355 |
| DAP | 1.106231485 | 0.003439143 | 0.018268237 |
| LGALS1 | 1.112564748 | 0.000412946 | 0.003299193 |
| PSMB3 | 1.114765703 | 0.001114628 | 0.007344269 |
| NDUFB3 | 1.115791588 | 0.009645526 | 0.041019615 |
| REEP5 | 1.116137704 | 0.003560309 | 0.018739372 |
| PSMA7 | 1.118377039 | 0.000188035 | 0.001680159 |
| MDM2 | 1.124504442 | 0.002386876 | 0.013551877 |
| GBP5 | 1.126407472 | 0.000288499 | 0.002413016 |
| SELT | 1.12720896 | 3.63E-05 | 0.000404118 |
| SRF | 1.128727715 | 0.000813613 | 0.005730334 |
| OAZ1 | 1.12939482 | 2.09E-05 | 0.00025018 |
| RPN1 | 1.131083826 | 0.00321226 | 0.017287703 |
| PHB | 1.133943819 | 0.004243616 | 0.021574191 |
| EIF4A1 | 1.134445139 | 0.000131117 | 0.001234536 |
| PARK7 | 1.134469247 | 0.000818661 | 0.005745267 |

| | | | |
|---|---|---|---|
| PDE6D | 1.138174372 | 0.001363097 | 0.008616843 |
| LIN7C | 1.141722257 | 0.007906189 | 0.035290092 |
| ATP5H | 1.143282445 | 0.000364409 | 0.002954724 |
| HINT1 | 1.145936732 | 3.93E-05 | 0.000433403 |
| PREPL | 1.145959089 | 0.008581471 | 0.037545302 |
| NPM1 | 1.146078535 | 0.000156246 | 0.001440719 |
| NDUFB6 | 1.148370045 | 0.00312194 | 0.016903618 |
| MCTS1 | 1.153418525 | 0.008186904 | 0.036197259 |
| NARS | 1.1572168 | 0.001864094 | 0.011070598 |
| KIAA0907 | 1.158445228 | 0.001842804 | 0.010957426 |
| ANXA7 | 1.162444714 | 0.000930016 | 0.00635403 |
| SRP19 | 1.170049985 | 0.000193689 | 0.001724399 |
| S100A10 | 1.170553753 | 0.000941722 | 0.006415105 |
| ID2 | 1.170717229 | 0.000277508 | 0.002335014 |
| XBP1 | 1.171332346 | 6.49E-05 | 0.000667781 |
| TNIP1 | 1.172356708 | 0.00714288 | 0.032579635 |
| ZNF37A | 1.174111521 | 0.003085053 | 0.016767407 |
| ESF1 | 1.176601615 | 0.006091686 | 0.028609638 |
| SNRPD1 | 1.180395519 | 0.003354936 | 0.01795705 |
| ISCU | 1.184699494 | 0.001768161 | 0.010616543 |
| ANKRD40 | 1.189766082 | 0.0012959 | 0.008288185 |
| GABARAPL2 | 1.1900089 | 0.00902572 | 0.039009402 |
| NOL11 | 1.19071167 | 0.007324196 | 0.033205957 |
| NDUFA1 | 1.200899872 | 4.14E-05 | 0.00045247 |
| SNRPB2 | 1.201870877 | 0.005114455 | 0.024951726 |
| GRPEL1 | 1.203451272 | 0.007441908 | 0.033522756 |
| TMEM50A | 1.205557837 | 0.000291476 | 0.00243376 |
| ABHD3 | 1.20720221 | 0.006121505 | 0.028694737 |
| ASNSD1 | 1.20938253 | 0.003823789 | 0.019913602 |
| PPP1R15B | 1.209628386 | 0.001097225 | 0.007258851 |
| EIF1AD | 1.211164189 | 0.007365694 | 0.03330176 |
| MCCC2 | 1.212135403 | 0.00391918 | 0.020269414 |
| NDUFS2 | 1.212432375 | 0.008265376 | 0.036511305 |
| RNF19A | 1.218428819 | 0.000151016 | 0.001403048 |
| PSMB2 | 1.21871862 | 0.009901309 | 0.0418534 |
| GNG5 | 1.21988104 | 0.000561207 | 0.004251741 |
| TMEM43 | 1.223121319 | 0.00236591 | 0.013465886 |
| ROMO1 | 1.224031543 | 0.007175734 | 0.032699084 |
| CCT8 | 1.225716402 | 8.05E-06 | 0.00010839 |
| COIL | 1.235324654 | 0.007186148 | 0.032711688 |
| LSM12 | 1.236066473 | 0.009055363 | 0.039103067 |
| TMEM64 | 1.236660631 | 0.00466866 | 0.023250876 |
| TOP1 | 1.236686533 | 1.15E-06 | 1.94E-05 |
| SNRPE | 1.240307225 | 0.003092144 | 0.016767407 |
| VDAC1 | 1.243070695 | 0.000682762 | 0.004969275 |
| EWSR1 | 1.243160615 | 2.63E-05 | 0.000308852 |
| CTSC | 1.243322404 | 0.000163374 | 0.001497998 |
| SDHB | 1.24387332 | 0.008760873 | 0.03809899 |
| PPIB | 1.243956319 | 0.003426334 | 0.018229809 |
| CCT3 | 1.24410254 | 0.004423222 | 0.02228877 |

| | | | |
|---|---|---|---|
| EHBP1 | 1.245960016 | 0.008051435 | 0.035711236 |
| COX7B | 1.246786784 | 0.000119115 | 0.001135705 |
| CREB1 | 1.248312518 | 0.000388962 | 0.003125394 |
| GABPA | 1.249038985 | 0.004255109 | 0.021608113 |
| TGS1 | 1.25032288 | 0.006564799 | 0.030337843 |
| SERPINB9 | 1.258119448 | 0.001882457 | 0.011172887 |
| ATXN1 | 1.258149435 | 0.00031287 | 0.002581641 |
| TMED9 | 1.260505456 | 1.01E-07 | 2.28E-06 |
| CEBPG | 1.262724633 | 0.002999482 | 0.016437722 |
| TTC39C | 1.263912227 | 0.001457256 | 0.009101105 |
| YWHAE | 1.264250476 | 0.001022467 | 0.006861436 |
| KRCC1 | 1.265948563 | 0.004396751 | 0.022189572 |
| AHSA1 | 1.266326692 | 0.008370802 | 0.036827774 |
| CDC123 | 1.266517346 | 0.003598824 | 0.018921794 |
| HGSNAT | 1.267002597 | 0.004163118 | 0.021328645 |
| CD48 | 1.267332411 | 6.02E-05 | 0.00062662 |
| SLC30A7 | 1.268162558 | 0.00060207 | 0.004478784 |
| CCZ1 | 1.269620409 | 0.006500921 | 0.030119508 |
| EIF3J | 1.27252916 | 6.34E-06 | 8.80E-05 |
| TBL1XR1 | 1.272595109 | 0.000503638 | 0.003896837 |
| C1orf43 | 1.275214336 | 0.002095415 | 0.012164568 |
| CYCS | 1.275390804 | 0.000629372 | 0.00463571 |
| SCAF4 | 1.278756838 | 0.004434614 | 0.022334703 |
| CDC42SE1 | 1.279275393 | 2.92E-05 | 0.00033804 |
| SEC22B | 1.279432558 | 6.03E-06 | 8.42E-05 |
| TMED5 | 1.280896712 | 0.000110363 | 0.001062585 |
| PIGY | 1.283149432 | 0.007364009 | 0.03330176 |
| BTBD10 | 1.283755925 | 0.004109125 | 0.02111819 |
| FAM126B | 1.286720474 | 3.28E-07 | 6.48E-06 |
| NDUFV2 | 1.289762951 | 0.002078032 | 0.012119201 |
| NDUFB4 | 1.291509839 | 0.001636992 | 0.009979811 |
| QKI | 1.292587406 | 5.12E-06 | 7.30E-05 |
| SART3 | 1.293172516 | 0.005975945 | 0.02821463 |
| PPP1CB | 1.296318587 | 6.37E-06 | 8.82E-05 |
| GNG10 | 1.300469601 | 0.006021731 | 0.028362555 |
| NDUFA11 | 1.301431222 | 0.007064 | 0.032234838 |
| GTF3C6 | 1.302004012 | 0.002067881 | 0.012080059 |
| NOP58 | 1.306647029 | 4.95E-05 | 0.000530587 |
| LYSMD2 | 1.307992966 | 9.36E-05 | 0.000922713 |
| TSPYL1 | 1.31041965 | 2.02E-06 | 3.22E-05 |
| AK2 | 1.310959047 | 0.000528893 | 0.004044403 |
| NDUFAB1 | 1.311814098 | 0.002923741 | 0.01611507 |
| SUSD6 | 1.315137991 | 0.000140708 | 0.001317262 |
| CD96 | 1.315460154 | 0.000992899 | 0.006685887 |
| SPTY2D1 | 1.315636294 | 0.001332247 | 0.008465462 |
| NUDT5 | 1.319485014 | 0.00738842 | 0.033363173 |
| ERI1 | 1.320008686 | 0.001021008 | 0.006856341 |
| FTSJ1 | 1.321714453 | 0.000724214 | 0.005217686 |
| USO1 | 1.32201125 | 0.000788073 | 0.005590585 |
| UBL5 | 1.32366249 | 0.000102401 | 0.000996684 |

| | | | |
|---|---|---|---|
| FAM107B | 1.323793893 | 7.51E-06 | 0.000101573 |
| NOMO2 | 1.323856254 | 0.002681321 | 0.014958583 |
| BRIX1 | 1.32469511 | 4.31E-05 | 0.000468398 |
| NCBP2 | 1.324715021 | 0.000724871 | 0.005217686 |
| PPP1R16B | 1.326110015 | 0.002092576 | 0.012162478 |
| SENP5 | 1.331213534 | 0.003168864 | 0.017091656 |
| PCNA | 1.331899375 | 0.003431011 | 0.018235622 |
| ATP5J2 | 1.333710744 | 0.001171288 | 0.007608863 |
| GOLGA8J | 1.333756498 | 0.005327735 | 0.025787079 |
| DDX18 | 1.333912494 | 0.000567292 | 0.004281309 |
| PDCD10 | 1.336250644 | 0.000575015 | 0.004322921 |
| LBH | 1.338806267 | 0.000182457 | 0.001636279 |
| BCCIP | 1.339059426 | 0.001489783 | 0.009250801 |
| EIF1AX | 1.341225836 | 0.000765898 | 0.00545695 |
| ZC3H4 | 1.344181819 | 0.006461363 | 0.02997278 |
| PRKCQ-AS1 | 1.344960687 | 0.001830066 | 0.010901505 |
| NEDD9 | 1.349035592 | 0.003554557 | 0.018719142 |
| PITPNB | 1.350994887 | 0.000861057 | 0.005980049 |
| HSP90AA1 | 1.352698549 | 8.07E-09 | 2.30E-07 |
| NASP | 1.355003221 | 0.000914144 | 0.006263032 |
| PDPK1 | 1.355312544 | 0.000639821 | 0.004695055 |
| IFRD1 | 1.356101616 | 0.001663957 | 0.010102158 |
| KCNQ1OT1 | 1.357754782 | 0.009382554 | 0.040232621 |
| GBP2 | 1.359692568 | 2.93E-06 | 4.43E-05 |
| LFNG | 1.360617592 | 0.002083992 | 0.012119769 |
| SOD1 | 1.36075461 | 1.44E-05 | 0.000180713 |
| YWHAG | 1.360905012 | 0.003511261 | 0.018566657 |
| CBX5 | 1.361280824 | 0.000336009 | 0.002744863 |
| CTNNB1 | 1.36175185 | 0.000162339 | 0.001489901 |
| FLOT1 | 1.363133707 | 0.000974091 | 0.006581823 |
| VASP | 1.366615836 | 7.46E-05 | 0.000754772 |
| BCL10 | 1.369950443 | 0.002008006 | 0.011803804 |
| ENO1 | 1.373013179 | 1.65E-06 | 2.67E-05 |
| LDLR | 1.373564297 | 0.002188456 | 0.012585548 |
| NOLC1 | 1.373740114 | 0.000772313 | 0.005490696 |
| SMIM15 | 1.383012168 | 0.000279016 | 0.002343685 |
| CAAP1 | 1.383144868 | 0.009868075 | 0.041748894 |
| ARFIP1 | 1.384500484 | 0.002607519 | 0.014660382 |
| STARD3 | 1.386437698 | 0.004308894 | 0.02180225 |
| METAP2 | 1.387573213 | 0.001166065 | 0.007586379 |
| PDCL3 | 1.387667872 | 0.008422578 | 0.037038956 |
| OST4 | 1.388287652 | 9.89E-05 | 0.000968704 |
| ATP13A3 | 1.391143395 | 0.00261624 | 0.014700991 |
| NDUFS5 | 1.392885355 | 0.009766771 | 0.041427491 |
| RNF145 | 1.393927556 | 3.78E-06 | 5.55E-05 |
| DHRS3 | 1.397035872 | 1.81E-05 | 0.00022005 |
| HNRNPF | 1.397139522 | 0.00088327 | 0.006094065 |
| HCP5 | 1.39906486 | 0.005317848 | 0.025751929 |
| DCUN1D1 | 1.400684456 | 0.000709098 | 0.005126723 |
| ATF4 | 1.401437162 | 4.08E-05 | 0.000448053 |

| | | | |
|---|---|---|---|
| TMEM258 | 1.403384884 | 7.89E-05 | 0.000795124 |
| SLMO2 | 1.403784829 | 0.000746456 | 0.005345609 |
| RTCB | 1.411444096 | 0.002151784 | 0.012425634 |
| ZNF354A | 1.414219886 | 0.009527287 | 0.040710893 |
| RNF219 | 1.415453493 | 0.005821875 | 0.027660249 |
| IARS | 1.420981991 | 0.001643612 | 0.00999271 |
| LPXN | 1.423116268 | 8.26E-08 | 1.90E-06 |
| MED17 | 1.424467118 | 0.002249354 | 0.012897963 |
| SRSF7 | 1.425375133 | 6.07E-07 | 1.11E-05 |
| CCT7 | 1.428804379 | 7.62E-07 | 1.36E-05 |
| MTF1 | 1.429649993 | 0.004689612 | 0.023293977 |
| FTH1 | 1.431433155 | 2.29E-07 | 4.71E-06 |
| CD53 | 1.432376796 | 1.43E-07 | 3.07E-06 |
| GPATCH4 | 1.433251156 | 0.005475192 | 0.026331915 |
| HPRT1 | 1.435498319 | 0.005047627 | 0.024724047 |
| PPARA | 1.435816791 | 0.008635926 | 0.037740342 |
| ETNK1 | 1.438375073 | 0.000424506 | 0.003377801 |
| MAPK6 | 1.439529645 | 0.006079548 | 0.028584217 |
| TNFAIP3 | 1.443067847 | 0.002366241 | 0.013465886 |
| SPAG1 | 1.443620638 | 0.000917594 | 0.006282283 |
| PSMD1 | 1.447440871 | 1.75E-05 | 0.000214093 |
| ZNF562 | 1.44765126 | 0.009348096 | 0.040102393 |
| C12orf5 | 1.452912747 | 0.006989138 | 0.031952672 |
| CCT6A | 1.454097991 | 1.19E-07 | 2.64E-06 |
| NOB1 | 1.454216512 | 0.004862699 | 0.023985891 |
| CMC2 | 1.454409988 | 0.003980254 | 0.02053381 |
| FURIN | 1.455535365 | 0.001260648 | 0.008094385 |
| SLA | 1.456004533 | 0.000934813 | 0.006377922 |
| CCDC43 | 1.4565844 | 0.001813015 | 0.010832819 |
| SLC39A6 | 1.463051107 | 0.001257681 | 0.008085915 |
| GPX4 | 1.464421008 | 0.001328431 | 0.008452162 |
| VPS72 | 1.46544572 | 0.001610711 | 0.009845913 |
| NOL7 | 1.465473395 | 0.001452594 | 0.009094706 |
| FAM102B | 1.465900921 | 0.005112985 | 0.024951726 |
| SPCS2 | 1.466407141 | 5.69E-05 | 0.000596838 |
| PSMD11 | 1.467104089 | 9.45E-06 | 0.000125567 |
| CCT4 | 1.468281172 | 8.97E-05 | 0.000890247 |
| GTF2H5 | 1.470007383 | 0.008431875 | 0.03706323 |
| TAB2 | 1.472467781 | 0.000174695 | 0.001586975 |
| CWC25 | 1.473365511 | 0.000776505 | 0.005516503 |
| WRNIP1 | 1.479918232 | 0.000269726 | 0.002277353 |
| ARPC1A | 1.479994938 | 0.002496032 | 0.014107495 |
| KPNA1 | 1.481688026 | 0.000181913 | 0.001636279 |
| HS6ST1 | 1.482580912 | 0.005093703 | 0.024883592 |
| FDFT1 | 1.482700788 | 3.23E-05 | 0.000365389 |
| SSR3 | 1.483410085 | 0.000219193 | 0.001904782 |
| C19orf10 | 1.486920436 | 0.001381276 | 0.008709318 |
| RPS6KA3 | 1.48824375 | 2.52E-06 | 3.91E-05 |
| SLC30A5 | 1.488513113 | 0.001629031 | 0.009939318 |
| MAT2A | 1.488754448 | 4.79E-05 | 0.000516029 |

| | | | |
|---|---|---|---|
| KIN | 1.490123019 | 0.008288652 | 0.036581179 |
| NFE2L1 | 1.49130731 | 0.000565361 | 0.004273311 |
| LMAN1 | 1.493103344 | 0.001230108 | 0.007937399 |
| ZNF720 | 1.495620486 | 0.001512738 | 0.009351905 |
| CKS1B | 1.495784153 | 0.007750367 | 0.034720937 |
| CACYBP | 1.498195581 | 5.80E-06 | 8.14E-05 |
| CRAMP1L | 1.50076813 | 0.006211396 | 0.029045767 |
| GBP1 | 1.501110187 | 0.003237196 | 0.017402813 |
| PABPC4 | 1.511239124 | 7.89E-05 | 0.000795124 |
| SNX10 | 1.51334044 | 0.003087485 | 0.016767407 |
| IMPA1 | 1.513675527 | 0.009811935 | 0.041547213 |
| SNHG15 | 1.515333969 | 0.007934094 | 0.035334268 |
| NUTF2 | 1.516648722 | 0.002694099 | 0.015009544 |
| PIGX | 1.516928677 | 0.000175896 | 0.00159641 |
| MRPL57 | 1.52146887 | 0.000416589 | 0.003322888 |
| RCC2 | 1.521521992 | 6.52E-07 | 1.19E-05 |
| PLEK | 1.524138669 | 3.56E-05 | 0.000397351 |
| TCP1 | 1.528796579 | 6.81E-06 | 9.31E-05 |
| EMC7 | 1.529158294 | 0.001797133 | 0.01075102 |
| FAM98B | 1.529949398 | 0.002532699 | 0.0143053 |
| ENY2 | 1.53079806 | 1.46E-06 | 2.40E-05 |
| EIF2AK4 | 1.530971111 | 0.00338694 | 0.018079034 |
| PRNP | 1.533770555 | 0.00027598 | 0.002324153 |
| TMED7 | 1.536566699 | 0.002847887 | 0.015767844 |
| WWC3 | 1.537557827 | 0.000303472 | 0.002518922 |
| RRP15 | 1.537943048 | 0.006343453 | 0.029537549 |
| HYPK | 1.538687858 | 0.003809646 | 0.019881083 |
| EDEM3 | 1.540369011 | 0.001262498 | 0.008100959 |
| HIPK2 | 1.541965394 | 1.60E-09 | 5.25E-08 |
| STXBP3 | 1.542951462 | 0.008313864 | 0.036643001 |
| WBSCR16 | 1.545321178 | 0.009575547 | 0.040845953 |
| DGUOK | 1.546289546 | 3.05E-05 | 0.000349486 |
| RYBP | 1.546482256 | 1.67E-05 | 0.000204794 |
| RAB21 | 1.54723539 | 0.000913618 | 0.006263032 |
| ATP8B4 | 1.547821565 | 0.003477695 | 0.018423144 |
| PSMD2 | 1.551051435 | 0.000818183 | 0.005745267 |
| PRDX6 | 1.554707522 | 0.000122762 | 0.001163686 |
| RRN3 | 1.556332183 | 4.25E-06 | 6.14E-05 |
| DNAJC3 | 1.556622249 | 2.00E-07 | 4.20E-06 |
| BTG3 | 1.557177193 | 0.003045331 | 0.016607972 |
| GNAQ | 1.557198499 | 0.003822296 | 0.019913602 |
| SIPA1L1 | 1.558175747 | 0.001406363 | 0.008861806 |
| SC5D | 1.558735783 | 0.003450638 | 0.018309467 |
| MIS12 | 1.558755828 | 0.004751464 | 0.023543742 |
| TIGIT | 1.560077838 | 9.40E-05 | 0.000926285 |
| CCNJ | 1.560392116 | 0.008305794 | 0.036623884 |
| ATG5 | 1.563029522 | 0.004246071 | 0.021574191 |
| DNAJC16 | 1.563040639 | 0.001953007 | 0.011501169 |
| TESPA1 | 1.566376627 | 0.009822344 | 0.041573346 |
| TM6SF1 | 1.569344635 | 0.001715444 | 0.01035706 |

| | | | |
|---|---|---|---|
| SAMD10 | 1.572693083 | 0.008059716 | 0.035731531 |
| HSPA5 | 1.577244311 | 1.38E-11 | 6.76E-10 |
| FTL | 1.577264069 | 9.06E-10 | 3.13E-08 |
| CPSF4 | 1.578865259 | 0.003882423 | 0.020117322 |
| PPP1R10 | 1.580001587 | 0.002117609 | 0.012271626 |
| SQSTM1 | 1.58127924 | 5.45E-07 | 1.01E-05 |
| CCT2 | 1.584828337 | 1.96E-06 | 3.13E-05 |
| KIAA0020 | 1.585294735 | 9.65E-05 | 0.00094849 |
| PSMA2 | 1.585810198 | 0.000134518 | 0.001261717 |
| NUS1 | 1.586335815 | 2.19E-05 | 0.000261349 |
| H2AFZ | 1.586443686 | 2.90E-07 | 5.76E-06 |
| KDM5B | 1.588051316 | 0.000835822 | 0.005844794 |
| TSG101 | 1.588259459 | 0.000760031 | 0.005419091 |
| DNAJB9 | 1.589404262 | 4.58E-06 | 6.60E-05 |
| EML2 | 1.591292942 | 0.003012459 | 0.016456144 |
| ZFP36L1 | 1.592040304 | 3.97E-07 | 7.63E-06 |
| PTGES3 | 1.593258631 | 4.98E-07 | 9.34E-06 |
| TRIM13 | 1.596310933 | 2.12E-05 | 0.000253703 |
| CHM | 1.596874328 | 0.00718652 | 0.032711688 |
| NOSIP | 1.597013974 | 0.009416606 | 0.040360995 |
| BPGM | 1.599743704 | 2.87E-05 | 0.000333341 |
| PDIA4 | 1.60203741 | 7.13E-05 | 0.000724626 |
| FARSB | 1.60304144 | 0.008583642 | 0.037545302 |
| PRPF19 | 1.60401675 | 0.004531466 | 0.022729149 |
| SKIV2L2 | 1.604571995 | 0.000159859 | 0.001468518 |
| CCDC47 | 1.608045232 | 0.001831992 | 0.010906358 |
| CYP51A1 | 1.612114052 | 5.08E-05 | 0.000541971 |
| RNF34 | 1.612258229 | 0.000154445 | 0.001426704 |
| TBCA | 1.615208232 | 1.06E-05 | 0.000138911 |
| IER3IP1 | 1.61849051 | 3.69E-07 | 7.18E-06 |
| IFNAR1 | 1.61872416 | 0.003073177 | 0.016740379 |
| EI24 | 1.620029667 | 4.10E-06 | 5.97E-05 |
| HDAC2 | 1.622479569 | 2.99E-05 | 0.000343959 |
| CHST11 | 1.625172764 | 0.000121356 | 0.001152589 |
| CHORDC1 | 1.625849245 | 1.57E-06 | 2.57E-05 |
| USP14 | 1.628467966 | 4.28E-05 | 0.00046677 |
| ARHGEF2 | 1.62891696 | 0.000675457 | 0.004924343 |
| NIFK | 1.629041455 | 0.002773459 | 0.015388355 |
| TP53RK | 1.629876569 | 1.89E-05 | 0.000227878 |
| TOB2 | 1.631354376 | 0.000415083 | 0.003313575 |
| KRTCAP2 | 1.633124697 | 5.47E-06 | 7.75E-05 |
| UQCRQ | 1.633303287 | 7.18E-05 | 0.000729127 |
| PSMD14 | 1.634599809 | 0.000237995 | 0.002039276 |
| ATP6V1D | 1.635131498 | 0.00442053 | 0.022286649 |
| CREM | 1.638057942 | 0.000600469 | 0.004476596 |
| LINC01004 | 1.638277382 | 0.004790347 | 0.023682896 |
| PFKP | 1.638990729 | 0.002764103 | 0.015347263 |
| SGPL1 | 1.639384482 | 1.38E-05 | 0.000174773 |
| MRPL32 | 1.641898345 | 8.40E-05 | 0.000840533 |
| MDH1 | 1.641941831 | 0.00082493 | 0.00578099 |

| | | | |
|---|---|---|---|
| ZSWIM6 | 1.642305392 | 0.000815029 | 0.005732076 |
| HNRNPAB | 1.644856706 | 2.94E-05 | 0.000338819 |
| CMTM6 | 1.645831198 | 9.23E-08 | 2.11E-06 |
| TMEM263 | 1.650266147 | 0.000131069 | 0.001234536 |
| FHOD1 | 1.656436841 | 0.000261879 | 0.002226425 |
| OSTC | 1.656756007 | 4.76E-05 | 0.000513594 |
| KPNB1 | 1.659745886 | 3.85E-12 | 2.02E-10 |
| POC1B-GALNT4 | 1.660760138 | 3.01E-05 | 0.000345873 |
| AP1AR | 1.661292846 | 6.79E-05 | 0.000695282 |
| SFPQ | 1.663632503 | 1.21E-06 | 2.02E-05 |
| ANKRD20A9P | 1.667495788 | 0.000234387 | 0.002011874 |
| PPP2CB | 1.672187702 | 0.000178335 | 0.001614066 |
| MPHOSPH6 | 1.678023652 | 0.006141323 | 0.028773886 |
| MCTP1 | 1.678603825 | 0.001069621 | 0.007119441 |
| EAF1 | 1.681835129 | 1.26E-06 | 2.10E-05 |
| LDHA | 1.681842844 | 5.40E-07 | 1.01E-05 |
| CAMTA1 | 1.682218187 | 0.001640354 | 0.009983566 |
| EIF4G1 | 1.683283139 | 0.000107417 | 0.001038292 |
| HYOU1 | 1.68474903 | 1.51E-05 | 0.000188115 |
| NIP7 | 1.685760485 | 0.000802252 | 0.005670672 |
| SEC61A1 | 1.685773222 | 0.000107941 | 0.001042329 |
| PSMC6 | 1.686220893 | 3.14E-06 | 4.70E-05 |
| UBE2D1 | 1.687752963 | 0.005950892 | 0.028136967 |
| ODC1 | 1.689705586 | 0.003090615 | 0.016767407 |
| TOMM5 | 1.691696495 | 1.44E-05 | 0.000180713 |
| COMT | 1.692721503 | 0.000319237 | 0.002629756 |
| CLTA | 1.694150843 | 0.000587846 | 0.004401418 |
| EIF5A | 1.694863043 | 2.45E-09 | 7.73E-08 |
| SMARCAD1 | 1.696990176 | 0.00063666 | 0.004675356 |
| C16orf72 | 1.700927065 | 1.46E-05 | 0.000183151 |
| ZNF680 | 1.70146204 | 0.003494801 | 0.018503776 |
| FAM210A | 1.70416244 | 0.006354025 | 0.029545133 |
| APOL2 | 1.705436606 | 0.000593732 | 0.004433111 |
| CISD2 | 1.705686221 | 0.000307962 | 0.002549713 |
| SLAMF7 | 1.707273116 | 1.76E-07 | 3.72E-06 |
| TAP1 | 1.707681172 | 0.000159391 | 0.001466962 |
| ZNF518B | 1.708393713 | 0.001156572 | 0.007549042 |
| RQCD1 | 1.71126153 | 1.61E-09 | 5.25E-08 |
| TRMT6 | 1.713338159 | 0.001979549 | 0.011650485 |
| KDELR2 | 1.714325607 | 0.000116563 | 0.001117889 |
| UNC50 | 1.714469956 | 0.001505782 | 0.009329893 |
| DCP1A | 1.716982352 | 9.10E-07 | 1.59E-05 |
| PMF1 | 1.717572972 | 0.00379816 | 0.019842252 |
| HSPA8 | 1.718787849 | 7.34E-10 | 2.56E-08 |
| POMP | 1.720076398 | 1.23E-07 | 2.72E-06 |
| NAA15 | 1.723177323 | 3.50E-06 | 5.19E-05 |
| RAB10 | 1.724102349 | 9.00E-05 | 0.000892995 |
| SFXN1 | 1.725516204 | 0.002150974 | 0.012425634 |
| PDIA6 | 1.727660066 | 6.85E-11 | 3.03E-09 |
| FCHSD2 | 1.730805089 | 0.000384922 | 0.003098002 |

| | | | |
|---|---|---|---|
| EIF2S2 | 1.732992873 | 3.82E-07 | 7.40E-06 |
| RIOK1 | 1.737844328 | 0.00026688 | 0.002257206 |
| RDX | 1.738330721 | 1.58E-05 | 0.00019537 |
| PRPSAP2 | 1.739288493 | 1.47E-05 | 0.000183151 |
| GAR1 | 1.743156688 | 0.000924347 | 0.006324107 |
| PIP5K1C | 1.745841895 | 0.000724725 | 0.005217686 |
| ZNF430 | 1.750754633 | 0.000879537 | 0.006072795 |
| BCL3 | 1.751803724 | 0.00723836 | 0.032892798 |
| DCTPP1 | 1.751950219 | 0.002067826 | 0.012080059 |
| NOM1 | 1.752351906 | 0.007707117 | 0.034542954 |
| SNHG16 | 1.752561175 | 0.000158918 | 0.001463989 |
| DKC1 | 1.754874083 | 2.80E-07 | 5.59E-06 |
| NDC1 | 1.755432686 | 0.006342285 | 0.029537549 |
| WDR89 | 1.756870914 | 0.000214449 | 0.001870186 |
| DBR1 | 1.757641019 | 0.00608336 | 0.028584217 |
| SLC39A8 | 1.760906848 | 0.002364738 | 0.013465886 |
| PDE4B | 1.762539565 | 0.009615155 | 0.04096148 |
| PROSC | 1.762540283 | 0.000721691 | 0.005207325 |
| TAF8 | 1.762735985 | 0.009569137 | 0.040845953 |
| SNRNP35 | 1.76407465 | 0.002050824 | 0.012019493 |
| NCOA3 | 1.767532021 | 1.91E-06 | 3.07E-05 |
| NUPL1 | 1.767616872 | 0.00181226 | 0.010832819 |
| ST8SIA4 | 1.768186858 | 0.000249458 | 0.002128203 |
| GNL3 | 1.769178647 | 2.93E-05 | 0.000338444 |
| TMEM208 | 1.770945084 | 0.002149941 | 0.012425634 |
| IRF2BP2 | 1.772017681 | 3.24E-09 | 1.01E-07 |
| MRPS23 | 1.772744531 | 0.005065279 | 0.024773407 |
| DEGS1 | 1.773078149 | 0.000460216 | 0.003612146 |
| CD44 | 1.773956499 | 2.91E-08 | 7.41E-07 |
| CCT5 | 1.775335869 | 3.83E-07 | 7.40E-06 |
| DNAJA1 | 1.780315847 | 3.87E-05 | 0.000428941 |
| GLO1 | 1.781097704 | 0.000146935 | 0.00137032 |
| TIMM8B | 1.785146137 | 0.005068742 | 0.024777989 |
| HSPA13 | 1.785714881 | 0.000742101 | 0.005322187 |
| ASH2L | 1.791765205 | 0.000833803 | 0.005834838 |
| PSMA5 | 1.796874852 | 8.36E-06 | 0.000112042 |
| DUSP11 | 1.799228365 | 0.000483877 | 0.00377407 |
| SQRDL | 1.804486914 | 0.00107652 | 0.007155652 |
| GTPBP10 | 1.804847369 | 6.09E-05 | 0.000633644 |
| FZD3 | 1.809925356 | 0.008438502 | 0.037075748 |
| AFG3L1P | 1.810832254 | 6.77E-05 | 0.000693784 |
| BACH2 | 1.813350912 | 0.001903205 | 0.011261968 |
| COPS7A | 1.81545227 | 0.005541516 | 0.026559755 |
| ABCE1 | 1.818001262 | 2.86E-06 | 4.35E-05 |
| EIF4A3 | 1.818327473 | 0.000134388 | 0.001261717 |
| ING5 | 1.819164938 | 0.007689383 | 0.034494987 |
| SMIM3 | 1.819361456 | 0.005204331 | 0.025320624 |
| VIMP | 1.822765087 | 5.47E-06 | 7.75E-05 |
| SHFM1 | 1.829268736 | 2.47E-06 | 3.83E-05 |
| GSTO1 | 1.832459091 | 0.000891404 | 0.006137236 |

| | | | |
|---|---|---|---|
| IER5 | 1.832729423 | 2.25E-06 | 3.54E-05 |
| RAN | 1.833398517 | 2.23E-07 | 4.61E-06 |
| NFAT5 | 1.848368398 | 7.54E-08 | 1.75E-06 |
| DNAJA3 | 1.849704946 | 0.000800887 | 0.005665106 |
| NUDT9 | 1.852084612 | 0.005392996 | 0.026051541 |
| TRIM4 | 1.852823442 | 6.21E-05 | 0.000642796 |
| ZNF738 | 1.8545317 | 0.007844588 | 0.035094962 |
| MEX3C | 1.855933095 | 2.67E-10 | 1.03E-08 |
| SDF2L1 | 1.85709512 | 5.55E-05 | 0.000583557 |
| MPC1 | 1.858543447 | 0.001709969 | 0.010330359 |
| WDR75 | 1.873638403 | 0.00033743 | 0.002751894 |
| NAA20 | 1.87536004 | 6.51E-06 | 8.95E-05 |
| IPO5 | 1.876828001 | 5.69E-05 | 0.000596672 |
| MRPL13 | 1.878178288 | 0.003573221 | 0.018797248 |
| SLC3A2 | 1.878460464 | 4.49E-07 | 8.51E-06 |
| NMD3 | 1.878920692 | 0.000201686 | 0.001787082 |
| TOMM70A | 1.879929173 | 0.000541347 | 0.004123567 |
| TCEB3 | 1.8804371 | 3.16E-05 | 0.000359765 |
| DPM1 | 1.882129268 | 0.00012411 | 0.001175328 |
| IMMP1L | 1.884779734 | 0.002697942 | 0.015022425 |
| STK17B | 1.886564803 | 1.27E-11 | 6.21E-10 |
| SS18L2 | 1.895853302 | 0.00257133 | 0.014505487 |
| TMEM243 | 1.896295279 | 0.005773357 | 0.027496312 |
| MARS | 1.899798177 | 1.63E-08 | 4.36E-07 |
| MGC27345 | 1.900287343 | 0.001910302 | 0.011292817 |
| SNRPF | 1.9030533 | 3.76E-07 | 7.30E-06 |
| HERPUD1 | 1.903854931 | 1.22E-05 | 0.000156474 |
| TRAF3 | 1.903986426 | 0.001154816 | 0.007548232 |
| ME2 | 1.904507898 | 2.51E-05 | 0.000295486 |
| RPRD1B | 1.90688965 | 0.009145929 | 0.039407424 |
| LSM5 | 1.910075278 | 1.83E-05 | 0.000221644 |
| SEC23B | 1.918171038 | 0.000336352 | 0.002745386 |
| PSMC3 | 1.919218673 | 6.16E-05 | 0.000639217 |
| PPAP2A | 1.931547975 | 0.001337161 | 0.00849119 |
| PTGER4 | 1.934982578 | 0.000251537 | 0.002144072 |
| TRMT1L | 1.935430909 | 0.002865693 | 0.015839611 |
| NDUFB5 | 1.938243158 | 0.001418637 | 0.008916237 |
| BCL2L11 | 1.943306807 | 0.00078991 | 0.005599574 |
| IFIT3 | 1.944017591 | 0.000605461 | 0.004493327 |
| FAM174A | 1.945670508 | 0.004215875 | 0.021475571 |
| GGCT | 1.946518228 | 0.000248128 | 0.002118698 |
| SLIRP | 1.949175936 | 0.000902219 | 0.006207342 |
| SPRYD3 | 1.950223628 | 0.004469621 | 0.022476399 |
| IDH3A | 1.952118972 | 0.005853034 | 0.02778984 |
| MCM6 | 1.95311884 | 0.001735228 | 0.010457199 |
| PSMA6 | 1.954188412 | 3.11E-05 | 0.000355715 |
| LOC100505549 | 1.954335615 | 0.000317805 | 0.002620155 |
| TIFA | 1.954738891 | 0.000444092 | 0.003502404 |
| ADPRHL2 | 1.955962377 | 0.001769668 | 0.010619089 |
| TGIF1 | 1.956259018 | 0.00559386 | 0.026780048 |

| | | | |
|---|---|---|---|
| TMEM33 | 1.957298101 | 3.32E-05 | 0.000374282 |
| FASN | 1.960408685 | 0.000254545 | 0.002167831 |
| PRDM1 | 1.960998361 | 4.82E-09 | 1.42E-07 |
| MCMBP | 1.966019709 | 1.66E-05 | 0.000204006 |
| GABPB1 | 1.968068805 | 1.97E-06 | 3.15E-05 |
| MARCKSL1 | 1.97158206 | 0.001355512 | 0.008591036 |
| LTB | 1.971767472 | 0.00011091 | 0.001066806 |
| GPR180 | 1.972243212 | 9.89E-06 | 0.000130608 |
| HSPH1 | 1.972468303 | 5.25E-10 | 1.90E-08 |
| MED19 | 1.978176978 | 0.000555789 | 0.004215885 |
| CDCA4 | 1.988727985 | 0.000798979 | 0.005655685 |
| TIMM17A | 1.9894063 | 0.007355969 | 0.03330176 |
| PDDC1 | 1.990232398 | 0.008326652 | 0.036682885 |
| RABGGTB | 1.99166629 | 0.000433627 | 0.003430898 |
| SESN2 | 1.997166326 | 5.53E-05 | 0.000581913 |
| C15orf57 | 1.999014088 | 0.003163552 | 0.017077306 |
| GTF2H2C | 2.001060482 | 0.005095417 | 0.024883592 |
| NANS | 2.002146107 | 0.001049368 | 0.007000066 |
| IGFLR1 | 2.008037961 | 0.002854604 | 0.015787213 |
| CALR | 2.010195385 | 3.71E-12 | 1.95E-10 |
| UTP6 | 2.012203771 | 1.79E-05 | 0.000217212 |
| GTF2H1 | 2.01222642 | 0.00018441 | 0.001649264 |
| SELK | 2.014363457 | 6.69E-07 | 1.21E-05 |
| GARS | 2.027539145 | 1.25E-05 | 0.000159111 |
| PSMD5 | 2.027763752 | 4.07E-08 | 1.00E-06 |
| SERF1A | 2.028056477 | 0.003794153 | 0.019831877 |
| SNRPG | 2.028329879 | 2.51E-05 | 0.000295486 |
| ARCN1 | 2.028922671 | 1.82E-09 | 5.87E-08 |
| PSMA1 | 2.034200069 | 1.18E-11 | 5.88E-10 |
| SLC25A44 | 2.038699123 | 0.000208846 | 0.001834371 |
| SOCS2 | 2.04026815 | 0.001250609 | 0.00804572 |
| TUBGCP4 | 2.0403147 | 0.007580385 | 0.034099567 |
| HSP90AB1 | 2.040995015 | 8.47E-12 | 4.26E-10 |
| TUBA1A | 2.043338348 | 1.10E-08 | 3.03E-07 |
| TDP1 | 2.045168973 | 0.000852011 | 0.00593263 |
| ARF4 | 2.046071899 | 3.62E-08 | 8.96E-07 |
| NDUFC1 | 2.046450475 | 0.000350382 | 0.002848077 |
| APOBEC3G | 2.054046386 | 1.16E-05 | 0.000149216 |
| ADAM17 | 2.054266965 | 3.89E-06 | 5.70E-05 |
| POLH | 2.055884638 | 0.001144364 | 0.007494894 |
| ZNF274 | 2.058964075 | 0.009079304 | 0.039171966 |
| TEX10 | 2.063271879 | 0.001952939 | 0.011501169 |
| MRPL3 | 2.063880703 | 5.45E-07 | 1.01E-05 |
| COX17 | 2.066499411 | 2.60E-10 | 1.01E-08 |
| CIRH1A | 2.068674428 | 0.000948943 | 0.006447497 |
| DFFA | 2.069172245 | 0.003847875 | 0.020004554 |
| MRPS14 | 2.070387781 | 0.002967187 | 0.016290471 |
| KCNA3 | 2.072191597 | 1.26E-07 | 2.75E-06 |
| RPA3 | 2.078022701 | 0.006110208 | 0.02866918 |
| SEC11C | 2.084893296 | 1.05E-09 | 3.56E-08 |

| | | | |
|---|---|---|---|
| MED28 | 2.090986028 | 4.57E-05 | 0.000494003 |
| RPF2 | 2.091290576 | 0.000608432 | 0.004506756 |
| ASAP1 | 2.094351243 | 2.62E-09 | 8.20E-08 |
| ZC3H12A | 2.096998762 | 2.47E-08 | 6.45E-07 |
| IL6ST | 2.097440773 | 8.73E-06 | 0.000116655 |
| GID4 | 2.099176169 | 0.001828603 | 0.010899406 |
| LEO1 | 2.104567795 | 3.88E-07 | 7.50E-06 |
| EIF2S1 | 2.109230388 | 3.12E-05 | 0.000356344 |
| PNO1 | 2.110874306 | 0.001111316 | 0.007332297 |
| USP31 | 2.111680259 | 0.00146011 | 0.009101105 |
| TMEM165 | 2.113012556 | 1.08E-06 | 1.85E-05 |
| MGAT2 | 2.114220169 | 2.20E-08 | 5.79E-07 |
| AKAP1 | 2.11884255 | 0.00354379 | 0.018682495 |
| GCLM | 2.122026971 | 2.81E-06 | 4.28E-05 |
| APOBEC3D | 2.130671608 | 8.16E-06 | 0.0001097 |
| MFHAS1 | 2.131108143 | 0.005200291 | 0.025320127 |
| LARP4 | 2.132157029 | 6.79E-07 | 1.23E-05 |
| RBL1 | 2.138267315 | 0.000371217 | 0.003002479 |
| FKBP2 | 2.140765224 | 8.43E-09 | 2.39E-07 |
| LETM1 | 2.142147389 | 2.33E-06 | 3.64E-05 |
| TMEM185B | 2.142725399 | 0.00799346 | 0.035566364 |
| RPL7L1 | 2.143561122 | 0.002239197 | 0.012847228 |
| HSPD1 | 2.143722118 | 1.67E-09 | 5.45E-08 |
| AARS | 2.146697995 | 0.006648263 | 0.030665777 |
| UPP1 | 2.15695526 | 1.47E-07 | 3.15E-06 |
| MRPS36 | 2.162577703 | 4.71E-06 | 6.77E-05 |
| PRDX1 | 2.163165805 | 0.000182007 | 0.001636279 |
| ZC3H8 | 2.165845705 | 5.19E-05 | 0.000552478 |
| PRMT6 | 2.166554101 | 0.004627415 | 0.023127647 |
| CCNYL1 | 2.167155063 | 0.004080621 | 0.02098269 |
| SEC22A | 2.173076558 | 0.001944199 | 0.011469956 |
| CHSY1 | 2.17318953 | 3.81E-09 | 1.15E-07 |
| MAML2 | 2.175601742 | 0.002681903 | 0.014958583 |
| ELL | 2.177939703 | 0.000242861 | 0.002075534 |
| HN1L | 2.178499093 | 0.001444565 | 0.009052866 |
| NDUFAF4 | 2.179403566 | 3.46E-05 | 0.000388058 |
| TFRC | 2.187726779 | 0.000705341 | 0.005110856 |
| NCOA5 | 2.196320008 | 5.32E-05 | 0.00056376 |
| TDG | 2.197876043 | 1.36E-06 | 2.26E-05 |
| XCL2 | 2.199377867 | 1.40E-07 | 3.03E-06 |
| SCD | 2.201703378 | 0.000516724 | 0.003980659 |
| PITRM1 | 2.203960347 | 0.000294036 | 0.002448889 |
| HMGCS1 | 2.211886011 | 1.78E-05 | 0.000216328 |
| PDP2 | 2.21790921 | 0.003394213 | 0.018092388 |
| PUS7 | 2.220153276 | 0.001520144 | 0.009380159 |
| SEC61G | 2.223976707 | 3.90E-11 | 1.76E-09 |
| IRAK2 | 2.226743349 | 2.93E-06 | 4.43E-05 |
| HOXB2 | 2.229148495 | 0.00094814 | 0.006447497 |
| SURF4 | 2.229163502 | 4.64E-11 | 2.09E-09 |
| SLC4A10 | 2.232630427 | 0.00589046 | 0.027905023 |

| | | | |
|---|---|---|---|
| TXNRD1 | 2.232689763 | 3.50E-09 | 1.07E-07 |
| TFE3 | 2.232720991 | 0.001128318 | 0.0074245 |
| PSMD5-AS1 | 2.233079864 | 0.000942225 | 0.006415105 |
| LRIF1 | 2.235608086 | 3.54E-05 | 0.00039562 |
| G3BP1 | 2.241926158 | 3.46E-08 | 8.64E-07 |
| IKBIP | 2.243466745 | 1.73E-05 | 0.000211077 |
| LRR1 | 2.244408464 | 0.001331903 | 0.008465462 |
| HMOX1 | 2.250436615 | 0.009520839 | 0.040701068 |
| ZNF268 | 2.256961294 | 0.003171953 | 0.017098919 |
| KLHL15 | 2.258339965 | 0.000486357 | 0.003790027 |
| MSMO1 | 2.25989766 | 4.98E-08 | 1.20E-06 |
| HSPB1 | 2.27033682 | 4.70E-09 | 1.39E-07 |
| LCP2 | 2.271419888 | 4.52E-18 | 5.47E-16 |
| GTPBP4 | 2.274183977 | 0.000903773 | 0.006213676 |
| ITGAE | 2.275082707 | 0.000483924 | 0.00377407 |
| LRRC8C | 2.281437911 | 8.03E-10 | 2.78E-08 |
| MPZL3 | 2.283107747 | 0.009323078 | 0.040012561 |
| LMNB2 | 2.283646398 | 0.000265376 | 0.002248366 |
| BHLHE40-AS1 | 2.285581076 | 0.003609926 | 0.018969996 |
| LSM6 | 2.290422652 | 2.71E-05 | 0.000316683 |
| FBXO30 | 2.290443546 | 2.63E-07 | 5.32E-06 |
| CBWD5 | 2.292621656 | 6.31E-06 | 8.76E-05 |
| EYA3 | 2.296259058 | 1.63E-06 | 2.65E-05 |
| ISOC1 | 2.297037084 | 0.00040827 | 0.003264496 |
| NABP1 | 2.301071537 | 1.44E-09 | 4.82E-08 |
| MAP3K8 | 2.304108282 | 3.15E-14 | 2.34E-12 |
| DDX21 | 2.304990991 | 1.60E-12 | 9.01E-11 |
| SLC30A6 | 2.308098842 | 7.50E-05 | 0.000757837 |
| PPA1 | 2.320429606 | 9.18E-05 | 0.000906154 |
| CD38 | 2.320674529 | 0.000347745 | 0.002833663 |
| RNF121 | 2.323984555 | 0.009060731 | 0.039109033 |
| CLSPN | 2.324265214 | 0.001123118 | 0.007395242 |
| LMLN | 2.332058849 | 0.001440794 | 0.009041941 |
| PARD6B | 2.335594838 | 0.005999443 | 0.028298332 |
| IVNS1ABP | 2.33665742 | 1.64E-12 | 9.18E-11 |
| POP1 | 2.338638693 | 0.000530423 | 0.004049787 |
| FAM102A | 2.343057365 | 0.008012632 | 0.035619364 |
| DNAJB11 | 2.344438963 | 1.26E-13 | 8.67E-12 |
| SMS | 2.348229658 | 1.01E-06 | 1.74E-05 |
| RBM14-RBM4 | 2.348491122 | 3.22E-06 | 4.80E-05 |
| C16orf52 | 2.358073504 | 9.10E-05 | 0.000900449 |
| SVIP | 2.365315018 | 0.000167296 | 0.001526826 |
| SYNC | 2.365428552 | 0.008672271 | 0.0378486 |
| NOP10 | 2.368109446 | 4.47E-08 | 1.08E-06 |
| TFB1M | 2.369475043 | 0.001196465 | 0.007748197 |
| MED11 | 2.370632286 | 1.97E-06 | 3.15E-05 |
| TXNDC15 | 2.370983419 | 1.68E-10 | 6.66E-09 |
| SLC5A3 | 2.378143044 | 5.10E-08 | 1.22E-06 |
| FAS | 2.395259784 | 0.00052627 | 0.004030628 |
| TUBB | 2.395478714 | 6.00E-20 | 9.06E-18 |

| | | | |
|---|---|---|---|
| NFKBIA | 2.397462419 | 4.97E-21 | 9.03E-19 |
| HSBP1 | 2.397535688 | 0.000182014 | 0.001636279 |
| SEH1L | 2.399976867 | 1.05E-09 | 3.56E-08 |
| RCC1 | 2.403436384 | 0.001931019 | 0.011405914 |
| ZNF681 | 2.407913579 | 0.009263078 | 0.039809688 |
| EED | 2.41244904 | 1.04E-06 | 1.80E-05 |
| MCM5 | 2.420744638 | 6.06E-07 | 1.11E-05 |
| EBNA1BP2 | 2.421700851 | 2.98E-05 | 0.000343309 |
| PLAGL1 | 2.423551159 | 0.000269032 | 0.002273447 |
| ALAS1 | 2.428241396 | 0.000306942 | 0.002543415 |
| HSPE1 | 2.432436113 | 1.20E-10 | 4.94E-09 |
| TUBE1 | 2.434633704 | 4.13E-05 | 0.00045247 |
| LACTB | 2.43804026 | 2.18E-09 | 6.95E-08 |
| FAM3C | 2.438332195 | 8.76E-06 | 0.000116868 |
| CLDND1 | 2.439253154 | 3.33E-20 | 5.26E-18 |
| HDGFRP3 | 2.447050864 | 5.67E-07 | 1.05E-05 |
| GNPNAT1 | 2.449296569 | 2.87E-05 | 0.00033328 |
| GCH1 | 2.458286928 | 2.21E-11 | 1.04E-09 |
| RHOH | 2.466285669 | 5.93E-06 | 8.30E-05 |
| BCL6 | 2.470593453 | 3.86E-06 | 5.66E-05 |
| KLHL6 | 2.474130101 | 0.009573025 | 0.040845953 |
| HAVCR2 | 2.478239734 | 7.28E-13 | 4.30E-11 |
| GPR183 | 2.483082876 | 0.00068435 | 0.004977134 |
| ZNF436 | 2.487824752 | 0.003382683 | 0.018075984 |
| TBCCD1 | 2.490400335 | 0.000641261 | 0.004702099 |
| GNB4 | 2.491182808 | 0.008773199 | 0.038102638 |
| THAP5 | 2.49395323 | 5.92E-10 | 2.12E-08 |
| SEC61B | 2.496058999 | 1.16E-20 | 1.97E-18 |
| RRS1 | 2.496619841 | 0.000850754 | 0.00592809 |
| SLC16A3 | 2.497359212 | 1.73E-08 | 4.61E-07 |
| FAIM | 2.501810094 | 0.001682551 | 0.01018982 |
| ARL8B | 2.510637304 | 1.41E-08 | 3.80E-07 |
| STAT5A | 2.517399246 | 1.08E-07 | 2.44E-06 |
| ELL2 | 2.519644539 | 1.79E-06 | 2.90E-05 |
| C1orf216 | 2.521284123 | 0.000948954 | 0.006447497 |
| ACTL6A | 2.5224658 | 0.000906834 | 0.006230356 |
| EOGT | 2.523556332 | 2.73E-06 | 4.17E-05 |
| DCUN1D3 | 2.524829029 | 0.001087115 | 0.007211416 |
| IMPDH2 | 2.528906982 | 0.00283264 | 0.015701145 |
| LIMK2 | 2.535712753 | 2.26E-06 | 3.55E-05 |
| C1QBP | 2.536514318 | 0.000861259 | 0.005980049 |
| AFAP1 | 2.538009577 | 0.000553125 | 0.004201824 |
| STT3A | 2.543008562 | 5.73E-06 | 8.07E-05 |
| WDR3 | 2.544843961 | 5.65E-06 | 7.97E-05 |
| TBC1D24 | 2.550262295 | 8.61E-07 | 1.51E-05 |
| RAB3GAP1 | 2.559445315 | 0.007926256 | 0.035330877 |
| PMAIP1 | 2.559761828 | 2.20E-09 | 6.99E-08 |
| DHX33 | 2.563856389 | 1.08E-05 | 0.000140042 |
| TMEM189 | 2.567609987 | 9.72E-06 | 0.000128668 |
| NFKBIZ | 2.581294944 | 1.83E-12 | 1.02E-10 |

| | | | |
|---|---|---|---|
| ZNF200 | 2.588100867 | 0.002451766 | 0.013880135 |
| ARSB | 2.58908419 | 0.00042287 | 0.003367518 |
| PPAT | 2.600973454 | 0.005208136 | 0.025320624 |
| C1GALT1C1 | 2.605932685 | 0.000132331 | 0.00124477 |
| AP1S3 | 2.607427506 | 0.004791616 | 0.023682896 |
| IRF5 | 2.615862526 | 6.38E-06 | 8.82E-05 |
| DNAJA4 | 2.623737676 | 5.95E-05 | 0.000621228 |
| ZNF45 | 2.62787476 | 0.009263635 | 0.039809688 |
| SRPRB | 2.628307983 | 0.000213865 | 0.001866753 |
| TUBB4B | 2.633652573 | 2.97E-15 | 2.68E-13 |
| GRIP1 | 2.635935483 | 0.000591699 | 0.004421294 |
| MRPL14 | 2.64044831 | 1.21E-06 | 2.02E-05 |
| AGPAT5 | 2.641892608 | 0.005812686 | 0.027646533 |
| GOT1 | 2.644457316 | 9.98E-05 | 0.000975372 |
| HAT1 | 2.645823687 | 1.75E-07 | 3.71E-06 |
| SNX9 | 2.654471454 | 0.000142098 | 0.001327738 |
| DUSP10 | 2.657718614 | 5.19E-08 | 1.24E-06 |
| C14orf1 | 2.661563065 | 2.14E-06 | 3.37E-05 |
| TVP23B | 2.662068291 | 2.46E-07 | 5.03E-06 |
| IRGQ | 2.66408201 | 1.53E-05 | 0.000190696 |
| ZNF780B | 2.664474978 | 0.000647208 | 0.004742162 |
| STIP1 | 2.66504947 | 0.000100873 | 0.000984745 |
| STK32C | 2.665431686 | 0.001566958 | 0.009618053 |
| PDCD5 | 2.672987785 | 0.00016725 | 0.001526826 |
| PIM3 | 2.675874391 | 7.85E-06 | 0.000105728 |
| APOO | 2.679842826 | 0.00763659 | 0.034289513 |
| AK6 | 2.680452086 | 1.65E-07 | 3.51E-06 |
| EPB41L2 | 2.680879595 | 0.001369584 | 0.00864671 |
| SERTAD1 | 2.680936778 | 0.001863487 | 0.011070598 |
| FAM129A | 2.681632519 | 8.73E-12 | 4.37E-10 |
| SLC25A33 | 2.683978913 | 0.000203927 | 0.001800532 |
| B4GALT5 | 2.684199184 | 6.88E-09 | 2.00E-07 |
| APIP | 2.685711428 | 0.00116142 | 0.007562554 |
| AGO2 | 2.691280251 | 6.27E-05 | 0.000647185 |
| ADO | 2.699804014 | 1.54E-13 | 1.04E-11 |
| MCM2 | 2.701508569 | 1.07E-05 | 0.000139249 |
| EYS | 2.704898707 | 0.004200086 | 0.02142276 |
| VMP1 | 2.713176091 | 3.43E-17 | 3.78E-15 |
| ERCC4 | 2.71549148 | 5.18E-05 | 0.000552442 |
| PHF23 | 2.721695616 | 0.00472841 | 0.023453202 |
| BCOR | 2.725527882 | 4.04E-09 | 1.21E-07 |
| EXOSC3 | 2.726720223 | 0.00010706 | 0.001035859 |
| MRPS26 | 2.727177945 | 0.006688206 | 0.030812076 |
| BTG2 | 2.729658779 | 2.27E-19 | 3.32E-17 |
| SQLE | 2.733457108 | 4.25E-08 | 1.04E-06 |
| NEDD4 | 2.738388328 | 0.00845367 | 0.037125764 |
| ANK3 | 2.743115593 | 8.30E-08 | 1.91E-06 |
| MOB3C | 2.743133399 | 3.22E-05 | 0.000365389 |
| MLLT4 | 2.74434281 | 7.35E-05 | 0.000744753 |
| CEBPB | 2.746900907 | 1.25E-11 | 6.15E-10 |

| | | | |
|---|---|---|---|
| HMGCR | 2.76321855 | 6.42E-15 | 5.29E-13 |
| SRGAP2 | 2.766351792 | 0.008888473 | 0.038535046 |
| ZDHHC16 | 2.773322565 | 0.001604641 | 0.009814922 |
| DPAGT1 | 2.774806783 | 0.004061581 | 0.020895738 |
| ACP2 | 2.774910876 | 0.008659779 | 0.037827736 |
| SEC24D | 2.78090699 | 9.21E-07 | 1.61E-05 |
| GZMB | 2.787448954 | 2.68E-18 | 3.51E-16 |
| IL18R1 | 2.791721916 | 1.30E-12 | 7.44E-11 |
| NAMPT | 2.792591184 | 3.44E-07 | 6.75E-06 |
| RNF24 | 2.794992707 | 0.007959974 | 0.035433441 |
| SKIL | 2.799069669 | 1.10E-23 | 2.71E-21 |
| GTF2H3 | 2.802774823 | 4.20E-06 | 6.09E-05 |
| DUSP14 | 2.803176458 | 0.001893944 | 0.011227483 |
| NT5DC3 | 2.812007262 | 0.007851336 | 0.035109143 |
| SNHG12 | 2.816066282 | 7.92E-11 | 3.44E-09 |
| FICD | 2.817988821 | 5.50E-05 | 0.000581268 |
| DNAH5 | 2.821391407 | 0.005603139 | 0.026802728 |
| MUC6 | 2.831989878 | 0.003514898 | 0.018570096 |
| GADD45B | 2.83380864 | 1.91E-10 | 7.53E-09 |
| PIM2 | 2.834917743 | 3.56E-13 | 2.22E-11 |
| SLC7A1 | 2.837058905 | 6.54E-06 | 8.96E-05 |
| RNF170 | 2.84579197 | 0.000569561 | 0.00429513 |
| BCL2L2 | 2.854432408 | 0.001045823 | 0.006984731 |
| UBTD2 | 2.867441361 | 0.00274933 | 0.015273884 |
| NANOS1 | 2.867885572 | 0.001623552 | 0.009918225 |
| ACSL1 | 2.875803573 | 0.000230017 | 0.001981296 |
| SCO1 | 2.878297688 | 4.56E-05 | 0.000493699 |
| ENKUR | 2.884870717 | 0.006667959 | 0.030742175 |
| GPN3 | 2.892584516 | 0.000232551 | 0.001997866 |
| CCDC58 | 2.904338419 | 0.000467024 | 0.003656803 |
| SGMS1 | 2.909348638 | 1.17E-07 | 2.61E-06 |
| TEX30 | 2.913376707 | 1.99E-06 | 3.17E-05 |
| CDK2AP1 | 2.914893939 | 9.52E-08 | 2.17E-06 |
| TP53I11 | 2.914992427 | 0.001049546 | 0.007000066 |
| PKM | 2.915831788 | 7.77E-22 | 1.52E-19 |
| MRPL35 | 2.916746222 | 0.001508181 | 0.009334674 |
| SOD2 | 2.916809001 | 2.35E-11 | 1.10E-09 |
| MT2A | 2.923409466 | 3.01E-07 | 5.97E-06 |
| TXNDC5 | 2.940307844 | 1.80E-10 | 7.12E-09 |
| PSMA3 | 2.942421913 | 3.75E-15 | 3.31E-13 |
| GINS4 | 2.944546724 | 0.008972733 | 0.038842431 |
| APBB2 | 2.94549868 | 0.007343869 | 0.033279767 |
| VIM | 2.946209983 | 1.04E-17 | 1.23E-15 |
| DNAH14 | 2.947905294 | 0.009130975 | 0.03936028 |
| ALG2 | 2.94830645 | 3.37E-08 | 8.45E-07 |
| TNFAIP2 | 2.949509061 | 0.004440561 | 0.022341713 |
| SRM | 2.949629651 | 3.75E-09 | 1.13E-07 |
| SLC35F2 | 2.954598727 | 3.10E-05 | 0.000354702 |
| CPOX | 2.955742468 | 0.004655373 | 0.023220062 |
| KLHL2 | 2.955750849 | 3.41E-07 | 6.70E-06 |

| | | | |
|---|---|---|---|
| RNF19B | 2.956293402 | 6.81E-05 | 0.00069662 |
| LOC101928381 | 2.957764542 | 0.007273738 | 0.033022972 |
| CCDC86 | 2.958240466 | 4.31E-05 | 0.000468398 |
| TNFAIP8 | 2.961636046 | 4.96E-18 | 5.93E-16 |
| MANF | 2.967107674 | 2.39E-19 | 3.45E-17 |
| PGAP1 | 2.975893359 | 2.18E-05 | 0.000260347 |
| POLR3K | 2.977390613 | 0.000706759 | 0.005113845 |
| E2F6 | 2.978770917 | 0.000231271 | 0.001990347 |
| RASGEF1B | 2.994850062 | 1.05E-10 | 4.42E-09 |
| GABRA4 | 3.004073025 | 0.004816301 | 0.023792915 |
| RRAS2 | 3.014661167 | 1.90E-12 | 1.05E-10 |
| MCUR1 | 3.014874544 | 0.000392565 | 0.00314919 |
| ARHGAP31 | 3.018703078 | 4.15E-07 | 7.92E-06 |
| STK3 | 3.021108196 | 9.12E-05 | 0.000901049 |
| NME1 | 3.021918055 | 5.62E-06 | 7.95E-05 |
| SETMAR | 3.031010281 | 0.004892009 | 0.024045841 |
| MEFV | 3.03110509 | 0.005681061 | 0.027122575 |
| TCEB1 | 3.031878536 | 1.95E-17 | 2.26E-15 |
| TNFSF10 | 3.038341447 | 1.83E-12 | 1.02E-10 |
| IRAK3 | 3.042029629 | 0.001408785 | 0.008862983 |
| NOP16 | 3.047685547 | 0.000325655 | 0.002666947 |
| SLC43A3 | 3.049414185 | 0.000431684 | 0.003422297 |
| MIR22HG | 3.049437441 | 9.05E-11 | 3.84E-09 |
| SAMD4A | 3.050115243 | 0.008729817 | 0.038015194 |
| FKBP4 | 3.051661546 | 1.74E-09 | 5.66E-08 |
| FDPSP2 | 3.069821872 | 8.67E-05 | 0.000864012 |
| CNTLN | 3.070467082 | 0.001356643 | 0.008592655 |
| TARS | 3.076869474 | 7.55E-23 | 1.72E-20 |
| LOC284454 | 3.083323398 | 0.00039618 | 0.003173001 |
| GTPBP8 | 3.085409946 | 1.18E-06 | 1.99E-05 |
| FRMD4B | 3.089149873 | 6.09E-10 | 2.17E-08 |
| PSMB5 | 3.095376451 | 1.42E-05 | 0.000179018 |
| CSMD1 | 3.103598318 | 0.006943293 | 0.03175788 |
| POLR2K | 3.121368856 | 4.14E-14 | 2.99E-12 |
| HIVEP2 | 3.124734806 | 2.57E-09 | 8.09E-08 |
| TNFRSF25 | 3.140699712 | 4.15E-05 | 0.00045247 |
| C21orf91 | 3.145859479 | 2.66E-15 | 2.41E-13 |
| MRPL17 | 3.148936623 | 9.69E-05 | 0.000952116 |
| BHLHE40 | 3.155262669 | 2.10E-20 | 3.43E-18 |
| SLFN11 | 3.164662297 | 6.96E-13 | 4.17E-11 |
| IRF8 | 3.171740416 | 1.45E-12 | 8.28E-11 |
| MOB1A | 3.173822655 | 0.000624971 | 0.004610221 |
| ACVR1B | 3.175697889 | 7.21E-05 | 0.000731832 |
| APOLD1 | 3.179826462 | 0.009781192 | 0.041470732 |
| LOC101926955 | 3.184831698 | 0.006360514 | 0.029560873 |
| APP | 3.190196846 | 0.009660244 | 0.04106441 |
| BCL2L1 | 3.19524912 | 5.35E-13 | 3.26E-11 |
| LOC101928514 | 3.195338742 | 0.004582973 | 0.022952297 |
| NR4A3 | 3.19664542 | 0.003394974 | 0.018092388 |
| PPIF | 3.203932616 | 0.000154263 | 0.001426704 |

| | | | |
|---|---|---|---|
| IL12RB2 | 3.207220762 | 0.002257435 | 0.012929183 |
| FEN1 | 3.21261963 | 3.20E-13 | 2.02E-11 |
| UMPS | 3.213418877 | 2.78E-07 | 5.56E-06 |
| CREB5 | 3.221524435 | 0.000105686 | 0.001024593 |
| CDKL1 | 3.227965805 | 0.002676085 | 0.014951634 |
| CKS2 | 3.22877733 | 0.000874798 | 0.006052643 |
| ABCA13 | 3.229385906 | 3.97E-05 | 0.000437208 |
| DENND5A | 3.22947131 | 1.11E-06 | 1.89E-05 |
| CSRNP2 | 3.230206051 | 8.02E-07 | 1.42E-05 |
| TUBA1B | 3.232146631 | 1.39E-24 | 3.90E-22 |
| TTC5 | 3.241544367 | 0.00596518 | 0.028177363 |
| RGS1 | 3.246952205 | 1.05E-06 | 1.81E-05 |
| RPL23AP53 | 3.257796104 | 2.93E-05 | 0.000338444 |
| KDM6B | 3.25824295 | 2.19E-09 | 6.98E-08 |
| SUSD5 | 3.271786865 | 0.003202151 | 0.017242755 |
| ZNF542P | 3.277552445 | 6.58E-08 | 1.54E-06 |
| PPP3R1 | 3.280789826 | 5.48E-08 | 1.30E-06 |
| TNFSF8 | 3.281145248 | 5.22E-05 | 0.00055498 |
| UTP11L | 3.285000767 | 3.71E-05 | 0.000412381 |
| POM121L10P | 3.297918275 | 0.004211958 | 0.021466762 |
| XCL1 | 3.305687714 | 2.03E-07 | 4.25E-06 |
| TOP1P1 | 3.310787301 | 0.001519584 | 0.009380159 |
| RBPJ | 3.311431437 | 8.95E-33 | 3.82E-30 |
| MDFIC | 3.31237532 | 9.98E-15 | 8.16E-13 |
| APITD1 | 3.31291817 | 0.00057935 | 0.004343568 |
| FEZ1 | 3.314188847 | 1.92E-22 | 4.10E-20 |
| LIMCH1 | 3.339826765 | 0.000117956 | 0.001130143 |
| WDR17 | 3.341614829 | 0.007369362 | 0.033302998 |
| ZNF71 | 3.345073342 | 0.000292565 | 0.002440775 |
| SLC39A14 | 3.348356925 | 1.36E-05 | 0.000173249 |
| RALA | 3.358155236 | 2.24E-15 | 2.05E-13 |
| LRP12 | 3.358681977 | 9.08E-05 | 0.000900191 |
| SLC38A6 | 3.364609918 | 0.004887133 | 0.024045841 |
| LINC01296 | 3.376662001 | 0.000262907 | 0.002233226 |
| LCMT2 | 3.378226548 | 0.002641022 | 0.014823267 |
| ITIH5 | 3.387314197 | 0.005369065 | 0.025974309 |
| RELB | 3.387509623 | 3.97E-05 | 0.000437415 |
| NDUFAF4P1 | 3.395584076 | 0.000620238 | 0.004585651 |
| SHMT2 | 3.39911586 | 4.73E-07 | 8.93E-06 |
| MAGI2-AS3 | 3.40550249 | 0.003093365 | 0.016767407 |
| ADCY3 | 3.411750906 | 0.009276056 | 0.039845617 |
| SOCS3 | 3.411903061 | 2.45E-09 | 7.73E-08 |
| KCTD17 | 3.414364497 | 0.000854227 | 0.005943844 |
| RUNX1T1 | 3.416664367 | 9.52E-05 | 0.000937002 |
| SV2C | 3.416767742 | 0.00352833 | 0.018621004 |
| CCBE1 | 3.41752318 | 0.004676515 | 0.023255211 |
| MAK | 3.419370695 | 0.001459984 | 0.009101105 |
| C10orf2 | 3.425048715 | 0.002946813 | 0.016214909 |
| CD109 | 3.432625045 | 0.000102179 | 0.000995509 |
| NAB2 | 3.435916563 | 0.000328506 | 0.002688049 |

| | | | |
|---|---|---|---|
| FAM161A | 3.441206239 | 6.21E-05 | 0.000642796 |
| TUBA1C | 3.442276934 | 1.23E-10 | 5.03E-09 |
| SCN7A | 3.458509623 | 0.006511068 | 0.030132117 |
| SAP30 | 3.471395241 | 0.000292864 | 0.002441195 |
| BCAT1 | 3.477937944 | 0.000808838 | 0.005704897 |
| NALCN | 3.481887063 | 0.004148392 | 0.021275416 |
| LOC101927257 | 3.504450795 | 0.009209345 | 0.039645847 |
| NDST3 | 3.505046121 | 0.001638238 | 0.009979811 |
| C9orf40 | 3.511250014 | 0.000119245 | 0.001135842 |
| RNF144A | 3.519088352 | 3.69E-10 | 1.39E-08 |
| RCAN3 | 3.525381817 | 0.001039935 | 0.006950139 |
| loc920 | 3.535161137 | 2.59E-13 | 1.65E-11 |
| PELO | 3.541304558 | 0.000360041 | 0.002921726 |
| ZDHHC9 | 3.560923496 | 0.007585571 | 0.034107259 |
| PFKFB3 | 3.574346103 | 3.58E-15 | 3.19E-13 |
| SAMD5 | 3.577464295 | 0.003535225 | 0.018647359 |
| AGFG2 | 3.585149428 | 0.005399738 | 0.026071276 |
| STXBP4 | 3.591416867 | 0.000388347 | 0.003123013 |
| CCND2 | 3.602769991 | 1.44E-38 | 1.01E-35 |
| MTHFD2 | 3.609350369 | 9.49E-19 | 1.29E-16 |
| STX11 | 3.610057206 | 7.61E-08 | 1.76E-06 |
| LOC440300 | 3.611031876 | 5.68E-07 | 1.05E-05 |
| FAM89A | 3.642458635 | 0.00028123 | 0.002360265 |
| HAPLN1 | 3.6472646 | 0.001690884 | 0.010227661 |
| GNG4 | 3.648090793 | 0.003422882 | 0.018221323 |
| ZNF619 | 3.649931405 | 0.004365178 | 0.022052915 |
| PRDX3 | 3.656935618 | 1.79E-09 | 5.79E-08 |
| PAICS | 3.661506847 | 9.89E-13 | 5.74E-11 |
| MRTO4 | 3.664188192 | 1.78E-05 | 0.000216328 |
| PSMC3IP | 3.671965446 | 1.29E-07 | 2.82E-06 |
| CHEK1 | 3.676035604 | 2.91E-08 | 7.41E-07 |
| SLC35G5 | 3.692666506 | 0.004299031 | 0.021763566 |
| BATF | 3.692737402 | 1.15E-06 | 1.94E-05 |
| IL4R | 3.694319068 | 2.31E-16 | 2.36E-14 |
| CD274 | 3.710996375 | 1.40E-05 | 0.000177376 |
| SLC16A1 | 3.714788589 | 8.34E-05 | 0.000834964 |
| LOC101928372 | 3.731628092 | 0.006692542 | 0.030812076 |
| FSD2 | 3.741479353 | 0.0028385 | 0.015724746 |
| FABP5 | 3.742121864 | 4.88E-08 | 1.18E-06 |
| ZNF267 | 3.747121447 | 2.54E-18 | 3.36E-16 |
| INPP5E | 3.76177693 | 1.08E-10 | 4.51E-09 |
| OLR1 | 3.765052322 | 0.005193849 | 0.025301315 |
| TNFRSF4 | 3.774309148 | 1.25E-07 | 2.73E-06 |
| ETS2 | 3.786651624 | 5.89E-05 | 0.000614912 |
| DUSP5 | 3.789195684 | 3.50E-08 | 8.70E-07 |
| IL1R1 | 3.816031699 | 0.005876755 | 0.027867008 |
| FASLG | 3.81621742 | 3.89E-21 | 7.34E-19 |
| AHR | 3.816602939 | 5.38E-22 | 1.12E-19 |
| DLEU2 | 3.823693099 | 0.000456801 | 0.003588208 |
| TANK | 3.827536972 | 1.88E-21 | 3.62E-19 |

| | | | |
|---|---|---|---|
| GPR34 | 3.828273806 | 0.00368507 | 0.019302841 |
| GPR82 | 3.832221839 | 5.14E-06 | 7.30E-05 |
| MEIS2 | 3.835826606 | 0.003824194 | 0.019913602 |
| ZNF663P | 3.83788659 | 0.006828037 | 0.031347622 |
| DIAPH3 | 3.848734325 | 0.000721839 | 0.005207325 |
| CDK6 | 3.858973755 | 3.28E-39 | 2.68E-36 |
| LTBP1 | 3.872490019 | 0.000911467 | 0.006257803 |
| HIF1A-AS2 | 3.886437693 | 0.003162236 | 0.017077306 |
| HSPA1A | 3.906203652 | 2.13E-37 | 1.30E-34 |
| MEG3 | 3.911634728 | 0.001502159 | 0.009315856 |
| LRRTM2 | 3.916883377 | 1.14E-05 | 0.000147878 |
| EPB41L4B | 3.918797259 | 0.001183919 | 0.007677086 |
| NDUFAF1 | 3.920154934 | 2.46E-06 | 3.83E-05 |
| LOC102724612 | 3.924086598 | 0.001213056 | 0.00784528 |
| SRXN1 | 3.927919624 | 3.76E-17 | 4.10E-15 |
| TAX1BP3 | 3.930926649 | 4.44E-18 | 5.44E-16 |
| NDFIP2 | 3.930958823 | 4.03E-16 | 4.04E-14 |
| BAIAP2 | 3.935815836 | 0.000814759 | 0.005732076 |
| LOC101927580 | 3.938990359 | 0.000321838 | 0.002644515 |
| HSPA6 | 3.942686916 | 7.41E-06 | 0.000100635 |
| GPT2 | 3.948837869 | 1.98E-05 | 0.000238357 |
| PIM1 | 3.951288259 | 6.98E-24 | 1.76E-21 |
| TMEM254-AS1 | 3.959852662 | 0.00093212 | 0.006363973 |
| KSR1 | 3.962978492 | 0.002069027 | 0.012080059 |
| LACC1 | 3.96347542 | 2.57E-06 | 3.98E-05 |
| GRIA3 | 3.987280331 | 0.000807721 | 0.005701117 |
| PPIL1 | 3.995580796 | 0.000669656 | 0.00489203 |
| C9orf84 | 4.010981952 | 0.000182136 | 0.001636279 |
| NFKB2 | 4.02199759 | 2.19E-10 | 8.58E-09 |
| TRHDE | 4.02707228 | 0.000514842 | 0.003971004 |
| LOC100506472 | 4.029601459 | 7.46E-05 | 0.000754772 |
| CDH18 | 4.037477333 | 0.000536671 | 0.004094311 |
| AK4 | 4.045116119 | 0.000109186 | 0.001052282 |
| DNPH1 | 4.062348109 | 0.000220514 | 0.001914567 |
| NPIPB3 | 4.081204677 | 1.47E-08 | 3.96E-07 |
| RAB39B | 4.093128662 | 3.80E-05 | 0.000421338 |
| FAM227A | 4.098651695 | 0.001765492 | 0.010607007 |
| BCL2A1 | 4.106333118 | 1.78E-07 | 3.75E-06 |
| EFR3B | 4.113212368 | 0.000953947 | 0.006472116 |
| SNX24 | 4.142522906 | 0.001108282 | 0.007317195 |
| TNFRSF18 | 4.151350131 | 6.32E-22 | 1.29E-19 |
| NFKB1 | 4.157836668 | 2.89E-29 | 9.76E-27 |
| ZSWIM4 | 4.164615137 | 0.000395513 | 0.003170241 |
| MAFG | 4.165690544 | 3.06E-11 | 1.41E-09 |
| TRPV1 | 4.169379183 | 0.00091299 | 0.006263032 |
| EBF3 | 4.177688273 | 0.001139093 | 0.007475343 |
| LINC01215 | 4.195567784 | 1.56E-05 | 0.000193733 |
| CYTIP | 4.202037903 | 1.44E-58 | 3.54E-55 |
| FAM83B | 4.209416655 | 0.00039105 | 0.003139602 |
| HEY1 | 4.241006025 | 0.002909031 | 0.016061061 |

| | | | |
|---|---|---|---|
| SLC41A2 | 4.248927901 | 3.98E-07 | 7.63E-06 |
| TGFB2 | 4.253485181 | 0.000504054 | 0.003896986 |
| GFRAL | 4.25736198 | 0.005437952 | 0.026204196 |
| MBL2 | 4.265991484 | 0.005488386 | 0.026382436 |
| PDC | 4.270552196 | 0.004928243 | 0.024199694 |
| SGCB | 4.271923722 | 7.06E-05 | 0.000718962 |
| FLJ35934 | 4.272889407 | 2.56E-05 | 0.000300764 |
| TXN | 4.275415797 | 5.04E-24 | 1.30E-21 |
| SMIM17 | 4.286078462 | 0.00473459 | 0.023471985 |
| SRSF12 | 4.29460714 | 0.000272773 | 0.002299124 |
| TWSG1 | 4.316182908 | 1.12E-07 | 2.51E-06 |
| DYX1C1 | 4.31764207 | 0.001029086 | 0.006896424 |
| HLF | 4.353698037 | 0.000969339 | 0.006554226 |
| PDGFA | 4.357527432 | 0.000124502 | 0.001177904 |
| DLG2 | 4.378812411 | 0.000756844 | 0.00540049 |
| PPAN | 4.398048472 | 0.00531519 | 0.025751766 |
| DPP4 | 4.399907057 | 0.000495187 | 0.003840534 |
| SLC7A11 | 4.412101351 | 3.88E-05 | 0.00042939 |
| LNP1 | 4.421103236 | 0.001360093 | 0.008603397 |
| LINC00824 | 4.427171389 | 0.007824084 | 0.035019199 |
| LOC101929680 | 4.441704695 | 0.000541214 | 0.004123567 |
| TRIP10 | 4.460879273 | 0.000118646 | 0.001134535 |
| P2RY13 | 4.47069003 | 0.000311645 | 0.002573692 |
| SLC35F1 | 4.503359153 | 4.14E-05 | 0.00045247 |
| DCAF4L1 | 4.506175054 | 8.56E-05 | 0.000855439 |
| LOC101930452 | 4.511024412 | 3.63E-05 | 0.000404118 |
| SLC7A5 | 4.579907217 | 2.00E-15 | 1.85E-13 |
| SNCA | 4.607236415 | 0.000189376 | 0.001690597 |
| PGR | 4.620066504 | 0.001552987 | 0.009558568 |
| SCARNA10 | 4.635478778 | 0.002123668 | 0.01229947 |
| SEMA7A | 4.651331027 | 0.000978571 | 0.006602998 |
| BUB1B | 4.651708539 | 0.000192392 | 0.001714402 |
| ACKR4 | 4.702485478 | 0.001687656 | 0.010214432 |
| FAM138C | 4.703836843 | 0.002064284 | 0.012080059 |
| CEP55 | 4.712029697 | 0.006018455 | 0.028360743 |
| TRAF1 | 4.715195007 | 4.88E-28 | 1.60E-25 |
| CDC6 | 4.716219804 | 5.50E-05 | 0.000581268 |
| HOTTIP | 4.72814827 | 0.000670854 | 0.004897135 |
| IL15RA | 4.731599873 | 3.58E-09 | 1.09E-07 |
| KLHL31 | 4.743175426 | 0.000695679 | 0.005048306 |
| SCN8A | 4.743657712 | 3.08E-06 | 4.62E-05 |
| LMNA | 4.770170309 | 5.30E-08 | 1.26E-06 |
| IL6R | 4.789972104 | 2.89E-08 | 7.39E-07 |
| CCL4 | 4.792628445 | 6.13E-83 | 3.01E-79 |
| CCL3L1 | 4.799711013 | 7.67E-07 | 1.37E-05 |
| GRB10 | 4.800290358 | 0.003007509 | 0.016453594 |
| MIR17HG | 4.801109864 | 4.11E-08 | 1.01E-06 |
| FAM157C | 4.802353497 | 0.000566623 | 0.004279553 |
| THAP6 | 4.870654122 | 2.66E-06 | 4.08E-05 |
| DNAJC6 | 4.880392526 | 6.49E-05 | 0.000667781 |

| | | | |
|---|---|---|---|
| WARS | 4.886951761 | 3.65E-20 | 5.60E-18 |
| GPR132 | 4.887808486 | 2.37E-06 | 3.71E-05 |
| C17orf96 | 4.8945408 | 1.66E-05 | 0.000204726 |
| CTH | 4.917609459 | 0.000224424 | 0.001943356 |
| SLC4A5 | 4.931679777 | 2.36E-17 | 2.69E-15 |
| FLJ31356 | 4.979580168 | 0.000205264 | 0.001807758 |
| HAPLN3 | 5.03656518 | 2.80E-06 | 4.28E-05 |
| OR4F29 | 5.03954991 | 0.001776629 | 0.010654347 |
| FAM129B | 5.040050447 | 1.13E-06 | 1.92E-05 |
| WDR11-AS1 | 5.041553662 | 0.000204695 | 0.001804366 |
| FAM138A | 5.050173904 | 0.000111204 | 0.001068582 |
| LOC654841 | 5.064377426 | 0.000163946 | 0.001501843 |
| ANKS4B | 5.070807469 | 0.000149927 | 0.001395572 |
| IL1RN | 5.076633485 | 5.99E-05 | 0.000624572 |
| SOCS1 | 5.090129285 | 1.11E-14 | 8.93E-13 |
| SDC4 | 5.093859832 | 7.36E-14 | 5.20E-12 |
| MTHFD1L | 5.093993563 | 2.85E-10 | 1.10E-08 |
| CMAHP | 5.121773518 | 2.40E-36 | 1.27E-33 |
| LGALS3 | 5.135985621 | 4.83E-12 | 2.51E-10 |
| ZSWIM3 | 5.137893106 | 0.001142937 | 0.007490553 |
| KLHL23 | 5.13963347 | 3.31E-06 | 4.93E-05 |
| BUD13 | 5.151841127 | 1.05E-05 | 0.000137374 |
| BIRC3 | 5.171821537 | 2.77E-51 | 3.40E-48 |
| AXIN2 | 5.173455941 | 0.001558178 | 0.009578501 |
| TRPA1 | 5.191354935 | 7.41E-07 | 1.33E-05 |
| RGCC | 5.210299436 | 1.84E-07 | 3.87E-06 |
| ICAM1 | 5.212657444 | 1.05E-08 | 2.91E-07 |
| SAMSN1 | 5.231628252 | 3.48E-24 | 9.23E-22 |
| DCTN1-AS1 | 5.284013425 | 4.14E-06 | 6.00E-05 |
| C15orf48 | 5.284028777 | 1.43E-37 | 9.33E-35 |
| CCL4L2 | 5.290932036 | 3.65E-20 | 5.60E-18 |
| IRF4 | 5.292986668 | 2.43E-13 | 1.59E-11 |
| HK2 | 5.303107962 | 4.78E-07 | 9.00E-06 |
| ZNRF1 | 5.321639199 | 3.73E-06 | 5.52E-05 |
| IDO1 | 5.346687186 | 1.41E-06 | 2.32E-05 |
| KIAA0895 | 5.357245493 | 0.000429247 | 0.003409995 |
| TNF | 5.360536134 | 0.000320779 | 0.002640233 |
| EFCAB5 | 5.390473579 | 6.45E-06 | 8.89E-05 |
| TMEM217 | 5.420135799 | 5.09E-07 | 9.52E-06 |
| IER3 | 5.42718031 | 1.42E-22 | 3.18E-20 |
| DLGAP1 | 5.438866885 | 1.14E-05 | 0.000147878 |
| PSAT1 | 5.450247928 | 3.64E-11 | 1.65E-09 |
| PYCR1 | 5.518835763 | 4.61E-05 | 0.000498019 |
| B3GNT5 | 5.530054156 | 1.73E-11 | 8.30E-10 |
| CDKN1A | 5.535519121 | 2.14E-12 | 1.17E-10 |
| CXCL8 | 5.544326527 | 0.001670368 | 0.010128544 |
| TNFRSF12A | 5.556668826 | 0.000443178 | 0.003498005 |
| GADD45G | 5.566140836 | 1.15E-07 | 2.57E-06 |
| HOXC8 | 5.595755577 | 0.000190077 | 0.001695312 |
| CFHR5 | 5.625751235 | 0.000119043 | 0.001135705 |

| | | | |
|---|---|---|---|
| ERC2 | 5.681104367 | 1.74E-07 | 3.70E-06 |
| PHLDA1 | 5.687460566 | 1.66E-30 | 6.02E-28 |
| ABTB2 | 5.700115529 | 8.74E-09 | 2.46E-07 |
| SLC35G4 | 5.773817291 | 5.26E-05 | 0.000558331 |
| MTTP | 5.783482321 | 1.56E-05 | 0.00019397 |
| LOC100507140 | 5.789769397 | 3.30E-08 | 8.29E-07 |
| ADORA2A | 5.791520043 | 4.95E-05 | 0.000530587 |
| CCL3 | 5.834593221 | 1.08E-73 | 3.52E-70 |
| CCDC173 | 5.846058453 | 1.03E-06 | 1.78E-05 |
| CLEC4D | 5.876073582 | 3.70E-05 | 0.000412088 |
| CD83 | 5.913096639 | 1.13E-09 | 3.79E-08 |
| LPP-AS2 | 5.978237521 | 4.63E-10 | 1.70E-08 |
| VNN1 | 5.997315617 | 1.16E-07 | 2.59E-06 |
| LOC283299 | 5.999439105 | 1.57E-06 | 2.57E-05 |
| CYP1B1 | 6.012953939 | 7.46E-09 | 2.14E-07 |
| NETO2 | 6.059037175 | 3.45E-09 | 1.06E-07 |
| LINC00515 | 6.081957338 | 4.63E-06 | 6.66E-05 |
| DLGAP1-AS4 | 6.11835158 | 9.55E-06 | 0.000126794 |
| IRS1 | 6.159494816 | 5.88E-06 | 8.24E-05 |
| C4orf32 | 6.159551368 | 1.07E-16 | 1.13E-14 |
| MYC | 6.189631926 | 9.44E-11 | 3.99E-09 |
| ZNF547 | 6.211679329 | 1.65E-06 | 2.67E-05 |
| LOC100134368 | 6.22955771 | 2.47E-06 | 3.83E-05 |
| GPR171 | 6.240548729 | 1.08E-29 | 3.78E-27 |
| CISH | 6.258765221 | 4.35E-18 | 5.41E-16 |
| SPHK1 | 6.267252637 | 3.98E-06 | 5.80E-05 |
| NR5A2 | 6.271187926 | 2.88E-06 | 4.38E-05 |
| RBKS | 6.273060412 | 3.85E-06 | 5.65E-05 |
| KCNA7 | 6.367450353 | 4.40E-07 | 8.35E-06 |
| RRAD | 6.416461414 | 6.18E-07 | 1.13E-05 |
| HSPA1B | 6.439570457 | 1.96E-17 | 2.26E-15 |
| ADM2 | 6.469898273 | 1.68E-06 | 2.73E-05 |
| ARHGEF35 | 6.514723764 | 1.16E-07 | 2.60E-06 |
| PMEPA1 | 6.54731179 | 7.90E-11 | 3.44E-09 |
| AGPAT9 | 6.601897796 | 6.50E-12 | 3.30E-10 |
| CCR7 | 6.611597535 | 2.81E-12 | 1.52E-10 |
| C9orf153 | 6.644074719 | 2.21E-06 | 3.47E-05 |
| TICAM2 | 6.668862293 | 3.89E-07 | 7.50E-06 |
| LOC650293 | 6.713749473 | 1.10E-06 | 1.87E-05 |
| CCR1 | 6.722648919 | 7.11E-08 | 1.65E-06 |
| LOC101928401 | 6.763389974 | 1.83E-07 | 3.86E-06 |
| PERP | 6.781091893 | 1.21E-07 | 2.67E-06 |
| AGMAT | 6.892553599 | 3.92E-15 | 3.44E-13 |
| ANKRD26P3 | 6.932272957 | 1.74E-08 | 4.62E-07 |
| EBI3 | 6.977281645 | 9.79E-09 | 2.73E-07 |
| CCL1 | 7.011714792 | 5.13E-08 | 1.23E-06 |
| SLC1A5 | 7.047548441 | 6.23E-06 | 8.69E-05 |
| FCGR1B | 7.057781611 | 4.84E-10 | 1.77E-08 |
| CSF2 | 7.099209244 | 3.23E-10 | 1.23E-08 |
| MAMLD1 | 7.151928144 | 2.36E-13 | 1.55E-11 |

| | | | |
|---|---|---|---|
| CSF1 | 7.180285369 | 7.87E-09 | 2.25E-07 |
| CD82 | 7.207736636 | 4.42E-17 | 4.76E-15 |
| IL23R | 7.226885146 | 7.93E-11 | 3.44E-09 |
| KIAA0226L | 7.239484016 | 5.01E-11 | 2.25E-09 |
| GPR26 | 7.246961849 | 3.15E-17 | 3.51E-15 |
| LIF | 7.272095898 | 9.20E-09 | 2.58E-07 |
| ULBP1 | 7.346814452 | 1.26E-12 | 7.29E-11 |
| DTNA | 7.348559587 | 1.09E-10 | 4.52E-09 |
| CXCL2 | 7.381443776 | 3.47E-09 | 1.06E-07 |
| NCS1 | 7.416693625 | 4.68E-23 | 1.09E-20 |
| P2RX5 | 7.454807132 | 1.52E-09 | 5.05E-08 |
| LTA | 7.550826019 | 2.08E-07 | 4.35E-06 |
| DUSP4 | 7.589266396 | 6.53E-10 | 2.32E-08 |
| LOC653513 | 7.677836355 | 4.79E-13 | 2.96E-11 |
| loc1578 | 7.682896484 | 5.29E-41 | 5.19E-38 |
| TNFSF4 | 7.733900237 | 2.57E-24 | 7.01E-22 |
| SLC27A2 | 7.919971268 | 5.31E-15 | 4.53E-13 |
| CXCR6 | 7.963271834 | 1.78E-15 | 1.66E-13 |
| EGR2 | 7.979678989 | 4.47E-16 | 4.43E-14 |
| CCR5 | 8.260382035 | 2.12E-14 | 1.59E-12 |
| IL26 | 8.436420689 | 1.95E-14 | 1.51E-12 |
| ITGA1 | 8.464548998 | 1.26E-24 | 3.64E-22 |
| CXCL9 | 8.499072421 | 1.84E-14 | 1.43E-12 |
| CHAC1 | 8.58067294 | 3.53E-19 | 5.02E-17 |
| MFSD2A | 8.883003196 | 7.21E-21 | 1.29E-18 |
| TNFRSF9 | 9.239844914 | 1.24E-11 | 6.15E-10 |
| ADAM19 | 9.376468347 | 5.08E-48 | 5.54E-45 |
| IFNG | 10.34284363 | 4.92E-160 | 4.83E-156 |
| PLAU | 10.42002416 | 2.73E-36 | 1.29E-33 |
| IL2RA | 10.54167467 | 6.88E-31 | 2.70E-28 |
| HNRNPA1P10 | 11.05220753 | 9.25E-39 | 6.98E-36 |
| RGS16 | 12.67284376 | 3.38E-58 | 6.63E-55 |
| MIR155HG | 12.7148808 | 2.97E-56 | 4.86E-53 |
| **CD56$^{neg}$NK stimulated vs CD56$^{neg}$NK unstimulated** | | | |
| ID | log2FC | pval | padj |
| FCRL6 | -10.07054349 | 1.40E-25 | 1.10E-22 |
| TMEM71 | -8.791337968 | 3.73E-17 | 9.72E-15 |
| TKTL1 | -8.65321568 | 5.54E-20 | 2.55E-17 |
| MYOM2 | -8.164665656 | 6.17E-10 | 3.78E-08 |
| FOS | -8.15398089 | 4.94E-17 | 1.21E-14 |
| CD52 | -8.031266696 | 1.45E-15 | 2.57E-13 |
| PLEKHG3 | -7.982720312 | 7.41E-14 | 1.04E-11 |
| TRANK1 | -7.949884697 | 1.10E-15 | 2.01E-13 |
| CDADC1 | -7.930586663 | 4.30E-14 | 6.24E-12 |
| RAB37 | -7.735708626 | 8.70E-16 | 1.62E-13 |
| KLRF1 | -7.558744199 | 3.88E-27 | 3.80E-24 |
| VNN2 | -7.406143028 | 4.73E-12 | 4.81E-10 |
| HVCN1 | -7.13762302 | 5.07E-09 | 2.29E-07 |

| | | | |
|---|---|---|---|
| ACSS1 | -6.888725965 | 2.07E-11 | 1.72E-09 |
| DDX19B | -6.857537008 | 9.02E-08 | 2.78E-06 |
| FAM21A | -6.762369144 | 3.98E-11 | 3.15E-09 |
| AMY2B | -6.720588603 | 1.68E-10 | 1.18E-08 |
| PIP4K2C | -6.665783616 | 1.54E-09 | 8.18E-08 |
| CACNA2D2 | -6.641353887 | 1.43E-09 | 7.79E-08 |
| CTBP1-AS2 | -6.638546192 | 1.62E-09 | 8.53E-08 |
| SBK1 | -6.633577063 | 1.06E-18 | 3.97E-16 |
| PDLIM2 | -6.619312104 | 1.81E-09 | 9.29E-08 |
| TPST2 | -6.60900503 | 1.95E-10 | 1.35E-08 |
| SIGLEC7 | -6.515639504 | 2.12E-09 | 1.06E-07 |
| CECR1 | -6.500833083 | 6.63E-16 | 1.27E-13 |
| TMC8 | -6.43548334 | 4.93E-19 | 1.93E-16 |
| CDKN2C | -6.400970704 | 1.56E-08 | 6.16E-07 |
| RAB27B | -6.347145821 | 7.72E-13 | 8.90E-11 |
| GSTM1 | -6.346552254 | 5.55E-07 | 1.41E-05 |
| FAM213B | -6.344196927 | 1.04E-07 | 3.14E-06 |
| GGACT | -6.275531685 | 3.37E-07 | 8.94E-06 |
| TMCC3 | -6.224885959 | 4.31E-07 | 1.11E-05 |
| THRA | -6.19815948 | 1.96E-08 | 7.51E-07 |
| KLF2 | -6.176683892 | 2.86E-11 | 2.31E-09 |
| PTPLAD2 | -6.049112787 | 1.37E-08 | 5.55E-07 |
| ZBTB46 | -5.932708888 | 6.44E-07 | 1.59E-05 |
| IQCE | -5.920502007 | 8.08E-07 | 1.92E-05 |
| SIK1 | -5.877978534 | 5.63E-11 | 4.24E-09 |
| FOSL2 | -5.845816867 | 5.71E-08 | 1.84E-06 |
| FCN1 | -5.767949399 | 2.80E-05 | 0.000426807 |
| ZBED6CL | -5.760078675 | 6.43E-07 | 1.59E-05 |
| USP30 | -5.735197793 | 4.19E-07 | 1.08E-05 |
| NME8 | -5.711892819 | 3.76E-07 | 9.81E-06 |
| MAP2K6 | -5.70534829 | 1.37E-06 | 3.11E-05 |
| TFIP11 | -5.679135873 | 7.08E-07 | 1.72E-05 |
| TNRC6C-AS1 | -5.640557175 | 3.68E-06 | 7.36E-05 |
| CTNNBIP1 | -5.62534373 | 8.11E-09 | 3.55E-07 |
| DUSP28 | -5.611924029 | 1.20E-06 | 2.74E-05 |
| ZNF710 | -5.592456762 | 8.01E-06 | 0.000145945 |
| RAP1GAP2 | -5.578120536 | 1.31E-07 | 3.91E-06 |
| P2RY10 | -5.550285146 | 7.54E-06 | 0.000138281 |
| SOX13 | -5.533450709 | 2.42E-05 | 0.000377238 |
| ATXN7L1 | -5.520152089 | 1.11E-08 | 4.69E-07 |
| GPX7 | -5.495252253 | 1.96E-05 | 0.000317218 |
| FARS2 | -5.465855472 | 1.33E-05 | 0.000223845 |
| TSPAN32 | -5.405707361 | 4.58E-12 | 4.72E-10 |
| ITPKB | -5.373405198 | 6.97E-10 | 4.23E-08 |
| ZNF23 | -5.345750465 | 1.02E-05 | 0.000179227 |
| SPON2 | -5.329022538 | 8.39E-13 | 9.52E-11 |
| SLC39A11 | -5.29029123 | 8.02E-06 | 0.000145945 |
| MTERF4 | -5.287007926 | 1.29E-05 | 0.000218471 |
| LINC00565 | -5.263183112 | 1.08E-05 | 0.000189049 |
| SLC2A4RG | -5.255242722 | 3.41E-08 | 1.21E-06 |

| | | | |
|---|---|---|---|
| HOXB3 | -5.227950395 | 3.53E-05 | 0.000515119 |
| C1orf162 | -5.210064425 | 1.96E-10 | 1.35E-08 |
| CAMKMT | -5.202709259 | 0.00030512 | 0.003028381 |
| TMEM229B | -5.199630119 | 1.12E-05 | 0.000195549 |
| CD160 | -5.194881584 | 1.55E-09 | 8.20E-08 |
| LOC101927780 | -5.172205549 | 0.000189077 | 0.002058203 |
| RGS9 | -5.171422592 | 3.01E-05 | 0.000450361 |
| MSH5-SAPCD1 | -5.169268758 | 4.75E-05 | 0.000661634 |
| ALDOC | -5.145182096 | 0.000105614 | 0.001274371 |
| LOC653160 | -5.116241794 | 0.000291103 | 0.002915126 |
| HIST1H1B | -5.077829012 | 0.000500979 | 0.004572457 |
| MIR646HG | -5.038626252 | 1.82E-05 | 0.000295994 |
| LOC643733 | -5.019146751 | 8.18E-05 | 0.001036269 |
| NAGLU | -5.008991644 | 5.70E-06 | 0.000108404 |
| ATP7A | -4.999121448 | 1.22E-05 | 0.000208451 |
| LOC728392 | -4.976974557 | 6.34E-05 | 0.000848086 |
| LGALS9C | -4.965201936 | 7.88E-12 | 7.52E-10 |
| GOLGA8B | -4.947716329 | 3.11E-08 | 1.12E-06 |
| OGFRL1 | -4.944268241 | 9.76E-05 | 0.001195611 |
| LRFN1 | -4.937085254 | 0.000708483 | 0.006103552 |
| ENPP5 | -4.930192928 | 0.00016616 | 0.001848296 |
| HIST1H2AH | -4.920501475 | 7.41E-05 | 0.000954364 |
| LRRC37B | -4.885085536 | 0.000206696 | 0.002199235 |
| AGTRAP | -4.876023332 | 0.001066489 | 0.008595713 |
| C16orf54 | -4.872903928 | 2.96E-07 | 8.10E-06 |
| ADHFE1 | -4.864769922 | 2.71E-07 | 7.51E-06 |
| CYTH4 | -4.854900536 | 1.45E-05 | 0.000241739 |
| PVRIG | -4.849339727 | 1.67E-09 | 8.71E-08 |
| SCARNA5 | -4.840554545 | 0.000130473 | 0.001509209 |
| SLC9A9 | -4.825731525 | 7.05E-05 | 0.000923448 |
| TADA1 | -4.820439467 | 0.000176459 | 0.001943533 |
| LINC00852 | -4.812716784 | 0.000118587 | 0.001396477 |
| SOX12 | -4.795872645 | 7.52E-05 | 0.000965045 |
| NPEPL1 | -4.793526409 | 0.000106761 | 0.001286226 |
| KIR2DL4 | -4.787468956 | 0.000260342 | 0.002668509 |
| FAM160A2 | -4.771528475 | 1.63E-05 | 0.000268273 |
| VAV3 | -4.748298818 | 2.50E-09 | 1.22E-07 |
| ZUFSP | -4.747996085 | 7.20E-05 | 0.000936939 |
| CTSF | -4.732288876 | 0.000562993 | 0.005059641 |
| ZNF552 | -4.719644646 | 0.000312669 | 0.003087653 |
| GPRIN3 | -4.716206828 | 1.20E-09 | 6.89E-08 |
| PLAC8 | -4.704487917 | 2.49E-06 | 5.18E-05 |
| CXCR4 | -4.700173761 | 9.48E-09 | 4.06E-07 |
| SAYSD1 | -4.685500006 | 7.21E-05 | 0.000936989 |
| FGR | -4.682485427 | 4.23E-08 | 1.45E-06 |
| FOXO4 | -4.680086439 | 0.000105209 | 0.001271434 |
| TSPAN2 | -4.672628432 | 3.17E-07 | 8.52E-06 |
| AGER | -4.669804721 | 0.000298241 | 0.002975188 |
| PARP15 | -4.664546558 | 5.36E-05 | 0.000736541 |
| BMP6 | -4.657238679 | 0.000988148 | 0.008094341 |

| | | | |
|---|---|---|---|
| WDR54 | -4.656033915 | 0.000572318 | 0.005104585 |
| C19orf68 | -4.614307686 | 0.000574158 | 0.00511517 |
| KIAA1841 | -4.612962574 | 9.50E-05 | 0.001172026 |
| ERGIC1 | -4.607339792 | 5.88E-08 | 1.89E-06 |
| COX10-AS1 | -4.606503836 | 5.34E-05 | 0.000734679 |
| PDGFRB | -4.599768178 | 0.000447564 | 0.004158174 |
| PPIL3 | -4.59475878 | 2.37E-05 | 0.000371491 |
| OAS2 | -4.585839075 | 1.97E-07 | 5.66E-06 |
| IFITM3 | -4.582567601 | 1.39E-09 | 7.67E-08 |
| NAA38 | -4.574265034 | 2.48E-07 | 6.96E-06 |
| DZANK1 | -4.56566085 | 0.000111193 | 0.001324393 |
| SYTL1 | -4.552197085 | 2.08E-08 | 7.83E-07 |
| SIGIRR | -4.532286745 | 6.24E-11 | 4.65E-09 |
| ZNF296 | -4.526986956 | 0.000857933 | 0.007185533 |
| NUB1 | -4.524856751 | 2.50E-08 | 9.19E-07 |
| FAIM3 | -4.507862503 | 3.34E-08 | 1.19E-06 |
| ITGB2-AS1 | -4.491040975 | 1.98E-06 | 4.27E-05 |
| STX16-NPEPL1 | -4.459995108 | 0.000834349 | 0.007010503 |
| IRS2 | -4.455293576 | 2.26E-09 | 1.11E-07 |
| PLXND1 | -4.444087569 | 0.002243685 | 0.015687764 |
| FAM179A | -4.442929538 | 3.53E-09 | 1.68E-07 |
| SMG1P3 | -4.424482881 | 0.000730595 | 0.006273345 |
| COX11 | -4.417955299 | 1.79E-06 | 3.93E-05 |
| ZNF76 | -4.397690119 | 0.000239682 | 0.002502602 |
| THEM6 | -4.397058098 | 0.000908225 | 0.007542217 |
| SPTBN1 | -4.381721253 | 1.94E-13 | 2.54E-11 |
| LTB4R | -4.377522932 | 5.18E-06 | 9.94E-05 |
| VILL | -4.373467963 | 0.001362172 | 0.010500813 |
| ASCL2 | -4.368747892 | 0.000305043 | 0.003028381 |
| IQSEC3 | -4.349701432 | 0.002393862 | 0.016502052 |
| NRM | -4.349397359 | 0.000404013 | 0.00382567 |
| LSP1 | -4.346693884 | 3.88E-08 | 1.35E-06 |
| TMCO4 | -4.343870963 | 0.003324347 | 0.02148683 |
| B3GNT7 | -4.335088037 | 5.02E-08 | 1.67E-06 |
| MAST3 | -4.333845881 | 4.80E-08 | 1.63E-06 |
| ABLIM1 | -4.328222771 | 2.72E-09 | 1.32E-07 |
| GPATCH3 | -4.326459028 | 2.99E-05 | 0.000449146 |
| DLG3 | -4.325417324 | 2.62E-05 | 0.000403893 |
| CCDC141 | -4.325383148 | 0.000303602 | 0.003020972 |
| TC2N | -4.31647124 | 1.17E-05 | 0.000202667 |
| CCBL1 | -4.299347891 | 0.001817132 | 0.013224871 |
| CTSO | -4.287096304 | 3.00E-07 | 8.19E-06 |
| PATL2 | -4.28577204 | 4.98E-06 | 9.68E-05 |
| ZNF831 | -4.282033355 | 3.21E-06 | 6.55E-05 |
| CX3CR1 | -4.279434655 | 2.93E-05 | 0.000441091 |
| LOC115110 | -4.274681585 | 0.001742106 | 0.01274994 |
| TMEM91 | -4.273359362 | 0.002337148 | 0.016254178 |
| GSTK1 | -4.272082052 | 4.02E-07 | 1.05E-05 |
| SAMHD1 | -4.25156107 | 1.80E-06 | 3.95E-05 |
| ITGB2 | -4.23534779 | 1.90E-16 | 4.02E-14 |

| | | | |
|---|---|---|---|
| RNF103 | -4.221444122 | 2.52E-05 | 0.000391207 |
| ATP6V1E2 | -4.216677291 | 0.003755489 | 0.023717126 |
| CEP85 | -4.216223859 | 0.002388896 | 0.016496864 |
| PDPR | -4.215815172 | 0.000406078 | 0.003840575 |
| HIST1H1C | -4.21329634 | 6.08E-06 | 0.000114265 |
| CD84 | -4.204581906 | 1.57E-05 | 0.000259112 |
| ZBTB25 | -4.202910808 | 0.004758779 | 0.028360731 |
| IFT46 | -4.191512174 | 0.004665596 | 0.027932938 |
| STMN1 | -4.184320881 | 5.13E-08 | 1.69E-06 |
| PRR3 | -4.169646158 | 0.000274611 | 0.002781992 |
| LGALS9 | -4.167363722 | 3.99E-09 | 1.86E-07 |
| ATF7 | -4.161434039 | 3.44E-06 | 6.99E-05 |
| LLGL2 | -4.157791887 | 1.20E-08 | 4.98E-07 |
| FBXO25 | -4.152825407 | 5.83E-06 | 0.000110617 |
| DTX3 | -4.12852423 | 2.61E-06 | 5.41E-05 |
| ZNF160 | -4.11680559 | 0.00569484 | 0.032083663 |
| BAALC | -4.102692678 | 0.005356606 | 0.030861523 |
| TACC3 | -4.088641023 | 0.000565618 | 0.005073713 |
| ZDHHC4 | -4.077179511 | 0.000134365 | 0.001549655 |
| CCDC88C | -4.05378577 | 3.42E-14 | 5.05E-12 |
| ZNF337 | -4.052587608 | 0.001828669 | 0.013296478 |
| KAT7 | -4.045527233 | 2.41E-06 | 5.09E-05 |
| RAD9A | -4.041244535 | 0.003828481 | 0.024088421 |
| TAPBPL | -4.03836325 | 2.78E-07 | 7.65E-06 |
| ANKAR | -4.026090933 | 0.000121791 | 0.001430572 |
| HIST1H2AM | -4.026042296 | 0.000188689 | 0.002057973 |
| S100A4 | -4.022861924 | 1.42E-06 | 3.18E-05 |
| LOC440434 | -4.014122627 | 3.92E-05 | 0.000560015 |
| HIST1H1D | -4.012619403 | 6.23E-05 | 0.000839748 |
| BLOC1S5 | -4.009107688 | 0.000447624 | 0.004158174 |
| EXTL2 | -3.997822634 | 0.000885186 | 0.007380206 |
| IFT20 | -3.995905103 | 1.66E-05 | 0.00027248 |
| GAB3 | -3.990484857 | 1.97E-08 | 7.53E-07 |
| SPDYE3 | -3.972113519 | 0.006019148 | 0.033374021 |
| PSEN2 | -3.961988518 | 0.004796781 | 0.02852209 |
| ANKMY2 | -3.957856384 | 6.02E-05 | 0.000814683 |
| NMUR1 | -3.955283042 | 8.84E-05 | 0.001098848 |
| ZNF318 | -3.951259651 | 8.39E-05 | 0.00105661 |
| C19orf52 | -3.932503765 | 0.006819993 | 0.036617966 |
| RBM38 | -3.924843443 | 2.05E-06 | 4.38E-05 |
| FBXO33 | -3.914512902 | 5.59E-05 | 0.000765148 |
| INVS | -3.896493552 | 0.004849571 | 0.028792262 |
| FAM49A | -3.89520302 | 4.60E-05 | 0.00064329 |
| SFT2D2 | -3.89107763 | 0.00066067 | 0.005748565 |
| TTC38 | -3.881509195 | 3.91E-05 | 0.000560015 |
| AHNAK | -3.878458333 | 1.11E-06 | 2.60E-05 |
| FAM189B | -3.874802455 | 0.00984789 | 0.047870159 |
| LEPROTL1 | -3.861386089 | 0.000130251 | 0.001508867 |
| PTPN22 | -3.856718235 | 1.48E-13 | 2.00E-11 |
| PRORSD1P | -3.842147545 | 0.000264178 | 0.002697236 |

| | | | |
|---|---|---|---|
| A2M-AS1 | -3.838773337 | 0.001659166 | 0.012257484 |
| UCP2 | -3.838419438 | 8.86E-12 | 8.07E-10 |
| FIS1 | -3.836971599 | 6.13E-05 | 0.000827411 |
| AK1 | -3.820785715 | 0.00044229 | 0.004123302 |
| GALT | -3.820462654 | 0.005608806 | 0.03178188 |
| ITPK1-AS1 | -3.817490699 | 0.007607054 | 0.039477032 |
| IPO13 | -3.809008657 | 0.008423268 | 0.042666633 |
| RNF214 | -3.796699819 | 0.001410914 | 0.010771375 |
| BZRAP1 | -3.796623352 | 0.00192456 | 0.013890535 |
| TCEAL8 | -3.784797436 | 0.000636208 | 0.005581916 |
| RNF125 | -3.781986205 | 9.23E-08 | 2.82E-06 |
| HNRNPA1L2 | -3.781709157 | 0.001338019 | 0.010364022 |
| SPIN2B | -3.781166302 | 0.007170065 | 0.037785183 |
| UVRAG | -3.77119033 | 9.37E-07 | 2.21E-05 |
| TTN-AS1 | -3.770450204 | 0.005380019 | 0.030861523 |
| STK10 | -3.766326935 | 6.68E-07 | 1.64E-05 |
| CALML4 | -3.763359321 | 0.000162999 | 0.001815716 |
| MYRF | -3.758585856 | 0.000828273 | 0.006995838 |
| PIKFYVE | -3.752188109 | 6.60E-06 | 0.000122717 |
| DKFZP586I1420 | -3.751091968 | 2.59E-06 | 5.37E-05 |
| SEMA4C | -3.750203217 | 0.000545238 | 0.004913419 |
| GOLIM4 | -3.744272645 | 0.000121905 | 0.001430572 |
| TSC22D3 | -3.74402484 | 1.80E-09 | 9.26E-08 |
| VAMP1 | -3.739753918 | 9.09E-05 | 0.001125177 |
| ATG9A | -3.738152554 | 0.000280027 | 0.002825888 |
| KCNQ5 | -3.718461035 | 0.004122566 | 0.025500642 |
| OBFC1 | -3.715650755 | 0.000253494 | 0.002615428 |
| CBX3P2 | -3.695535029 | 0.001127305 | 0.008976841 |
| KIAA1586 | -3.672609815 | 0.00859147 | 0.043294597 |
| AHSA2 | -3.671957518 | 1.17E-07 | 3.52E-06 |
| SLC16A6 | -3.671346621 | 2.46E-07 | 6.96E-06 |
| RNMTL1 | -3.668497168 | 0.007069538 | 0.03747836 |
| ZNF92 | -3.663826002 | 0.000502053 | 0.004576926 |
| PYHIN1 | -3.658754071 | 5.50E-13 | 6.65E-11 |
| GZMM | -3.647016912 | 0.000312168 | 0.003086598 |
| MADD | -3.645899426 | 5.63E-07 | 1.41E-05 |
| CHAF1A | -3.64130976 | 2.15E-05 | 0.000340136 |
| COLGALT2 | -3.636267875 | 0.000421807 | 0.003965391 |
| MAP4K1 | -3.626560179 | 1.80E-08 | 6.93E-07 |
| WHAMMP3 | -3.622226323 | 0.000172682 | 0.0019073 |
| VPS51 | -3.621997285 | 8.04E-07 | 1.92E-05 |
| MPPE1 | -3.62020307 | 0.000743032 | 0.006352275 |
| TSC22D1 | -3.619279456 | 0.000203323 | 0.002167033 |
| FCGR3A | -3.604709648 | 5.58E-06 | 0.000106565 |
| CRYZ | -3.604065266 | 0.000644403 | 0.00563205 |
| ITGAM | -3.594675151 | 6.48E-06 | 0.00012082 |
| NCALD | -3.593424353 | 7.22E-07 | 1.75E-05 |
| RASSF1 | -3.590234283 | 8.00E-08 | 2.50E-06 |
| BCL9L | -3.567920408 | 0.000129212 | 0.001499054 |
| ADAMTS10 | -3.560882362 | 0.004295051 | 0.026276989 |

| | | | |
|---|---|---|---|
| PREX1 | -3.558603361 | 1.33E-06 | 3.04E-05 |
| IFITM1 | -3.55781372 | 2.06E-19 | 8.51E-17 |
| NHLRC3 | -3.552315875 | 0.000261202 | 0.002670326 |
| METTL3 | -3.549862672 | 0.000805292 | 0.006817559 |
| DTHD1 | -3.540663704 | 0.00624869 | 0.034291165 |
| ARSD | -3.52863489 | 0.001375602 | 0.010569839 |
| ADD3 | -3.528583492 | 3.63E-09 | 1.72E-07 |
| RBL2 | -3.521411365 | 2.11E-07 | 6.06E-06 |
| SMYD3 | -3.51701314 | 0.000832956 | 0.007006312 |
| DUSP1 | -3.508176185 | 3.22E-05 | 0.000476716 |
| AKAP2 | -3.506852675 | 0.000810106 | 0.006850911 |
| ITFG2 | -3.502109719 | 0.006074002 | 0.033567756 |
| SLC35A4 | -3.499641443 | 0.000885889 | 0.007380206 |
| DOPEY1 | -3.495021123 | 0.000339277 | 0.003312816 |
| ADA | -3.491849222 | 0.000323534 | 0.003178913 |
| TBC1D14 | -3.481109202 | 0.005959862 | 0.033171058 |
| SAMD3 | -3.475683571 | 1.39E-06 | 3.12E-05 |
| PLCL2 | -3.455383275 | 2.88E-05 | 0.000435683 |
| NDST2 | -3.445373909 | 0.000115601 | 0.001365415 |
| PRSS23 | -3.435447616 | 6.70E-05 | 0.000885102 |
| PLSCR3 | -3.431196556 | 0.000940993 | 0.007764929 |
| TFDP2 | -3.426799083 | 6.82E-11 | 5.04E-09 |
| ZMYM3 | -3.422229071 | 0.001377842 | 0.010569839 |
| H1FX | -3.421988135 | 7.32E-05 | 0.000945529 |
| LAIR2 | -3.421263586 | 9.10E-05 | 0.001125177 |
| VPS26B | -3.420607812 | 9.64E-05 | 0.001184825 |
| ADH5 | -3.418566071 | 3.85E-05 | 0.000553762 |
| AUTS2 | -3.412325358 | 2.17E-10 | 1.46E-08 |
| DCTN3 | -3.405616984 | 9.19E-05 | 0.001135659 |
| AREG | -3.389874695 | 0.000376715 | 0.003595154 |
| TRIM5 | -3.369470462 | 0.003155242 | 0.020694052 |
| SLA2 | -3.354651332 | 0.000524553 | 0.004759878 |
| HSH2D | -3.351038354 | 1.22E-06 | 2.79E-05 |
| SYNE1 | -3.345775573 | 2.44E-11 | 2.01E-09 |
| BNIP3L | -3.3457001 | 8.02E-08 | 2.50E-06 |
| GIMAP4 | -3.342927654 | 0.002097711 | 0.01486622 |
| NR1D2 | -3.342641373 | 7.24E-05 | 0.000937978 |
| RRN3P1 | -3.338027864 | 0.00197048 | 0.014143748 |
| RASGRP2 | -3.335297024 | 0.000142939 | 0.001626967 |
| TRERF1 | -3.32689487 | 0.000197411 | 0.002122615 |
| NDRG1 | -3.316620676 | 7.16E-06 | 0.000132221 |
| IFITM2 | -3.315795483 | 2.00E-06 | 4.30E-05 |
| ADRB2 | -3.315342571 | 0.00108442 | 0.008709842 |
| SORL1 | -3.293194131 | 1.48E-07 | 4.37E-06 |
| ERCC5 | -3.291100378 | 1.07E-07 | 3.24E-06 |
| LOC100130872 | -3.290285283 | 0.001917059 | 0.013861948 |
| RASA3 | -3.287981469 | 5.61E-07 | 1.41E-05 |
| TRMT13 | -3.270777293 | 4.02E-05 | 0.000572607 |
| SCML4 | -3.253239047 | 0.001500783 | 0.011311482 |
| OSBPL5 | -3.252631917 | 0.000381085 | 0.003621695 |

| | | | |
|---|---|---|---|
| TLE1 | -3.246535902 | 0.000135896 | 0.001564997 |
| GYPC | -3.235773497 | 4.03E-05 | 0.000573106 |
| ABCA7 | -3.231583072 | 0.00483144 | 0.028706378 |
| ESYT1 | -3.224544907 | 1.28E-05 | 0.000216641 |
| KRT72 | -3.224451573 | 0.008068569 | 0.041243451 |
| CHST2 | -3.202475308 | 3.75E-13 | 4.67E-11 |
| MVD | -3.201475396 | 0.003380657 | 0.021735569 |
| CCND3 | -3.197542832 | 0.000485535 | 0.004467948 |
| MRS2P2 | -3.193464305 | 0.002565708 | 0.017441024 |
| RBM15 | -3.193088524 | 0.001166002 | 0.009251225 |
| SEC14L1P1 | -3.189501615 | 0.003473621 | 0.022241969 |
| DGKZ | -3.17856445 | 8.23E-06 | 0.000148848 |
| GPR56 | -3.167445444 | 2.56E-12 | 2.74E-10 |
| BEX2 | -3.158990193 | 0.003309462 | 0.02141851 |
| MYLIP | -3.151474493 | 1.21E-07 | 3.62E-06 |
| HSPA7 | -3.149209214 | 0.000110796 | 0.001324244 |
| ILF3-AS1 | -3.148348514 | 2.81E-05 | 0.000427201 |
| LPCAT1 | -3.147404547 | 2.18E-09 | 1.08E-07 |
| SS18L1 | -3.142815454 | 0.000698358 | 0.006022957 |
| C10orf54 | -3.132727313 | 0.001389086 | 0.010636054 |
| IFI6 | -3.119643692 | 4.75E-06 | 9.27E-05 |
| PEPD | -3.115712893 | 0.003390485 | 0.021780878 |
| RNF166 | -3.111749278 | 1.14E-08 | 4.80E-07 |
| SLC25A42 | -3.111177772 | 0.001562993 | 0.01169764 |
| PML | -3.109297543 | 0.001891552 | 0.013728214 |
| HPS3 | -3.100355996 | 0.001060118 | 0.008558541 |
| DENND1C | -3.092066044 | 1.17E-05 | 0.000202466 |
| INTS4 | -3.090856594 | 0.000457003 | 0.00423025 |
| KAT8 | -3.08201767 | 0.000657431 | 0.005726744 |
| TYROBP | -3.079451893 | 1.03E-13 | 1.42E-11 |
| ATG4A | -3.078016383 | 0.003488883 | 0.022321441 |
| ATHL1 | -3.075389218 | 0.001035397 | 0.008402275 |
| PPP2R2B | -3.071883296 | 6.90E-05 | 0.000905429 |
| CEP290 | -3.06784504 | 2.48E-05 | 0.000385364 |
| PPM1M | -3.059767841 | 0.00011179 | 0.001324393 |
| NCR3 | -3.056969149 | 0.004429837 | 0.026849886 |
| VPS37B | -3.052027182 | 8.78E-05 | 0.001094676 |
| RASSF4 | -3.048075067 | 0.001447182 | 0.010992123 |
| ELF4 | -3.041837622 | 0.005311249 | 0.030688306 |
| DENND1A | -3.03875115 | 0.008445145 | 0.042722178 |
| ETFB | -3.037398437 | 0.002233813 | 0.015646678 |
| UVSSA | -3.032588884 | 0.000906567 | 0.007536443 |
| ATP8A1 | -3.026145958 | 2.47E-11 | 2.02E-09 |
| FKTN | -3.015833992 | 0.00034581 | 0.003359846 |
| TMEM9 | -3.015821013 | 0.008476231 | 0.042851753 |
| TTC39B | -3.011846518 | 0.006299696 | 0.034522688 |
| CXXC5 | -3.010929204 | 2.03E-05 | 0.000325617 |
| ICAM3 | -3.009045076 | 0.001632805 | 0.012138483 |
| MAPK8IP3 | -3.002841997 | 0.005398007 | 0.030879112 |
| SIPA1 | -3.002310504 | 0.002024419 | 0.014451437 |

| | | | |
|---|---|---|---|
| CPT1A | -2.994458149 | 6.47E-12 | 6.33E-10 |
| NDUFAF5 | -2.988451561 | 0.003177123 | 0.020802718 |
| DOK2 | -2.9862829 | 0.003015108 | 0.019958845 |
| FBXO32 | -2.983578816 | 0.00015707 | 0.001759683 |
| RABGAP1L | -2.981068247 | 4.83E-07 | 1.23E-05 |
| TRIB2 | -2.978109575 | 9.64E-06 | 0.000171602 |
| ADCK3 | -2.977108529 | 0.004589566 | 0.027583187 |
| SUN2 | -2.962248042 | 1.09E-10 | 7.69E-09 |
| CROCCP2 | -2.940743129 | 0.000368762 | 0.003532988 |
| TCF7 | -2.938360339 | 0.00376426 | 0.023753359 |
| CTSW | -2.936272812 | 2.45E-06 | 5.13E-05 |
| KIR2DL3 | -2.935888268 | 0.002910675 | 0.019365759 |
| MYO1F | -2.929275689 | 3.62E-08 | 1.27E-06 |
| ADAMTS1 | -2.923167832 | 0.00388664 | 0.024349019 |
| APOBR | -2.922898191 | 0.000229699 | 0.002414465 |
| DCK | -2.908976364 | 2.26E-05 | 0.000355884 |
| PI4KA | -2.908551925 | 3.92E-05 | 0.000560015 |
| MED20 | -2.908373389 | 0.007183112 | 0.03782848 |
| HPS1 | -2.907993522 | 5.90E-06 | 0.000111533 |
| MARK4 | -2.906120291 | 0.006112128 | 0.033730849 |
| CCDC144B | -2.905722005 | 0.00370606 | 0.023480708 |
| TBC1D7 | -2.893897677 | 0.008502789 | 0.042930587 |
| PLEKHA2 | -2.887788456 | 3.40E-07 | 9.00E-06 |
| PPP2R5A | -2.885810048 | 5.40E-08 | 1.76E-06 |
| BTN3A1 | -2.88253799 | 6.01E-06 | 0.000113107 |
| FAM134A | -2.881186027 | 0.003968303 | 0.024820909 |
| KIR3DL1 | -2.870063542 | 0.008315693 | 0.042230991 |
| ARHGAP27 | -2.867306987 | 1.94E-06 | 4.21E-05 |
| CDK5R1 | -2.8559286 | 0.001411241 | 0.010771375 |
| IL10RB | -2.855279222 | 0.002510414 | 0.017124611 |
| GLIPR1 | -2.854163682 | 3.42E-05 | 0.000499459 |
| WDR13 | -2.853046968 | 0.002898992 | 0.019337313 |
| SNX15 | -2.848008926 | 0.004572627 | 0.027502488 |
| RAE1 | -2.845022821 | 0.008146387 | 0.041586936 |
| KIAA1407 | -2.840757942 | 0.005845354 | 0.032743179 |
| KDM1B | -2.838798899 | 0.006197042 | 0.034055461 |
| DDX28 | -2.835460463 | 0.008003595 | 0.0409916 |
| CUTA | -2.833521495 | 0.006825427 | 0.036617966 |
| DST | -2.82976052 | 0.003608641 | 0.022956349 |
| COQ4 | -2.828153198 | 0.007305281 | 0.038189355 |
| ZNF394 | -2.820057634 | 0.00208695 | 0.014803353 |
| NSD1 | -2.814611773 | 0.000199692 | 0.002139244 |
| MTSS1 | -2.802255454 | 0.007052561 | 0.037417752 |
| CENPJ | -2.795454945 | 0.005379188 | 0.030861523 |
| SPSB3 | -2.795040922 | 0.00096185 | 0.007912022 |
| MKI67 | -2.789144445 | 8.03E-06 | 0.000145945 |
| FGL2 | -2.787204232 | 0.004249485 | 0.026018546 |
| MGEA5 | -2.786136297 | 4.35E-05 | 0.000614105 |
| PDE4A | -2.780051338 | 0.004007875 | 0.024892083 |
| FGFBP2 | -2.767024176 | 1.34E-05 | 0.000225039 |

| | | | |
|---|---|---|---|
| ADCY7 | -2.766067576 | 1.40E-08 | 5.65E-07 |
| ZNF600 | -2.759575816 | 0.001514665 | 0.011383246 |
| RETSAT | -2.758293269 | 0.007780334 | 0.040163347 |
| RNF126 | -2.743904853 | 0.006882912 | 0.036867361 |
| ATP10A | -2.731363149 | 0.006183392 | 0.034052137 |
| TMEM161B | -2.726712978 | 0.006062367 | 0.033527113 |
| ATG2A | -2.722896554 | 0.002722207 | 0.018361419 |
| SLC2A3 | -2.715567307 | 2.81E-08 | 1.02E-06 |
| DZIP3 | -2.710658294 | 1.73E-05 | 0.000282568 |
| SELPLG | -2.709873417 | 0.000376915 | 0.003595154 |
| HCST | -2.709330086 | 2.36E-06 | 5.00E-05 |
| LOC283070 | -2.687850087 | 0.008329513 | 0.042273761 |
| TNIK | -2.68660148 | 1.33E-05 | 0.000223845 |
| TMEM120B | -2.679871116 | 0.00366234 | 0.023222496 |
| RREB1 | -2.675143389 | 0.005637804 | 0.031821178 |
| LINC00861 | -2.661348183 | 0.000990573 | 0.008105724 |
| BIN2 | -2.654621814 | 1.86E-05 | 0.000301381 |
| UCHL5 | -2.646781629 | 0.003286961 | 0.021343444 |
| TMIGD2 | -2.642946089 | 0.003715077 | 0.023518813 |
| ZNF263 | -2.64229915 | 0.009930849 | 0.048243471 |
| ITGA6 | -2.638231638 | 0.006964448 | 0.037151632 |
| LIMA1 | -2.63608223 | 0.006190528 | 0.034055461 |
| HERC5 | -2.635709836 | 0.004703287 | 0.028115602 |
| HLA-DPB1 | -2.635451034 | 0.001897978 | 0.013762101 |
| NLRC5 | -2.634330193 | 3.62E-05 | 0.00052658 |
| ICAM2 | -2.617823654 | 0.001108665 | 0.008850106 |
| PDK3 | -2.613058189 | 0.002460095 | 0.016825329 |
| ARHGAP9 | -2.609030506 | 2.59E-05 | 0.0003994 |
| FLOT2 | -2.603331819 | 0.000274349 | 0.002781992 |
| CXorf65 | -2.580067697 | 0.004556154 | 0.027445573 |
| LOC101928673 | -2.573658177 | 0.008878097 | 0.044354369 |
| TRAPPC2 | -2.571111378 | 0.00355191 | 0.022650656 |
| IL12RB1 | -2.571065251 | 0.00017781 | 0.001955657 |
| TSPO | -2.55654731 | 0.00502712 | 0.029621803 |
| CTAGE5 | -2.553792954 | 0.00509916 | 0.029835795 |
| AP2A2 | -2.551626519 | 0.001227836 | 0.0096538 |
| MLC1 | -2.542804453 | 0.001767854 | 0.012902204 |
| KLRG1 | -2.539891558 | 0.008798849 | 0.044056131 |
| RNPEPL1 | -2.535543167 | 0.005985512 | 0.033242941 |
| PXN | -2.529178439 | 3.22E-10 | 2.10E-08 |
| FAM117A | -2.520915957 | 0.00654347 | 0.035683784 |
| USPL1 | -2.513182313 | 0.00110069 | 0.008807176 |
| NNT | -2.504232297 | 0.002336882 | 0.016254178 |
| HCFC1 | -2.496443504 | 0.000145536 | 0.001646951 |
| UNC13D | -2.49608705 | 0.003243257 | 0.021112177 |
| CDC25B | -2.490484966 | 0.000168731 | 0.001871573 |
| ZAP70 | -2.489403987 | 0.000111576 | 0.001324393 |
| TGFB1 | -2.476822457 | 0.007265287 | 0.038090857 |
| ACTN1 | -2.475960601 | 0.006740488 | 0.03632812 |
| IQSEC1 | -2.47266922 | 4.37E-06 | 8.63E-05 |

| | | | |
|---|---|---|---|
| TXNIP | -2.468265922 | 5.60E-10 | 3.45E-08 |
| IFNAR2 | -2.459242762 | 0.000389987 | 0.003701803 |
| PDP1 | -2.458682557 | 0.008019566 | 0.04104655 |
| HAUS3 | -2.455242572 | 0.004001665 | 0.024892083 |
| CYBA | -2.435995933 | 0.003878584 | 0.024318009 |
| NMRAL1 | -2.429406867 | 0.002534248 | 0.01725713 |
| PITPNC1 | -2.424191817 | 0.002561013 | 0.017424231 |
| CTSS | -2.417666124 | 0.003293423 | 0.021357445 |
| ACADSB | -2.415621998 | 0.009971971 | 0.04841321 |
| PLEKHM3 | -2.408587204 | 0.00967917 | 0.047284831 |
| ADAT2 | -2.401691338 | 0.003325857 | 0.02148683 |
| C19orf66 | -2.394620065 | 2.92E-05 | 0.000440375 |
| ZNF91 | -2.393684733 | 0.00151845 | 0.011400748 |
| IPP | -2.388408492 | 0.001170459 | 0.009267814 |
| CYTH1 | -2.382382903 | 9.59E-07 | 2.25E-05 |
| LAPTM5 | -2.381366354 | 0.000107 | 0.001287125 |
| CCDC50 | -2.3793182 | 0.003431269 | 0.022006771 |
| UBA7 | -2.375408128 | 0.006559761 | 0.035698691 |
| MAPK13 | -2.367625716 | 0.001579081 | 0.011788161 |
| GLIPR2 | -2.360444385 | 0.007008205 | 0.037258149 |
| APBB1IP | -2.359447629 | 0.000140379 | 0.001604834 |
| SLC11A2 | -2.348693326 | 6.69E-05 | 0.000885102 |
| DGKQ | -2.348422693 | 0.005483209 | 0.031251096 |
| PRMT2 | -2.342797228 | 6.64E-06 | 0.000123216 |
| APLP2 | -2.338007155 | 0.002972629 | 0.019694295 |
| CASP10 | -2.326368185 | 0.005311461 | 0.030688306 |
| SLC25A46 | -2.322061249 | 0.000153414 | 0.001723651 |
| RNF44 | -2.320000296 | 3.04E-05 | 0.000453786 |
| PIK3R1 | -2.305151108 | 8.24E-08 | 2.56E-06 |
| RABL2B | -2.30077143 | 0.007132327 | 0.037618334 |
| AGO1 | -2.299669632 | 0.007102771 | 0.037531578 |
| COPS8 | -2.297620455 | 0.000975865 | 0.008010478 |
| MAZ | -2.295832273 | 0.008268606 | 0.042101074 |
| ZFP36L2 | -2.291865851 | 0.000622789 | 0.005486006 |
| GPCPD1 | -2.291095446 | 0.005968651 | 0.03317282 |
| C12orf75 | -2.2829897 | 2.07E-05 | 0.000331599 |
| GTPBP1 | -2.273157632 | 0.005043162 | 0.029671679 |
| EVL | -2.271717555 | 1.40E-07 | 4.16E-06 |
| SH3BGRL | -2.270933588 | 0.002400691 | 0.016531374 |
| TRAM2 | -2.270558742 | 0.006753134 | 0.036371248 |
| LCK | -2.270138472 | 0.007004389 | 0.037258149 |
| ACAP2 | -2.268038647 | 0.000315868 | 0.003115318 |
| GIMAP2 | -2.261302833 | 0.001397911 | 0.010690467 |
| CCL28 | -2.261172186 | 0.009055471 | 0.044938777 |
| AES | -2.257326752 | 5.29E-08 | 1.73E-06 |
| SUPT16H | -2.251836865 | 2.68E-05 | 0.000410968 |
| AP1G2 | -2.243565785 | 0.003404892 | 0.021855501 |
| PDE7A | -2.239106938 | 0.00043516 | 0.004074682 |
| C14orf159 | -2.236902721 | 0.005554707 | 0.031520952 |
| STX8 | -2.235111365 | 0.008078549 | 0.041267524 |

| | | | |
|---|---|---|---|
| CBX4 | -2.23322922 | 0.006908403 | 0.036953348 |
| ARF3 | -2.23066192 | 0.006602202 | 0.035750236 |
| FAM208A | -2.220886443 | 0.000320716 | 0.003155182 |
| IL10RA | -2.197870392 | 0.001040544 | 0.0084353 |
| PFKL | -2.197283818 | 0.005965443 | 0.03317282 |
| TRIM8 | -2.193775785 | 0.009565804 | 0.046847912 |
| KIAA1109 | -2.182152868 | 6.69E-05 | 0.000885102 |
| SSH2 | -2.182120758 | 0.000584811 | 0.005186472 |
| TAF1 | -2.181270383 | 0.009707437 | 0.047393351 |
| ESYT2 | -2.180413851 | 0.003294572 | 0.021357445 |
| P2RY8 | -2.180384242 | 5.07E-06 | 9.81E-05 |
| ARHGAP4 | -2.177619301 | 0.007639045 | 0.039590574 |
| DBNL | -2.175409907 | 0.00199929 | 0.014324279 |
| NLRC3 | -2.173816344 | 0.00134805 | 0.010421106 |
| CAMK1D | -2.167824076 | 0.002450322 | 0.01679035 |
| ARHGEF6 | -2.165206719 | 0.003184238 | 0.020831883 |
| MATK | -2.160497877 | 0.000452087 | 0.004189698 |
| PIP4K2A | -2.150657912 | 1.00E-05 | 0.000177038 |
| TBCC | -2.146705861 | 0.008001496 | 0.0409916 |
| MAPK3 | -2.145125363 | 0.00421111 | 0.025864475 |
| SPECC1L | -2.11035808 | 0.005375555 | 0.030861523 |
| PAPD5 | -2.109105033 | 0.000111719 | 0.001324393 |
| PCNT | -2.102138226 | 0.000487682 | 0.004479888 |
| CCDC115 | -2.099698951 | 0.007244833 | 0.038025663 |
| EFHD2 | -2.093379695 | 1.21E-05 | 0.000207527 |
| NIN | -2.091050409 | 2.49E-07 | 6.96E-06 |
| PRKCB | -2.089357427 | 1.52E-06 | 3.39E-05 |
| SHISA5 | -2.076067797 | 0.009726451 | 0.047427047 |
| WHAMMP2 | -2.07543376 | 0.000230564 | 0.002420308 |
| ARHGEF18 | -2.070980189 | 0.00460893 | 0.027635934 |
| MAPK14 | -2.065453032 | 0.005091364 | 0.029835795 |
| TRAF3IP3 | -2.064448227 | 2.23E-06 | 4.73E-05 |
| CDKN2D | -2.05071526 | 0.006569381 | 0.035700777 |
| HMGB2 | -2.046537203 | 1.18E-05 | 0.000203032 |
| MRPS18B | -2.04582603 | 0.001003137 | 0.008197897 |
| S1PR5 | -2.045523239 | 0.002910552 | 0.019365759 |
| OPTN | -2.039510688 | 4.07E-05 | 0.00057722 |
| GMFG | -2.030164648 | 5.84E-05 | 0.000794765 |
| PTPRCAP | -2.029029962 | 0.000198445 | 0.002128793 |
| MRPL10 | -2.02082547 | 0.004153498 | 0.025651456 |
| FBXO42 | -2.016378369 | 0.007587326 | 0.039453088 |
| DEF6 | -2.008073954 | 0.001720449 | 0.012634736 |
| WHAMM | -2.005490604 | 9.89E-06 | 0.000175244 |
| ODF2L | -2.005121562 | 0.00363127 | 0.023052541 |
| ADD1 | -2.003176055 | 5.49E-05 | 0.000752323 |
| SETD2 | -1.997459701 | 8.02E-05 | 0.001020167 |
| GZMA | -1.991457081 | 3.52E-08 | 1.24E-06 |
| SHOC2 | -1.987739901 | 0.008682567 | 0.043557454 |
| KAT6B | -1.985456406 | 2.02E-05 | 0.000325095 |
| EIF2AK2 | -1.974522291 | 1.21E-05 | 0.000207587 |

| | | | |
|---|---|---|---|
| FMR1 | -1.973586627 | 9.63E-05 | 0.001184825 |
| CNOT1 | -1.972148214 | 0.007001552 | 0.037258149 |
| RAB11FIP2 | -1.954976746 | 0.001553908 | 0.011644645 |
| NKG7 | -1.953962392 | 0.000158904 | 0.001777682 |
| FAM21C | -1.940875566 | 0.002206974 | 0.015500166 |
| HTT | -1.939200449 | 0.000974954 | 0.008010478 |
| C6orf106 | -1.93665751 | 0.00592649 | 0.033032275 |
| CTBP1 | -1.930692283 | 0.000129 | 0.00149881 |
| KANSL1 | -1.928645914 | 0.00137011 | 0.010548257 |
| FAM214A | -1.924392295 | 5.51E-06 | 0.000105506 |
| FAM76B | -1.924217189 | 0.001250007 | 0.009808425 |
| ACAD8 | -1.922903137 | 0.009771233 | 0.047586148 |
| CEP85L | -1.912975243 | 0.001252027 | 0.00981444 |
| SMARCA2 | -1.9032384 | 0.001646906 | 0.012212996 |
| MAPRE2 | -1.900947509 | 0.000189499 | 0.002058203 |
| ZNF548 | -1.885672311 | 0.006700916 | 0.036189566 |
| NIPBL | -1.884452395 | 0.001131819 | 0.008989121 |
| ULK3 | -1.879682662 | 0.005227796 | 0.030392626 |
| ZNF276 | -1.877739817 | 0.007991384 | 0.040982665 |
| ARHGDIB | -1.869841746 | 0.001007149 | 0.008215606 |
| CALM1 | -1.868351625 | 6.70E-05 | 0.000885102 |
| FAM65B | -1.86438653 | 5.03E-06 | 9.75E-05 |
| ANXA6 | -1.847470811 | 0.000380602 | 0.003621496 |
| PSPC1 | -1.841105353 | 0.002907136 | 0.019365759 |
| DOCK11 | -1.83001141 | 0.001057938 | 0.008549751 |
| DDX6 | -1.829468743 | 0.001077444 | 0.0086716 |
| PSAP | -1.811445826 | 1.10E-05 | 0.000191498 |
| RCOR3 | -1.808268078 | 0.005127991 | 0.029901187 |
| SRSF5 | -1.805023301 | 4.73E-06 | 9.27E-05 |
| SPN | -1.796563184 | 0.007620008 | 0.039518068 |
| MYBL1 | -1.793303314 | 0.000221602 | 0.002337865 |
| CAPN2 | -1.784282126 | 0.002784377 | 0.018649375 |
| CNOT8 | -1.774914915 | 0.004003366 | 0.024892083 |
| IKZF5 | -1.76147013 | 0.006214468 | 0.034127278 |
| FLNA | -1.754827448 | 0.000241836 | 0.002518377 |
| KLRK1 | -1.747352286 | 0.000859161 | 0.007188133 |
| PIK3C2A | -1.747096805 | 0.002237228 | 0.0156566 |
| CST7 | -1.741256342 | 3.65E-05 | 0.000530034 |
| USP25 | -1.734293631 | 0.002420826 | 0.016643979 |
| LITAF | -1.732257413 | 0.000505594 | 0.004603847 |
| COMMD6 | -1.730922071 | 0.006915903 | 0.036968213 |
| CASP8 | -1.729747805 | 0.005451583 | 0.031138835 |
| CCDC69 | -1.728834565 | 3.77E-05 | 0.000543156 |
| MXD4 | -1.728458956 | 0.009404362 | 0.046259775 |
| CD247 | -1.727131073 | 2.15E-05 | 0.000339834 |
| IL16 | -1.710577593 | 0.002367725 | 0.016407666 |
| FYB | -1.700985906 | 7.25E-05 | 0.000937978 |
| IQGAP1 | -1.696442313 | 2.88E-05 | 0.000435683 |
| SMCHD1 | -1.687959842 | 3.34E-05 | 0.000488827 |
| ITGAL | -1.687817024 | 0.00019641 | 0.002115658 |

| | | | |
|---|---|---|---|
| ARHGAP30 | -1.665673879 | 0.000650039 | 0.005674975 |
| PSD4 | -1.66050081 | 0.003227461 | 0.021044333 |
| ITGB1 | -1.657828732 | 8.70E-05 | 0.00108868 |
| MBP | -1.642817765 | 0.002369694 | 0.016407666 |
| JARID2 | -1.635395298 | 0.007915961 | 0.040675783 |
| PTPRA | -1.623871173 | 0.009551996 | 0.046843647 |
| MYSM1 | -1.618940466 | 0.009824665 | 0.047816624 |
| ESCO1 | -1.616737484 | 0.005520254 | 0.031370907 |
| SIN3A | -1.616361626 | 0.000272922 | 0.002772053 |
| DCAF8 | -1.612315492 | 0.000946544 | 0.007802511 |
| JOSD1 | -1.610165825 | 0.000239351 | 0.002502485 |
| CNOT6L | -1.606526597 | 0.000100865 | 0.001226345 |
| FAM46A | -1.604118583 | 0.004749859 | 0.028350723 |
| DPP8 | -1.59934233 | 0.002004811 | 0.014350708 |
| BTN3A2 | -1.594528924 | 0.002166522 | 0.015257225 |
| CLK1 | -1.566030507 | 0.002769397 | 0.01858169 |
| HDAC7 | -1.544737218 | 0.002366038 | 0.016407666 |
| HADHA | -1.523859811 | 0.004162579 | 0.025670976 |
| LOC606724 | -1.516912534 | 0.001262546 | 0.009867263 |
| SLC9A3R1 | -1.506491393 | 0.000266838 | 0.002717305 |
| CMIP | -1.495565018 | 0.002402337 | 0.016531374 |
| HLA-E | -1.495456959 | 0.000344205 | 0.003348413 |
| SH2D1B | -1.481496357 | 0.00044841 | 0.004160543 |
| C1orf21 | -1.479315553 | 0.001633758 | 0.012138483 |
| DIAPH1 | -1.476608204 | 0.002839844 | 0.0189751 |
| HERPUD2 | -1.475956252 | 0.008779048 | 0.044013268 |
| ABHD17A | -1.473465829 | 0.002603898 | 0.017624134 |
| PNRC1 | -1.447334513 | 0.003278933 | 0.021308983 |
| EPB41 | -1.44206674 | 0.002939718 | 0.019509266 |
| HMHA1 | -1.435470522 | 0.003106001 | 0.020456763 |
| CNN2 | -1.425155637 | 0.007913077 | 0.040675783 |
| TAPBP | -1.419038066 | 0.003994606 | 0.024892083 |
| CD81 | -1.411688125 | 0.00333416 | 0.021507256 |
| RORA | -1.41025812 | 0.001563967 | 0.01169764 |
| STAT6 | -1.391266568 | 0.001326924 | 0.010303935 |
| CAST | -1.39122615 | 0.004009912 | 0.024892083 |
| WAS | -1.383614022 | 0.002044732 | 0.014569878 |
| SYF2 | -1.382332569 | 0.00525765 | 0.030543513 |
| INPP5D | -1.381147079 | 0.00444868 | 0.02691936 |
| GCC2 | -1.368384984 | 0.00201886 | 0.0144249 |
| UTRN | -1.365924084 | 0.006959475 | 0.037150406 |
| JAK1 | -1.346359692 | 0.001639495 | 0.012169558 |
| KLRB1 | -1.342620145 | 0.004687316 | 0.028041538 |
| ATP2B4 | -1.337232097 | 0.001930306 | 0.013919174 |
| MALAT1 | -1.334437713 | 0.000150556 | 0.001696412 |
| USP28 | -1.329908858 | 0.004707906 | 0.028121751 |
| SYAP1 | -1.312894804 | 0.007890785 | 0.040674367 |
| PTPRC | -1.310475287 | 0.000170105 | 0.001884147 |
| ZNF451 | -1.284000348 | 0.001486836 | 0.011227975 |
| PDCD4 | -1.266865706 | 0.005398225 | 0.030879112 |

| | | | |
|---|---|---|---|
| PTPN12 | -1.254547617 | 0.005360217 | 0.030861523 |
| RPL3 | -1.245495544 | 0.006192828 | 0.034055461 |
| XAF1 | -1.240398946 | 0.002070894 | 0.014716125 |
| MBNL1 | -1.232661812 | 0.007721227 | 0.039962379 |
| EID1 | -1.231003411 | 0.009396756 | 0.04625141 |
| BPTF | -1.205378746 | 0.000568123 | 0.005078783 |
| GNG2 | -1.197252721 | 0.005639067 | 0.031821178 |
| PCM1 | -1.195360896 | 0.004929123 | 0.029176085 |
| ZC3HAV1 | -1.190228769 | 0.001226674 | 0.0096538 |
| TTC3 | -1.17675658 | 0.007390024 | 0.038547772 |
| ARL4C | -1.168795512 | 0.004000856 | 0.024892083 |
| MYH9 | -1.164830807 | 0.009450408 | 0.046457091 |
| SERPINB1 | -1.152882018 | 0.006593803 | 0.035750236 |
| BCLAF1 | -1.137211562 | 0.005553892 | 0.031520952 |
| MYCBP2 | -1.119467395 | 0.005866413 | 0.032767392 |
| MACF1 | -1.114780532 | 0.007354841 | 0.038397174 |
| SAMD9L | -1.108901205 | 0.008881069 | 0.044354369 |
| U2SURP | -1.107656025 | 0.007812222 | 0.04030139 |
| NKTR | -1.080946214 | 0.008242124 | 0.041993541 |
| IQGAP2 | -1.030042406 | 0.005907721 | 0.032951113 |
| PCBP1 | 1.014864135 | 0.008681959 | 0.043557454 |
| EIF4A1 | 1.055509204 | 0.005213565 | 0.030332412 |
| CTSC | 1.063205646 | 0.008381744 | 0.042483779 |
| RBM8A | 1.088947293 | 0.006826999 | 0.036617966 |
| QKI | 1.118921796 | 0.004178806 | 0.025706384 |
| NAP1L1 | 1.121129486 | 0.00789491 | 0.040674367 |
| U2AF1 | 1.126123033 | 0.009559177 | 0.046844751 |
| PPP1CB | 1.143804312 | 0.001514336 | 0.011383246 |
| FAM107B | 1.165538638 | 0.003145478 | 0.020647308 |
| NFAT5 | 1.198915809 | 0.001729438 | 0.012680926 |
| LDHA | 1.223859227 | 0.008931718 | 0.0445505 |
| NOP56 | 1.229941832 | 0.001782724 | 0.012998613 |
| NPM1 | 1.252322097 | 0.002918628 | 0.019385733 |
| KDELR2 | 1.253340913 | 0.002917043 | 0.019385733 |
| LMO4 | 1.256139184 | 0.007073556 | 0.03747836 |
| UBA2 | 1.257059428 | 0.00702268 | 0.037309777 |
| HSPA5 | 1.261294077 | 0.000984901 | 0.008076188 |
| TCP1 | 1.273412727 | 0.001329531 | 0.010308475 |
| PIGY | 1.277302862 | 0.008822802 | 0.044147835 |
| PLEK | 1.284673187 | 0.001577206 | 0.011785401 |
| RQCD1 | 1.291041126 | 0.001470749 | 0.011127957 |
| NOL11 | 1.297238495 | 0.007912838 | 0.040675783 |
| ATP5J | 1.314752652 | 0.003814041 | 0.024028764 |
| RPL36AL | 1.316185026 | 0.000187427 | 0.002049914 |
| H2AFZ | 1.317015686 | 0.006618499 | 0.035793829 |
| HNRNPC | 1.334507031 | 0.000539159 | 0.004875465 |
| IWS1 | 1.339421005 | 0.005140978 | 0.029954614 |
| RYBP | 1.33949194 | 0.003224687 | 0.021043772 |
| CCT7 | 1.342561085 | 0.007595257 | 0.039468121 |
| IRF2BP2 | 1.348728947 | 0.001177014 | 0.009310302 |

| | | | |
|---|---|---|---|
| RCC2 | 1.354834437 | 0.002635616 | 0.017823408 |
| PPP1R16B | 1.355317673 | 0.007744052 | 0.040002419 |
| HNRNPF | 1.363535704 | 0.006423425 | 0.035077994 |
| SNRPF | 1.364446931 | 0.006857029 | 0.036753866 |
| ATP2A2 | 1.365957478 | 0.008990733 | 0.044759334 |
| SAR1B | 1.373754196 | 0.005396616 | 0.030879112 |
| EIF4H | 1.385970773 | 0.000619486 | 0.0054664 |
| POMP | 1.389015446 | 0.001989833 | 0.01426958 |
| CD53 | 1.413401885 | 0.000110931 | 0.001324244 |
| SFPQ | 1.415557279 | 0.006563126 | 0.035698691 |
| CD96 | 1.417904711 | 0.006351638 | 0.034741009 |
| SNX10 | 1.429675373 | 0.004229799 | 0.025938572 |
| LPAR6 | 1.432076942 | 0.004596723 | 0.027605014 |
| GCH1 | 1.434738644 | 0.001914166 | 0.01385382 |
| TRIM13 | 1.442038167 | 0.000302165 | 0.003010497 |
| HUWE1 | 1.447707193 | 0.001169123 | 0.009266601 |
| PPIB | 1.452850815 | 0.004314975 | 0.026337153 |
| ARL8B | 1.456470114 | 0.004945932 | 0.029209347 |
| RAN | 1.457364803 | 0.000838022 | 0.007033817 |
| NDUFV2 | 1.459497742 | 0.007090276 | 0.037490853 |
| NUDT21 | 1.466373261 | 0.005120167 | 0.029880485 |
| EXOC1 | 1.469126607 | 0.005291128 | 0.030669743 |
| PTPLAD1 | 1.471204675 | 0.008959706 | 0.044633245 |
| VBP1 | 1.47143805 | 0.006090955 | 0.033637707 |
| SRSF1 | 1.477438813 | 0.000829882 | 0.006995838 |
| HMGCS1 | 1.478624439 | 0.003753838 | 0.023717126 |
| SLC33A1 | 1.49017638 | 0.003237835 | 0.021094414 |
| MAP2K1 | 1.495182162 | 0.002754462 | 0.018515187 |
| LSM12 | 1.495367137 | 0.00195107 | 0.014030144 |
| APOBEC3D | 1.496366667 | 0.006732097 | 0.036307888 |
| ILF2 | 1.499554989 | 0.006048981 | 0.03347673 |
| MCM5 | 1.501717144 | 0.008652152 | 0.043488446 |
| CDK2AP2 | 1.503326898 | 0.001248863 | 0.009808425 |
| FDFT1 | 1.503924857 | 0.009089884 | 0.045052457 |
| MAGT1 | 1.505242292 | 0.004572431 | 0.027502488 |
| FURIN | 1.506153332 | 0.001663026 | 0.012274416 |
| TUBA1A | 1.512357014 | 0.002318598 | 0.016168246 |
| HIPK2 | 1.512536586 | 0.000101302 | 0.001228017 |
| XBP1 | 1.514170699 | 0.000619865 | 0.0054664 |
| FOXK2 | 1.516311832 | 0.003542931 | 0.022611812 |
| EIF5A | 1.51983914 | 7.96E-05 | 0.001013363 |
| GLUD1 | 1.520254158 | 0.00134943 | 0.010421487 |
| CD48 | 1.522534863 | 0.001288854 | 0.010052804 |
| SEH1L | 1.529753769 | 0.002849273 | 0.019021871 |
| HIGD1A | 1.53323804 | 0.005707834 | 0.032110664 |
| SF3B4 | 1.541392896 | 0.003509287 | 0.022433652 |
| ATP5G3 | 1.543189351 | 0.000357262 | 0.003449717 |
| KDM5B | 1.54560667 | 0.005006901 | 0.02952488 |
| SQSTM1 | 1.559216499 | 0.000127764 | 0.001486655 |
| HAX1 | 1.577757641 | 0.005495537 | 0.031281175 |

| | | | |
|---|---|---|---|
| EIF2S2 | 1.578140016 | 0.000347628 | 0.003370318 |
| TDG | 1.581071947 | 0.005099102 | 0.029835795 |
| COX6A1 | 1.582771851 | 0.001441886 | 0.010977723 |
| SIPA1L1 | 1.583935568 | 0.009635647 | 0.04716047 |
| GBP2 | 1.588531805 | 0.000951577 | 0.007835752 |
| LPXN | 1.588722187 | 0.000362821 | 0.003494771 |
| CAB39 | 1.589531294 | 0.000112075 | 0.001325773 |
| HM13 | 1.589980286 | 0.007393533 | 0.038547772 |
| CD44 | 1.591088609 | 0.005863513 | 0.032767392 |
| SMS | 1.596694184 | 0.00110262 | 0.008810835 |
| HINT1 | 1.598487244 | 5.73E-05 | 0.000782023 |
| KPNB1 | 1.599207945 | 4.51E-05 | 0.000634549 |
| PGS1 | 1.604809536 | 0.001731281 | 0.012682563 |
| GSTP1 | 1.607834954 | 0.005496475 | 0.031281175 |
| NUTF2 | 1.610235919 | 0.006955942 | 0.037150406 |
| PDIA4 | 1.619220853 | 0.0002517 | 0.002600348 |
| ATF4 | 1.623617151 | 0.000193876 | 0.002094125 |
| MIER1 | 1.625600789 | 0.004998135 | 0.029495401 |
| FMNL3 | 1.627728716 | 0.002570598 | 0.017459112 |
| TALDO1 | 1.628306394 | 0.002580378 | 0.017495189 |
| RAB10 | 1.628523041 | 0.009461612 | 0.046482987 |
| LCORL | 1.631166555 | 0.005691837 | 0.032083663 |
| SEC23B | 1.633337699 | 0.004476566 | 0.027049371 |
| TMEM127 | 1.635988687 | 0.007088277 | 0.037490853 |
| TOB2 | 1.636758379 | 0.000364301 | 0.00350042 |
| TMEM33 | 1.640528142 | 0.005797795 | 0.032538984 |
| CCT5 | 1.640691557 | 0.0001905 | 0.002066217 |
| TTC39C | 1.641645838 | 0.002140884 | 0.015117463 |
| ATP13A3 | 1.646895805 | 0.002150863 | 0.015160582 |
| TARBP1 | 1.650650954 | 0.005797112 | 0.032538984 |
| ELAVL1 | 1.651947867 | 0.00103538 | 0.008402275 |
| ANKRD20A9P | 1.657507256 | 0.002970465 | 0.019694295 |
| TIMMDC1 | 1.661857829 | 0.006890274 | 0.036881567 |
| UTP6 | 1.669839044 | 0.000830817 | 0.006995838 |
| TIMM10B | 1.671882253 | 0.009234462 | 0.045653456 |
| FTL | 1.674313279 | 3.19E-05 | 0.00047436 |
| COX7B | 1.675218168 | 0.000137952 | 0.001584016 |
| TBCA | 1.677698845 | 0.00542563 | 0.031013216 |
| FOXN2 | 1.67842244 | 0.004793321 | 0.02852209 |
| ATP6V1B2 | 1.686305002 | 0.006379683 | 0.034863431 |
| TXLNA | 1.686695634 | 0.003071937 | 0.020266506 |
| SNHG16 | 1.69215846 | 3.30E-05 | 0.000485518 |
| THOC7 | 1.692585569 | 0.00076411 | 0.006525352 |
| UBE2I | 1.692975615 | 0.000122385 | 0.001432578 |
| OSTC | 1.696094479 | 0.001082336 | 0.008702023 |
| NIPA2 | 1.697913707 | 0.002786332 | 0.018649375 |
| IER3IP1 | 1.70006168 | 0.004011468 | 0.024892083 |
| RAPGEF6 | 1.701210662 | 0.004375694 | 0.026623303 |
| STK17B | 1.720450195 | 1.21E-05 | 0.000207587 |
| CCT2 | 1.723614344 | 4.08E-05 | 0.000577778 |

| | | | |
|---|---|---|---|
| WDR36 | 1.727477211 | 0.003744478 | 0.023685789 |
| TMEM258 | 1.728322 | 0.000256885 | 0.002639982 |
| REEP5 | 1.729602045 | 0.001253895 | 0.009818082 |
| C10orf32 | 1.735287459 | 0.009049938 | 0.044938777 |
| PDIA6 | 1.737931893 | 4.56E-05 | 0.000640179 |
| BOLA2 | 1.738159912 | 0.006005121 | 0.033328208 |
| ASAP1 | 1.739479476 | 0.001066918 | 0.008595713 |
| PSMD14 | 1.741120992 | 0.001628432 | 0.012121912 |
| SESN2 | 1.745522742 | 0.004378855 | 0.026623303 |
| CHORDC1 | 1.745604302 | 0.000734874 | 0.006303177 |
| FTH1 | 1.748460541 | 1.70E-05 | 0.000278867 |
| TMEM165 | 1.749570603 | 0.003246755 | 0.021117388 |
| PCMT1 | 1.749979682 | 0.002493692 | 0.017025373 |
| YWHAG | 1.751996362 | 0.007125123 | 0.037618334 |
| KATNAL1 | 1.754037952 | 0.006712922 | 0.03622942 |
| HDAC2 | 1.760423286 | 5.67E-05 | 0.000774239 |
| UXS1 | 1.767949869 | 0.006334519 | 0.034689246 |
| SART3 | 1.768057592 | 0.000922213 | 0.007626032 |
| B4GALT5 | 1.77407835 | 0.000342846 | 0.003339335 |
| PCGF5 | 1.777554531 | 0.007266983 | 0.038090857 |
| NHP2L1 | 1.78191311 | 0.004528246 | 0.027298456 |
| HSP90AA1 | 1.782935602 | 1.17E-05 | 0.000202667 |
| TESPA1 | 1.785732651 | 0.004142657 | 0.025604691 |
| PPP1R15B | 1.790102908 | 0.001939317 | 0.01397129 |
| IVNS1ABP | 1.795596777 | 0.00074147 | 0.006345853 |
| MRPS35 | 1.799236477 | 0.007197646 | 0.037854106 |
| HYPK | 1.801072631 | 0.001207184 | 0.009529694 |
| RDX | 1.804998401 | 7.11E-05 | 0.000926584 |
| MXD1 | 1.808143322 | 0.000590216 | 0.00522849 |
| SLC39A8 | 1.808221967 | 0.006596748 | 0.035750236 |
| SARS | 1.808641987 | 0.001128943 | 0.008976841 |
| TMEM209 | 1.810494135 | 0.007904681 | 0.040675783 |
| ATP8B4 | 1.812264492 | 0.001309661 | 0.010194785 |
| PRDM1 | 1.812514761 | 0.001923425 | 0.013890535 |
| DDX55 | 1.813764915 | 0.001672382 | 0.012328804 |
| XIAP | 1.818765394 | 0.005106571 | 0.029842955 |
| ACSL4 | 1.821926851 | 0.005351162 | 0.030857843 |
| KXD1 | 1.824040984 | 0.001378088 | 0.010569839 |
| CAMKK2 | 1.826798855 | 0.008794054 | 0.044056131 |
| BPGM | 1.828414841 | 0.003052759 | 0.02017397 |
| GADD45A | 1.828690944 | 0.008376263 | 0.042483492 |
| VDAC2 | 1.833420733 | 0.000487977 | 0.004479888 |
| PITRM1 | 1.833609888 | 0.009258851 | 0.04574515 |
| STYX | 1.836762056 | 0.007211092 | 0.037899368 |
| NDUFB3 | 1.836896135 | 0.00146501 | 0.011095256 |
| SHFM1 | 1.840227456 | 0.008288252 | 0.042118948 |
| FAM3C | 1.841437213 | 0.006762223 | 0.036395169 |
| RHEB | 1.845771794 | 8.54E-05 | 0.001073287 |
| SLC5A3 | 1.848240557 | 0.002189541 | 0.015405477 |
| LINC01004 | 1.853256525 | 0.002103186 | 0.014891545 |

| | | | |
|---|---|---|---|
| BLOC1S4 | 1.854962983 | 0.002450692 | 0.01679035 |
| DDIT4 | 1.860547892 | 9.70E-05 | 0.001191214 |
| UNC50 | 1.864904796 | 0.001458655 | 0.011057816 |
| C2CD5 | 1.871804269 | 0.006532688 | 0.035649814 |
| MMADHC | 1.875389423 | 0.009081568 | 0.045039746 |
| TMEM14B | 1.878497015 | 0.000728116 | 0.006258916 |
| RPN1 | 1.878558181 | 0.001101038 | 0.008807176 |
| PROSC | 1.890146005 | 0.006120448 | 0.033752977 |
| UBR1 | 1.890635394 | 0.002350415 | 0.016317466 |
| PPP4R1 | 1.894120723 | 0.008509338 | 0.04293597 |
| NADK2 | 1.89442567 | 0.001809411 | 0.013180927 |
| COPB2 | 1.895417622 | 0.004433709 | 0.026852574 |
| EPG5 | 1.897562233 | 4.79E-05 | 0.000666252 |
| GNL3 | 1.900389262 | 0.009096834 | 0.045058383 |
| E2F4 | 1.902477541 | 0.002453283 | 0.01679341 |
| LLPH | 1.903031016 | 0.000317061 | 0.003123147 |
| C19orf10 | 1.90407335 | 0.008434884 | 0.042697853 |
| ZC3H12A | 1.908189456 | 0.009510364 | 0.046693206 |
| DEGS1 | 1.911711416 | 0.004163215 | 0.025670976 |
| POC1B-GALNT4 | 1.917197737 | 0.000446906 | 0.004158174 |
| CCDC47 | 1.926195054 | 0.001595844 | 0.011890629 |
| HIC2 | 1.92640112 | 0.004101906 | 0.025392905 |
| PSMA5 | 1.928276242 | 0.001655801 | 0.012244175 |
| APOBEC3G | 1.931302767 | 0.001701877 | 0.012513991 |
| ZFX | 1.935335232 | 0.009345972 | 0.046088356 |
| GSTO1 | 1.935438532 | 0.000209261 | 0.002220491 |
| MCTS1 | 1.937860013 | 0.006974166 | 0.037170478 |
| SDF2L1 | 1.955672662 | 0.000255043 | 0.002624499 |
| CHD7 | 1.958394677 | 0.001953574 | 0.014035262 |
| ZCCHC10 | 1.964068488 | 0.004194369 | 0.02578187 |
| NUS1 | 1.967964807 | 0.002671222 | 0.01803305 |
| MAP3K5 | 1.9731596 | 0.00828381 | 0.042118948 |
| ERCC6 | 1.974727754 | 0.003856193 | 0.024213103 |
| CHSY1 | 1.975454605 | 0.000210462 | 0.002230215 |
| C11orf98 | 1.976109502 | 0.004780825 | 0.028470449 |
| VPS35 | 1.98578482 | 0.001948717 | 0.014026109 |
| PTGES3 | 1.986558848 | 1.14E-06 | 2.63E-05 |
| JMJD6 | 1.98877953 | 0.009016104 | 0.044809041 |
| SEC61A1 | 1.989746664 | 0.005289298 | 0.030669743 |
| TOR1A | 1.99457291 | 0.004757715 | 0.028360731 |
| CISD2 | 1.998434194 | 0.005639913 | 0.031821178 |
| MDH1 | 2.002410133 | 0.001225762 | 0.0096538 |
| GTPBP8 | 2.003190302 | 0.000608922 | 0.005382018 |
| PSMB3 | 2.003482287 | 0.003982328 | 0.024876885 |
| SRF | 2.004536536 | 1.41E-06 | 3.17E-05 |
| ARF4 | 2.006281588 | 0.000430405 | 0.004041365 |
| COPS3 | 2.011055477 | 0.005860774 | 0.032767392 |
| DNAJB11 | 2.012119398 | 1.99E-05 | 0.000321124 |
| EIF4E | 2.015331006 | 9.39E-06 | 0.000167575 |
| SNRPE | 2.017233569 | 0.000232272 | 0.002434972 |

| | | | |
|---|---|---|---|
| GOSR1 | 2.021708539 | 0.003632593 | 0.023052541 |
| RRP36 | 2.022164437 | 0.004606429 | 0.027635934 |
| FAM98B | 2.028739092 | 0.002640684 | 0.017842278 |
| HSPB1 | 2.031816223 | 0.005318289 | 0.030690877 |
| DCTN4 | 2.033593075 | 0.00507655 | 0.02980095 |
| CALR | 2.038105165 | 1.64E-06 | 3.64E-05 |
| ZNF830 | 2.040139485 | 0.004363529 | 0.026571382 |
| TMEM106B | 2.041306846 | 0.007285105 | 0.038140498 |
| HSPA8 | 2.041370643 | 1.74E-08 | 6.79E-07 |
| NOMO2 | 2.049507766 | 0.000669087 | 0.005802456 |
| NCOA3 | 2.05186872 | 0.000681553 | 0.005897503 |
| PPARA | 2.054282244 | 0.005502649 | 0.031293569 |
| RBM14-RBM4 | 2.056162322 | 0.000203393 | 0.002167033 |
| FKBP2 | 2.061181315 | 8.16E-05 | 0.001035037 |
| NGRN | 2.063299802 | 0.002085489 | 0.014803353 |
| SKIL | 2.065671124 | 1.57E-05 | 0.000259112 |
| NOP10 | 2.068375717 | 0.000145944 | 0.001649183 |
| EIF5B | 2.069222195 | 9.26E-06 | 0.000165859 |
| RCN1 | 2.076289662 | 0.005379541 | 0.030861523 |
| DCTPP1 | 2.079996178 | 0.000347748 | 0.003370318 |
| DNAJB9 | 2.080758571 | 0.000187348 | 0.002049914 |
| HYOU1 | 2.083598937 | 5.25E-05 | 0.000726373 |
| MRPL33 | 2.090633414 | 0.00430345 | 0.026287296 |
| AAAS | 2.093753962 | 0.000636528 | 0.005581916 |
| CCDC6 | 2.097523982 | 0.000207558 | 0.002205406 |
| ZDHHC5 | 2.101685334 | 0.00972504 | 0.047427047 |
| FASTKD2 | 2.102645461 | 0.005856231 | 0.032767392 |
| C15orf57 | 2.104594118 | 0.007133601 | 0.037618334 |
| HINT3 | 2.105135696 | 0.003829662 | 0.024088421 |
| HADH | 2.106994981 | 0.003021915 | 0.019987006 |
| TUBA1C | 2.108241497 | 0.001428709 | 0.010894083 |
| PRPS2 | 2.108325002 | 0.000665336 | 0.005782737 |
| MEX3C | 2.114024291 | 2.91E-06 | 5.98E-05 |
| XPO4 | 2.121169116 | 0.003983605 | 0.024876885 |
| C17orf58 | 2.127178252 | 0.00385876 | 0.024213103 |
| MRPL3 | 2.133266328 | 0.000494471 | 0.004528892 |
| ATP5G1 | 2.13651774 | 0.00147824 | 0.011173837 |
| PDP2 | 2.137544075 | 0.008671371 | 0.043557091 |
| ENO1 | 2.138255934 | 2.61E-08 | 9.57E-07 |
| TAB2 | 2.147262928 | 0.000188246 | 0.002056003 |
| FOPNL | 2.148971069 | 0.00019589 | 0.002112969 |
| MT2A | 2.154852706 | 0.001371233 | 0.010548257 |
| TUBB | 2.158690738 | 3.18E-07 | 8.53E-06 |
| GFPT1 | 2.161835447 | 8.52E-06 | 0.000153744 |
| HSP90AB1 | 2.166050994 | 2.06E-08 | 7.77E-07 |
| PXK | 2.166770936 | 0.003341157 | 0.021516944 |
| HNRNPAB | 2.167678452 | 4.55E-07 | 1.17E-05 |
| NAMPT | 2.170626779 | 0.0010424 | 0.008441609 |
| MAGOH | 2.179090657 | 0.000111698 | 0.001324393 |
| HMGCR | 2.180814911 | 0.000338704 | 0.003311352 |

| | | | |
|---|---|---|---|
| ARCN1 | 2.181878364 | 8.18E-06 | 0.00014828 |
| ACOT13 | 2.183731958 | 0.0001232 | 0.001439974 |
| GCA | 2.186100856 | 0.008873028 | 0.044354369 |
| POLR2K | 2.190051397 | 5.30E-05 | 0.000732589 |
| OPA3 | 2.195239598 | 0.002529248 | 0.01723807 |
| DHRS3 | 2.203295484 | 8.02E-06 | 0.000145945 |
| COX17 | 2.203313749 | 1.94E-06 | 4.21E-05 |
| CCT3 | 2.206903672 | 8.26E-05 | 0.001043593 |
| MARCKSL1 | 2.211684089 | 0.007497719 | 0.039013047 |
| TRMT1L | 2.216670568 | 0.00332825 | 0.02148683 |
| MARS | 2.220331164 | 2.42E-06 | 5.10E-05 |
| GTF2H5 | 2.220841967 | 0.001497328 | 0.011296316 |
| FDPSP2 | 2.228242499 | 0.00571521 | 0.032129081 |
| SLC3A2 | 2.240576158 | 3.22E-06 | 6.56E-05 |
| IL6ST | 2.241949102 | 0.00069178 | 0.0059728 |
| AARS | 2.247931592 | 0.001534904 | 0.011513251 |
| MPC1 | 2.256601656 | 0.00132763 | 0.010303935 |
| PMAIP1 | 2.256633741 | 0.000717362 | 0.006173254 |
| LSM6 | 2.260144059 | 0.000118441 | 0.001396477 |
| NDUFB4 | 2.262271473 | 0.000310138 | 0.003074292 |
| STAMBP | 2.268439263 | 0.001748603 | 0.012773612 |
| RHOH | 2.272330109 | 0.000530461 | 0.00480236 |
| EYA3 | 2.276751753 | 0.005187519 | 0.030203316 |
| PSMA3 | 2.288908429 | 0.000143448 | 0.001628034 |
| SMG5 | 2.289326404 | 0.001456851 | 0.011054847 |
| LCP2 | 2.29616455 | 1.76E-09 | 9.13E-08 |
| DNAJA1 | 2.297637728 | 0.001505861 | 0.011338841 |
| KATNA1 | 2.303228269 | 0.009552969 | 0.046843647 |
| ANKRD27 | 2.303741718 | 0.00930863 | 0.045933132 |
| CCNJ | 2.308513394 | 6.86E-05 | 0.000902557 |
| PSMD11 | 2.311918947 | 0.000341147 | 0.003326924 |
| TMEM68 | 2.312421629 | 0.000279936 | 0.002825888 |
| LTB | 2.314799948 | 5.60E-08 | 1.81E-06 |
| MCCC2 | 2.31517407 | 0.000104653 | 0.001266676 |
| BRCA1 | 2.316507045 | 0.004407574 | 0.026735641 |
| KLHL9 | 2.317668722 | 0.004072342 | 0.025249811 |
| PTGER4 | 2.32010479 | 0.000297798 | 0.002974562 |
| SS18L2 | 2.322601282 | 0.003462725 | 0.022190342 |
| LMAN1 | 2.323748741 | 6.84E-05 | 0.000901507 |
| ACER3 | 2.324652443 | 0.007968722 | 0.040901177 |
| CNIH1 | 2.327530286 | 0.000775855 | 0.006604041 |
| PRDX4 | 2.337002145 | 0.009112321 | 0.045106565 |
| GOLGA8J | 2.347950489 | 0.005380988 | 0.030861523 |
| EBPL | 2.350567148 | 0.005270247 | 0.030593999 |
| ZNF445 | 2.356923878 | 0.008154373 | 0.041600581 |
| SOD2 | 2.360235679 | 0.007286194 | 0.038140498 |
| ECHDC1 | 2.363654491 | 0.000281422 | 0.00283267 |
| SLAMF7 | 2.370701326 | 1.88E-06 | 4.10E-05 |
| HDGFRP3 | 2.372450565 | 2.83E-05 | 0.000429125 |
| MAT2A | 2.381850761 | 1.45E-08 | 5.79E-07 |

| | | | |
|---|---|---|---|
| AK6 | 2.393844406 | 3.22E-05 | 0.000476716 |
| LSMEM2 | 2.398600099 | 0.002590068 | 0.017545696 |
| B3GALT6 | 2.39919187 | 0.005638821 | 0.031821178 |
| SRFBP1 | 2.400825298 | 0.009647671 | 0.047189826 |
| NDC1 | 2.40089959 | 0.002759134 | 0.018530688 |
| PTS | 2.409122497 | 0.005313924 | 0.030688306 |
| TXNRD1 | 2.409874679 | 4.01E-05 | 0.000571323 |
| KIF27 | 2.410943489 | 0.005940256 | 0.033085451 |
| PIP5K1C | 2.411836466 | 2.08E-05 | 0.000331704 |
| SURF4 | 2.414103238 | 3.55E-06 | 7.13E-05 |
| NIFK | 2.420072832 | 0.000288616 | 0.002893922 |
| NXT1 | 2.423748457 | 0.003849515 | 0.024193863 |
| ST8SIA4 | 2.428787615 | 0.00063359 | 0.005568622 |
| CEP162 | 2.431403663 | 0.009202407 | 0.04552372 |
| ESF1 | 2.431526854 | 4.13E-06 | 8.19E-05 |
| TCOF1 | 2.432244797 | 0.00741702 | 0.038618805 |
| KCTD5 | 2.440858508 | 0.001652347 | 0.012241752 |
| PIGP | 2.442953664 | 0.002111997 | 0.014940423 |
| GCLM | 2.445795111 | 0.00019289 | 0.002086353 |
| KIAA0020 | 2.445869164 | 0.004853281 | 0.028792455 |
| HSPH1 | 2.447698765 | 1.75E-07 | 5.08E-06 |
| FDX1 | 2.450438687 | 3.24E-07 | 8.63E-06 |
| AGO2 | 2.451048076 | 0.000509339 | 0.004632557 |
| CSE1L | 2.459964559 | 0.006588615 | 0.035750236 |
| HOXB2 | 2.460314698 | 0.006261221 | 0.034335872 |
| RRAS2 | 2.461975944 | 0.000189473 | 0.002058203 |
| PSMC6 | 2.469789437 | 1.47E-05 | 0.000243316 |
| IER5 | 2.470610949 | 6.20E-07 | 1.54E-05 |
| NUP188 | 2.474529243 | 0.002053782 | 0.014607782 |
| ANKS1A | 2.477061978 | 0.00899815 | 0.044767799 |
| SRGAP2 | 2.478950704 | 0.001003931 | 0.008197897 |
| SEC61B | 2.480954025 | 2.52E-10 | 1.69E-08 |
| CDCA4 | 2.483958141 | 3.58E-05 | 0.000521283 |
| ZNF25 | 2.490438394 | 0.006605873 | 0.035750236 |
| IPO5 | 2.494058933 | 2.31E-05 | 0.0003632 |
| IL18R1 | 2.49896733 | 6.03E-07 | 1.50E-05 |
| HSPE1 | 2.49933445 | 5.57E-07 | 1.41E-05 |
| ZNF2 | 2.504937724 | 0.00859791 | 0.043299185 |
| EBNA1BP2 | 2.518315529 | 0.00054153 | 0.004891261 |
| TNFSF10 | 2.519506715 | 0.000146543 | 0.001653572 |
| POLR2I | 2.521978123 | 0.004396262 | 0.026708402 |
| NUDC | 2.524276625 | 0.000334226 | 0.003271651 |
| TMEM39A | 2.524447566 | 0.002771474 | 0.01858169 |
| RNF145 | 2.533855369 | 0.000265944 | 0.002711725 |
| NFKBIA | 2.540103621 | 1.10E-11 | 9.80E-10 |
| ANKRD32 | 2.548518955 | 0.000140032 | 0.001603199 |
| PGAP1 | 2.552212765 | 0.005471252 | 0.031228407 |
| HTRA2 | 2.555196441 | 0.004939068 | 0.029190824 |
| SMIM19 | 2.557010693 | 0.00378362 | 0.023856304 |
| USP45 | 2.560287079 | 0.00448512 | 0.027080164 |

| | | | |
|---|---|---|---|
| NUDCD1 | 2.563475612 | 0.000784779 | 0.006670518 |
| PLGRKT | 2.564649042 | 0.001747195 | 0.012773612 |
| C19orf60 | 2.569172418 | 0.009835551 | 0.047839876 |
| TMEM167A | 2.569382322 | 0.000139006 | 0.001593791 |
| IRGQ | 2.569505306 | 0.000937012 | 0.007740234 |
| HRSP12 | 2.570231274 | 0.007222159 | 0.037932078 |
| GADD45B | 2.575219862 | 1.13E-06 | 2.62E-05 |
| STXBP3 | 2.578847307 | 0.001592574 | 0.011877567 |
| MAP3K8 | 2.580922407 | 7.63E-10 | 4.56E-08 |
| SQLE | 2.590037376 | 0.000197598 | 0.002122615 |
| NFIA | 2.590331108 | 0.000670075 | 0.005804597 |
| SFXN1 | 2.599483724 | 4.36E-05 | 0.000614399 |
| ZNF585A | 2.599785775 | 0.005873723 | 0.032784835 |
| BTG2 | 2.601178154 | 4.33E-06 | 8.56E-05 |
| TXNDC17 | 2.609142998 | 0.004086475 | 0.025317395 |
| C11orf84 | 2.610270674 | 0.002463113 | 0.016831275 |
| WDR89 | 2.612116597 | 0.000626563 | 0.005513052 |
| PRKCQ-AS1 | 2.616023354 | 4.60E-05 | 0.00064329 |
| GTPBP10 | 2.621063643 | 0.001279275 | 0.009988037 |
| SLC25A22 | 2.624798822 | 0.000281033 | 0.002832395 |
| ZNRF2 | 2.629317749 | 0.004243714 | 0.02600354 |
| SEC61G | 2.635899523 | 8.26E-09 | 3.59E-07 |
| SNRPG | 2.640236584 | 4.19E-09 | 1.92E-07 |
| RFC3 | 2.642906156 | 0.008056318 | 0.041207724 |
| CFAP20 | 2.649314893 | 0.004008796 | 0.024892083 |
| DDX1 | 2.654706498 | 0.000439387 | 0.004101117 |
| HAUS6 | 2.654727372 | 8.81E-05 | 0.001096742 |
| GZMB | 2.662427917 | 2.48E-09 | 1.21E-07 |
| USP37 | 2.665421362 | 0.002050717 | 0.014599239 |
| GTF2H3 | 2.672800663 | 0.000159821 | 0.001785387 |
| RPP14 | 2.680650411 | 0.000247692 | 0.002569105 |
| PFKFB3 | 2.681868668 | 0.000203236 | 0.002167033 |
| SPPL3 | 2.685110013 | 0.008618852 | 0.043348897 |
| SBNO2 | 2.689138666 | 1.74E-07 | 5.06E-06 |
| PAK1IP1 | 2.689319025 | 9.95E-05 | 0.001211798 |
| ARSB | 2.694165684 | 0.000477581 | 0.004399927 |
| TVP23B | 2.695644256 | 0.006977474 | 0.037170478 |
| FASLG | 2.696695142 | 5.54E-09 | 2.48E-07 |
| LINC00472 | 2.700305376 | 0.002731532 | 0.01840846 |
| PGM2L1 | 2.701417364 | 0.009353696 | 0.046097417 |
| BCOR | 2.702524169 | 0.006821846 | 0.036617966 |
| UPP1 | 2.715888609 | 7.51E-05 | 0.000965045 |
| C7orf60 | 2.722568145 | 0.008611552 | 0.043340015 |
| DUSP5 | 2.723764933 | 3.32E-05 | 0.000487572 |
| TXNDC15 | 2.724346519 | 3.94E-09 | 1.85E-07 |
| ALKBH8 | 2.728092109 | 0.00580059 | 0.032538984 |
| QTRTD1 | 2.73612575 | 0.000355545 | 0.003437375 |
| SEC24D | 2.736875362 | 0.000795575 | 0.006749893 |
| ASNS | 2.737016988 | 0.00223003 | 0.015634169 |
| FBXO28 | 2.73830855 | 0.000396507 | 0.003759132 |

| | | | |
|---|---|---|---|
| PMM2 | 2.741372715 | 0.002424423 | 0.016654084 |
| TUBA1B | 2.744263107 | 3.13E-13 | 3.95E-11 |
| UBE2Q2P1 | 2.747144332 | 0.007305043 | 0.038189355 |
| ATAD3A | 2.751890951 | 0.005120625 | 0.029880485 |
| ADAM9 | 2.754998883 | 5.96E-05 | 0.000807124 |
| ZNF267 | 2.763137697 | 7.84E-08 | 2.46E-06 |
| PDDC1 | 2.763516977 | 0.003113704 | 0.020490266 |
| SRM | 2.765274333 | 0.000175739 | 0.001938325 |
| FAM129A | 2.765860041 | 1.24E-05 | 0.000210463 |
| SNIP1 | 2.766558219 | 0.007603532 | 0.039477032 |
| RASGEF1B | 2.780097556 | 0.003341076 | 0.021516944 |
| HAT1 | 2.786579287 | 0.007344343 | 0.038367943 |
| BCL2L11 | 2.788916916 | 0.001721525 | 0.012634736 |
| LDLR | 2.789929072 | 0.000143172 | 0.001627256 |
| PDE8A | 2.791564236 | 0.005063452 | 0.029746357 |
| BCL2L1 | 2.794917947 | 0.000472034 | 0.004364228 |
| BCL2A1 | 2.798330692 | 0.000107954 | 0.00129661 |
| TUBGCP4 | 2.798984774 | 0.004938388 | 0.029190824 |
| ZNF268 | 2.809447499 | 0.006021899 | 0.033374021 |
| PIM2 | 2.81043832 | 2.95E-08 | 1.07E-06 |
| DLG2 | 2.810815383 | 0.005621171 | 0.031821178 |
| DNAJA3 | 2.814400388 | 3.16E-05 | 0.000470317 |
| VIMP | 2.815676343 | 2.18E-05 | 0.000343141 |
| KIAA1958 | 2.822698624 | 0.005093377 | 0.029835795 |
| PKM | 2.824458973 | 1.05E-10 | 7.51E-09 |
| PYGO2 | 2.826320378 | 0.007679078 | 0.039771734 |
| GCNT1 | 2.830532115 | 0.00322146 | 0.021040244 |
| RCC1 | 2.841914337 | 0.000245433 | 0.002549048 |
| GK | 2.844024062 | 0.003910882 | 0.02448131 |
| C1orf131 | 2.844146072 | 1.37E-05 | 0.000229034 |
| DDX21 | 2.845043715 | 3.48E-11 | 2.78E-09 |
| PCNA | 2.847286232 | 2.39E-08 | 8.92E-07 |
| NFKBIZ | 2.853213481 | 5.34E-11 | 4.10E-09 |
| METTL2A | 2.863829534 | 0.006135742 | 0.033813506 |
| MPP5 | 2.865951267 | 0.007411162 | 0.038613978 |
| HAVCR2 | 2.869402982 | 3.04E-07 | 8.20E-06 |
| IRF4 | 2.872013586 | 0.005818676 | 0.032617071 |
| PCDH11Y | 2.881655116 | 0.007970272 | 0.040901177 |
| TCEB1 | 2.8841464 | 2.48E-06 | 5.18E-05 |
| SLC4A10 | 2.896680073 | 0.004175663 | 0.025706384 |
| WWC2 | 2.896680401 | 0.000155758 | 0.001747476 |
| SPATS2 | 2.901934525 | 0.000529868 | 0.00480236 |
| NMNAT1 | 2.906398313 | 0.003143632 | 0.020647308 |
| CTPS1 | 2.916657123 | 0.004216674 | 0.025878352 |
| ING5 | 2.918755437 | 0.000221815 | 0.002337865 |
| COA4 | 2.931447838 | 0.00100971 | 0.008227928 |
| PCDH7 | 2.933994543 | 0.006564438 | 0.035698691 |
| RBPJ | 2.936526459 | 1.37E-11 | 1.18E-09 |
| FEZ1 | 2.93775787 | 3.55E-10 | 2.26E-08 |
| HPS6 | 2.942059324 | 0.003515782 | 0.022456842 |

| | | | |
|---|---|---|---|
| FASN | 2.942797286 | 3.22E-07 | 8.61E-06 |
| FGFBP3 | 2.95344786 | 0.005101536 | 0.029835795 |
| SHMT2 | 2.958614224 | 0.000310982 | 0.003078758 |
| TARS | 2.975041554 | 6.08E-09 | 2.71E-07 |
| MCM2 | 2.978472482 | 8.74E-05 | 0.001091407 |
| MEGF8 | 2.981515701 | 0.004407395 | 0.026735641 |
| TBC1D24 | 2.984718342 | 0.001443477 | 0.010977723 |
| VIM | 2.995497905 | 5.45E-14 | 7.75E-12 |
| PAICS | 2.999908238 | 2.42E-07 | 6.87E-06 |
| ADCY10P1 | 3.000908973 | 0.000241832 | 0.002518377 |
| THAP5 | 3.001850373 | 2.13E-05 | 0.000338427 |
| LACTB | 3.01328598 | 0.000351383 | 0.003401332 |
| BCL9 | 3.013372072 | 0.009375301 | 0.046174829 |
| STIP1 | 3.014640702 | 3.70E-05 | 0.000535608 |
| CSTF2T | 3.017728468 | 4.52E-06 | 8.90E-05 |
| CCND2 | 3.0228689 | 2.77E-16 | 5.56E-14 |
| STAT5A | 3.0244473 | 1.19E-11 | 1.05E-09 |
| MTFP1 | 3.024599052 | 0.005037246 | 0.029659151 |
| SLC1A4 | 3.025191918 | 3.72E-06 | 7.42E-05 |
| HSPD1 | 3.026595928 | 1.09E-12 | 1.22E-10 |
| MAK16 | 3.030383396 | 0.000140709 | 0.001606261 |
| LINC01531 | 3.030499371 | 0.004911532 | 0.029115978 |
| CEBPB | 3.0481017 | 9.14E-10 | 5.38E-08 |
| SHISA9 | 3.050029642 | 0.008214061 | 0.041877809 |
| ADRM1 | 3.067029288 | 0.000250644 | 0.002592857 |
| MRPL18 | 3.068482408 | 0.00010913 | 0.001308728 |
| TWSG1 | 3.070234639 | 0.00091396 | 0.007570873 |
| SQRDL | 3.083458417 | 2.22E-07 | 6.32E-06 |
| IL4R | 3.086542016 | 0.000369044 | 0.003532988 |
| TMEM243 | 3.088317542 | 0.002385006 | 0.016484542 |
| CLDND1 | 3.093950036 | 9.31E-09 | 4.01E-07 |
| ZNF557 | 3.094288418 | 0.000378796 | 0.003608696 |
| loc468 | 3.096435159 | 0.001319473 | 0.010260969 |
| LSM2 | 3.098932199 | 0.000912488 | 0.007569592 |
| MRPL41 | 3.100468108 | 0.000192593 | 0.00208603 |
| ZNF124 | 3.102464208 | 0.004660727 | 0.027925137 |
| BOLA1 | 3.131534707 | 0.008569586 | 0.043212126 |
| PDCD5 | 3.141014808 | 2.13E-05 | 0.000338427 |
| ZNF578 | 3.145968618 | 0.006682285 | 0.036113852 |
| GLO1 | 3.147816436 | 7.65E-07 | 1.84E-05 |
| VKORC1L1 | 3.148964145 | 0.000420437 | 0.003957265 |
| LRP8 | 3.151203542 | 2.08E-06 | 4.44E-05 |
| CMSS1 | 3.15227185 | 0.001905941 | 0.013807052 |
| IARS | 3.154105078 | 5.02E-09 | 2.28E-07 |
| ADO | 3.154181184 | 0.000152858 | 0.001719868 |
| CSMD3 | 3.155924853 | 0.004178652 | 0.025706384 |
| SGMS1 | 3.157973439 | 0.00058258 | 0.005172541 |
| DPAGT1 | 3.164959973 | 0.001362384 | 0.010500813 |
| SVIP | 3.168907621 | 5.17E-06 | 9.94E-05 |
| IL15RA | 3.170907137 | 0.001854574 | 0.013472326 |

| | | | |
|---|---|---|---|
| FAIM | 3.172523017 | 0.00058141 | 0.005168012 |
| DLAT | 3.173010965 | 0.000249923 | 0.002588822 |
| RPF2 | 3.174886937 | 1.63E-08 | 6.36E-07 |
| SENP3-EIF4A1 | 3.181037686 | 0.007735497 | 0.039984604 |
| VMP1 | 3.181084589 | 1.01E-07 | 3.08E-06 |
| SLC39A1 | 3.182916711 | 0.000124069 | 0.001445815 |
| BET1 | 3.183176593 | 0.00530613 | 0.030688306 |
| FBXO30 | 3.184176915 | 0.004354253 | 0.026535532 |
| EXPH5 | 3.19485973 | 0.000904568 | 0.007527812 |
| HIST1H2BK | 3.202990717 | 0.001022509 | 0.008314919 |
| URB1 | 3.204128142 | 6.52E-05 | 0.000867911 |
| ANK3 | 3.219337435 | 1.60E-06 | 3.56E-05 |
| YIF1A | 3.225355366 | 0.000543531 | 0.004903675 |
| FAM186A | 3.226292383 | 0.006039544 | 0.033448138 |
| NABP1 | 3.22826466 | 8.19E-05 | 0.001036269 |
| GOT1 | 3.232897791 | 3.23E-05 | 0.000477908 |
| RSPH4A | 3.252726321 | 0.001680128 | 0.012365681 |
| PRDX1 | 3.25388299 | 0.000657311 | 0.005726744 |
| ZNF714 | 3.254547986 | 0.000771811 | 0.006576772 |
| CNTN5 | 3.255130552 | 0.000281954 | 0.002834379 |
| FOXP4 | 3.263712047 | 0.00849541 | 0.042921004 |
| COX16 | 3.2690611 | 7.08E-05 | 0.000925435 |
| NLN | 3.270704908 | 0.007726094 | 0.039962379 |
| EMX2OS | 3.273278143 | 0.003624711 | 0.023039866 |
| BOLA3-AS1 | 3.27508052 | 0.003135229 | 0.020614592 |
| FAM114A2 | 3.27528356 | 6.54E-08 | 2.08E-06 |
| HNRNPLL | 3.277123196 | 0.001121685 | 0.008944923 |
| LAGE3 | 3.277795372 | 0.001047186 | 0.008471604 |
| SEMA3A | 3.288472098 | 0.008930418 | 0.0445505 |
| TIFA | 3.298148137 | 0.000211488 | 0.002238053 |
| LOC439994 | 3.299492732 | 0.006352844 | 0.034741009 |
| TANK | 3.299808618 | 0.000685117 | 0.005921803 |
| SRXN1 | 3.307716495 | 3.07E-09 | 1.48E-07 |
| TMEM208 | 3.330495957 | 5.06E-05 | 0.00070182 |
| RBM28 | 3.332243789 | 0.000254868 | 0.002624499 |
| TEX30 | 3.348509934 | 0.000563403 | 0.005059641 |
| ALAS1 | 3.34923821 | 0.003095746 | 0.020406387 |
| PDE4C | 3.358443241 | 0.002341442 | 0.016269591 |
| loc920 | 3.359558514 | 5.09E-06 | 9.81E-05 |
| TUBG1 | 3.372018181 | 0.000603958 | 0.005344176 |
| IRF8 | 3.380012865 | 1.33E-08 | 5.44E-07 |
| SLC7A1 | 3.3855287 | 4.70E-07 | 1.20E-05 |
| PPM1D | 3.387715475 | 0.000363769 | 0.003499596 |
| MEFV | 3.392670522 | 0.004302939 | 0.026287296 |
| TRDMT1 | 3.398065376 | 0.006605345 | 0.035750236 |
| BIRC3 | 3.399435126 | 2.60E-13 | 3.33E-11 |
| SRD5A3 | 3.417822064 | 0.003374646 | 0.021714754 |
| BHLHE40 | 3.418002417 | 4.28E-08 | 1.46E-06 |
| SI | 3.426902398 | 0.001292143 | 0.010068429 |
| EPHA3 | 3.431050729 | 0.009666742 | 0.047253591 |

| | | | |
|---|---|---|---|
| MDFIC | 3.431823858 | 5.52E-12 | 5.47E-10 |
| CD274 | 3.445786431 | 0.000494092 | 0.004528892 |
| NKIRAS1 | 3.455737844 | 0.001340203 | 0.010370685 |
| SLC31A1 | 3.456859585 | 8.89E-07 | 2.10E-05 |
| MINOS1P1 | 3.468569231 | 0.002150514 | 0.015160582 |
| KDM6B | 3.471515673 | 9.98E-10 | 5.83E-08 |
| NFKB1 | 3.472787405 | 3.66E-08 | 1.28E-06 |
| PIH1D2 | 3.482932925 | 0.007029106 | 0.037318597 |
| EPHA1-AS1 | 3.491549681 | 0.000830802 | 0.006995838 |
| MYH15 | 3.493550288 | 0.008274589 | 0.042104165 |
| TNFAIP8 | 3.508958522 | 8.97E-12 | 8.07E-10 |
| SLC35F2 | 3.520571831 | 0.007193508 | 0.037854106 |
| CPM | 3.521992878 | 0.002736812 | 0.01841335 |
| PDSS1 | 3.526409341 | 0.003160648 | 0.020712165 |
| IRAK2 | 3.530240322 | 5.63E-06 | 0.000107183 |
| PRRG4 | 3.536312194 | 0.005706391 | 0.032110664 |
| RCL1 | 3.570344672 | 0.000438927 | 0.004101117 |
| KCNA3 | 3.581932281 | 2.04E-08 | 7.74E-07 |
| URB2 | 3.582054381 | 0.002811218 | 0.018799873 |
| AACS | 3.596482541 | 0.004343985 | 0.026493571 |
| RFC5 | 3.599329451 | 9.83E-05 | 0.001199176 |
| FUT4 | 3.605040771 | 0.000416647 | 0.00392631 |
| SHARPIN | 3.613465573 | 0.000640506 | 0.005610519 |
| PPP1R14B | 3.614780348 | 0.000569826 | 0.005088151 |
| LOC100335030 | 3.615484378 | 0.009747275 | 0.04749901 |
| RAB3IP | 3.615825163 | 7.70E-07 | 1.85E-05 |
| SC5D | 3.620349721 | 2.63E-06 | 5.43E-05 |
| HMCN1 | 3.627455365 | 0.002428051 | 0.016664391 |
| F3 | 3.630417987 | 0.004451612 | 0.02691936 |
| CDH6 | 3.637025737 | 0.00110061 | 0.008807176 |
| FABP5 | 3.639047478 | 8.08E-07 | 1.92E-05 |
| RAD54B | 3.639742871 | 2.96E-05 | 0.000444302 |
| ATCAY | 3.649752138 | 0.009017884 | 0.044809041 |
| ENKD1 | 3.651103043 | 0.002208937 | 0.015500166 |
| SLC4A5 | 3.651190455 | 4.64E-05 | 0.000647358 |
| RALA | 3.653430785 | 2.65E-14 | 4.15E-12 |
| SLC9B1 | 3.654892832 | 0.002736961 | 0.01841335 |
| NSDHL | 3.65583676 | 0.001204151 | 0.009515346 |
| WDFY3 | 3.660715277 | 0.000765944 | 0.006533888 |
| COQ3 | 3.662492509 | 0.000495938 | 0.004537019 |
| CDK6 | 3.664515772 | 4.00E-24 | 2.61E-21 |
| LRP12 | 3.683698297 | 0.004925534 | 0.029176085 |
| TPTE2P6 | 3.684779601 | 0.008952829 | 0.044627373 |
| ZNF681 | 3.686524343 | 0.002375507 | 0.016433391 |
| RMI1 | 3.695066092 | 0.00028566 | 0.002867949 |
| LINC01560 | 3.702017654 | 0.000123956 | 0.001445815 |
| LMO3 | 3.703139399 | 0.005383328 | 0.030861523 |
| CD109 | 3.709092962 | 7.37E-05 | 0.000950349 |
| L2HGDH | 3.709997944 | 0.003568136 | 0.022717131 |
| IRAK1BP1 | 3.712114953 | 0.002012475 | 0.014392412 |

| | | | |
|---|---|---|---|
| DNAJA4 | 3.713822806 | 0.000522275 | 0.004744706 |
| HK2 | 3.7236201 | 0.005300191 | 0.030688306 |
| LINC01296 | 3.729181902 | 0.002250195 | 0.015719251 |
| PIM1 | 3.73905024 | 1.56E-17 | 4.54E-15 |
| GLRX2 | 3.746064713 | 1.49E-07 | 4.38E-06 |
| DCAF12 | 3.750004682 | 7.11E-05 | 0.000926584 |
| HS3ST3B1 | 3.753430549 | 2.61E-07 | 7.26E-06 |
| ZNF790 | 3.760026164 | 0.002137433 | 0.015106714 |
| C6orf203 | 3.780904216 | 0.001443884 | 0.010977723 |
| MAFG | 3.783913239 | 5.52E-13 | 6.65E-11 |
| FAM210A | 3.786649899 | 1.95E-05 | 0.000315134 |
| UBXN2A | 3.788174249 | 0.003297809 | 0.021360745 |
| CCL3L1 | 3.796286859 | 0.002316419 | 0.016167451 |
| MACC1 | 3.796888729 | 0.000170668 | 0.001887712 |
| HIF1A-AS2 | 3.801209865 | 4.02E-10 | 2.54E-08 |
| ZNF490 | 3.819575731 | 0.000560742 | 0.005047324 |
| XCL1 | 3.821782551 | 8.29E-05 | 0.001044935 |
| WDR3 | 3.824972048 | 3.70E-07 | 9.69E-06 |
| APITD1 | 3.826136863 | 0.005580027 | 0.031641703 |
| BUB1B | 3.826187451 | 0.001254999 | 0.009818082 |
| LRRC8B | 3.827614802 | 3.05E-05 | 0.000455332 |
| DOCK9 | 3.831758059 | 6.29E-05 | 0.000845037 |
| NETO2 | 3.837718524 | 0.003555505 | 0.022655133 |
| TNFRSF4 | 3.838304416 | 0.001389437 | 0.010636054 |
| HSPA1A | 3.847116885 | 2.15E-14 | 3.44E-12 |
| FLT1 | 3.860430526 | 0.005480043 | 0.031251096 |
| STX11 | 3.860682659 | 7.09E-17 | 1.63E-14 |
| GFOD2 | 3.86130757 | 9.77E-05 | 0.001195611 |
| MANF | 3.873543858 | 2.86E-15 | 4.98E-13 |
| FRMD4B | 3.878339736 | 1.45E-08 | 5.79E-07 |
| PPIP5K1 | 3.882793071 | 0.000854083 | 0.007160946 |
| TNFRSF18 | 3.886580371 | 5.14E-08 | 1.69E-06 |
| FGB | 3.890519522 | 0.006805082 | 0.036600687 |
| MTHFD2 | 3.893517324 | 2.79E-17 | 7.80E-15 |
| SYNPO2 | 3.898567129 | 0.0030666 | 0.020248353 |
| ZBTB8A | 3.902198295 | 0.000435514 | 0.004074682 |
| ZNRD1 | 3.905423281 | 3.27E-05 | 0.000482982 |
| RFPL1S | 3.914005446 | 0.002206445 | 0.015500166 |
| PCAT7 | 3.92136809 | 0.00165566 | 0.012244175 |
| ZCCHC4 | 3.922788713 | 0.000297742 | 0.002974562 |
| HIVEP2 | 3.927428367 | 2.76E-05 | 0.000421022 |
| RAB3GAP1 | 3.928166608 | 0.000237674 | 0.002488274 |
| ZNF234 | 3.941650447 | 0.000184209 | 0.002023202 |
| DAND5 | 3.945492751 | 0.002391165 | 0.016497987 |
| EPB41L5 | 3.971993749 | 2.56E-05 | 0.000395477 |
| SEMA5A | 3.977768345 | 0.006549086 | 0.035689557 |
| MCF2 | 3.97882837 | 0.000567561 | 0.005078783 |
| WARS | 3.981705993 | 3.63E-07 | 9.54E-06 |
| FAS | 3.98270325 | 6.87E-05 | 0.000902557 |
| MED12L | 4.009829211 | 0.00319786 | 0.020903539 |

| | | | |
|---|---|---|---|
| KGFLP2 | 4.012159857 | 0.000260935 | 0.002670326 |
| TUBB4B | 4.013681711 | 5.04E-08 | 1.67E-06 |
| ARHGAP31 | 4.014244468 | 8.60E-05 | 0.001077578 |
| POLR3D | 4.015757939 | 3.80E-05 | 0.000546876 |
| HTATSF1P2 | 4.021514006 | 0.000579387 | 0.005155885 |
| C1orf109 | 4.03400658 | 0.000412917 | 0.003895848 |
| RNF144A | 4.044391096 | 4.81E-06 | 9.37E-05 |
| IMPDH2 | 4.057282546 | 3.98E-08 | 1.37E-06 |
| TSHZ2 | 4.082084018 | 8.94E-06 | 0.000160492 |
| AK4 | 4.08318276 | 0.00036962 | 0.003534181 |
| RPS10-NUDT3 | 4.08882216 | 0.000259383 | 0.002662164 |
| PRDM8 | 4.10021732 | 0.000914576 | 0.007570873 |
| LOC403312 | 4.113345243 | 0.005640122 | 0.031821178 |
| FASTKD3 | 4.115667614 | 6.37E-05 | 0.000851768 |
| DNPH1 | 4.125483521 | 0.000668579 | 0.005802456 |
| CMAHP | 4.131635062 | 4.52E-15 | 7.53E-13 |
| USP54 | 4.132968432 | 0.00073816 | 0.006324432 |
| PDE10A | 4.134230869 | 0.00203747 | 0.014531352 |
| LRR1 | 4.143473598 | 0.000185822 | 0.002038055 |
| MSMO1 | 4.144428657 | 6.69E-05 | 0.000885102 |
| ALDH1B1 | 4.197032072 | 6.29E-05 | 0.000845037 |
| LOC440300 | 4.204725341 | 0.000476277 | 0.004393082 |
| CREB5 | 4.213101265 | 0.002337076 | 0.016254178 |
| PSMC3IP | 4.238064526 | 0.000860236 | 0.007189443 |
| GHRLOS | 4.24761163 | 0.000110854 | 0.001324244 |
| BCAT1 | 4.264956738 | 0.001673543 | 0.012328804 |
| PPIF | 4.268146849 | 4.82E-08 | 1.63E-06 |
| B3GNT5 | 4.274478626 | 3.28E-05 | 0.000483203 |
| MBL2 | 4.287834969 | 0.001013199 | 0.008247776 |
| MUC15 | 4.301136838 | 0.000643478 | 0.005630256 |
| CTCFL | 4.302428556 | 0.004521565 | 0.02727918 |
| BCL11A | 4.312855456 | 6.31E-05 | 0.000846404 |
| NYAP2 | 4.337099646 | 0.000785368 | 0.006670518 |
| SKIDA1 | 4.337161249 | 0.000203314 | 0.002167033 |
| LRRC8C | 4.346590483 | 9.34E-11 | 6.77E-09 |
| PPAN | 4.362150472 | 0.002573968 | 0.017466848 |
| MRPL24 | 4.362989178 | 1.02E-05 | 0.000179227 |
| RPS6KA6 | 4.372038473 | 0.000798062 | 0.00676366 |
| BATF | 4.398996719 | 9.73E-06 | 0.000172813 |
| LIG4 | 4.401269687 | 7.64E-05 | 0.000977468 |
| TXN | 4.405908654 | 4.64E-17 | 1.17E-14 |
| DNM3OS | 4.408747255 | 1.26E-05 | 0.000214221 |
| TSIX | 4.410497458 | 0.001129126 | 0.008976841 |
| C21orf91 | 4.416696827 | 5.66E-13 | 6.71E-11 |
| DTNA | 4.417545436 | 9.35E-06 | 0.000167231 |
| NPIPB3 | 4.419656085 | 0.000161897 | 0.001806001 |
| CYTIP | 4.425480406 | 1.65E-25 | 1.18E-22 |
| EBF1 | 4.427610276 | 4.19E-05 | 0.00059207 |
| EDN1 | 4.448726171 | 0.000866767 | 0.007236307 |
| CCL4 | 4.465246285 | 1.42E-27 | 1.59E-24 |

| | | | |
|---|---|---|---|
| NBEA | 4.472438474 | 0.000275284 | 0.0027852 |
| SLC7A11 | 4.475044577 | 1.44E-05 | 0.000239693 |
| PARD6B | 4.496794008 | 7.70E-05 | 0.00098428 |
| TMCC1-AS1 | 4.496912888 | 0.00047472 | 0.004383885 |
| KCNK6 | 4.505706838 | 0.00035989 | 0.003470813 |
| GAS2L3 | 4.506240835 | 0.000137276 | 0.001578575 |
| CNTNAP3B | 4.509848891 | 2.52E-05 | 0.000391207 |
| LINC00965 | 4.535095615 | 9.02E-05 | 0.001119904 |
| GPR171 | 4.554209976 | 1.01E-22 | 5.67E-20 |
| LOC101929567 | 4.571302192 | 9.81E-05 | 0.00119857 |
| TRAF3 | 4.576102644 | 2.14E-08 | 8.00E-07 |
| KGFLP1 | 4.578313671 | 0.000500662 | 0.004572457 |
| KPTN | 4.580231819 | 0.000166567 | 0.001850198 |
| PPFIA2 | 4.596692253 | 0.000568128 | 0.005078783 |
| ROR1-AS1 | 4.598082134 | 0.000212843 | 0.002249362 |
| MEG3 | 4.613314788 | 1.15E-08 | 4.82E-07 |
| SGCD | 4.618094099 | 0.000133211 | 0.001538608 |
| CDC20B | 4.619197878 | 0.000225632 | 0.002374897 |
| ACVR2B | 4.620770779 | 0.000244542 | 0.002543174 |
| AGPAT9 | 4.662528807 | 0.000412529 | 0.003895848 |
| MPDZ | 4.662589146 | 5.91E-06 | 0.000111576 |
| LINC01410 | 4.666491044 | 7.57E-05 | 0.000969874 |
| RBMS3 | 4.700958578 | 0.000143976 | 0.001631659 |
| FLJ35934 | 4.736603338 | 1.02E-05 | 0.000179811 |
| PHLDA1 | 4.739391531 | 4.95E-08 | 1.66E-06 |
| CDH2 | 4.74073594 | 8.55E-05 | 0.001073474 |
| PCK2 | 4.76852172 | 0.000327531 | 0.00321013 |
| DENND5B | 4.770118193 | 1.22E-06 | 2.78E-05 |
| TAF4B | 4.774633875 | 6.40E-07 | 1.59E-05 |
| PCDH15 | 4.808082015 | 0.000140915 | 0.001606266 |
| KCNA7 | 4.829828146 | 0.000368745 | 0.003532988 |
| CCL4L2 | 4.835748257 | 1.95E-09 | 9.85E-08 |
| GUCY1A2 | 4.838193853 | 2.11E-05 | 0.000335882 |
| ZNRF1 | 4.863656806 | 6.40E-05 | 0.000853823 |
| NOX1 | 4.87074948 | 0.001214689 | 0.009579286 |
| LINC00550 | 4.876840047 | 5.32E-05 | 0.000732814 |
| IDO1 | 4.902383598 | 0.000431969 | 0.004051199 |
| EPAS1 | 4.915001086 | 0.000268971 | 0.002735474 |
| TTC4 | 4.918700719 | 1.38E-06 | 3.11E-05 |
| ABCA13 | 4.919035798 | 5.85E-05 | 0.000794765 |
| HSPA1B | 4.946815043 | 7.75E-08 | 2.45E-06 |
| LIF | 4.966111637 | 1.72E-05 | 0.000281211 |
| SCN1A | 4.979898805 | 0.000101008 | 0.001226345 |
| OR7E14P | 4.998870368 | 0.001101036 | 0.008807176 |
| LRAT | 5.000694254 | 4.09E-07 | 1.06E-05 |
| SLC43A3 | 5.005377723 | 2.63E-05 | 0.000404041 |
| TIMD4 | 5.005436182 | 3.76E-05 | 0.000543156 |
| SAMSN1 | 5.011778761 | 1.94E-18 | 6.59E-16 |
| AFAP1 | 5.012636389 | 7.09E-06 | 0.00013127 |
| FAM133A | 5.021258723 | 2.63E-05 | 0.000404402 |

| | | | |
|---|---|---|---|
| LGALS3 | 5.02573027 | 1.45E-16 | 3.14E-14 |
| MYC | 5.032781147 | 7.30E-06 | 0.000134393 |
| LPHN2 | 5.035517072 | 0.000122031 | 0.001430572 |
| KLHL6 | 5.052550498 | 3.84E-09 | 1.81E-07 |
| LINC01193 | 5.055596756 | 0.000327478 | 0.00321013 |
| DNAH7 | 5.074762764 | 8.86E-07 | 2.10E-05 |
| BCAR3 | 5.089356897 | 3.55E-06 | 7.13E-05 |
| ACVR1B | 5.158616835 | 1.21E-05 | 0.000207587 |
| PRELID2 | 5.162294971 | 1.37E-05 | 0.000228551 |
| ESRRG | 5.166051852 | 2.04E-05 | 0.000326297 |
| loc869 | 5.168723331 | 5.87E-05 | 0.000796334 |
| LOC101927151 | 5.174890519 | 3.84E-06 | 7.65E-05 |
| ZNF280B | 5.176690036 | 2.73E-05 | 0.000416902 |
| AHR | 5.178515641 | 6.40E-09 | 2.83E-07 |
| SLITRK4 | 5.228216419 | 6.47E-06 | 0.00012082 |
| OPRM1 | 5.249796249 | 1.78E-07 | 5.14E-06 |
| SOCS3 | 5.278195363 | 1.47E-07 | 4.36E-06 |
| ICAM1 | 5.288378933 | 4.18E-09 | 1.92E-07 |
| C15orf48 | 5.289539581 | 1.96E-06 | 4.24E-05 |
| EREG | 5.292667025 | 2.21E-07 | 6.32E-06 |
| OR7D2 | 5.294037197 | 2.42E-05 | 0.000377238 |
| TNF | 5.350315526 | 6.17E-08 | 1.97E-06 |
| PARM1 | 5.352470546 | 9.12E-08 | 2.80E-06 |
| LINC01441 | 5.387214333 | 3.43E-07 | 9.05E-06 |
| HAUS8 | 5.394258299 | 1.14E-05 | 0.000197694 |
| TADA2A | 5.401086149 | 2.90E-06 | 5.97E-05 |
| SLC7A5 | 5.435023663 | 2.48E-12 | 2.72E-10 |
| RGCC | 5.445976092 | 3.82E-08 | 1.33E-06 |
| KCNN3 | 5.48953835 | 1.54E-07 | 4.51E-06 |
| ANKRD62 | 5.535246031 | 1.22E-05 | 0.000208451 |
| SLC9C1 | 5.538416536 | 5.56E-07 | 1.41E-05 |
| KCNT2 | 5.544993362 | 7.31E-06 | 0.000134393 |
| PPIL6 | 5.558913132 | 1.43E-09 | 7.79E-08 |
| LOC285740 | 5.585587173 | 7.83E-05 | 0.000998322 |
| CRX | 5.622074906 | 3.05E-08 | 1.10E-06 |
| IER3 | 5.665257472 | 1.18E-08 | 4.93E-07 |
| PROS1 | 5.672319308 | 1.74E-05 | 0.000282568 |
| CCR7 | 5.698035684 | 1.22E-09 | 7.00E-08 |
| CXCL8 | 5.705054173 | 8.74E-06 | 0.000157265 |
| FOXD4L5 | 5.716091441 | 6.81E-07 | 1.67E-05 |
| PLCH1 | 5.768086928 | 5.06E-05 | 0.00070182 |
| OSMR | 5.768174596 | 2.17E-06 | 4.61E-05 |
| LCMT2 | 5.785586466 | 1.68E-07 | 4.90E-06 |
| ITPRIPL2 | 5.804654314 | 6.94E-08 | 2.20E-06 |
| C17orf96 | 5.81240295 | 4.66E-06 | 9.14E-05 |
| NFKB2 | 5.826574506 | 8.71E-12 | 8.03E-10 |
| CADM1 | 5.844897076 | 5.73E-07 | 1.43E-05 |
| PLCXD2-AS1 | 5.849947642 | 5.57E-11 | 4.24E-09 |
| CCL3 | 5.865513076 | 9.66E-20 | 4.20E-17 |
| CETN3 | 5.871917125 | 2.13E-09 | 1.06E-07 |

| | | | |
|---|---|---|---|
| TM4SF1 | 5.886774916 | 6.22E-06 | 0.000116453 |
| FBN2 | 5.957048086 | 9.51E-07 | 2.24E-05 |
| SOCS1 | 5.98420075 | 1.27E-08 | 5.21E-07 |
| LOC440896 | 5.992570057 | 2.44E-06 | 5.12E-05 |
| SLC27A2 | 5.999926196 | 4.09E-09 | 1.90E-07 |
| ST8SIA2 | 6.001099011 | 3.45E-06 | 6.99E-05 |
| NEGR1 | 6.062476762 | 1.77E-06 | 3.90E-05 |
| CYP1B1 | 6.064125944 | 7.58E-07 | 1.83E-05 |
| ELTD1 | 6.078159536 | 1.70E-05 | 0.000278867 |
| ADM2 | 6.109593783 | 1.15E-06 | 2.65E-05 |
| LOC100129434 | 6.129724567 | 3.02E-06 | 6.19E-05 |
| RUNX1T1 | 6.148497052 | 2.46E-08 | 9.09E-07 |
| VSIG10 | 6.163561443 | 3.55E-06 | 7.13E-05 |
| ZNF594 | 6.175521912 | 3.02E-07 | 8.20E-06 |
| GMPPB | 6.183770074 | 7.55E-10 | 4.55E-08 |
| FOXO3B | 6.220795565 | 1.37E-06 | 3.11E-05 |
| LOC731424 | 6.253854572 | 1.09E-06 | 2.54E-05 |
| CCR1 | 6.261927341 | 1.31E-09 | 7.37E-08 |
| LOC101929563 | 6.27703425 | 1.88E-06 | 4.11E-05 |
| ADAMTS5 | 6.285253033 | 9.29E-11 | 6.77E-09 |
| ANKRD30A | 6.299380525 | 1.98E-06 | 4.27E-05 |
| ZNF347 | 6.346235063 | 1.07E-10 | 7.65E-09 |
| LOC101928597 | 6.349185226 | 2.87E-07 | 7.89E-06 |
| EN2 | 6.349863923 | 1.14E-06 | 2.63E-05 |
| ABCA9 | 6.381300478 | 3.90E-06 | 7.76E-05 |
| TMOD2 | 6.423583562 | 3.18E-06 | 6.49E-05 |
| LMNA | 6.442237325 | 5.17E-09 | 2.33E-07 |
| LTA | 6.455409407 | 4.96E-12 | 4.98E-10 |
| LOC101926960 | 6.456822441 | 4.22E-08 | 1.45E-06 |
| C18orf63 | 6.529850934 | 3.22E-08 | 1.15E-06 |
| SLITRK1 | 6.548412133 | 1.50E-09 | 8.06E-08 |
| LINC01160 | 6.552780539 | 3.02E-07 | 8.20E-06 |
| SCARNA10 | 6.655075533 | 1.25E-08 | 5.14E-07 |
| SDC4 | 6.663954101 | 1.94E-09 | 9.85E-08 |
| SLC1A5 | 6.670976512 | 7.66E-17 | 1.71E-14 |
| FGF9 | 6.672066107 | 1.54E-11 | 1.29E-09 |
| FZD5 | 6.717810772 | 2.56E-07 | 7.12E-06 |
| SDK1 | 6.730737686 | 1.65E-06 | 3.64E-05 |
| FGF2 | 6.745057002 | 5.49E-08 | 1.78E-06 |
| LOC100131047 | 6.803590506 | 7.08E-07 | 1.72E-05 |
| TNFSF4 | 6.864605773 | 3.14E-12 | 3.28E-10 |
| EPS8 | 6.899693384 | 8.60E-08 | 2.66E-06 |
| C4orf32 | 6.958589004 | 3.26E-15 | 5.54E-13 |
| OR4F29 | 6.968393473 | 1.60E-08 | 6.28E-07 |
| OTOGL | 6.970677872 | 9.86E-09 | 4.20E-07 |
| C8orf44 | 6.971779004 | 3.43E-10 | 2.20E-08 |
| CREG2 | 7.033889683 | 9.19E-09 | 3.98E-07 |
| TMEM182 | 7.041001182 | 2.48E-07 | 6.96E-06 |
| CD82 | 7.048781712 | 3.28E-10 | 2.13E-08 |
| CISH | 7.075805742 | 6.47E-16 | 1.27E-13 |

| | | | |
|---|---|---|---|
| LINC01366 | 7.101844969 | 1.33E-09 | 7.38E-08 |
| ITGB8 | 7.102337802 | 2.85E-10 | 1.87E-08 |
| PSAT1 | 7.10844463 | 4.80E-10 | 2.98E-08 |
| USP44 | 7.225019494 | 2.75E-10 | 1.83E-08 |
| CD83 | 7.256853281 | 1.40E-28 | 1.83E-25 |
| PTPRQ | 7.311171393 | 4.36E-11 | 3.41E-09 |
| MIR155HG | 7.348736897 | 1.30E-09 | 7.37E-08 |
| UGT2B15 | 7.354215563 | 6.86E-09 | 3.02E-07 |
| EPHB2 | 7.358799305 | 4.84E-08 | 1.63E-06 |
| VDR | 7.365365456 | 1.85E-13 | 2.46E-11 |
| PDE6A | 7.448468046 | 4.64E-10 | 2.90E-08 |
| RIMS1 | 7.459309539 | 5.96E-17 | 1.41E-14 |
| NT5DC2 | 7.511652595 | 1.34E-11 | 1.16E-09 |
| STC1 | 7.514192645 | 2.46E-08 | 9.09E-07 |
| C2orf15 | 7.518234909 | 2.26E-16 | 4.66E-14 |
| TMEM217 | 7.567471041 | 2.00E-14 | 3.27E-12 |
| WDR87 | 7.574432591 | 1.14E-09 | 6.63E-08 |
| POM121L10P | 7.617218391 | 1.49E-08 | 5.93E-07 |
| FAS-AS1 | 7.624144284 | 1.48E-09 | 8.01E-08 |
| MFSD2A | 7.634343185 | 6.20E-13 | 7.25E-11 |
| LOC100507140 | 7.648143822 | 8.95E-10 | 5.31E-08 |
| P2RX5 | 7.71647904 | 8.80E-21 | 4.31E-18 |
| NHLH2 | 7.717057121 | 1.77E-08 | 6.88E-07 |
| ADAM12 | 7.787004977 | 7.98E-12 | 7.52E-10 |
| LOC643387 | 7.832209417 | 2.91E-14 | 4.38E-12 |
| NUP210L | 7.889335652 | 2.79E-12 | 2.95E-10 |
| GOLGA6L22 | 7.915654495 | 6.69E-12 | 6.47E-10 |
| NKAPL | 7.925740848 | 4.72E-11 | 3.66E-09 |
| DUSP4 | 8.0630914 | 2.73E-14 | 4.19E-12 |
| MITF | 8.091022051 | 2.09E-10 | 1.42E-08 |
| FAM81A | 8.195100975 | 8.18E-12 | 7.62E-10 |
| IL26 | 8.238999265 | 1.47E-11 | 1.25E-09 |
| loc1578 | 8.286050862 | 1.41E-22 | 7.36E-20 |
| KIAA0226L | 8.485667243 | 5.67E-23 | 3.41E-20 |
| ACVR1C | 8.509897365 | 2.50E-12 | 2.72E-10 |
| PLAU | 8.636759291 | 1.15E-18 | 4.10E-16 |
| ADAM19 | 8.655011139 | 9.20E-27 | 8.00E-24 |
| NCS1 | 8.65646468 | 3.45E-17 | 9.32E-15 |
| DMRTC2 | 8.660552518 | 1.32E-09 | 7.37E-08 |
| TXNDC5 | 8.700178995 | 9.24E-18 | 2.89E-15 |
| GPT2 | 8.828765664 | 2.35E-18 | 7.67E-16 |
| ITGA1 | 9.087552262 | 1.17E-17 | 3.53E-15 |
| TNFRSF9 | 9.100632574 | 9.25E-44 | 3.62E-40 |
| EGR2 | 9.853657948 | 2.33E-29 | 3.65E-26 |
| IFNG | 10.53550297 | 1.18E-31 | 2.32E-28 |
| RGS16 | 10.83734478 | 1.52E-33 | 3.96E-30 |
| IL2RA | 12.29186191 | 7.91E-47 | 6.19E-43 |

| Protein class analysis | | | | | |
|---|---|---|---|---|---|
| CD56neg stim vs CD56neg | | | | | |
| Homo sapiens (REF) | | | | | |
| PANTHER Protein Class | expected | Fold Enrichment | +/- | raw P value | FDR |
| chaperonin | 0.54 | 12.99 | + | 7.61E-06 | 7.35E-04 |
| tubulin | 1 | 6.03 | + | 1.02E-03 | 3.95E-02 |
| transaminase | 1.04 | 5.79 | + | 1.23E-03 | 3.95E-02 |
| chaperone | 7.75 | 3.74 | + | 1.04E-08 | 2.02E-06 |
| transferase | 27.58 | 1.7 | + | 8.25E-04 | 3.98E-02 |
| metabolite interconversion enzyme | 76.42 | 1.41 | + | 4.14E-04 | 2.66E-02 |
| CD56dim stim vs CD56dim | | | | | |
| Homo sapiens (REF) | | | | | |
| PANTHER Protein Class | expected | Fold Enrichment | +/- | raw P value | FDR |
| chaperonin | 0.72 | 13.95 | + | 8.98E-08 | 8.66E-06 |
| tubulin | 1.32 | 4.53 | + | 4.06E-03 | 4.12E-02 |
| vesicle coat protein | 2.54 | 3.94 | + | 5.79E-04 | 1.02E-02 |
| aminoacyl-tRNA synthetase | 2.26 | 3.54 | + | 3.63E-03 | 4.38E-02 |
| chaperone | 10.31 | 3.49 | + | 1.55E-09 | 2.99E-07 |
| translation initiation factor | 4.36 | 3.21 | + | 3.26E-04 | 6.30E-03 |
| kinase modulator | 4.74 | 2.74 | + | 1.93E-03 | 2.48E-02 |
| translation factor | 6.23 | 2.41 | + | 3.84E-03 | 4.36E-02 |
| translational protein | 18.48 | 2 | + | 2.15E-04 | 4.60E-03 |
| RNA processing factor | 13.02 | 2 | + | 1.80E-03 | 2.48E-02 |
| RNA metabolism protein | 37.78 | 1.72 | + | 7.57E-05 | 2.09E-03 |
| nucleic acid metabolism protein | 50.42 | 1.67 | + | 1.54E-05 | 9.92E-04 |
| transferase | 36.68 | 1.58 | + | 1.18E-03 | 1.75E-02 |
| metabolite interconversion enzyme | 101.66 | 1.34 | + | 8.35E-04 | 1.34E-02 |
| protein class | 664.28 | 1.1 | + | 4.29E-05 | 1.66E-03 |
| Unclassified | 471.72 | 0.85 | - | 4.29E-05 | 1.38E-03 |
| transmembrane signal receptor | 60.67 | 0.64 | - | 3.98E-03 | 4.27E-02 |
| defense/immunity protein | 27.47 | 0.33 | - | 1.05E-04 | 2.53E-03 |
| homeodomain transcription factor | 10.37 | 0.19 | - | 4.70E-03 | 4.54E-02 |
| immunoglobulin | 11.42 | < 0.01 | - | 2.54E-05 | 1.23E-03 |

| Supplemental Table 9 \| GSEA of NK cell in different culture reference list and ranked genes | |
|---|---|
| Ref:CD56$^{dim}$NK in PBMC_vs_CD56$^{neg}$NK in PBMCs enriched in CD56$^{dim}$NK | |
| Gene | log2fold_change |
| APOBEC3C | 2.040288868 |
| MTRNR2L8 | 1.988226763 |
| TNFRSF1B | 1.67254545 |
| SELPLG | 1.59900128 |
| CIRBP | 1.335937171 |
| TRMT112 | 1.327414041 |
| BTN3A1 | 1.29468915 |
| DCTN1 | 1.256459665 |
| UBA52 | 1.254704031 |
| TAPBP | 1.209423277 |
| PKM | 1.175148319 |
| ST6GAL1 | 1.173221175 |
| CTSD | 1.154142774 |
| PRF1 | 1.139880329 |
| AHNAK | 1.11339889 |
| S100A6 | 1.103841158 |
| PSMB10 | 1.102635689 |
| GNLY | 1.095388002 |
| HLA-E | 1.094686182 |
| IL2RG | 1.090660701 |
| IRF1 | 1.087888406 |
| LY6E | 1.085020246 |
| LAT2 | 1.080241259 |
| LOC606724 | 1.079875944 |
| GZMB | 1.075899214 |
| NAGK | 1.063635487 |
| CD247 | 1.062657738 |
| CD2 | 1.046932002 |
| XCL2 | 1.037407091 |
| ANXA6 | 1.036733423 |
| YPEL3 | 1.033691747 |
| MAST3 | 1.015388085 |
| GLTSCR2 | 1.007378356 |
| ARHGAP9 | 1.003525787 |
| KIAA0226 | 1.000707971 |

| Supplemental Table 10 \| Primers for bulk RNA-Seq and ATAC-Seq |
|---|
| **Barcoded primers for reverse transcription and final library amplication in bulk RNA-Seq** |
| Reverse transcription primers for Lin$^{-}$CD45$^{+}$CD56$^{-}$CD127$^{-}$ and Lin$^{-}$CD45$^{+}$CD56$^{-}$CD127$^{+}$ cells: |
| donor 1 Lin$^{-}$CD45$^{+}$CD56$^{-}$CD127$^{-}$ : GCCGGTAATACGACTCACTATAGGGAGTTCTACAGTCCGACGATCNNNNNNAGACTCTTTTTTTTTTTTTTTTTTTTTTTTV |
| donor 2 Lin$^{-}$CD45$^{+}$CD56$^{-}$CD127$^{-}$ : GCCGGTAATACGACTCACTATAGGGAGTTCTACAGTCCGACGATCNNNNNNCATGAGTTTTTTTTTTTTTTTTTTTTTTTTV |
| donor 3 Lin$^{-}$CD45$^{+}$CD56$^{-}$CD127$^{-}$ : GCCGGTAATACGACTCACTATAGGGAGTTCTACAGTCCGACGATCNNNNNNCAGATCTTTTTTTTTTTTTTTTTTTTTTTTV |
| donor 4 Lin$^{-}$CD45$^{+}$CD56$^{-}$CD127$^{-}$ : GCCGGTAATACGACTCACTATAGGGAGTTCTACAGTCCGACGATCNNNNNNTCACAGTTTTTTTTTTTTTTTTTTTTTTTTV |
| donor 5 Lin$^{-}$CD45$^{+}$CD56$^{-}$CD127$^{-}$ : GCCGGTAATACGACTCACTATAGGGAGTTCTACAGTCCGACGATCNNNNNNGTCTAGTTTTTTTTTTTTTTTTTTTTTTTTV |
| donor 1 Lin$^{-}$CD45$^{+}$CD56$^{-}$CD127$^{+}$ : GCCGGTAATACGACTCACTATAGGGAGTTCTACAGTCCGACGATCNNNNNNGTTGCATTTTTTTTTTTTTTTTTTTTTTTTV |
| donor 2 Lin$^{-}$CD45$^{+}$CD56$^{-}$CD127$^{+}$ : GCCGGTAATACGACTCACTATAGGGAGTTCTACAGTCCGACGATCNNNNNNACCATGTTTTTTTTTTTTTTTTTTTTTTTTV |
| donor 3 Lin$^{-}$CD45$^{+}$CD56$^{-}$CD127$^{+}$ : GCCGGTAATACGACTCACTATAGGGAGTTCTACAGTCCGACGATCNNNNNNACTCGATTTTTTTTTTTTTTTTTTTTTTTTV |
| donor 4 Lin$^{-}$CD45$^{+}$CD56$^{-}$CD127$^{+}$ : GCCGGTAATACGACTCACTATAGGGAGTTCTACAGTCCGACGATCNNNNNNACGTACTTTTTTTTTTTTTTTTTTTTTTTTV |
| donor 5 Lin$^{-}$CD45$^{+}$CD56$^{-}$CD127$^{+}$ : GCCGGTAATACGACTCACTATAGGGAGTTCTACAGTCCGACGATCNNNNNNCTGTGATTTTTTTTTTTTTTTTTTTTTTTTV |
| Final library amplication primers for Lin$^{-}$CD45$^{+}$CD56$^{-}$CD127$^{-}$ and Lin$^{-}$CD45$^{+}$CD56$^{-}$CD127$^{+}$ cells: |
| RP1: AAT GAT ACG GCG ACC ACC GAG ATC TAC ACG TTC AGA GTT CTA CAG TCC GA |
| RPI1: CAA GCA GAA GAC GGC ATA CGA GAT CGT GAT GTG ACT GGA GTT CCT TGG CAC CCG AGA ATT CCA |
| Reverse transcription primers for CD56$^{hi}$, CD56$^{dim}$, CD56$^{neg}$ NK cells and ILCs: |
| donor1 CD56$^{hi}$NK cells: GCCGGTAATACGACTCACTATAGGGAGTTCTACAGTCCGACGATCNNNNNNAGCTTCTTTTTTTTTTTTTTTTTTTTTTTTV |
| donor2 CD56$^{hi}$NK cells: GCCGGTAATACGACTCACTATAGGGAGTTCTACAGTCCGACGATCNNNNNNCACTAGTTTTTTTTTTTTTTTTTTTTTTTTV |
| donor3 CD56$^{hi}$NK cells: GCCGGTAATACGACTCACTATAGGGAGTTCTACAGTCCGACGATCNNNNNNAGGATCTTTTTTTTTTTTTTTTTTTTTTTTV |
| donor4 CD56$^{hi}$NK cells: GCCGGTAATACGACTCACTATAGGGAGTTCTACAGTCCGACGATCNNNNNNAGTGCATTTTTTTTTTTTTTTTTTTTTTTTV |
| donor1 CD56$^{dim}$NK cells: GCCGGTAATACGACTCACTATAGGGAGTTCTACAGTCCGACGATCNNNNNNTCCTAGTTTTTTTTTTTTTTTTTTTTTTTTV |
| donor2 CD56$^{dim}$NK cells: GCCGGTAATACGACTCACTATAGGGAGTTCTACAGTCCGACGATCNNNNNNGTACAGTTTTTTTTTTTTTTTTTTTTTTTTV |
| donor3 CD56$^{dim}$NK cells: GCCGGTAATACGACTCACTATAGGGAGTTCTACAGTCCGACGATCNNNNNNGTGACATTTTTTTTTTTTTTTTTTTTTTTTV |
| donor4 CD56$^{dim}$NK cells: GCCGGTAATACGACTCACTATAGGGAGTTCTACAGTCCGACGATCNNNNNNCTAGACTTTTTTTTTTTTTTTTTTTTTTTTV |
| donor1 CD56$^{neg}$NK cells: GCCGGTAATACGACTCACTATAGGGAGTTCTACAGTCCGACGATCNNNNNNCTCAGATTTTTTTTTTTTTTTTTTTTTTTTV |
| donor2 CD56$^{neg}$NK cells: GCCGGTAATACGACTCACTATAGGGAGTTCTACAGTCCGACGATCNNNNNNGACATGTTTTTTTTTTTTTTTTTTTTTTTTV |
| donor3 CD56$^{neg}$NK cells: GCCGGTAATACGACTCACTATAGGGAGTTCTACAGTCCGACGATCNNNNNNCACTTCTTTTTTTTTTTTTTTTTTTTTTTTV |
| donor4 CD56$^{neg}$NK cells: GCCGGTAATACGACTCACTATAGGGAGTTCTACAGTCCGACGATCNNNNNNCAGAAGTTTTTTTTTTTTTTTTTTTTTTTTV |
| donor1 ILCs: GCCGGTAATACGACTCACTATAGGGAGTTCTACAGTCCGACGATCNNNNNNTCACCATTTTTTTTTTTTTTTTTTTTTTTTV |
| donor2 ILCs: GCCGGTAATACGACTCACTATAGGGAGTTCTACAGTCCGACGATCNNNNNNTCTGTCTTTTTTTTTTTTTTTTTTTTTTTTV |
| donor3 ILCs: GCCGGTAATACGACTCACTATAGGGAGTTCTACAGTCCGACGATCNNNNNNGTTGAGTTTTTTTTTTTTTTTTTTTTTTTTV |
| donor4 ILCs: GCCGGTAATACGACTCACTATAGGGAGTTCTACAGTCCGACGATCNNNNNNACAGACTTTTTTTTTTTTTTTTTTTTTTTTV |
| Final library amplication primers for CD56$^{hi}$, CD56$^{dim}$, CD56$^{neg}$ NK cells and ILCs: |
| RP1: AAT GAT ACG GCG ACC ACC GAG ATC TAC ACG TTC AGA GTT CTA CAG TCC GA |
| RPI1: CAA GCA GAA GAC GGC ATA CGA GAT CGT GAT GTG ACT GGA GTT CCT TGG CAC CCG AGA ATT CCA |
| Reverse transcription primers for CD56$^{hi}$, CD56$^{dim}$, CD56$^{neg}$ NK cells stimulated with IL-12+IL15+IL-18: |
| donor2 CD56$^{hi}$NK cells: GCCGGTAATACGACTCACTATAGGGAGTTCTACAGTCCGACGATCNNNNNNCATGAGTTTTTTTTTTTTTTTTTTTTTTTTV |
| donor3 CD56$^{hi}$NK cells: GCCGGTAATACGACTCACTATAGGGAGTTCTACAGTCCGACGATCNNNNNNCAGATCTTTTTTTTTTTTTTTTTTTTTTTTV |
| donor4 CD56$^{hi}$NK cells: GCCGGTAATACGACTCACTATAGGGAGTTCTACAGTCCGACGATCNNNNNNTCACAGTTTTTTTTTTTTTTTTTTTTTTTTV |
| donor2 CD56$^{dim}$NK cells: GCCGGTAATACGACTCACTATAGGGAGTTCTACAGTCCGACGATCNNNNNNGTTGCATTTTTTTTTTTTTTTTTTTTTTTTV |
| donor3 CD56$^{dim}$NK cells: GCCGGTAATACGACTCACTATAGGGAGTTCTACAGTCCGACGATCNNNNNNACCATGTTTTTTTTTTTTTTTTTTTTTTTTV |
| donor4 CD56$^{dim}$NK cells: GCCGGTAATACGACTCACTATAGGGAGTTCTACAGTCCGACGATCNNNNNNACTCGATTTTTTTTTTTTTTTTTTTTTTTTV |
| donor2 CD56$^{neg}$NK cells: GCCGGTAATACGACTCACTATAGGGAGTTCTACAGTCCGACGATCNNNNNNCTGTGATTTTTTTTTTTTTTTTTTTTTTTTV |
| donor3 CD56$^{neg}$NK cells: GCCGGTAATACGACTCACTATAGGGAGTTCTACAGTCCGACGATCNNNNNNTGCAGATTTTTTTTTTTTTTTTTTTTTTTTV |
| donor4 CD56$^{neg}$NK cells: GCCGGTAATACGACTCACTATAGGGAGTTCTACAGTCCGACGATCNNNNNNCAACCATTTTTTTTTTTTTTTTTTTTTTTTV |
| Final library amplication primers for CD56$^{hi}$, CD56$^{dim}$, CD56$^{neg}$ NK cells stimulated with IL-12+IL-15+IL-18: |
| RP1: AAT GAT ACG GCG ACC ACC GAG ATC TAC ACG TTC AGA GTT CTA CAG TCC GA |
| RPI2: CAA GCA GAA GAC GGC ATA CGA GAT ACA TCG GTG ACT GGA GTT CCT TGG CAC CCG AGA ATT CCA |
| Reverse transcription primers for NK cells sorted from different culture: |
| donor1 CD56$^{dim}$NK cells sorted from PBMC culture: GCCGGTAATACGACTCACTATAGGGAGTTCTACAGTCCGACGATCNNNNNNAGACTCTTTTTTTTTTTTTTTTTTTTTTTTV |

| |
|---|
| donor2 $CD56^{dim}$NK cells sorted from PBMC culture: GCCGGTAATACGACTCACTATAGGGAGTTCTACAGTCCGACGATCNNNNNNCATGAGTTTTTTTTTTTTTTTTTTTTTTTTV |
| donor3 $CD56^{dim}$NK cells sorted from PBMC culture: GCCGGTAATACGACTCACTATAGGGAGTTCTACAGTCCGACGATCNNNNNNCAGATCTTTTTTTTTTTTTTTTTTTTTTTTV |
| donor4 $CD56^{dim}$NK cells sorted from PBMC culture: GCCGGTAATACGACTCACTATAGGGAGTTCTACAGTCCGACGATCNNNNNNTCACAGTTTTTTTTTTTTTTTTTTTTTTTTV |
| donor5 $CD56^{dim}$NK cells sorted from PBMC culture: GCCGGTAATACGACTCACTATAGGGAGTTCTACAGTCCGACGATCNNNNNNGTCTAGTTTTTTTTTTTTTTTTTTTTTTTTV |
| donor6 $CD56^{dim}$NK cells sorted from PBMC culture: GCCGGTAATACGACTCACTATAGGGAGTTCTACAGTCCGACGATCNNNNNNGTTGCATTTTTTTTTTTTTTTTTTTTTTTTV |
| donor1 $CD56^{dim}$NK cells sorted from $CD4^{-}$PBMC culture: GCCGGTAATACGACTCACTATAGGGAGTTCTACAGTCCGACGATCNNNNNNACCATGTTTTTTTTTTTTTTTTTTTTTTTTV |
| donor2 $CD56^{dim}$NK cells sorted from $CD4^{-}$PBMC culture: GCCGGTAATACGACTCACTATAGGGAGTTCTACAGTCCGACGATCNNNNNNACTCGATTTTTTTTTTTTTTTTTTTTTTTTV |
| donor4 $CD56^{dim}$NK cells sorted from $CD4^{-}$PBMC culture: GCCGGTAATACGACTCACTATAGGGAGTTCTACAGTCCGACGATCNNNNNNCTGTGATTTTTTTTTTTTTTTTTTTTTTTTV |
| donor1 $CD56^{dim}$NK cells sorted from $CD4^{-}$PBMC+IL-2 culture: GCCGGTAATACGACTCACTATAGGGAGTTCTACAGTCCGACGATCNNNNNNAGCTTCTTTTTTTTTTTTTTTTTTTTTTTTV |
| donor2 $CD56^{dim}$NK cells sorted from $CD4^{-}$PBMC+IL-2 culture: GCCGGTAATACGACTCACTATAGGGAGTTCTACAGTCCGACGATCNNNNNNCACTAGTTTTTTTTTTTTTTTTTTTTTTTTV |
| donor3 $CD56^{dim}$NK cells sorted from $CD4^{-}$PBMC+IL-2 culture: GCCGGTAATACGACTCACTATAGGGAGTTCTACAGTCCGACGATCNNNNNNAGGATCTTTTTTTTTTTTTTTTTTTTTTTTV |
| donor4 $CD56^{dim}$NK cells sorted from $CD4^{-}$PBMC+IL-2 culture: GCCGGTAATACGACTCACTATAGGGAGTTCTACAGTCCGACGATCNNNNNNAGTGCATTTTTTTTTTTTTTTTTTTTTTTTV |
| donor5 $CD56^{dim}$NK cells sorted from $CD4^{-}$PBMC+IL-2 culture: GCCGGTAATACGACTCACTATAGGGAGTTCTACAGTCCGACGATCNNNNNNTCCTAGTTTTTTTTTTTTTTTTTTTTTTTTV |
| donor6 $CD56^{dim}$NK cells sorted from $CD4^{-}$PBMC+IL-2 culture: GCCGGTAATACGACTCACTATAGGGAGTTCTACAGTCCGACGATCNNNNNNGTACAGTTTTTTTTTTTTTTTTTTTTTTTTV |
| donor1 $CD56^{neg}$NK cells sorted from PBMC culture: GCCGGTAATACGACTCACTATAGGGAGTTCTACAGTCCGACGATCNNNNNNAGACTCTTTTTTTTTTTTTTTTTTTTTTTTV |
| donor2 $CD56^{neg}$NK cells sorted from PBMC culture: GCCGGTAATACGACTCACTATAGGGAGTTCTACAGTCCGACGATCNNNNNNCATGAGTTTTTTTTTTTTTTTTTTTTTTTTV |
| donor3 $CD56^{neg}$NK cells sorted from PBMC culture: GCCGGTAATACGACTCACTATAGGGAGTTCTACAGTCCGACGATCNNNNNNCAGATCTTTTTTTTTTTTTTTTTTTTTTTTV |
| donor4 $CD56^{neg}$NK cells sorted from PBMC culture: GCCGGTAATACGACTCACTATAGGGAGTTCTACAGTCCGACGATCNNNNNNTCACAGTTTTTTTTTTTTTTTTTTTTTTTTV |
| donor5 $CD56^{neg}$NK cells sorted from PBMC culture: GCCGGTAATACGACTCACTATAGGGAGTTCTACAGTCCGACGATCNNNNNNGTCTAGTTTTTTTTTTTTTTTTTTTTTTTTV |
| donor6 $CD56^{neg}$NK cells sorted from PBMC culture: GCCGGTAATACGACTCACTATAGGGAGTTCTACAGTCCGACGATCNNNNNNGTTGCATTTTTTTTTTTTTTTTTTTTTTTTV |
| Final library amplication primers for NK cells sorted from different cutlure: |
| RP1: AAT GAT ACG GCG ACC ACC GAG ATC TAC ACG TTC AGA GTT CTA CAG TCC GA |
| RPI1 for $CD56^{dim}$NK cells: CAA GCA GAA GAC GGC ATA CGA GAT CGT GAT GTG ACT GGA GTT CCT TGG CAC CCG AGA ATT CCA |
| RPI2 for $CD56^{neg}$NK cells: CAA GCA GAA GAC GGC ATA CGA GAT ACA TCG GTG ACT GGA GTT CCT TGG CAC CCG AGA ATT CCA |
| **Primers for ATAC-Seq** |
| donor1 $CD56^{hi}$NK cells: CAAGCAGAAGACGGCATACGAGATAGGAGTCCGTCTCGTGGGCTCGGAGATGT |
| donor1 $CD56^{dim}$NK cells: CAAGCAGAAGACGGCATACGAGATCATGCCTAGTCTCGTGGGCTCGGAGATGT |
| donor1 $CD56^{neg}$NK cells: CAAGCAGAAGACGGCATACGAGATGTAGAGAGGTCTCGTGGGCTCGGAGATGT |
| donor1 ILCs: CAAGCAGAAGACGGCATACGAGATCCTCTCTGGTCTCGTGGGCTCGGAGATGT |
| donor2 $CD56^{hi}$NK cells: CAAGCAGAAGACGGCATACGAGATTCGCCTTAGTCTCGTGGGCTCGGAGATGT |
| donor2 $CD56^{dim}$NK cells: CAAGCAGAAGACGGCATACGAGATCTAGTACGGTCTCGTGGGCTCGGAGATGT |
| donor2 $CD56^{neg}$NK cells: CAAGCAGAAGACGGCATACGAGATTTCTGCCTGTCTCGTGGGCTCGGAGATGT |
| donor2 ILCs: CAAGCAGAAGACGGCATACGAGATGCTCAGGAGTCTCGTGGGCTCGGAGATGT |
| common primer: AAT GAT CGG CGA CCA CCG ATA TCT ACA CTC GTC GGC AGC GTC AGA TGTG |
